## Supplemental Information for "Promoter selectivity of the RhlR quorum-sensing transcription factor receptor in *Pseudomonas aeruginosa* is coordinated by distinct and overlapping dependencies on C_4_-homoserine lactone and PqsE"

### Figure S1

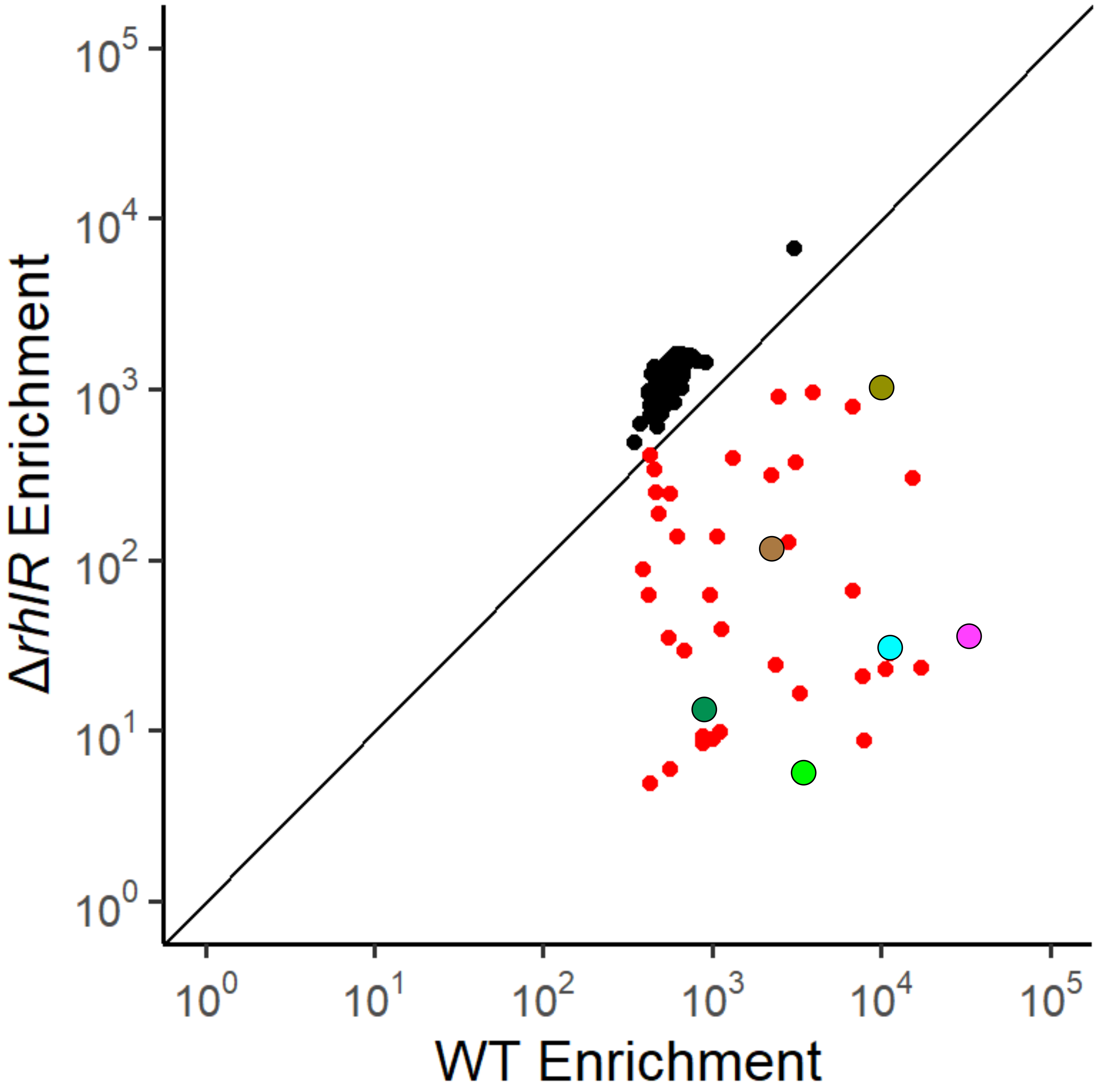

$10^5$
$10^4$
$10^3$
$10^2$
$10^1$
$10^0$
ΔrhlR Enrichment
$10^0$
$10^1$
$10^2$
$10^3$
$10^4$
$10^5$
WT Enrichment

Figure S2

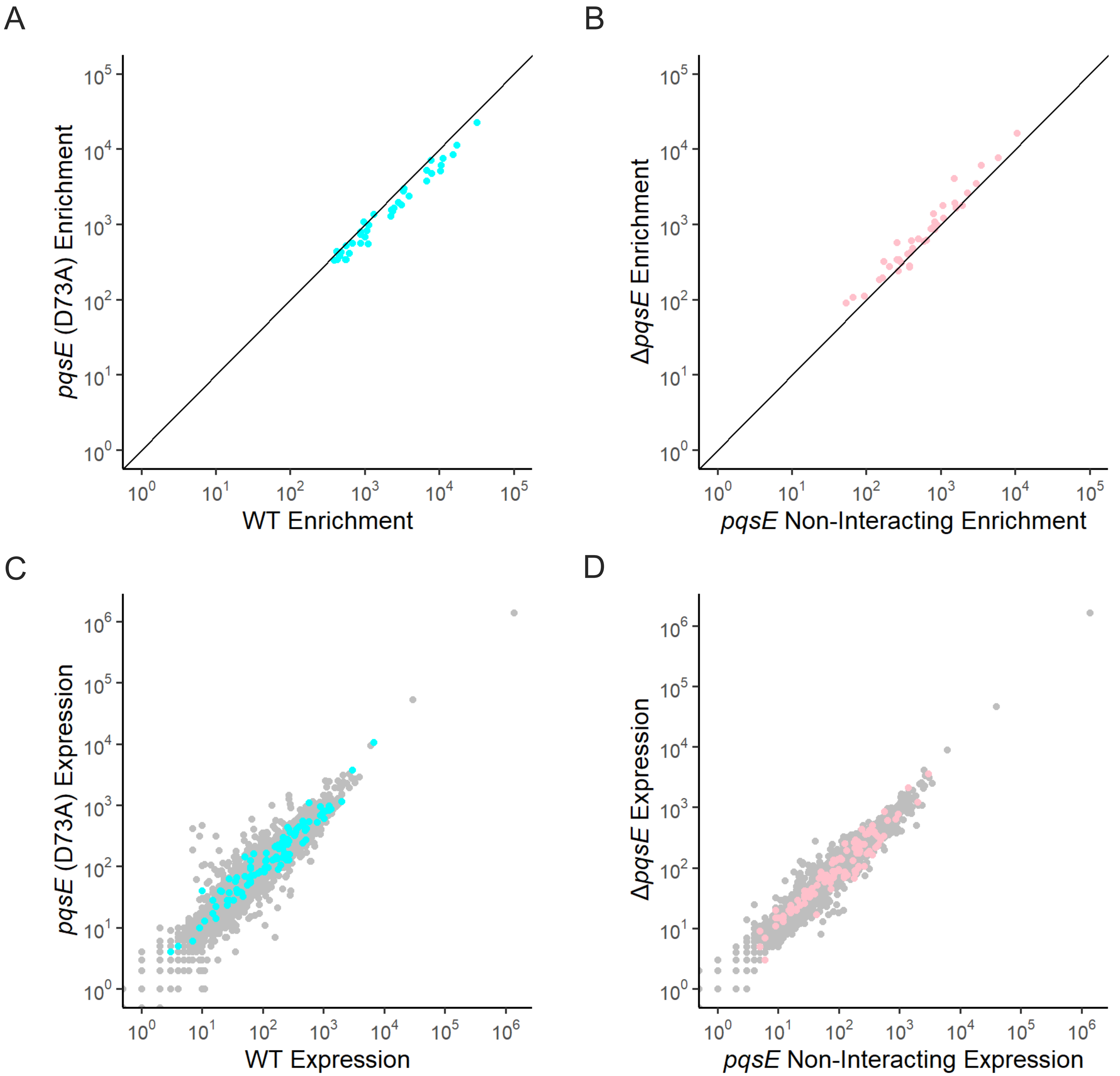

A
B
C
D
pqsE (D73A) Enrichment
WT Enrichment
ΔpqsE Enrichment
pqsE Non-Interacting Enrichment
pqsE (D73A) Expression
WT Expression
ΔpqsE Expression
pqsE Non-Interacting Expression

Table 1

| | A | B | C | D | E | F | G | H | I | J |
|---|---|---|---|---|---|---|---|---|---|---|
| 1 | **Site Location[a]** | **Synonym[b]** | **Motif[c]** | **WT Binding** | **Δ*rhlR* Binding** | **Δ*pqsE* Binding** | **Δ*rhlI* Binding** | **RhlR Expression Change** | **PqsE Expression Change** | **RhlI Expression Chang** |
| 2 | 64211 | *phzH* | CAACTATTGAATATAAGAGTT | 2437.4 | 906.1 | 906.8 | 962.1 | 5.8 | 3.0 | 5.8 |
| 3 | 139733 | *PA14_01490 (rahU)* | - | 2364.4 | 24.5 | 640.1 | 2111.9 | 18.4 | 1.4 | 2.9 |
| 4 | 560072 | *PA14_06320* | CCCCTACTAGAATTCACAGGT | 1010.4 | 8.9 | 596.4 | 1461.2 | 0.8 | 0.8 | 0.6 |
| 5 | 754615 | *rpsL operon* | GCCCTGCCAATTTTCGGGGTA | 552.9 | 35.1 | 194.2 | 144.2 | 0.6 | 1.4 | 1.1 |
| 6 | 812427 | *phzB1*, phzC1* | - | 3931.3 | 966.2 | 1641.1 | 3353.5 | 264.5, 138.3 | 29.3, 20.8 | 44.1, 31.1 |
| 7 | 813529 | *phzA1* operon, *phzM* | CACCTACCAGATCTTGTAGTT | 10446.1 | 995.9 | 1676.3 | 5496.0 | 124, 19 | 15.5, 8.4 | 15.5, 7.0 |
| 8 | 895433 | *PA14_10360* operon | GAACTGCCAGGATTAGCGGTT | 6809.1 | 800.8 | 1376.8 | 794.7 | 28.9 | 1.9 | 12.9 |
| 9 | 1372370 | *PA14_16100*, PA14_16110* | CACCTGCTTGAAATTGCAGTA | 6758.9 | 66.6 | 1914.7 | 1152.0 | 1.5, 1.3 | 1.4, 0.7 | 1.6, 1.1 |
| 10 | 1387361 | *lasB* | AACCTGCCAGAACTGGCAGGT | 878.9 | 9.4 | 605.7 | 296.7 | 2.4 | 0.7 | 3.2 |
| 11 | 1620628 | *PA14_18800* | TACCTGAGAGATTTATGAGTT | 1315.4 | 395.0 | 863.9 | 629.5 | 4.0 | 1.1 | 3.6 |
| 12 | 1621028 | *PA14_18810* operon | - | 481.3 | 189.0 | 481.7 | 257.6 | 0.6 | 1.3 | 1.3 |
| 13 | 1648391 | *rhlA* operon | TAACTGCCAGATTTCACAGGA | 11271.2 | 28.5 | 7736.8 | 2586.2 | 45.3 | 1.0 | 5.1 |
| 14 | 1651804 | *rhlI* | - | 1112.2 | 9.8 | 1080.2 | 1021.5 | 1.5 | 1.0 | 35.2 |
| 15 | 1735823 | *PA14_20130, PA14_20140* (*fpr*) | - | 414.7 | 62.4 | 307.4 | 233.3 | 1.0, 1.2 | 1.0, 1.2 | 1.0, 1.1 |
| 16 | 1774336 | *lecB, PA14_20620* operon | CCACTGCTAGAGTTCGCAGGA | 32231.7 | 35.0 | 577.6 | 33814.6 | 5.6, 1.4 | 2.9, 1.1 | 2.2, 1.0 |
| 17 | 1816906 | *PA14_21020* (*azeB*) operon, *PA14_21030* (*azeA*) | TACCTACCAGAATTAACAGTT | 15193.0 | 301.7 | 16297.0 | 6827.0 | 11.2, 4.3 | 2.3, 0.9 | 5.3, 2.2 |
| 18 | 1924465 | *PA14_22090* | - | 390.1 | 89.1 | 242.1 | 142.1 | 1.0 | 0.9 | 1.0 |
| 19 | 2444398 | *PA14_28250*, PA14_28260* | AAGCTGCCGGATCTGGTAGGC | 885.1 | 8.4 | 321.1 | 166.8 | 0.9, 1.2 | 0.8, 0.8 | 0.8, 1.1 |
| 20 | 2568048 | *PA14_29620, fhp* operon | AAACTACCAGAATTCACGGGC | 7797.7 | 20.9 | 1780.0 | 1602.7 | 0.7, 0.02 | 0.9, 0.03 | 0.8, 1.1 |
| 21 | 2647472 | *PA14_30570, PA14_30580* (*vqsR*) | CACCTACCAGAACTGGTAGTT | 2294.2 | 118.2 | 3484.9 | 1094.3 | 1.8, 0.8 | 1.0, 0.6 | 1.4, 0.8 |
| 22 | 2677542 | *PA14_30840** | - | 17187.9 | 23.2 | 90.7 | 9811.6 | 1.3 | 0.8 | 0.9 |
| 23 | 2721761 | *pa1L* | CTCCTGCATGAATTGATAGGC | 431.0 | 410.1 | 278.4 | 315.4 | 5.5 | 1.7 | 6.0 |
| 24 | 3188223 | *tnpS, tnpT* operon | - | 556.8 | 243.6 | 339.1 | 258.0 | 1.3, 1.0 | 1.0, 1.2 | 1.0, 1.2 |
| 25 | 3236436 | *hcnA* operon, *exoY* | - | 3335.0 | 5.4 | 342.1 | 4885.0 | 10.2, 1.4 | 3.6, 0.8 | 2.4, 0.9 |
| 26 | 3364761 | *PA14_37745* operon | GCCCTGCCAGATTTCGCAGGC | 423.7 | 4.9 | 112.0 | 37.0 | 34.2 | 1.5 | 14.6 |
| 27 | 3561157 | *phzB2*, phzC2* | - | 453.4 | 340.6 | 420.0 | 336.5 | 100.1, 132.4 | 8.3, 14.9 | 40.0, 34.0 |
| 28 | 3561969 | *phzA2* operon | CACCTGTAATTTTTAAGGGGT | 2238.4 | 314.8 | 1218.8 | 348.0 | 88.5 | 8.4 | 29.5 |
| 29 | 3600666 | *lasA* | CAACTATCAGCTTTTGCAGTA | 555.5 | 6.0 | 277.4 | 206.6 | 2.3 | 0.9 | 2.5 |
| 30 | 3831541 | *hsiA2* operon, *hcpD* | - | 3119.5 | 373.5 | 1767.9 | 629.0 | 2.7, 2.2 | 1.9, 1.8 | 2.3, 2.7 |
| 31 | 4285523 | *PA14_48140* operon | CACCTGGCAGAACTGACAGGT | 1059.8 | 136.6 | 405.7 | 272.9 | 0.9 | 0.7 | 0.9 |
| 32 | 4313855 | *PA14_48530* operon | CAACTATGAGAATTGGTAGTT | 2800.5 | 127.7 | 2597.9 | 2645.5 | 2.7 | 1.0 | 1.3 |
| 33 | 4314560 | *PA14_48530** | - | 620.4 | 137.3 | 974.9 | 651.9 | 2.7 | 1.0 | 1.3 |
| 34 | 4382169 | *PA14_49310* | CAACTGCCAGATCTGGCAGGC | 1136.8 | 39.3 | 880.7 | 1398.4 | 0.9 | 0.8 | 1.2 |
| 35 | 4425570 | *PA14_49740, PA14_49750* operon | AAACTACCGGAATTCACAGGT | 7939.2 | 8.7 | 6100.9 | 3189.2 | 0.9, 6.8 | 1.0, 0.9 | 0.8, 2.1 |
| 36 | 5159548 | *PA14_57970*, PA14_57980* | CAACTGTTACATATGAGCGGT | 881.2 | 12.7 | 106.4 | 236.3 | 0.9, 1.0 | 1.2, 1.5 | 0.9, 1.1 |
| 37 | 5268881 | *PA14_59180** | CAACTCGCAGAACTGGTGGGG | 679.5 | 29.4 | 185.2 | 247.1 | 0.9 | 0.5 | 0.7 |
| 38 | 6082366 | *rmlB* operon | CACCTACCAGATCTGGGGTTG | 10477.0 | 23.2 | 4067.5 | 1291.5 | 2.3 | 0.9 | 1.8 |
| 39 | 6093032 | *arcD* operon | TCCCTATAGGAATTGAGAGTG | 3285.2 | 16.5 | 333.9 | 205.1 | 0.9 | 2.6 | 0.7 |

[a]Site location refers to the genomic position of the base in the center of a given ChIP peak.
[b]Two genes are listed where a site is upstream of a gene on both strands, or within one gene and upstream of an adjacent gene on the same strand.
*Gene is associated with an internal binding site.
[c]Binding motifs were identified using MEME analysis [63] and the sequence contributing to the consensus motif is listed in line with its associated ChIP site and gene(s) where identified.

Table S1

| | A | B | C | D | E | F | G | H |
|---|---|---|---|---|---|---|---|---|
| 1 | **Site Location** | **WT** | **Δ*rhlR*** | **Δ*rhlI*** | **Δ*pqsE*** | **Δ*rhlI* Δ*pqsE*** | ***pqsE*- NI** | ***pqsE* (D73A)** |
| 2 | 53066 | 591.4 | 1026.9 | 580.9 | 417.0 | 331.8 | 681.8 | 507.6 |
| 3 | 54148 | 432.0 | 1241.6 | 489.9 | 311.3 | 221.4 | 776.1 | 385.1 |
| 4 | 64211 | 2437.4 | 906.1 | 962.1 | 906.8 | 337.2 | 779.4 | 1647.0 |
| 5 | 139733 | 2364.4 | 24.5 | 2111.9 | 640.1 | 82.2 | 501.9 | 1509.8 |
| 6 | 560072 | 1010.4 | 8.9 | 1461.2 | 596.4 | 238.6 | 597.8 | 686.4 |
| 7 | 711431 | 465.4 | 751.6 | 588.4 | 415.7 | 315.1 | 644.9 | 441.2 |
| 8 | 711675 | 550.1 | 952.4 | 657.0 | 410.0 | 267.6 | 627.3 | 535.0 |
| 9 | 733156 | 600.0 | 1105.2 | 758.7 | 683.3 | 505.2 | 836.8 | 598.4 |
| 10 | 736253 | 598.9 | 838.4 | 660.0 | 615.1 | 424.6 | 726.3 | 638.4 |
| 11 | 736928 | 660.5 | 1160.2 | 873.9 | 724.1 | 517.1 | 920.8 | 671.3 |
| 12 | 737180 | 555.3 | 923.5 | 663.7 | 592.1 | 363.0 | 717.8 | 592.4 |
| 13 | 754615 | 552.9 | 35.1 | 144.2 | 194.2 | 253.8 | 164.9 | 341.4 |
| 14 | 812427 | 3931.3 | 966.2 | 3353.5 | 1641.1 | 263.2 | 1602.2 | 2375.6 |
| 15 | 813529 | 10446.1 | 995.9 | 5496.0 | 1676.3 | 352.0 | 1598.7 | 5131.9 |
| 16 | 895433 | 6809.1 | 800.8 | 794.7 | 1376.8 | 157.7 | 792.7 | 3790.1 |
| 17 | 898184 | 465.6 | 1088.9 | 613.8 | 470.2 | 210.5 | 686.5 | 418.8 |
| 18 | 956560 | 568.5 | 931.5 | 832.7 | 458.8 | 297.1 | 561.5 | 471.9 |
| 19 | 1195684 | 735.5 | 1580.2 | 900.4 | 713.5 | 390.2 | 876.1 | 666.6 |
| 20 | 1195963 | 599.2 | 1305.3 | 862.5 | 698.3 | 399.1 | 995.2 | 575.3 |
| 21 | 1231146 | 630.4 | 1040.8 | 668.5 | 508.8 | 282.5 | 819.9 | 570.8 |
| 22 | 1372370 | 6758.9 | 66.6 | 1152.0 | 1914.7 | 183.9 | 1534.5 | 5258.8 |
| 23 | 1387361 | 878.9 | 9.4 | 296.7 | 605.7 | 42.0 | 404.6 | 796.4 |
| 24 | 1620628 | 1315.4 | 395.0 | 629.5 | 863.9 | 174.4 | 831.3 | 1366.9 |
| 25 | 1621028 | 481.3 | 189.0 | 257.6 | 481.7 | 207.4 | 419.2 | 428.9 |
| 26 | 1635342 | 521.0 | 792.8 | 550.4 | 429.5 | 382.3 | 657.8 | 505.1 |
| 27 | 1648391 | 11271.2 | 28.5 | 2586.2 | 7736.8 | 460.1 | 5875.2 | 7577.1 |
| 28 | 1651804 | 1112.2 | 9.8 | 1021.5 | 1080.2 | 247.9 | 827.1 | 553.6 |
| 29 | 1735823 | 414.7 | 62.4 | 233.3 | 307.4 | 2048.2 | 295.2 | 437.7 |
| 30 | 1762030 | 518.2 | 967.0 | 657.8 | 498.6 | 239.7 | 743.8 | 460.9 |
| 31 | 1765892 | 589.0 | 1008.6 | 564.8 | 487.3 | 390.8 | 742.2 | 587.8 |
| 32 | 1766203 | 526.7 | 971.0 | 522.2 | 421.4 | 240.5 | 744.5 | 452.3 |
| 33 | 1767099 | 510.0 | 913.0 | 624.9 | 431.1 | 338.3 | 658.8 | 520.3 |
| 34 | 1774336 | 32231.7 | 35.0 | 33814.6 | 577.6 | 45.8 | 259.5 | 22562.8 |
| 35 | 1816906 | 15193.0 | 301.7 | 6827.0 | 16297.0 | 3344.4 | 10539.4 | 8477.9 |
| 36 | 1861853 | 517.0 | 994.2 | 662.6 | 522.7 | 310.2 | 763.8 | 467.5 |
| 37 | 1863055 | 461.7 | 860.8 | 562.3 | 388.6 | 208.2 | 677.0 | 430.4 |
| 38 | 1863312 | 690.9 | 1403.3 | 761.5 | 549.9 | 360.8 | 876.9 | 674.6 |
| 39 | 1921598 | 457.3 | 252.1 | 195.7 | 267.8 | 1086.3 | 379.8 | 385.2 |
| 40 | 1924465 | 390.1 | 89.1 | 142.1 | 242.1 | 1145.9 | 269.9 | 331.7 |
| 41 | 1925890 | 527.8 | 1209.1 | 699.9 | 610.6 | 402.1 | 800.7 | 513.3 |
| 42 | 1928093 | 595.5 | 1119.2 | 741.9 | 621.9 | 326.3 | 831.7 | 544.0 |
| 43 | 1928610 | 656.2 | 1466.2 | 862.6 | 672.8 | 372.0 | 942.0 | 666.3 |
| 44 | 1929125 | 486.8 | 1127.1 | 738.4 | 564.4 | 268.1 | 752.7 | 480.4 |
| 45 | 1929320 | 590.3 | 1307.9 | 868.1 | 711.2 | 327.9 | 862.6 | 627.4 |
| 46 | 1929423 | 550.5 | 1285.7 | 838.0 | 743.3 | 295.5 | 957.6 | 560.2 |
| 47 | 1931650 | 648.3 | 1276.1 | 759.1 | 660.1 | 267.7 | 884.6 | 654.3 |
| 48 | 1931828 | 617.5 | 1335.5 | 789.2 | 653.1 | 301.5 | 980.3 | 598.2 |
| 49 | 1931996 | 579.0 | 1157.7 | 767.1 | 667.7 | 239.4 | 919.0 | 531.3 |
| 50 | 1932145 | 519.1 | 1207.3 | 792.9 | 659.3 | 223.6 | 755.9 | 543.3 |
| 51 | 1932413 | 643.3 | 1339.9 | 908.4 | 729.8 | 281.7 | 1026.5 | 626.6 |
| 52 | 1933063 | 501.9 | 1141.3 | 731.7 | 567.3 | 219.3 | 853.5 | 496.4 |
| 53 | 1934437 | 639.6 | 1469.1 | 796.6 | 695.6 | 223.9 | 981.2 | 549.4 |
| 54 | 1934549 | 577.3 | 1433.1 | 717.8 | 678.2 | 199.3 | 981.4 | 569.0 |
| 55 | 1938348 | 666.5 | 1244.9 | 749.0 | 653.2 | 256.2 | 921.3 | 621.8 |
| 56 | 1938815 | 510.9 | 1226.5 | 657.5 | 535.0 | 239.3 | 813.3 | 513.2 |
| 57 | 1939178 | 545.9 | 1050.0 | 654.6 | 629.8 | 317.1 | 833.8 | 512.3 |
| 58 | 2027979 | 581.5 | 1020.4 | 722.0 | 592.9 | 398.2 | 830.4 | 561.1 |

| | A | B | C | D | E | F | G | H |
|---|---|---|---|---|---|---|---|---|
| 59 | 2028457 | 472.6 | 1284.7 | 717.8 | 592.8 | 253.4 | 832.0 | 500.2 |
| 60 | 2028987 | 463.8 | 987.5 | 803.7 | 535.8 | 217.1 | 714.2 | 438.4 |
| 61 | 2029197 | 578.2 | 1186.1 | 873.7 | 563.8 | 312.2 | 860.7 | 537.6 |
| 62 | 2029507 | 534.7 | 1124.4 | 722.7 | 571.1 | 259.9 | 702.4 | 480.5 |
| 63 | 2029944 | 504.1 | 948.0 | 766.5 | 504.7 | 275.9 | 608.9 | 489.6 |
| 64 | 2030348 | 493.6 | 960.2 | 558.0 | 453.2 | 295.4 | 766.6 | 490.7 |
| 65 | 2030753 | 812.0 | 1477.7 | 819.3 | 520.3 | 374.5 | 1053.9 | 784.0 |
| 66 | 2031255 | 772.7 | 1565.8 | 737.5 | 524.6 | 295.5 | 1107.4 | 723.0 |
| 67 | 2031678 | 535.7 | 1305.9 | 618.6 | 457.1 | 374.2 | 964.1 | 478.1 |
| 68 | 2032697 | 608.6 | 1623.3 | 651.2 | 451.6 | 482.2 | 1345.6 | 551.5 |
| 69 | 2034193 | 541.8 | 1064.3 | 650.6 | 529.4 | 294.1 | 883.0 | 504.3 |
| 70 | 2035624 | 554.3 | 1141.3 | 775.0 | 649.4 | 298.8 | 894.6 | 592.9 |
| 71 | 2035849 | 489.7 | 973.7 | 683.2 | 587.5 | 224.5 | 798.6 | 470.0 |
| 72 | 2036100 | 552.9 | 1087.9 | 619.3 | 519.2 | 183.5 | 688.9 | 507.9 |
| 73 | 2036498 | 515.4 | 1045.3 | 587.0 | 478.0 | 248.7 | 731.6 | 438.9 |
| 74 | 2037512 | 515.9 | 1078.0 | 699.7 | 627.9 | 225.3 | 693.9 | 480.9 |
| 75 | 2039630 | 612.3 | 1048.8 | 677.9 | 556.4 | 262.3 | 824.8 | 550.3 |
| 76 | 2040895 | 517.2 | 1189.2 | 730.0 | 585.3 | 330.4 | 827.3 | 496.6 |
| 77 | 2041043 | 596.3 | 1237.6 | 762.7 | 606.4 | 361.8 | 874.2 | 590.4 |
| 78 | 2422185 | 655.4 | 1030.1 | 812.0 | 544.5 | 349.5 | 829.2 | 601.5 |
| 79 | 2444398 | 885.1 | 8.4 | 166.8 | 321.1 | 27.9 | 170.7 | 736.4 |
| 80 | 2477339 | 526.0 | 983.5 | 634.5 | 493.6 | 243.2 | 703.7 | 526.3 |
| 81 | 2477468 | 510.3 | 890.2 | 601.2 | 447.9 | 376.6 | 585.4 | 533.6 |
| 82 | 2478717 | 494.3 | 727.6 | 611.8 | 478.8 | 265.3 | 611.9 | 481.8 |
| 83 | 2478901 | 547.5 | 947.3 | 808.1 | 684.8 | 334.3 | 868.2 | 544.1 |
| 84 | 2568048 | 7797.7 | 20.9 | 1602.7 | 1780.0 | 283.8 | 1939.8 | 7130.4 |
| 85 | 2647472 | 2294.2 | 118.2 | 1094.3 | 3484.9 | 761.6 | 2963.5 | 1546.8 |
| 86 | 2677542 | 17187.9 | 23.2 | 9811.6 | 90.7 | 136.7 | 53.7 | 11243.3 |
| 87 | 2721761 | 431.0 | 410.1 | 315.4 | 278.4 | 168.4 | 382.9 | 420.0 |
| 88 | 2859603 | 492.7 | 721.2 | 462.9 | 365.8 | 263.7 | 481.8 | 433.7 |
| 89 | 2866123 | 623.9 | 1320.1 | 791.0 | 597.3 | 221.5 | 837.6 | 635.8 |
| 90 | 2867864 | 592.6 | 1287.8 | 529.8 | 454.3 | 178.9 | 788.5 | 539.2 |
| 91 | 2870549 | 587.3 | 1546.6 | 704.2 | 616.3 | 363.9 | 935.2 | 577.3 |
| 92 | 2870709 | 601.0 | 1377.3 | 696.8 | 585.0 | 303.8 | 980.6 | 586.9 |
| 93 | 2870815 | 603.2 | 1120.5 | 642.0 | 528.4 | 293.8 | 811.4 | 513.0 |
| 94 | 2910738 | 347.0 | 487.4 | 210.6 | 248.9 | 387.6 | 308.4 | 427.6 |
| 95 | 3174873 | 533.9 | 1024.4 | 616.2 | 444.4 | 286.6 | 705.2 | 510.8 |
| 96 | 3175027 | 602.4 | 1068.5 | 585.8 | 443.2 | 216.5 | 776.6 | 567.7 |
| 97 | 3181581 | 538.8 | 1246.5 | 806.2 | 670.6 | 324.4 | 993.3 | 564.8 |
| 98 | 3182110 | 497.4 | 941.3 | 642.8 | 480.5 | 301.3 | 725.2 | 493.6 |
| 99 | 3183400 | 576.6 | 1201.2 | 590.0 | 570.8 | 251.6 | 907.1 | 563.0 |
| 100 | 3183679 | 621.3 | 1317.2 | 768.4 | 635.1 | 344.1 | 1025.3 | 669.8 |
| 101 | 3183968 | 544.1 | 1154.4 | 827.1 | 653.8 | 252.0 | 890.3 | 525.4 |
| 102 | 3184251 | 513.4 | 998.2 | 670.4 | 537.1 | 248.2 | 845.0 | 479.8 |
| 103 | 3188223 | 556.8 | 243.6 | 258.0 | 339.1 | 969.3 | 259.0 | 338.6 |
| 104 | 3236436 | 3335.0 | 5.4 | 4885.0 | 342.1 | 111.0 | 274.7 | 2999.7 |
| 105 | 3364761 | 423.7 | 4.9 | 37.0 | 112.0 | 18.8 | 94.0 | 343.8 |
| 106 | 3405263 | 478.9 | 973.4 | 584.7 | 494.4 | 217.8 | 672.4 | 468.5 |
| 107 | 3405951 | 597.1 | 1047.7 | 719.8 | 495.3 | 337.0 | 835.6 | 540.4 |
| 108 | 3407018 | 492.6 | 871.5 | 669.0 | 527.5 | 262.0 | 772.4 | 482.6 |
| 109 | 3512549 | 509.6 | 1282.9 | 644.9 | 510.6 | 223.0 | 1051.2 | 447.9 |
| 110 | 3515027 | 451.6 | 964.7 | 581.2 | 543.1 | 230.4 | 629.1 | 495.3 |
| 111 | 3561157 | 453.4 | 340.6 | 336.5 | 420.0 | 174.7 | 388.6 | 392.9 |
| 112 | 3561969 | 2238.4 | 314.8 | 348.0 | 1218.8 | 106.0 | 1093.1 | 1287.0 |
| 113 | 3600666 | 555.5 | 6.0 | 206.6 | 277.4 | 35.8 | 205.1 | 517.6 |
| 114 | 3831541 | 3119.5 | 373.5 | 629.0 | 1767.9 | 169.5 | 1056.1 | 1815.4 |
| 115 | 4133306 | 470.6 | 1013.1 | 639.9 | 523.3 | 214.9 | 689.6 | 424.9 |
| 116 | 4134459 | 452.7 | 880.3 | 706.6 | 521.6 | 275.4 | 681.2 | 459.7 |

| | A | B | C | D | E | F | G | H |
|---|---|---|---|---|---|---|---|---|
| 117 | 4134700 | 509.7 | 1241.0 | 808.8 | 708.4 | 348.4 | 870.5 | 537.9 |
| 118 | 4285523 | 1059.8 | 136.6 | 272.9 | 405.7 | 99.7 | 359.3 | 826.5 |
| 119 | 4313855 | 2800.5 | 127.7 | 2645.5 | 2597.9 | 556.1 | 2295.9 | 1970.4 |
| 120 | 4314560 | 620.4 | 137.3 | 651.9 | 974.9 | 377.1 | 885.9 | 416.6 |
| 121 | 4354082 | 528.0 | 1369.7 | 703.9 | 482.6 | 224.5 | 1015.7 | 483.8 |
| 122 | 4354199 | 498.2 | 1302.6 | 634.3 | 450.2 | 237.1 | 898.9 | 466.0 |
| 123 | 4355373 | 659.5 | 1611.5 | 755.9 | 594.2 | 280.1 | 1203.2 | 657.3 |
| 124 | 4382169 | 1136.8 | 39.3 | 1398.4 | 880.7 | 232.5 | 735.6 | 979.2 |
| 125 | 4400553 | 613.8 | 1442.0 | 750.1 | 518.2 | 302.6 | 1118.8 | 567.3 |
| 126 | 4400847 | 613.1 | 1405.4 | 897.9 | 578.3 | 363.1 | 1110.2 | 538.0 |
| 127 | 4402913 | 456.2 | 1352.4 | 588.0 | 452.5 | 221.7 | 883.4 | 430.6 |
| 128 | 4405414 | 534.5 | 1072.4 | 659.1 | 573.4 | 223.7 | 808.5 | 465.3 |
| 129 | 4425570 | 7939.2 | 8.7 | 3189.2 | 6100.9 | 393.9 | 3470.3 | 4751.6 |
| 130 | 4576933 | 428.1 | 811.0 | 568.0 | 444.9 | 253.4 | 654.0 | 423.0 |
| 131 | 4587198 | 543.3 | 963.7 | 557.5 | 486.1 | 233.3 | 700.2 | 486.4 |
| 132 | 4587495 | 631.1 | 1261.4 | 651.1 | 537.6 | 231.8 | 914.8 | 587.4 |
| 133 | 4752954 | 563.2 | 1294.7 | 650.9 | 537.6 | 230.7 | 950.3 | 554.1 |
| 134 | 4753616 | 539.0 | 1431.8 | 707.2 | 507.9 | 343.5 | 963.8 | 515.3 |
| 135 | 4754176 | 578.5 | 1480.2 | 723.0 | 586.9 | 297.5 | 1001.7 | 523.9 |
| 136 | 4754652 | 609.7 | 1406.1 | 706.1 | 560.8 | 355.0 | 929.2 | 559.5 |
| 137 | 4754878 | 632.4 | 1389.6 | 738.0 | 544.3 | 337.7 | 908.1 | 595.3 |
| 138 | 4755307 | 580.7 | 1512.1 | 738.7 | 475.1 | 294.5 | 1001.9 | 564.9 |
| 139 | 4755608 | 414.9 | 951.9 | 461.7 | 381.2 | 143.4 | 699.8 | 406.7 |
| 140 | 5159548 | 881.2 | 12.7 | 236.3 | 106.4 | 62.3 | 66.0 | 558.3 |
| 141 | 5219733 | 912.2 | 1442.4 | 534.7 | 541.4 | 635.0 | 879.4 | 910.6 |
| 142 | 5233092 | 662.5 | 1203.9 | 746.0 | 614.7 | 306.3 | 819.3 | 610.2 |
| 143 | 5236558 | 491.5 | 1006.2 | 606.4 | 488.4 | 220.0 | 777.4 | 445.1 |
| 144 | 5236864 | 471.7 | 781.1 | 525.1 | 490.1 | 214.4 | 540.0 | 380.3 |
| 145 | 5236985 | 487.4 | 762.7 | 493.8 | 454.9 | 237.7 | 499.6 | 409.2 |
| 146 | 5268881 | 679.5 | 29.4 | 247.1 | 185.2 | 60.9 | 150.2 | 566.6 |
| 147 | 5270523 | 434.3 | 693.4 | 476.1 | 374.1 | 211.7 | 496.6 | 377.5 |
| 148 | 5275584 | 632.2 | 1362.5 | 740.9 | 609.0 | 386.2 | 944.8 | 619.3 |
| 149 | 5276703 | 465.9 | 1107.3 | 905.5 | 558.8 | 241.1 | 742.3 | 452.2 |
| 150 | 5291283 | 439.0 | 915.8 | 578.8 | 470.8 | 255.2 | 623.9 | 443.2 |
| 151 | 5300407 | 536.5 | 1074.0 | 588.7 | 466.1 | 254.4 | 837.3 | 528.0 |
| 152 | 5300796 | 654.3 | 1528.1 | 733.0 | 577.7 | 269.5 | 961.0 | 644.1 |
| 153 | 5301027 | 521.8 | 1146.9 | 568.7 | 524.4 | 225.8 | 799.6 | 498.9 |
| 154 | 5305066 | 596.3 | 1400.2 | 761.5 | 579.3 | 320.6 | 870.8 | 596.4 |
| 155 | 5330676 | 557.9 | 1243.0 | 762.2 | 521.6 | 332.2 | 947.2 | 521.0 |
| 156 | 5331461 | 598.2 | 1245.1 | 941.0 | 679.1 | 373.4 | 919.2 | 548.3 |
| 157 | 5331638 | 611.9 | 1201.4 | 852.4 | 648.7 | 398.2 | 794.7 | 540.9 |
| 158 | 5352651 | 427.5 | 707.7 | 483.5 | 440.1 | 218.4 | 490.2 | 381.7 |
| 159 | 5352994 | 525.2 | 1181.7 | 681.5 | 637.8 | 203.9 | 882.8 | 482.3 |
| 160 | 5403392 | 3058.5 | 6709.5 | 3102.8 | 3161.8 | 4584.5 | 3222.5 | 2620.0 |
| 161 | 5414219 | 967.4 | 62.9 | 508.3 | 620.9 | 186.0 | 644.1 | 1090.7 |
| 162 | 5515082 | 369.2 | 632.3 | 186.3 | 225.1 | 165.8 | 301.8 | 347.1 |
| 163 | 5996355 | 536.5 | 1225.5 | 824.3 | 550.2 | 292.3 | 796.0 | 504.9 |
| 164 | 5996882 | 416.3 | 982.0 | 826.9 | 513.5 | 184.6 | 677.8 | 450.1 |
| 165 | 5998097 | 509.8 | 1009.1 | 746.2 | 555.3 | 300.6 | 698.2 | 520.2 |
| 166 | 5999094 | 567.7 | 1448.3 | 910.4 | 678.5 | 249.7 | 1114.7 | 518.7 |
| 167 | 6082366 | 10477.0 | 23.2 | 1291.5 | 4067.5 | 120.7 | 1507.1 | 6109.5 |
| 168 | 6093032 | 3285.2 | 16.5 | 205.1 | 333.9 | 83.0 | 269.2 | 2756.2 |
| 169 | 6199144 | 472.3 | 607.7 | 468.8 | 364.8 | 355.0 | 517.5 | 432.8 |

[a]Site Location refers to the genomic position of the base in the center of a given ChIP peak.

Table S2

| | A | B | C | D | E | F | G | H | I | J | K | L | M | N |
|---|---|---|---|---|---|---|---|---|---|---|---|---|---|---|
| 1 | **Associated Binding Site[a]** | **Gene Start[b]** | **Gene End[c]** | **Strand[d]** | **Name** | **Synonym** | **Product** | **Expression WT** | **Expression Δ*rhlR*** | **Expression Δ*rhlI*** | **Expression Δ*pqsE*** | **Expression *pqsE*- NI** | **Expression *pqsE* (D73A)** | **qValue WT vs Δ*rhlR*** |
| 2 | - | 483 | 2027 | + | *dnaA* | *PA14_00010* | chromosomal replication initiation protein | 257 | 240 | 252 | 245 | 233 | 229 | 1 |
| 3 | - | 2056 | 3159 | + | *dnaN* | *PA14_00020* | DNA polymerase III subunit beta | 194 | 256 | 197 | 192 | 202 | 190 | 1 |
| 4 | - | 3169 | 4278 | + | *recF* | *PA14_00030* | recombination protein F | 251 | 223 | 304 | 214 | 235 | 217 | 1 |
| 5 | - | 4275 | 6695 | + | *gyrB* | *PA14_00050* | DNA gyrase subunit B | 244 | 282 | 219 | 216 | 257 | 222 | 1 |
| 6 | - | 7791 | 7018 | - | - | *PA14_00060* | acyltransferase | 39 | 36 | 39 | 32 | 33 | 35 | 1 |
| 7 | - | 8339 | 7803 | - | - | *PA14_00070* | D,D-heptose 1,7-bisphosphate phosphatase | 41 | 32 | 52 | 33 | 35 | 35 | 1 |
| 8 | - | 8671 | 10377 | + | - | *PA14_00080* | hypothetical protein | 110 | 85 | 117 | 119 | 97 | 110 | 1 |
| 9 | - | 12488 | 10434 | - | *glyS* | *PA14_00090* | glycyl-tRNA synthetase subunit beta | 67 | 95 | 68 | 68 | 77 | 50 | 1 |
| 10 | - | 13435 | 12488 | - | *glyQ* | *PA14_00100* | glycyl-tRNA synthetase subunit alpha | 45 | 60 | 42 | 47 | 51 | 33 | 1 |
| 11 | - | 13540 | 14091 | + | *tag* | *PA14_00110* | DNA-3-methyladenine glycosidase I | 16 | 16 | 16 | 14 | 16 | 13 | 1 |
| 12 | - | 14236 | 15123 | + | - | *PA14_00120* | lipid A biosynthesis lauroyl acyltransferase | 51 | 50 | 62 | 55 | 63 | 52 | 1 |
| 13 | - | 15208 | 15474 | + | - | *PA14_00130* | hypothetical protein | 44 | 47 | 59 | 45 | 47 | 44 | 1 |
| 14 | - | 15621 | 16274 | + | - | *PA14_00140* | hypothetical protein | 11 | 14 | 13 | 10 | 9 | 9 | 1 |
| 15 | - | 16608 | 16336 | - | - | *PA14_00150* | hypothetical protein | 6 | 9 | 8 | 5 | 6 | 4 | 1 |
| 16 | - | 17240 | 16923 | - | - | *PA14_00160* | hypothetical protein | 130 | 123 | 141 | 124 | 132 | 139 | 1 |
| 17 | - | 18762 | 17389 | - | *trkA* | *PA14_00170* | potassium transporter peripheral membrane protein | 87 | 85 | 81 | 73 | 83 | 70 | 1 |
| 18 | - | 20093 | 18789 | - | - | *PA14_00180* | tRNA and rRNA cytosine-C5-methylases | 67 | 68 | 68 | 54 | 62 | 53 | 1 |
| 19 | - | 21034 | 20090 | - | *fmt* | *PA14_00190* | methionyl-tRNA formyltransferase | 70 | 58 | 66 | 50 | 63 | 52 | 1 |
| 20 | - | 21594 | 21088 | - | *def* | *PA14_00200* | peptide deformylase | 342 | 301 | 343 | 220 | 259 | 288 | 1 |
| 21 | - | 21733 | 22758 | + | - | *PA14_00210* | lysin domain-containing protein | 71 | 95 | 76 | 94 | 85 | 66 | 1 |
| 22 | - | 22893 | 23981 | + | - | *PA14_00230* | Rossmann fold nucleotide-binding protein | 9 | 13 | 9 | 12 | 10 | 9 | 1 |
| 23 | - | 24022 | 24579 | + | - | *PA14_00240* | hypothetical protein | 71 | 81 | 74 | 78 | 69 | 70 | 1 |
| 24 | - | 25566 | 24589 | - | *qor* | *PA14_00250* | quinone oxidoreductase | 73 | 85 | 67 | 77 | 73 | 64 | 1 |
| 25 | - | 25757 | 26674 | + | *hemF* | *PA14_00280* | coproporphyrinogen III oxidase | 81 | 95 | 74 | 111 | 192 | 149 | 1 |
| 26 | - | 26732 | 27556 | + | *aroE* | *PA14_00290* | shikimate 5-dehydrogenase | 74 | 98 | 81 | 106 | 146 | 121 | 1 |
| 27 | - | 27667 | 28653 | + | *plcB* | *PA14_00300* | phospholipase C, PlcB | 80 | 132 | 110 | 139 | 115 | 100 | 1 |
| 28 | - | 28634 | 29920 | + | - | *PA14_00310* | peptidyl-prolyl isomerase | 194 | 304 | 286 | 370 | 262 | 238 | 1 |
| 29 | - | 29917 | 30519 | + | - | *PA14_00320* | hypothetical protein | 112 | 201 | 148 | 221 | 157 | 149 | 0.87182855 |
| 30 | - | 32076 | 30523 | - | - | *PA14_00340* | sulfate transporter | 36 | 61 | 49 | 82 | 57 | 57 | 0.853730745 |
| 31 | - | 33004 | 32081 | - | - | *PA14_00360* | hypothetical protein | 8 | 7 | 7 | 9 | 7 | 7 | 1 |
| 32 | - | 34532 | 33021 | - | *betC* | *PA14_00380* | choline sulfatase | 21 | 19 | 23 | 22 | 17 | 19 | 1 |
| 33 | - | 34645 | 35559 | + | - | *PA14_00400* | LysR family transcriptional regulator | 18 | 19 | 18 | 16 | 16 | 15 | 1 |
| 34 | - | 35570 | 35947 | + | - | *PA14_00410* | dioxygenase | 131 | 118 | 109 | 97 | 112 | 103 | 1 |
| 35 | - | 36291 | 35926 | - | - | *PA14_00420* | histidine phosphotransfer domain-containing protein | 51 | 51 | 51 | 41 | 45 | 39 | 1 |
| 36 | - | 36922 | 36299 | - | - | *PA14_00430* | two-component response regulator | 31 | 23 | 27 | 24 | 24 | 23 | 1 |
| 37 | - | 37914 | 37108 | - | *trpA* | *PA14_00440* | tryptophan synthase subunit alpha | 55 | 49 | 42 | 43 | 45 | 43 | 1 |
| 38 | - | 39119 | 37911 | - | *trpB* | *PA14_00450* | tryptophan synthase subunit beta | 80 | 64 | 63 | 63 | 64 | 63 | 1 |
| 39 | - | 39223 | 40110 | + | *trpI* | *PA14_00460* | transcriptional regulator TrpI | 19 | 17 | 16 | 15 | 14 | 14 | 1 |
| 40 | - | 40211 | 40426 | + | - | *PA14_00470* | hypothetical protein | 773 | 957 | 1090 | 933 | 1030 | 1009 | 1 |
| 41 | - | 40453 | 40609 | ? | - | predicted RNA | - | 1520 | 2020 | 1667 | 1909 | 1651 | 1921 | 1 |
| 42 | - | 40610 | 40837 | + | - | *PA14_00480* | hypothetical protein | 2080 | 2747 | 2724 | 3329 | 2669 | 3108 | 1 |
| 43 | - | 41133 | 42821 | + | - | *PA14_00490* | hemolysin activation/secretion protein | 26 | 25 | 28 | 26 | 25 | 21 | 1 |
| 44 | - | 42934 | 53265 | + | - | *PA14_00510* | hemagglutinin | 36 | 29 | 31 | 25 | 30 | 24 | 1 |
| 45 | - | 54577 | 54699 | + | - | *PA14_00520* | hypothetical protein | 32 | 27 | 22 | 31 | 43 | 39 | 1 |
| 46 | - | 54909 | 55304 | + | - | *PA14_00530* | hypothetical protein | 86 | 77 | 83 | 107 | 70 | 76 | 1 |
| 47 | - | 56984 | 55575 | - | - | *PA14_00550* | hypothetical protein | 18 | 16 | 17 | 14 | 17 | 14 | 1 |
| 48 | - | 57148 | 58476 | + | *exoT* | *PA14_00560* | exoenzyme T | 176 | 174 | 208 | 194 | 168 | 183 | 1 |
| 49 | - | 59018 | 59704 | + | - | *PA14_00570* | lipoprotein | 12 | 17 | 13 | 10 | 14 | 8 | 1 |
| 50 | - | 59735 | 60088 | + | - | *PA14_00580* | lipoprotein | 16 | 24 | 17 | 14 | 26 | 12 | 1 |
| 51 | - | 60241 | 60750 | + | - | *PA14_00590* | lipoprotein | 17 | 19 | 16 | 13 | 18 | 11 | 1 |
| 52 | - | 61148 | 60765 | - | - | *PA14_00600* | transcriptional regulator | 9 | 15 | 12 | 9 | 14 | 6 | 1 |
| 53 | - | 63091 | 61430 | - | - | *PA14_00620* | hypothetical protein | 6 | 12 | 8 | 12 | 11 | 9 | 0.258262081 |
| 54 | - | 63701 | 63841 | + | - | *PA14_00630* | hypothetical protein | 131 | 82 | 103 | 136 | 115 | 161 | 1 |
| 55 | 64211 | 64665 | 66497 | + | *phzH* | *PA14_00640* | potential phenazine-modifying enzyme | 63 | 10 | 10 | 20 | 20 | 57 | 3.40517E-17 |
| 56 | - | 66978 | 66550 | - | - | *PA14_00650* | hypothetical protein | 105 | 83 | 96 | 135 | 110 | 209 | 1 |
| 57 | - | 67735 | 67187 | - | - | *PA14_00660* | RNA 2'-phosphotransferase-like protein | 17 | 16 | 12 | 12 | 20 | 12 | 1 |
| 58 | - | 68280 | 67774 | - | - | *PA14_00670* | hypothetical protein | 45 | 52 | 34 | 38 | 40 | 32 | 1 |
| 59 | - | 69266 | 68346 | - | - | *PA14_00680* | LysR family transcriptional regulator | 10 | 10 | 9 | 10 | 10 | 9 | 1 |
| 60 | - | 69374 | 70261 | + | - | *PA14_00690* | hypothetical protein | 10 | 6 | 8 | 8 | 7 | 9 | 0.41668339 |
| 61 | - | 70324 | 71028 | + | - | *PA14_00700* | hypothetical protein | 7 | 7 | 7 | 8 | 6 | 8 | 1 |
| 62 | - | 71112 | 71567 | + | *osmC* | *PA14_00710* | osmotically inducible protein OsmC | 131 | 178 | 140 | 165 | 194 | 180 | 1 |
| 63 | - | 71678 | 71911 | + | - | *PA14_00720* | hypothetical protein | 93 | 118 | 105 | 107 | 148 | 145 | 1 |
| 64 | - | 72360 | 71923 | - | - | *PA14_00730* | hypothetical protein | 141 | 115 | 151 | 125 | 134 | 126 | 1 |
| 65 | - | 72833 | 72417 | - | - | *PA14_00740* | lipoprotein | 24 | 28 | 31 | 32 | 29 | 24 | 1 |
| 66 | - | 72925 | 74052 | + | - | *PA14_00750* | hypothetical protein | 75 | 69 | 68 | 68 | 64 | 70 | 1 |
| 67 | - | 75043 | 74060 | - | - | *PA14_00760* | hypothetical protein | 82 | 70 | 65 | 73 | 65 | 63 | 1 |
| 68 | - | 75741 | 75076 | - | - | *PA14_00770* | hypothetical protein | 48 | 50 | 48 | 49 | 51 | 46 | 1 |
| 69 | - | 76276 | 75734 | - | - | *PA14_00780* | hypothetical protein | 82 | 81 | 78 | 67 | 71 | 71 | 1 |
| 70 | - | 76354 | 78399 | + | *prlC* | *PA14_00790* | oligopeptidase A | 216 | 174 | 156 | 142 | 166 | 186 | 1 |
| 71 | - | 78396 | 78671 | + | - | *PA14_00800* | hypothetical protein | 117 | 123 | 81 | 92 | 107 | 102 | 1 |
| 72 | - | 78760 | 79818 | + | - | *PA14_00810* | DNA repair photolyase | 18 | 17 | 17 | 18 | 19 | 22 | 1 |
| 73 | - | 80963 | 80049 | - | - | *PA14_00820* | hypothetical protein | 130 | 136 | 79 | 46 | 122 | 73 | 1 |
| 74 | - | 82737 | 81025 | - | - | *PA14_00830* | hypothetical protein | 63 | 51 | 46 | 41 | 54 | 48 | 1 |
| 75 | - | 83929 | 82730 | - | - | *PA14_00850* | hypothetical protein | 32 | 20 | 21 | 14 | 24 | 18 | 1 |
| 76 | - | 84648 | 83929 | - | - | *PA14_00860* | ABC transporter ATP-binding protein | 40 | 24 | 21 | 16 | 28 | 24 | 0.925024982 |
| 77 | - | 87743 | 84645 | - | *ppkA* | *PA14_00875* | serine/threonine protein kinase PpkA | 58 | 40 | 36 | 33 | 54 | 45 | 1 |
| 78 | - | 88479 | 87751 | - | - | *PA14_00890* | phosphoprotein phosphatase | 59 | 38 | 44 | 27 | 54 | 45 | 1 |
| 79 | - | 89169 | 88489 | - | - | *PA14_00900* | hypothetical protein | 59 | 40 | 40 | 28 | 50 | 46 | 1 |
| 80 | - | 92474 | 89166 | - | - | *PA14_00910* | hypothetical protein | 62 | 40 | 40 | 36 | 48 | 45 | 1 |
| 81 | - | 94045 | 92696 | - | - | *PA14_00925* | hypothetical protein | 81 | 54 | 62 | 54 | 60 | 54 | 1 |
| 82 | - | 95386 | 94052 | - | - | *PA14_00940* | hypothetical protein | 60 | 47 | 54 | 46 | 51 | 45 | 1 |
| 83 | - | 95866 | 95402 | - | - | *PA14_00960* | lipoprotein | 37 | 32 | 27 | 24 | 30 | 24 | 1 |
| 84 | - | 96453 | 95911 | - | - | *PA14_00970* | hypothetical protein | 95 | 90 | 78 | 74 | 70 | 81 | 1 |

| | A | B | C | D | E | F | G | H | I | J | K | L | M | N |
|---|---|---|---|---|---|---|---|---|---|---|---|---|---|---|
| 85 | - | 97409 | 96450 | - | - | *PA14_00980* | hypothetical protein | 79 | 71 | 68 | 82 | 81 | 80 | 1 |
| 86 | - | 97777 | 98811 | + | - | *PA14_00990* | hypothetical protein | 42 | 25 | 33 | 22 | 24 | 19 | 1 |
| 87 | - | 98900 | 99418 | + | - | *PA14_01010* | hypothetical protein | 186 | 137 | 126 | 117 | 195 | 107 | 1 |
| 88 | - | 99431 | 100927 | + | - | *PA14_01020* | hypothetical protein | 103 | 85 | 70 | 55 | 94 | 57 | 1 |
| 89 | - | 101003 | 101491 | + | - | *PA14_01030* | hypothetical protein | 324 | 239 | 180 | 116 | 280 | 158 | 1 |
| 90 | - | 101659 | 102504 | + | - | *PA14_01040* | secretion protein | 64 | 38 | 44 | 28 | 47 | 34 | 1 |
| 91 | - | 102506 | 103015 | + | - | *PA14_01060* | hypothetical protein | 24 | 18 | 19 | 16 | 23 | 17 | 1 |
| 92 | - | 103012 | 104871 | + | - | *PA14_01070* | hypothetical protein | 24 | 19 | 18 | 13 | 22 | 15 | 1 |
| 93 | - | 104835 | 105881 | + | - | *PA14_01080* | hypothetical protein | 27 | 16 | 17 | 12 | 27 | 14 | 0.803084159 |
| 94 | - | 105874 | 108582 | + | - | *PA14_01100* | ClpA/B-type chaperone | 59 | 37 | 37 | 29 | 57 | 35 | 1 |
| 95 | - | 108629 | 110560 | + | - | *PA14_01110* | hypothetical protein | 43 | 27 | 24 | 19 | 37 | 23 | 1 |
| 96 | - | 110959 | 110675 | - | - | *PA14_01120* | hypothetical protein | 50 | 53 | 37 | 36 | 57 | 35 | 1 |
| 97 | - | 111205 | 110972 | - | - | *PA14_01130* | hypothetical protein | 72 | 69 | 65 | 61 | 95 | 69 | 1 |
| 98 | - | 112514 | 111195 | - | - | *PA14_01140* | hypothetical protein | 27 | 23 | 24 | 19 | 23 | 16 | 1 |
| 99 | - | 113027 | 112593 | - | - | *PA14_01150* | hypothetical protein | 28 | 27 | 24 | 28 | 27 | 24 | 1 |
| 100 | - | 113281 | 115506 | + | - | *PA14_01160* | hypothetical protein | 83 | 70 | 87 | 89 | 79 | 75 | 1 |
| 101 | - | 115534 | 115983 | + | - | *PA14_01170* | hypothetical protein | 44 | 34 | 33 | 31 | 34 | 34 | 1 |
| 102 | - | 115913 | 117112 | + | - | *PA14_01180* | hypothetical protein | 24 | 20 | 22 | 20 | 25 | 19 | 1 |
| 103 | - | 117109 | 118146 | + | - | *PA14_01190* | 3-oxoacyl-ACP synthase | 28 | 23 | 24 | 20 | 26 | 19 | 1 |
| 104 | - | 118146 | 119237 | + | - | *PA14_01200* | hypothetical protein | 43 | 33 | 38 | 33 | 41 | 34 | 1 |
| 105 | - | 119248 | 120168 | + | - | *PA14_01220* | hypothetical protein | 81 | 57 | 63 | 53 | 70 | 61 | 1 |
| 106 | - | 120178 | 121410 | + | - | *PA14_01230* | hypothetical protein | 37 | 27 | 27 | 23 | 27 | 24 | 1 |
| 107 | - | 121786 | 122514 | + | - | *PA14_01240* | carbonic anhydrase | 103 | 107 | 86 | 80 | 98 | 88 | 1 |
| 108 | - | 122725 | 124296 | + | - | *PA14_01250* | sulfate transporter | 17 | 16 | 19 | 21 | 18 | 16 | 1 |
| 109 | - | 124433 | 125029 | + | - | *PA14_01270* | hypothetical protein | 13 | 15 | 19 | 15 | 14 | 12 | 1 |
| 110 | - | 125293 | 126417 | + | *coxB* | *PA14_01290* | cytochrome c oxidase subunit II | 133 | 95 | 185 | 117 | 68 | 106 | 1 |
| 111 | - | 126427 | 128019 | + | *coxA* | *PA14_01300* | cytochrome c oxidase subunit I | 73 | 49 | 68 | 71 | 42 | 75 | 1 |
| 112 | - | 128030 | 128584 | + | - | *PA14_01310* | cytochrome C oxidase assembly protein | 45 | 37 | 58 | 68 | 36 | 68 | 1 |
| 113 | - | 128595 | 129482 | + | *colII* | *PA14_01320* | cytochrome c oxidase subunit III | 161 | 166 | 209 | 207 | 164 | 238 | 1 |
| 114 | - | 129707 | 129498 | - | - | *PA14_01330* | hypothetical protein | 432 | 489 | 604 | 646 | 666 | 715 | 1 |
| 115 | - | 129723 | 130517 | + | - | *PA14_01340* | hypothetical protein | 31 | 24 | 40 | 33 | 23 | 32 | 1 |
| 116 | - | 130492 | 131070 | + | - | *PA14_01350* | hypothetical protein | 18 | 18 | 24 | 25 | 14 | 26 | 1 |
| 117 | - | 131135 | 132208 | + | - | *PA14_01360* | hypothetical protein | 18 | 15 | 16 | 21 | 12 | 21 | 1 |
| 118 | - | 132234 | 133148 | + | - | *PA14_01380* | protoheme IX farnesyltransferase | 40 | 39 | 44 | 48 | 40 | 54 | 1 |
| 119 | - | 133174 | 133809 | + | - | *PA14_01390* | hypothetical protein | 106 | 73 | 101 | 87 | 104 | 105 | 1 |
| 120 | - | 134301 | 133849 | - | - | *PA14_01400* | hypothetical protein | 33 | 37 | 35 | 30 | 36 | 28 | 1 |
| 121 | - | 134433 | 134906 | + | - | *PA14_01410* | hypothetical protein | 75 | 88 | 82 | 91 | 77 | 89 | 1 |
| 122 | - | 135162 | 135890 | + | - | *PA14_01430* | short chain dehydrogenase | 13 | 10 | 10 | 11 | 10 | 12 | 1 |
| 123 | - | 135915 | 136502 | + | - | *PA14_01440* | hypothetical protein | 15 | 17 | 11 | 12 | 13 | 14 | 1 |
| 124 | - | 136732 | 138081 | + | - | *PA14_01460* | C4-dicarboxylate transporter DctA | 10 | 14 | 9 | 10 | 13 | 10 | 1 |
| 125 | - | 138130 | 138816 | + | - | *PA14_01470* | transcriptional regulator | 68 | 58 | 50 | 58 | 60 | 60 | 1 |
| 126 | - | 138917 | 139651 | + | - | *PA14_01480* | hypothetical protein | 22 | 25 | 18 | 20 | 18 | 19 | 1 |
| 127 | 139733 | 139831 | 140241 | + | - | *PA14_01490* | hemolysin | 2956 | 160 | 1024 | 2108 | 1401 | 3762 | 2.0605E-123 |
| 128 | - | 141181 | 140273 | - | - | *PA14_01500* | transcriptional regulator | 189 | 30 | 58 | 127 | 84 | 196 | 2.46944E-13 |
| 129 | - | 141762 | 141481 | - | - | *PA14_01510* | hypothetical protein | 11 | 8 | 8 | 7 | 7 | 7 | 1 |
| 130 | - | 141986 | 141759 | - | - | *PA14_01520* | hypothetical protein | 10 | 8 | 9 | 9 | 9 | 10 | 1 |
| 131 | - | 142785 | 142162 | - | - | *PA14_01540* | hypothetical protein | 16 | 17 | 13 | 8 | 14 | 9 | 1 |
| 132 | - | 142886 | 143386 | + | - | *PA14_01550* | lipoprotein | 25 | 31 | 27 | 24 | 27 | 19 | 1 |
| 133 | - | 143458 | 143799 | + | - | *PA14_01560* | hypothetical protein | 98 | 73 | 154 | 58 | 48 | 59 | 1 |
| 134 | - | 145308 | 143881 | - | *gabP* | *PA14_01580* | gamma-aminobutyrate permease | 39 | 55 | 47 | 60 | 55 | 66 | 1 |
| 135 | - | 146970 | 145477 | - | - | *PA14_01600* | aldehyde dehydrogenase | 263 | 323 | 268 | 377 | 335 | 401 | 1 |
| 136 | - | 147341 | 147054 | - | - | *PA14_01610* | hypothetical protein | 96 | 171 | 173 | 290 | 220 | 343 | 0.605197743 |
| 137 | - | 148687 | 147341 | - | *aptA* | *PA14_01620* | beta alanine--pyruvate transaminase | 161 | 266 | 260 | 396 | 349 | 569 | 1 |
| 138 | - | 148823 | 149740 | + | - | *PA14_01640* | LysR family transcriptional regulator | 17 | 20 | 19 | 22 | 18 | 23 | 1 |
| 139 | - | 151223 | 149853 | - | - | *PA14_01660* | guanine deaminase | 52 | 45 | 45 | 48 | 50 | 51 | 1 |
| 140 | - | 152381 | 153952 | + | - | *PA14_01670* | ABC transporter ATP-binding protein | 12 | 10 | 13 | 13 | 11 | 12 | 1 |
| 141 | - | 153952 | 155049 | + | - | *PA14_01680* | ABC transporter permease | 4 | 4 | 3 | 4 | 5 | 4 | 1 |
| 142 | - | 155072 | 156052 | + | - | *PA14_01690* | ABC transporter permease | 16 | 18 | 18 | 15 | 16 | 15 | 1 |
| 143 | - | 156164 | 156727 | + | *ahpC* | *PA14_01710* | alkyl hydroperoxide reductase | 1247 | 649 | 297 | 276 | 440 | 1483 | 1 |
| 144 | - | 156872 | 158437 | + | *ahpF* | *PA14_01720* | alkyl hydroperoxide reductase | 288 | 98 | 98 | 59 | 74 | 280 | 0.579361359 |
| 145 | - | 159413 | 158517 | - | - | *PA14_01730* | hypothetical protein | 562 | 683 | 744 | 382 | 542 | 498 | 1 |
| 146 | - | 159868 | 161217 | + | - | *PA14_01750* | hydroxydechloroatrazine ethylaminohydrolase | 19 | 31 | 24 | 29 | 30 | 22 | 0.952168061 |
| 147 | - | 161387 | 162376 | + | *nuh* | *PA14_01760* | nonspecific ribonucleoside hydrolase | 79 | 97 | 98 | 103 | 90 | 88 | 1 |
| 148 | - | 163261 | 162413 | - | - | *PA14_01770* | nucleoside-binding outer membrane protein | 35 | 56 | 47 | 59 | 47 | 41 | 0.944453713 |
| 149 | - | 163922 | 163296 | - | - | *PA14_01780* | nucleoside 2-deoxyribosyltransferase | 85 | 136 | 133 | 158 | 116 | 117 | 0.940967938 |
| 150 | - | 164590 | 164072 | - | - | *PA14_01790* | nucleoside 2-deoxyribosyltransferase | 23 | 26 | 28 | 23 | 23 | 22 | 1 |
| 151 | - | 164828 | 165925 | + | - | *PA14_01800* | hypothetical protein | 5 | 7 | 6 | 7 | 4 | 5 | 1 |
| 152 | - | 165998 | 166960 | + | - | *PA14_01810* | oxidoreductase | 23 | 21 | 22 | 22 | 21 | 23 | 1 |
| 153 | - | 167065 | 168015 | + | - | *PA14_01830* | adenosine deaminase | 33 | 33 | 36 | 34 | 26 | 29 | 1 |
| 154 | - | 168212 | 168757 | + | - | *PA14_01840* | RNA polymerase ECF-subfamily sigma-70 factor | 17 | 22 | 21 | 44 | 29 | 18 | 1 |
| 155 | - | 168754 | 169749 | + | - | *PA14_01860* | transmembrane sensor | 11 | 15 | 12 | 18 | 15 | 8 | 1 |
| 156 | - | 169897 | 172284 | + | - | *PA14_01870* | TonB-dependent receptor | 7 | 6 | 7 | 9 | 7 | 5 | 1 |
| 157 | - | 172662 | 173489 | + | *pcaQ* | *PA14_01890* | transcriptional regulator PcaQ | 39 | 38 | 44 | 32 | 37 | 38 | 1 |
| 158 | - | 173624 | 174343 | + | *pcaH* | *PA14_01900* | protocatechuate 3,4-dioxygenase subunit beta | 10 | 11 | 11 | 11 | 9 | 10 | 1 |
| 159 | - | 174354 | 174959 | + | *pcaG* | *PA14_01910* | protocatechuate 3,4-dioxygenase subunit alpha | 11 | 13 | 14 | 12 | 12 | 14 | 1 |
| 160 | - | 175175 | 176014 | + | *pcaR* | *PA14_01930* | transcriptional regulator PcaR | 14 | 14 | 14 | 15 | 13 | 16 | 1 |
| 161 | - | 176167 | 177318 | + | - | *PA14_01940* | RND efflux membrane fusion protein | 97 | 113 | 124 | 155 | 113 | 106 | 1 |
| 162 | - | 177315 | 178385 | + | - | *PA14_01960* | RND efflux membrane fusion protein | 84 | 124 | 114 | 145 | 115 | 107 | 1 |
| 163 | - | 178382 | 181429 | + | - | *PA14_01970* | RND efflux transporter | 120 | 178 | 173 | 207 | 180 | 173 | 1 |
| 164 | - | 181628 | 182566 | + | - | *PA14_01980* | LysR family transcriptional regulator | 45 | 52 | 51 | 49 | 53 | 51 | 1 |
| 165 | - | 182682 | 182867 | + | - | *PA14_01990* | hypothetical protein | 31 | 33 | 29 | 29 | 26 | 28 | 1 |
| 166 | - | 183147 | 183299 | + | - | *PA14_02010* | hypothetical protein | 14 | 6 | 7 | 6 | 8 | 5 | 0.25264025 |
| 167 | - | 183454 | 184788 | + | - | *PA14_02020* | outer membrane porin | 32 | 34 | 28 | 26 | 30 | 25 | 1 |
| 168 | - | 185614 | 184817 | - | - | *PA14_02030* | AraC family transcriptional regulator | 19 | 19 | 14 | 17 | 21 | 17 | 1 |

| | A | B | C | D | E | F | G | H | I | J | K | L | M | N |
|---|---|---|---|---|---|---|---|---|---|---|---|---|---|---|
| 169 | - | 185692 | 187308 | + | - | PA14_02050 | gamma-glutamyltranspeptidase | 30 | 21 | 31 | 18 | 22 | 24 | 1 |
| 170 | - | 187980 | 188816 | + | - | PA14_02060 | hypothetical protein | 5 | 7 | 6 | 5 | 5 | 5 | 1 |
| 171 | - | 189058 | 190464 | + | - | PA14_02070 | transporter | 10 | 13 | 10 | 10 | 9 | 9 | 1 |
| 172 | - | 190557 | 191222 | + | - | PA14_02090 | TetR family transcriptional regulator | 39 | 49 | 39 | 29 | 36 | 26 | 1 |
| 173 | - | 191228 | 191818 | + | - | PA14_02100 | hypothetical protein | 116 | 130 | 100 | 105 | 127 | 105 | 1 |
| 174 | - | 192532 | 191825 | - | - | PA14_02110 | diguanylate cyclase | 43 | 40 | 43 | 44 | 46 | 44 | 1 |
| 175 | - | 193039 | 192659 | - | - | PA14_02130 | hypothetical protein | 10 | 9 | 10 | 8 | 8 | 8 | 1 |
| 176 | - | 193608 | 193066 | - | - | PA14_02140 | hypothetical protein | 6 | 6 | 6 | 7 | 5 | 3 | 1 |
| 177 | - | 195608 | 193617 | - | - | PA14_02150 | hypothetical protein | 11 | 9 | 11 | 11 | 8 | 9 | 1 |
| 178 | - | 196920 | 195871 | - | cheB | PA14_02180 | chemotaxis-specific methylesterase | 29 | 28 | 36 | 40 | 31 | 48 | 1 |
| 179 | - | 197542 | 196940 | - | - | PA14_02190 | hypothetical protein | 47 | 41 | 55 | 52 | 56 | 70 | 1 |
| 180 | - | 198390 | 197548 | - | - | PA14_02200 | chemotaxis protein methyltransferase | 83 | 88 | 108 | 104 | 102 | 146 | 1 |
| 181 | - | 200499 | 198460 | - | - | PA14_02220 | chemotaxis transducer | 406 | 421 | 485 | 568 | 490 | 659 | 1 |
| 182 | - | 201020 | 200535 | - | cheW | PA14_02230 | purine-binding chemotaxis protein | 316 | 318 | 387 | 473 | 376 | 504 | 1 |
| 183 | - | 202926 | 201007 | - | cheA | PA14_02250 | two-component sensor | 252 | 225 | 295 | 300 | 261 | 324 | 1 |
| 184 | - | 203319 | 202954 | - | - | PA14_02260 | two-component response regulator | 753 | 756 | 1089 | 880 | 863 | 997 | 1 |
| 185 | - | 204689 | 203517 | - | - | PA14_02270 | chemotaxis transducer | 106 | 96 | 133 | 137 | 101 | 121 | 1 |
| 186 | - | 205815 | 204880 | - | - | PA14_02290 | LysR family transcriptional regulator | 42 | 38 | 36 | 44 | 40 | 38 | 1 |
| 187 | - | 205932 | 206684 | + | fabG | PA14_02300 | 3-ketoacyl-ACP reductase | 16 | 16 | 13 | 15 | 17 | 11 | 1 |
| 188 | - | 208394 | 206784 | - | atsA | PA14_02310 | arylsulfatase | 7 | 8 | 6 | 6 | 7 | 6 | 1 |
| 189 | - | 209321 | 208482 | - | - | PA14_02330 | ABC transporter ATP-binding protein | 8 | 6 | 6 | 6 | 6 | 6 | 1 |
| 190 | - | 210934 | 209318 | - | - | PA14_02340 | ABC transporter permease | 6 | 6 | 4 | 6 | 5 | 5 | 1 |
| 191 | - | 211201 | 212262 | + | - | PA14_02360 | ABC transporter substrate-binding protein | 10 | 10 | 10 | 10 | 11 | 12 | 1 |
| 192 | - | 213643 | 212285 | - | - | PA14_02370 | porin | 14 | 15 | 15 | 14 | 13 | 14 | 1 |
| 193 | - | 213891 | 214616 | + | - | PA14_02380 | acid phosphatase | 8 | 9 | 9 | 12 | 8 | 8 | 1 |
| 194 | - | 215557 | 214640 | - | - | PA14_02390 | transcriptional regulator | 8 | 14 | 11 | 12 | 10 | 10 | 0.956796655 |
| 195 | - | 215908 | 218280 | + | - | PA14_02410 | TonB-dependent receptor | 7 | 9 | 8 | 10 | 7 | 8 | 1 |
| 196 | - | 218321 | 219223 | + | - | PA14_02420 | hypothetical protein | 4 | 3 | 4 | 3 | 4 | 3 | 1 |
| 197 | - | 219291 | 220190 | + | - | PA14_02435 | hypothetical protein | 5 | 5 | 6 | 6 | 5 | 5 | 1 |
| 198 | - | 220837 | 221955 | + | - | PA14_02450 | NAD(P) transhydrogenase subunit alpha part 1 | 56 | 59 | 71 | 67 | 62 | 73 | 1 |
| 199 | - | 222031 | 222339 | + | - | PA14_02460 | NAD(P) transhydrogenase subunit alpha part 2 | 27 | 28 | 33 | 37 | 34 | 44 | 1 |
| 200 | - | 222339 | 223775 | + | pntB | PA14_02470 | pyridine nucleotide transhydrogenase subunit beta | 30 | 36 | 34 | 36 | 33 | 45 | 1 |
| 201 | - | 224119 | 224931 | + | tonB2 | PA14_02490 | hypothetical protein | 11 | 9 | 9 | 5 | 10 | 14 | 1 |
| 202 | - | 224960 | 225679 | + | exbB1 | PA14_02500 | transport protein ExbB | 17 | 16 | 18 | 17 | 19 | 29 | 1 |
| 203 | - | 225681 | 226082 | + | exbD1 | PA14_02510 | transport protein ExbD | 76 | 54 | 45 | 23 | 63 | 133 | 1 |
| 204 | - | 226475 | 226263 | - | - | PA14_02520 | hypothetical protein | 215 | 189 | 144 | 75 | 268 | 504 | 1 |
| 205 | - | 226691 | 227272 | + | - | PA14_02530 | hypothetical protein | 34 | 68 | 45 | 27 | 45 | 55 | 0.317658547 |
| 206 | - | 227383 | 229047 | + | mdcA | PA14_02550 | malonate decarboxylase subunit alpha | 78 | 116 | 90 | 68 | 92 | 96 | 1 |
| 207 | - | 229047 | 229928 | + | - | PA14_02560 | triphosphoribosyl-dephospho-CoA synthase | 37 | 70 | 60 | 43 | 62 | 80 | 0.559494168 |
| 208 | - | 229930 | 230229 | + | mdcC | PA14_02570 | malonate decarboxylase subunit delta | 23 | 57 | 45 | 30 | 56 | 61 | 0.044254247 |
| 209 | - | 230222 | 231085 | + | mdcD | PA14_02580 | malonate decarboxylase subunit beta | 38 | 82 | 66 | 49 | 75 | 84 | 0.320506198 |
| 210 | - | 231082 | 231888 | + | mdcE | PA14_02590 | malonate decarboxylase subunit gamma | 28 | 56 | 47 | 40 | 60 | 64 | 0.343217685 |
| 211 | - | 231904 | 232596 | + | - | PA14_02610 | phosphoribosyl-dephospho-CoA transferase | 31 | 42 | 57 | 31 | 41 | 39 | 1 |
| 212 | - | 232593 | 233525 | + | - | PA14_02620 | epsilon subunit of malonate decarboxylase | 48 | 55 | 63 | 44 | 54 | 51 | 1 |
| 213 | - | 233582 | 233986 | + | - | PA14_02630 | malonate carrier protein | 20 | 25 | 27 | 21 | 23 | 22 | 1 |
| 214 | - | 233992 | 234756 | + | - | PA14_02640 | malonate transporter subunit MadM | 18 | 25 | 24 | 24 | 25 | 25 | 1 |
| 215 | - | 235770 | 234841 | - | - | PA14_02650 | malonate utilization transcriptional regulator | 21 | 20 | 20 | 22 | 21 | 18 | 1 |
| 216 | - | 237019 | 236099 | - | - | PA14_02660 | LysR family transcriptional regulator | 22 | 19 | 21 | 22 | 22 | 23 | 1 |
| 217 | - | 237370 | 238860 | + | - | PA14_02680 | aldehyde dehydrogenase | 4 | 4 | 4 | 6 | 4 | 5 | 1 |
| 218 | - | 238918 | 240351 | + | - | PA14_02690 | amino acid permease | 12 | 11 | 12 | 16 | 10 | 12 | 1 |
| 219 | - | 240382 | 241764 | + | - | PA14_02700 | aminotransferase | 6 | 7 | 5 | 6 | 6 | 8 | 1 |
| 220 | - | 241928 | 242986 | + | - | PA14_02720 | hypothetical protein | 2 | 2 | 2 | 2 | 2 | 1 | 1 |
| 221 | - | 243945 | 243064 | - | - | PA14_02730 | dihydrodipicolinate synthetase | 17 | 10 | 10 | 11 | 8 | 8 | 0.527092266 |
| 222 | - | 244787 | 244005 | - | - | PA14_02740 | class II aldolase/adducin domain-containing protein | 39 | 29 | 34 | 40 | 30 | 34 | 1 |
| 223 | - | 244942 | 245481 | + | - | PA14_02750 | transcriptional regulator | 43 | 41 | 39 | 35 | 41 | 38 | 1 |
| 224 | - | 245632 | 246483 | + | - | PA14_02760 | CoA transferase, subunit A | 5 | 6 | 5 | 4 | 5 | 6 | 1 |
| 225 | - | 246480 | 247262 | + | - | PA14_02770 | CoA transferase subunit B | 8 | 8 | 9 | 8 | 6 | 7 | 1 |
| 226 | - | 247259 | 248464 | + | pcaF | PA14_02790 | beta-ketoadipyl CoA thiolase | 16 | 15 | 12 | 14 | 10 | 12 | 1 |
| 227 | - | 248614 | 249912 | + | pcaT | PA14_02810 | dicarboxylic acid transporter PcaT | 31 | 25 | 28 | 28 | 28 | 27 | 1 |
| 228 | - | 249935 | 251314 | + | pcaB | PA14_02830 | 3-carboxy-cis,cis-muconate cycloisomerase | 64 | 62 | 59 | 59 | 61 | 60 | 1 |
| 229 | - | 251329 | 252120 | + | pcaD | PA14_02840 | beta-ketoadipate enol-lactone hydrolase | 46 | 49 | 41 | 38 | 50 | 40 | 1 |
| 230 | - | 252131 | 252532 | + | pcaC | PA14_02850 | gamma-carboxymuconolactone decarboxylase | 49 | 67 | 47 | 59 | 65 | 72 | 1 |
| 231 | - | 252660 | 253601 | + | - | PA14_02870 | transcriptional regulator | 52 | 44 | 42 | 48 | 44 | 47 | 1 |
| 232 | - | 253792 | 254625 | + | - | PA14_02890 | hypothetical protein | 6 | 6 | 6 | 6 | 6 | 5 | 1 |
| 233 | - | 256110 | 254764 | - | pcaK | PA14_02900 | 4-hydroxybenzoate transporter PcaK | 2 | 2 | 3 | 2 | 2 | 2 | 1 |
| 234 | - | 256416 | 257195 | + | - | PA14_02910 | IclR family transcriptional regulator | 52 | 15 | 26 | 32 | 19 | 42 | 9.68945E-05 |
| 235 | - | 257439 | 258482 | + | - | PA14_02930 | oxidoreductase | 9 | 7 | 8 | 7 | 7 | 7 | 1 |
| 236 | - | 258505 | 259320 | + | - | PA14_02960 | hypothetical protein | 11 | 12 | 13 | 13 | 11 | 12 | 1 |
| 237 | - | 259470 | 260348 | + | - | PA14_02970 | hypothetical protein | 12 | 12 | 15 | 11 | 11 | 9 | 1 |
| 238 | - | 261639 | 260374 | - | - | PA14_02980 | porin | 14 | 13 | 15 | 15 | 13 | 15 | 1 |
| 239 | - | 263030 | 261705 | - | - | PA14_02990 | MFS transporter | 4 | 5 | 5 | 4 | 5 | 5 | 1 |
| 240 | - | 263577 | 265481 | + | - | PA14_03000 | hypothetical protein | 8 | 8 | 8 | 8 | 7 | 8 | 1 |
| 241 | - | 265574 | 266242 | + | - | PA14_03010 | TetR family transcriptional regulator | 35 | 24 | 27 | 23 | 22 | 32 | 1 |
| 242 | - | 267136 | 266282 | - | - | PA14_03020 | shikimate 5-dehydrogenase | 9 | 5 | 5 | 4 | 6 | 6 | 0.43335915 |
| 243 | - | 267579 | 267133 | - | aroQ2 | PA14_03030 | 3-dehydroquinate dehydratase | 5 | 5 | 6 | 4 | 4 | 5 | 1 |
| 244 | - | 269201 | 267696 | - | - | PA14_03040 | MFS transporter | 20 | 20 | 19 | 17 | 20 | 14 | 1 |
| 245 | - | 270560 | 269376 | - | pobA | PA14_03050 | 4-hydroxybenzoate 3-monooxygenase | 9 | 6 | 6 | 6 | 6 | 7 | 1 |
| 246 | - | 270738 | 271604 | + | - | PA14_03070 | transcriptional regulator | 28 | 39 | 30 | 25 | 32 | 29 | 1 |
| 247 | - | 272047 | 271601 | - | - | PA14_03080 | acetyltransferase | 80 | 46 | 33 | 45 | 39 | 93 | 0.810957915 |
| 248 | - | 272559 | 272125 | - | - | PA14_03090 | hypothetical protein | 423 | 121 | 124 | 144 | 129 | 377 | 0.015343712 |
| 249 | - | 273365 | 272724 | - | - | PA14_03100 | hypothetical protein | 8 | 7 | 8 | 7 | 6 | 5 | 1 |
| 250 | - | 273920 | 273633 | - | - | PA14_03110 | hypothetical protein | 5 | 4 | 5 | 3 | 4 | 4 | 1 |
| 251 | - | 274488 | 274012 | - | - | PA14_03120 | transcriptional regulator | 35 | 28 | 30 | 31 | 24 | 30 | 1 |
| 252 | - | 276019 | 274529 | - | - | PA14_03130 | hypothetical protein | 56 | 48 | 48 | 57 | 49 | 46 | 1 |

| | A | B | C | D | E | F | G | H | I | J | K | L | M | N |
|---|---|---|---|---|---|---|---|---|---|---|---|---|---|---|
| 253 | - | 276839 | 276153 | - | - | *PA14_03150* | hypothetical protein | 113 | 111 | 115 | 123 | 113 | 100 | 1 |
| 254 | - | 277935 | 277003 | - | - | *PA14_03160* | hypothetical protein | 110 | 106 | 137 | 142 | 109 | 137 | 1 |
| 255 | - | 278510 | 279133 | + | - | *PA14_03163* | hypothetical protein | 715 | 666 | 627 | 614 | 540 | 645 | 1 |
| 256 | - | 279133 | 279531 | + | - | *PA14_03166* | hypothetical protein | 119 | 111 | 119 | 116 | 155 | 140 | 1 |
| 257 | - | 280557 | 279718 | - | - | *PA14_03170* | hypothetical protein | 46 | 46 | 38 | 42 | 48 | 39 | 1 |
| 258 | - | 280853 | 280590 | - | - | *PA14_03180* | hypothetical protein | 108 | 124 | 97 | 118 | 137 | 112 | 1 |
| 259 | - | 282415 | 280973 | - | - | *PA14_03190* | hypothetical protein | 87 | 79 | 76 | 80 | 86 | 79 | 1 |
| 260 | - | 284715 | 282565 | - | - | *PA14_03200* | hypothetical protein | 31 | 30 | 28 | 26 | 35 | 26 | 1 |
| 261 | - | 285209 | 284712 | - | - | *PA14_03210* | hypothetical protein | 24 | 20 | 22 | 18 | 27 | 19 | 1 |
| 262 | - | 288272 | 285213 | - | - | *PA14_03220* | hypothetical protein | 31 | 26 | 24 | 23 | 28 | 21 | 1 |
| 263 | - | 288972 | 288454 | - | *hcpC* | *PA14_03240* | secreted protein Hcp | 33 | 19 | 18 | 20 | 21 | 22 | 0.580848337 |
| 264 | - | 291568 | 293184 | + | - | *PA14_03250* | hypothetical protein | 70 | 67 | 63 | 58 | 64 | 56 | 1 |
| 265 | - | 293187 | 293936 | + | - | *PA14_03265* | hypothetical protein | 82 | 82 | 66 | 65 | 86 | 69 | 1 |
| 266 | - | 294101 | 297238 | + | - | *PA14_03270* | hypothetical protein | 101 | 94 | 77 | 81 | 98 | 85 | 1 |
| 267 | - | 297222 | 298367 | + | - | *PA14_03285* | hypothetical protein | 63 | 57 | 50 | 51 | 62 | 55 | 1 |
| 268 | - | 298567 | 298875 | + | - | *PA14_03290* | hypothetical protein | 461 | 431 | 441 | 441 | 406 | 406 | 1 |
| 269 | - | 298872 | 299717 | + | - | *PA14_03300* | hypothetical protein | 123 | 111 | 115 | 119 | 115 | 111 | 1 |
| 270 | - | 300333 | 301931 | + | - | *PA14_03310* | hypothetical protein | 41 | 47 | 45 | 46 | 46 | 44 | 1 |
| 271 | - | 303804 | 302683 | - | - | *PA14_03320* | hypothetical protein | 80 | 78 | 66 | 70 | 85 | 69 | 1 |
| 272 | - | 304676 | 303810 | - | - | *PA14_03330* | hypothetical protein | 26 | 30 | 24 | 22 | 34 | 25 | 1 |
| 273 | - | 305599 | 304673 | - | - | *PA14_03340* | hypothetical protein | 41 | 33 | 37 | 34 | 41 | 34 | 1 |
| 274 | - | 306903 | 305950 | - | - | *PA14_03350* | hypothetical protein | 165 | 185 | 128 | 148 | 182 | 151 | 1 |
| 275 | - | 307666 | 306896 | - | - | *PA14_03360* | hypothetical protein | 174 | 193 | 138 | 133 | 171 | 136 | 1 |
| 276 | - | 308983 | 307985 | - | - | *PA14_03370* | hypothetical protein | 88 | 88 | 79 | 83 | 101 | 79 | 1 |
| 277 | - | 309749 | 311728 | + | - | *PA14_03380* | hypothetical protein | 65 | 59 | 60 | 66 | 62 | 60 | 1 |
| 278 | - | 312174 | 311665 | - | - | *PA14_03390* | hypothetical protein | 11 | 13 | 12 | 9 | 11 | 9 | 1 |
| 279 | - | 312499 | 312167 | - | - | *PA14_03400* | hypothetical protein | 3 | 6 | 5 | 3 | 4 | 2 | 1 |
| 280 | - | 313034 | 312961 | - | - | *PA14_03410* | Arg tRNA | 18 | 14 | 15 | 8 | 12 | 13 | 1 |
| 281 | - | 313638 | 313222 | - | - | *PA14_03420* | hypothetical protein | 44 | 34 | 42 | 39 | 43 | 35 | 1 |
| 282 | - | 313663 | 315114 | + | *gabD* | *PA14_03430* | succinate-semialdehyde dehydrogenase I | 158 | 188 | 140 | 164 | 153 | 162 | 1 |
| 283 | - | 315359 | 316639 | + | *gabT* | *PA14_03450* | 4-aminobutyrate aminotransferase | 174 | 219 | 122 | 164 | 187 | 184 | 1 |
| 284 | - | 316965 | 318164 | + | - | *PA14_03470* | hypothetical protein | 75 | 81 | 76 | 80 | 82 | 69 | 1 |
| 285 | - | 319753 | 318332 | - | - | *PA14_03480* | GntR family transcriptional regulator | 15 | 20 | 16 | 18 | 19 | 17 | 1 |
| 286 | - | 319880 | 320317 | + | - | *PA14_03490* | hypothetical protein | 78 | 56 | 42 | 52 | 46 | 102 | 1 |
| 287 | - | 320329 | 320736 | + | - | *PA14_03510* | hypothetical protein | 98 | 72 | 52 | 74 | 83 | 145 | 1 |
| 288 | - | 320770 | 321054 | + | - | *PA14_03520* | hypothetical protein | 127 | 63 | 62 | 91 | 83 | 183 | 0.321350893 |
| 289 | - | 321983 | 321051 | - | - | *PA14_03530* | transcriptional regulator | 57 | 30 | 32 | 45 | 38 | 90 | 0.8338713 |
| 290 | - | 323247 | 322033 | - | - | *PA14_03550* | MFS transporter | 7 | 5 | 6 | 6 | 4 | 6 | 1 |
| 291 | - | 324180 | 323410 | - | - | *PA14_03560* | hypothetical protein | 5 | 5 | 4 | 4 | 6 | 4 | 1 |
| 292 | - | 324975 | 324289 | - | - | *PA14_03580* | transcriptional regulator | 44 | 35 | 41 | 32 | 31 | 37 | 1 |
| 293 | - | 325051 | 325566 | + | - | *PA14_03590* | hypothetical protein | 81 | 47 | 67 | 53 | 45 | 75 | 1 |
| 294 | - | 326364 | 325606 | - | - | *PA14_03610* | Zn-dependent protease with chaperone function | 31 | 20 | 29 | 23 | 23 | 35 | 1 |
| 295 | - | 327288 | 326536 | - | - | *PA14_03620* | hypothetical protein | 9 | 7 | 8 | 8 | 8 | 8 | 1 |
| 296 | - | 327382 | 328080 | + | - | *PA14_03630* | ArsR family transcriptional regulator | 39 | 34 | 37 | 32 | 33 | 31 | 1 |
| 297 | - | 329082 | 328093 | - | *cysA* | *PA14_03650* | sulfate transport protein CysA | 40 | 35 | 14 | 15 | 26 | 15 | 1 |
| 298 | - | 329955 | 329086 | - | *cysW* | *PA14_03670* | sulfate transport protein CysW | 33 | 28 | 14 | 17 | 25 | 12 | 1 |
| 299 | - | 330784 | 329966 | - | *cysT* | *PA14_03680* | sulfate transport protein CysT | 19 | 18 | 9 | 10 | 11 | 10 | 1 |
| 300 | - | 331944 | 330946 | - | *sbp* | *PA14_03700* | sulfate-binding protein | 14 | 35 | 12 | 10 | 19 | 10 | 0.070186394 |
| 301 | - | 332303 | 332121 | - | - | *PA14_03710* | hypothetical protein | 40 | 105 | 49 | 42 | 54 | 37 | 0.022359242 |
| 302 | - | 334749 | 332467 | - | - | *PA14_03720* | sensory box GGDEF domain-containing protein | 23 | 25 | 22 | 23 | 23 | 20 | 1 |
| 303 | - | 334920 | 336098 | + | - | *PA14_03730* | fatty acid desaturase | 215 | 181 | 203 | 132 | 153 | 140 | 1 |
| 304 | - | 336327 | 337712 | + | - | *PA14_03760* | sodium:solute symporter | 7 | 6 | 6 | 6 | 8 | 6 | 1 |
| 305 | - | 337768 | 338724 | + | *speB1* | *PA14_03770* | agmatinase | 5 | 7 | 5 | 7 | 5 | 4 | 1 |
| 306 | - | 338777 | 339739 | + | - | *PA14_03780* | transcriptional regulator | 34 | 29 | 31 | 30 | 29 | 24 | 1 |
| 307 | - | 339852 | 340823 | + | - | *PA14_03790* | sensory box GGDEF domain-containing protein | 106 | 112 | 143 | 146 | 98 | 107 | 1 |
| 308 | - | 341437 | 342828 | + | *oprE* | *PA14_03800* | anaerobically-induced outer membrane porin OprE precursor | 169 | 253 | 124 | 120 | 186 | 134 | 1 |
| 309 | - | 344032 | 342926 | - | *aguA* | *PA14_03810* | agmatine deiminase | 45 | 55 | 35 | 32 | 47 | 38 | 1 |
| 310 | - | 344996 | 344118 | - | *aguB* | *PA14_03830* | N-carbamoylputrescine amidohydrolase | 11 | 12 | 11 | 8 | 9 | 9 | 1 |
| 311 | - | 345824 | 345159 | - | *aguR* | *PA14_03840* | transcriptional regulator AguR | 17 | 19 | 20 | 20 | 19 | 20 | 1 |
| 312 | - | 346948 | 345887 | - | - | *PA14_03855* | periplasmic polyamine binding protein | 38 | 39 | 46 | 34 | 31 | 30 | 1 |
| 313 | - | 348580 | 347204 | - | - | *PA14_03860* | glutamine synthetase | 167 | 187 | 157 | 181 | 149 | 173 | 1 |
| 314 | - | 348859 | 349611 | + | *spuA* | *PA14_03870* | glutamine amidotransferase | 92 | 94 | 101 | 81 | 79 | 85 | 1 |
| 315 | - | 349652 | 351010 | + | *spuB* | *PA14_03880* | glutamine synthetase | 107 | 136 | 95 | 130 | 101 | 104 | 1 |
| 316 | - | 351076 | 352446 | + | *spuC* | *PA14_03900* | aminotransferase | 327 | 398 | 274 | 335 | 314 | 289 | 1 |
| 317 | - | 352562 | 353665 | + | *spuD* | *PA14_03920* | polyamine transport protein | 272 | 320 | 206 | 249 | 278 | 248 | 1 |
| 318 | - | 354084 | 355181 | + | *spuE* | *PA14_03930* | polyamine transport protein | 117 | 130 | 108 | 106 | 111 | 108 | 1 |
| 319 | - | 355236 | 356390 | + | *spuF* | *PA14_03940* | polyamine transport protein PotG | 136 | 181 | 117 | 154 | 169 | 148 | 1 |
| 320 | - | 356417 | 357298 | + | *spuG* | *PA14_03950* | polyamine transport protein PotH | 119 | 142 | 99 | 122 | 134 | 126 | 1 |
| 321 | - | 357381 | 358250 | + | *spuH* | *PA14_03960* | polyamine transport protein PotI | 191 | 187 | 149 | 161 | 175 | 163 | 1 |
| 322 | - | 360835 | 358448 | - | - | *PA14_03980* | hypothetical protein | 24 | 27 | 21 | 21 | 25 | 22 | 1 |
| 323 | - | 360946 | 361998 | + | - | *PA14_04000* | transcriptional regulator | 52 | 68 | 57 | 53 | 67 | 55 | 1 |
| 324 | - | 362480 | 361977 | - | - | *PA14_04010* | hypothetical protein | 169 | 241 | 194 | 141 | 183 | 159 | 1 |
| 325 | - | 363195 | 364214 | + | - | *PA14_04030* | hypothetical protein | 43 | 43 | 37 | 37 | 34 | 31 | 1 |
| 326 | - | 363196 | 362585 | - | - | *PA14_04020* | hypothetical protein | 50 | 44 | 56 | 52 | 42 | 43 | 1 |
| 327 | - | 364986 | 364234 | - | - | *PA14_04040* | hypothetical protein | 73 | 63 | 68 | 60 | 55 | 57 | 1 |
| 328 | - | 365035 | 365733 | + | - | *PA14_04050* | proline hydroxylase | 54 | 54 | 65 | 62 | 52 | 56 | 1 |
| 329 | - | 366309 | 365755 | - | - | *PA14_04060* | hypothetical protein | 7 | 5 | 9 | 7 | 5 | 5 | 1 |
| 330 | - | 366575 | 367069 | + | - | *PA14_04070* | hypothetical protein | 149 | 154 | 184 | 195 | 141 | 149 | 1 |
| 331 | - | 367766 | 367074 | - | - | *PA14_04080* | ABC transporter permease | 61 | 68 | 60 | 71 | 62 | 62 | 1 |
| 332 | - | 368608 | 367838 | - | - | *PA14_04090* | ABC transporter substrate-binding protein | 51 | 56 | 55 | 50 | 65 | 54 | 1 |
| 333 | - | 368901 | 369338 | + | - | *PA14_04100* | hypothetical protein | 185 | 141 | 120 | 153 | 153 | 212 | 1 |
| 334 | - | 370625 | 369396 | - | *serA* | *PA14_04110* | D-3-phosphoglycerate dehydrogenase | 105 | 119 | 94 | 88 | 103 | 88 | 1 |
| 335 | - | 370829 | 372223 | + | - | *PA14_04140* | hypothetical protein | 88 | 92 | 97 | 85 | 91 | 75 | 1 |
| 336 | - | 372315 | 372980 | + | - | *PA14_04150* | hypothetical protein | 216 | 226 | 186 | 155 | 167 | 180 | 1 |

| | A | B | C | D | E | F | G | H | I | J | K | L | M | N |
|---|---|---|---|---|---|---|---|---|---|---|---|---|---|---|
| 337 | - | 373079 | 374068 | + | - | *PA14_04160* | transcriptional regulator | 57 | 58 | 58 | 60 | 59 | 57 | 1 |
| 338 | - | 374152 | 374502 | + | - | *PA14_04180* | hypothetical protein | 19 | 27 | 32 | 17 | 12 | 8 | 1 |
| 339 | - | 375617 | 374583 | - | - | *PA14_04190* | acetylpolyamine aminohydrolase | 18 | 17 | 20 | 19 | 17 | 18 | 1 |
| 340 | - | 377027 | 375633 | - | - | *PA14_04210* | transporter | 10 | 9 | 11 | 10 | 9 | 9 | 1 |
| 341 | - | 378491 | 377448 | - | - | *PA14_04220* | ABC transporter substrate-binding protein | 12 | 12 | 12 | 13 | 10 | 11 | 1 |
| 342 | - | 379327 | 378539 | - | - | *PA14_04230* | ABC transporter permease | 13 | 15 | 14 | 17 | 12 | 15 | 1 |
| 343 | - | 380253 | 379324 | - | - | *PA14_04240* | ABC transporter permease | 42 | 41 | 44 | 49 | 44 | 50 | 1 |
| 344 | - | 381299 | 380250 | - | - | *PA14_04250* | ABC transporter ATP-binding protein | 5 | 4 | 4 | 4 | 5 | 4 | 1 |
| 345 | - | 381627 | 382592 | + | - | *PA14_04270* | transcriptional regulator | 38 | 34 | 43 | 42 | 36 | 40 | 1 |
| 346 | - | 384574 | 382631 | - | - | *PA14_04290* | hypothetical protein | 37 | 41 | 40 | 42 | 39 | 35 | 1 |
| 347 | - | 385207 | 384875 | - | - | *PA14_04300* | hypothetical protein | 199 | 271 | 222 | 214 | 190 | 231 | 1 |
| 348 | - | 386001 | 385330 | - | *rpiA* | *PA14_04310* | ribose-5-phosphate isomerase A | 71 | 62 | 87 | 59 | 49 | 53 | 1 |
| 349 | - | 386260 | 387774 | + | *ilvA1* | *PA14_04320* | threonine dehydratase | 64 | 62 | 67 | 56 | 63 | 48 | 1 |
| 350 | - | 387894 | 388361 | + | - | *PA14_04330* | hypothetical protein | 213 | 198 | 252 | 215 | 194 | 197 | 1 |
| 351 | - | 388412 | 389683 | + | - | *PA14_04340* | hypothetical protein | 68 | 55 | 68 | 61 | 63 | 63 | 1 |
| 352 | - | 389680 | 390147 | + | - | *PA14_04350* | integral membrane protein | 96 | 100 | 94 | 107 | 104 | 93 | 1 |
| 353 | - | 391361 | 390120 | - | - | *PA14_04370* | MFS transporter | 45 | 40 | 45 | 35 | 37 | 40 | 1 |
| 354 | - | 392064 | 391411 | - | - | *PA14_04380* | hypothetical protein | 54 | 44 | 55 | 46 | 43 | 37 | 1 |
| 355 | - | 392268 | 392747 | + | *ygdP* | *PA14_04390* | dinucleoside polyphosphate hydrolase | 402 | 373 | 413 | 418 | 416 | 379 | 1 |
| 356 | - | 392770 | 395049 | + | *ptsP* | *PA14_04410* | phosphoenolpyruvate-protein phosphotransferase PtsP | 405 | 427 | 364 | 411 | 376 | 392 | 1 |
| 357 | - | 395075 | 396205 | + | - | *PA14_04420* | hypothetical protein | 51 | 47 | 52 | 50 | 41 | 41 | 1 |
| 358 | - | 396964 | 396209 | - | - | *PA14_04430* | hypothetical protein | 44 | 41 | 49 | 43 | 39 | 39 | 1 |
| 359 | - | 397086 | 397889 | + | - | *PA14_04440* | permease | 27 | 21 | 30 | 28 | 24 | 24 | 1 |
| 360 | - | 397899 | 398699 | + | *lgt* | *PA14_04460* | prolipoprotein diacylglyceryl transferase | 55 | 73 | 60 | 60 | 65 | 49 | 1 |
| 361 | - | 398905 | 399699 | + | *thyA* | *PA14_04480* | thymidylate synthase | 66 | 65 | 75 | 68 | 71 | 67 | 1 |
| 362 | - | 400523 | 399726 | - | - | *PA14_04490* | hypothetical protein | 73 | 73 | 77 | 69 | 82 | 74 | 1 |
| 363 | - | 401937 | 400558 | - | - | *PA14_04510* | hypothetical protein | 104 | 116 | 102 | 123 | 118 | 120 | 1 |
| 364 | - | 403315 | 401930 | - | - | *PA14_04520* | hypothetical protein | 68 | 60 | 67 | 76 | 64 | 68 | 1 |
| 365 | - | 403507 | 403869 | + | - | *PA14_04530* | hypothetical protein | 53 | 55 | 50 | 44 | 48 | 41 | 1 |
| 366 | - | 405057 | 403906 | - | *glpQ* | *PA14_04550* | glycerophosphoryl diester phosphodiesterase, periplasmic | 19 | 21 | 20 | 21 | 21 | 19 | 1 |
| 367 | - | 405267 | 406361 | + | - | *PA14_04560* | hypothetical protein | 18 | 20 | 21 | 21 | 17 | 17 | 1 |
| 368 | - | 406358 | 407398 | + | - | *PA14_04570* | hypothetical protein | 14 | 14 | 15 | 15 | 14 | 14 | 1 |
| 369 | - | 407469 | 407975 | + | *folA* | *PA14_04580* | dihydrofolate reductase | 22 | 33 | 28 | 23 | 27 | 23 | 1 |
| 370 | - | 408464 | 407991 | - | - | *PA14_04590* | hypothetical protein | 51 | 70 | 64 | 45 | 60 | 46 | 1 |
| 371 | - | 409988 | 408603 | - | - | *PA14_04610* | transporter | 11 | 12 | 15 | 10 | 11 | 8 | 1 |
| 372 | - | 412056 | 410218 | - | *ilvD* | *PA14_04630* | dihydroxy-acid dehydratase | 102 | 118 | 101 | 87 | 112 | 99 | 1 |
| 373 | - | 413580 | 412384 | - | - | *PA14_04640* | hypothetical protein | 59 | 52 | 69 | 55 | 55 | 60 | 1 |
| 374 | - | 414192 | 413653 | - | *pfpI* | *PA14_04650* | protease PfpI | 110 | 149 | 114 | 97 | 142 | 115 | 1 |
| 375 | - | 414409 | 415233 | + | - | *PA14_04660* | hypothetical protein | 28 | 23 | 33 | 19 | 20 | 18 | 1 |
| 376 | - | 415292 | 416104 | + | *mutM* | *PA14_04670* | formamidopyrimidine-DNA glycosylase | 62 | 56 | 60 | 55 | 58 | 45 | 1 |
| 377 | - | 416181 | 416759 | + | - | *PA14_04680* | hypothetical protein | 24 | 24 | 24 | 25 | 22 | 21 | 1 |
| 378 | - | 416842 | 417186 | + | - | *PA14_04690* | hypothetical protein | 136 | 161 | 184 | 98 | 131 | 123 | 1 |
| 379 | - | 417456 | 418442 | + | - | *PA14_04700* | hypothetical protein | 80 | 83 | 79 | 84 | 87 | 80 | 1 |
| 380 | - | 418986 | 418555 | - | - | *PA14_04710* | hypothetical protein | 76 | 78 | 80 | 59 | 79 | 64 | 1 |
| 381 | - | 420871 | 419138 | - | - | *PA14_04730* | gamma-glutamyltranspeptidase | 57 | 50 | 62 | 57 | 47 | 48 | 1 |
| 382 | - | 421250 | 420999 | - | *fdx1* | *PA14_04750* | ferredoxin (4Fe-4S) | 74 | 63 | 80 | 53 | 54 | 48 | 1 |
| 383 | - | 421850 | 421371 | - | *coaD* | *PA14_04760* | phosphopantetheine adenylyltransferase | 28 | 23 | 30 | 26 | 21 | 29 | 1 |
| 384 | - | 423594 | 421999 | - | - | *PA14_04780* | oxidoreductase | 33 | 27 | 27 | 36 | 27 | 38 | 1 |
| 385 | - | 424198 | 423650 | - | - | *PA14_04790* | hypothetical protein | 42 | 43 | 42 | 57 | 39 | 58 | 1 |
| 386 | - | 425683 | 424253 | - | - | *PA14_04810* | aldehyde dehydrogenase | 73 | 64 | 65 | 91 | 61 | 94 | 1 |
| 387 | - | 425972 | 426619 | + | - | *PA14_04820* | TetR family transcriptional regulator | 66 | 66 | 75 | 76 | 64 | 70 | 1 |
| 388 | - | 427014 | 426604 | - | - | *PA14_04830* | acetyltransferase | 207 | 211 | 181 | 209 | 254 | 229 | 1 |
| 389 | - | 428075 | 427077 | - | - | *PA14_04840* | hypothetical protein | 65 | 47 | 54 | 52 | 56 | 52 | 1 |
| 390 | - | 428402 | 428112 | - | - | *PA14_04850* | hypothetical protein | 83 | 77 | 82 | 75 | 93 | 85 | 1 |
| 391 | - | 429277 | 428681 | - | - | *PA14_04860* | methyltransferase | 98 | 107 | 85 | 95 | 101 | 83 | 1 |
| 392 | - | 430764 | 429277 | - | - | *PA14_04870* | hypothetical protein | 93 | 118 | 94 | 96 | 114 | 95 | 1 |
| 393 | - | 432154 | 430757 | - | - | *PA14_04890* | zinc protease | 53 | 50 | 56 | 46 | 50 | 50 | 1 |
| 394 | - | 432275 | 433630 | + | *ftsY* | *PA14_04900* | signal recognition particle receptor FtsY | 152 | 127 | 165 | 153 | 145 | 133 | 1 |
| 395 | - | 433627 | 434298 | + | *ftsE* | *PA14_04910* | cell division ATP-binding protein FtsE | 181 | 197 | 197 | 211 | 209 | 184 | 1 |
| 396 | - | 434298 | 435305 | + | *ftsX* | *PA14_04920* | cell division protein FtsX | 203 | 182 | 198 | 212 | 209 | 180 | 1 |
| 397 | - | 435419 | 436273 | + | *rpoH* | *PA14_04930* | RNA polymerase factor sigma-32 | 677 | 557 | 773 | 497 | 455 | 473 | 1 |
| 398 | - | 436338 | 436943 | + | - | *PA14_04940* | hypothetical protein | 97 | 105 | 101 | 108 | 114 | 87 | 1 |
| 399 | - | 437679 | 436948 | - | *mtgA* | *PA14_04950* | monofunctional biosynthetic peptidoglycan transglycosylase | 43 | 45 | 52 | 45 | 42 | 36 | 1 |
| 400 | - | 437716 | 438093 | + | - | *PA14_04960* | hypothetical protein | 41 | 51 | 41 | 53 | 41 | 40 | 1 |
| 401 | - | 438196 | 438396 | + | - | *PA14_04970* | sulfur carrier protein ThiS | 33 | 26 | 31 | 28 | 22 | 20 | 1 |
| 402 | - | 438455 | 439252 | + | *thiG* | *PA14_04980* | thiazole synthase | 34 | 39 | 34 | 24 | 24 | 22 | 1 |
| 403 | - | 439406 | 440077 | + | *trmB* | *PA14_05000* | tRNA (guanine-N(7)-)-methyltransferase | 37 | 32 | 43 | 33 | 33 | 25 | 1 |
| 404 | - | 440240 | 441586 | + | - | *PA14_05010* | hypothetical protein | 31 | 34 | 37 | 43 | 31 | 35 | 1 |
| 405 | - | 441856 | 441599 | - | - | *PA14_05020* | hypothetical protein | 22 | 29 | 27 | 25 | 26 | 36 | 1 |
| 406 | - | 442241 | 441918 | - | - | *PA14_05030* | hypothetical protein | 14 | 15 | 19 | 10 | 10 | 12 | 1 |
| 407 | - | 443476 | 442322 | - | - | *PA14_05040* | coproporphyrinogen III oxidase | 20 | 15 | 23 | 17 | 14 | 16 | 1 |
| 408 | - | 444190 | 443597 | - | - | *PA14_05050* | deoxyribonucleotide triphosphate pyrophosphatase | 226 | 240 | 238 | 201 | 225 | 218 | 1 |
| 409 | - | 444606 | 444187 | - | - | *PA14_05060* | hypothetical protein | 320 | 340 | 412 | 274 | 301 | 316 | 1 |
| 410 | - | 445358 | 444738 | - | - | *PA14_05070* | methionine biosynthesis protein | 21 | 28 | 34 | 21 | 24 | 23 | 1 |
| 411 | - | 446505 | 445366 | - | *metX* | *PA14_05080* | homoserine O-acetyltransferase | 18 | 17 | 21 | 19 | 16 | 16 | 1 |
| 412 | - | 448561 | 446594 | - | - | *PA14_05110* | hypothetical protein | 51 | 75 | 58 | 68 | 52 | 46 | 1 |
| 413 | - | 448839 | 448669 | - | - | *PA14_05120* | hypothetical protein | 42 | 44 | 34 | 26 | 39 | 36 | 1 |
| 414 | - | 449555 | 448962 | - | - | *PA14_05130* | hypothetical protein | 99 | 111 | 91 | 80 | 101 | 85 | 1 |
| 415 | - | 450387 | 449566 | - | *proC* | *PA14_05150* | pyrroline-5-carboxylate reductase | 186 | 192 | 182 | 149 | 151 | 151 | 1 |
| 416 | - | 451091 | 450399 | - | - | *PA14_05160* | hypothetical protein | 86 | 78 | 80 | 58 | 66 | 61 | 1 |
| 417 | - | 451306 | 452340 | + | *pilT* | *PA14_05180* | twitching motility protein PilT | 123 | 122 | 146 | 147 | 133 | 136 | 1 |
| 418 | - | 452518 | 453666 | + | *pilU* | *PA14_05190* | twitching motility protein PilU | 252 | 218 | 307 | 283 | 213 | 246 | 1 |
| 419 | - | 454573 | 453674 | - | - | *PA14_05200* | cation efflux system protein | 76 | 78 | 96 | 95 | 78 | 87 | 1 |
| 420 | - | 454728 | 455132 | + | - | *PA14_05210* | hypothetical protein | 31 | 46 | 30 | 20 | 24 | 33 | 1 |

| | A | B | C | D | E | F | G | H | I | J | K | L | M | N |
|---|---|---|---|---|---|---|---|---|---|---|---|---|---|---|
| 421 | - | 455375 | 456748 | + | - | *PA14_05220* | cystathionine beta-synthase | 119 | 154 | 101 | 101 | 144 | 106 | 1 |
| 422 | - | 456745 | 457929 | + | - | *PA14_05230* | cystathionine gamma-lyase | 111 | 151 | 82 | 86 | 131 | 94 | 1 |
| 423 | - | 459428 | 458157 | - | *pyrC* | *PA14_05250* | dihydroorotase | 100 | 107 | 88 | 85 | 102 | 85 | 1 |
| 424 | - | 460429 | 459425 | - | *pyrB* | *PA14_05260* | aspartate carbamoyltransferase | 139 | 136 | 131 | 119 | 131 | 115 | 1 |
| 425 | - | 460965 | 460453 | - | *pyrR* | *PA14_05270* | bifunctional pyrimidine regulatory protein PyrR/uracil phosphoribosyltransferase | 123 | 136 | 130 | 96 | 110 | 119 | 1 |
| 426 | - | 461511 | 461077 | - | *yqgF* | *PA14_05280* | Holliday junction resolvase-like protein | 67 | 68 | 63 | 62 | 64 | 57 | 1 |
| 427 | - | 462080 | 461511 | - | - | *PA14_05290* | hypothetical protein | 63 | 73 | 68 | 58 | 57 | 59 | 1 |
| 428 | - | 463088 | 462129 | - | - | *PA14_05300* | TonB domain-containing protein | 37 | 37 | 37 | 40 | 31 | 32 | 1 |
| 429 | - | 464122 | 463169 | - | *gshB* | *PA14_05310* | glutathione synthetase | 316 | 416 | 483 | 436 | 339 | 282 | 1 |
| 430 | - | 464377 | 464784 | + | *pilG* | *PA14_05320* | twitching motility protein PilG | 418 | 400 | 545 | 438 | 469 | 543 | 1 |
| 431 | - | 464831 | 465196 | + | *pilH* | *PA14_05330* | twitching motility protein PilH | 191 | 206 | 257 | 212 | 214 | 229 | 1 |
| 432 | - | 465247 | 465783 | + | *pilI* | *PA14_05340* | twitching motility protein PilI | 150 | 146 | 202 | 162 | 166 | 156 | 1 |
| 433 | - | 465868 | 467916 | + | *pilJ* | *PA14_05360* | twitching motility protein PilJ | 288 | 377 | 313 | 299 | 333 | 285 | 1 |
| 434 | - | 467977 | 468852 | + | *pilK* | *PA14_05380* | methyltransferase PilK | 199 | 220 | 218 | 253 | 241 | 219 | 1 |
| 435 | - | 468864 | 476294 | + | *chpA* | *PA14_05390* | ChpA | 339 | 383 | 350 | 374 | 402 | 364 | 1 |
| 436 | - | 476287 | 477318 | + | - | *PA14_05400* | methylesterase | 274 | 299 | 297 | 277 | 350 | 304 | 1 |
| 437 | - | 477315 | 477821 | + | - | *PA14_05410* | chemotaxis protein | 131 | 182 | 143 | 161 | 197 | 160 | 1 |
| 438 | - | 477829 | 478623 | + | - | *PA14_05420* | transcriptional regulator | 51 | 62 | 56 | 55 | 67 | 55 | 1 |
| 439 | - | 478700 | 479311 | + | - | *PA14_05430* | chemotaxis protein | 15 | 18 | 20 | 17 | 19 | 14 | 1 |
| 440 | - | 480734 | 479319 | - | - | *PA14_05440* | hypothetical protein | 21 | 26 | 20 | 22 | 24 | 17 | 1 |
| 441 | - | 481491 | 480769 | - | - | *PA14_05450* | 16S ribosomal RNA methyltransferase RsmE | 33 | 31 | 38 | 24 | 26 | 24 | 1 |
| 442 | - | 482987 | 481584 | - | *bioA* | *PA14_05460* | adenosylmethionine-8-amino-7-oxononanoate aminotransferase | 29 | 32 | 25 | 29 | 29 | 27 | 1 |
| 443 | - | 483161 | 484651 | + | - | *PA14_05480* | hypothetical protein | 63 | 68 | 63 | 61 | 60 | 49 | 1 |
| 444 | - | 484832 | 485401 | + | - | *PA14_05500* | hypothetical protein | 111 | 62 | 108 | 55 | 71 | 67 | 0.956402005 |
| 445 | - | 485413 | 485988 | + | - | *PA14_05510* | hypothetical protein | 316 | 488 | 235 | 201 | 302 | 259 | 1 |
| 446 | - | 486500 | 486057 | - | *mexR* | *PA14_05520* | multidrug resistance operon repressor MexR | 215 | 205 | 251 | 350 | 229 | 271 | 1 |
| 447 | - | 486775 | 487926 | + | *mexA* | *PA14_05530* | RND multidrug efflux membrane fusion protein MexA | 264 | 212 | 230 | 200 | 179 | 195 | 1 |
| 448 | - | 487942 | 491082 | + | *mexB* | *PA14_05540* | RND multidrug efflux transporter MexB | 499 | 401 | 467 | 420 | 394 | 387 | 1 |
| 449 | - | 491084 | 492541 | + | *oprM* | *PA14_05550* | major intrinsic multiple antibiotic resistance efflux outer membrane protein OprM precursor | 599 | 504 | 695 | 446 | 415 | 415 | 1 |
| 450 | - | 494556 | 492637 | - | - | *PA14_05560* | ATP-dependent RNA helicase | 238 | 184 | 221 | 171 | 165 | 186 | 1 |
| 451 | - | 495877 | 494807 | - | - | *PA14_05580* | hypothetical protein | 124 | 105 | 118 | 97 | 92 | 118 | 1 |
| 452 | - | 496819 | 495947 | - | *metF* | *PA14_05590* | 5,10-methylenetetrahydrofolate reductase | 192 | 111 | 131 | 111 | 98 | 155 | 1 |
| 453 | - | 497416 | 496862 | - | - | *PA14_05600* | hypothetical protein | 169 | 98 | 101 | 78 | 80 | 121 | 1 |
| 454 | - | 498866 | 497457 | - | *sahH* | *PA14_05620* | S-adenosyl-L-homocysteine hydrolase | 471 | 305 | 282 | 264 | 257 | 392 | 1 |
| 455 | - | 499155 | 499589 | + | - | *PA14_05630* | hypothetical protein | 20 | 16 | 20 | 18 | 15 | 13 | 1 |
| 456 | - | 499715 | 501907 | + | - | *PA14_05640* | hypothetical protein | 23 | 18 | 22 | 20 | 17 | 17 | 1 |
| 457 | - | 501918 | 503396 | + | - | *PA14_05650* | hypothetical protein | 12 | 9 | 12 | 9 | 10 | 8 | 1 |
| 458 | - | 503481 | 504101 | + | - | *PA14_05660* | transcriptional regulator | 203 | 133 | 156 | 171 | 187 | 272 | 1 |
| 459 | - | 505409 | 504138 | - | *codA* | *PA14_05690* | cytosine deaminase | 55 | 55 | 53 | 56 | 63 | 60 | 1 |
| 460 | - | 506649 | 505399 | - | *codB* | *PA14_05700* | cytosine permease | 14 | 13 | 12 | 12 | 14 | 12 | 1 |
| 461 | - | 508108 | 506831 | - | - | *PA14_05740* | dihydropyrimidine dehydrogenase | 21 | 7 | 17 | 9 | 6 | 20 | 0.003564822 |
| 462 | - | 509472 | 508105 | - | - | *PA14_05750* | oxidoreductase | 25 | 13 | 22 | 16 | 11 | 24 | 0.391278558 |
| 463 | - | 511006 | 509567 | - | *dhT* | *PA14_05770* | phenylhydantoinase | 46 | 17 | 34 | 18 | 11 | 30 | 0.053355548 |
| 464 | - | 511229 | 511113 | - | - | *PA14_05775* | hypothetical protein | 1 | 1 | 0 | 0 | 0 | 0 | 1 |
| 465 | - | 511622 | 513112 | + | - | *PA14_05790* | transporter | 10 | 6 | 9 | 8 | 7 | 11 | 1 |
| 466 | - | 513171 | 514454 | + | *amaB* | *PA14_05810* | allantoate amidohydrolase | 56 | 39 | 63 | 43 | 38 | 65 | 1 |
| 467 | - | 515878 | 514655 | - | - | *PA14_05820* | hypothetical protein | 79 | 116 | 84 | 92 | 107 | 122 | 1 |
| 468 | - | 517179 | 515998 | - | *gcdH* | *PA14_05840* | glutaryl-CoA dehydrogenase | 219 | 365 | 262 | 213 | 325 | 313 | 1 |
| 469 | - | 517400 | 518308 | + | - | *PA14_05850* | LysR family transcriptional regulator | 33 | 32 | 31 | 34 | 30 | 31 | 1 |
| 470 | - | 518460 | 518864 | + | - | *PA14_05860* | hypothetical protein | 271 | 244 | 135 | 142 | 206 | 281 | 1 |
| 471 | - | 520529 | 518907 | - | - | *PA14_05870* | phosphate transporter | 17 | 15 | 13 | 12 | 14 | 18 | 1 |
| 472 | - | 520909 | 522240 | + | - | *PA14_05880* | membrane-bound protease | 225 | 214 | 518 | 274 | 287 | 315 | 1 |
| 473 | - | 522237 | 523031 | + | - | *PA14_05890* | stomatin-like protein | 28 | 34 | 45 | 34 | 37 | 44 | 1 |
| 474 | - | 523139 | 523870 | + | - | *PA14_05910* | periplasmic transport system | 42 | 43 | 49 | 46 | 44 | 45 | 1 |
| 475 | - | 526068 | 523867 | - | - | *PA14_05920* | hypothetical protein | 36 | 38 | 42 | 43 | 38 | 37 | 1 |
| 476 | - | 527706 | 526330 | - | *dbpA* | *PA14_05950* | ATP-dependent RNA helicase DbpA | 41 | 39 | 46 | 34 | 33 | 34 | 1 |
| 477 | - | 528054 | 528263 | + | - | *PA14_05960* | cold-shock protein | 1537 | 1697 | 1142 | 976 | 1077 | 994 | 1 |
| 478 | - | 528414 | 528938 | + | - | *PA14_05970* | hypothetical protein | 96 | 90 | 106 | 90 | 89 | 85 | 1 |
| 479 | - | 530474 | 529041 | - | - | *PA14_05990* | EmrB/QacA family drug resistance transporter | 51 | 50 | 55 | 50 | 49 | 43 | 1 |
| 480 | - | 531095 | 533647 | + | - | *PA14_06000* | ClpA/B protease ATP binding subunit | 539 | 578 | 757 | 555 | 456 | 565 | 1 |
| 481 | - | 533749 | 534327 | + | - | *PA14_06010* | hypothetical protein | 534 | 674 | 639 | 601 | 635 | 674 | 1 |
| 482 | - | 535279 | 535983 | + | - | *PA14_06040* | hypothetical protein | 82 | 70 | 75 | 59 | 75 | 70 | 1 |
| 483 | - | 535331 | 534444 | - | - | *PA14_06030* | acyltransferase | 129 | 136 | 158 | 143 | 136 | 138 | 1 |
| 484 | - | 536068 | 536757 | + | *creB* | *PA14_06060* | DNA-binding response regulator CreB | 22 | 17 | 20 | 17 | 18 | 18 | 1 |
| 485 | - | 536757 | 538181 | + | *creC* | *PA14_06070* | sensory histidine kinase CreC | 32 | 30 | 31 | 29 | 32 | 25 | 1 |
| 486 | - | 538283 | 539641 | + | *creD* | *PA14_06080* | hypothetical protein | 18 | 16 | 16 | 15 | 14 | 15 | 1 |
| 487 | - | 539691 | 539993 | + | - | *PA14_06090* | hypothetical protein | 13 | 13 | 13 | 15 | 13 | 11 | 1 |
| 488 | - | 540758 | 540138 | - | - | *PA14_06120* | hypothetical protein | 37 | 27 | 23 | 25 | 32 | 46 | 1 |
| 489 | - | 541727 | 540777 | - | - | *PA14_06130* | hypothetical protein | 144 | 82 | 93 | 86 | 93 | 129 | 1 |
| 490 | - | 541813 | 542670 | + | - | *PA14_06150* | hypothetical protein | 142 | 149 | 138 | 125 | 156 | 134 | 1 |
| 491 | - | 545236 | 542843 | - | - | *PA14_06160* | hydroxamate-type ferrisiderophore receptor | 20 | 23 | 24 | 37 | 29 | 22 | 1 |
| 492 | - | 546311 | 545340 | - | - | *PA14_06170* | transmembrane sensor | 9 | 28 | 14 | 90 | 53 | 17 | 0.002852411 |
| 493 | - | 546826 | 546308 | - | - | *PA14_06180* | RNA polymerase sigma factor | 10 | 58 | 19 | 235 | 178 | 30 | 5.65402E-21 |
| 494 | - | 546995 | 547744 | + | - | *PA14_06190* | glutathione S-transferase | 23 | 30 | 17 | 17 | 23 | 18 | 1 |
| 495 | - | 547884 | 548288 | + | - | *PA14_06200* | hypothetical protein | 2 | 2 | 2 | 2 | 2 | 1 | 1 |
| 496 | - | 548338 | 548907 | + | - | *PA14_06210* | transcriptional regulator | 22 | 21 | 16 | 19 | 19 | 17 | 1 |
| 497 | - | 550668 | 548941 | - | - | *PA14_06230* | permease | 40 | 36 | 36 | 48 | 38 | 45 | 1 |
| 498 | - | 551942 | 551016 | - | - | *PA14_06240* | LysR family transcriptional regulator | 20 | 17 | 20 | 17 | 18 | 18 | 1 |
| 499 | - | 552030 | 552506 | + | - | *PA14_06250* | GNAT family acetyltransferase | 41 | 43 | 43 | 42 | 45 | 42 | 1 |
| 500 | - | 553523 | 552585 | - | - | *PA14_06260* | LysR family transcriptional regulator | 61 | 56 | 54 | 56 | 49 | 51 | 1 |
| 501 | - | 553627 | 554424 | + | - | *PA14_06270* | hydrolase | 61 | 64 | 75 | 63 | 57 | 58 | 1 |
| 502 | - | 554467 | 554910 | + | - | *PA14_06280* | hypothetical protein | 48 | 47 | 56 | 67 | 45 | 50 | 1 |
| 503 | - | 557135 | 554958 | - | *glcB* | *PA14_06290* | malate synthase G | 228 | 182 | 148 | 145 | 144 | 142 | 1 |
| 504 | - | 557442 | 557885 | + | - | *PA14_06300* | GNAT family acetyltransferase | 133 | 106 | 130 | 124 | 113 | 129 | 1 |

| | A | B | C | D | E | F | G | H | I | J | K | L | M | N |
|---|---|---|---|---|---|---|---|---|---|---|---|---|---|---|
| 505 | - | 557917 | 558432 | + | - | *PA14_06310* | ACT domain-containing protein | 202 | 198 | 266 | 265 | 192 | 263 | 1 |
| 506 | 560072 | 560017 | 559121 | - | - | *PA14_06320* | hypothetical protein | 26 | 34 | 41 | 35 | 28 | 23 | 1 |
| 507 | - | 561194 | 560220 | - | - | *PA14_06330* | serine/threonine protein kinase | 25 | 25 | 27 | 20 | 21 | 21 | 1 |
| 508 | - | 562014 | 561256 | - | - | *PA14_06340* | molybdenum transport regulator | 49 | 37 | 43 | 32 | 44 | 37 | 1 |
| 509 | - | 562402 | 562082 | - | - | *PA14_06350* | hypothetical protein | 43 | 39 | 44 | 48 | 42 | 39 | 1 |
| 510 | - | 563169 | 562444 | - | - | *PA14_06360* | phosphoribosyl transferase | 18 | 13 | 12 | 12 | 15 | 13 | 1 |
| 511 | - | 563601 | 563308 | - | - | *PA14_06390* | hypothetical protein | 166 | 149 | 197 | 157 | 190 | 169 | 1 |
| 512 | - | 564581 | 563655 | - | - | *PA14_06400* | LysR family transcriptional regulator | 17 | 15 | 14 | 12 | 15 | 13 | 1 |
| 513 | - | 564699 | 565457 | + | - | *PA14_06420* | hypothetical protein | 29 | 18 | 16 | 18 | 17 | 18 | 1 |
| 514 | - | 565534 | 565782 | + | - | *PA14_06430* | hypothetical protein | 25 | 13 | 8 | 11 | 10 | 7 | 0.148048673 |
| 515 | - | 565793 | 567169 | + | - | *PA14_06450* | acetyl-CoA carboxylase biotin carboxylase subunit | 31 | 13 | 10 | 11 | 12 | 9 | 0.11934908 |
| 516 | - | 567171 | 568049 | + | - | *PA14_06460* | hypothetical protein | 40 | 19 | 14 | 16 | 19 | 17 | 0.215275599 |
| 517 | - | 568039 | 569016 | + | - | *PA14_06480* | hydrolase | 54 | 22 | 24 | 25 | 23 | 22 | 0.140519471 |
| 518 | - | 569617 | 570675 | + | *bioB* | *PA14_06500* | biotin synthase | 89 | 181 | 50 | 70 | 128 | 81 | 0.843937152 |
| 519 | - | 570781 | 571986 | + | *bioF* | *PA14_06510* | 8-amino-7-oxononanoate synthase | 49 | 63 | 36 | 35 | 48 | 45 | 1 |
| 520 | - | 571979 | 572701 | + | - | *PA14_06530* | biotin biosynthesis protein bioH | 82 | 90 | 59 | 60 | 72 | 75 | 1 |
| 521 | - | 572694 | 573518 | + | - | *PA14_06540* | biotin synthesis protein BioC | 66 | 71 | 40 | 47 | 63 | 58 | 1 |
| 522 | - | 573522 | 574208 | + | *bioD* | *PA14_06570* | dithiobiotin synthetase | 76 | 89 | 60 | 62 | 79 | 83 | 1 |
| 523 | - | 574317 | 574547 | + | - | *PA14_06580* | hypothetical protein | 268 | 283 | 380 | 231 | 271 | 262 | 1 |
| 524 | - | 574888 | 576693 | + | - | *PA14_06600* | acyl-CoA dehydrogenase | 326 | 263 | 364 | 222 | 277 | 333 | 1 |
| 525 | - | 576906 | 578702 | + | - | *PA14_06620* | acyl-CoA dehydrogenase | 21 | 21 | 17 | 17 | 17 | 17 | 1 |
| 526 | - | 578977 | 580755 | + | - | *PA14_06640* | acyl-CoA dehydrogenase | 24 | 23 | 19 | 22 | 25 | 22 | 1 |
| 527 | - | 582569 | 581088 | - | *nirN* | *PA14_06650* | c-type cytochrome | 100 | 177 | 265 | 304 | 428 | 258 | 1 |
| 528 | - | 583399 | 582560 | - | *nirE* | *PA14_06660* | uroporphyrin-III c-methyltransferase | 106 | 253 | 261 | 340 | 538 | 308 | 0.391278558 |
| 529 | - | 584573 | 583410 | - | *nirJ* | *PA14_06670* | heme d1 biosynthesis protein NirJ | 116 | 280 | 256 | 389 | 558 | 361 | 0.723542393 |
| 530 | - | 585082 | 584567 | - | *nirH* | *PA14_06680* | hypothetical protein | 114 | 266 | 246 | 362 | 489 | 318 | 0.232924335 |
| 531 | - | 585500 | 585057 | - | *nirG* | *PA14_06690* | transcriptional regulator | 89 | 223 | 183 | 324 | 424 | 280 | 0.115616918 |
| 532 | - | 586017 | 585493 | - | *nirL* | *PA14_06700* | heme d1 biosynthesis protein NirL | 65 | 180 | 136 | 193 | 275 | 179 | 0.036442662 |
| 533 | - | 586466 | 586014 | - | - | *PA14_06710* | transcriptional regulator | 118 | 312 | 256 | 361 | 476 | 342 | 0.063886935 |
| 534 | - | 587653 | 586475 | - | *nirF* | *PA14_06720* | heme d1 biosynthesis protein NirF | 148 | 361 | 283 | 450 | 598 | 453 | 0.791445601 |
| 535 | - | 588009 | 587650 | - | *nirC* | *PA14_06730* | c-type cytochrome | 109 | 306 | 171 | 265 | 366 | 240 | 0.034724611 |
| 536 | - | 588320 | 588006 | - | *nirM* | *PA14_06740* | cytochrome c-551 | 196 | 810 | 447 | 565 | 890 | 535 | 3.65613E-05 |
| 537 | - | 588321 | 588369 | ? | - | predicted RNA | - | 283 | 1331 | 774 | 890 | 1411 | 1049 | 1.97712E-09 |
| 538 | - | 590077 | 588371 | - | *nirS* | *PA14_06750* | nitrite reductase | 286 | 1211 | 715 | 927 | 1254 | 898 | 0.434308967 |
| 539 | - | 590293 | 591075 | + | *nirQ* | *PA14_06770* | regulatory protein NirQ | 89 | 217 | 64 | 220 | 343 | 280 | 0.242362109 |
| 540 | - | 591110 | 591637 | + | - | *PA14_06790* | cytochrome c oxidase subunit | 22 | 65 | 11 | 81 | 134 | 112 | 0.010140343 |
| 541 | - | 591645 | 591902 | + | - | *PA14_06800* | hypothetical protein | 18 | 35 | 9 | 61 | 96 | 74 | 0.52741753 |
| 542 | - | 591992 | 592432 | + | *norC* | *PA14_06810* | nitric-oxide reductase subunit C | 266 | 702 | 47 | 1090 | 2368 | 1275 | 0.432571835 |
| 543 | - | 592432 | 593829 | + | *norB* | *PA14_06830* | nitric-oxide reductase subunit B | 266 | 554 | 101 | 1082 | 2050 | 1455 | 1 |
| 544 | - | 593831 | 595669 | + | - | *PA14_06840* | dinitrification protein NorD | 141 | 306 | 152 | 627 | 1150 | 1003 | 1 |
| 545 | - | 595870 | 595676 | - | - | *PA14_06860* | hypothetical protein | 510 | 764 | 472 | 1113 | 1036 | 948 | 1 |
| 546 | - | 596637 | 595954 | - | *dnr* | *PA14_06870* | transcriptional regulator Dnr | 73 | 60 | 85 | 95 | 154 | 160 | 1 |
| 547 | - | 596840 | 596963 | + | - | *PA14_06875* | rsmYregulatoryRNA | 135 | 141 | 221 | 90 | 54 | 48 | 1 |
| 548 | - | 597868 | 596990 | - | - | *PA14_06880* | LysR family transcriptional regulator | 10 | 11 | 13 | 8 | 9 | 8 | 1 |
| 549 | - | 597970 | 598671 | + | - | *PA14_06890* | hypothetical protein | 302 | 116 | 260 | 111 | 164 | 114 | 0.499266146 |
| 550 | - | 598668 | 598904 | + | - | *PA14_06900* | hypothetical protein | 118 | 52 | 93 | 43 | 61 | 51 | 0.081032313 |
| 551 | - | 598897 | 600078 | + | - | *PA14_06920* | class III pyridoxal phosphate-dependent aminotransferase | 104 | 42 | 84 | 37 | 57 | 41 | 0.321350893 |
| 552 | - | 600078 | 600794 | + | - | *PA14_06930* | glutamine amidotransferase | 67 | 35 | 60 | 37 | 37 | 37 | 0.828318293 |
| 553 | - | 601375 | 600869 | - | - | *PA14_06940* | hypothetical protein | 44 | 32 | 45 | 46 | 40 | 36 | 1 |
| 554 | - | 601418 | 602908 | + | - | *PA14_06950* | LuxR family transcriptional regulator | 44 | 42 | 42 | 42 | 39 | 38 | 1 |
| 555 | - | 603247 | 604536 | + | - | *PA14_06960* | hypothetical protein | 48 | 48 | 31 | 106 | 93 | 328 | 1 |
| 556 | - | 604554 | 605108 | + | - | *PA14_06970* | Cro/CI family transcriptional regulator | 57 | 45 | 51 | 82 | 65 | 152 | 1 |
| 557 | - | 606244 | 605219 | - | - | *PA14_06980* | hypothetical protein | 122 | 119 | 126 | 107 | 119 | 85 | 1 |
| 558 | - | 606856 | 606248 | - | - | *PA14_06990* | hypothetical protein | 95 | 89 | 89 | 58 | 78 | 53 | 1 |
| 559 | - | 607453 | 606944 | - | *dsbB* | *PA14_07000* | disulfide bond formation protein | 24 | 20 | 22 | 14 | 11 | 10 | 1 |
| 560 | - | 607682 | 608512 | + | - | *PA14_07010* | hypothetical protein | 12 | 10 | 12 | 14 | 15 | 9 | 1 |
| 561 | - | 608582 | 608968 | + | - | *PA14_07020* | hypothetical protein | 25 | 32 | 30 | 30 | 26 | 28 | 1 |
| 562 | - | 609082 | 609540 | + | - | *PA14_07030* | cytochrome c' | 153 | 189 | 176 | 206 | 170 | 134 | 1 |
| 563 | - | 610151 | 609732 | - | - | *PA14_07040* | hypothetical protein | 104 | 120 | 134 | 106 | 104 | 93 | 1 |
| 564 | - | 610401 | 611369 | + | - | *PA14_07050* | hypothetical protein | 75 | 71 | 88 | 95 | 81 | 91 | 1 |
| 565 | - | 612116 | 611373 | - | - | *PA14_07060* | hypothetical protein | 35 | 42 | 52 | 55 | 42 | 53 | 1 |
| 566 | - | 612321 | 613625 | + | - | *PA14_07070* | hypothetical protein | 393 | 272 | 491 | 212 | 171 | 170 | 1 |
| 567 | - | 614872 | 613682 | - | *metK* | *PA14_07090* | S-adenosylmethionine synthetase | 401 | 159 | 147 | 115 | 130 | 280 | 0.981513735 |
| 568 | - | 615889 | 614888 | - | - | *PA14_07110* | ArsR family transcriptional regulator | 435 | 114 | 249 | 154 | 117 | 340 | 0.05276977 |
| 569 | - | 616135 | 618132 | + | *tktA* | *PA14_07130* | transketolase | 191 | 192 | 183 | 167 | 174 | 165 | 1 |
| 570 | - | 618132 | 619196 | + | - | *PA14_07140* | hypothetical protein | 120 | 112 | 118 | 109 | 119 | 108 | 1 |
| 571 | - | 619974 | 619177 | - | - | *PA14_07150* | hypothetical protein | 44 | 40 | 51 | 45 | 46 | 43 | 1 |
| 572 | - | 620189 | 621250 | + | *epd* | *PA14_07170* | D-erythrose 4-phosphate dehydrogenase | 37 | 29 | 37 | 29 | 27 | 30 | 1 |
| 573 | - | 621256 | 622419 | + | *pgk* | *PA14_07190* | phosphoglycerate kinase | 225 | 234 | 210 | 202 | 250 | 173 | 1 |
| 574 | - | 622492 | 622692 | + | - | *PA14_07200* | hypothetical protein | 477 | 516 | 543 | 667 | 508 | 549 | 1 |
| 575 | - | 622852 | 623193 | + | - | *PA14_07210* | hypothetical protein | 150 | 141 | 178 | 174 | 191 | 182 | 1 |
| 576 | - | 623313 | 624377 | + | *fda* | *PA14_07230* | fructose-1,6-bisphosphate aldolase | 192 | 248 | 188 | 187 | 250 | 190 | 1 |
| 577 | - | 624443 | 624925 | + | - | *PA14_07240* | hypothetical protein | 41 | 57 | 46 | 45 | 49 | 45 | 1 |
| 578 | - | 625582 | 626349 | + | - | *PA14_07260* | hypothetical protein | 74 | 74 | 75 | 82 | 73 | 61 | 1 |
| 579 | - | 625583 | 624927 | - | - | *PA14_07250* | hypothetical protein | 15 | 27 | 21 | 26 | 25 | 25 | 0.723249369 |
| 580 | - | 627524 | 626346 | - | - | *PA14_07280* | hypothetical protein | 54 | 52 | 57 | 59 | 55 | 52 | 1 |
| 581 | - | 627637 | 628125 | + | - | *PA14_07290* | hypothetical protein | 35 | 27 | 43 | 29 | 27 | 33 | 1 |
| 582 | - | 628151 | 629305 | + | - | *PA14_07300* | hypothetical protein | 26 | 21 | 23 | 19 | 21 | 21 | 1 |
| 583 | - | 629981 | 629307 | - | - | *PA14_07310* | hydrolase | 28 | 19 | 25 | 23 | 24 | 17 | 1 |
| 584 | - | 630463 | 630110 | - | - | *PA14_07330* | hypothetical protein | 27 | 25 | 20 | 19 | 23 | 18 | 1 |
| 585 | - | 631552 | 630641 | - | - | *PA14_07340* | transcriptional regulator | 17 | 15 | 14 | 12 | 12 | 12 | 1 |
| 586 | - | 631670 | 632008 | + | - | *PA14_07355* | hypothetical protein | 69 | 89 | 68 | 79 | 75 | 83 | 1 |
| 587 | - | 632513 | 631998 | - | - | *PA14_07360* | hypothetical protein | 63 | 79 | 69 | 74 | 69 | 66 | 1 |
| 588 | - | 632701 | 632859 | + | - | *PA14_07370* | hypothetical protein | 179 | 226 | 176 | 191 | 214 | 185 | 1 |

| | A | B | C | D | E | F | G | H | I | J | K | L | M | N |
|---|---|---|---|---|---|---|---|---|---|---|---|---|---|---|
| 589 | - | 633327 | 632869 | - | - | *PA14_07380* | hypothetical protein | 77 | 95 | 69 | 91 | 87 | 79 | 1 |
| 590 | - | 633813 | 633343 | - | - | *PA14_07400* | hypothetical protein | 39 | 37 | 31 | 32 | 28 | 30 | 1 |
| 591 | - | 634174 | 633827 | - | - | *PA14_07410* | hypothetical protein | 15 | 13 | 13 | 15 | 15 | 11 | 1 |
| 592 | - | 634778 | 634164 | - | - | *PA14_07420* | hypothetical protein | 36 | 34 | 33 | 33 | 33 | 31 | 1 |
| 593 | - | 637740 | 634969 | - | - | *PA14_07430* | hypothetical protein | 106 | 199 | 97 | 144 | 158 | 141 | 1 |
| 594 | - | 638644 | 638309 | - | - | *PA14_07440* | hypothetical protein | 19 | 23 | 20 | 24 | 18 | 22 | 1 |
| 595 | - | 639858 | 638737 | - | - | *PA14_07450* | hypothetical protein | 21 | 22 | 20 | 22 | 22 | 18 | 1 |
| 596 | - | 640278 | 640117 | - | - | *PA14_07460* | hypothetical protein | 24 | 30 | 30 | 24 | 23 | 25 | 1 |
| 597 | - | 640483 | 640410 | - | - | *PA14_07470* | Met tRNA | 57 | 49 | 49 | 30 | 26 | 40 | 1 |
| 598 | - | 642266 | 640581 | - | - | *PA14_07480* | reverse transcriptase | 383 | 258 | 176 | 164 | 220 | 258 | 1 |
| 599 | - | 647084 | 643347 | - | - | *PA14_07500* | hypothetical protein | 32 | 33 | 35 | 45 | 39 | 52 | 1 |
| 600 | - | 649044 | 647191 | - | *rpoD* | *PA14_07520* | RNA polymerase sigma factor RpoD | 666 | 623 | 774 | 518 | 589 | 559 | 1 |
| 601 | - | 651118 | 649124 | - | *dnaG* | *PA14_07530* | DNA primase | 80 | 76 | 85 | 79 | 83 | 65 | 1 |
| 602 | - | 651650 | 651201 | - | - | *PA14_07550* | hypothetical protein | 31 | 27 | 37 | 24 | 19 | 15 | 1 |
| 603 | - | 651935 | 651720 | - | *rpsU* | *PA14_07560* | 30S ribosomal protein S21 | 80 | 78 | 86 | 64 | 44 | 36 | 1 |
| 604 | - | 652136 | 653161 | + | *gcp* | *PA14_07570* | DNA-binding/iron metalloprotein/AP endonuclease | 82 | 68 | 91 | 80 | 65 | 63 | 1 |
| 605 | - | 653809 | 653240 | - | - | *PA14_07580* | glycerol-3-phosphate acyltransferase PlsY | 64 | 58 | 73 | 65 | 62 | 53 | 1 |
| 606 | - | 653893 | 654246 | + | *folB* | *PA14_07590* | dihydroneopterin aldolase | 60 | 49 | 67 | 62 | 48 | 56 | 1 |
| 607 | - | 654237 | 654779 | + | - | *PA14_07600* | 2-amino-4-hydroxy-6-hydroxymethyldihydropteridine pyrophosphokinase | 30 | 27 | 34 | 27 | 28 | 28 | 1 |
| 608 | - | 655984 | 654752 | - | *cca* | *PA14_07620* | multifunctional tRNA nucleotidyl transferase/2'3'-cyclic phosphodiesterase/2'nucleotidase/phosphatase | 24 | 24 | 23 | 22 | 26 | 22 | 1 |
| 609 | - | 656534 | 656028 | - | - | *PA14_07630* | hypothetical protein | 11 | 14 | 14 | 18 | 16 | 16 | 1 |
| 610 | - | 658182 | 656629 | - | - | *PA14_07650* | SpoVR family protein | 241 | 286 | 243 | 316 | 283 | 405 | 1 |
| 611 | - | 659450 | 658179 | - | - | *PA14_07660* | hypothetical protein | 297 | 332 | 295 | 352 | 339 | 445 | 1 |
| 612 | - | 661473 | 659551 | - | - | *PA14_07680* | hypothetical protein | 588 | 610 | 591 | 611 | 618 | 747 | 1 |
| 613 | - | 662084 | 661752 | - | *glpE* | *PA14_07690* | thiosulfate sulfurtransferase | 55 | 58 | 52 | 57 | 57 | 65 | 1 |
| 614 | - | 662979 | 662128 | - | *apaH* | *PA14_07700* | diadenosine tetraphosphatase | 86 | 89 | 85 | 68 | 86 | 82 | 1 |
| 615 | - | 663359 | 662979 | - | *apaG* | *PA14_07710* | ApaG protein | 137 | 138 | 133 | 120 | 135 | 143 | 1 |
| 616 | - | 664202 | 663396 | - | *ksgA* | *PA14_07730* | dimethyladenosine transferase | 35 | 29 | 38 | 34 | 36 | 29 | 1 |
| 617 | - | 665304 | 664318 | - | *pdxA* | *PA14_07740* | 4-hydroxythreonine-4-phosphate dehydrogenase | 77 | 90 | 75 | 87 | 91 | 81 | 1 |
| 618 | - | 666593 | 665301 | - | *surA* | *PA14_07760* | peptidyl-prolyl cis-trans isomerase SurA | 267 | 299 | 242 | 273 | 301 | 236 | 1 |
| 619 | - | 669348 | 666574 | - | *ostA* | *PA14_07770* | organic solvent tolerance protein OstA | 253 | 270 | 250 | 262 | 273 | 227 | 1 |
| 620 | - | 669475 | 670491 | + | - | *PA14_07780* | hypothetical protein | 179 | 156 | 186 | 152 | 187 | 196 | 1 |
| 621 | - | 670488 | 671162 | + | - | *PA14_07790* | nucleotidyl transferase | 190 | 198 | 216 | 188 | 205 | 213 | 1 |
| 622 | - | 671164 | 671922 | + | - | *PA14_07800* | hypothetical protein | 192 | 188 | 249 | 229 | 239 | 245 | 1 |
| 623 | - | 672984 | 671923 | - | - | *PA14_07810* | hypothetical protein | 50 | 53 | 55 | 65 | 49 | 52 | 1 |
| 624 | - | 673136 | 675529 | + | - | *PA14_07820* | histidine kinase | 56 | 50 | 56 | 58 | 50 | 46 | 1 |
| 625 | - | 675575 | 676207 | + | - | *PA14_07840* | two-component response regulator | 116 | 99 | 108 | 113 | 99 | 96 | 1 |
| 626 | - | 676336 | 677370 | + | - | *PA14_07850* | ABC transporter substrate-binding protein | 41 | 73 | 44 | 50 | 53 | 38 | 0.718815586 |
| 627 | - | 677604 | 678713 | + | - | *PA14_07860* | ABC transporter ATP-binding protein | 26 | 44 | 29 | 28 | 26 | 22 | 0.76759159 |
| 628 | - | 678769 | 679815 | + | - | *PA14_07870* | ABC transporter substrate-binding protein | 61 | 79 | 61 | 57 | 45 | 47 | 1 |
| 629 | - | 679930 | 681177 | + | - | *PA14_07890* | ABC transporter permease | 20 | 25 | 20 | 19 | 18 | 18 | 1 |
| 630 | - | 681283 | 682113 | + | - | *PA14_07900* | ABC transporter permease | 13 | 15 | 12 | 12 | 11 | 9 | 1 |
| 631 | - | 682237 | 682911 | + | *rpe* | *PA14_07910* | ribulose-phosphate 3-epimerase | 42 | 30 | 36 | 31 | 32 | 26 | 1 |
| 632 | - | 682911 | 683729 | + | - | *PA14_07930* | phosphoglycolate phosphatase | 40 | 37 | 40 | 37 | 39 | 31 | 1 |
| 633 | - | 683802 | 685280 | + | *trpE* | *PA14_07940* | anthranilate synthase component I | 108 | 83 | 99 | 98 | 92 | 97 | 1 |
| 634 | - | 685781 | 685467 | - | *prtN* | *PA14_07950* | transcriptional regulator PrtN | 124 | 186 | 138 | 202 | 186 | 166 | 1 |
| 635 | - | 686651 | 685881 | - | *prtR* | *PA14_07960* | transcriptional regulator PrtR | 60 | 67 | 58 | 59 | 53 | 52 | 1 |
| 636 | - | 687109 | 687309 | + | - | *PA14_07970* | hypothetical protein | 86 | 152 | 89 | 130 | 123 | 87 | 0.580848337 |
| 637 | - | 687357 | 687716 | + | - | *PA14_07980* | hypothetical protein | 69 | 136 | 76 | 114 | 125 | 75 | 0.333226212 |
| 638 | - | 688080 | 688529 | + | - | *PA14_07990* | hypothetical protein | 139 | 217 | 141 | 286 | 331 | 157 | 1 |
| 639 | - | 688551 | 689066 | + | - | *PA14_08000* | hypothetical protein | 284 | 402 | 309 | 506 | 556 | 320 | 1 |
| 640 | - | 689063 | 689620 | + | - | *PA14_08010* | hypothetical protein | 190 | 379 | 232 | 428 | 501 | 241 | 0.986282904 |
| 641 | - | 689773 | 690099 | + | - | *PA14_08020* | bacteriophage protein | 174 | 442 | 245 | 486 | 543 | 246 | 0.10997121 |
| 642 | - | 690096 | 690983 | + | - | *PA14_08030* | phage baseplate assembly protein | 309 | 592 | 336 | 765 | 904 | 415 | 1 |
| 643 | - | 690976 | 691509 | + | - | *PA14_08040* | phage tail protein | 371 | 672 | 390 | 825 | 988 | 469 | 1 |
| 644 | - | 691511 | 693586 | + | - | *PA14_08050* | tail fiber protein | 393 | 749 | 468 | 904 | 1049 | 535 | 1 |
| 645 | - | 693583 | 694041 | + | - | *PA14_08060* | tail fiber assembly protein | 200 | 533 | 329 | 645 | 746 | 348 | 0.221092106 |
| 646 | - | 694084 | 695244 | + | - | *PA14_08070* | phage tail sheath protein | 665 | 1247 | 798 | 1180 | 1409 | 814 | 1 |
| 647 | - | 695257 | 695760 | + | - | *PA14_08090* | phage tail tube protein | 647 | 1219 | 767 | 1157 | 1369 | 784 | 1 |
| 648 | - | 695775 | 696119 | + | - | *PA14_08100* | hypothetical protein | 479 | 854 | 514 | 827 | 976 | 533 | 1 |
| 649 | - | 696088 | 696198 | + | - | *PA14_08110* | hypothetical protein | 337 | 648 | 372 | 723 | 851 | 392 | 0.549291923 |
| 650 | - | 696279 | 696288 | ? | - | predicted RNA | - | 335 | 696 | 386 | 734 | 858 | 393 | 0.214768135 |
| 651 | - | 696289 | 698526 | + | - | *PA14_08120* | tail length determinator protein | 384 | 702 | 456 | 782 | 937 | 442 | 1 |
| 652 | - | 698536 | 699408 | + | - | *PA14_08130* | hypothetical protein | 329 | 558 | 351 | 622 | 757 | 347 | 1 |
| 653 | - | 699383 | 699589 | + | - | *PA14_08140* | hypothetical protein | 246 | 486 | 294 | 524 | 644 | 272 | 0.480861202 |
| 654 | - | 699647 | 700636 | + | - | *PA14_08150* | phage late control gene D protein | 228 | 436 | 252 | 481 | 624 | 281 | 1 |
| 655 | - | 700669 | 701298 | + | - | *PA14_08160* | lytic enzyme | 193 | 305 | 207 | 350 | 425 | 214 | 1 |
| 656 | - | 701295 | 701657 | + | - | *PA14_08180* | hypothetical protein | 266 | 337 | 222 | 383 | 475 | 258 | 1 |
| 657 | - | 701654 | 701911 | + | - | *PA14_08190* | hypothetical protein | 198 | 316 | 184 | 367 | 443 | 216 | 0.934854092 |
| 658 | - | 701926 | 702213 | + | - | *PA14_08200* | hypothetical protein | 176 | 343 | 201 | 289 | 339 | 174 | 0.503207228 |
| 659 | - | 702227 | 702721 | + | - | *PA14_08210* | hypothetical protein | 480 | 863 | 516 | 757 | 869 | 553 | 1 |
| 660 | - | 702733 | 703080 | + | - | *PA14_08220* | hypothetical protein | 442 | 823 | 540 | 797 | 974 | 601 | 1 |
| 661 | - | 703092 | 703109 | ? | - | predicted RNA | - | 542 | 817 | 499 | 811 | 1008 | 607 | 1 |
| 662 | - | 703110 | 703364 | + | - | *PA14_08230* | hypothetical protein | 600 | 938 | 570 | 968 | 1180 | 683 | 1 |
| 663 | - | 703411 | 705246 | + | - | *PA14_08240* | hypothetical protein | 257 | 519 | 310 | 581 | 700 | 342 | 1 |
| 664 | - | 705239 | 705580 | + | - | *PA14_08250* | hypothetical protein | 352 | 667 | 398 | 897 | 1072 | 499 | 1 |
| 665 | - | 705588 | 706283 | + | - | *PA14_08260* | minor tail protein L | 208 | 341 | 217 | 369 | 442 | 236 | 1 |
| 666 | - | 706286 | 707056 | + | - | *PA14_08270* | hypothetical protein | 252 | 392 | 257 | 451 | 555 | 286 | 1 |
| 667 | - | 707111 | 707713 | + | - | *PA14_08280* | bacteriophage protein | 211 | 384 | 231 | 461 | 560 | 264 | 1 |
| 668 | - | 707772 | 711431 | + | - | *PA14_08300* | phage-related protein, tail component | 208 | 343 | 211 | 370 | 469 | 252 | 1 |
| 669 | - | 712461 | 713513 | + | - | *PA14_08310* | hypothetical protein | 162 | 312 | 191 | 274 | 338 | 221 | 1 |
| 670 | - | 713513 | 713815 | + | - | *PA14_08320* | hypothetical protein | 221 | 348 | 249 | 342 | 435 | 314 | 1 |
| 671 | - | 713812 | 714042 | + | - | *PA14_08330* | hypothetical protein | 115 | 212 | 164 | 179 | 234 | 148 | 0.499266146 |
| 672 | - | 714461 | 715066 | + | *trpG* | *PA14_08340* | anthranilate synthase component II | 132 | 117 | 119 | 147 | 141 | 109 | 1 |

| | A | B | C | D | E | F | G | H | I | J | K | L | M | N |
|---|---|---|---|---|---|---|---|---|---|---|---|---|---|---|
| 673 | - | 715068 | 716117 | + | *trpD* | *PA14_08350* | anthranilate phosphoribosyltransferase | 154 | 147 | 159 | 166 | 149 | 156 | 1 |
| 674 | - | 716114 | 716950 | + | *trpC* | *PA14_08360* | indole-3-glycerol-phosphate synthase | 107 | 109 | 83 | 99 | 110 | 92 | 1 |
| 675 | - | 717656 | 717012 | - | *vfr* | *PA14_08370* | cAMP-regulatory protein | 163 | 202 | 144 | 155 | 186 | 129 | 1 |
| 676 | - | 717928 | 718350 | + | - | *PA14_08380* | hypothetical protein | 96 | 85 | 101 | 103 | 100 | 88 | 1 |
| 677 | - | 718670 | 719464 | + | *speD* | *PA14_08390* | S-adenosylmethionine decarboxylase | 32 | 33 | 30 | 33 | 35 | 28 | 1 |
| 678 | - | 720166 | 719519 | - | - | *PA14_08400* | hypothetical protein | 114 | 108 | 125 | 97 | 127 | 120 | 1 |
| 679 | - | 720604 | 720266 | - | - | *PA14_08420* | HIT family protein | 139 | 146 | 175 | 192 | 181 | 224 | 1 |
| 680 | - | 720683 | 722164 | + | - | *PA14_08430* | ATPase | 101 | 116 | 109 | 120 | 108 | 103 | 1 |
| 681 | - | 723004 | 722204 | - | - | *PA14_08440* | short chain alcohol dehydrogenase | 45 | 50 | 42 | 43 | 51 | 37 | 1 |
| 682 | - | 724147 | 723065 | - | - | *PA14_08450* | hypothetical protein | 35 | 43 | 29 | 33 | 45 | 29 | 1 |
| 683 | - | 725237 | 724269 | - | - | *PA14_08460* | hypothetical protein | 48 | 64 | 39 | 53 | 62 | 46 | 1 |
| 684 | - | 725676 | 725254 | - | - | *PA14_08470* | hypothetical protein | 22 | 26 | 19 | 21 | 25 | 19 | 1 |
| 685 | - | 725967 | 727001 | + | *argC* | *PA14_08480* | N-acetyl-gamma-glutamyl-phosphate reductase | 107 | 89 | 104 | 105 | 92 | 93 | 1 |
| 686 | - | 727001 | 727720 | + | - | *PA14_08490* | hypothetical protein | 65 | 72 | 69 | 68 | 72 | 51 | 1 |
| 687 | - | 727721 | 728143 | + | - | *PA14_08500* | hypothetical protein | 128 | 139 | 152 | 127 | 134 | 120 | 1 |
| 688 | - | 728221 | 728571 | + | - | *PA14_08510* | iron-sulfur cluster insertion protein ErpA | 189 | 186 | 121 | 231 | 405 | 288 | 1 |
| 689 | - | 729716 | 728625 | - | *anmK* | *PA14_08520* | anhydro-N-acetylmuramic acid kinase | 86 | 99 | 87 | 100 | 98 | 82 | 1 |
| 690 | - | 731062 | 729719 | - | - | *PA14_08540* | hypothetical protein | 125 | 130 | 119 | 140 | 111 | 108 | 1 |
| 691 | - | 731347 | 732546 | + | *tyrZ* | *PA14_08560* | tyrosyl-tRNA synthetase | 61 | 64 | 67 | 58 | 62 | 50 | 1 |
| 692 | - | 733095 | 734620 | + | - | *PA14_08570* | 16S ribosomal RNA | 237 | 360 | 900 | 310 | 553 | 68 | 1 |
| 693 | - | 734686 | 734759 | + | - | *PA14_08580* | Ile tRNA | 710 | 1071 | 928 | 568 | 435 | 216 | 1 |
| 694 | - | 734791 | 734863 | + | - | *PA14_08590* | Ala tRNA | 696 | 956 | 932 | 643 | 489 | 240 | 1 |
| 695 | - | 735093 | 737983 | + | - | *PA14_08600* | 23S ribosomal RNA | 612 | 872 | 726 | 418 | 753 | 121 | 1 |
| 696 | - | 738120 | 738251 | + | - | *PA14_08610* | 5S ribosomal RNA | 11 | 11 | 27 | 11 | 5 | 2 | 1 |
| 697 | - | 738405 | 739343 | + | *birA* | *PA14_08620* | biotin--protein ligase | 50 | 51 | 54 | 56 | 51 | 42 | 1 |
| 698 | - | 739340 | 740086 | + | - | *PA14_08630* | pantothenate kinase | 52 | 51 | 49 | 48 | 47 | 42 | 1 |
| 699 | - | 740095 | 740799 | + | - | *PA14_08640* | hypothetical protein | 64 | 68 | 72 | 77 | 72 | 61 | 1 |
| 700 | - | 740947 | 741028 | + | - | *PA14_08650* | Tyr tRNA | 409 | 188 | 348 | 211 | 145 | 145 | 0.134543655 |
| 701 | - | 741058 | 741128 | + | - | *PA14_08660* | Gly tRNA | 415 | 195 | 381 | 262 | 173 | 179 | 0.14959202 |
| 702 | - | 741151 | 741223 | + | - | *PA14_08670* | Thr tRNA | 198 | 124 | 206 | 152 | 99 | 167 | 0.771447696 |
| 703 | - | 741311 | 742504 | + | *tufB* | *PA14_08680* | elongation factor Tu | 1616 | 2292 | 1236 | 1010 | 1620 | 947 | 1 |
| 704 | - | 742560 | 742632 | + | - | *PA14_08690* | Trp tRNA | 93 | 146 | 114 | 72 | 68 | 47 | 1 |
| 705 | - | 742681 | 743049 | + | *secE* | *PA14_08695* | preprotein translocase subunit SecE | 283 | 249 | 305 | 207 | 200 | 158 | 1 |
| 706 | - | 743059 | 743592 | + | *nusG* | *PA14_08710* | transcription antitermination protein NusG | 518 | 477 | 571 | 386 | 423 | 328 | 1 |
| 707 | - | 743709 | 744140 | + | *rplK* | *PA14_08720* | 50S ribosomal protein L11 | 559 | 889 | 447 | 358 | 611 | 307 | 1 |
| 708 | - | 744140 | 744835 | + | *rplA* | *PA14_08730* | 50S ribosomal protein L1 | 568 | 893 | 465 | 394 | 679 | 324 | 1 |
| 709 | - | 745022 | 745033 | ? | - | predicted RNA | - | 522 | 790 | 483 | 376 | 673 | 324 | 1 |
| 710 | - | 745034 | 745534 | + | *rplJ* | *PA14_08740* | 50S ribosomal protein L10 | 427 | 611 | 320 | 241 | 422 | 219 | 1 |
| 711 | - | 745613 | 745981 | + | *rplL* | *PA14_08750* | 50S ribosomal protein L7/L12 | 638 | 1049 | 557 | 402 | 597 | 300 | 1 |
| 712 | - | 746203 | 750276 | + | *rpoB* | *PA14_08760* | DNA-directed RNA polymerase subunit beta | 693 | 754 | 556 | 495 | 620 | 437 | 1 |
| 713 | - | 750342 | 754541 | + | *rpoC* | *PA14_08780* | DNA-directed RNA polymerase subunit beta' | 863 | 950 | 603 | 558 | 738 | 533 | 1 |
| 714 | 754615 | 754693 | 755064 | + | *rpsL* | *PA14_08790* | 30S ribosomal protein S12 | 507 | 858 | 453 | 370 | 395 | 271 | 1 |
| 715 | 754615 | 755164 | 755634 | + | *rpsG* | *PA14_08810* | 30S ribosomal protein S7 | 789 | 1192 | 736 | 646 | 865 | 528 | 1 |
| 716 | 754615 | 755665 | 757785 | + | *fusA1* | *PA14_08820* | elongation factor G | 1032 | 1350 | 864 | 775 | 953 | 611 | 1 |
| 717 | 754615 | 757816 | 759009 | + | *tufA* | *PA14_08830* | elongation factor Tu | 1996 | 2750 | 1478 | 1221 | 1985 | 1143 | 1 |
| 718 | - | 759165 | 759476 | + | *rpsJ* | *PA14_08840* | 30S ribosomal protein S10 | 474 | 753 | 498 | 416 | 369 | 276 | 1 |
| 719 | - | 759559 | 760194 | + | *rplC* | *PA14_08850* | 50S ribosomal protein L3 | 653 | 929 | 746 | 531 | 657 | 405 | 1 |
| 720 | - | 760208 | 760810 | + | *rplD* | *PA14_08860* | 50S ribosomal protein L4 | 626 | 925 | 621 | 487 | 653 | 366 | 1 |
| 721 | - | 760807 | 761106 | + | *rplW* | *PA14_08870* | 50S ribosomal protein L23 | 328 | 655 | 309 | 255 | 319 | 191 | 0.912927393 |
| 722 | - | 761118 | 761939 | + | *rplB* | *PA14_08880* | 50S ribosomal protein L2 | 603 | 806 | 494 | 459 | 653 | 346 | 1 |
| 723 | - | 761956 | 762231 | + | *rpsS* | *PA14_08890* | 30S ribosomal protein S19 | 534 | 833 | 433 | 393 | 564 | 302 | 1 |
| 724 | - | 762244 | 762576 | + | *rplV* | *PA14_08900* | 50S ribosomal protein L22 | 630 | 840 | 524 | 461 | 575 | 363 | 1 |
| 725 | - | 762589 | 763275 | + | *rpsC* | *PA14_08910* | 30S ribosomal protein S3 | 682 | 875 | 568 | 402 | 594 | 360 | 1 |
| 726 | - | 763287 | 763700 | + | *rplP* | *PA14_08920* | 50S ribosomal protein L16 | 615 | 804 | 471 | 317 | 557 | 302 | 1 |
| 727 | - | 763700 | 763891 | + | *rpmC* | *PA14_08930* | 50S ribosomal protein L29 | 786 | 933 | 543 | 424 | 762 | 372 | 1 |
| 728 | - | 763894 | 764160 | + | *rpsQ* | *PA14_08940* | 30S ribosomal protein S17 | 438 | 611 | 304 | 230 | 392 | 215 | 1 |
| 729 | - | 764184 | 764552 | + | *rplN* | *PA14_08950* | 50S ribosomal protein L14 | 880 | 1131 | 655 | 564 | 908 | 488 | 1 |
| 730 | - | 764565 | 764879 | + | *rplX* | *PA14_08960* | 50S ribosomal protein L24 | 525 | 677 | 351 | 274 | 391 | 233 | 1 |
| 731 | - | 764899 | 765438 | + | *rplE* | *PA14_08970* | 50S ribosomal protein L5 | 864 | 942 | 597 | 479 | 739 | 465 | 1 |
| 732 | - | 765452 | 765757 | + | *rpsN* | *PA14_08980* | 30S ribosomal protein S14 | 900 | 1116 | 665 | 390 | 638 | 394 | 1 |
| 733 | - | 765762 | 765784 | ? | - | predicted RNA | - | 430 | 589 | 361 | 153 | 210 | 151 | 1 |
| 734 | - | 765947 | 766339 | + | *rpsH* | *PA14_08990* | 30S ribosomal protein S8 | 377 | 402 | 352 | 288 | 291 | 219 | 1 |
| 735 | - | 766351 | 766884 | + | *rplF* | *PA14_09000* | 50S ribosomal protein L6 | 573 | 686 | 537 | 515 | 566 | 405 | 1 |
| 736 | - | 766895 | 767245 | + | *rplR* | *PA14_09010* | 50S ribosomal protein L18 | 744 | 1027 | 730 | 636 | 683 | 466 | 1 |
| 737 | - | 767249 | 767749 | + | *rpsE* | *PA14_09020* | 30S ribosomal protein S5 | 1418 | 1910 | 1362 | 1400 | 1696 | 1071 | 1 |
| 738 | - | 767752 | 767928 | + | *rpmD* | *PA14_09030* | 50S ribosomal protein L30 | 1094 | 1443 | 890 | 883 | 1004 | 664 | 1 |
| 739 | - | 767932 | 768366 | + | *rplO* | *PA14_09040* | 50S ribosomal protein L15 | 1189 | 1506 | 941 | 924 | 1102 | 694 | 1 |
| 740 | - | 768367 | 769695 | + | *secY* | *PA14_09050* | preprotein translocase subunit SecY | 946 | 1024 | 832 | 729 | 839 | 563 | 1 |
| 741 | - | 769724 | 769840 | + | *rpmJ* | *PA14_09070* | 50S ribosomal protein L36 | 1672 | 1985 | 1811 | 1465 | 1394 | 1119 | 1 |
| 742 | - | 769850 | 769902 | ? | - | predicted RNA | - | 1195 | 1431 | 1095 | 827 | 889 | 677 | 1 |
| 743 | - | 769971 | 770327 | + | *rpsM* | *PA14_09080* | 30S ribosomal protein S13 | 762 | 831 | 602 | 496 | 608 | 467 | 1 |
| 744 | - | 770346 | 770735 | + | *rpsK* | *PA14_09090* | 30S ribosomal protein S11 | 524 | 766 | 415 | 343 | 459 | 321 | 1 |
| 745 | - | 770752 | 771372 | + | *rpsD* | *PA14_09100* | 30S ribosomal protein S4 | 609 | 884 | 448 | 412 | 501 | 369 | 1 |
| 746 | - | 771395 | 772396 | + | *rpoA* | *PA14_09115* | DNA-directed RNA polymerase subunit alpha | 1067 | 1192 | 952 | 814 | 835 | 628 | 1 |
| 747 | - | 772440 | 772829 | + | *rplQ* | *PA14_09130* | 50S ribosomal protein L17 | 468 | 674 | 326 | 296 | 385 | 282 | 1 |
| 748 | - | 773110 | 774558 | + | *katA* | *PA14_09150* | catalase | 460 | 464 | 341 | 182 | 326 | 565 | 1 |
| 749 | - | 774689 | 775153 | + | *bfrA* | *PA14_09160* | bacterioferritin | 164 | 193 | 135 | 196 | 221 | 182 | 1 |
| 750 | - | 778062 | 775225 | - | *uvrA* | *PA14_09180* | excinuclease ABC subunit A | 70 | 87 | 77 | 69 | 81 | 61 | 1 |
| 751 | - | 778276 | 779664 | + | - | *PA14_09195* | major facilitator transporter | 52 | 43 | 54 | 47 | 44 | 37 | 1 |
| 752 | - | 779681 | 780178 | + | *ssb* | *PA14_09200* | single-stranded DNA-binding protein | 168 | 214 | 152 | 151 | 175 | 135 | 1 |
| 753 | - | 781697 | 780267 | - | *pchA* | *PA14_09210* | salicylate biosynthesis isochorismate synthase | 86 | 227 | 85 | 251 | 249 | 207 | 0.460197005 |
| 754 | - | 781999 | 781694 | - | *pchB* | *PA14_09220* | isochorismate-pyruvate lyase | 74 | 255 | 61 | 287 | 254 | 217 | 0.000294392 |
| 755 | - | 782754 | 781999 | - | *pchC* | *PA14_09230* | pyochelin biosynthetic protein PchC | 35 | 97 | 33 | 176 | 134 | 96 | 0.01946313 |
| 756 | - | 784394 | 782751 | - | *pchD* | *PA14_09240* | pyochelin biosynthesis protein PchD | 40 | 132 | 42 | 294 | 194 | 118 | 0.007836446 |

| | A | B | C | D | E | F | G | H | I | J | K | L | M | N |
|---|---|---|---|---|---|---|---|---|---|---|---|---|---|---|
| 757 | - | 784624 | 785514 | + | *pchR* | *PA14_09260* | transcriptional regulator PchR | 19 | 54 | 23 | 62 | 50 | 33 | 0.007773202 |
| 758 | - | 785708 | 790024 | + | *pchE* | *PA14_09270* | dihydroaeruginoic acid synthetase | 93 | 238 | 99 | 295 | 291 | 193 | 1 |
| 759 | - | 790021 | 795450 | + | *pchF* | *PA14_09280* | pyochelin synthetase | 119 | 233 | 111 | 192 | 242 | 210 | 1 |
| 760 | - | 795447 | 796496 | + | *pchG* | *PA14_09290* | pyochelin biosynthetic protein PchG | 153 | 255 | 120 | 203 | 253 | 257 | 1 |
| 761 | - | 796493 | 798205 | + | - | *PA14_09300* | ABC transporter ATP-binding protein | 97 | 147 | 72 | 117 | 159 | 145 | 1 |
| 762 | - | 798202 | 799926 | + | - | *PA14_09320* | ABC transporter ATP-binding protein | 173 | 223 | 129 | 171 | 239 | 216 | 1 |
| 763 | - | 800018 | 802180 | + | *fptA* | *PA14_09340* | Fe(III)-pyochelin outer membrane receptor | 101 | 329 | 102 | 665 | 468 | 190 | 0.401171332 |
| 764 | - | 802180 | 802461 | + | - | *PA14_09350* | hypothetical protein | 153 | 314 | 162 | 561 | 477 | 256 | 0.395977757 |
| 765 | - | 802793 | 803968 | + | - | *PA14_09370* | hypothetical protein | 59 | 92 | 45 | 145 | 154 | 78 | 1 |
| 766 | - | 803961 | 805205 | + | - | *PA14_09380* | transporter | 110 | 137 | 56 | 137 | 170 | 123 | 1 |
| 767 | - | 806518 | 805310 | - | *phzS* | *PA14_09400* | hypothetical protein | 943 | 51 | 51 | 63 | 97 | 678 | 5.8118E-125 |
| 768 | 813529 | 807503 | 806856 | - | *phzG1* | *PA14_09410* | pyrodoxamine 5'-phosphate oxidase | 907 | 10 | 30 | 58 | 70 | 705 | 0 |
| 769 | 813529 | 808362 | 807526 | - | *phzF1* | *PA14_09420* | phenazine biosynthesis protein | 899 | 10 | 27 | 57 | 72 | 708 | 0 |
| 770 | 813529 | 810259 | 808376 | - | *phzE1* | *PA14_09440* | phenazine biosynthesis protein PhzE | 1250 | 15 | 44 | 94 | 98 | 969 | 0 |
| 771 | 813529 | 810879 | 810256 | - | *phzD1* | *PA14_09450* | phenazine biosynthesis protein PhzD | 968 | 6 | 21 | 69 | 82 | 748 | 0 |
| 772 | 812427 | 812093 | 810876 | - | *phzC1* | *PA14_09460* | phenazine biosynthesis protein PhzC | 1244 | 8 | 39 | 59 | 72 | 832 | 0 |
| 773 | 812427 | 812617 | 812129 | - | *phzB1* | *PA14_09470* | phenazine biosynthesis protein | 528 | 1 | 11 | 17 | 43 | 415 | 0 |
| 774 | 813529 | 813135 | 812647 | - | *phzA1* | *PA14_09480* | phenazine biosynthesis protein | 123 | 0 | 7 | 7 | 6 | 95 | 0 |
| 775 | 813529 | 813831 | 814835 | + | *phzM* | *PA14_09490* | phenazine-specific methyltransferase | 284 | 14 | 40 | 33 | 33 | 358 | 2.1957E-299 |
| 776 | - | 816309 | 814846 | - | *opmD* | *PA14_09500* | outer membrane protein | 488 | 22 | 35 | 30 | 31 | 605 | 8.6398E-244 |
| 777 | - | 819395 | 816306 | - | *mexI* | *PA14_09520* | RND efflux transporter | 340 | 17 | 16 | 17 | 17 | 366 | 3.1109E-142 |
| 778 | - | 820520 | 819408 | - | *mexH* | *PA14_09530* | RND efflux membrane fusion protein | 286 | 14 | 11 | 12 | 11 | 301 | 2.0372E-284 |
| 779 | - | 820974 | 820528 | - | *mexG* | *PA14_09540* | hypothetical protein | 330 | 22 | 15 | 18 | 16 | 315 | 1.4046E-178 |
| 780 | - | 822791 | 821625 | - | - | *PA14_09550* | hypothetical protein | 48 | 39 | 39 | 37 | 48 | 42 | 1 |
| 781 | - | 823725 | 822871 | - | - | *PA14_09570* | LysR family transcriptional regulator | 25 | 22 | 20 | 21 | 22 | 28 | 1 |
| 782 | - | 823833 | 824888 | + | - | *PA14_09580* | hypothetical protein | 32 | 38 | 26 | 23 | 35 | 33 | 1 |
| 783 | - | 824984 | 826024 | + | *ddlA* | *PA14_09600* | D-alanine--D-alanine ligase | 190 | 149 | 147 | 130 | 138 | 189 | 1 |
| 784 | - | 826930 | 826064 | - | - | *PA14_09610* | hypothetical protein | 65 | 58 | 57 | 32 | 54 | 52 | 1 |
| 785 | - | 828919 | 827138 | - | - | *PA14_09630* | acyl-CoA dehydrogenase | 154 | 118 | 120 | 59 | 88 | 96 | 1 |
| 786 | - | 830659 | 829037 | - | - | *PA14_09660* | acyl-CoA synthetase | 105 | 97 | 92 | 50 | 78 | 65 | 1 |
| 787 | - | 830905 | 833181 | + | - | *PA14_09680* | two-component sensor | 71 | 70 | 62 | 41 | 69 | 58 | 1 |
| 788 | - | 833184 | 833828 | + | - | *PA14_09690* | two-component response regulator | 109 | 109 | 100 | 81 | 117 | 121 | 1 |
| 789 | - | 835038 | 833833 | - | - | *PA14_09700* | monooxygenase | 57 | 73 | 52 | 46 | 68 | 58 | 1 |
| 790 | - | 836696 | 835206 | - | - | *PA14_09710* | aldehyde dehydrogenase | 7 | 6 | 5 | 5 | 6 | 6 | 1 |
| 791 | - | 837600 | 836689 | - | - | *PA14_09730* | dihydrodipicolinate synthase | 6 | 6 | 5 | 3 | 5 | 5 | 1 |
| 792 | - | 838993 | 837665 | - | - | *PA14_09740* | MFS transporter | 7 | 7 | 7 | 7 | 6 | 7 | 1 |
| 793 | - | 840448 | 839129 | - | - | *PA14_09750* | oxidoreductase | 70 | 64 | 85 | 74 | 60 | 62 | 1 |
| 794 | - | 841270 | 840542 | - | - | *PA14_09760* | GntR family transcriptional regulator | 22 | 20 | 22 | 19 | 17 | 18 | 1 |
| 795 | - | 841579 | 842601 | + | - | *PA14_09770* | AraC family transcriptional regulator | 37 | 37 | 33 | 37 | 41 | 38 | 1 |
| 796 | - | 842627 | 843094 | + | - | *PA14_09780* | hypothetical protein | 36 | 40 | 39 | 29 | 36 | 37 | 1 |
| 797 | - | 843736 | 843098 | - | - | *PA14_09790* | transcriptional regulator | 54 | 72 | 53 | 52 | 58 | 52 | 1 |
| 798 | - | 844483 | 843764 | - | - | *PA14_09810* | hypothetical protein | 26 | 22 | 23 | 22 | 20 | 18 | 1 |
| 799 | - | 844784 | 846427 | + | - | *PA14_09820* | acetolactate synthase | 84 | 81 | 68 | 73 | 84 | 76 | 1 |
| 800 | - | 847765 | 846470 | - | - | *PA14_09850* | porin | 40 | 31 | 31 | 29 | 32 | 36 | 1 |
| 801 | - | 848746 | 848003 | - | - | *PA14_09870* | hypothetical protein | 5 | 5 | 6 | 5 | 3 | 4 | 1 |
| 802 | - | 849247 | 848846 | - | - | *PA14_09880* | hypothetical protein | 4 | 4 | 4 | 5 | 5 | 4 | 1 |
| 803 | - | 849390 | 849671 | + | *ppiC2* | *PA14_09890* | peptidyl-prolyl cis-trans isomerase C2 | 230 | 336 | 233 | 214 | 268 | 196 | 1 |
| 804 | - | 851380 | 849992 | - | *prpL* | *PA14_09900* | Pvds-regulated endoprotease, lysyl class | 93 | 141 | 93 | 141 | 125 | 148 | 1 |
| 805 | - | 852836 | 851913 | - | - | *PA14_09910* | transcriptional regulator | 36 | 55 | 44 | 63 | 51 | 57 | 1 |
| 806 | - | 852977 | 853375 | + | - | *PA14_09920* | translation initiation inhibitor | 4 | 5 | 2 | 3 | 3 | 3 | 1 |
| 807 | - | 854207 | 853407 | - | - | *PA14_09930* | exonuclease III | 9 | 12 | 11 | 12 | 12 | 9 | 1 |
| 808 | - | 854793 | 854230 | - | - | *PA14_09940* | protease | 19 | 32 | 32 | 28 | 27 | 22 | 1 |
| 809 | - | 855861 | 854929 | - | - | *PA14_09950* | oxidoreductase | 17 | 13 | 15 | 20 | 18 | 16 | 1 |
| 810 | - | 855942 | 856370 | + | - | *PA14_09960* | hypothetical protein | 12 | 13 | 14 | 13 | 12 | 9 | 1 |
| 811 | - | 858843 | 856435 | - | *fpvB* | *PA14_09970* | type I ferripyoverdine receptor, FpvB | 27 | 28 | 28 | 40 | 26 | 29 | 1 |
| 812 | - | 859186 | 860004 | + | *dkgB* | *PA14_09980* | 2,5-diketo-D-gluconate reductase B | 3 | 3 | 2 | 6 | 3 | 2 | 1 |
| 813 | - | 860512 | 860054 | - | - | *PA14_09990* | acetyltransferase | 3 | 4 | 3 | 4 | 4 | 6 | 1 |
| 814 | - | 860644 | 862134 | + | - | *PA14_10010* | transcriptional regulator | 28 | 27 | 22 | 23 | 26 | 24 | 1 |
| 815 | - | 862476 | 862150 | - | - | *PA14_10020* | hypothetical protein | 61 | 76 | 53 | 52 | 74 | 66 | 1 |
| 816 | - | 864247 | 862538 | - | - | *PA14_10040* | amidase | 66 | 86 | 61 | 54 | 75 | 56 | 1 |
| 817 | - | 864931 | 864434 | - | - | *PA14_10050* | hypothetical protein | 3 | 4 | 3 | 4 | 4 | 4 | 1 |
| 818 | - | 865931 | 864945 | - | - | *PA14_10070* | zinc-dependent oxidoreductase | 6 | 5 | 4 | 5 | 6 | 6 | 1 |
| 819 | - | 866321 | 865944 | - | - | *PA14_10080* | hypothetical protein | 24 | 23 | 22 | 33 | 21 | 26 | 1 |
| 820 | - | 866440 | 867351 | + | - | *PA14_10090* | LysR family transcriptional regulator | 46 | 40 | 47 | 41 | 44 | 45 | 1 |
| 821 | - | 868546 | 867362 | - | - | *PA14_10110* | transporter | 10 | 10 | 10 | 11 | 8 | 12 | 1 |
| 822 | - | 869571 | 868642 | - | - | *PA14_10120* | LysR family transcriptional regulator | 24 | 21 | 24 | 21 | 18 | 19 | 1 |
| 823 | - | 869711 | 870427 | + | - | *PA14_10130* | short chain dehydrogenase | 33 | 29 | 31 | 33 | 31 | 30 | 1 |
| 824 | - | 871468 | 870437 | - | *fepG* | *PA14_10140* | ferric enterobactin transport protein FepG | 16 | 13 | 13 | 19 | 14 | 11 | 1 |
| 825 | - | 872487 | 871465 | - | *fepD* | *PA14_10160* | ferric enterobactin transport protein FepD | 2 | 4 | 2 | 11 | 6 | 1 | 0.829483661 |
| 826 | - | 873423 | 872518 | - | *fepB* | *PA14_10170* | iron-enterobactin transporter periplasmic binding protein | 5 | 12 | 6 | 49 | 20 | 5 | 0.059785741 |
| 827 | - | 873549 | 874346 | + | *fepC* | *PA14_10180* | ferric enterobactin transport protein FepC | 5 | 10 | 4 | 14 | 12 | 7 | 0.760328835 |
| 828 | - | 875141 | 874353 | - | - | *PA14_10190* | transcriptional regulator | 23 | 26 | 22 | 25 | 23 | 20 | 1 |
| 829 | - | 875397 | 877481 | + | - | *PA14_10200* | TonB-dependent receptor protein | 7 | 11 | 7 | 19 | 11 | 8 | 1 |
| 830 | - | 877549 | 878856 | + | - | *PA14_10210* | oxidoreductase | 3 | 5 | 3 | 6 | 5 | 4 | 1 |
| 831 | - | 879293 | 879961 | + | - | *PA14_10220* | SH3 domain-containing protein | 9 | 7 | 13 | 7 | 9 | 5 | 1 |
| 832 | - | 881136 | 880045 | - | *adh* | *PA14_10230* | 2,3-butanediol dehydrogenase | 5 | 6 | 6 | 5 | 6 | 5 | 1 |
| 833 | - | 882262 | 881150 | - | - | *PA14_10240* | branched-chain alpha-keto acid dehydrogenase subunit E2 | 10 | 9 | 8 | 11 | 9 | 11 | 1 |
| 834 | - | 883278 | 882259 | - | *acoB* | *PA14_10250* | acetoin catabolism protein AcoB | 11 | 12 | 11 | 14 | 11 | 10 | 1 |
| 835 | - | 884285 | 883311 | - | - | *PA14_10260* | dehydrogenase E1 component | 27 | 25 | 29 | 43 | 30 | 32 | 1 |
| 836 | - | 885348 | 884308 | - | - | *PA14_10270* | hypothetical protein | 12 | 12 | 12 | 11 | 12 | 10 | 1 |
| 837 | - | 886145 | 885345 | - | - | *PA14_10280* | short-chain dehydrogenase | 8 | 9 | 9 | 7 | 8 | 10 | 1 |
| 838 | - | 886477 | 888354 | + | *acoR* | *PA14_10290* | transcriptional regulator AcoR | 68 | 79 | 81 | 82 | 73 | 96 | 1 |
| 839 | 895433 | 888559 | 889170 | + | - | *PA14_10300* | efflux protein | 15 | 12 | 12 | 13 | 12 | 17 | 1 |
| 840 | 895433 | 890047 | 889157 | - | - | *PA14_10320* | transcriptional regulator | 33 | 27 | 23 | 24 | 23 | 28 | 1 |

| | A | B | C | D | E | F | G | H | I | J | K | L | M | N |
|---|---|---|---|---|---|---|---|---|---|---|---|---|---|---|
| 841 | 895433 | 891558 | 890143 | - | - | *PA14_10330* | outer membrane protein | 28 | 6 | 6 | 20 | 15 | 37 | 3.37386E-14 |
| 842 | 895433 | 893717 | 891558 | - | - | *PA14_10340* | toxin transporter | 38 | 7 | 10 | 24 | 16 | 37 | 6.42463E-16 |
| 843 | 895433 | 894983 | 893727 | - | - | *PA14_10350* | secretion protein | 84 | 7 | 13 | 41 | 30 | 78 | 1.8387E-103 |
| 844 | 895433 | 895379 | 895080 | - | - | *PA14_10360* | hypothetical protein | 6793 | 234 | 525 | 3591 | 3008 | 10535 | 0 |
| 845 | - | 895384 | 895404 | ? | - | predicted RNA | - | 1121 | 33 | 42 | 324 | 234 | 1288 | 0 |
| 846 | - | 897481 | 895688 | - | - | *PA14_10370* | hypothetical protein | 11 | 6 | 6 | 7 | 5 | 10 | 0.452603163 |
| 847 | - | 897912 | 897613 | - | - | *PA14_10380* | hypothetical protein | 32 | 26 | 11 | 17 | 21 | 37 | 1 |
| 848 | - | 898251 | 898045 | - | - | *PA14_10400* | hypothetical protein | 1 | 0 | 0 | 1 | 0 | 1 | 3.0938E-17 |
| 849 | - | 898562 | 898269 | - | - | *PA14_10410* | hypothetical protein | 0 | 0 | 0 | 0 | 0 | 0 | 0.013995908 |
| 850 | - | 898671 | 899909 | + | *tyrS* | *PA14_10420* | tyrosyl-tRNA synthetase | 5 | 6 | 4 | 6 | 5 | 4 | 1 |
| 851 | - | 901194 | 899938 | - | - | *PA14_10440* | porin | 15 | 11 | 14 | 15 | 12 | 13 | 1 |
| 852 | - | 901559 | 902767 | + | - | *PA14_10470* | MFS transporter | 60 | 36 | 53 | 50 | 43 | 45 | 1 |
| 853 | - | 902853 | 903275 | + | - | *PA14_10480* | transcriptional regulator | 69 | 52 | 65 | 66 | 57 | 58 | 1 |
| 854 | - | 903533 | 903291 | - | - | *PA14_10490* | hypothetical protein | 331 | 279 | 304 | 347 | 333 | 302 | 1 |
| 855 | - | 905044 | 903617 | - | - | *PA14_10500* | cbb3-type cytochrome c oxidase subunit I | 624 | 653 | 556 | 611 | 750 | 522 | 1 |
| 856 | - | 906723 | 905308 | - | - | *PA14_10530* | GntR family transcriptional regulator | 895 | 854 | 805 | 1222 | 1038 | 972 | 1 |
| 857 | - | 906810 | 906820 | ? | - | predicted RNA | - | 689 | 713 | 542 | 814 | 811 | 737 | 1 |
| 858 | - | 908542 | 906821 | - | - | *PA14_10540* | iron-sulfur cluster-binding protein | 305 | 369 | 253 | 379 | 451 | 312 | 1 |
| 859 | - | 908648 | 910321 | + | - | *PA14_10550* | sulfite or nitrite reductas | 615 | 556 | 458 | 711 | 689 | 676 | 1 |
| 860 | - | 910318 | 910806 | + | - | *PA14_10560* | hypothetical protein | 899 | 769 | 695 | 1053 | 932 | 977 | 1 |
| 861 | - | 911694 | 910888 | - | - | *PA14_10570* | 2,4-dihydroxyhept-2-ene-1,7-dioic acid aldolase | 341 | 261 | 280 | 387 | 352 | 401 | 1 |
| 862 | - | 912510 | 911707 | - | *hpcG* | *PA14_10590* | 2-oxo-hepta-3-ene-1,7-dioic acid hydratase | 124 | 98 | 113 | 145 | 132 | 151 | 1 |
| 863 | - | 913832 | 912528 | - | - | *PA14_10600* | MFS transporter | 50 | 39 | 50 | 68 | 70 | 79 | 1 |
| 864 | - | 914317 | 913925 | - | *hpcD* | *PA14_10610* | 5-carboxymethyl-2-hydroxymuconate isomerase | 39 | 33 | 38 | 41 | 32 | 34 | 1 |
| 865 | - | 915251 | 914328 | - | *hpcB* | *PA14_10620* | homoprotocatechuate 2,3-dioxygenase | 21 | 20 | 22 | 28 | 20 | 20 | 1 |
| 866 | - | 916843 | 915383 | - | *hpcC* | *PA14_10630* | 5-carboxy-2-hydroxymuconate semialdehyde dehydrogenase | 12 | 11 | 11 | 13 | 9 | 13 | 1 |
| 867 | - | 917619 | 916840 | - | - | *PA14_10640* | hypothetical protein | 7 | 6 | 6 | 8 | 7 | 7 | 1 |
| 868 | - | 918289 | 917630 | - | - | *PA14_10650* | hypothetical protein | 7 | 7 | 9 | 7 | 5 | 6 | 1 |
| 869 | - | 918509 | 919420 | + | - | *PA14_10660* | transcriptional regulator | 13 | 14 | 14 | 14 | 12 | 14 | 1 |
| 870 | - | 919472 | 920275 | + | *aph* | *PA14_10670* | aminoglycoside 3'-phosphotransferase type IIB | 17 | 20 | 22 | 22 | 18 | 17 | 1 |
| 871 | - | 920968 | 920399 | - | - | *PA14_10680* | hypothetical protein | 52 | 48 | 56 | 57 | 55 | 58 | 1 |
| 872 | - | 923232 | 921046 | - | - | *PA14_10700* | bacteriophytochrome | 151 | 184 | 194 | 212 | 185 | 209 | 1 |
| 873 | - | 923847 | 923260 | - | - | *PA14_10710* | hypothetical protein | 216 | 236 | 327 | 390 | 266 | 284 | 1 |
| 874 | - | 925350 | 923965 | - | - | *PA14_10730* | hypothetical protein | 132 | 111 | 137 | 120 | 135 | 146 | 1 |
| 875 | - | 925770 | 926279 | + | - | *PA14_10740* | spermidine acetyltransferase | 36 | 42 | 32 | 30 | 36 | 40 | 1 |
| 876 | - | 926433 | 927623 | + | - | *PA14_10750* | sugar efflux transporter | 11 | 7 | 13 | 10 | 10 | 8 | 0.543961598 |
| 877 | - | 927751 | 932004 | + | - | *PA14_10770* | sensor/response regulator hybrid | 121 | 112 | 149 | 183 | 131 | 168 | 1 |
| 878 | - | 932566 | 932168 | - | - | *PA14_10780* | hypothetical protein | 65 | 77 | 91 | 86 | 82 | 78 | 1 |
| 879 | - | 933824 | 932631 | - | *ampC* | *PA14_10790* | beta-lactamase | 13 | 14 | 14 | 14 | 13 | 14 | 1 |
| 880 | - | 933973 | 934863 | + | *ampR* | *PA14_10800* | transcriptional regulator AmpR | 42 | 43 | 43 | 56 | 47 | 46 | 1 |
| 881 | - | 936666 | 935422 | - | - | *PA14_10820* | HDIG domain-containing protein | 133 | 121 | 154 | 105 | 138 | 146 | 1 |
| 882 | - | 937824 | 936949 | - | - | *PA14_10830* | LysR family transcriptional regulator | 74 | 73 | 78 | 65 | 79 | 73 | 1 |
| 883 | - | 937983 | 938735 | + | - | *PA14_10840* | dehydrogenase | 38 | 39 | 36 | 34 | 45 | 36 | 1 |
| 884 | - | 940431 | 938752 | - | - | *PA14_10850* | dehydrogenase | 26 | 29 | 24 | 27 | 27 | 25 | 1 |
| 885 | - | 941890 | 940586 | - | - | *PA14_10870* | hypothetical protein | 11 | 9 | 9 | 13 | 7 | 9 | 1 |
| 886 | - | 942700 | 941975 | - | - | *PA14_10890* | short-chain dehydrogenase | 9 | 11 | 12 | 14 | 10 | 10 | 1 |
| 887 | - | 943773 | 942715 | - | - | *PA14_10900* | alcohol dehydrogenase | 3 | 3 | 3 | 5 | 3 | 2 | 1 |
| 888 | - | 945029 | 943770 | - | - | *PA14_10910* | major facilitator transporter | 3 | 4 | 3 | 3 | 3 | 2 | 1 |
| 889 | - | 945633 | 945100 | - | - | *PA14_10920* | hypothetical protein | 3 | 3 | 4 | 5 | 3 | 4 | 1 |
| 890 | - | 945954 | 946913 | + | - | *PA14_10940* | AraC family transcriptional regulator | 25 | 26 | 26 | 28 | 27 | 30 | 1 |
| 891 | - | 946961 | 947377 | + | - | *PA14_10950* | hypothetical protein | 10 | 11 | 6 | 10 | 7 | 12 | 1 |
| 892 | - | 948499 | 947387 | - | - | *PA14_10960* | ferredoxin oxidoreductase subunit | 10 | 9 | 8 | 10 | 9 | 11 | 1 |
| 893 | - | 949493 | 948528 | - | - | *PA14_10970* | flavodoxin reductase | 10 | 8 | 10 | 10 | 9 | 10 | 1 |
| 894 | - | 949604 | 950458 | + | - | *PA14_10980* | hypothetical protein | 11 | 9 | 11 | 9 | 10 | 11 | 1 |
| 895 | - | 951014 | 950502 | - | *hpaC* | *PA14_10990* | 4-hydroxyphenylacetate 3-monooxygenase small subunit | 4 | 5 | 3 | 5 | 4 | 4 | 1 |
| 896 | - | 952606 | 951044 | - | *hpaA* | *PA14_11000* | 4-hydroxyphenylacetate 3-monooxygenase large chain | 5 | 6 | 6 | 6 | 6 | 6 | 1 |
| 897 | - | 953062 | 952772 | - | - | *PA14_11010* | hypothetical protein | 16 | 27 | 15 | 33 | 33 | 21 | 1 |
| 898 | - | 954126 | 953365 | - | - | *PA14_11020* | 3-ketoacyl-ACP reductase | 16 | 17 | 17 | 21 | 13 | 17 | 1 |
| 899 | - | 955508 | 954123 | - | - | *PA14_11030* | aminotransferase | 5 | 7 | 6 | 8 | 5 | 7 | 1 |
| 900 | - | 956341 | 955517 | - | - | *PA14_11050* | hypothetical protein | 1 | 2 | 1 | 2 | 2 | 2 | 0.129649142 |
| 901 | - | 956812 | 957381 | + | *cupB1* | *PA14_11060* | fimbrial subunit CupB1 | 12 | 9 | 8 | 6 | 11 | 10 | 1 |
| 902 | - | 957412 | 958158 | + | *cupB2* | *PA14_11070* | chaperone CupB2 | 1 | 1 | 2 | 1 | 1 | 1 | 0.842469071 |
| 903 | - | 958480 | 961014 | + | *cupB3* | *PA14_11080* | usher CupB3 | 16 | 20 | 21 | 24 | 21 | 25 | 1 |
| 904 | - | 961011 | 961751 | + | *cupB4* | *PA14_11090* | chaperone CupB4 | 2 | 2 | 2 | 1 | 1 | 1 | 1 |
| 905 | - | 961839 | 964895 | + | *cupB5* | *PA14_11100* | adhesive protein CupB5 | 5 | 6 | 5 | 6 | 6 | 5 | 1 |
| 906 | - | 964995 | 966140 | + | *cupB6* | *PA14_11110* | fimbrial protein cupB6 | 6 | 6 | 6 | 6 | 6 | 6 | 1 |
| 907 | - | 966785 | 967429 | + | - | *PA14_11120* | response regulator | 40 | 46 | 35 | 47 | 50 | 56 | 1 |
| 908 | - | 968133 | 967444 | - | - | *PA14_11130* | short chain dehydrogenase | 135 | 70 | 78 | 97 | 88 | 178 | 0.785036578 |
| 909 | - | 971198 | 968223 | - | - | *PA14_11140* | nonribosomal peptide synthetase | 109 | 38 | 42 | 68 | 45 | 141 | 0.452109894 |
| 910 | - | 971615 | 971409 | - | - | *PA14_11150* | transcriptional regulator | 7 | 11 | 7 | 6 | 9 | 9 | 1 |
| 911 | - | 971986 | 971612 | - | - | *PA14_11160* | hypothetical protein | 5 | 7 | 5 | 5 | 8 | 6 | 1 |
| 912 | - | 972093 | 972929 | + | - | *PA14_11170* | hypothetical protein | 78 | 80 | 70 | 78 | 79 | 68 | 1 |
| 913 | - | 973594 | 972926 | - | - | *PA14_11180* | transcriptional regulator | 22 | 25 | 22 | 23 | 19 | 21 | 1 |
| 914 | - | 973823 | 975310 | + | - | *PA14_11190* | aldehyde dehydrogenase | 8 | 12 | 8 | 14 | 12 | 10 | 1 |
| 915 | - | 975501 | 976991 | + | - | *PA14_11210* | amino acid permease | 6 | 8 | 7 | 8 | 8 | 7 | 1 |
| 916 | - | 976966 | 977544 | + | - | *PA14_11230* | hypothetical protein | 7 | 6 | 9 | 12 | 8 | 8 | 1 |
| 917 | - | 977705 | 978655 | + | - | *PA14_11240* | DNA-binding transcriptional activator FeaR | 29 | 28 | 25 | 34 | 24 | 33 | 1 |
| 918 | - | 978924 | 979808 | + | - | *PA14_11250* | hypothetical protein | 83 | 71 | 80 | 66 | 73 | 64 | 1 |
| 919 | - | 979801 | 980730 | + | - | *PA14_11260* | epimerase | 192 | 205 | 190 | 189 | 204 | 178 | 1 |
| 920 | - | 981869 | 981171 | - | *oprG* | *PA14_11270* | outer membrane protein OprG precursor | 1993 | 2754 | 2204 | 947 | 1496 | 1484 | 1 |
| 921 | - | 982535 | 982017 | - | - | *PA14_11280* | hypothetical protein | 29 | 30 | 32 | 36 | 32 | 32 | 1 |
| 922 | - | 983818 | 982553 | - | - | *PA14_11290* | permease | 29 | 24 | 25 | 27 | 25 | 30 | 1 |
| 923 | - | 984525 | 983821 | - | - | *PA14_11310* | ABC transporter ATP-binding protein | 17 | 13 | 17 | 15 | 14 | 14 | 1 |
| 924 | - | 985204 | 984614 | - | - | *PA14_11320* | hypothetical protein | 24 | 17 | 23 | 24 | 31 | 43 | 1 |

| | A | B | C | D | E | F | G | H | I | J | K | L | M | N |
|---|---|---|---|---|---|---|---|---|---|---|---|---|---|---|
| 925 | - | 985337 | 985681 | + | - | *PA14_11330* | hypothetical protein | 73 | 60 | 72 | 63 | 63 | 59 | 1 |
| 926 | - | 986587 | 985718 | - | - | *PA14_11340* | thioredoxin | 135 | 126 | 178 | 79 | 114 | 92 | 1 |
| 927 | - | 987321 | 986659 | - | - | *PA14_11350* | hypothetical protein | 34 | 29 | 37 | 30 | 28 | 28 | 1 |
| 928 | - | 987788 | 987318 | - | - | *PA14_11370* | lipoprotein | 21 | 21 | 23 | 20 | 21 | 21 | 1 |
| 929 | - | 987974 | 988438 | + | *nrdR* | *PA14_11380* | transcriptional regulator NrdR | 67 | 54 | 67 | 50 | 47 | 47 | 1 |
| 930 | - | 988435 | 989556 | + | *ribD* | *PA14_11400* | riboflavin-specific deaminase/reductase | 310 | 265 | 313 | 318 | 267 | 259 | 1 |
| 931 | - | 989984 | 990643 | + | *ribC* | *PA14_11410* | riboflavin synthase subunit alpha | 49 | 42 | 48 | 35 | 39 | 41 | 1 |
| 932 | - | 990670 | 991767 | + | *ribB* | *PA14_11420* | bifunctional 3,4-dihydroxy-2-butanone 4-phosphate synthase/GTP cyclohydrolase II-like protein | 55 | 56 | 55 | 40 | 56 | 44 | 1 |
| 933 | - | 991885 | 992361 | + | *ribH* | *PA14_11430* | 6,7-dimethyl-8-ribityllumazine synthase | 134 | 143 | 160 | 132 | 116 | 109 | 1 |
| 934 | - | 992358 | 992837 | + | *nusB* | *PA14_11450* | transcription antitermination protein NusB | 142 | 171 | 168 | 131 | 133 | 109 | 1 |
| 935 | - | 992856 | 993824 | + | *thiL* | *PA14_11460* | thiamine monophosphate kinase | 72 | 64 | 78 | 55 | 67 | 48 | 1 |
| 936 | - | 993817 | 994332 | + | *pgpA* | *PA14_11470* | phosphatidylglycerophosphatase A | 74 | 69 | 94 | 70 | 78 | 64 | 1 |
| 937 | - | 994341 | 995087 | + | - | *PA14_11480* | hypothetical protein | 93 | 79 | 119 | 105 | 98 | 98 | 1 |
| 938 | - | 995090 | 995734 | + | - | *PA14_11490* | hypothetical protein | 104 | 100 | 142 | 107 | 109 | 103 | 1 |
| 939 | - | 995865 | 996482 | + | *ribA* | *PA14_11510* | GTP cyclohydrolase II | 86 | 65 | 95 | 62 | 66 | 71 | 1 |
| 940 | - | 996479 | 996898 | + | - | *PA14_11520* | hypothetical protein | 56 | 54 | 63 | 41 | 52 | 44 | 1 |
| 941 | - | 996898 | 997695 | + | - | *PA14_11530* | hypothetical protein | 58 | 48 | 63 | 51 | 51 | 48 | 1 |
| 942 | - | 999702 | 997819 | - | *dxs* | *PA14_11550* | 1-deoxy-D-xylulose-5-phosphate synthase | 120 | 112 | 122 | 112 | 111 | 99 | 1 |
| 943 | - | 1000698 | 999811 | - | *ispA* | *PA14_11560* | geranyltranstransferase | 60 | 54 | 67 | 70 | 59 | 62 | 1 |
| 944 | - | 1000937 | 1000695 | - | *xseB* | *PA14_11570* | exodeoxyribonuclease VII small subunit | 34 | 26 | 40 | 30 | 29 | 28 | 1 |
| 945 | - | 1001095 | 1002264 | + | - | *PA14_11580* | hypothetical protein | 79 | 73 | 89 | 93 | 70 | 82 | 1 |
| 946 | - | 1003503 | 1002496 | - | - | *PA14_11590* | hypothetical protein | 24 | 24 | 24 | 27 | 21 | 28 | 1 |
| 947 | - | 1005307 | 1003505 | - | - | *PA14_11600* | ABC transporter | 47 | 45 | 49 | 52 | 42 | 50 | 1 |
| 948 | - | 1006051 | 1005317 | - | - | *PA14_11610* | ABC transporter permease | 6 | 5 | 8 | 8 | 7 | 6 | 1 |
| 949 | - | 1006974 | 1006048 | - | - | *PA14_11620* | ABC transporter | 7 | 7 | 9 | 9 | 6 | 8 | 1 |
| 950 | - | 1009679 | 1007379 | - | - | *PA14_11630* | two-component sensor | 31 | 36 | 36 | 38 | 38 | 35 | 1 |
| 951 | - | 1010699 | 1009698 | - | - | *PA14_11650* | hypothetical protein | 36 | 39 | 36 | 32 | 29 | 26 | 1 |
| 952 | - | 1011554 | 1010865 | - | *aqpZ* | *PA14_11660* | aquaporin Z | 9 | 9 | 8 | 8 | 7 | 7 | 1 |
| 953 | - | 1012170 | 1011901 | - | - | *PA14_11670* | hypothetical protein | 12 | 11 | 11 | 14 | 8 | 12 | 1 |
| 954 | - | 1012365 | 1013081 | + | - | *PA14_11680* | two-component regulator | 87 | 73 | 95 | 71 | 63 | 70 | 1 |
| 955 | - | 1013699 | 1013172 | - | *ppa* | *PA14_11690* | inorganic pyrophosphatase | 89 | 127 | 81 | 54 | 79 | 49 | 1 |
| 956 | - | 1014609 | 1013779 | - | - | *PA14_11700* | hypothetical protein | 26 | 24 | 34 | 22 | 22 | 21 | 1 |
| 957 | - | 1015277 | 1014612 | - | - | *PA14_11720* | hypothetical protein | 31 | 24 | 36 | 20 | 21 | 17 | 1 |
| 958 | - | 1015471 | 1017054 | + | - | *PA14_11730* | protein kinase | 24 | 22 | 24 | 23 | 22 | 22 | 1 |
| 959 | - | 1017840 | 1016947 | - | - | *PA14_11740* | hypothetical protein | 64 | 73 | 71 | 64 | 73 | 81 | 1 |
| 960 | - | 1018351 | 1017890 | - | - | *PA14_11750* | acetyltransferase | 125 | 119 | 111 | 92 | 111 | 120 | 1 |
| 961 | - | 1019336 | 1018515 | - | - | *PA14_11760* | ethanolamine ammonia-lyase small subunit | 51 | 58 | 58 | 67 | 47 | 63 | 1 |
| 962 | - | 1020741 | 1019347 | - | *eutB* | *PA14_11770* | ethanolamine ammonia-lyase large subunit | 54 | 69 | 74 | 73 | 58 | 73 | 1 |
| 963 | - | 1022248 | 1020800 | - | - | *PA14_11790* | amino acid transporter | 22 | 29 | 33 | 29 | 23 | 24 | 1 |
| 964 | - | 1023978 | 1022458 | - | - | *PA14_11810* | aldehyde dehydrogenase | 157 | 285 | 203 | 174 | 177 | 162 | 1 |
| 965 | - | 1024317 | 1026248 | + | - | *PA14_11830* | transcriptional regulator | 101 | 108 | 94 | 103 | 91 | 90 | 1 |
| 966 | - | 1026406 | 1027761 | + | *mpl* | *PA14_11845* | UDP-N-acetylmuramate:L-alanyl-gamma-D-glutamyl-meso-diaminopimelate ligase | 117 | 111 | 114 | 114 | 117 | 114 | 1 |
| 967 | - | 1027770 | 1028399 | + | - | *PA14_11860* | aromatic acid decarboxylase | 160 | 170 | 166 | 152 | 163 | 148 | 1 |
| 968 | - | 1028401 | 1028676 | + | - | *PA14_11880* | hypothetical protein | 157 | 176 | 188 | 165 | 166 | 185 | 1 |
| 969 | - | 1029337 | 1028696 | - | - | *PA14_11890* | hypothetical protein | 264 | 309 | 366 | 298 | 364 | 386 | 1 |
| 970 | - | 1031123 | 1029384 | - | - | *PA14_11900* | hypothetical protein | 39 | 35 | 39 | 36 | 31 | 32 | 1 |
| 971 | - | 1031769 | 1031314 | - | - | *PA14_11910* | hypothetical protein | 78 | 97 | 73 | 82 | 105 | 89 | 1 |
| 972 | - | 1031915 | 1032325 | + | - | *PA14_11920* | hypothetical protein | 43 | 38 | 48 | 39 | 35 | 29 | 1 |
| 973 | - | 1032325 | 1033035 | + | - | *PA14_11930* | hypothetical protein | 80 | 68 | 88 | 78 | 70 | 61 | 1 |
| 974 | - | 1033055 | 1033648 | + | - | *PA14_11940* | hypothetical protein | 248 | 239 | 278 | 289 | 264 | 257 | 1 |
| 975 | - | 1034972 | 1033659 | - | - | *PA14_11960* | hypothetical protein | 79 | 75 | 79 | 77 | 75 | 72 | 1 |
| 976 | - | 1035688 | 1034969 | - | - | *PA14_11970* | 3-methyladenine DNA glycosylase | 67 | 45 | 63 | 58 | 51 | 51 | 1 |
| 977 | - | 1035752 | 1036609 | + | - | *PA14_11980* | hypothetical protein | 23 | 21 | 24 | 27 | 22 | 23 | 1 |
| 978 | - | 1037840 | 1036617 | - | - | *PA14_11990* | hydrolase | 25 | 26 | 31 | 31 | 25 | 34 | 1 |
| 979 | - | 1038138 | 1039403 | + | *proA* | *PA14_12010* | gamma-glutamyl phosphate reductase | 117 | 114 | 129 | 120 | 97 | 90 | 1 |
| 980 | - | 1039403 | 1040047 | + | *nadD* | *PA14_12020* | nicotinic acid mononucleotide adenylyltransferase | 229 | 225 | 285 | 248 | 221 | 189 | 1 |
| 981 | - | 1040070 | 1040426 | + | - | *PA14_12030* | hypothetical protein | 275 | 365 | 360 | 284 | 286 | 276 | 1 |
| 982 | - | 1040434 | 1040901 | + | - | *PA14_12050* | rRNA large subunit methyltransferase | 206 | 188 | 253 | 181 | 181 | 165 | 1 |
| 983 | - | 1040914 | 1042854 | + | *pbpA* | *PA14_12060* | penicillin-binding protein 2 | 73 | 64 | 84 | 68 | 72 | 60 | 1 |
| 984 | - | 1042886 | 1043989 | + | *rodA* | *PA14_12070* | rod shape-determining protein | 74 | 65 | 82 | 81 | 79 | 67 | 1 |
| 985 | - | 1043998 | 1045020 | + | *sltB1* | *PA14_12080* | soluble lytic transglycosylase B | 179 | 126 | 174 | 185 | 182 | 165 | 1 |
| 986 | - | 1045017 | 1046042 | + | - | *PA14_12090* | RlpA family lipoprotein | 301 | 277 | 327 | 319 | 279 | 272 | 1 |
| 987 | - | 1046108 | 1047268 | + | *dacC* | *PA14_12100* | D-ala-D-ala-carboxypeptidase | 368 | 402 | 355 | 361 | 380 | 332 | 1 |
| 988 | - | 1047339 | 1047620 | + | - | *PA14_12110* | hypothetical protein | 324 | 354 | 336 | 285 | 306 | 282 | 1 |
| 989 | - | 1047620 | 1048273 | + | *lipB* | *PA14_12120* | lipoate-protein ligase B | 260 | 229 | 253 | 237 | 225 | 197 | 1 |
| 990 | - | 1048270 | 1049253 | + | *lis* | *PA14_12130* | lipoyl synthase | 523 | 453 | 542 | 508 | 544 | 525 | 1 |
| 991 | - | 1050291 | 1049398 | - | - | *PA14_12140* | transcriptional regulator | 40 | 35 | 42 | 40 | 39 | 44 | 1 |
| 992 | - | 1050399 | 1051289 | + | - | *PA14_12150* | epoxide hydrolase | 9 | 9 | 9 | 11 | 10 | 9 | 1 |
| 993 | - | 1052838 | 1051492 | - | - | *PA14_12160* | murein transglycosylase | 57 | 62 | 63 | 52 | 65 | 54 | 1 |
| 994 | - | 1053067 | 1053285 | + | - | *PA14_12170* | hypothetical protein | 9 | 9 | 13 | 10 | 10 | 10 | 1 |
| 995 | - | 1053553 | 1053401 | - | - | *PA14_12180* | hypothetical protein | 7 | 7 | 9 | 7 | 9 | 7 | 1 |
| 996 | - | 1054682 | 1053645 | - | *holA* | *PA14_12200* | DNA polymerase III subunit delta | 65 | 68 | 53 | 47 | 59 | 52 | 1 |
| 997 | - | 1055343 | 1054720 | - | - | *PA14_12210* | hypothetical protein | 137 | 148 | 112 | 112 | 147 | 108 | 1 |
| 998 | - | 1058031 | 1055410 | - | *leuS* | *PA14_12230* | leucyl-tRNA synthetase | 192 | 205 | 182 | 175 | 185 | 164 | 1 |
| 999 | - | 1058202 | 1058633 | + | - | *PA14_12260* | hypothetical protein | 434 | 361 | 586 | 420 | 394 | 424 | 1 |
| 1000 | - | 1058706 | 1059467 | + | - | *PA14_12270* | hypothetical protein | 62 | 54 | 61 | 57 | 56 | 54 | 1 |
| 1001 | - | 1061052 | 1059517 | - | *lnt* | *PA14_12280* | apolipoprotein N-acyltransferase | 41 | 44 | 33 | 36 | 36 | 29 | 1 |
| 1002 | - | 1062077 | 1061238 | - | - | *PA14_12300* | hypothetical protein | 147 | 150 | 142 | 114 | 126 | 125 | 1 |
| 1003 | - | 1062556 | 1062074 | - | - | *PA14_12310* | metalloprotease | 165 | 172 | 191 | 171 | 218 | 186 | 1 |
| 1004 | - | 1063568 | 1062546 | - | - | *PA14_12330* | hypothetical protein | 299 | 308 | 308 | 248 | 280 | 290 | 1 |
| 1005 | - | 1065077 | 1063737 | - | - | *PA14_12350* | (dimethylallyl)adenosine tRNA methylthiotransferase | 62 | 61 | 70 | 67 | 57 | 51 | 1 |
| 1006 | - | 1065206 | 1065538 | + | - | *PA14_12360* | hypothetical protein | 30 | 27 | 38 | 28 | 28 | 31 | 1 |
| 1007 | - | 1066102 | 1065554 | - | - | *PA14_12370* | hypothetical protein | 244 | 243 | 309 | 213 | 283 | 249 | 1 |
| 1008 | - | 1067594 | 1066311 | - | *hemL* | *PA14_12390* | glutamate-1-semialdehyde aminotransferase | 123 | 131 | 106 | 95 | 115 | 89 | 1 |

| | A | B | C | D | E | F | G | H | I | J | K | L | M | N |
|---|---|---|---|---|---|---|---|---|---|---|---|---|---|---|
| 1009 | - | 1068261 | 1067632 | - | *thiE* | *PA14_12400* | thiamine-phosphate pyrophosphorylase | 49 | 50 | 41 | 44 | 38 | 36 | 1 |
| 1010 | - | 1069069 | 1068272 | - | *thiD* | *PA14_12410* | phosphomethylpyrimidine kinase | 96 | 66 | 83 | 71 | 70 | 67 | 1 |
| 1011 | - | 1071822 | 1072469 | + | - | *PA14_12440* | AcrR family transcriptional regulator | 170 | 136 | 145 | 145 | 185 | 225 | 1 |
| 1012 | - | 1072466 | 1074115 | + | - | *PA14_12450* | acyl-CoA dehydrogenase | 126 | 126 | 112 | 120 | 178 | 194 | 1 |
| 1013 | - | 1074127 | 1074558 | + | - | *PA14_12470* | hypothetical protein | 71 | 61 | 79 | 64 | 81 | 91 | 1 |
| 1014 | - | 1074613 | 1076112 | + | - | *PA14_12490* | AMP nucleosidase | 75 | 81 | 65 | 62 | 77 | 70 | 1 |
| 1015 | - | 1077872 | 1076787 | - | - | *PA14_12530* | hypothetical protein | 56 | 55 | 52 | 43 | 52 | 47 | 1 |
| 1016 | - | 1078011 | 1078580 | + | - | *PA14_12540* | pseudouridine synthase | 29 | 30 | 27 | 25 | 23 | 18 | 1 |
| 1017 | - | 1079006 | 1078614 | - | - | *PA14_12550* | hypothetical protein | 31 | 23 | 27 | 20 | 24 | 20 | 1 |
| 1018 | - | 1079316 | 1079107 | - | - | *PA14_12560* | hypothetical protein | 160 | 78 | 199 | 126 | 131 | 162 | 0.243629287 |
| 1019 | - | 1080027 | 1079518 | - | - | *PA14_12570* | transcription regulator AsnC | 20 | 34 | 26 | 23 | 22 | 22 | 1 |
| 1020 | - | 1081194 | 1080409 | - | - | *PA14_12590* | hypothetical protein | 63 | 74 | 69 | 112 | 51 | 49 | 1 |
| 1021 | - | 1082047 | 1081151 | - | - | *PA14_12610* | transporter | 35 | 41 | 35 | 46 | 34 | 30 | 1 |
| 1022 | - | 1082552 | 1082136 | - | - | *PA14_12620* | hypothetical protein | 40 | 43 | 33 | 26 | 40 | 31 | 1 |
| 1023 | - | 1082719 | 1085211 | + | - | *PA14_12630* | ATP-dependent helicase | 66 | 67 | 66 | 60 | 59 | 49 | 1 |
| 1024 | - | 1085629 | 1085222 | - | - | *PA14_12640* | hypothetical protein | 30 | 38 | 33 | 36 | 37 | 37 | 1 |
| 1025 | - | 1086387 | 1085653 | - | - | *PA14_12650* | hypothetical protein | 11 | 9 | 13 | 14 | 9 | 8 | 1 |
| 1026 | - | 1087616 | 1086477 | - | - | *PA14_12670* | hypothetical protein | 39 | 30 | 40 | 39 | 34 | 39 | 1 |
| 1027 | - | 1087678 | 1088514 | + | - | *PA14_12680* | short chain dehydrogenase | 119 | 117 | 144 | 141 | 140 | 149 | 1 |
| 1028 | - | 1088955 | 1088521 | - | - | *PA14_12690* | hypothetical protein | 51 | 60 | 75 | 62 | 50 | 52 | 1 |
| 1029 | - | 1089678 | 1089022 | - | - | *PA14_12700* | hypothetical protein | 23 | 21 | 26 | 27 | 19 | 21 | 1 |
| 1030 | - | 1090856 | 1089762 | - | - | *PA14_12710* | hypothetical protein | 15 | 16 | 15 | 15 | 15 | 12 | 1 |
| 1031 | - | 1091595 | 1091014 | - | - | *PA14_12730* | hypothetical protein | 10 | 6 | 13 | 7 | 5 | 8 | 0.499266146 |
| 1032 | - | 1091836 | 1092411 | + | - | *PA14_12740* | hypothetical protein | 284 | 240 | 461 | 155 | 232 | 158 | 1 |
| 1033 | - | 1092477 | 1092980 | + | - | *PA14_12750* | hypothetical protein | 79 | 66 | 108 | 52 | 71 | 57 | 1 |
| 1034 | - | 1093058 | 1094407 | + | - | *PA14_12760* | ATP-dependent RNA helicase | 61 | 60 | 63 | 58 | 57 | 49 | 1 |
| 1035 | - | 1095605 | 1094430 | - | - | *PA14_12770* | hypothetical protein | 78 | 59 | 74 | 62 | 53 | 53 | 1 |
| 1036 | - | 1096593 | 1095964 | - | - | *PA14_12780* | two-component response regulator | 94 | 73 | 106 | 71 | 82 | 70 | 1 |
| 1037 | - | 1096875 | 1098053 | + | - | *PA14_12810* | two-component response regulator | 22 | 25 | 23 | 22 | 21 | 23 | 1 |
| 1038 | - | 1098155 | 1101793 | + | - | *PA14_12820* | two-component sensor | 10 | 10 | 11 | 10 | 10 | 10 | 1 |
| 1039 | - | 1102008 | 1102598 | + | - | *PA14_12840* | acetyltransferase | 367 | 345 | 443 | 364 | 385 | 440 | 1 |
| 1040 | - | 1102625 | 1103203 | + | - | *PA14_12850* | acetyltransferase | 116 | 103 | 112 | 91 | 98 | 99 | 1 |
| 1041 | - | 1103206 | 1104801 | + | - | *PA14_12860* | hypothetical protein | 93 | 79 | 87 | 75 | 85 | 88 | 1 |
| 1042 | - | 1104835 | 1105704 | + | *tesB* | *PA14_12870* | acyl-CoA thioesterase | 53 | 44 | 50 | 41 | 49 | 46 | 1 |
| 1043 | - | 1105701 | 1106297 | + | - | *PA14_12890* | hypothetical protein | 55 | 68 | 63 | 56 | 64 | 62 | 1 |
| 1044 | - | 1106702 | 1106421 | - | - | *PA14_12900* | DNA binding protein | 205 | 517 | 218 | 177 | 212 | 193 | 0.11934908 |
| 1045 | - | 1107072 | 1107251 | + | - | *PA14_12910* | hypothetical protein | 15 | 11 | 6 | 6 | 10 | 9 | 1 |
| 1046 | - | 1107458 | 1108471 | + | - | *PA14_12920* | taurine ABC transporter periplasmic protein | 10 | 12 | 11 | 11 | 8 | 10 | 1 |
| 1047 | - | 1108535 | 1109326 | + | - | *PA14_12940* | taurine ABC transporter ATP-binding protein | 6 | 7 | 7 | 7 | 6 | 7 | 1 |
| 1048 | - | 1109316 | 1110134 | + | - | *PA14_12960* | taurine ABC transporter permease | 6 | 5 | 5 | 6 | 4 | 4 | 1 |
| 1049 | - | 1110241 | 1111074 | + | *tauD* | *PA14_12970* | taurine dioxygenase | 12 | 10 | 11 | 12 | 9 | 10 | 1 |
| 1050 | - | 1111237 | 1113273 | + | - | *PA14_12980* | hypothetical protein | 68 | 88 | 75 | 110 | 84 | 74 | 1 |
| 1051 | - | 1115342 | 1113381 | - | - | *PA14_12990* | choline transporter | 85 | 59 | 72 | 70 | 72 | 78 | 1 |
| 1052 | - | 1116506 | 1115532 | - | - | *PA14_13000* | transcriptional regulator | 11 | 15 | 10 | 10 | 12 | 13 | 1 |
| 1053 | - | 1117461 | 1116682 | - | - | *PA14_13010* | hypothetical protein | 15 | 50 | 15 | 12 | 23 | 14 | 0.000919853 |
| 1054 | - | 1117899 | 1119365 | + | *cioA* | *PA14_13030* | CioA, cyanide insensitive terminal oxidase | 227 | 165 | 136 | 273 | 277 | 227 | 1 |
| 1055 | - | 1119369 | 1120376 | + | *cioB* | *PA14_13040* | CioB, cyanide insensitive terminal oxidase | 310 | 208 | 197 | 321 | 311 | 295 | 1 |
| 1056 | - | 1120390 | 1120560 | + | - | *PA14_13050* | hypothetical protein | 102 | 81 | 65 | 113 | 121 | 108 | 1 |
| 1057 | - | 1121623 | 1120835 | - | - | *PA14_13060* | transcriptional regulator | 18 | 22 | 17 | 17 | 19 | 18 | 1 |
| 1058 | - | 1121725 | 1122906 | + | - | *PA14_13070* | MFS family transporter | 12 | 11 | 14 | 16 | 11 | 10 | 1 |
| 1059 | - | 1124138 | 1122963 | - | - | *PA14_13090* | acyl-CoA thiolase | 35 | 39 | 31 | 33 | 31 | 29 | 1 |
| 1060 | - | 1124320 | 1126002 | + | - | *PA14_13110* | long-chain-fatty-acid--CoA ligase | 41 | 51 | 46 | 66 | 46 | 62 | 1 |
| 1061 | - | 1126214 | 1128139 | + | - | *PA14_13130* | hypothetical protein | 70 | 91 | 71 | 129 | 79 | 116 | 1 |
| 1062 | - | 1128200 | 1129567 | + | - | *PA14_13140* | hypothetical protein | 61 | 92 | 62 | 122 | 78 | 109 | 1 |
| 1063 | - | 1129901 | 1132621 | + | - | *PA14_13150* | transcriptional regulator | 132 | 142 | 136 | 164 | 129 | 154 | 1 |
| 1064 | - | 1135031 | 1132653 | - | - | *PA14_13170* | metal transporting P-type ATPase | 42 | 61 | 29 | 67 | 69 | 55 | 1 |
| 1065 | - | 1135535 | 1136926 | + | - | *PA14_13190* | hypothetical protein | 206 | 211 | 187 | 185 | 207 | 264 | 1 |
| 1066 | - | 1137043 | 1137819 | + | - | *PA14_13200* | hypothetical protein | 33 | 29 | 32 | 35 | 34 | 52 | 1 |
| 1067 | - | 1138011 | 1137829 | - | - | *PA14_13210* | hypothetical protein | 73 | 43 | 53 | 98 | 83 | 166 | 0.43335915 |
| 1068 | - | 1138602 | 1138174 | - | - | *PA14_13220* | protein-tyrosine-phosphatase | 26 | 18 | 23 | 16 | 17 | 19 | 1 |
| 1069 | - | 1138820 | 1139302 | + | *moaC* | *PA14_13230* | molybdenum cofactor biosynthesis protein MoaC | 59 | 61 | 68 | 91 | 58 | 233 | 1 |
| 1070 | - | 1139299 | 1139550 | + | *moaD* | *PA14_13240* | molybdopterin converting factor, small subunit | 99 | 92 | 105 | 136 | 139 | 350 | 1 |
| 1071 | - | 1139555 | 1140007 | + | *moaE* | *PA14_13250* | molybdopterin converting factor, large subunit | 53 | 51 | 47 | 60 | 64 | 141 | 1 |
| 1072 | - | 1140109 | 1140666 | + | *moaB1* | *PA14_13260* | MoaB1 | 7 | 25 | 11 | 125 | 98 | 417 | 1.45127E-05 |
| 1073 | - | 1140663 | 1141886 | + | *moeA1* | *PA14_13280* | molybdenum cofactor biosynthetic protein A1 | 7 | 27 | 9 | 30 | 55 | 185 | 4.73358E-05 |
| 1074 | - | 1142001 | 1142996 | + | - | *PA14_13290* | protease | 41 | 63 | 51 | 78 | 91 | 103 | 1 |
| 1075 | - | 1143008 | 1143898 | + | - | *PA14_13300* | hypothetical protein | 49 | 102 | 71 | 96 | 139 | 101 | 0.391278558 |
| 1076 | - | 1143892 | 1144407 | + | - | *PA14_13320* | hypothetical protein | 44 | 88 | 71 | 83 | 138 | 86 | 0.328493033 |
| 1077 | - | 1144659 | 1146221 | + | - | *PA14_13330* | hypothetical protein | 9 | 10 | 11 | 13 | 11 | 11 | 1 |
| 1078 | - | 1146258 | 1148600 | + | - | *PA14_13340* | extracellular nuclease | 12 | 7 | 7 | 9 | 7 | 12 | 0.711763148 |
| 1079 | - | 1149641 | 1148922 | - | - | *PA14_13350* | hypothetical protein | 142 | 217 | 164 | 165 | 250 | 149 | 1 |
| 1080 | - | 1150423 | 1149638 | - | - | *PA14_13360* | hypothetical protein | 140 | 165 | 148 | 143 | 193 | 115 | 1 |
| 1081 | - | 1150803 | 1150420 | - | - | *PA14_13370* | hypothetical protein | 159 | 180 | 174 | 190 | 219 | 129 | 1 |
| 1082 | - | 1151323 | 1150796 | - | - | *PA14_13380* | hypothetical protein | 110 | 130 | 125 | 121 | 137 | 89 | 1 |
| 1083 | - | 1151715 | 1151320 | - | - | *PA14_13390* | hypothetical protein | 112 | 133 | 137 | 154 | 112 | 99 | 1 |
| 1084 | - | 1153683 | 1152100 | - | *prfC* | *PA14_13410* | peptide chain release factor 3 | 47 | 51 | 63 | 52 | 48 | 37 | 1 |
| 1085 | - | 1153904 | 1154365 | + | - | *PA14_13420* | hypothetical protein | 40 | 46 | 42 | 35 | 47 | 35 | 1 |
| 1086 | - | 1156819 | 1154465 | - | *fecA* | *PA14_13430* | Fe(III) dicitrate transport protein FecA | 16 | 17 | 17 | 19 | 20 | 14 | 1 |
| 1087 | - | 1157867 | 1156914 | - | - | *PA14_13450* | transmembrane sensor | 18 | 29 | 25 | 67 | 43 | 20 | 0.934705131 |
| 1088 | - | 1158373 | 1157864 | - | - | *PA14_13460* | RNA polymerase sigma factor | 7 | 31 | 10 | 80 | 70 | 12 | 3.18925E-09 |
| 1089 | - | 1159355 | 1158474 | - | - | *PA14_13470* | AraC family transcriptional regulator | 21 | 19 | 21 | 23 | 19 | 19 | 1 |
| 1090 | - | 1159481 | 1160383 | + | - | *PA14_13490* | hypothetical protein | 15 | 16 | 14 | 16 | 14 | 12 | 1 |
| 1091 | - | 1161378 | 1160401 | - | - | *PA14_13500* | 2-hydroxyacid dehydrogenase | 22 | 25 | 21 | 20 | 24 | 18 | 1 |
| 1092 | - | 1162426 | 1161473 | - | - | *PA14_13510* | LysR family transcriptional regulator | 31 | 31 | 32 | 28 | 31 | 31 | 1 |

| | A | B | C | D | E | F | G | H | I | J | K | L | M | N |
|---|---|---|---|---|---|---|---|---|---|---|---|---|---|---|
| 1093 | - | 1162672 | 1164162 | + | - | PA14_13520 | outer membrane protein | 20 | 13 | 17 | 15 | 13 | 15 | 1 |
| 1094 | - | 1164159 | 1166354 | + | - | PA14_13530 | outer membrane protein | 12 | 10 | 10 | 10 | 10 | 10 | 1 |
| 1095 | - | 1166581 | 1167489 | + | - | PA14_13560 | hypothetical protein | 11 | 8 | 10 | 8 | 13 | 9 | 1 |
| 1096 | - | 1168780 | 1167617 | - | - | PA14_13580 | ABC transporter ATP-binding protein | 64 | 87 | 73 | 71 | 98 | 81 | 1 |
| 1097 | - | 1169455 | 1168793 | - | - | PA14_13590 | ABC transporter permease | 63 | 65 | 67 | 63 | 84 | 84 | 1 |
| 1098 | - | 1170390 | 1169455 | - | - | PA14_13600 | ABC transporter substrate-binding protein | 50 | 51 | 51 | 53 | 66 | 57 | 1 |
| 1099 | - | 1171112 | 1170390 | - | - | PA14_13610 | ABC transporter permease | 63 | 48 | 61 | 51 | 64 | 60 | 1 |
| 1100 | - | 1171554 | 1172828 | + | nhaP | PA14_13620 | Na+/H+ antiporter NhaP | 41 | 38 | 38 | 40 | 36 | 35 | 1 |
| 1101 | - | 1173132 | 1172887 | - | - | PA14_13630 | hypothetical protein | 43 | 51 | 54 | 52 | 55 | 53 | 1 |
| 1102 | - | 1174062 | 1173163 | - | - | PA14_13650 | hypothetical protein | 31 | 30 | 26 | 24 | 26 | 25 | 1 |
| 1103 | - | 1174823 | 1174167 | - | - | PA14_13660 | hypothetical protein | 3 | 6 | 6 | 4 | 4 | 4 | 1 |
| 1104 | - | 1175304 | 1174918 | - | - | PA14_13670 | hypothetical protein | 9 | 9 | 11 | 9 | 9 | 8 | 1 |
| 1105 | - | 1176155 | 1175325 | - | - | PA14_13680 | short chain dehydrogenase | 29 | 23 | 30 | 26 | 28 | 29 | 1 |
| 1106 | - | 1176992 | 1176243 | - | - | PA14_13690 | methyltransferase | 35 | 43 | 31 | 38 | 43 | 39 | 1 |
| 1107 | - | 1177472 | 1177008 | - | - | PA14_13710 | hypothetical protein | 33 | 32 | 31 | 32 | 38 | 37 | 1 |
| 1108 | - | 1178033 | 1177638 | - | - | PA14_13720 | hypothetical protein | 128 | 135 | 125 | 168 | 223 | 172 | 1 |
| 1109 | - | 1178777 | 1178118 | - | narL | PA14_13730 | transcriptional regulator NarL | 235 | 203 | 299 | 138 | 148 | 249 | 1 |
| 1110 | - | 1180642 | 1178774 | - | narX | PA14_13740 | two-component sensor NarX | 59 | 50 | 67 | 49 | 50 | 60 | 1 |
| 1111 | - | 1180817 | 1182112 | + | narK1 | PA14_13750 | nitrite extrusion protein 1 | 17 | 59 | 18 | 80 | 79 | 316 | 0.000376823 |
| 1112 | - | 1182125 | 1183531 | + | narK2 | PA14_13770 | nitrite extrusion protein 2 | 10 | 49 | 13 | 17 | 67 | 102 | 1.71783E-09 |
| 1113 | - | 1183607 | 1187392 | + | narG | PA14_13780 | respiratory nitrate reductase alpha subun | 21 | 60 | 23 | 19 | 82 | 50 | 0.066717397 |
| 1114 | - | 1187404 | 1188945 | + | narH | PA14_13800 | respiratory nitrate reductase beta subuni | 25 | 76 | 27 | 18 | 98 | 35 | 0.016650042 |
| 1115 | - | 1188951 | 1189691 | + | narJ | PA14_13810 | respiratory nitrate reductase delta chain | 13 | 51 | 12 | 8 | 51 | 15 | 2.63546E-06 |
| 1116 | - | 1189694 | 1190377 | + | narI | PA14_13830 | respiratory nitrate reductase gamma chain | 15 | 49 | 12 | 8 | 51 | 15 | 0.000417016 |
| 1117 | - | 1190432 | 1191250 | + | - | PA14_13840 | peptidyl-prolyl cis-trans isomerase, PpiC-type | 19 | 54 | 20 | 13 | 54 | 22 | 0.006997432 |
| 1118 | - | 1191307 | 1192296 | + | moaA | PA14_13850 | molybdenum cofactor biosynthesis protein A | 20 | 34 | 17 | 15 | 35 | 17 | 0.65533104 |
| 1119 | - | 1192482 | 1192303 | - | - | PA14_13860 | hypothetical protein | 14 | 19 | 13 | 14 | 19 | 13 | 1 |
| 1120 | - | 1193000 | 1192617 | - | - | PA14_13870 | hypothetical protein | 13 | 13 | 16 | 19 | 10 | 12 | 1 |
| 1121 | - | 1193138 | 1193043 | - | - | PA14_13880 | hypothetical protein | 7 | 8 | 7 | 8 | 6 | 4 | 1 |
| 1122 | - | 1194201 | 1193191 | - | - | PA14_13890 | integrase | 8 | 8 | 6 | 6 | 7 | 6 | 1 |
| 1123 | - | 1195465 | 1195623 | + | - | PA14_13920 | hypothetical protein | 5 | 5 | 7 | 5 | 4 | 7 | 1 |
| 1124 | - | 1196245 | 1197924 | + | - | PA14_13940 | S-type pyocin protein | 70 | 90 | 66 | 73 | 91 | 74 | 1 |
| 1125 | - | 1197924 | 1198262 | + | - | PA14_13950 | hypothetical protein | 112 | 133 | 98 | 74 | 130 | 100 | 1 |
| 1126 | - | 1198426 | 1198334 | - | - | PA14_13960 | hypothetical protein | 96 | 90 | 86 | 75 | 108 | 80 | 1 |
| 1127 | - | 1198694 | 1198431 | - | - | PA14_13970 | hypothetical protein | 43 | 36 | 38 | 43 | 46 | 43 | 1 |
| 1128 | - | 1199242 | 1200033 | + | - | PA14_13990 | amino acid ABC transporter | 35 | 43 | 30 | 90 | 40 | 42 | 1 |
| 1129 | - | 1200689 | 1200057 | - | - | PA14_14000 | hypothetical protein | 27 | 29 | 27 | 59 | 25 | 26 | 1 |
| 1130 | - | 1201879 | 1200752 | - | - | PA14_14010 | amino acid oxidase | 19 | 23 | 18 | 43 | 20 | 18 | 1 |
| 1131 | - | 1202849 | 1201902 | - | - | PA14_14020 | hypothetical protein | 26 | 29 | 25 | 48 | 26 | 25 | 1 |
| 1132 | - | 1204533 | 1203010 | - | rhl | PA14_14040 | ATP-dependent RNA helicase RhlB | 103 | 97 | 94 | 95 | 88 | 84 | 1 |
| 1133 | - | 1204817 | 1206715 | + | - | PA14_14060 | AMP-binding protein | 51 | 69 | 53 | 64 | 62 | 57 | 1 |
| 1134 | - | 1207378 | 1206731 | - | - | PA14_14080 | carboxylesterase | 62 | 74 | 64 | 69 | 74 | 62 | 1 |
| 1135 | - | 1207659 | 1208684 | + | - | PA14_14100 | amino acid-binding protein | 178 | 147 | 211 | 248 | 167 | 220 | 1 |
| 1136 | - | 1208798 | 1209514 | + | pcs | PA14_14110 | phosphatidylserine synthase | 108 | 80 | 107 | 75 | 88 | 92 | 1 |
| 1137 | - | 1209585 | 1210067 | + | - | PA14_14130 | hypothetical protein | 68 | 55 | 66 | 66 | 67 | 69 | 1 |
| 1138 | - | 1210134 | 1210835 | + | - | PA14_14140 | hypothetical protein | 50 | 46 | 50 | 52 | 46 | 47 | 1 |
| 1139 | - | 1210828 | 1211142 | + | - | PA14_14150 | GIY-YIG nuclease superfamily protein | 64 | 56 | 72 | 72 | 63 | 63 | 1 |
| 1140 | - | 1211171 | 1211860 | + | - | PA14_14160 | acetyltransferase | 53 | 51 | 49 | 60 | 60 | 59 | 1 |
| 1141 | - | 1211981 | 1212916 | + | - | PA14_14170 | hypothetical protein | 73 | 75 | 75 | 64 | 73 | 67 | 1 |
| 1142 | - | 1212920 | 1213672 | + | - | PA14_14200 | hypothetical protein | 54 | 54 | 53 | 55 | 59 | 57 | 1 |
| 1143 | - | 1213761 | 1214672 | + | - | PA14_14210 | hypothetical protein | 33 | 28 | 26 | 30 | 34 | 25 | 1 |
| 1144 | - | 1215719 | 1214715 | - | - | PA14_14220 | nucleoid-associated protein NdpA | 83 | 78 | 81 | 86 | 85 | 86 | 1 |
| 1145 | - | 1217167 | 1215812 | - | - | PA14_14230 | hypothetical protein | 58 | 60 | 52 | 50 | 61 | 58 | 1 |
| 1146 | - | 1217711 | 1217241 | - | - | PA14_14250 | hypothetical protein | 36 | 34 | 38 | 34 | 32 | 38 | 1 |
| 1147 | - | 1217815 | 1218357 | + | - | PA14_14270 | isochorismatase family hydrolase | 380 | 340 | 403 | 355 | 352 | 459 | 1 |
| 1148 | - | 1219243 | 1218347 | - | - | PA14_14280 | LysR family transcriptional regulator | 80 | 77 | 87 | 88 | 84 | 95 | 1 |
| 1149 | - | 1219367 | 1219978 | + | - | PA14_14290 | hypothetical protein | 28 | 30 | 31 | 34 | 27 | 31 | 1 |
| 1150 | - | 1221262 | 1220135 | - | - | PA14_14300 | zinc-binding oxidoreductase | 11 | 9 | 10 | 11 | 10 | 10 | 1 |
| 1151 | - | 1221495 | 1221193 | - | - | PA14_14310 | transcriptional regulator | 14 | 14 | 15 | 15 | 10 | 10 | 1 |
| 1152 | - | 1221589 | 1222182 | + | - | PA14_14320 | hypothetical protein | 12 | 4 | 6 | 6 | 4 | 13 | 0.001057312 |
| 1153 | - | 1222824 | 1222474 | - | - | PA14_14330 | chaperone | 13 | 17 | 14 | 13 | 11 | 14 | 1 |
| 1154 | - | 1224022 | 1223012 | - | - | PA14_14340 | SAM-dependent methyltransferase | 22 | 20 | 25 | 19 | 21 | 27 | 1 |
| 1155 | - | 1225946 | 1224114 | - | - | PA14_14360 | sodium:sulfate symporter | 98 | 84 | 129 | 52 | 62 | 109 | 1 |
| 1156 | - | 1226859 | 1226065 | - | - | PA14_14370 | ABC-transporter ATP-binding component | 70 | 82 | 54 | 58 | 68 | 56 | 1 |
| 1157 | - | 1227752 | 1226862 | - | - | PA14_14380 | ABC transporter permease | 24 | 30 | 20 | 21 | 25 | 17 | 1 |
| 1158 | - | 1228813 | 1227836 | - | - | PA14_14390 | ABC-type transport protein, periplasmic c | 73 | 122 | 61 | 62 | 86 | 61 | 1 |
| 1159 | - | 1228936 | 1229070 | + | - | PA14_14400 | hypothetical protein | 5 | 2 | 4 | 4 | 3 | 3 | 0.56521933 |
| 1160 | - | 1229163 | 1229642 | + | - | PA14_14420 | hypothetical protein | 14 | 17 | 11 | 12 | 15 | 9 | 1 |
| 1161 | - | 1231207 | 1231584 | + | - | PA14_14430 | hypothetical protein | 36 | 47 | 39 | 43 | 37 | 31 | 1 |
| 1162 | - | 1234433 | 1231581 | - | valS | PA14_14440 | valyl-tRNA synthetase | 76 | 88 | 80 | 79 | 85 | 65 | 1 |
| 1163 | - | 1234922 | 1234554 | - | - | PA14_14450 | hypothetical protein | 41 | 42 | 42 | 37 | 50 | 43 | 1 |
| 1164 | - | 1235362 | 1234934 | - | holC | PA14_14460 | DNA polymerase III subunit chi | 88 | 92 | 76 | 73 | 71 | 68 | 1 |
| 1165 | - | 1236846 | 1235359 | - | pepA | PA14_14470 | leucyl aminopeptidase | 118 | 129 | 103 | 97 | 106 | 93 | 1 |
| 1166 | - | 1237183 | 1237995 | + | - | PA14_14480 | transcriptional regulator | 57 | 44 | 56 | 60 | 48 | 50 | 1 |
| 1167 | - | 1238079 | 1239002 | + | - | PA14_14490 | hydrolase | 29 | 22 | 22 | 27 | 20 | 24 | 1 |
| 1168 | - | 1239195 | 1240313 | + | - | PA14_14500 | hypothetical protein | 71 | 43 | 72 | 60 | 55 | 59 | 1 |
| 1169 | - | 1240306 | 1241373 | + | - | PA14_14510 | hypothetical protein | 90 | 85 | 87 | 97 | 84 | 76 | 1 |
| 1170 | - | 1242002 | 1241505 | - | - | PA14_14520 | hypothetical protein | 26 | 23 | 28 | 25 | 24 | 26 | 1 |
| 1171 | - | 1242132 | 1243703 | + | - | PA14_14530 | hypothetical protein | 45 | 40 | 56 | 64 | 46 | 52 | 1 |
| 1172 | - | 1243940 | 1244128 | + | - | PA14_14540 | hypothetical protein | 325 | 640 | 387 | 515 | 488 | 522 | 0.580848337 |
| 1173 | - | 1244088 | 1244495 | + | - | PA14_14550 | hypothetical protein | 1239 | 2105 | 1599 | 2551 | 2280 | 2463 | 1 |
| 1174 | - | 1244649 | 1244524 | - | - | PA14_14560 | hypothetical protein | 106 | 118 | 110 | 142 | 132 | 103 | 1 |
| 1175 | - | 1244826 | 1244743 | - | - | PA14_14570 | Leu tRNA | 15 | 9 | 10 | 5 | 8 | 8 | 0.760328835 |
| 1176 | - | 1244929 | 1245972 | + | queA | PA14_14590 | S-adenosylmethionine--tRNA ribosyltransferase-isomerase | 54 | 52 | 54 | 56 | 48 | 42 | 1 |

| | A | B | C | D | E | F | G | H | I | J | K | L | M | N |
|---|---|---|---|---|---|---|---|---|---|---|---|---|---|---|
| 1177 | - | 1245985 | 1247103 | + | *tgt* | *PA14_14600* | queuine tRNA-ribosyltransferase | 64 | 66 | 77 | 60 | 64 | 54 | 1 |
| 1178 | - | 1247147 | 1247485 | + | *yajC* | *PA14_14610* | preprotein translocase subunit YajC | 74 | 95 | 71 | 50 | 56 | 42 | 1 |
| 1179 | - | 1247545 | 1249407 | + | *secD* | *PA14_14630* | preprotein translocase subunit SecD | 99 | 130 | 105 | 90 | 124 | 69 | 1 |
| 1180 | - | 1249418 | 1250338 | + | *secF* | *PA14_14650* | preprotein translocase subunit SecF | 168 | 209 | 157 | 121 | 187 | 105 | 1 |
| 1181 | - | 1250522 | 1251070 | + | - | *PA14_14660* | hypothetical protein | 1065 | 1016 | 1310 | 1071 | 1249 | 1239 | 1 |
| 1182 | - | 1251990 | 1251175 | - | - | *PA14_14680* | extragenic suppressor protein SuhB | 48 | 49 | 66 | 57 | 42 | 39 | 1 |
| 1183 | - | 1252142 | 1252915 | + | - | *PA14_14690* | methyltransferase | 62 | 42 | 52 | 45 | 43 | 49 | 1 |
| 1184 | - | 1252915 | 1253691 | + | *cysE* | *PA14_14700* | serine O-acetyltransferase | 241 | 212 | 215 | 212 | 210 | 200 | 1 |
| 1185 | - | 1253836 | 1254327 | + | - | *PA14_14710* | Rrf2 family protein | 56 | 150 | 47 | 223 | 263 | 76 | 0.033575691 |
| 1186 | - | 1254357 | 1255571 | + | *iscS* | *PA14_14730* | cysteine desulfurase | 244 | 367 | 136 | 591 | 674 | 363 | 1 |
| 1187 | - | 1255607 | 1255993 | + | - | *PA14_14740* | scaffold protein | 383 | 393 | 209 | 635 | 727 | 554 | 1 |
| 1188 | - | 1255994 | 1256014 | ? | - | predicted RNA | - | 412 | 402 | 200 | 735 | 861 | 643 | 1 |
| 1189 | - | 1256021 | 1256344 | + | - | *PA14_14750* | iron-binding protein IscA | 423 | 386 | 222 | 719 | 872 | 639 | 1 |
| 1190 | - | 1256352 | 1256873 | + | *hscB* | *PA14_14770* | co-chaperone HscB | 194 | 158 | 115 | 286 | 397 | 267 | 1 |
| 1191 | - | 1256916 | 1258775 | + | *hscA* | *PA14_14780* | chaperone protein HscA | 229 | 155 | 138 | 271 | 370 | 298 | 1 |
| 1192 | - | 1258782 | 1259120 | + | *fdx2* | *PA14_14800* | ferredoxin 2Fe-2S | 251 | 152 | 127 | 260 | 332 | 339 | 1 |
| 1193 | - | 1259146 | 1259346 | + | - | *PA14_14810* | hypothetical protein | 382 | 293 | 315 | 379 | 507 | 534 | 1 |
| 1194 | - | 1259590 | 1260021 | + | *ndk* | *PA14_14820* | nucleoside diphosphate kinase | 363 | 458 | 332 | 247 | 247 | 266 | 1 |
| 1195 | - | 1260046 | 1261185 | + | - | *PA14_14830* | hypothetical protein | 283 | 256 | 254 | 240 | 225 | 220 | 1 |
| 1196 | - | 1261203 | 1261961 | + | *pilF* | *PA14_14850* | type 4 fimbrial biogenesis protein PilF | 216 | 200 | 224 | 270 | 233 | 234 | 1 |
| 1197 | - | 1261958 | 1263001 | + | - | *PA14_14860* | hypothetical protein | 232 | 224 | 227 | 251 | 230 | 234 | 1 |
| 1198 | - | 1262998 | 1264113 | + | *ispG* | *PA14_14880* | 4-hydroxy-3-methylbut-2-en-1-yl diphosphate synthase | 414 | 372 | 401 | 392 | 363 | 394 | 1 |
| 1199 | - | 1264132 | 1265421 | + | *hisS* | *PA14_14890* | histidyl-tRNA synthetase | 345 | 324 | 345 | 321 | 318 | 345 | 1 |
| 1200 | - | 1265448 | 1266092 | + | - | *PA14_14900* | hypothetical protein | 580 | 528 | 570 | 558 | 572 | 573 | 1 |
| 1201 | - | 1266094 | 1267236 | + | - | *PA14_14910* | hypothetical protein | 480 | 441 | 455 | 459 | 448 | 454 | 1 |
| 1202 | - | 1267314 | 1268795 | + | *engA* | *PA14_14930* | GTP-binding protein EngA | 170 | 149 | 185 | 178 | 167 | 184 | 1 |
| 1203 | - | 1269092 | 1270240 | + | - | *PA14_14940* | aminotransferase | 30 | 28 | 29 | 25 | 32 | 25 | 1 |
| 1204 | - | 1270228 | 1271022 | + | - | *PA14_14960* | hypothetical protein | 164 | 163 | 151 | 154 | 147 | 156 | 1 |
| 1205 | - | 1271156 | 1271743 | + | - | *PA14_14975* | hypothetical protein | 331 | 284 | 357 | 401 | 365 | 385 | 1 |
| 1206 | - | 1272782 | 1271832 | - | - | *PA14_14990* | oxidoreductase | 84 | 82 | 82 | 78 | 84 | 74 | 1 |
| 1207 | - | 1272963 | 1273418 | + | - | *PA14_15000* | hypothetical protein | 38 | 36 | 39 | 45 | 36 | 39 | 1 |
| 1208 | - | 1273472 | 1273804 | + | - | *PA14_15020* | hypothetical protein | 27 | 31 | 34 | 26 | 26 | 23 | 1 |
| 1209 | - | 1275652 | 1273874 | - | *leuA* | *PA14_15030* | 2-isopropylmalate synthase | 104 | 109 | 92 | 86 | 95 | 97 | 1 |
| 1210 | - | 1276025 | 1276486 | + | - | *PA14_15050* | hypothetical protein | 230 | 208 | 268 | 288 | 230 | 185 | 1 |
| 1211 | - | 1276607 | 1278778 | + | *oprC* | *PA14_15070* | outer membrane copper receptor OprC | 192 | 234 | 171 | 219 | 230 | 166 | 1 |
| 1212 | - | 1278843 | 1280258 | + | - | *PA14_15080* | hypothetical protein | 263 | 240 | 251 | 277 | 275 | 245 | 1 |
| 1213 | - | 1280671 | 1280276 | - | - | *PA14_15090* | hypothetical protein | 178 | 192 | 188 | 195 | 200 | 192 | 1 |
| 1214 | - | 1281606 | 1280758 | - | - | *PA14_15100* | hypothetical protein | 82 | 85 | 107 | 111 | 95 | 91 | 1 |
| 1215 | - | 1281778 | 1282167 | + | - | *PA14_15110* | hypothetical protein | 222 | 203 | 218 | 235 | 202 | 244 | 1 |
| 1216 | - | 1282296 | 1282772 | + | - | *PA14_15120* | hypothetical protein | 154 | 112 | 111 | 180 | 148 | 202 | 1 |
| 1217 | - | 1282829 | 1283401 | + | - | *PA14_15130* | hypothetical protein | 291 | 267 | 287 | 447 | 337 | 412 | 1 |
| 1218 | - | 1284065 | 1283436 | - | - | *PA14_15140* | hypothetical protein | 301 | 208 | 241 | 405 | 330 | 389 | 1 |
| 1219 | - | 1284186 | 1285139 | + | - | *PA14_15150* | transcriptional regulator | 70 | 62 | 70 | 79 | 65 | 67 | 1 |
| 1220 | - | 1285843 | 1285136 | - | - | *PA14_15160* | hypothetical protein | 27 | 24 | 23 | 20 | 22 | 24 | 1 |
| 1221 | - | 1287182 | 1285902 | - | - | *PA14_15180* | transporter | 17 | 15 | 19 | 18 | 19 | 19 | 1 |
| 1222 | - | 1287673 | 1287179 | - | - | *PA14_15190* | hypothetical protein | 15 | 12 | 12 | 13 | 14 | 14 | 1 |
| 1223 | - | 1288786 | 1287746 | - | - | *PA14_15200* | hypothetical protein | 12 | 13 | 11 | 11 | 13 | 10 | 1 |
| 1224 | - | 1289873 | 1288938 | - | - | *PA14_15210* | LysR family transcriptional regulator | 32 | 30 | 29 | 27 | 22 | 22 | 1 |
| 1225 | - | 1291259 | 1289880 | - | *xseA* | *PA14_15230* | exodeoxyribonuclease VII large subunit | 54 | 47 | 48 | 43 | 41 | 37 | 1 |
| 1226 | - | 1292204 | 1291296 | - | - | *PA14_15240* | LysR family transcriptional regulator | 6 | 6 | 6 | 6 | 5 | 5 | 1 |
| 1227 | - | 1292312 | 1293094 | + | - | *PA14_15250* | hypothetical protein | 8 | 7 | 7 | 7 | 8 | 7 | 1 |
| 1228 | - | 1294221 | 1293079 | - | - | *PA14_15260* | acetylpolyamine aminohydrolase | 3 | 3 | 3 | 2 | 2 | 2 | 1 |
| 1229 | - | 1295408 | 1294251 | - | - | *PA14_15270* | hypothetical protein | 3 | 3 | 4 | 4 | 3 | 3 | 1 |
| 1230 | - | 1296317 | 1295421 | - | - | *PA14_15280* | hypothetical protein | 9 | 7 | 6 | 10 | 6 | 5 | 1 |
| 1231 | - | 1297429 | 1296452 | - | - | *PA14_15290* | transcriptional regulator | 20 | 20 | 21 | 24 | 17 | 22 | 1 |
| 1232 | - | 1297693 | 1299162 | + | *guaB* | *PA14_15310* | inosine 5'-monophosphate dehydrogenase | 192 | 234 | 241 | 175 | 179 | 128 | 1 |
| 1233 | - | 1299242 | 1300819 | + | *guaA* | *PA14_15340* | GMP synthase | 182 | 206 | 195 | 176 | 185 | 123 | 1 |
| 1234 | - | 1301126 | 1302328 | + | - | *PA14_15350* | integrase | 152 | 163 | 158 | 147 | 128 | 127 | 1 |
| 1235 | - | 1303723 | 1302743 | - | - | *PA14_15360* | hypothetical protein | 173 | 162 | 161 | 136 | 153 | 161 | 1 |
| 1236 | - | 1304052 | 1304930 | + | *repA* | *PA14_15380* | replicative helicase, RepA | 8 | 5 | 9 | 7 | 6 | 7 | 0.791445601 |
| 1237 | - | 1304084 | 1303857 | - | - | *PA14_15370* | hypothetical protein | 21 | 29 | 21 | 23 | 22 | 19 | 1 |
| 1238 | - | 1304902 | 1305789 | + | - | *PA14_15400* | replication protein, RepC | 25 | 21 | 25 | 27 | 18 | 22 | 1 |
| 1239 | - | 1306970 | 1306410 | - | - | *PA14_15430* | resolvase, essential for transposition | 17 | 17 | 17 | 17 | 11 | 14 | 1 |
| 1240 | - | 1308090 | 1307101 | - | - | *PA14_15435* | hypothetical protein | 37 | 38 | 33 | 37 | 35 | 35 | 1 |
| 1241 | - | 1308323 | 1308087 | - | *merE* | *PA14_15445* | mercury resistance protein | 38 | 36 | 32 | 39 | 33 | 35 | 1 |
| 1242 | - | 1308685 | 1308320 | - | *merD* | *PA14_15450* | transcriptional regulator MerD | 43 | 48 | 41 | 50 | 36 | 40 | 1 |
| 1243 | - | 1310388 | 1308703 | - | *merA* | *PA14_15460* | mercuric reductase | 46 | 50 | 42 | 50 | 40 | 40 | 1 |
| 1244 | - | 1310735 | 1310460 | - | *merP* | *PA14_15470* | periplasmic mercuric ion binding protein, MerP | 75 | 79 | 66 | 80 | 59 | 53 | 1 |
| 1245 | - | 1311098 | 1310748 | - | *merT* | *PA14_15475* | mercuric transport protein | 33 | 42 | 30 | 42 | 32 | 29 | 1 |
| 1246 | - | 1311170 | 1311604 | + | *merR* | *PA14_15480* | transcriptional regulator MerR | 168 | 195 | 167 | 151 | 126 | 113 | 1 |
| 1247 | - | 1311858 | 1312034 | + | - | *PA14_15490* | hypothetical protein | 12 | 12 | 13 | 12 | 10 | 10 | 1 |
| 1248 | - | 1312074 | 1312361 | + | - | *PA14_15500* | oriT-binding protein, TraK | 27 | 23 | 21 | 19 | 19 | 27 | 1 |
| 1249 | - | 1312655 | 1313026 | + | *traJ* | *PA14_15510* | conjugal transfer relaxosome component TraJ | 23 | 22 | 24 | 27 | 19 | 23 | 1 |
| 1250 | - | 1313291 | 1314076 | + | *trbJ* | *PA14_15520* | conjugal transfer protein TrbJ | 16 | 15 | 17 | 17 | 14 | 11 | 1 |
| 1251 | - | 1314094 | 1314369 | + | - | *PA14_15530* | entry/exclusion protein TrbK | 42 | 27 | 48 | 15 | 11 | 23 | 0.615940863 |
| 1252 | - | 1314366 | 1315988 | + | - | *PA14_15540* | mating pair formation protein TrbL | 45 | 35 | 51 | 27 | 24 | 35 | 1 |
| 1253 | - | 1316812 | 1316504 | - | - | *PA14_15560* | hypothetical protein | 89 | 100 | 84 | 74 | 70 | 64 | 1 |
| 1254 | - | 1317148 | 1317348 | + | - | *PA14_15570* | hypothetical protein | 66 | 79 | 82 | 68 | 81 | 61 | 1 |
| 1255 | - | 1317407 | 1320187 | + | - | *PA14_15580* | Type II restriction enzyme, methylase subunit | 133 | 140 | 123 | 143 | 149 | 126 | 1 |
| 1256 | - | 1321218 | 1323335 | + | - | *PA14_15590* | hypothetical protein | 196 | 223 | 178 | 203 | 205 | 199 | 1 |
| 1257 | - | 1323328 | 1324512 | + | - | *PA14_15600* | hypothetical protein | 252 | 276 | 230 | 265 | 269 | 268 | 1 |
| 1258 | - | 1324805 | 1327411 | + | - | *PA14_15610* | hypothetical protein | 234 | 253 | 221 | 240 | 226 | 248 | 1 |
| 1259 | - | 1328012 | 1328974 | + | - | *PA14_15620* | oxidoreductase | 9 | 9 | 9 | 8 | 7 | 5 | 1 |
| 1260 | - | 1328895 | 1330193 | + | - | *PA14_15630* | hypothetical protein | 17 | 17 | 20 | 21 | 18 | 16 | 1 |

| | A | B | C | D | E | F | G | H | I | J | K | L | M | N |
|---|---|---|---|---|---|---|---|---|---|---|---|---|---|---|
| 1261 | - | 1331201 | 1330245 | - | - | PA14_15650 | hypothetical protein | 31 | 27 | 26 | 27 | 22 | 23 | 1 |
| 1262 | - | 1331350 | 1332345 | + | - | PA14_15660 | oxidoreductase | 21 | 18 | 20 | 18 | 15 | 16 | 1 |
| 1263 | - | 1332509 | 1333900 | + | - | PA14_15670 | metallo-oxidoreductase | 59 | 69 | 76 | 70 | 88 | 71 | 1 |
| 1264 | - | 1333891 | 1334439 | + | - | PA14_15680 | hypothetical protein | 64 | 84 | 94 | 109 | 105 | 94 | 1 |
| 1265 | - | 1334828 | 1336084 | + | - | PA14_15700 | amino acid permease | 14 | 11 | 13 | 11 | 11 | 10 | 1 |
| 1266 | - | 1336210 | 1336794 | + | - | PA14_15710 | hypothetical protein | 39 | 43 | 43 | 35 | 30 | 31 | 1 |
| 1267 | - | 1338163 | 1336808 | - | - | PA14_15720 | transglycosylase | 51 | 47 | 52 | 52 | 47 | 47 | 1 |
| 1268 | - | 1338564 | 1342460 | + | purL | PA14_15740 | phosphoribosylformylglycinamidine synthase | 147 | 141 | 140 | 134 | 135 | 112 | 1 |
| 1269 | - | 1342467 | 1342745 | + | - | PA14_15750 | hypothetical protein | 142 | 181 | 236 | 230 | 198 | 229 | 1 |
| 1270 | - | 1342925 | 1343242 | + | - | PA14_15770 | hypothetical protein | 175 | 172 | 213 | 175 | 137 | 139 | 1 |
| 1271 | - | 1345156 | 1343444 | - | - | PA14_15780 | PTS system N-acetylglucosamine-specific IIBC component | 30 | 30 | 30 | 26 | 29 | 31 | 1 |
| 1272 | - | 1347709 | 1345181 | - | - | PA14_15790 | phosphoenolpyruvate-protein phosphotransferase | 61 | 54 | 56 | 55 | 54 | 52 | 1 |
| 1273 | - | 1348748 | 1347726 | - | - | PA14_15810 | aminotransferase | 20 | 25 | 20 | 19 | 23 | 20 | 1 |
| 1274 | - | 1349836 | 1348745 | - | - | PA14_15820 | N-acetylglucosamine-6-phosphate deacetylase | 11 | 16 | 12 | 13 | 14 | 13 | 1 |
| 1275 | - | 1350596 | 1349853 | - | - | PA14_15830 | GntR family transcriptional regulator | 9 | 9 | 10 | 11 | 10 | 8 | 1 |
| 1276 | - | 1351339 | 1350839 | - | - | PA14_15840 | hypothetical protein | 69 | 59 | 79 | 56 | 56 | 62 | 1 |
| 1277 | - | 1351956 | 1351402 | - | - | PA14_15850 | hypothetical protein | 43 | 40 | 48 | 36 | 39 | 43 | 1 |
| 1278 | - | 1352097 | 1352708 | + | - | PA14_15860 | hypothetical protein | 109 | 112 | 119 | 85 | 109 | 105 | 1 |
| 1279 | - | 1352746 | 1353270 | + | - | PA14_15870 | hypothetical protein | 462 | 435 | 463 | 496 | 449 | 506 | 1 |
| 1280 | - | 1353267 | 1353461 | + | - | PA14_15880 | hypothetical protein | 155 | 154 | 141 | 141 | 156 | 128 | 1 |
| 1281 | - | 1353499 | 1354680 | + | purT | PA14_15890 | phosphoribosylglycinamide formyltransferase 2 | 127 | 125 | 108 | 103 | 117 | 111 | 1 |
| 1282 | - | 1354682 | 1355431 | + | - | PA14_15910 | hypothetical protein | 221 | 174 | 210 | 210 | 214 | 207 | 1 |
| 1283 | - | 1356752 | 1355439 | - | - | PA14_15920 | major facilitator transporter | 13 | 12 | 15 | 15 | 14 | 13 | 1 |
| 1284 | - | 1358348 | 1357056 | - | - | PA14_15930 | hemolysin | 35 | 33 | 37 | 43 | 36 | 35 | 1 |
| 1285 | - | 1359161 | 1358361 | - | - | PA14_15940 | hypothetical protein | 41 | 30 | 43 | 36 | 30 | 34 | 1 |
| 1286 | - | 1359462 | 1360835 | + | ffh | PA14_15960 | signal recognition particle protein Ffh | 132 | 125 | 146 | 120 | 109 | 97 | 1 |
| 1287 | - | 1361045 | 1361296 | + | rpsP | PA14_15970 | 30S ribosomal protein S16 | 548 | 746 | 775 | 573 | 490 | 343 | 1 |
| 1288 | - | 1361312 | 1361839 | + | rimM | PA14_15980 | 16S rRNA-processing protein RimM | 528 | 630 | 702 | 541 | 431 | 311 | 1 |
| 1289 | - | 1361846 | 1362604 | + | trmD | PA14_15990 | tRNA (guanine-N(1)-)-methyltransferase | 272 | 319 | 393 | 300 | 266 | 199 | 1 |
| 1290 | - | 1362646 | 1362996 | + | rplS | PA14_16000 | 50S ribosomal protein L19 | 322 | 453 | 317 | 183 | 256 | 185 | 1 |
| 1291 | - | 1363148 | 1363573 | + | - | PA14_16010 | hypothetical protein | 40 | 45 | 48 | 45 | 40 | 39 | 1 |
| 1292 | - | 1364329 | 1363643 | - | - | PA14_16020 | hypothetical protein | 64 | 117 | 89 | 68 | 77 | 70 | 0.641536103 |
| 1293 | - | 1366277 | 1364442 | - | - | PA14_16030 | sodium/hydrogen antiporter | 35 | 30 | 34 | 35 | 33 | 31 | 1 |
| 1294 | - | 1366492 | 1367388 | + | xerD | PA14_16040 | site-specific tyrosine recombinase XerD | 33 | 23 | 26 | 22 | 23 | 19 | 1 |
| 1295 | - | 1367515 | 1368243 | + | dsbC | PA14_16050 | thiol:disulfide interchange protein DsbC | 110 | 108 | 115 | 91 | 118 | 93 | 1 |
| 1296 | - | 1368457 | 1369761 | + | hom | PA14_16070 | homoserine dehydrogenase | 123 | 122 | 121 | 106 | 124 | 104 | 1 |
| 1297 | - | 1369814 | 1371223 | + | thrC | PA14_16090 | threonine synthase | 96 | 114 | 84 | 80 | 89 | 83 | 1 |
| 1298 | 1372370 | 1372497 | 1371277 | - | - | PA14_16100 | hypothetical protein | 55 | 36 | 35 | 38 | 31 | 50 | 1 |
| 1299 | 1372370 | 1372594 | 1372878 | + | - | PA14_16110 | hypothetical protein | 50 | 38 | 44 | 68 | 47 | 69 | 1 |
| 1300 | - | 1374294 | 1373065 | - | - | PA14_16130 | hypothetical protein | 51 | 41 | 48 | 50 | 39 | 49 | 1 |
| 1301 | - | 1374776 | 1375219 | + | - | PA14_16140 | hypothetical protein | 112 | 104 | 94 | 84 | 122 | 84 | 1 |
| 1302 | - | 1375234 | 1375929 | + | - | PA14_16150 | hypothetical protein | 110 | 104 | 79 | 78 | 106 | 81 | 1 |
| 1303 | - | 1375985 | 1376626 | + | - | PA14_16160 | hypothetical protein | 34 | 29 | 23 | 20 | 32 | 19 | 1 |
| 1304 | - | 1376662 | 1378728 | + | - | PA14_16180 | hypothetical protein | 83 | 68 | 58 | 48 | 73 | 57 | 1 |
| 1305 | - | 1378878 | 1384118 | + | - | PA14_16190 | hypothetical protein | 65 | 49 | 43 | 39 | 54 | 45 | 1 |
| 1306 | - | 1384137 | 1384829 | + | - | PA14_16200 | hypothetical protein | 47 | 38 | 30 | 26 | 41 | 37 | 1 |
| 1307 | - | 1384936 | 1385475 | + | - | PA14_16210 | hypothetical protein | 19 | 13 | 21 | 16 | 16 | 14 | 1 |
| 1308 | - | 1385519 | 1387234 | + | recJ | PA14_16220 | single-stranded-DNA-specific exonuclease RecJ | 28 | 24 | 27 | 27 | 27 | 25 | 1 |
| 1309 | 1387361 | 1387521 | 1389017 | + | lasB | PA14_16250 | elastase LasB | 583 | 241 | 181 | 863 | 566 | 1089 | 1 |
| 1310 | - | 1389187 | 1390293 | + | - | PA14_16260 | FMN oxidoreductase | 247 | 252 | 331 | 248 | 283 | 314 | 1 |
| 1311 | - | 1390481 | 1390831 | + | - | PA14_16270 | hypothetical protein | 110 | 88 | 102 | 65 | 72 | 74 | 1 |
| 1312 | - | 1391485 | 1390844 | - | - | PA14_16280 | transcriptional regulator | 291 | 232 | 252 | 121 | 208 | 209 | 1 |
| 1313 | - | 1391688 | 1392113 | + | - | PA14_16290 | hypothetical protein | 272 | 85 | 176 | 88 | 99 | 86 | 0.012540033 |
| 1314 | - | 1392121 | 1392282 | + | - | PA14_16300 | hypothetical protein | 137 | 50 | 95 | 54 | 59 | 46 | 0.000861338 |
| 1315 | - | 1392430 | 1393671 | + | - | PA14_16310 | MFS permease | 34 | 9 | 14 | 10 | 7 | 31 | 2.90451E-05 |
| 1316 | - | 1394099 | 1393758 | - | - | PA14_16320 | peptidyl-prolyl cis-trans isomerase, FkbP-type | 23 | 35 | 21 | 24 | 29 | 22 | 1 |
| 1317 | - | 1394310 | 1396016 | + | - | PA14_16330 | hypothetical protein | 61 | 73 | 64 | 56 | 61 | 52 | 1 |
| 1318 | - | 1396898 | 1396095 | - | - | PA14_16340 | hypothetical protein | 31 | 29 | 29 | 24 | 24 | 25 | 1 |
| 1319 | - | 1397660 | 1397019 | - | - | PA14_16350 | two-component response regulator | 9 | 7 | 10 | 11 | 8 | 7 | 1 |
| 1320 | - | 1399846 | 1397984 | - | - | PA14_16360 | hypothetical protein | 19 | 19 | 20 | 20 | 18 | 19 | 1 |
| 1321 | - | 1400168 | 1400863 | + | - | PA14_16370 | hypothetical protein | 193 | 237 | 205 | 162 | 211 | 172 | 1 |
| 1322 | - | 1401805 | 1400900 | - | - | PA14_16380 | LysR family transcriptional regulator | 78 | 93 | 87 | 80 | 76 | 81 | 1 |
| 1323 | - | 1401977 | 1403650 | + | - | PA14_16390 | GMC-type oxidoreductase | 64 | 100 | 84 | 137 | 84 | 96 | 1 |
| 1324 | - | 1403759 | 1405378 | + | - | PA14_16410 | MFS transporter | 34 | 51 | 40 | 61 | 40 | 45 | 1 |
| 1325 | - | 1405645 | 1407273 | + | wspA | PA14_16430 | chemotaxis transducer | 232 | 233 | 271 | 300 | 253 | 269 | 1 |
| 1326 | - | 1407278 | 1407793 | + | - | PA14_16440 | hypothetical protein | 232 | 258 | 270 | 288 | 250 | 269 | 1 |
| 1327 | - | 1407790 | 1409058 | + | wspC | PA14_16450 | methyltransferase | 145 | 136 | 149 | 160 | 136 | 155 | 1 |
| 1328 | - | 1409051 | 1409740 | + | wspD | PA14_16460 | CheW domain-containing protein | 84 | 92 | 84 | 111 | 93 | 97 | 1 |
| 1329 | - | 1409737 | 1412046 | + | - | PA14_16470 | chemotaxis sensor/effector fusion protein | 175 | 171 | 186 | 202 | 180 | 183 | 1 |
| 1330 | - | 1412043 | 1413050 | + | wspF | PA14_16480 | chemotaxis-specific methylesterase | 207 | 195 | 239 | 238 | 218 | 239 | 1 |
| 1331 | - | 1413158 | 1414201 | + | wspR | PA14_16500 | two-component response regulator | 177 | 176 | 189 | 217 | 171 | 218 | 1 |
| 1332 | - | 1414421 | 1415419 | + | - | PA14_16510 | peptide chain release factor 2 | 131 | 135 | 147 | 145 | 127 | 133 | 1 |
| 1333 | - | 1415600 | 1417105 | + | lysS | PA14_16530 | lysyl-tRNA synthetase | 62 | 84 | 74 | 56 | 66 | 50 | 1 |
| 1334 | - | 1417317 | 1418030 | + | - | PA14_16550 | TetR family transcriptional regulator | 43 | 41 | 40 | 39 | 38 | 31 | 1 |
| 1335 | - | 1418127 | 1418678 | + | - | PA14_16560 | lipoprotein | 162 | 127 | 156 | 178 | 144 | 156 | 1 |
| 1336 | - | 1418687 | 1419982 | + | - | PA14_16580 | hypothetical protein | 94 | 103 | 91 | 116 | 117 | 112 | 1 |
| 1337 | - | 1420051 | 1420797 | + | - | PA14_16590 | hypothetical protein | 140 | 135 | 119 | 113 | 129 | 125 | 1 |
| 1338 | - | 1420794 | 1421699 | + | - | PA14_16600 | alpha/beta hydrolase | 123 | 123 | 119 | 111 | 133 | 117 | 1 |
| 1339 | - | 1421696 | 1422019 | + | - | PA14_16610 | hypothetical protein | 123 | 136 | 111 | 126 | 114 | 126 | 1 |
| 1340 | - | 1422016 | 1422537 | + | - | PA14_16620 | hypothetical protein | 140 | 127 | 144 | 129 | 157 | 130 | 1 |
| 1341 | - | 1423398 | 1422613 | - | - | PA14_16630 | outer membrane protein, OmpA | 873 | 1210 | 912 | 806 | 1347 | 1018 | 1 |
| 1342 | - | 1423851 | 1423447 | - | - | PA14_16640 | lipoprotein | 890 | 1240 | 1126 | 832 | 1318 | 1058 | 1 |
| 1343 | - | 1426481 | 1424253 | - | - | PA14_16660 | metal-transporting P-type ATPase | 77 | 78 | 53 | 61 | 56 | 73 | 1 |
| 1344 | - | 1426595 | 1427065 | + | - | PA14_16670 | transcriptional regulator CadR | 230 | 276 | 257 | 275 | 287 | 301 | 1 |

| | A | B | C | D | E | F | G | H | I | J | K | L | M | N |
|---|---|---|---|---|---|---|---|---|---|---|---|---|---|---|
| 1345 | - | 1427418 | 1427074 | - | - | *PA14_16680* | hypothetical protein | 378 | 460 | 497 | 447 | 427 | 607 | 1 |
| 1346 | - | 1427626 | 1430262 | + | *ppc* | *PA14_16690* | phosphoenolpyruvate carboxylase | 182 | 156 | 147 | 174 | 160 | 185 | 1 |
| 1347 | - | 1430423 | 1431070 | + | *adk* | *PA14_16700* | adenylate kinase | 206 | 188 | 189 | 158 | 168 | 168 | 1 |
| 1348 | - | 1431174 | 1431854 | + | - | *PA14_16710* | hypothetical protein | 36 | 39 | 43 | 31 | 34 | 29 | 1 |
| 1349 | - | 1431928 | 1432275 | + | - | *PA14_16720* | hypothetical protein | 77 | 70 | 107 | 70 | 68 | 75 | 1 |
| 1350 | - | 1432294 | 1433160 | + | - | *PA14_16730* | hypothetical protein | 40 | 26 | 44 | 30 | 33 | 33 | 1 |
| 1351 | - | 1433233 | 1434057 | + | - | *PA14_16740* | hypothetical protein | 24 | 27 | 24 | 26 | 22 | 24 | 1 |
| 1352 | - | 1434062 | 1434757 | + | - | *PA14_16750* | hypothetical protein | 16 | 18 | 18 | 22 | 17 | 19 | 1 |
| 1353 | - | 1434813 | 1435598 | + | - | *PA14_16770* | hypothetical protein | 34 | 30 | 38 | 29 | 27 | 24 | 1 |
| 1354 | - | 1435627 | 1436904 | + | - | *PA14_16780* | hypothetical protein | 47 | 97 | 69 | 71 | 78 | 51 | 0.490513327 |
| 1355 | - | 1437549 | 1436911 | - | - | *PA14_16790* | TetR family transcriptional regulator | 42 | 262 | 162 | 170 | 214 | 130 | 6.13255E-17 |
| 1356 | - | 1437645 | 1438748 | + | - | *PA14_16800* | efflux transmembrane protein | 15 | 186 | 92 | 94 | 94 | 43 | 2.97801E-94 |
| 1357 | - | 1438753 | 1441830 | + | - | *PA14_16820* | efflux transmembrane protein | 27 | 237 | 135 | 141 | 132 | 64 | 2.36603E-25 |
| 1358 | - | 1442553 | 1441873 | - | - | *PA14_16830* | hypothetical protein | 36 | 139 | 88 | 68 | 90 | 38 | 1.87587E-05 |
| 1359 | - | 1443088 | 1442690 | - | - | *PA14_16840* | lipoprotein | 54 | 80 | 39 | 38 | 63 | 46 | 1 |
| 1360 | - | 1445735 | 1443231 | - | *plsB* | *PA14_16860* | glycerol-3-phosphate acyltransferase | 40 | 43 | 41 | 38 | 41 | 33 | 1 |
| 1361 | - | 1446019 | 1446942 | + | - | *PA14_16870* | ABC transporter ATP-binding protein | 30 | 30 | 30 | 32 | 33 | 27 | 1 |
| 1362 | - | 1446939 | 1447673 | + | - | *PA14_16880* | ABC transporter permease | 24 | 26 | 22 | 25 | 29 | 19 | 1 |
| 1363 | - | 1447684 | 1449531 | + | - | *PA14_16890* | auxiliary component of ABC transporter | 38 | 43 | 33 | 34 | 39 | 29 | 1 |
| 1364 | - | 1449535 | 1450515 | + | - | *PA14_16910* | hypothetical protein | 106 | 110 | 116 | 111 | 110 | 93 | 1 |
| 1365 | - | 1450944 | 1450522 | - | - | *PA14_16920* | hypothetical protein | 150 | 147 | 185 | 150 | 141 | 148 | 1 |
| 1366 | - | 1452146 | 1450941 | - | - | *PA14_16930* | pyridoxal-phosphate dependent protein | 87 | 78 | 105 | 87 | 87 | 89 | 1 |
| 1367 | - | 1453275 | 1452241 | - | *dapD* | *PA14_16950* | tetrahydrodipicolinate succinylase | 104 | 110 | 91 | 85 | 90 | 80 | 1 |
| 1368 | - | 1453928 | 1453305 | - | - | *PA14_16960* | amino acid transporter LysE | 43 | 29 | 43 | 31 | 36 | 33 | 1 |
| 1369 | - | 1454288 | 1453941 | - | - | *PA14_16970* | arsenate reductase | 53 | 44 | 56 | 35 | 35 | 38 | 1 |
| 1370 | - | 1454591 | 1454944 | + | - | *PA14_16980* | hypothetical protein | 104 | 119 | 131 | 112 | 111 | 98 | 1 |
| 1371 | - | 1455252 | 1454959 | - | - | *PA14_16990* | hypothetical protein | 792 | 879 | 846 | 1050 | 1104 | 869 | 1 |
| 1372 | - | 1455933 | 1455574 | - | - | *PA14_17000* | hypothetical protein | 7 | 7 | 10 | 11 | 5 | 6 | 1 |
| 1373 | - | 1456171 | 1457916 | + | - | *PA14_17010* | Na(+)/H(+) exchanger protein | 78 | 54 | 63 | 52 | 50 | 53 | 1 |
| 1374 | - | 1459175 | 1457967 | - | - | *PA14_17030* | succinyldiaminopimelate transaminase | 86 | 72 | 72 | 75 | 63 | 69 | 1 |
| 1375 | - | 1461895 | 1459193 | - | *glnD* | *PA14_17040* | PII uridylyl-transferase | 59 | 58 | 64 | 65 | 57 | 62 | 1 |
| 1376 | - | 1462847 | 1462062 | - | *map* | *PA14_17050* | methionine aminopeptidase | 172 | 142 | 195 | 142 | 131 | 142 | 1 |
| 1377 | - | 1463112 | 1463852 | + | *rpsB* | *PA14_17060* | 30S ribosomal protein S2 | 278 | 331 | 278 | 189 | 259 | 153 | 1 |
| 1378 | - | 1463983 | 1464852 | + | *tsf* | *PA14_17070* | elongation factor Ts | 145 | 241 | 149 | 94 | 143 | 73 | 1 |
| 1379 | - | 1465051 | 1465788 | + | *pyrH* | *PA14_17080* | uridylate kinase | 256 | 281 | 277 | 266 | 267 | 175 | 1 |
| 1380 | - | 1465791 | 1466348 | + | *frr* | *PA14_17100* | ribosome recycling factor | 279 | 400 | 295 | 276 | 285 | 215 | 1 |
| 1381 | - | 1466364 | 1467119 | + | *uppS* | *PA14_17110* | UDP pyrophosphate synthetase | 155 | 178 | 170 | 136 | 155 | 110 | 1 |
| 1382 | - | 1467113 | 1467928 | + | *cdsA* | *PA14_17120* | phosphatidate cytidylyltransferase | 150 | 151 | 176 | 146 | 157 | 106 | 1 |
| 1383 | - | 1467925 | 1469115 | + | *dxr* | *PA14_17130* | 1-deoxy-D-xylulose 5-phosphate reductoisomerase | 201 | 204 | 207 | 187 | 190 | 146 | 1 |
| 1384 | - | 1469141 | 1470493 | + | - | *PA14_17140* | membrane-associated zinc metalloprotease | 181 | 150 | 198 | 177 | 178 | 146 | 1 |
| 1385 | - | 1470564 | 1472948 | + | - | *PA14_17150* | outer membrane antigen | 158 | 170 | 158 | 186 | 170 | 161 | 1 |
| 1386 | - | 1472999 | 1473505 | + | - | *PA14_17170* | hypothetical protein | 548 | 582 | 556 | 524 | 493 | 481 | 1 |
| 1387 | - | 1473505 | 1474566 | + | *lpxD* | *PA14_17180* | UDP-3-O-[3-hydroxymyristoyl] glucosamine N-acyltransferase | 292 | 279 | 264 | 231 | 264 | 234 | 1 |
| 1388 | - | 1474612 | 1475052 | + | *fabZ* | *PA14_17190* | (3R)-hydroxymyristoyl-ACP dehydratase | 197 | 154 | 199 | 180 | 174 | 178 | 1 |
| 1389 | - | 1475049 | 1475825 | + | *lpxA* | *PA14_17210* | UDP-N-acetylglucosamine acyltransferase | 250 | 236 | 266 | 225 | 237 | 220 | 1 |
| 1390 | - | 1475829 | 1476965 | + | *lpxB* | *PA14_17220* | lipid-A-disaccharide synthase | 87 | 85 | 84 | 80 | 94 | 71 | 1 |
| 1391 | - | 1476965 | 1477570 | + | *rnhB* | *PA14_17230* | ribonuclease HII | 125 | 124 | 126 | 112 | 134 | 98 | 1 |
| 1392 | - | 1477787 | 1479202 | + | - | *PA14_17250* | amino acid permease | 49 | 61 | 56 | 41 | 55 | 36 | 1 |
| 1393 | - | 1479332 | 1482853 | + | *dnaE* | *PA14_17260* | DNA polymerase III subunit alpha | 121 | 117 | 119 | 118 | 120 | 110 | 1 |
| 1394 | - | 1483003 | 1483953 | + | *accA* | *PA14_17270* | acetyl-CoA carboxylase carboxyltransferase subunit alpha | 125 | 121 | 126 | 96 | 103 | 93 | 1 |
| 1395 | - | 1484023 | 1485351 | + | - | *PA14_17280* | hypothetical protein | 80 | 75 | 81 | 73 | 76 | 60 | 1 |
| 1396 | - | 1485522 | 1487150 | + | *pyrG* | *PA14_17290* | CTP synthetase | 220 | 189 | 230 | 216 | 173 | 180 | 1 |
| 1397 | - | 1487153 | 1487998 | + | *kdsA* | *PA14_17310* | 2-dehydro-3-deoxyphosphooctonate aldolase | 266 | 261 | 266 | 267 | 236 | 230 | 1 |
| 1398 | - | 1488044 | 1489333 | + | *eno* | *PA14_17320* | phosphopyruvate hydratase | 349 | 337 | 326 | 297 | 305 | 264 | 1 |
| 1399 | - | 1489398 | 1489682 | + | - | *PA14_17330* | hypothetical protein | 188 | 163 | 182 | 154 | 161 | 168 | 1 |
| 1400 | - | 1489702 | 1490406 | + | *ispD* | *PA14_17340* | 2-C-methyl-D-erythritol 4-phosphate cytidylyltransferase | 49 | 31 | 49 | 28 | 35 | 40 | 1 |
| 1401 | - | 1490715 | 1490467 | - | - | *PA14_17350* | hypothetical protein | 45 | 52 | 54 | 40 | 43 | 42 | 1 |
| 1402 | - | 1491907 | 1490681 | - | - | *PA14_17370* | inner membrane protein | 51 | 38 | 61 | 36 | 32 | 43 | 1 |
| 1403 | - | 1492891 | 1491983 | - | - | *PA14_17380* | LysR family transcriptional regulator | 39 | 32 | 39 | 32 | 33 | 44 | 1 |
| 1404 | - | 1493023 | 1494135 | + | *adhC* | *PA14_17400* | alcohol dehydrogenase | 145 | 159 | 162 | 142 | 139 | 138 | 1 |
| 1405 | - | 1494189 | 1495040 | + | - | *PA14_17410* | esterase | 89 | 95 | 84 | 83 | 74 | 83 | 1 |
| 1406 | - | 1495111 | 1495584 | + | *ispF* | *PA14_17420* | 2-C-methyl-D-erythritol 2,4-cyclodiphosphate synthase | 113 | 113 | 127 | 121 | 109 | 100 | 1 |
| 1407 | - | 1495581 | 1496648 | + | *truD* | *PA14_17440* | tRNA pseudouridine synthase D | 83 | 81 | 87 | 84 | 81 | 72 | 1 |
| 1408 | - | 1496636 | 1497385 | + | *surE* | *PA14_17450* | stationary phase survival protein SurE | 106 | 101 | 103 | 109 | 112 | 97 | 1 |
| 1409 | - | 1497418 | 1498053 | + | *pcm* | *PA14_17460* | protein-L-isoaspartate O-methyltransferase | 249 | 256 | 255 | 298 | 296 | 233 | 1 |
| 1410 | - | 1498099 | 1498992 | + | - | *PA14_17470* | hypothetical protein | 1134 | 1168 | 1154 | 1376 | 1060 | 1279 | 1 |
| 1411 | - | 1499097 | 1500101 | + | *rpoS* | *PA14_17480* | RNA polymerase sigma factor RpoS | 641 | 827 | 492 | 742 | 707 | 818 | 1 |
| 1412 | - | 1500850 | 1500527 | - | *fdxA* | *PA14_17490* | ferredoxin I | 61 | 76 | 73 | 48 | 71 | 58 | 1 |
| 1413 | - | 1503484 | 1500917 | - | *mutS* | *PA14_17500* | DNA mismatch repair protein MutS | 62 | 60 | 69 | 54 | 54 | 49 | 1 |
| 1414 | - | 1504617 | 1503610 | - | - | *PA14_17510* | hypothetical protein | 31 | 24 | 34 | 32 | 26 | 23 | 1 |
| 1415 | - | 1504765 | 1505271 | + | - | *PA14_17520* | hypothetical protein | 138 | 143 | 147 | 142 | 123 | 119 | 1 |
| 1416 | - | 1505405 | 1506445 | + | *recA* | *PA14_17530* | recombinase A | 219 | 256 | 220 | 211 | 253 | 229 | 1 |
| 1417 | - | 1506451 | 1506912 | + | *recX* | *PA14_17540* | recombination regulator RecX | 96 | 109 | 91 | 90 | 89 | 92 | 1 |
| 1418 | - | 1508004 | 1506934 | - | - | *PA14_17550* | hypothetical protein | 184 | 192 | 185 | 149 | 156 | 184 | 1 |
| 1419 | - | 1509490 | 1508087 | - | - | *PA14_17570* | hypothetical protein | 230 | 193 | 235 | 126 | 138 | 193 | 1 |
| 1420 | - | 1509670 | 1512075 | + | - | *PA14_17580* | hypothetical protein | 212 | 197 | 256 | 141 | 167 | 164 | 1 |
| 1421 | - | 1512420 | 1512199 | - | - | *PA14_17590* | hypothetical protein | 99 | 99 | 131 | 95 | 89 | 90 | 1 |
| 1422 | - | 1512849 | 1512439 | - | - | *PA14_17600* | hypothetical protein | 99 | 96 | 117 | 85 | 93 | 82 | 1 |
| 1423 | - | 1514089 | 1513025 | - | *potD* | *PA14_17610* | polyamine ABC transporter | 7 | 7 | 8 | 8 | 7 | 6 | 1 |
| 1424 | - | 1514948 | 1514178 | - | *potC* | *PA14_17620* | polyamine transport protein PotC | 7 | 7 | 9 | 8 | 7 | 8 | 1 |
| 1425 | - | 1515834 | 1514941 | - | *potB* | *PA14_17630* | polyamine transport protein PotB | 5 | 5 | 6 | 5 | 5 | 4 | 1 |
| 1426 | - | 1516930 | 1515839 | - | *potA* | *PA14_17640* | polyamine transport protein PotA | 6 | 4 | 6 | 5 | 4 | 5 | 0.859738761 |
| 1427 | - | 1517189 | 1517905 | + | - | *PA14_17650* | hypothetical protein | 29 | 27 | 33 | 28 | 23 | 21 | 1 |
| 1428 | - | 1518835 | 1517909 | - | - | *PA14_17660* | hypothetical protein | 24 | 29 | 27 | 23 | 23 | 20 | 1 |

| | A | B | C | D | E | F | G | H | I | J | K | L | M | N |
|---|---|---|---|---|---|---|---|---|---|---|---|---|---|---|
| 1429 | - | 1518968 | 1519621 | + | - | *PA14_17670* | LuxR family transcriptional regulator | 62 | 49 | 55 | 48 | 53 | 48 | 1 |
| 1430 | - | 1519693 | 1520064 | + | *dgkA* | *PA14_17675* | diacylglycerol kinase | 25 | 23 | 20 | 20 | 20 | 18 | 1 |
| 1431 | - | 1521744 | 1520134 | - | - | *PA14_17690* | hypothetical protein | 42 | 43 | 36 | 36 | 40 | 36 | 1 |
| 1432 | - | 1521988 | 1522251 | + | *rpmE2* | *PA14_17700* | 50S ribosomal protein L31 | 22 | 38 | 23 | 64 | 51 | 34 | 0.818857487 |
| 1433 | - | 1522251 | 1522403 | + | *rpmJ* | *PA14_17710* | 50S ribosomal protein L36 | 35 | 51 | 28 | 75 | 58 | 42 | 1 |
| 1434 | - | 1523216 | 1522431 | - | - | *PA14_17720* | LuxR family transcriptional regulator | 32 | 30 | 30 | 37 | 30 | 25 | 1 |
| 1435 | - | 1523344 | 1524159 | + | - | *PA14_17730* | hypothetical protein | 18 | 16 | 20 | 19 | 20 | 16 | 1 |
| 1436 | - | 1524359 | 1525681 | + | - | *PA14_17740* | amino acid ABC transporter permease | 8 | 8 | 11 | 10 | 8 | 9 | 1 |
| 1437 | - | 1525771 | 1526844 | + | - | *PA14_17760* | methylated-DNA--protein-cysteine methyltransferase | 15 | 16 | 18 | 20 | 14 | 16 | 1 |
| 1438 | - | 1528250 | 1526841 | - | - | *PA14_17780* | major facilitator transporter | 11 | 9 | 11 | 11 | 11 | 9 | 1 |
| 1439 | - | 1529356 | 1528469 | - | - | *PA14_17790* | LysR family transcriptional regulator | 25 | 22 | 24 | 24 | 18 | 24 | 1 |
| 1440 | - | 1529470 | 1531197 | + | - | *PA14_17810* | acyl-CoA dehydrogenase | 10 | 8 | 10 | 10 | 7 | 8 | 1 |
| 1441 | - | 1531236 | 1532423 | + | - | *PA14_17820* | hypothetical protein | 5 | 3 | 5 | 5 | 3 | 4 | 0.568108676 |
| 1442 | - | 1532483 | 1533280 | + | - | *PA14_17850* | enoyl-CoA hydratase | 2 | 1 | 2 | 1 | 1 | 1 | 1 |
| 1443 | - | 1533277 | 1534809 | + | - | *PA14_17860* | 3-hydroxyacyl-CoA dehydrogenase | 10 | 10 | 8 | 10 | 8 | 8 | 1 |
| 1444 | - | 1534832 | 1536037 | + | - | *PA14_17880* | acetyl-CoA acetyltransferase | 6 | 6 | 7 | 6 | 5 | 4 | 1 |
| 1445 | - | 1536090 | 1537340 | + | - | *PA14_17890* | porin | 10 | 11 | 9 | 10 | 12 | 11 | 1 |
| 1446 | - | 1538266 | 1537346 | - | *metR* | *PA14_17900* | transcriptional regulator MetR | 89 | 61 | 81 | 59 | 93 | 113 | 1 |
| 1447 | - | 1538565 | 1539551 | + | - | *PA14_17910* | alpha/beta hydrolase | 9 | 10 | 10 | 10 | 7 | 9 | 1 |
| 1448 | - | 1539897 | 1539568 | - | *glpM* | *PA14_17920* | membrane protein GlpM | 15 | 16 | 11 | 9 | 9 | 12 | 1 |
| 1449 | - | 1541568 | 1540030 | - | *glpD* | *PA14_17930* | glycerol-3-phosphate dehydrogenase | 19 | 41 | 38 | 84 | 196 | 183 | 0.275790074 |
| 1450 | - | 1542602 | 1541847 | - | *glpR* | *PA14_17940* | glycerol-3-phosphate regulon repressor | 32 | 31 | 30 | 33 | 43 | 39 | 1 |
| 1451 | - | 1544325 | 1542808 | - | *glpK* | *PA14_17960* | glycerol kinase | 35 | 65 | 48 | 72 | 154 | 109 | 0.654227233 |
| 1452 | - | 1545204 | 1544365 | - | *glpF* | *PA14_17980* | glycerol uptake facilitator protein | 56 | 111 | 99 | 151 | 252 | 184 | 0.447575716 |
| 1453 | - | 1545469 | 1545939 | + | - | *PA14_17990* | hypothetical protein | 22 | 19 | 20 | 21 | 20 | 22 | 1 |
| 1454 | - | 1546003 | 1547487 | + | - | *PA14_18010* | glycerol kinase | 23 | 27 | 21 | 27 | 34 | 23 | 1 |
| 1455 | - | 1547683 | 1548468 | + | - | *PA14_18020* | PhzF family phenazine biosynthesis protein | 89 | 90 | 69 | 81 | 100 | 109 | 1 |
| 1456 | - | 1549283 | 1549459 | + | - | *PA14_18040* | hypothetical protein | 180 | 183 | 177 | 196 | 148 | 170 | 1 |
| 1457 | - | 1549993 | 1549571 | - | - | *PA14_18050* | hypothetical protein | 177 | 159 | 203 | 198 | 177 | 181 | 1 |
| 1458 | - | 1550622 | 1550092 | - | - | *PA14_18060* | hypothetical protein | 36 | 48 | 40 | 29 | 22 | 11 | 1 |
| 1459 | - | 1550805 | 1551002 | + | - | *PA14_18070* | periplasmic metal-binding protein | 158 | 312 | 151 | 257 | 222 | 167 | 0.410206905 |
| 1460 | - | 1551619 | 1550981 | - | - | *PA14_18080* | TetR family transcriptional regulator | 76 | 111 | 80 | 101 | 91 | 75 | 1 |
| 1461 | - | 1551985 | 1553163 | + | - | *PA14_18090* | major facilitator subfamily transporter protein | 68 | 73 | 116 | 48 | 47 | 38 | 1 |
| 1462 | - | 1553347 | 1553171 | - | - | *PA14_18100* | hypothetical protein | 93 | 113 | 149 | 72 | 59 | 58 | 1 |
| 1463 | - | 1554397 | 1553474 | - | *mmsR* | *PA14_18110* | transcriptional regulator MmsR | 22 | 18 | 20 | 21 | 17 | 17 | 1 |
| 1464 | - | 1554529 | 1556022 | + | *mmsA* | *PA14_18120* | methylmalonate-semialdehyde dehydrogenase | 160 | 254 | 258 | 506 | 249 | 413 | 1 |
| 1465 | - | 1556038 | 1556934 | + | *mmsB* | *PA14_18140* | 3-hydroxyisobutyrate dehydrogenase | 78 | 113 | 107 | 243 | 116 | 195 | 1 |
| 1466 | - | 1557036 | 1558922 | + | - | *PA14_18150* | acetyl-coa synthetase | 41 | 54 | 43 | 91 | 50 | 76 | 1 |
| 1467 | - | 1559993 | 1558980 | - | - | *PA14_18160* | oxidoreductase | 32 | 52 | 28 | 31 | 42 | 35 | 0.885197681 |
| 1468 | - | 1560308 | 1560015 | - | - | *PA14_18180* | hypothetical protein | 30 | 46 | 31 | 19 | 48 | 44 | 1 |
| 1469 | - | 1560404 | 1561324 | + | - | *PA14_18200* | LysR family transcriptional regulator | 25 | 24 | 21 | 18 | 21 | 19 | 1 |
| 1470 | - | 1562289 | 1561612 | - | - | *PA14_18210* | TatD family deoxyribonuclease | 33 | 34 | 34 | 31 | 32 | 28 | 1 |
| 1471 | - | 1563391 | 1562402 | - | *fruR* | *PA14_18230* | DNA-binding transcriptional regulator FruR | 39 | 36 | 40 | 32 | 39 | 31 | 1 |
| 1472 | - | 1563719 | 1566589 | + | - | *PA14_18250* | phosphotransferase system enzyme I | 16 | 52 | 42 | 23 | 50 | 32 | 0.007467257 |
| 1473 | - | 1566589 | 1567533 | + | *fruK* | *PA14_18260* | 1-phosphofructokinase | 20 | 66 | 49 | 26 | 63 | 44 | 0.001044072 |
| 1474 | - | 1567535 | 1569292 | + | *fruA* | *PA14_18275* | phosphotransferase system, fructose-specific IIBC component | 25 | 68 | 59 | 30 | 67 | 48 | 0.044138282 |
| 1475 | - | 1570839 | 1569445 | - | - | *PA14_18300* | nucleotide sugar dehydrogenase | 24 | 47 | 41 | 47 | 47 | 54 | 0.41668339 |
| 1476 | - | 1571249 | 1570836 | - | - | *PA14_18310* | hypothetical protein | 12 | 24 | 23 | 34 | 31 | 32 | 0.366758507 |
| 1477 | - | 1571593 | 1571246 | - | - | *PA14_18320* | hypothetical protein | 16 | 20 | 20 | 24 | 23 | 26 | 1 |
| 1478 | - | 1573239 | 1571590 | - | *arnT* | *PA14_18330* | 4-amino-4-deoxy-L-arabinose transferase | 8 | 11 | 12 | 16 | 14 | 16 | 1 |
| 1479 | - | 1574123 | 1573236 | - | - | *PA14_18340* | hypothetical protein | 12 | 21 | 19 | 21 | 20 | 21 | 1 |
| 1480 | - | 1576108 | 1574120 | - | - | *PA14_18350* | bifunctional UDP-glucuronic acid decarboxylase/UDP-4-amino-4-deoxy-L-arabinose formyltransferase | 20 | 31 | 32 | 34 | 32 | 40 | 1 |
| 1481 | - | 1577124 | 1576105 | - | - | *PA14_18360* | glycosyl transferase family protein | 14 | 22 | 23 | 24 | 22 | 26 | 1 |
| 1482 | - | 1578269 | 1577121 | - | - | *PA14_18370* | UDP-4-amino-4-deoxy-L-arabinose--oxoglutarate aminotransferase | 10 | 12 | 14 | 13 | 13 | 16 | 1 |
| 1483 | - | 1580102 | 1578657 | - | *algA* | *PA14_18380* | mannose-1-phosphate guanylyltransferase | 11 | 11 | 11 | 11 | 12 | 12 | 1 |
| 1484 | - | 1580949 | 1580299 | - | *algF* | *PA14_18410* | alginate o-acetyltransferase AlgF | 7 | 8 | 7 | 6 | 5 | 6 | 1 |
| 1485 | - | 1582197 | 1581022 | - | *algJ* | *PA14_18430* | alginate o-acetyltransferase AlgJ | 3 | 4 | 5 | 3 | 4 | 3 | 1 |
| 1486 | - | 1583774 | 1582212 | - | *algI* | *PA14_18450* | alginate o-acetyltransferase AlgI | 8 | 7 | 8 | 8 | 6 | 7 | 1 |
| 1487 | - | 1585119 | 1584016 | - | *algL* | *PA14_18470* | poly(beta-D-mannuronate) lyase | 5 | 5 | 5 | 5 | 6 | 6 | 1 |
| 1488 | - | 1586547 | 1585123 | - | *algX* | *PA14_18480* | alginate biosynthesis protein AlgX | 4 | 4 | 4 | 4 | 3 | 4 | 1 |
| 1489 | - | 1588191 | 1586560 | - | *algG* | *PA14_18500* | alginate-c5-mannuronan-epimerase AlgG | 68 | 67 | 67 | 64 | 58 | 61 | 1 |
| 1490 | - | 1589684 | 1588212 | - | *algE* | *PA14_18510* | alginate production outer membrane protein AlgE | 7 | 7 | 8 | 7 | 6 | 5 | 1 |
| 1491 | - | 1591108 | 1589681 | - | *algK* | *PA14_18520* | alginate biosynthetic protein AlgK | 8 | 8 | 10 | 9 | 7 | 6 | 1 |
| 1492 | - | 1592291 | 1591122 | - | *alg44* | *PA14_18550* | alginate biosynthesis protein Alg44 | 6 | 6 | 6 | 6 | 6 | 5 | 1 |
| 1493 | - | 1593867 | 1592374 | - | *alg8* | *PA14_18565* | alginate biosynthesis protein Alg8 | 17 | 14 | 17 | 16 | 15 | 17 | 1 |
| 1494 | - | 1595308 | 1593998 | - | *algD* | *PA14_18580* | GDP-mannose 6-dehydrogenase AlgD | 8 | 8 | 9 | 8 | 7 | 6 | 1 |
| 1495 | - | 1596990 | 1596211 | - | - | *PA14_18590* | hypothetical protein | 34 | 16 | 14 | 10 | 14 | 18 | 0.210107695 |
| 1496 | - | 1598179 | 1597097 | - | - | *PA14_18600* | ABC transporter ATP-binding protein | 53 | 47 | 49 | 36 | 49 | 44 | 1 |
| 1497 | - | 1599101 | 1598184 | - | *argF* | *PA14_18610* | ornithine carbamoyltransferase | 32 | 27 | 35 | 25 | 29 | 31 | 1 |
| 1498 | - | 1599380 | 1599847 | + | - | *PA14_18620* | hypothetical protein | 164 | 151 | 158 | 183 | 132 | 140 | 1 |
| 1499 | - | 1603228 | 1600241 | - | - | *PA14_18630* | serine protease | 48 | 70 | 55 | 64 | 62 | 64 | 1 |
| 1500 | - | 1603527 | 1605635 | + | - | *PA14_18640* | oxidoreductase | 14 | 14 | 13 | 16 | 15 | 16 | 1 |
| 1501 | - | 1605748 | 1606074 | + | - | *PA14_18650* | hypothetical protein | 198 | 283 | 130 | 156 | 291 | 263 | 1 |
| 1502 | - | 1606157 | 1607308 | + | - | *PA14_18660* | hypothetical protein | 46 | 36 | 41 | 38 | 46 | 47 | 1 |
| 1503 | - | 1607852 | 1607376 | - | *bfrB* | *PA14_18670* | bacterioferritin | 723 | 395 | 988 | 356 | 162 | 538 | 1 |
| 1504 | - | 1608283 | 1608062 | - | - | *PA14_18680* | hypothetical protein | 56 | 136 | 42 | 132 | 86 | 60 | 0.070160504 |
| 1505 | - | 1608543 | 1609145 | + | - | *PA14_18690* | peroxidase | 891 | 1221 | 725 | 549 | 837 | 607 | 1 |
| 1506 | - | 1609998 | 1609324 | - | *rnt* | *PA14_18700* | ribonuclease T | 92 | 139 | 111 | 125 | 134 | 101 | 1 |
| 1507 | - | 1611041 | 1609995 | - | *pyrC* | *PA14_18710* | dihydroorotase | 61 | 66 | 71 | 51 | 54 | 43 | 1 |
| 1508 | - | 1611174 | 1612139 | + | - | *PA14_18720* | OmpA family membrane protein | 96 | 87 | 114 | 84 | 103 | 101 | 1 |
| 1509 | - | 1612257 | 1613474 | + | *argG* | *PA14_18740* | argininosuccinate synthase | 83 | 70 | 86 | 61 | 65 | 59 | 1 |
| 1510 | - | 1613952 | 1613566 | - | *gloA1* | *PA14_18750* | lactoylglutathione lyase | 70 | 97 | 78 | 47 | 58 | 56 | 1 |
| 1511 | - | 1614329 | 1615486 | + | - | *PA14_18760* | RND efflux membrane fusion protein | 6 | 5 | 5 | 5 | 5 | 4 | 1 |
| 1512 | - | 1615483 | 1618644 | + | - | *PA14_18780* | RND efflux transporter | 8 | 8 | 6 | 7 | 7 | 6 | 1 |

| | A | B | C | D | E | F | G | H | I | J | K | L | M | N |
|---|---|---|---|---|---|---|---|---|---|---|---|---|---|---|
| 1513 | - | 1618641 | 1620116 | + | - | *PA14_18790* | outer membrane efflux protein | 5 | 5 | 4 | 5 | 5 | 5 | 1 |
| 1514 | 1620628 | 1620687 | 1620881 | + | - | *PA14_18800* | hypothetical protein | 35 | 8 | 9 | 32 | 27 | 57 | 3.95499E-10 |
| 1515 | 1621028 | 1621282 | 1622319 | + | - | *PA14_18810* | hypothetical protein | 4 | 7 | 3 | 3 | 6 | 5 | 0.630688527 |
| 1516 | 1621028 | 1622352 | 1623338 | + | - | *PA14_18820* | hypothetical protein | 3 | 6 | 2 | 5 | 5 | 4 | 1 |
| 1517 | 1621028 | 1623369 | 1624802 | + | - | *PA14_18830* | adenylosuccinate lyase | 11 | 11 | 15 | 15 | 10 | 13 | 1 |
| 1518 | 1621028 | 1624802 | 1626253 | + | - | *PA14_18850* | adenylosuccinate lyase | 7 | 7 | 6 | 9 | 5 | 6 | 1 |
| 1519 | 1621028 | 1626303 | 1627376 | + | - | *PA14_18860* | hypothetical protein | 28 | 33 | 27 | 35 | 27 | 28 | 1 |
| 1520 | - | 1627755 | 1627576 | - | - | *PA14_18870* | hypothetical protein | 80 | 132 | 70 | 67 | 43 | 53 | 0.903714641 |
| 1521 | - | 1628502 | 1627864 | - | *nth* | *PA14_18880* | endonuclease III | 35 | 49 | 43 | 36 | 43 | 33 | 1 |
| 1522 | - | 1629215 | 1628499 | - | - | *PA14_18890* | electron transport complex protein RsxE | 42 | 39 | 44 | 45 | 45 | 38 | 1 |
| 1523 | - | 1629852 | 1629208 | - | - | *PA14_18900* | hypothetical protein | 44 | 43 | 52 | 56 | 55 | 47 | 1 |
| 1524 | - | 1630886 | 1629852 | - | - | *PA14_18910* | hypothetical protein | 31 | 36 | 33 | 40 | 38 | 34 | 1 |
| 1525 | - | 1633213 | 1630889 | - | - | *PA14_18920* | electron transport complex protein RnfC | 42 | 47 | 46 | 47 | 45 | 37 | 1 |
| 1526 | - | 1633776 | 1633210 | - | - | *PA14_18930* | electron transport complex protein RnfB | 44 | 38 | 49 | 50 | 40 | 35 | 1 |
| 1527 | - | 1634357 | 1633773 | - | - | *PA14_18950* | Na(+)-translocating NADH-quinone reductase subunit E | 27 | 22 | 28 | 24 | 20 | 20 | 1 |
| 1528 | - | 1635233 | 1634535 | - | - | *PA14_18960* | hypothetical protein | 44 | 42 | 53 | 64 | 53 | 50 | 1 |
| 1529 | - | 1638973 | 1635662 | - | *pldA* | *PA14_18970* | phospholipase D | 34 | 28 | 24 | 28 | 32 | 27 | 1 |
| 1530 | - | 1641403 | 1638977 | - | - | *PA14_18985* | hypothetical protein | 20 | 19 | 15 | 18 | 22 | 15 | 1 |
| 1531 | - | 1642135 | 1641698 | - | - | *PA14_19010* | hypothetical protein | 26 | 22 | 20 | 18 | 17 | 16 | 1 |
| 1532 | - | 1643358 | 1642132 | - | - | *PA14_19020* | hypothetical protein | 31 | 27 | 23 | 24 | 29 | 23 | 1 |
| 1533 | - | 1644245 | 1643445 | - | - | *PA14_19030* | hypothetical protein | 66 | 62 | 65 | 51 | 56 | 40 | 1 |
| 1534 | - | 1646315 | 1644279 | - | *metG* | *PA14_19050* | methionyl-tRNA synthetase | 105 | 115 | 108 | 94 | 103 | 90 | 1 |
| 1535 | - | 1646446 | 1647540 | + | - | *PA14_19065* | hypothetical protein | 64 | 66 | 59 | 49 | 66 | 50 | 1 |
| 1536 | - | 1647642 | 1648208 | + | *dcd* | *PA14_19090* | deoxycytidine triphosphate deaminase | 58 | 46 | 49 | 46 | 43 | 42 | 1 |
| 1537 | 1648391 | 1648632 | 1649519 | + | *rhlA* | *PA14_19100* | rhamnosyltransferase chain A | 452 | 9 | 87 | 436 | 240 | 550 | 0 |
| 1538 | 1648391 | 1649585 | 1650865 | + | *rhlB* | *PA14_19110* | rhamnosyltransferase chain B | 217 | 18 | 37 | 255 | 124 | 301 | 3.12603E-64 |
| 1539 | - | 1650990 | 1651715 | + | *rhlR* | *PA14_19120* | transcriptional regulator RhlR | 707 | 1 | 615 | 994 | 731 | 898 | 0 |
| 1540 | 1651804 | 1651896 | 1652501 | + | *rhlI* | *PA14_19130* | autoinducer synthesis protein RhlI | 210 | 142 | 5 | 206 | 191 | 193 | 1 |
| 1541 | - | 1652656 | 1653462 | + | *pheC* | *PA14_19140* | cyclohexadienyl dehydratase | 45 | 39 | 141 | 48 | 40 | 42 | 1 |
| 1542 | - | 1653556 | 1654416 | + | - | *PA14_19150* | hypothetical protein | 10 | 12 | 25 | 11 | 13 | 10 | 1 |
| 1543 | - | 1654413 | 1655312 | + | - | *PA14_19160* | hypothetical protein | 18 | 26 | 30 | 23 | 22 | 17 | 1 |
| 1544 | - | 1655929 | 1655333 | - | - | *PA14_19170* | hypothetical protein | 47 | 61 | 68 | 49 | 43 | 42 | 1 |
| 1545 | - | 1656387 | 1658081 | + | - | *PA14_19190* | malate dehydrogenase | 93 | 137 | 93 | 85 | 111 | 71 | 1 |
| 1546 | - | 1658661 | 1658203 | - | - | *PA14_19205* | hypothetical protein | 47 | 47 | 47 | 42 | 44 | 35 | 1 |
| 1547 | - | 1659200 | 1658658 | - | - | *PA14_19210* | hypothetical protein | 34 | 39 | 32 | 27 | 36 | 24 | 1 |
| 1548 | - | 1660546 | 1659218 | - | - | *PA14_19230* | hypothetical protein | 26 | 32 | 26 | 22 | 29 | 20 | 1 |
| 1549 | - | 1662081 | 1660714 | - | - | *PA14_19270* | MFS transporter | 6 | 7 | 6 | 7 | 6 | 7 | 1 |
| 1550 | - | 1662267 | 1663607 | + | - | *PA14_19290* | ATP-dependent RNA helicase | 57 | 54 | 65 | 59 | 56 | 56 | 1 |
| 1551 | - | 1665444 | 1663729 | - | - | *PA14_19310* | hypothetical protein | 258 | 225 | 305 | 204 | 214 | 274 | 1 |
| 1552 | - | 1665661 | 1666986 | + | - | *PA14_19320* | hypothetical protein | 40 | 36 | 37 | 41 | 33 | 38 | 1 |
| 1553 | - | 1667345 | 1667115 | - | - | *PA14_19330* | hypothetical protein | 30 | 32 | 32 | 33 | 28 | 32 | 1 |
| 1554 | - | 1670152 | 1667396 | - | - | *PA14_19340* | sensor/response regulator hybrid | 22 | 21 | 25 | 28 | 21 | 29 | 1 |
| 1555 | - | 1671485 | 1670289 | - | - | *PA14_19350* | hypothetical protein | 67 | 100 | 91 | 114 | 125 | 130 | 1 |
| 1556 | - | 1673239 | 1671482 | - | - | *PA14_19360* | GNAT family acetyltransferase | 101 | 140 | 133 | 166 | 180 | 178 | 1 |
| 1557 | - | 1675049 | 1673280 | - | - | *PA14_19370* | asparagine synthetase | 262 | 240 | 278 | 202 | 263 | 278 | 1 |
| 1558 | - | 1675192 | 1675665 | + | - | *PA14_19380* | transcriptional regulator | 282 | 212 | 270 | 146 | 181 | 304 | 1 |
| 1559 | - | 1676673 | 1675921 | - | - | *PA14_19390* | hypothetical protein | 100 | 87 | 117 | 69 | 67 | 72 | 1 |
| 1560 | - | 1678738 | 1676774 | - | *mnmC* | *PA14_19400* | 5-methylaminomethyl-2-thiouridine methyltransferase | 38 | 32 | 32 | 34 | 29 | 25 | 1 |
| 1561 | - | 1678882 | 1680372 | + | - | *PA14_19410* | hypothetical protein | 100 | 101 | 97 | 93 | 96 | 98 | 1 |
| 1562 | - | 1680525 | 1681709 | + | - | *PA14_19430* | acyl-CoA thiolase | 26 | 24 | 21 | 20 | 29 | 27 | 1 |
| 1563 | - | 1681838 | 1682503 | + | - | *PA14_19450* | hypothetical protein | 19 | 21 | 18 | 14 | 18 | 13 | 1 |
| 1564 | - | 1684298 | 1682727 | - | *mqoA* | *PA14_19470* | malate:quinone oxidoreductase | 17 | 13 | 16 | 14 | 16 | 14 | 1 |
| 1565 | - | 1684710 | 1684501 | - | - | *PA14_19480* | hypothetical protein | 40 | 36 | 46 | 60 | 37 | 58 | 1 |
| 1566 | - | 1684916 | 1685554 | + | - | *PA14_19490* | antioxidant protein | 14 | 62 | 21 | 17 | 31 | 11 | 3.07409E-09 |
| 1567 | - | 1685805 | 1686809 | + | - | *PA14_19500* | hypothetical protein | 1 | 2 | 1 | 1 | 1 | 1 | 1 |
| 1568 | - | 1686813 | 1687637 | + | - | *PA14_19510* | ABC transporter permease | 3 | 5 | 3 | 4 | 3 | 3 | 1 |
| 1569 | - | 1687634 | 1688383 | + | - | *PA14_19520* | ABC transporter ATP-binding protein | 2 | 3 | 3 | 2 | 1 | 4 | 1 |
| 1570 | - | 1688537 | 1689130 | + | - | *PA14_19530* | NAD(P)H-dependent FMN reductase | 6 | 40 | 12 | 11 | 20 | 9 | 2.45612E-25 |
| 1571 | - | 1689249 | 1690220 | + | - | *PA14_19540* | hypothetical protein | 9 | 15 | 9 | 9 | 9 | 7 | 0.940015681 |
| 1572 | - | 1690298 | 1691446 | + | *ssuD* | *PA14_19560* | alkanesulfonate monooxygenase | 2 | 3 | 1 | 2 | 2 | 1 | 1 |
| 1573 | - | 1691467 | 1692255 | + | - | *PA14_19570* | ABC transporter permease | 3 | 3 | 2 | 2 | 2 | 1 | 1 |
| 1574 | - | 1692252 | 1693076 | + | *ssuB* | *PA14_19580* | aliphatic sulfonates transport ATP-binding subunit | 7 | 8 | 7 | 6 | 6 | 5 | 1 |
| 1575 | - | 1693117 | 1693332 | + | - | *PA14_19590* | molybdopterin-binding protein | 14 | 19 | 9 | 11 | 12 | 11 | 1 |
| 1576 | - | 1693850 | 1693512 | - | - | *PA14_19600* | hypothetical protein | 60 | 75 | 59 | 50 | 59 | 43 | 1 |
| 1577 | - | 1694213 | 1693902 | - | - | *PA14_19610* | hypothetical protein | 52 | 68 | 70 | 48 | 57 | 59 | 1 |
| 1578 | - | 1694713 | 1694342 | - | *folX* | *PA14_19620* | D-erythro-7,8-dihydroneopterin triphosphate 2'-epimerase | 102 | 127 | 81 | 106 | 152 | 137 | 1 |
| 1579 | - | 1695275 | 1694715 | - | *folE1* | *PA14_19630* | GTP cyclohydrolase I | 120 | 116 | 101 | 107 | 132 | 123 | 1 |
| 1580 | - | 1695997 | 1695293 | - | - | *PA14_19640* | short chain dehydrogenase | 34 | 35 | 30 | 26 | 43 | 39 | 1 |
| 1581 | - | 1696665 | 1696108 | - | - | *PA14_19650* | hypothetical protein | 5 | 5 | 6 | 6 | 5 | 4 | 1 |
| 1582 | - | 1696945 | 1697397 | + | - | *PA14_19660* | flavodoxin | 75 | 99 | 106 | 93 | 87 | 72 | 1 |
| 1583 | - | 1698429 | 1697536 | - | - | *PA14_19670* | LysR family transcriptional regulator | 23 | 27 | 24 | 22 | 20 | 21 | 1 |
| 1584 | - | 1698525 | 1698914 | + | - | *PA14_19680* | hypothetical protein | 27 | 25 | 34 | 38 | 19 | 40 | 1 |
| 1585 | - | 1698901 | 1699587 | + | - | *PA14_19690* | hypothetical protein | 33 | 28 | 41 | 44 | 25 | 47 | 1 |
| 1586 | - | 1699716 | 1700495 | + | - | *PA14_19700* | aldolase | 43 | 45 | 49 | 43 | 50 | 59 | 1 |
| 1587 | - | 1700492 | 1701388 | + | - | *PA14_19710* | alpha/beta hydrolase | 23 | 20 | 19 | 17 | 21 | 26 | 1 |
| 1588 | - | 1701487 | 1701780 | + | - | *PA14_19720* | hypothetical protein | 16 | 13 | 21 | 18 | 13 | 20 | 1 |
| 1589 | - | 1701896 | 1702807 | + | - | *PA14_19730* | oxidoreductase | 25 | 28 | 31 | 29 | 43 | 38 | 1 |
| 1590 | - | 1702924 | 1703694 | + | - | *PA14_19740* | enoyl-CoA hydratase | 135 | 130 | 163 | 176 | 138 | 147 | 1 |
| 1591 | - | 1704058 | 1703714 | - | - | *PA14_19750* | hypothetical protein | 58 | 67 | 75 | 91 | 69 | 72 | 1 |
| 1592 | - | 1705499 | 1704093 | - | - | *PA14_19770* | hypothetical protein | 16 | 18 | 18 | 20 | 16 | 22 | 1 |
| 1593 | - | 1706333 | 1705590 | - | - | *PA14_19800* | AraC family transcriptional regulator | 27 | 24 | 28 | 24 | 23 | 23 | 1 |
| 1594 | - | 1706618 | 1708375 | + | - | *PA14_19810* | hypothetical protein | 15 | 14 | 18 | 19 | 14 | 17 | 1 |
| 1595 | - | 1708432 | 1709802 | + | - | *PA14_19830* | hypothetical protein | 13 | 13 | 15 | 13 | 10 | 14 | 1 |
| 1596 | - | 1709952 | 1712435 | + | - | *PA14_19850* | transcriptional regulator | 9 | 8 | 8 | 9 | 9 | 11 | 1 |

| | A | B | C | D | E | F | G | H | I | J | K | L | M | N |
|---|---|---|---|---|---|---|---|---|---|---|---|---|---|---|
| 1597 | - | 1713254 | 1712442 | - | - | PA14_19860 | hypothetical protein | 73 | 72 | 78 | 86 | 69 | 64 | 1 |
| 1598 | - | 1713500 | 1714525 | + | ldh | PA14_19870 | leucine dehydrogenase | 278 | 334 | 374 | 314 | 330 | 352 | 1 |
| 1599 | - | 1714667 | 1715764 | + | - | PA14_19900 | pyruvate dehydrogenase E1 component subunit alpha | 136 | 173 | 256 | 220 | 190 | 238 | 1 |
| 1600 | - | 1715757 | 1716758 | + | - | PA14_19910 | pyruvate dehydrogenase E1 component, beta chain | 129 | 164 | 218 | 230 | 202 | 268 | 1 |
| 1601 | - | 1716776 | 1717888 | + | - | PA14_19920 | branched-chain alpha-keto acid dehydrogenase subunit E2 | 56 | 81 | 96 | 106 | 83 | 126 | 1 |
| 1602 | - | 1718347 | 1717889 | - | - | PA14_19930 | hypothetical protein | 23 | 40 | 40 | 53 | 43 | 60 | 0.882653101 |
| 1603 | - | 1718965 | 1718381 | - | - | PA14_19940 | hypothetical protein | 61 | 83 | 90 | 93 | 88 | 92 | 1 |
| 1604 | - | 1719267 | 1719001 | - | - | PA14_19950 | hypothetical protein | 271 | 325 | 326 | 380 | 354 | 338 | 1 |
| 1605 | - | 1719477 | 1719746 | + | - | PA14_19960 | hypothetical protein | 15 | 17 | 12 | 27 | 13 | 14 | 1 |
| 1606 | - | 1719866 | 1720084 | + | - | PA14_19970 | hypothetical protein | 4 | 13 | 3 | 13 | 8 | 2 | 9.20065E-07 |
| 1607 | - | 1720161 | 1720676 | + | - | PA14_19990 | RNA polymerase ECF-subfamily sigma-70 factor | 5 | 9 | 4 | 19 | 10 | 5 | 1 |
| 1608 | - | 1720741 | 1721727 | + | - | PA14_20000 | transmembrane sensor | 12 | 12 | 13 | 21 | 12 | 13 | 1 |
| 1609 | - | 1721937 | 1724612 | + | hasR | PA14_20010 | heme uptake outer membrane receptor HasR | 8 | 9 | 8 | 11 | 8 | 8 | 1 |
| 1610 | - | 1724698 | 1725315 | + | hasAp | PA14_20020 | heme acquisition protein HasAp | 46 | 43 | 53 | 51 | 34 | 39 | 1 |
| 1611 | - | 1725533 | 1727323 | + | hasD | PA14_20030 | transport protein HasD | 3 | 2 | 3 | 2 | 1 | 2 | 1 |
| 1612 | - | 1727320 | 1728651 | + | hasE | PA14_20040 | metalloprotease secretion protein | 3 | 2 | 4 | 2 | 2 | 2 | 0.925024982 |
| 1613 | - | 1728648 | 1730003 | + | - | PA14_20050 | outer membrane protein | 6 | 6 | 7 | 7 | 7 | 7 | 1 |
| 1614 | - | 1730283 | 1729996 | - | - | PA14_20060 | hypothetical protein | 21 | 19 | 26 | 22 | 19 | 18 | 1 |
| 1615 | - | 1730458 | 1730943 | + | - | PA14_20070 | hypothetical protein | 14 | 14 | 13 | 15 | 13 | 12 | 1 |
| 1616 | - | 1731056 | 1732027 | + | - | PA14_20080 | hypothetical protein | 20 | 18 | 19 | 16 | 15 | 13 | 1 |
| 1617 | - | 1732031 | 1733203 | + | - | PA14_20100 | hypothetical protein | 27 | 27 | 25 | 26 | 23 | 23 | 1 |
| 1618 | - | 1733200 | 1734333 | + | - | PA14_20110 | hypothetical protein | 25 | 25 | 23 | 27 | 21 | 22 | 1 |
| 1619 | - | 1734387 | 1734728 | + | - | PA14_20120 | hypothetical protein | 129 | 145 | 125 | 148 | 135 | 136 | 1 |
| 1620 | 1735823 | 1735656 | 1734730 | - | - | PA14_20130 | LysR family transcriptional regulator | 61 | 64 | 63 | 64 | 64 | 60 | 1 |
| 1621 | 1735823 | 1735930 | 1736706 | + | fpr | PA14_20140 | ferredoxin--NADP+ reductase | 215 | 176 | 193 | 176 | 215 | 232 | 1 |
| 1622 | - | 1737325 | 1736789 | - | nosL | PA14_20150 | NosL protein | 26 | 53 | 75 | 80 | 120 | 102 | 0.285762132 |
| 1623 | - | 1738169 | 1737342 | - | nosY | PA14_20170 | NosY protein | 8 | 32 | 38 | 51 | 70 | 61 | 4.22828E-06 |
| 1624 | - | 1739074 | 1738160 | - | nosF | PA14_20180 | NosF protein | 18 | 51 | 62 | 68 | 93 | 84 | 0.010969112 |
| 1625 | - | 1740357 | 1739071 | - | nosD | PA14_20190 | copper ABC transporter periplasmic substrate-binding protein | 36 | 103 | 118 | 122 | 171 | 178 | 0.024246819 |
| 1626 | - | 1742264 | 1740354 | - | nosZ | PA14_20200 | nitrous-oxide reductase | 49 | 214 | 147 | 188 | 254 | 264 | 0.000219048 |
| 1627 | - | 1744455 | 1742308 | - | nosR | PA14_20230 | regulatory protein NosR | 60 | 88 | 63 | 164 | 237 | 190 | 1 |
| 1628 | - | 1745097 | 1744684 | - | - | PA14_20240 | ring-cleaving dioxygenase | 4 | 4 | 4 | 4 | 4 | 3 | 1 |
| 1629 | - | 1745158 | 1745460 | + | - | PA14_20250 | hypothetical protein | 6 | 7 | 9 | 8 | 9 | 7 | 1 |
| 1630 | - | 1746162 | 1745467 | - | - | PA14_20260 | hypothetical protein | 8 | 9 | 9 | 9 | 8 | 9 | 1 |
| 1631 | - | 1746985 | 1746215 | - | rhlG | PA14_20270 | beta-ketoacyl reductase | 6 | 7 | 8 | 8 | 7 | 7 | 1 |
| 1632 | - | 1747448 | 1747083 | - | - | PA14_20280 | hypothetical protein | 15 | 25 | 20 | 28 | 31 | 25 | 0.844537532 |
| 1633 | - | 1747928 | 1747602 | - | - | PA14_20290 | DNA binding-protein | 581 | 839 | 601 | 777 | 982 | 897 | 1 |
| 1634 | - | 1748289 | 1749125 | + | phnC | PA14_20300 | ABC phosphonate transporter ATP-binding protein | 15 | 11 | 14 | 15 | 13 | 13 | 1 |
| 1635 | - | 1749178 | 1750182 | + | phnD | PA14_20320 | phosphonate ABC transporter substrate-binding protein | 4 | 3 | 3 | 4 | 3 | 3 | 1 |
| 1636 | - | 1750250 | 1751044 | + | phnE | PA14_20330 | phosphonate ABC transporter permease | 2 | 3 | 4 | 3 | 2 | 2 | 1 |
| 1637 | - | 1751065 | 1751787 | + | phnF | PA14_20350 | transcriptional regulator | 5 | 6 | 6 | 6 | 6 | 6 | 1 |
| 1638 | - | 1751797 | 1752255 | + | phnG | PA14_20360 | phosphonate metabolism protein PhnG | 42 | 31 | 42 | 37 | 30 | 33 | 1 |
| 1639 | - | 1752255 | 1752884 | + | phnH | PA14_20370 | carbon-phosphorus lyase complex subunit | 28 | 24 | 26 | 22 | 15 | 19 | 1 |
| 1640 | - | 1752884 | 1753984 | + | phnI | PA14_20380 | hypothetical protein | 9 | 9 | 10 | 9 | 9 | 10 | 1 |
| 1641 | - | 1753981 | 1754865 | + | phnJ | PA14_20390 | hypothetical protein | 8 | 8 | 10 | 11 | 8 | 9 | 1 |
| 1642 | - | 1754862 | 1755677 | + | phnK | PA14_20400 | phosphonate C-P lyase system protein PhnK | 8 | 8 | 8 | 9 | 8 | 10 | 1 |
| 1643 | - | 1755728 | 1756444 | + | phnL | PA14_20420 | phosphonate ABC transporter ATPase | 5 | 6 | 7 | 5 | 6 | 6 | 1 |
| 1644 | - | 1756434 | 1757597 | + | phnM | PA14_20430 | phosphonate metabolism protein | 19 | 19 | 22 | 29 | 19 | 23 | 1 |
| 1645 | - | 1757597 | 1758154 | + | phnN | PA14_20440 | phosphonate transport ATP-binding protein | 96 | 92 | 107 | 121 | 102 | 108 | 1 |
| 1646 | - | 1758145 | 1758915 | + | phnP | PA14_20450 | carbon-phosphorus lyase complex accessory protein | 49 | 36 | 47 | 47 | 46 | 48 | 1 |
| 1647 | - | 1759139 | 1758954 | - | - | PA14_20460 | hypothetical protein | 180 | 105 | 140 | 139 | 131 | 120 | 0.87182855 |
| 1648 | - | 1759337 | 1759191 | - | - | PA14_20470 | hypothetical protein | 182 | 114 | 135 | 139 | 124 | 133 | 1 |
| 1649 | - | 1759704 | 1759426 | - | - | PA14_20480 | hypothetical protein | 67 | 51 | 46 | 47 | 47 | 52 | 1 |
| 1650 | - | 1760415 | 1759897 | - | - | PA14_20491 | acetyltransferase | 17 | 15 | 16 | 16 | 14 | 17 | 1 |
| 1651 | - | 1760535 | 1760608 | + | - | PA14_20500 | Arg tRNA | 30 | 30 | 28 | 25 | 22 | 27 | 1 |
| 1652 | - | 1762508 | 1763482 | + | - | PA14_20510 | hypothetical protein | 29 | 38 | 27 | 56 | 45 | 26 | 1 |
| 1653 | - | 1763497 | 1766208 | + | - | PA14_20520 | hypothetical protein | 41 | 46 | 37 | 70 | 56 | 38 | 1 |
| 1654 | - | 1766195 | 1766671 | + | - | PA14_20530 | hypothetical protein | 27 | 40 | 25 | 61 | 55 | 25 | 1 |
| 1655 | - | 1768341 | 1768003 | - | - | PA14_20550 | hypothetical protein | 9 | 5 | 6 | 5 | 4 | 3 | 0.738547973 |
| 1656 | - | 1768608 | 1769648 | + | amiE | PA14_20560 | acylamide amidohydrolase | 78 | 103 | 85 | 86 | 66 | 72 | 1 |
| 1657 | - | 1769732 | 1770847 | + | - | PA14_20570 | chaperone | 35 | 49 | 43 | 47 | 33 | 38 | 1 |
| 1658 | - | 1770873 | 1772030 | + | amiC | PA14_20580 | aliphatic amidase expression-regulating protein | 46 | 51 | 48 | 59 | 41 | 45 | 1 |
| 1659 | - | 1772027 | 1772617 | + | amiR | PA14_20590 | aliphatic amidase regulator | 60 | 71 | 61 | 82 | 62 | 79 | 1 |
| 1660 | - | 1773139 | 1773813 | + | - | PA14_20600 | hypothetical protein | 74 | 32 | 43 | 42 | 30 | 61 | 0.061837823 |
| 1661 | 1774336 | 1774207 | 1773860 | - | lecB | PA14_20610 | fucose-binding lectin PA-IIL | 393 | 69 | 176 | 137 | 96 | 449 | 7.47627E-12 |
| 1662 | 1774336 | 1774674 | 1775732 | + | - | PA14_20620 | HlyD family secretion protein | 17 | 12 | 18 | 16 | 12 | 14 | 1 |
| 1663 | 1774336 | 1775755 | 1776765 | + | - | PA14_20630 | hypothetical protein | 9 | 8 | 8 | 11 | 9 | 10 | 1 |
| 1664 | - | 1777648 | 1776770 | - | - | PA14_20640 | hypothetical protein | 10 | 11 | 8 | 9 | 10 | 7 | 1 |
| 1665 | - | 1779051 | 1777705 | - | dsdA | PA14_20650 | D-serine dehydratase | 33 | 34 | 31 | 34 | 31 | 36 | 1 |
| 1666 | - | 1779377 | 1780618 | + | - | PA14_20670 | glutamine synthetase | 152 | 122 | 108 | 104 | 118 | 126 | 1 |
| 1667 | - | 1780802 | 1782106 | + | - | PA14_20680 | MFS transporter | 44 | 41 | 40 | 47 | 38 | 44 | 1 |
| 1668 | - | 1782148 | 1782759 | + | - | PA14_20690 | hypothetical protein | 160 | 140 | 161 | 160 | 128 | 161 | 1 |
| 1669 | - | 1783554 | 1782763 | - | - | PA14_20700 | glycosyltransferase | 142 | 128 | 155 | 126 | 125 | 140 | 1 |
| 1670 | - | 1784053 | 1783583 | - | flgN | PA14_20720 | hypothetical protein | 294 | 322 | 333 | 230 | 313 | 288 | 1 |
| 1671 | - | 1784431 | 1784108 | - | flgM | PA14_20730 | hypothetical protein | 481 | 605 | 574 | 448 | 567 | 606 | 1 |
| 1672 | - | 1785274 | 1784576 | - | flgA | PA14_20740 | flagellar basal body P-ring biosynthesis protein FlgA | 77 | 61 | 79 | 78 | 62 | 60 | 1 |
| 1673 | - | 1785406 | 1786338 | + | - | PA14_20750 | chemotaxis protein | 963 | 897 | 1018 | 903 | 857 | 965 | 1 |
| 1674 | - | 1786415 | 1787239 | + | - | PA14_20760 | chemotaxis protein methyltransferase | 463 | 460 | 512 | 579 | 546 | 602 | 1 |
| 1675 | - | 1787554 | 1787859 | + | - | PA14_20770 | hypothetical protein | 279 | 381 | 413 | 478 | 440 | 499 | 1 |
| 1676 | - | 1787861 | 1789576 | + | - | PA14_20780 | two-component response regulator | 170 | 208 | 230 | 345 | 236 | 285 | 1 |
| 1677 | - | 1789642 | 1789992 | + | - | PA14_20800 | histidine phosphotransfer domain-containing protein | 106 | 91 | 115 | 142 | 101 | 124 | 1 |
| 1678 | - | 1790076 | 1792214 | + | recQ | PA14_20810 | ATP-dependent DNA helicase RecQ | 94 | 83 | 104 | 91 | 74 | 70 | 1 |
| 1679 | - | 1792303 | 1793472 | + | - | PA14_20820 | two-component response regulator | 63 | 78 | 74 | 88 | 75 | 73 | 1 |
| 1680 | - | 1793623 | 1794639 | + | - | PA14_20840 | hypothetical protein | 70 | 76 | 72 | 84 | 77 | 93 | 1 |

| | A | B | C | D | E | F | G | H | I | J | K | L | M | N |
|---|---|---|---|---|---|---|---|---|---|---|---|---|---|---|
| 1681 | - | 1794757 | 1795191 | + | - | *PA14_20850* | MarR family transcriptional regulator | 83 | 95 | 92 | 82 | 91 | 109 | 1 |
| 1682 | - | 1795468 | 1797513 | + | - | *PA14_20860* | Tfp pilus assembly protein FimV | 54 | 48 | 54 | 59 | 47 | 69 | 1 |
| 1683 | - | 1799707 | 1797521 | - | - | *PA14_20870* | hypothetical protein | 20 | 23 | 25 | 27 | 23 | 30 | 1 |
| 1684 | - | 1799776 | 1800066 | + | - | *PA14_20880* | hypothetical protein | 32 | 16 | 26 | 19 | 27 | 24 | 0.047727227 |
| 1685 | - | 1800156 | 1801154 | + | *rfaD* | *PA14_20890* | ADP-L-glycero-D-manno-heptose-6-epimerase | 336 | 338 | 439 | 109 | 151 | 314 | 1 |
| 1686 | - | 1802331 | 1801165 | - | - | *PA14_20900* | MFS transporter | 247 | 58 | 77 | 76 | 147 | 123 | 0.001792166 |
| 1687 | - | 1803175 | 1802423 | - | - | *PA14_20920* | hypothetical protein | 208 | 12 | 22 | 64 | 156 | 111 | 3.476E-252 |
| 1688 | 1816906 | 1803532 | 1803293 | - | - | *PA14_20940* | acyl carrier protein | 247 | 13 | 35 | 99 | 206 | 156 | 0 |
| 1689 | 1816906 | 1804538 | 1803546 | - | *fabH2* | *PA14_20950* | 3-oxoacyl-ACP synthase | 268 | 11 | 33 | 82 | 187 | 127 | 0 |
| 1690 | 1816906 | 1804967 | 1804542 | - | - | *PA14_20960* | isomerase | 195 | 11 | 20 | 68 | 179 | 105 | 0 |
| 1691 | 1816906 | 1806220 | 1804964 | - | *cyp23* | *PA14_20970* | cytochrome P450 | 274 | 17 | 39 | 107 | 260 | 159 | 7.8332E-152 |
| 1692 | 1816906 | 1807127 | 1806213 | - | - | *PA14_20980* | short chain dehydrogenas | 451 | 16 | 56 | 165 | 359 | 239 | 0 |
| 1693 | 1816906 | 1808457 | 1807129 | - | - | *PA14_21000* | hypothetical protein | 262 | 17 | 44 | 107 | 213 | 155 | 2.0973E-123 |
| 1694 | 1816906 | 1809620 | 1808454 | - | - | *PA14_21010* | FAD-dependent monooxygenase | 246 | 10 | 35 | 90 | 185 | 127 | 0 |
| 1695 | 1816906 | 1816675 | 1809617 | - | - | *PA14_21020* | non-ribosomal peptide synthetase | 178 | 15 | 33 | 76 | 136 | 89 | 2.36197E-32 |
| 1696 | 1816906 | 1817157 | 1817762 | + | - | *PA14_21030* | ATP-dependent Clp protease proteolytic subunit | 572 | 132 | 259 | 618 | 630 | 537 | 0.002328387 |
| 1697 | - | 1818994 | 1818107 | - | - | *PA14_21040* | hypothetical protein | 38 | 14 | 24 | 50 | 41 | 38 | 0.011702045 |
| 1698 | - | 1820782 | 1819004 | - | - | *PA14_21050* | short chain dehydrogenase | 16 | 16 | 13 | 19 | 15 | 18 | 1 |
| 1699 | - | 1821654 | 1820779 | - | - | *PA14_21060* | hypothetical protein | 12 | 12 | 12 | 13 | 13 | 14 | 1 |
| 1700 | - | 1822080 | 1822778 | + | - | *PA14_21070* | MerR family transcriptional regulator | 38 | 27 | 34 | 33 | 27 | 28 | 1 |
| 1701 | - | 1823700 | 1822786 | - | - | *PA14_21080* | LysR family transcriptional activator | 16 | 15 | 14 | 17 | 12 | 16 | 1 |
| 1702 | - | 1823789 | 1824244 | + | - | *PA14_21090* | hypothetical protein | 4 | 4 | 4 | 5 | 5 | 7 | 1 |
| 1703 | - | 1824476 | 1826554 | + | *plcN* | *PA14_21110* | non-hemolytic phospholipase C | 6 | 6 | 6 | 7 | 5 | 5 | 1 |
| 1704 | - | 1827112 | 1826612 | - | - | *PA14_21120* | hypothetical protein | 11 | 10 | 10 | 9 | 10 | 10 | 1 |
| 1705 | - | 1827906 | 1827178 | - | - | *PA14_21130* | outer membrane lipoprotein | 25 | 18 | 22 | 20 | 21 | 17 | 1 |
| 1706 | - | 1828783 | 1828013 | - | - | *PA14_21140* | ABC transporter permease | 17 | 20 | 16 | 21 | 20 | 13 | 1 |
| 1707 | - | 1829613 | 1828780 | - | - | *PA14_21150* | ABC transporter permease | 12 | 16 | 14 | 12 | 15 | 11 | 1 |
| 1708 | - | 1830410 | 1829607 | - | - | *PA14_21160* | ABC transporter ATP-binding protein | 59 | 61 | 55 | 50 | 52 | 46 | 1 |
| 1709 | - | 1831413 | 1832303 | + | - | *PA14_21180* | 3-hydroxyisobutyrate dehydrogenase | 42 | 41 | 39 | 36 | 43 | 31 | 1 |
| 1710 | - | 1831414 | 1830407 | - | - | *PA14_21175* | hypothetical protein | 38 | 51 | 37 | 37 | 41 | 32 | 1 |
| 1711 | - | 1834664 | 1832307 | - | - | *PA14_21190* | hypothetical protein | 31 | 31 | 37 | 43 | 36 | 41 | 1 |
| 1712 | - | 1834819 | 1836474 | + | - | *PA14_21210* | hypothetical protein | 67 | 67 | 84 | 48 | 54 | 75 | 1 |
| 1713 | - | 1836998 | 1836543 | - | - | *PA14_21220* | hypothetical protein | 2414 | 2570 | 2646 | 1105 | 1165 | 2305 | 1 |
| 1714 | - | 1840133 | 1837281 | - | *hepA* | *PA14_21230* | ATP-dependent helicase HepA | 45 | 56 | 53 | 51 | 50 | 34 | 1 |
| 1715 | - | 1840369 | 1840677 | + | - | *PA14_21240* | hypothetical protein | 170 | 196 | 191 | 239 | 241 | 240 | 1 |
| 1716 | - | 1840731 | 1841333 | + | *alkB* | *PA14_21250* | hypothetical protein | 77 | 69 | 80 | 70 | 73 | 71 | 1 |
| 1717 | - | 1841740 | 1841853 | + | - | *PA14_21260* | hypothetical protein | 2674 | 1368 | 3463 | 1900 | 1433 | 1957 | 1 |
| 1718 | - | 1842061 | 1844055 | + | - | *PA14_21280* | hypothetical protein | 41 | 36 | 47 | 41 | 45 | 43 | 1 |
| 1719 | - | 1844273 | 1845130 | + | - | *PA14_21290* | hypothetical protein | 34 | 30 | 29 | 33 | 36 | 39 | 1 |
| 1720 | - | 1846286 | 1845111 | - | - | *PA14_21300* | MFS transporte | 12 | 13 | 13 | 13 | 13 | 16 | 1 |
| 1721 | - | 1846907 | 1846437 | - | *phaJ1* | *PA14_21310* | hypothetical protein | 53 | 84 | 52 | 56 | 79 | 53 | 1 |
| 1722 | - | 1847854 | 1846904 | - | - | *PA14_21320* | hypothetical protein | 26 | 25 | 26 | 20 | 22 | 19 | 1 |
| 1723 | - | 1848101 | 1849789 | + | *fadD2* | *PA14_21340* | long-chain-fatty-acid--CoA ligase | 64 | 79 | 66 | 108 | 85 | 84 | 1 |
| 1724 | - | 1850024 | 1851712 | + | *fadD1* | *PA14_21370* | long-chain-fatty-acid--CoA ligase | 99 | 118 | 93 | 90 | 93 | 78 | 1 |
| 1725 | - | 1851742 | 1852044 | + | - | *PA14_21380* | hypothetical protein | 17 | 18 | 22 | 37 | 20 | 24 | 1 |
| 1726 | - | 1852220 | 1856200 | + | *hrpA* | *PA14_21400* | ATP-dependent helicase | 61 | 56 | 60 | 57 | 55 | 52 | 1 |
| 1727 | - | 1857703 | 1856273 | - | *phoA* | *PA14_21410* | alkaline phosphatase | 7 | 7 | 8 | 7 | 6 | 5 | 1 |
| 1728 | - | 1857908 | 1858345 | + | - | *PA14_21440* | HIT family protein | 42 | 50 | 43 | 40 | 33 | 37 | 1 |
| 1729 | - | 1858574 | 1860640 | + | - | *PA14_21450* | hypothetical protein | 20 | 17 | 14 | 14 | 21 | 12 | 1 |
| 1730 | - | 1860650 | 1861465 | + | - | *PA14_21460* | hypothetical protein | 11 | 7 | 8 | 8 | 9 | 6 | 0.674979992 |
| 1731 | - | 1861405 | 1861983 | + | - | *PA14_21470* | hypothetical protein | 8 | 4 | 5 | 5 | 8 | 6 | 0.347673727 |
| 1732 | - | 1862506 | 1863060 | + | - | *PA14_21480* | hypothetical protein | 10 | 10 | 6 | 8 | 13 | 7 | 1 |
| 1733 | - | 1863075 | 1865723 | + | - | *PA14_21490* | hypothetical protein | 58 | 53 | 52 | 58 | 60 | 61 | 1 |
| 1734 | - | 1866175 | 1865762 | - | - | *PA14_21510* | hypothetical protein | 49 | 40 | 49 | 42 | 38 | 47 | 1 |
| 1735 | - | 1866255 | 1866758 | + | - | *PA14_21520* | hypothetical protein | 92 | 45 | 44 | 45 | 41 | 104 | 0.452603163 |
| 1736 | - | 1867277 | 1866762 | - | - | *PA14_21530* | ankyrin domain-containing protein | 179 | 31 | 34 | 34 | 28 | 168 | 6.87207E-17 |
| 1737 | - | 1867837 | 1868889 | + | - | *PA14_21540* | 3-oxoacyl-ACP synthase | 81 | 76 | 67 | 72 | 62 | 77 | 1 |
| 1738 | - | 1868892 | 1869503 | + | - | *PA14_21550* | RNA polymerase ECF-subfamily sigma-70 factor | 105 | 110 | 103 | 115 | 96 | 102 | 1 |
| 1739 | - | 1869497 | 1869736 | + | - | *PA14_21560* | hypothetical protein | 128 | 114 | 120 | 123 | 103 | 102 | 1 |
| 1740 | - | 1870142 | 1870456 | + | - | *PA14_21570* | hypothetical protein | 291 | 92 | 190 | 52 | 24 | 40 | 0.01350491 |
| 1741 | - | 1870475 | 1871329 | + | - | *PA14_21580* | hypothetical protein | 210 | 66 | 147 | 81 | 23 | 43 | 0.07173263 |
| 1742 | - | 1871326 | 1872060 | + | - | *PA14_21590* | hypothetical protein | 25 | 11 | 21 | 7 | 5 | 7 | 0.05276977 |
| 1743 | - | 1872063 | 1872653 | + | - | *PA14_21600* | hypothetical protein | 24 | 11 | 19 | 6 | 4 | 7 | 0.050479413 |
| 1744 | - | 1872900 | 1874216 | + | *oprO* | *PA14_21610* | pyrophosphate-specific outer membrane porin OprO precursor | 9 | 8 | 10 | 9 | 7 | 7 | 1 |
| 1745 | - | 1874675 | 1875997 | + | *oprP* | *PA14_21620* | phosphate-specific outer membrane porin OprP precursor | 3 | 3 | 3 | 5 | 4 | 3 | 1 |
| 1746 | - | 1876166 | 1876465 | + | - | *PA14_21630* | hypothetical protein | 564 | 549 | 995 | 416 | 366 | 698 | 1 |
| 1747 | - | 1876590 | 1877402 | + | - | *PA14_21640* | short chain dehydrogenase | 57 | 46 | 78 | 42 | 52 | 62 | 1 |
| 1748 | - | 1877635 | 1878057 | + | - | *PA14_21650* | hypothetical protein | 14 | 17 | 23 | 17 | 16 | 15 | 1 |
| 1749 | - | 1878054 | 1878383 | + | - | *PA14_21660* | hypothetical protein | 32 | 48 | 45 | 45 | 48 | 40 | 1 |
| 1750 | - | 1878700 | 1879299 | + | - | *PA14_21680* | hypothetical protein | 32 | 42 | 42 | 43 | 39 | 34 | 1 |
| 1751 | - | 1878749 | 1878462 | - | - | *PA14_21670* | hypothetical protein | 49 | 96 | 91 | 82 | 88 | 79 | 0.360825712 |
| 1752 | - | 1879368 | 1883714 | + | - | *PA14_21690* | ATP-dependent DNA helicase | 68 | 67 | 76 | 70 | 73 | 73 | 1 |
| 1753 | - | 1887201 | 1883722 | - | - | *PA14_21700* | two-component sensor | 108 | 70 | 92 | 71 | 66 | 68 | 1 |
| 1754 | - | 1887467 | 1888054 | + | - | *PA14_21710* | hypothetical protein | 123 | 89 | 119 | 96 | 85 | 81 | 1 |
| 1755 | - | 1888121 | 1888975 | + | - | *PA14_21720* | AraC family transcriptional regulator | 18 | 14 | 18 | 21 | 19 | 13 | 1 |
| 1756 | - | 1889072 | 1891237 | + | - | *PA14_21730* | TonB-dependent receptor | 10 | 15 | 17 | 16 | 15 | 8 | 1 |
| 1757 | - | 1891412 | 1893286 | + | - | *PA14_21750* | hypothetical protein | 41 | 33 | 37 | 35 | 36 | 30 | 1 |
| 1758 | - | 1893556 | 1893347 | - | *capB* | *PA14_21760* | cold acclimation protein B | 685 | 511 | 494 | 347 | 423 | 349 | 1 |
| 1759 | - | 1894085 | 1893771 | - | - | *PA14_21770* | DMT family permease | 29 | 26 | 32 | 32 | 22 | 28 | 1 |
| 1760 | - | 1895160 | 1894225 | - | - | *PA14_21780* | transporter | 15 | 17 | 16 | 18 | 14 | 15 | 1 |
| 1761 | - | 1896167 | 1895247 | - | *rdgC* | *PA14_21790* | recombination associated protein | 28 | 35 | 41 | 36 | 34 | 31 | 1 |
| 1762 | - | 1896332 | 1896404 | + | - | *PA14_21800* | Val tRNA | 151 | 90 | 168 | 141 | 86 | 58 | 0.382662007 |
| 1763 | - | 1896424 | 1896497 | + | - | *PA14_21810* | Asp tRNA | 184 | 119 | 182 | 155 | 107 | 71 | 0.853730745 |
| 1764 | - | 1896757 | 1897518 | + | - | *PA14_21820* | peptidyl-prolyl cis-trans isomerase, FkbP-type | 128 | 123 | 111 | 80 | 103 | 87 | 1 |

| | A | B | C | D | E | F | G | H | I | J | K | L | M | N |
|---|---|---|---|---|---|---|---|---|---|---|---|---|---|---|
| 1765 | - | 1897888 | 1898319 | + | - | PA14_21830 | hypothetical protein | 403 | 310 | 435 | 431 | 326 | 364 | 1 |
| 1766 | - | 1899053 | 1898307 | - | - | PA14_21840 | hypothetical protein | 171 | 166 | 202 | 179 | 136 | 158 | 1 |
| 1767 | - | 1899157 | 1899468 | + | - | PA14_21850 | transcriptional regulator | 200 | 173 | 217 | 183 | 160 | 190 | 1 |
| 1768 | - | 1899826 | 1900305 | + | - | PA14_21860 | hypothetical protein | 171 | 175 | 203 | 185 | 142 | 173 | 1 |
| 1769 | - | 1902124 | 1900319 | - | - | PA14_21870 | EAL domain/GGDEF domain-containing protein | 23 | 23 | 28 | 28 | 21 | 24 | 1 |
| 1770 | - | 1904259 | 1902163 | - | tsp | PA14_21880 | periplasmic tail-specific protease | 375 | 389 | 384 | 353 | 378 | 423 | 1 |
| 1771 | - | 1904389 | 1905351 | + | - | PA14_21890 | oxidoreductase | 263 | 245 | 276 | 264 | 262 | 266 | 1 |
| 1772 | - | 1905996 | 1905418 | - | - | PA14_21900 | HAD-superfamily hydrolase | 88 | 88 | 88 | 103 | 90 | 99 | 1 |
| 1773 | - | 1907063 | 1906068 | - | - | PA14_21910 | ABC transporter ATP-binding protein | 20 | 20 | 20 | 26 | 20 | 24 | 1 |
| 1774 | - | 1907863 | 1907066 | - | - | PA14_21920 | ABC transporter permease | 9 | 11 | 13 | 13 | 11 | 13 | 1 |
| 1775 | - | 1908689 | 1907850 | - | - | PA14_21930 | ABC transporter permease | 12 | 11 | 11 | 13 | 9 | 13 | 1 |
| 1776 | - | 1909495 | 1908686 | - | - | PA14_21940 | hypothetical protein | 15 | 16 | 16 | 18 | 12 | 16 | 1 |
| 1777 | - | 1910608 | 1909550 | - | - | PA14_21960 | hypothetical protein | 37 | 40 | 41 | 43 | 36 | 48 | 1 |
| 1778 | - | 1910972 | 1911688 | + | - | PA14_21970 | transcriptional regulator | 31 | 33 | 42 | 52 | 33 | 35 | 1 |
| 1779 | - | 1911754 | 1912305 | + | - | PA14_21980 | hypothetical protein | 57 | 55 | 58 | 56 | 54 | 46 | 1 |
| 1780 | - | 1913637 | 1912348 | - | - | PA14_21990 | aminopeptidase 2 | 61 | 59 | 57 | 49 | 50 | 52 | 1 |
| 1781 | - | 1914441 | 1913806 | - | rluA | PA14_22000 | pseudouridine synthase | 18 | 21 | 22 | 27 | 26 | 23 | 1 |
| 1782 | - | 1914799 | 1914545 | - | minE | PA14_22010 | cell division topological specificity factor MinE | 92 | 139 | 98 | 87 | 85 | 59 | 1 |
| 1783 | - | 1915611 | 1914796 | - | minD | PA14_22020 | cell division inhibitor MinD | 102 | 107 | 99 | 100 | 97 | 86 | 1 |
| 1784 | - | 1916464 | 1915673 | - | minC | PA14_22040 | septum formation inhibitor | 46 | 35 | 48 | 44 | 34 | 35 | 1 |
| 1785 | - | 1916624 | 1917562 | + | htrB | PA14_22050 | lipid A biosynthesis lauroyl acyltransferase | 21 | 18 | 25 | 21 | 16 | 16 | 1 |
| 1786 | - | 1917630 | 1918799 | + | - | PA14_22060 | hypothetical protein | 68 | 65 | 63 | 75 | 67 | 66 | 1 |
| 1787 | - | 1919495 | 1918863 | - | - | PA14_22075 | pirin-related protein | 144 | 154 | 167 | 151 | 126 | 118 | 1 |
| 1788 | - | 1922695 | 1922000 | - | - | PA14_22080 | resolvase | 147 | 115 | 102 | 75 | 99 | 97 | 1 |
| 1789 | 1924465 | 1924132 | 1923032 | - | - | PA14_22090 | hypothetical protein | 43 | 45 | 42 | 46 | 36 | 38 | 1 |
| 1790 | - | 1927303 | 1926578 | - | - | PA14_22100 | hypothetical protein | 5 | 4 | 3 | 4 | 5 | 5 | 1 |
| 1791 | - | 1928419 | 1927460 | - | - | PA14_22110 | hypothetical protein | 5 | 5 | 5 | 5 | 6 | 4 | 1 |
| 1792 | - | 1931366 | 1929042 | - | - | PA14_22120 | hypothetical protein | 5 | 3 | 4 | 5 | 3 | 4 | 1 |
| 1793 | - | 1932050 | 1931382 | - | - | PA14_22130 | hypothetical protein | 1 | 0 | 1 | 1 | 0 | 0 | 0.002850335 |
| 1794 | - | 1933820 | 1933020 | - | - | PA14_22140 | hypothetical protein | 5 | 4 | 4 | 5 | 5 | 5 | 1 |
| 1795 | - | 1934861 | 1934355 | - | - | PA14_22160 | hypothetical protein | 0 | 0 | 0 | 0 | 0 | 1 | 1 |
| 1796 | - | 1935117 | 1934977 | - | - | PA14_22180 | hypothetical protein | 8 | 8 | 7 | 7 | 6 | 4 | 1 |
| 1797 | - | 1935500 | 1935114 | - | - | PA14_22190 | hypothetical protein | 14 | 14 | 11 | 12 | 13 | 14 | 1 |
| 1798 | - | 1935905 | 1935570 | - | - | PA14_22210 | hypothetical protein | 6 | 5 | 4 | 5 | 5 | 6 | 1 |
| 1799 | - | 1936104 | 1935907 | - | - | PA14_22220 | hypothetical protein | 7 | 7 | 4 | 3 | 6 | 5 | 1 |
| 1800 | - | 1936868 | 1936671 | - | - | PA14_22230 | hypothetical protein | 5 | 6 | 2 | 3 | 2 | 2 | 1 |
| 1801 | - | 1937691 | 1937290 | - | - | PA14_22240 | hypothetical protein | 6 | 6 | 4 | 10 | 7 | 7 | 1 |
| 1802 | - | 1938417 | 1937791 | - | - | PA14_22250 | hypothetical protein | 0 | 1 | 2 | 1 | 0 | 1 | 1 |
| 1803 | - | 1938800 | 1938450 | - | - | PA14_22260 | hypothetical protein | 2 | 1 | 0 | 1 | 1 | 1 | 1 |
| 1804 | - | 1941291 | 1939714 | - | - | PA14_22270 | recombinase | 42 | 34 | 36 | 31 | 27 | 30 | 1 |
| 1805 | - | 1941585 | 1941322 | - | - | PA14_22280 | pirin-related protein | 28 | 31 | 29 | 23 | 20 | 24 | 1 |
| 1806 | - | 1942528 | 1941725 | - | vacJ | PA14_22290 | hypothetical protein | 63 | 62 | 58 | 82 | 70 | 70 | 1 |
| 1807 | - | 1943893 | 1942532 | - | - | PA14_22310 | hypothetical protein | 45 | 39 | 42 | 53 | 39 | 53 | 1 |
| 1808 | - | 1944257 | 1944036 | - | - | PA14_22320 | hypothetical protein | 22 | 7 | 8 | 13 | 9 | 76 | 0.001722686 |
| 1809 | - | 1944551 | 1945408 | + | - | PA14_22330 | glycine betaine-binding protein | 19 | 15 | 16 | 20 | 14 | 22 | 1 |
| 1810 | - | 1945626 | 1945937 | + | - | PA14_22340 | hypothetical protein | 142 | 133 | 94 | 85 | 63 | 81 | 1 |
| 1811 | - | 1945934 | 1947589 | + | actP | PA14_22350 | acetate permease | 91 | 82 | 67 | 61 | 56 | 45 | 1 |
| 1812 | - | 1947679 | 1949478 | + | - | PA14_22370 | hypothetical protein | 87 | 69 | 68 | 76 | 64 | 58 | 1 |
| 1813 | - | 1949475 | 1950101 | + | - | PA14_22380 | DNA polymerase III subunit epsilon | 35 | 23 | 26 | 20 | 17 | 17 | 1 |
| 1814 | - | 1950210 | 1950368 | + | - | PA14_22400 | hypothetical protein | 84 | 177 | 147 | 188 | 158 | 149 | 0.23199809 |
| 1815 | - | 1950645 | 1951769 | + | - | PA14_22410 | hypothetical protein | 34 | 39 | 33 | 41 | 34 | 33 | 1 |
| 1816 | - | 1952068 | 1951802 | - | - | PA14_22420 | hypothetical protein | 26 | 34 | 25 | 30 | 34 | 30 | 1 |
| 1817 | - | 1954182 | 1952350 | - | - | PA14_22440 | ABC transporter ATP-binding protein/permease | 115 | 118 | 112 | 113 | 124 | 123 | 1 |
| 1818 | - | 1954808 | 1954245 | - | ppiA | PA14_22450 | peptidyl-prolyl cis-trans isomerase A | 148 | 200 | 156 | 136 | 166 | 153 | 1 |
| 1819 | - | 1955617 | 1954790 | - | - | PA14_22460 | alpha/beta hydrolase | 90 | 100 | 100 | 89 | 96 | 99 | 1 |
| 1820 | - | 1956553 | 1955624 | - | - | PA14_22470 | LysR family transcriptional regulator | 105 | 99 | 105 | 95 | 105 | 104 | 1 |
| 1821 | - | 1956753 | 1957070 | + | - | PA14_22480 | hypothetical protein | 141 | 187 | 171 | 156 | 147 | 139 | 1 |
| 1822 | - | 1957217 | 1957858 | + | acpD | PA14_22490 | ACP phosphodieterase | 3 | 2 | 3 | 4 | 2 | 3 | 1 |
| 1823 | - | 1958847 | 1958113 | - | - | PA14_22500 | protein-disulfide isomerase | 9 | 9 | 9 | 10 | 9 | 11 | 1 |
| 1824 | - | 1959333 | 1958941 | - | - | PA14_22510 | hypothetical protein | 15 | 13 | 11 | 13 | 11 | 11 | 1 |
| 1825 | - | 1960494 | 1959439 | - | - | PA14_22520 | hypothetical protein | 11 | 10 | 9 | 10 | 8 | 9 | 1 |
| 1826 | - | 1961041 | 1960424 | - | - | PA14_22530 | glutathione S-transferase | 27 | 30 | 27 | 25 | 21 | 23 | 1 |
| 1827 | - | 1961671 | 1961117 | - | - | PA14_22540 | flavodoxin | 47 | 44 | 44 | 46 | 44 | 50 | 1 |
| 1828 | - | 1961914 | 1962726 | + | - | PA14_22550 | LysR family transcriptional regulator | 50 | 52 | 49 | 56 | 51 | 54 | 1 |
| 1829 | - | 1963696 | 1962803 | - | - | PA14_22560 | permease | 58 | 53 | 55 | 51 | 50 | 40 | 1 |
| 1830 | - | 1964041 | 1963706 | - | csaA | PA14_22570 | CsaA protein | 61 | 48 | 54 | 41 | 44 | 29 | 1 |
| 1831 | - | 1964870 | 1964106 | - | - | PA14_22580 | AraC family transcriptional regulator | 35 | 28 | 34 | 28 | 24 | 26 | 1 |
| 1832 | - | 1965017 | 1965829 | + | - | PA14_22590 | hypothetical protein | 7 | 7 | 7 | 8 | 7 | 5 | 1 |
| 1833 | - | 1965826 | 1966881 | + | - | PA14_22600 | glycosyl transferase family protein | 14 | 19 | 21 | 20 | 15 | 14 | 1 |
| 1834 | - | 1968262 | 1966871 | - | cyaB | PA14_22620 | hypothetical protein | 27 | 29 | 40 | 35 | 27 | 25 | 1 |
| 1835 | - | 1968796 | 1968479 | - | - | PA14_22630 | hypothetical protein | 27 | 39 | 34 | 28 | 31 | 29 | 1 |
| 1836 | - | 1968939 | 1969952 | + | - | PA14_22640 | AraC family transcriptional regulator | 88 | 89 | 93 | 89 | 75 | 80 | 1 |
| 1837 | - | 1970619 | 1969975 | - | - | PA14_22650 | ABC transporter | 223 | 239 | 205 | 239 | 224 | 222 | 1 |
| 1838 | - | 1971554 | 1970616 | - | - | PA14_22660 | hypothetical protein | 125 | 114 | 112 | 132 | 116 | 118 | 1 |
| 1839 | - | 1972350 | 1971556 | - | - | PA14_22670 | ABC transporter ATP-binding protein | 140 | 120 | 140 | 153 | 143 | 127 | 1 |
| 1840 | - | 1973492 | 1972347 | - | - | PA14_22680 | ABC transporter permease | 132 | 133 | 144 | 158 | 126 | 132 | 1 |
| 1841 | - | 1975013 | 1973559 | - | trkH | PA14_22690 | potassium uptake protein TrkH | 24 | 21 | 25 | 25 | 20 | 26 | 1 |
| 1842 | - | 1975171 | 1975542 | + | - | PA14_22700 | ferredoxin | 5 | 3 | 6 | 7 | 5 | 4 | 1 |
| 1843 | - | 1975607 | 1976167 | + | - | PA14_22710 | hypothetical protein | 90 | 102 | 113 | 102 | 80 | 69 | 1 |
| 1844 | - | 1976624 | 1976148 | - | - | PA14_22720 | hypothetical protein | 81 | 95 | 97 | 88 | 86 | 74 | 1 |
| 1845 | - | 1978005 | 1976668 | - | - | PA14_22730 | two-component sensor | 43 | 43 | 50 | 47 | 40 | 38 | 1 |
| 1846 | - | 1978560 | 1978120 | - | - | PA14_22740 | hypothetical protein | 49 | 115 | 61 | 86 | 18 | 24 | 0.085032199 |
| 1847 | - | 1979366 | 1978689 | - | cpxR | PA14_22760 | two-component response regulator | 69 | 60 | 63 | 69 | 60 | 63 | 1 |
| 1848 | - | 1979745 | 1979377 | - | - | PA14_22770 | hypothetical protein | 122 | 133 | 128 | 143 | 125 | 122 | 1 |

| | A | B | C | D | E | F | G | H | I | J | K | L | M | N |
|---|---|---|---|---|---|---|---|---|---|---|---|---|---|---|
| 1849 | - | 1980220 | 1979921 | - | *yciI* | *PA14_22780* | YciI-like protein | 114 | 135 | 128 | 136 | 126 | 120 | 1 |
| 1850 | - | 1980809 | 1980222 | - | *yciB* | *PA14_22800* | intracellular septation protein A | 106 | 83 | 111 | 86 | 85 | 89 | 1 |
| 1851 | - | 1981024 | 1981911 | + | - | *PA14_22820* | PHP domain-containing protein | 74 | 57 | 82 | 74 | 62 | 84 | 1 |
| 1852 | - | 1981908 | 1982537 | + | - | *PA14_22830* | SUA5/yciO/yrdC family:Sua5/YciO/YrdC/YwlC family protein | 91 | 95 | 97 | 110 | 97 | 118 | 1 |
| 1853 | - | 1982710 | 1983462 | + | - | *PA14_22840* | hypothetical protein | 71 | 68 | 79 | 65 | 60 | 67 | 1 |
| 1854 | - | 1983607 | 1984599 | + | - | *PA14_22860* | transcriptional regulator | 62 | 37 | 60 | 50 | 40 | 46 | 1 |
| 1855 | - | 1984681 | 1985211 | + | - | *PA14_22870* | hypothetical protein | 18 | 25 | 23 | 22 | 27 | 18 | 1 |
| 1856 | - | 1985296 | 1985508 | + | - | *PA14_22880* | Fe-S protein | 101 | 103 | 98 | 100 | 93 | 119 | 1 |
| 1857 | - | 1986519 | 1985515 | - | *gapA* | *PA14_22890* | glyceraldehyde-3-phosphate dehydrogenase | 108 | 191 | 149 | 190 | 164 | 145 | 1 |
| 1858 | - | 1986649 | 1988475 | + | *edd* | *PA14_22910* | phosphogluconate dehydratase | 64 | 180 | 88 | 155 | 195 | 89 | 0.321350893 |
| 1859 | - | 1988578 | 1989573 | + | *glk* | *PA14_22930* | glucokinase | 103 | 196 | 127 | 165 | 216 | 112 | 1 |
| 1860 | - | 1989608 | 1990336 | + | *gltR* | *PA14_22940* | two-component response regulator GltR | 51 | 116 | 62 | 85 | 113 | 58 | 0.260298623 |
| 1861 | - | 1990360 | 1991781 | + | - | *PA14_22960* | two-component sensor | 25 | 63 | 37 | 57 | 61 | 35 | 0.104266321 |
| 1862 | - | 1992306 | 1993568 | + | - | *PA14_22980* | sugar ABC transporter substrate-binding protein | 10 | 679 | 387 | 547 | 594 | 472 | 0 |
| 1863 | - | 1993669 | 1994601 | + | - | *PA14_22990* | ABC sugar transporter permease | 11 | 183 | 101 | 145 | 153 | 109 | 3.4879E-157 |
| 1864 | - | 1994594 | 1995439 | + | - | *PA14_23000* | ABC sugar transporter permease | 14 | 178 | 88 | 128 | 131 | 105 | 6.23215E-97 |
| 1865 | - | 1995472 | 1996632 | + | *gltK* | *PA14_23010* | ABC transporter ATP-binding protein | 9 | 504 | 263 | 373 | 404 | 311 | 0 |
| 1866 | - | 1996676 | 1998040 | + | *oprB* | *PA14_23030* | glucose/carbohydrate outer membrane porin OprB precursor | 65 | 926 | 502 | 712 | 719 | 599 | 5.54754E-82 |
| 1867 | - | 1998255 | 1999106 | + | - | *PA14_23050* | hypothetical protein | 335 | 353 | 399 | 444 | 356 | 368 | 1 |
| 1868 | - | 1999972 | 1999115 | - | *hexR* | *PA14_23060* | DNA-binding transcriptional regulator HexR | 60 | 68 | 64 | 77 | 64 | 58 | 1 |
| 1869 | - | 2000159 | 2001628 | + | *zwf* | *PA14_23070* | glucose-6-phosphate 1-dehydrogenase | 49 | 125 | 59 | 134 | 155 | 78 | 0.158443239 |
| 1870 | - | 2001615 | 2002331 | + | *pgl* | *PA14_23080* | 6-phosphogluconolactonase | 44 | 147 | 56 | 131 | 181 | 72 | 0.001697282 |
| 1871 | - | 2002349 | 2003011 | + | - | *PA14_23090* | keto-hydroxyglutarate-aldolase/keto-deoxy-phosphogluconate aldolase | 77 | 113 | 99 | 112 | 112 | 81 | 1 |
| 1872 | - | 2003538 | 2003101 | - | - | *PA14_23100* | hypothetical protein | 38 | 34 | 34 | 45 | 32 | 34 | 1 |
| 1873 | - | 2003686 | 2004846 | + | - | *PA14_23110* | hypothetical protein | 51 | 47 | 59 | 62 | 47 | 55 | 1 |
| 1874 | - | 2004939 | 2005316 | + | - | *PA14_23120* | hypothetical protein | 98 | 90 | 99 | 104 | 86 | 87 | 1 |
| 1875 | - | 2006346 | 2005423 | - | - | *PA14_23130* | sensory box GGDEF domain-containing protein | 57 | 45 | 52 | 46 | 41 | 44 | 1 |
| 1876 | - | 2007789 | 2006575 | - | *gltS* | *PA14_23160* | sodium/glutamate symporter | 5 | 6 | 5 | 5 | 4 | 4 | 1 |
| 1877 | - | 2008763 | 2007828 | - | - | *PA14_23170* | formimidoylglutamase | 22 | 22 | 25 | 25 | 17 | 21 | 1 |
| 1878 | - | 2009507 | 2008779 | - | - | *PA14_23190* | transcriptional regulator | 11 | 8 | 11 | 10 | 8 | 7 | 1 |
| 1879 | - | 2010406 | 2009666 | - | - | *PA14_23200* | short chain dehydrogenase | 89 | 86 | 94 | 85 | 82 | 84 | 1 |
| 1880 | - | 2011151 | 2010471 | - | - | *PA14_23210* | phosphoglycolate phosphatase | 130 | 132 | 143 | 130 | 127 | 116 | 1 |
| 1881 | - | 2011846 | 2011148 | - | *ubiG* | *PA14_23220* | 3-demethylubiquinone-9 3-methyltransferase | 202 | 165 | 222 | 174 | 158 | 171 | 1 |
| 1882 | - | 2013331 | 2011997 | - | - | *PA14_23240* | N-ethylammeline chlorohydrolase | 44 | 38 | 46 | 40 | 38 | 38 | 1 |
| 1883 | - | 2013441 | 2014517 | + | *mtnA* | *PA14_23250* | methylthioribose-1-phosphate isomerase | 64 | 58 | 68 | 68 | 63 | 53 | 1 |
| 1884 | - | 2014754 | 2017519 | + | *gyrA* | *PA14_23260* | DNA gyrase subunit A | 285 | 292 | 294 | 283 | 297 | 257 | 1 |
| 1885 | - | 2017607 | 2018692 | + | *serC* | *PA14_23270* | phosphoserine aminotransferase | 270 | 276 | 277 | 257 | 285 | 256 | 1 |
| 1886 | - | 2018692 | 2019789 | + | *pheA* | *PA14_23280* | chorismate mutase | 257 | 259 | 246 | 272 | 300 | 265 | 1 |
| 1887 | - | 2019858 | 2020967 | + | *hisC2* | *PA14_23290* | histidinol-phosphate aminotransferase | 117 | 99 | 109 | 88 | 121 | 93 | 1 |
| 1888 | - | 2020960 | 2023200 | + | - | *PA14_23310* | bifunctional cyclohexadienyl dehydrogenase/ 3-phosphoshikimate 1-carboxyvinyltransferase | 163 | 169 | 151 | 131 | 181 | 121 | 1 |
| 1889 | - | 2023200 | 2023889 | + | *cmk* | *PA14_23320* | cytidylate kinase | 166 | 187 | 167 | 137 | 180 | 143 | 1 |
| 1890 | - | 2024157 | 2025836 | + | *rpsA* | *PA14_23330* | 30S ribosomal protein S1 | 599 | 641 | 623 | 473 | 536 | 388 | 1 |
| 1891 | - | 2025973 | 2026257 | + | *ihfB* | *PA14_23340* | integration host factor subunit beta | 3265 | 3404 | 3944 | 4185 | 2514 | 2683 | 1 |
| 1892 | - | 2026291 | 2026590 | + | *orfA* | *PA14_23350* | hypothetical protein | 326 | 396 | 263 | 314 | 367 | 259 | 1 |
| 1893 | - | 2026846 | 2027928 | + | *wzz* | *PA14_23360* | O-antigen chain length regulator | 59 | 48 | 50 | 40 | 49 | 43 | 1 |
| 1894 | - | 2028084 | 2029220 | + | *orfK* | *PA14_23370* | UDP-N-acetylglucosamine 2-epimerase | 242 | 193 | 172 | 162 | 252 | 183 | 1 |
| 1895 | - | 2029301 | 2030569 | + | *orfH* | *PA14_23380* | UDP-N-acetyl-D-mannosaminuronate dehydrogenase | 159 | 219 | 129 | 124 | 165 | 121 | 1 |
| 1896 | - | 2030647 | 2031909 | + | *orfE* | *PA14_23390* | polysaccharide biosynthesis protein | 53 | 44 | 46 | 46 | 71 | 54 | 1 |
| 1897 | - | 2031887 | 2033236 | + | - | *PA14_23400* | hypothetical protein | 92 | 70 | 73 | 82 | 133 | 108 | 1 |
| 1898 | - | 2033233 | 2034075 | + | *orfJ* | *PA14_23410* | glycosyl transferase family protein | 90 | 98 | 70 | 76 | 112 | 88 | 1 |
| 1899 | - | 2034189 | 2036357 | + | - | *PA14_23420* | zinc-binding dehydrogenase | 214 | 267 | 149 | 164 | 254 | 173 | 1 |
| 1900 | - | 2036366 | 2038018 | + | - | *PA14_23430* | heparinase | 135 | 143 | 101 | 94 | 163 | 120 | 1 |
| 1901 | - | 2038087 | 2039199 | + | *orfL* | *PA14_23440* | group 1 glycosyl transferase | 129 | 145 | 87 | 99 | 166 | 125 | 1 |
| 1902 | - | 2039196 | 2040149 | + | *orfM* | *PA14_23450* | NAD dependent epimerase/dehydratase | 319 | 282 | 268 | 275 | 329 | 297 | 1 |
| 1903 | - | 2040149 | 2041165 | + | *orfN* | *PA14_23460* | group 4 glycosyl transferase | 79 | 59 | 60 | 63 | 105 | 74 | 1 |
| 1904 | - | 2041610 | 2043607 | + | *wbpM* | *PA14_23470* | nucleotide sugar epimerase/dehydratase WbpM | 340 | 273 | 287 | 345 | 299 | 272 | 1 |
| 1905 | - | 2043798 | 2044127 | + | - | *PA14_23480* | hypothetical protein | 7 | 9 | 10 | 8 | 9 | 10 | 1 |
| 1906 | - | 2044275 | 2044203 | - | - | *PA14_23490* | Asn tRNA | 7 | 7 | 7 | 6 | 3 | 5 | 1 |
| 1907 | - | 2045540 | 2044344 | - | *tyrB* | *PA14_23500* | aromatic amino acid aminotransferase | 83 | 71 | 83 | 65 | 63 | 58 | 1 |
| 1908 | - | 2045728 | 2047740 | + | *uvrB* | *PA14_23510* | excinuclease ABC subunit B | 64 | 63 | 61 | 61 | 52 | 51 | 1 |
| 1909 | - | 2049243 | 2047744 | - | - | *PA14_23520* | MFS transporter | 47 | 38 | 51 | 42 | 32 | 38 | 1 |
| 1910 | - | 2050360 | 2049293 | - | - | *PA14_23530* | secretion protein | 4 | 4 | 6 | 4 | 2 | 4 | 1 |
| 1911 | - | 2050430 | 2051350 | + | *act* | *PA14_23540* | transcriptional regulator | 17 | 12 | 16 | 17 | 16 | 17 | 1 |
| 1912 | - | 2051388 | 2052872 | + | *gltX* | *PA14_23560* | glutamyl-tRNA synthetase | 59 | 69 | 53 | 48 | 60 | 46 | 1 |
| 1913 | - | 2053367 | 2053439 | + | - | *PA14_23570* | Ala tRNA | 279 | 72 | 107 | 66 | 38 | 34 | 4.59105E-11 |
| 1914 | - | 2053489 | 2053561 | + | - | *PA14_23580* | Glu tRNA | 157 | 9 | 20 | 7 | 6 | 7 | 7.7397E-255 |
| 1915 | - | 2053723 | 2054265 | + | - | *PA14_23590* | transcriptional regulator | 8 | 7 | 7 | 7 | 6 | 9 | 1 |
| 1916 | - | 2054277 | 2055134 | + | - | *PA14_23610* | hydrolase | 14 | 16 | 18 | 19 | 15 | 16 | 1 |
| 1917 | - | 2055247 | 2055894 | + | *kdgA* | *PA14_23620* | aldolase | 26 | 31 | 31 | 30 | 27 | 24 | 1 |
| 1918 | - | 2055894 | 2056331 | + | - | *PA14_23630* | hypothetical protein | 25 | 30 | 23 | 24 | 31 | 22 | 1 |
| 1919 | - | 2056424 | 2057383 | + | - | *PA14_23640* | hypothetical protein | 40 | 38 | 45 | 40 | 39 | 33 | 1 |
| 1920 | - | 2058144 | 2057398 | - | - | *PA14_23650* | short chain dehydrogenase | 67 | 61 | 55 | 62 | 59 | 63 | 1 |
| 1921 | - | 2058964 | 2058161 | - | - | *PA14_23670* | hypothetical protein | 31 | 31 | 27 | 24 | 29 | 25 | 1 |
| 1922 | - | 2059122 | 2059571 | + | *ibpA* | *PA14_23680* | heat-shock protein IbpA | 643 | 618 | 685 | 278 | 387 | 319 | 1 |
| 1923 | - | 2060943 | 2059615 | - | - | *PA14_23690* | secreted protein | 15 | 14 | 18 | 13 | 13 | 13 | 1 |
| 1924 | - | 2061055 | 2061966 | + | - | *PA14_23700* | LysR family transcriptional regulator | 46 | 46 | 41 | 40 | 41 | 40 | 1 |
| 1925 | - | 2062046 | 2062423 | + | - | *PA14_23720* | translation initiation inhibitor | 156 | 174 | 156 | 197 | 168 | 228 | 1 |
| 1926 | - | 2063409 | 2062516 | - | - | *PA14_23730* | LysR family transcriptional regulator | 76 | 75 | 82 | 100 | 81 | 102 | 1 |
| 1927 | - | 2063572 | 2064996 | + | *leuC* | *PA14_23750* | isopropylmalate isomerase large subunit | 80 | 68 | 73 | 78 | 63 | 77 | 1 |
| 1928 | - | 2065008 | 2065646 | + | *leuD* | *PA14_23760* | isopropylmalate isomerase small subunit | 106 | 114 | 104 | 126 | 97 | 120 | 1 |
| 1929 | - | 2065916 | 2066479 | + | - | *PA14_23770* | biotin synthesis protein | 48 | 34 | 44 | 36 | 33 | 41 | 1 |
| 1930 | - | 2066538 | 2067620 | + | *leuB* | *PA14_23790* | 3-isopropylmalate dehydrogenase | 68 | 73 | 72 | 68 | 63 | 65 | 1 |
| 1931 | - | 2067690 | 2068802 | + | *asd* | *PA14_23800* | aspartate-semialdehyde dehydrogenase | 147 | 128 | 145 | 150 | 164 | 152 | 1 |
| 1932 | - | 2068904 | 2069914 | + | - | *PA14_23810* | aspartate-semialdehyde dehydrogenase | 57 | 50 | 67 | 54 | 51 | 51 | 1 |

| | A | B | C | D | E | F | G | H | I | J | K | L | M | N |
|---|---|---|---|---|---|---|---|---|---|---|---|---|---|---|
| 1933 | - | 2070083 | 2072857 | + | *fimV* | *PA14_23830* | pilus assembly protein | 375 | 434 | 422 | 372 | 347 | 371 | 1 |
| 1934 | - | 2072879 | 2073736 | + | *truA* | *PA14_23840* | tRNA pseudouridine synthase A | 108 | 111 | 120 | 103 | 93 | 99 | 1 |
| 1935 | - | 2073825 | 2074460 | + | *trpF* | *PA14_23850* | N-(5'-phosphoribosyl)anthranilate isomerase | 372 | 326 | 465 | 366 | 318 | 299 | 1 |
| 1936 | - | 2074729 | 2075601 | + | *accD* | *PA14_23860* | acetyl-CoA carboxylase subunit beta | 229 | 214 | 238 | 208 | 193 | 192 | 1 |
| 1937 | - | 2075598 | 2076887 | + | *folC* | *PA14_23880* | folylpolyglutamate synthetase | 167 | 179 | 167 | 162 | 147 | 131 | 1 |
| 1938 | - | 2076891 | 2077538 | + | - | *PA14_23890* | hypothetical protein | 125 | 128 | 125 | 135 | 129 | 115 | 1 |
| 1939 | - | 2077612 | 2078154 | + | *cvpA* | *PA14_23900* | hypothetical protein | 44 | 35 | 53 | 48 | 41 | 35 | 1 |
| 1940 | - | 2078376 | 2079881 | + | *purF* | *PA14_23920* | amidophosphoribosyltransferase | 151 | 150 | 154 | 128 | 131 | 119 | 1 |
| 1941 | - | 2079898 | 2081109 | + | *metZ* | *PA14_23930* | O-succinylhomoserine sulfhydrylase | 238 | 219 | 231 | 210 | 230 | 204 | 1 |
| 1942 | - | 2081106 | 2081873 | + | - | *PA14_23950* | oxidoreductase | 168 | 150 | 190 | 156 | 176 | 146 | 1 |
| 1943 | - | 2083936 | 2081960 | - | *xcpQ* | *PA14_23970* | general secretion pathway protein D | 234 | 243 | 219 | 293 | 257 | 275 | 1 |
| 1944 | - | 2084648 | 2083941 | - | *xcpP* | *PA14_23980* | secretion protein XcpP | 118 | 109 | 132 | 148 | 117 | 127 | 1 |
| 1945 | - | 2084868 | 2086376 | + | *xcpR* | *PA14_23990* | general secretion pathway protein E | 191 | 190 | 188 | 246 | 189 | 235 | 1 |
| 1946 | - | 2086376 | 2087593 | + | *xcpS* | *PA14_24010* | general secretion pathway protein F | 142 | 135 | 143 | 185 | 156 | 176 | 1 |
| 1947 | - | 2087598 | 2088044 | + | *xcpT* | *PA14_24020* | general secretion pathway protein G | 248 | 335 | 255 | 350 | 375 | 363 | 1 |
| 1948 | - | 2088051 | 2088569 | + | *xcpU* | *PA14_24040* | general secretion pathway outer membrane protein H precursor | 177 | 198 | 187 | 210 | 205 | 210 | 1 |
| 1949 | - | 2088566 | 2088955 | + | *xcpV* | *PA14_24050* | general secretion pathway protein I | 134 | 149 | 144 | 170 | 149 | 164 | 1 |
| 1950 | - | 2088952 | 2089665 | + | *xcpW* | *PA14_24060* | general secretion pathway protein J | 171 | 181 | 180 | 240 | 208 | 214 | 1 |
| 1951 | - | 2089662 | 2090663 | + | *xcpX* | *PA14_24070* | general secretion pathway protein K | 95 | 92 | 112 | 129 | 116 | 116 | 1 |
| 1952 | - | 2090660 | 2091808 | + | *xcpY* | *PA14_24080* | general secretion pathway protein L | 301 | 294 | 310 | 378 | 331 | 332 | 1 |
| 1953 | - | 2091810 | 2092334 | + | *xcpZ* | *PA14_24100* | general secretion pathway protein M | 182 | 177 | 189 | 226 | 204 | 184 | 1 |
| 1954 | - | 2092473 | 2092545 | + | - | *PA14_24110* | Val tRNA | 263 | 172 | 269 | 235 | 162 | 109 | 1 |
| 1955 | - | 2092565 | 2092638 | + | - | *PA14_24120* | Asp tRNA | 785 | 749 | 868 | 774 | 576 | 489 | 1 |
| 1956 | - | 2092756 | 2092829 | + | - | *PA14_24130* | Asp tRNA | 344 | 334 | 372 | 294 | 223 | 189 | 1 |
| 1957 | - | 2094157 | 2093120 | - | - | *PA14_24140* | AraC family transcriptional regulator | 37 | 41 | 39 | 40 | 32 | 31 | 1 |
| 1958 | - | 2095272 | 2094154 | - | - | *PA14_24150* | hypothetical protein | 37 | 33 | 41 | 35 | 29 | 32 | 1 |
| 1959 | - | 2095487 | 2097526 | + | *fadH1* | *PA14_24170* | 2,4-dienoyl-CoA reductase | 26 | 28 | 26 | 25 | 25 | 22 | 1 |
| 1960 | - | 2097863 | 2099302 | + | - | *PA14_24180* | hypothetical protein | 98 | 107 | 120 | 131 | 83 | 94 | 1 |
| 1961 | - | 2099307 | 2100239 | + | - | *PA14_24190* | 1-aminocyclopropane-1-carboxylate deaminase | 50 | 58 | 57 | 55 | 49 | 55 | 1 |
| 1962 | - | 2101205 | 2100240 | - | - | *PA14_24210* | hypothetical protein | 69 | 75 | 77 | 80 | 85 | 111 | 1 |
| 1963 | - | 2101330 | 2102217 | + | *ppnK* | *PA14_24220* | inorganic polyphosphate/ATP-NAD kinase | 92 | 75 | 94 | 72 | 77 | 74 | 1 |
| 1964 | - | 2102217 | 2103197 | + | - | *PA14_24230* | hypothetical protein | 92 | 90 | 86 | 91 | 75 | 84 | 1 |
| 1965 | - | 2103194 | 2104054 | + | - | *PA14_24240* | hypothetical protein | 151 | 136 | 145 | 154 | 144 | 136 | 1 |
| 1966 | - | 2104138 | 2104401 | + | - | *PA14_24245* | hypothetical protein | 175 | 151 | 184 | 125 | 148 | 143 | 1 |
| 1967 | - | 2104434 | 2105234 | + | - | *PA14_24260* | hypothetical protein | 119 | 113 | 127 | 111 | 107 | 110 | 1 |
| 1968 | - | 2105253 | 2107910 | + | *pepN* | *PA14_24270* | aminopeptidase | 163 | 175 | 153 | 151 | 160 | 157 | 1 |
| 1969 | - | 2108357 | 2110321 | + | *gbt* | *PA14_24290* | glycine betaine transmethylase | 45 | 59 | 47 | 43 | 45 | 40 | 1 |
| 1970 | - | 2110380 | 2111747 | + | - | *PA14_24300* | hypothetical protein | 37 | 52 | 38 | 31 | 42 | 28 | 1 |
| 1971 | - | 2111927 | 2113024 | + | - | *PA14_24310* | BNR/Asp-box repeat-containing protein | 48 | 78 | 66 | 80 | 70 | 61 | 0.931437202 |
| 1972 | - | 2113036 | 2115417 | + | - | *PA14_24330* | hypothetical protein | 44 | 76 | 57 | 75 | 68 | 60 | 1 |
| 1973 | - | 2116784 | 2115489 | - | - | *PA14_24340* | two-component sensor | 20 | 24 | 24 | 25 | 23 | 21 | 1 |
| 1974 | - | 2117452 | 2116781 | - | - | *PA14_24350* | two-component response regulator | 15 | 10 | 17 | 14 | 14 | 13 | 0.969261494 |
| 1975 | - | 2117776 | 2120700 | + | - | *PA14_24360* | hypothetical protein | 11 | 11 | 12 | 12 | 10 | 12 | 1 |
| 1976 | - | 2121786 | 2120707 | - | - | *PA14_24370* | hypothetical protein | 69 | 65 | 68 | 72 | 65 | 74 | 1 |
| 1977 | - | 2123471 | 2121840 | - | - | *PA14_24380* | hypothetical protein | 66 | 60 | 64 | 74 | 62 | 73 | 1 |
| 1978 | - | 2125228 | 2123468 | - | - | *PA14_24390* | hypothetical protein | 44 | 36 | 44 | 45 | 38 | 43 | 1 |
| 1979 | - | 2126247 | 2125225 | - | - | *PA14_24400* | von Willebrand factor A | 34 | 29 | 32 | 36 | 29 | 33 | 1 |
| 1980 | - | 2126734 | 2126240 | - | - | *PA14_24410* | hypothetical protein | 29 | 33 | 32 | 33 | 25 | 31 | 1 |
| 1981 | - | 2127669 | 2126731 | - | - | *PA14_24420* | hypothetical protein | 61 | 54 | 63 | 60 | 47 | 54 | 1 |
| 1982 | - | 2128660 | 2127680 | - | - | *PA14_24430* | hypothetical protein | 107 | 90 | 119 | 99 | 92 | 103 | 1 |
| 1983 | - | 2129153 | 2129770 | + | - | *PA14_24440* | lipoprotein | 68 | 46 | 71 | 67 | 50 | 56 | 1 |
| 1984 | - | 2130044 | 2134906 | + | *gdhB* | *PA14_24445* | NAD-dependent glutamate dehydrogenase | 365 | 542 | 349 | 386 | 452 | 377 | 1 |
| 1985 | - | 2135468 | 2138314 | + | *pelA* | *PA14_24480* | hypothetical protein | 14 | 6 | 8 | 7 | 5 | 6 | 0.23199809 |
| 1986 | - | 2138292 | 2141873 | + | *pelB* | *PA14_24490* | hypothetical protein | 14 | 5 | 5 | 5 | 5 | 5 | 0.044138282 |
| 1987 | - | 2141913 | 2142431 | + | *pelC* | *PA14_24500* | lipoprotein | 18 | 5 | 4 | 4 | 5 | 3 | 1.46581E-05 |
| 1988 | - | 2142437 | 2143804 | + | *pelD* | *PA14_24510* | hypothetical protein | 12 | 4 | 4 | 4 | 4 | 3 | 0.000626932 |
| 1989 | - | 2143782 | 2144771 | + | *pelE* | *PA14_24530* | hypothetical protein | 24 | 14 | 18 | 20 | 14 | 14 | 0.915523483 |
| 1990 | - | 2144768 | 2146291 | + | *pelF* | *PA14_24550* | hypothetical protein | 16 | 8 | 8 | 7 | 7 | 8 | 0.480861202 |
| 1991 | - | 2146293 | 2147663 | + | *pelG* | *PA14_24560* | hypothetical protein | 11 | 7 | 6 | 7 | 7 | 8 | 1 |
| 1992 | - | 2147934 | 2147704 | - | - | *PA14_24570* | hypothetical protein | 11 | 9 | 10 | 7 | 10 | 8 | 1 |
| 1993 | - | 2148060 | 2148536 | + | - | *PA14_24580* | hypothetical protein | 30 | 31 | 30 | 22 | 23 | 24 | 1 |
| 1994 | - | 2148574 | 2149050 | + | - | *PA14_24590* | hypothetical protein | 62 | 66 | 64 | 57 | 60 | 58 | 1 |
| 1995 | - | 2151071 | 2149167 | - | - | *PA14_24600* | carboxypeptidase | 30 | 32 | 31 | 23 | 30 | 23 | 1 |
| 1996 | - | 2152240 | 2151239 | - | - | *PA14_24610* | hydrolytic enzyme | 26 | 35 | 28 | 26 | 35 | 26 | 1 |
| 1997 | - | 2153286 | 2153567 | + | - | *PA14_24630* | hypothetical protein | 86 | 87 | 94 | 102 | 82 | 84 | 1 |
| 1998 | - | 2153292 | 2152312 | - | - | *PA14_24620* | hypothetical protein | 59 | 58 | 70 | 53 | 61 | 58 | 1 |
| 1999 | - | 2153624 | 2154652 | + | *pyrD* | *PA14_24640* | dihydroorotate dehydrogenase 2 | 71 | 59 | 94 | 71 | 55 | 59 | 1 |
| 2000 | - | 2154952 | 2154740 | - | *rmf* | *PA14_24650* | ribosome modulation factor | 1022 | 1426 | 992 | 1646 | 1141 | 1471 | 1 |
| 2001 | - | 2155414 | 2156517 | + | - | *PA14_24665* | hypothetical protein | 48 | 41 | 51 | 45 | 40 | 40 | 1 |
| 2002 | - | 2156561 | 2157592 | + | - | *PA14_24675* | hypothetical protein | 40 | 38 | 46 | 44 | 38 | 37 | 1 |
| 2003 | - | 2159092 | 2157662 | - | - | *PA14_24690* | D-alanyl-D-alanine carboxypeptidase | 79 | 68 | 80 | 66 | 72 | 70 | 1 |
| 2004 | - | 2159403 | 2159747 | + | - | *PA14_24700* | hypothetical protein | 77 | 100 | 62 | 49 | 92 | 55 | 1 |
| 2005 | - | 2160174 | 2160797 | + | - | *PA14_24710* | two-component response regulator | 9 | 11 | 11 | 8 | 11 | 11 | 1 |
| 2006 | - | 2160811 | 2163036 | + | - | *PA14_24720* | two-component sensor | 11 | 13 | 12 | 12 | 12 | 13 | 1 |
| 2007 | - | 2164378 | 2163047 | - | - | *PA14_24730* | deoxyguanosinetriphosphate triphosphohydrolase-like protein | 91 | 96 | 88 | 77 | 89 | 85 | 1 |
| 2008 | - | 2164818 | 2164495 | - | - | *PA14_24740* | hypothetical protein | 308 | 385 | 364 | 281 | 312 | 260 | 1 |
| 2009 | - | 2165212 | 2164832 | - | - | *PA14_24760* | hypothetical protein | 445 | 565 | 448 | 385 | 539 | 452 | 1 |
| 2010 | - | 2165545 | 2165216 | - | - | *PA14_24770* | hypothetical protein | 601 | 858 | 605 | 472 | 649 | 608 | 1 |
| 2011 | - | 2165841 | 2167223 | + | - | *PA14_24780* | transporter | 6 | 4 | 6 | 5 | 5 | 6 | 1 |
| 2012 | - | 2168673 | 2167408 | - | - | *PA14_24790* | outer membrane porin | 54 | 91 | 62 | 61 | 56 | 56 | 1 |
| 2013 | - | 2169708 | 2168842 | - | - | *PA14_24810* | hypothetical protein | 26 | 18 | 17 | 21 | 16 | 29 | 1 |
| 2014 | - | 2170640 | 2169708 | - | - | *PA14_24820* | hypothetical protein | 13 | 10 | 11 | 12 | 9 | 14 | 1 |
| 2015 | - | 2171253 | 2170657 | - | - | *PA14_24830* | glutathione S-transferase | 11 | 11 | 15 | 12 | 8 | 10 | 1 |
| 2016 | - | 2171419 | 2171976 | + | - | *PA14_24840* | transcriptional regulator | 44 | 35 | 49 | 35 | 33 | 36 | 1 |

| | A | B | C | D | E | F | G | H | I | J | K | L | M | N |
|---|---|---|---|---|---|---|---|---|---|---|---|---|---|---|
| 2017 | - | 2171983 | 2172261 | + | - | *PA14_24850* | hypothetical protein | 31 | 30 | 30 | 23 | 28 | 27 | 1 |
| 2018 | - | 2172628 | 2174031 | + | *snr1* | *PA14_24860* | cytochrome c Snr1 | 89 | 12 | 13 | 17 | 12 | 160 | 9.5063E-30 |
| 2019 | - | 2174165 | 2174092 | - | - | *PA14_24870* | Pro tRNA | 2 | 1 | 3 | 3 | 1 | 1 | 0.005569024 |
| 2020 | - | 2174540 | 2174319 | - | - | *PA14_24880* | lipoprotein | 574 | 671 | 479 | 418 | 537 | 533 | 1 |
| 2021 | - | 2175268 | 2174672 | - | *mobA* | *PA14_24890* | molybdopterin-guanine dinucleotide biosynthesis protein MobA | 31 | 25 | 31 | 30 | 25 | 32 | 1 |
| 2022 | - | 2175335 | 2175874 | + | *moaB2* | *PA14_24900* | molybdopterin biosynthetic protein B2 | 88 | 98 | 80 | 71 | 79 | 63 | 1 |
| 2023 | - | 2175871 | 2177088 | + | *moeA2* | *PA14_24910* | molybdenum cofactor biosynthesis protein A2 | 121 | 123 | 99 | 99 | 118 | 94 | 1 |
| 2024 | - | 2178148 | 2177117 | - | - | *PA14_24920* | transcriptional regulator | 35 | 34 | 35 | 36 | 34 | 29 | 1 |
| 2025 | - | 2178325 | 2179920 | + | - | *PA14_24940* | hypothetical protein | 35 | 35 | 36 | 37 | 39 | 36 | 1 |
| 2026 | - | 2180121 | 2181506 | + | - | *PA14_24950* | FAD-dependent glycerol-3-phosphate dehydrogenase | 41 | 37 | 38 | 38 | 38 | 41 | 1 |
| 2027 | - | 2181527 | 2183086 | + | - | *PA14_24960* | carbohydrate kinase | 48 | 45 | 50 | 51 | 49 | 54 | 1 |
| 2028 | - | 2183161 | 2184069 | + | - | *PA14_24970* | lipid kinase | 70 | 60 | 81 | 67 | 70 | 69 | 1 |
| 2029 | - | 2184172 | 2184978 | + | - | *PA14_24980* | hypothetical protein | 287 | 217 | 268 | 324 | 256 | 269 | 1 |
| 2030 | - | 2185408 | 2185022 | - | - | *PA14_24990* | hypothetical protein | 191 | 177 | 160 | 226 | 215 | 182 | 1 |
| 2031 | - | 2187393 | 2185465 | - | - | *PA14_25000* | lytic transglycosylase | 73 | 70 | 77 | 82 | 74 | 83 | 1 |
| 2032 | - | 2187619 | 2189541 | + | - | *PA14_25020* | ABC transporter ATP-binding protein | 59 | 62 | 74 | 60 | 57 | 50 | 1 |
| 2033 | - | 2189692 | 2190378 | + | - | *PA14_25030* | hypothetical protein | 52 | 53 | 54 | 42 | 67 | 45 | 1 |
| 2034 | - | 2190845 | 2190408 | - | - | *PA14_25040* | hypothetical protein | 183 | 184 | 178 | 163 | 206 | 184 | 1 |
| 2035 | - | 2191053 | 2191487 | + | - | *PA14_25050* | hypothetical protein | 14 | 16 | 19 | 14 | 15 | 16 | 1 |
| 2036 | - | 2192322 | 2191504 | - | - | *PA14_25060* | hypothetical protein | 12 | 18 | 17 | 13 | 12 | 14 | 1 |
| 2037 | - | 2192750 | 2194897 | + | *fadB* | *PA14_25080* | multifunctional fatty acid oxidation complex subunit alpha | 192 | 247 | 219 | 176 | 209 | 181 | 1 |
| 2038 | - | 2194928 | 2196103 | + | *fadA* | *PA14_25090* | 3-ketoacyl-CoA thiolase | 193 | 278 | 197 | 190 | 248 | 207 | 1 |
| 2039 | - | 2196186 | 2196560 | + | - | *PA14_25100* | hypothetical protein | 59 | 45 | 74 | 66 | 55 | 63 | 1 |
| 2040 | - | 2196651 | 2199257 | + | *topA* | *PA14_25110* | DNA topoisomerase I | 152 | 177 | 164 | 174 | 171 | 152 | 1 |
| 2041 | - | 2199364 | 2199882 | + | - | *PA14_25130* | hypothetical protein | 140 | 153 | 163 | 151 | 142 | 142 | 1 |
| 2042 | - | 2200088 | 2200321 | + | - | *PA14_25140* | hypothetical protein | 115 | 77 | 155 | 96 | 97 | 113 | 1 |
| 2043 | - | 2200828 | 2200343 | - | - | *PA14_25150* | hypothetical protein | 73 | 73 | 87 | 67 | 67 | 75 | 1 |
| 2044 | - | 2201454 | 2200840 | - | *lexA* | *PA14_25160* | LexA repressor | 73 | 72 | 78 | 78 | 76 | 80 | 1 |
| 2045 | - | 2201686 | 2202387 | + | *psrA* | *PA14_25180* | transcriptional regulator PsrA | 972 | 774 | 1094 | 918 | 890 | 922 | 1 |
| 2046 | - | 2202589 | 2203587 | + | *nagZ* | *PA14_25195* | beta-hexosaminidase | 57 | 47 | 56 | 48 | 47 | 46 | 1 |
| 2047 | - | 2203599 | 2204336 | + | - | *PA14_25210* | 5'-methylthioadenosine phosphorylase | 98 | 103 | 88 | 108 | 108 | 95 | 1 |
| 2048 | - | 2205012 | 2204383 | - | - | *PA14_25220* | hypothetical protein | 112 | 108 | 110 | 113 | 94 | 111 | 1 |
| 2049 | - | 2208469 | 2205023 | - | *mfd* | *PA14_25230* | transcription-repair coupling factor | 112 | 114 | 108 | 114 | 99 | 99 | 1 |
| 2050 | - | 2208690 | 2210075 | + | - | *PA14_25250* | glyceraldehyde-3-phosphate dehydrogenase | 252 | 251 | 267 | 187 | 188 | 158 | 1 |
| 2051 | - | 2210177 | 2211586 | + | *aroP1* | *PA14_25270* | aromatic amino acid transport protein AroP1 | 63 | 56 | 74 | 55 | 54 | 47 | 1 |
| 2052 | - | 2211845 | 2213182 | + | *nqrA* | *PA14_25280* | Na(+)-translocating NADH-quinone reductase subunit A | 119 | 136 | 147 | 154 | 141 | 122 | 1 |
| 2053 | - | 2213186 | 2214397 | + | *nqrB* | *PA14_25305* | Na(+)-translocating NADH-quinone reductase subunit B | 123 | 168 | 139 | 157 | 140 | 114 | 1 |
| 2054 | - | 2214390 | 2215175 | + | *nqrC* | *PA14_25320* | Na(+)-translocating NADH-quinone reductase subunit C | 153 | 184 | 151 | 147 | 127 | 121 | 1 |
| 2055 | - | 2215172 | 2215846 | + | *nqrD* | *PA14_25330* | Na(+)-translocating NADH-quinone reductase subunit D | 169 | 219 | 178 | 166 | 159 | 152 | 1 |
| 2056 | - | 2215846 | 2216454 | + | *nqrE* | *PA14_25340* | Na(+)-translocating NADH-quinone reductase subunit E | 169 | 188 | 166 | 155 | 154 | 150 | 1 |
| 2057 | - | 2216466 | 2217689 | + | *nqrF* | *PA14_25350* | Na(+)-translocating NADH-quinone reductase subunit F | 212 | 222 | 221 | 212 | 219 | 199 | 1 |
| 2058 | - | 2217682 | 2218710 | + | - | *PA14_25360* | thiamine biosynthesis lipoprotein | 225 | 223 | 232 | 239 | 235 | 223 | 1 |
| 2059 | - | 2218707 | 2218934 | + | - | *PA14_25370* | hypothetical protein | 210 | 274 | 228 | 278 | 243 | 247 | 1 |
| 2060 | - | 2219113 | 2220507 | + | *sth* | *PA14_25390* | soluble pyridine nucleotide transhydrogenase | 221 | 198 | 201 | 186 | 198 | 177 | 1 |
| 2061 | - | 2221272 | 2220550 | - | - | *PA14_25400* | phosphodiesterase | 118 | 121 | 104 | 103 | 102 | 99 | 1 |
| 2062 | - | 2221577 | 2221269 | - | - | *PA14_25410* | hypothetical protein | 37 | 40 | 40 | 30 | 28 | 28 | 1 |
| 2063 | - | 2222289 | 2223539 | + | - | *PA14_25430* | hypothetical protein | 68 | 43 | 73 | 53 | 52 | 52 | 1 |
| 2064 | - | 2222351 | 2221587 | - | - | *PA14_25420* | hypothetical protein | 44 | 34 | 49 | 39 | 34 | 40 | 1 |
| 2065 | - | 2223532 | 2224215 | + | - | *PA14_25440* | lipoprotein releasing system, ATP-binding protein | 127 | 122 | 127 | 111 | 128 | 104 | 1 |
| 2066 | - | 2224228 | 2225529 | + | - | *PA14_25450* | hypothetical protein | 46 | 37 | 44 | 32 | 42 | 36 | 1 |
| 2067 | - | 2226150 | 2225617 | - | - | *PA14_25470* | hypothetical protein | 31 | 28 | 30 | 18 | 21 | 21 | 1 |
| 2068 | - | 2226281 | 2228506 | + | - | *PA14_25480* | competence protein | 36 | 29 | 35 | 39 | 33 | 35 | 1 |
| 2069 | - | 2228579 | 2229214 | + | - | *PA14_25490* | tolQ-type transport protein | 67 | 58 | 70 | 61 | 53 | 55 | 1 |
| 2070 | - | 2229211 | 2229651 | + | - | *PA14_25500* | hypothetical protein | 169 | 191 | 185 | 182 | 169 | 140 | 1 |
| 2071 | - | 2229651 | 2230649 | + | *lpxK* | *PA14_25510* | tetraacyldisaccharide 4'-kinase | 89 | 95 | 101 | 92 | 90 | 69 | 1 |
| 2072 | - | 2230687 | 2230872 | + | - | *PA14_25520* | hypothetical protein | 93 | 109 | 103 | 97 | 85 | 100 | 1 |
| 2073 | - | 2230872 | 2231636 | + | *kdsB* | *PA14_25530* | 3-deoxy-manno-octulosonate cytidylyltransferase | 105 | 114 | 109 | 100 | 95 | 92 | 1 |
| 2074 | - | 2231636 | 2232100 | + | *ptpA* | *PA14_25540* | phosphotyrosine protein phosphatase | 130 | 127 | 132 | 124 | 123 | 119 | 1 |
| 2075 | - | 2232097 | 2233116 | + | *murB* | *PA14_25550* | UDP-N-acetylenolpyruvoylglucosamine reductase | 131 | 114 | 134 | 125 | 115 | 105 | 1 |
| 2076 | - | 2236447 | 2233274 | - | *rne* | *PA14_25560* | ribonuclease E | 155 | 172 | 147 | 147 | 151 | 132 | 1 |
| 2077 | - | 2237024 | 2237980 | + | *rluC* | *PA14_25580* | ribosomal large subunit pseudouridine synthase C | 305 | 228 | 356 | 292 | 323 | 320 | 1 |
| 2078 | - | 2237973 | 2238665 | + | - | *PA14_25590* | hydrolase | 262 | 227 | 308 | 272 | 273 | 273 | 1 |
| 2079 | - | 2238658 | 2239638 | + | - | *PA14_25600* | peptidase | 295 | 295 | 328 | 292 | 302 | 285 | 1 |
| 2080 | - | 2240299 | 2239721 | - | - | *PA14_25610* | Maf-like protein | 21 | 26 | 23 | 22 | 19 | 21 | 1 |
| 2081 | - | 2240409 | 2240945 | + | - | *PA14_25620* | hypothetical protein | 240 | 212 | 341 | 220 | 147 | 135 | 1 |
| 2082 | - | 2240959 | 2241141 | + | *rpmF* | *PA14_25630* | 50S ribosomal protein L32 | 197 | 291 | 228 | 162 | 162 | 106 | 1 |
| 2083 | - | 2241175 | 2242155 | + | *plsX* | *PA14_25640* | glycerol-3-phosphate acyltransferase PlsX | 53 | 63 | 64 | 42 | 46 | 35 | 1 |
| 2084 | - | 2242261 | 2243199 | + | *fabD* | *PA14_25650* | malonyl-CoA-ACP transacylase | 89 | 107 | 108 | 87 | 85 | 66 | 1 |
| 2085 | - | 2243222 | 2243965 | + | *fabG* | *PA14_25660* | 3-ketoacyl-ACP reductase | 462 | 612 | 537 | 361 | 404 | 340 | 1 |
| 2086 | - | 2244062 | 2244090 | ? | - | predicted RNA | - | 619 | 775 | 547 | 421 | 732 | 501 | 1 |
| 2087 | - | 2244161 | 2244397 | + | *acpP* | *PA14_25670* | acyl carrier protein | 600 | 896 | 540 | 356 | 565 | 398 | 1 |
| 2088 | - | 2244526 | 2245770 | + | *fabF1* | *PA14_25690* | 3-oxoacyl-ACP synthase | 183 | 210 | 168 | 145 | 190 | 142 | 1 |
| 2089 | - | 2245770 | 2246585 | + | *pabC* | *PA14_25710* | 4-amino-4-deoxychorismate lyase | 170 | 164 | 179 | 136 | 160 | 131 | 1 |
| 2090 | - | 2246591 | 2247640 | + | - | *PA14_25730* | hypothetical protein | 77 | 57 | 83 | 73 | 63 | 59 | 1 |
| 2091 | - | 2247669 | 2248301 | + | *tmk* | *PA14_25740* | thymidylate kinase | 123 | 116 | 135 | 123 | 128 | 111 | 1 |
| 2092 | - | 2248294 | 2249280 | + | *holB* | *PA14_25760* | DNA polymerase III subunit delta' | 113 | 120 | 125 | 102 | 122 | 106 | 1 |
| 2093 | - | 2249314 | 2249670 | + | *pilZ* | *PA14_25770* | type 4 fimbrial biogenesis protein PilZ | 100 | 87 | 129 | 103 | 95 | 100 | 1 |
| 2094 | - | 2249685 | 2250461 | + | - | *PA14_25780* | TatD family deoxyribonuclease | 67 | 66 | 81 | 65 | 53 | 55 | 1 |
| 2095 | - | 2251820 | 2250687 | - | - | *PA14_25790* | hypothetical protein | 47 | 44 | 49 | 48 | 36 | 44 | 1 |
| 2096 | - | 2251906 | 2252544 | + | - | *PA14_25800* | TetR family transcriptional regulator | 99 | 117 | 118 | 111 | 91 | 91 | 1 |
| 2097 | - | 2252711 | 2253607 | + | - | *PA14_25810* | hypothetical protein | 54 | 56 | 61 | 59 | 49 | 43 | 1 |
| 2098 | - | 2253672 | 2254304 | + | - | *PA14_25820* | lipoprotein | 40 | 34 | 44 | 45 | 37 | 32 | 1 |
| 2099 | - | 2254297 | 2254866 | + | - | *PA14_25830* | hypothetical protein | 27 | 35 | 33 | 34 | 27 | 27 | 1 |
| 2100 | - | 2256570 | 2254915 | - | - | *PA14_25840* | electron transfer flavoprotein-ubiquinone oxidoreductase | 128 | 116 | 129 | 88 | 104 | 100 | 1 |

| | A | B | C | D | E | F | G | H | I | J | K | L | M | N |
|---|---|---|---|---|---|---|---|---|---|---|---|---|---|---|
| 2101 | - | 2256891 | 2257640 | + | *etfB* | *PA14_25860* | electron transfer flavoprotein subunit beta | 382 | 547 | 322 | 251 | 408 | 252 | 1 |
| 2102 | - | 2257640 | 2258569 | + | *etfA* | *PA14_25880* | electron transfer flavoprotein subunit alpha | 775 | 865 | 691 | 586 | 713 | 520 | 1 |
| 2103 | - | 2258754 | 2259950 | + | - | *PA14_25900* | trans-2-enoyl-CoA reductase | 77 | 102 | 84 | 76 | 87 | 60 | 1 |
| 2104 | - | 2260970 | 2260023 | - | - | *PA14_25910* | lipase | 19 | 21 | 24 | 26 | 19 | 20 | 1 |
| 2105 | - | 2261851 | 2261099 | - | *cobM* | *PA14_25920* | precorrin-3 methylase | 22 | 24 | 24 | 25 | 26 | 24 | 1 |
| 2106 | - | 2262276 | 2261848 | - | - | *PA14_25930* | hypothetical protein | 16 | 14 | 15 | 14 | 14 | 13 | 1 |
| 2107 | - | 2263443 | 2262349 | - | - | *PA14_25940* | hypothetical protein | 57 | 65 | 58 | 52 | 59 | 61 | 1 |
| 2108 | - | 2263799 | 2264926 | + | - | *PA14_25960* | cobalamin biosynthesis protein cobW | 40 | 48 | 35 | 38 | 44 | 36 | 1 |
| 2109 | - | 2265002 | 2268748 | + | *cobN* | *PA14_25970* | cobaltochelatase subunit CobN | 43 | 46 | 37 | 43 | 47 | 43 | 1 |
| 2110 | - | 2269871 | 2268777 | - | *aroF* | *PA14_25980* | phospho-2-dehydro-3-deoxyheptonate aldolase | 21 | 21 | 21 | 18 | 22 | 15 | 1 |
| 2111 | - | 2270442 | 2271455 | + | - | *PA14_25990* | magnesium chelatase | 60 | 47 | 53 | 45 | 45 | 43 | 1 |
| 2112 | - | 2271476 | 2272120 | + | - | *PA14_26000* | magnesium chelatase | 60 | 46 | 55 | 53 | 49 | 49 | 1 |
| 2113 | - | 2272242 | 2273381 | + | - | *PA14_26010* | acyl-CoA thiolase | 32 | 31 | 35 | 40 | 30 | 34 | 1 |
| 2114 | - | 2275078 | 2273468 | - | - | *PA14_26020* | aminopeptidase | 97 | 165 | 119 | 197 | 157 | 224 | 1 |
| 2115 | - | 2275579 | 2277036 | + | - | *PA14_26050* | transporter | 17 | 21 | 26 | 31 | 19 | 25 | 1 |
| 2116 | - | 2277400 | 2277092 | - | - | *PA14_26060* | hypothetical protein | 34 | 53 | 56 | 47 | 34 | 57 | 1 |
| 2117 | - | 2278077 | 2277496 | - | - | *PA14_26070* | hypothetical protein | 4 | 4 | 5 | 3 | 3 | 3 | 1 |
| 2118 | - | 2278467 | 2278667 | + | - | *PA14_26080* | hypothetical protein | 6 | 5 | 3 | 5 | 3 | 4 | 1 |
| 2119 | - | 2279646 | 2278687 | - | - | *PA14_26090* | hydrolase | 5 | 6 | 7 | 5 | 6 | 5 | 1 |
| 2120 | - | 2280846 | 2279668 | - | - | *PA14_26110* | MFS transporter | 3 | 3 | 5 | 3 | 3 | 4 | 1 |
| 2121 | - | 2282021 | 2280912 | - | *morB* | *PA14_26130* | morphinone reductase | 10 | 9 | 9 | 9 | 6 | 7 | 1 |
| 2122 | - | 2282147 | 2282737 | + | - | *PA14_26140* | transcriptional regulator | 149 | 108 | 139 | 149 | 119 | 136 | 1 |
| 2123 | - | 2283674 | 2282748 | - | - | *PA14_26150* | transcriptional regulator | 52 | 44 | 56 | 39 | 42 | 45 | 1 |
| 2124 | - | 2283740 | 2284354 | + | - | *PA14_26160* | hypothetical protein | 4 | 3 | 3 | 4 | 4 | 3 | 1 |
| 2125 | - | 2284423 | 2285985 | + | - | *PA14_26165* | hypothetical protein | 23 | 25 | 26 | 28 | 23 | 23 | 1 |
| 2126 | - | 2287367 | 2286036 | - | - | *PA14_26190* | hypothetical protein | 35 | 30 | 32 | 41 | 33 | 34 | 1 |
| 2127 | - | 2287497 | 2287679 | + | - | *PA14_26200* | hypothetical protein | 17 | 20 | 25 | 16 | 14 | 13 | 1 |
| 2128 | - | 2288456 | 2287689 | - | *hisP* | *PA14_26210* | histidine transport | 7 | 8 | 9 | 7 | 8 | 6 | 1 |
| 2129 | - | 2289190 | 2288477 | - | *hisM* | *PA14_26220* | histidine ABC transporter, inner membrane permease | 11 | 11 | 11 | 12 | 12 | 10 | 1 |
| 2130 | - | 2289876 | 2289187 | - | *hisQ* | *PA14_26230* | histidine transport system permease | 6 | 5 | 7 | 5 | 6 | 6 | 1 |
| 2131 | - | 2290717 | 2289932 | - | *hisJ* | *PA14_26240* | periplasmic histidine-binding protein HisJ | 5 | 4 | 3 | 3 | 4 | 2 | 1 |
| 2132 | - | 2291946 | 2290777 | - | - | *PA14_26260* | hydrolase | 4 | 3 | 3 | 4 | 4 | 3 | 1 |
| 2133 | - | 2292047 | 2293036 | + | - | *PA14_26270* | transcriptional regulator | 10 | 10 | 12 | 12 | 9 | 12 | 1 |
| 2134 | - | 2293121 | 2294758 | + | - | *PA14_26280* | chemotaxis transducer | 128 | 130 | 180 | 171 | 162 | 181 | 1 |
| 2135 | - | 2295027 | 2294770 | - | - | *PA14_26300* | hypothetical protein | 89 | 110 | 132 | 157 | 139 | 166 | 1 |
| 2136 | - | 2295854 | 2295081 | - | - | *PA14_26310* | short chain dehydrogenase | 32 | 38 | 42 | 47 | 45 | 53 | 1 |
| 2137 | - | 2296065 | 2296901 | + | - | *PA14_26330* | AraC family transcriptional regulator | 22 | 28 | 34 | 42 | 28 | 28 | 1 |
| 2138 | - | 2296952 | 2297545 | + | - | *PA14_26340* | hypothetical protein | 5 | 10 | 10 | 9 | 8 | 5 | 0.272776094 |
| 2139 | - | 2297665 | 2298531 | + | - | *PA14_26350* | hypothetical protein | 136 | 128 | 118 | 127 | 119 | 147 | 1 |
| 2140 | - | 2299563 | 2298532 | - | - | *PA14_26360* | ABC transporter permease | 56 | 47 | 51 | 62 | 53 | 70 | 1 |
| 2141 | - | 2300531 | 2299560 | - | - | *PA14_26390* | hypothetical protein | 15 | 14 | 15 | 18 | 15 | 21 | 1 |
| 2142 | - | 2301301 | 2300528 | - | - | *PA14_26400* | ABC transporter ATP-binding protein | 10 | 11 | 12 | 14 | 10 | 14 | 1 |
| 2143 | - | 2303510 | 2301354 | - | - | *PA14_26420* | TonB-dependent receptor | 15 | 15 | 19 | 20 | 15 | 15 | 1 |
| 2144 | - | 2304147 | 2304716 | + | - | *PA14_26450* | hypothetical protein | 4 | 6 | 6 | 8 | 5 | 5 | 1 |
| 2145 | - | 2305498 | 2304770 | - | *cobK* | *PA14_26460* | cobalt-precorrin-6x reductase | 49 | 39 | 46 | 41 | 45 | 38 | 1 |
| 2146 | - | 2306595 | 2305495 | - | *cbiD* | *PA14_26470* | cobalt-precorrin-6A synthase | 33 | 35 | 33 | 32 | 32 | 29 | 1 |
| 2147 | - | 2307835 | 2306588 | - | *cobL* | *PA14_26480* | precorrin-6y-dependent methyltransferase CobL | 33 | 33 | 31 | 30 | 30 | 26 | 1 |
| 2148 | - | 2308172 | 2309632 | + | - | *PA14_26485* | oxidoreductase | 36 | 30 | 34 | 26 | 28 | 27 | 1 |
| 2149 | - | 2309629 | 2310255 | + | *cobH* | *PA14_26500* | precorrin-8X methylmutase | 29 | 35 | 33 | 26 | 32 | 23 | 1 |
| 2150 | - | 2310252 | 2311004 | + | *cobI* | *PA14_26510* | precorrin-2 C(20)-methyltransferase | 39 | 49 | 40 | 34 | 43 | 36 | 1 |
| 2151 | - | 2310997 | 2312676 | + | *cobJ* | *PA14_26530* | precorrin-3 methylase CobJ | 55 | 61 | 56 | 47 | 63 | 48 | 1 |
| 2152 | - | 2312746 | 2313594 | + | - | *PA14_26540* | hypothetical protein | 70 | 68 | 76 | 72 | 59 | 59 | 1 |
| 2153 | - | 2313591 | 2313950 | + | - | *PA14_26550* | lipoprotein | 143 | 153 | 186 | 166 | 124 | 117 | 1 |
| 2154 | - | 2313950 | 2314759 | + | - | *PA14_26560* | outer membrane protein | 150 | 157 | 162 | 145 | 122 | 119 | 1 |
| 2155 | - | 2314953 | 2315591 | + | - | *PA14_26570* | transcriptional regulator | 90 | 66 | 86 | 85 | 79 | 88 | 1 |
| 2156 | - | 2315677 | 2316045 | + | - | *PA14_26580* | hypothetical protein | 14 | 8 | 10 | 9 | 8 | 9 | 0.283677495 |
| 2157 | - | 2316267 | 2317706 | + | - | *PA14_26590* | GntR family transcriptional regulator | 138 | 128 | 138 | 140 | 133 | 151 | 1 |
| 2158 | - | 2318037 | 2318621 | + | - | *PA14_26600* | RNA polymerase sigma factor | 88 | 78 | 97 | 75 | 80 | 78 | 1 |
| 2159 | - | 2318660 | 2319382 | + | - | *PA14_26610* | hypothetical protein | 126 | 103 | 157 | 115 | 116 | 122 | 1 |
| 2160 | - | 2319728 | 2320150 | + | - | *PA14_26620* | hypothetical protein | 72 | 83 | 92 | 96 | 80 | 89 | 1 |
| 2161 | - | 2321990 | 2320164 | - | - | *PA14_26640* | long-chain-acyl-CoA synthetase | 30 | 30 | 32 | 35 | 27 | 33 | 1 |
| 2162 | - | 2322977 | 2322153 | - | - | *PA14_26650* | short chain dehydrogenase | 13 | 14 | 13 | 14 | 12 | 17 | 1 |
| 2163 | - | 2325014 | 2323029 | - | - | *PA14_26670* | biotin carboxylase | 21 | 16 | 15 | 17 | 13 | 21 | 1 |
| 2164 | - | 2325854 | 2325060 | - | - | *PA14_26690* | enoyl-CoA hydratase/isomerase | 15 | 12 | 10 | 11 | 10 | 13 | 1 |
| 2165 | - | 2327035 | 2325875 | - | - | *PA14_26700* | acyl-CoA dehydrogenase | 22 | 25 | 18 | 24 | 21 | 30 | 1 |
| 2166 | - | 2328768 | 2327152 | - | - | *PA14_26720* | biotin-dependent carboxylase | 20 | 16 | 15 | 18 | 13 | 20 | 1 |
| 2167 | - | 2329648 | 2328770 | - | - | *PA14_26730* | short-chain dehydrogenase | 19 | 16 | 14 | 18 | 14 | 19 | 1 |
| 2168 | - | 2331523 | 2329721 | - | - | *PA14_26750* | hypothetical protein | 11 | 9 | 8 | 11 | 9 | 11 | 1 |
| 2169 | - | 2331804 | 2332400 | + | - | *PA14_26760* | transcriptional regulator | 205 | 192 | 195 | 164 | 199 | 180 | 1 |
| 2170 | - | 2333165 | 2332401 | - | - | *PA14_26770* | hypothetical protein | 65 | 70 | 90 | 69 | 78 | 75 | 1 |
| 2171 | - | 2333565 | 2333398 | - | - | *PA14_26780* | hypothetical protein | 694 | 774 | 947 | 775 | 829 | 767 | 1 |
| 2172 | - | 2333988 | 2335103 | + | - | *PA14_26810* | two-component sensor | 3 | 3 | 5 | 3 | 3 | 2 | 1 |
| 2173 | - | 2335105 | 2336016 | + | - | *PA14_26830* | two-component response regulator | 4 | 5 | 6 | 4 | 4 | 4 | 1 |
| 2174 | - | 2336540 | 2336025 | - | - | *PA14_26850* | hypothetical protein | 3 | 4 | 5 | 4 | 4 | 4 | 1 |
| 2175 | - | 2336587 | 2337477 | + | - | *PA14_26860* | LysR family transcriptional regulator | 102 | 102 | 88 | 73 | 103 | 98 | 1 |
| 2176 | - | 2338141 | 2337500 | - | - | *PA14_26870* | hypothetical protein | 47 | 41 | 47 | 35 | 49 | 48 | 1 |
| 2177 | - | 2338187 | 2339080 | + | - | *PA14_26880* | LysR family transcriptional regulator | 30 | 28 | 30 | 26 | 26 | 29 | 1 |
| 2178 | - | 2339177 | 2339875 | + | *pyrF* | *PA14_26890* | orotidine 5'-phosphate decarboxylase | 36 | 36 | 40 | 35 | 32 | 28 | 1 |
| 2179 | - | 2340124 | 2341041 | + | - | *PA14_26910* | hypothetical protein | 42 | 43 | 51 | 50 | 40 | 47 | 1 |
| 2180 | - | 2341041 | 2341997 | + | - | *PA14_26920* | hypothetical protein | 44 | 55 | 46 | 51 | 51 | 43 | 1 |
| 2181 | - | 2341994 | 2344000 | + | - | *PA14_26930* | transglutaminase | 41 | 47 | 42 | 51 | 47 | 42 | 1 |
| 2182 | - | 2343997 | 2344767 | + | - | *PA14_26940* | hypothetical protein | 73 | 82 | 86 | 81 | 71 | 70 | 1 |
| 2183 | - | 2344848 | 2345645 | + | - | *PA14_26960* | hypothetical protein | 59 | 58 | 65 | 67 | 63 | 51 | 1 |
| 2184 | - | 2347231 | 2345654 | - | - | *PA14_26970* | hypothetical protein | 29 | 28 | 34 | 32 | 27 | 30 | 1 |

| | A | B | C | D | E | F | G | H | I | J | K | L | M | N |
|---|---|---|---|---|---|---|---|---|---|---|---|---|---|---|
| 2185 | - | 2347729 | 2347241 | - | - | PA14_26980 | hypothetical protein | 16 | 11 | 15 | 11 | 10 | 17 | 1 |
| 2186 | - | 2348175 | 2347816 | - | - | PA14_26990 | hypothetical protein | 55 | 24 | 24 | 21 | 18 | 64 | 0.020457471 |
| 2187 | - | 2349773 | 2348301 | - | - | PA14_27000 | chemotaxis transducer | 223 | 181 | 261 | 182 | 148 | 198 | 1 |
| 2188 | - | 2350715 | 2349912 | - | mttC | PA14_27020 | secretion protein MttC | 27 | 22 | 29 | 25 | 22 | 25 | 1 |
| 2189 | - | 2350962 | 2352392 | + | - | PA14_27050 | Slt family transglycosylase | 37 | 30 | 43 | 36 | 31 | 33 | 1 |
| 2190 | - | 2352491 | 2352925 | + | - | PA14_27070 | hypothetical protein | 80 | 96 | 98 | 84 | 80 | 89 | 1 |
| 2191 | - | 2353959 | 2353093 | - | lipH | PA14_27090 | lipase chaperone | 5 | 8 | 8 | 8 | 8 | 10 | 1 |
| 2192 | - | 2355101 | 2354166 | - | lipA | PA14_27100 | lactonizing lipase | 5 | 9 | 7 | 7 | 7 | 6 | 1 |
| 2193 | - | 2355845 | 2355297 | - | ligT | PA14_27110 | 2'-5' RNA ligase | 7 | 6 | 8 | 5 | 5 | 7 | 1 |
| 2194 | - | 2355899 | 2356339 | + | - | PA14_27120 | hypothetical protein | 66 | 65 | 74 | 79 | 63 | 61 | 1 |
| 2195 | - | 2356858 | 2356352 | - | greB | PA14_27130 | transcription elongation factor GreB | 65 | 65 | 68 | 56 | 58 | 53 | 1 |
| 2196 | - | 2359402 | 2356910 | - | - | PA14_27140 | hypothetical protein | 40 | 37 | 37 | 38 | 36 | 32 | 1 |
| 2197 | - | 2360094 | 2359411 | - | - | PA14_27150 | ABC transporter ATP-binding protein | 41 | 28 | 36 | 27 | 25 | 26 | 1 |
| 2198 | - | 2360105 | 2360710 | + | tesA | PA14_27160 | acyl-CoA thioesterase | 71 | 80 | 87 | 88 | 80 | 68 | 1 |
| 2199 | - | 2360766 | 2361044 | + | - | PA14_27170 | hypothetical protein | 25 | 29 | 26 | 26 | 29 | 23 | 1 |
| 2200 | - | 2361122 | 2362093 | + | - | PA14_27180 | hypothetical protein | 189 | 142 | 191 | 135 | 139 | 164 | 1 |
| 2201 | - | 2362974 | 2362888 | - | - | PA14_27190 | Ser tRNA | 110 | 70 | 128 | 96 | 39 | 48 | 0.601884591 |
| 2202 | - | 2363130 | 2364260 | + | - | PA14_27200 | hypothetical protein | 42 | 30 | 39 | 34 | 30 | 29 | 1 |
| 2203 | - | 2364303 | 2364869 | + | efp | PA14_27210 | elongation factor P | 137 | 192 | 135 | 85 | 109 | 81 | 1 |
| 2204 | - | 2365385 | 2364957 | - | ohr | PA14_27220 | organic hydroperoxide resistance protein | 19 | 28 | 17 | 15 | 18 | 12 | 1 |
| 2205 | - | 2365985 | 2365530 | - | - | PA14_27230 | MarR family transcriptional regulator | 22 | 26 | 20 | 20 | 15 | 16 | 1 |
| 2206 | - | 2367090 | 2366071 | - | - | PA14_27250 | LysR family transcriptional regulator | 12 | 9 | 10 | 9 | 11 | 10 | 1 |
| 2207 | - | 2367128 | 2367877 | + | - | PA14_27270 | hypothetical protein | 36 | 24 | 30 | 30 | 31 | 29 | 1 |
| 2208 | - | 2368733 | 2367879 | - | - | PA14_27280 | LysR family transcriptional regulator | 30 | 29 | 26 | 25 | 28 | 28 | 1 |
| 2209 | - | 2368830 | 2369063 | + | - | PA14_27290 | hypothetical protein | 5 | 7 | 8 | 5 | 7 | 7 | 1 |
| 2210 | - | 2369146 | 2370354 | + | - | PA14_27310 | hypothetical protein | 29 | 29 | 34 | 34 | 27 | 30 | 1 |
| 2211 | - | 2371741 | 2370395 | - | - | PA14_27330 | phospho-2-dehydro-3-deoxyheptonate aldolase | 67 | 38 | 47 | 47 | 43 | 55 | 1 |
| 2212 | - | 2371917 | 2372666 | + | - | PA14_27350 | hypothetical protein | 40 | 45 | 47 | 49 | 43 | 36 | 1 |
| 2213 | - | 2373484 | 2372693 | - | - | PA14_27360 | enoyl-CoA hydratase | 35 | 38 | 43 | 40 | 35 | 32 | 1 |
| 2214 | - | 2373795 | 2375498 | + | - | PA14_27370 | ATP-dependent RNA helicase | 22 | 15 | 40 | 16 | 16 | 13 | 1 |
| 2215 | - | 2376635 | 2375865 | - | - | PA14_27390 | hypothetical protein | 11 | 12 | 12 | 7 | 10 | 10 | 1 |
| 2216 | - | 2377697 | 2376726 | - | - | PA14_27400 | LysR family transcriptional regulator | 11 | 12 | 10 | 11 | 7 | 10 | 1 |
| 2217 | - | 2379173 | 2377734 | - | - | PA14_27410 | outer membrane protein | 4 | 3 | 5 | 5 | 3 | 3 | 1 |
| 2218 | - | 2380234 | 2379170 | - | - | PA14_27420 | secretion protein | 5 | 5 | 7 | 8 | 4 | 6 | 1 |
| 2219 | - | 2381838 | 2380246 | - | - | PA14_27430 | multidrug efflux MFS transporter | 3 | 3 | 3 | 5 | 2 | 4 | 1 |
| 2220 | - | 2382941 | 2381976 | - | - | PA14_27440 | transcriptional regulator | 11 | 9 | 11 | 9 | 7 | 11 | 1 |
| 2221 | - | 2383065 | 2383511 | + | - | PA14_27450 | hypothetical protein | 31 | 32 | 31 | 30 | 27 | 24 | 1 |
| 2222 | - | 2383567 | 2384223 | + | tpm | PA14_27460 | thiopurine S-methyltransferase | 148 | 135 | 122 | 152 | 123 | 110 | 1 |
| 2223 | - | 2385382 | 2384255 | - | - | PA14_27470 | zinc carboxypeptidase | 46 | 49 | 45 | 43 | 45 | 37 | 1 |
| 2224 | - | 2386397 | 2385522 | - | htpX | PA14_27480 | heat shock protein HtpX | 196 | 183 | 230 | 125 | 174 | 145 | 1 |
| 2225 | - | 2386983 | 2386531 | - | - | PA14_27490 | hypothetical protein | 15 | 16 | 22 | 16 | 12 | 16 | 1 |
| 2226 | - | 2388245 | 2387034 | - | - | PA14_27500 | aminotransferase | 28 | 27 | 29 | 27 | 23 | 22 | 1 |
| 2227 | - | 2388430 | 2388828 | + | - | PA14_27510 | methionine sulfoxide reductase B | 256 | 254 | 269 | 245 | 204 | 264 | 1 |
| 2228 | - | 2388940 | 2389425 | + | - | PA14_27520 | glutathione peroxidase | 58 | 79 | 60 | 55 | 64 | 60 | 1 |
| 2229 | - | 2389422 | 2389913 | + | - | PA14_27530 | MarR family transcriptional regulator | 85 | 94 | 79 | 73 | 85 | 89 | 1 |
| 2230 | - | 2392292 | 2389932 | - | - | PA14_27550 | sensor/response regulator hybrid | 61 | 55 | 68 | 70 | 61 | 68 | 1 |
| 2231 | - | 2392375 | 2393265 | + | - | PA14_27560 | hypothetical protein | 48 | 51 | 48 | 47 | 40 | 39 | 1 |
| 2232 | - | 2393262 | 2393744 | + | - | PA14_27570 | GAF domain-containing protein | 108 | 120 | 103 | 111 | 93 | 92 | 1 |
| 2233 | - | 2394416 | 2393754 | - | - | PA14_27580 | glutathione S-transferase | 66 | 78 | 67 | 70 | 65 | 66 | 1 |
| 2234 | - | 2395251 | 2394478 | - | - | PA14_27590 | hypothetical protein | 77 | 64 | 96 | 53 | 62 | 72 | 1 |
| 2235 | - | 2395469 | 2395541 | + | - | PA14_27600 | Glu tRNA | 173 | 26 | 57 | 28 | 22 | 21 | 4.60686E-27 |
| 2236 | - | 2395547 | 2395619 | + | - | PA14_27610 | Gly tRNA | 245 | 101 | 202 | 98 | 49 | 70 | 0.00634821 |
| 2237 | - | 2395707 | 2395779 | + | - | PA14_27620 | Gly tRNA | 112 | 46 | 95 | 48 | 21 | 33 | 0.003088724 |
| 2238 | - | 2396312 | 2395992 | - | - | PA14_27630 | protein associated with synthesis and assembly of refractile inclusion bodies | 5 | 7 | 4 | 5 | 4 | 6 | 1 |
| 2239 | - | 2396695 | 2396375 | - | - | PA14_27640 | protein associated with synthesis and assembly of refractile inclusion bodies | 18 | 25 | 23 | 23 | 17 | 21 | 1 |
| 2240 | - | 2397391 | 2396846 | - | - | PA14_27650 | hypothetical protein | 18 | 17 | 21 | 22 | 15 | 19 | 1 |
| 2241 | - | 2397684 | 2397445 | - | - | PA14_27660 | hypothetical protein | 3 | 5 | 4 | 3 | 3 | 2 | 0.655993826 |
| 2242 | - | 2400054 | 2397913 | - | - | PA14_27675 | hypothetical protein | 9 | 9 | 9 | 10 | 8 | 9 | 1 |
| 2243 | - | 2400737 | 2400105 | - | - | PA14_27680 | hypothetical protein | 4 | 3 | 4 | 5 | 3 | 3 | 1 |
| 2244 | - | 2401672 | 2401079 | - | - | PA14_27690 | RNA polymerase sigma factor | 30 | 28 | 32 | 38 | 32 | 32 | 1 |
| 2245 | - | 2402463 | 2401738 | - | - | PA14_27700 | transcriptional regulator | 67 | 68 | 87 | 90 | 65 | 67 | 1 |
| 2246 | - | 2403137 | 2402727 | - | - | PA14_27710 | hypothetical protein | 43 | 45 | 55 | 47 | 42 | 39 | 1 |
| 2247 | - | 2403576 | 2403196 | - | - | PA14_27720 | hypothetical protein | 24 | 24 | 28 | 22 | 20 | 21 | 1 |
| 2248 | - | 2406152 | 2403705 | - | fadE | PA14_27730 | acyl-CoA dehydrogenase | 151 | 159 | 164 | 178 | 155 | 170 | 1 |
| 2249 | - | 2406305 | 2406976 | + | - | PA14_27740 | hypothetical protein | 38 | 38 | 47 | 29 | 42 | 43 | 1 |
| 2250 | - | 2407091 | 2407711 | + | - | PA14_27755 | glutathione S-transferase | 200 | 191 | 232 | 244 | 191 | 201 | 1 |
| 2251 | - | 2407787 | 2408719 | + | - | PA14_27770 | ABC transporter ATP-binding protein | 43 | 48 | 47 | 43 | 44 | 43 | 1 |
| 2252 | - | 2408716 | 2409495 | + | - | PA14_27780 | ABC transporter permease | 54 | 65 | 55 | 55 | 70 | 50 | 1 |
| 2253 | - | 2410876 | 2409545 | - | - | PA14_27800 | two-component sensor | 43 | 38 | 47 | 48 | 36 | 35 | 1 |
| 2254 | - | 2411553 | 2410873 | - | - | PA14_27810 | two-component response regulator | 28 | 26 | 24 | 27 | 22 | 19 | 1 |
| 2255 | - | 2411679 | 2411870 | + | - | PA14_27830 | hypothetical protein | 8 | 11 | 10 | 14 | 8 | 8 | 1 |
| 2256 | - | 2412042 | 2412659 | + | - | PA14_27840 | hypothetical protein | 6 | 6 | 7 | 7 | 5 | 4 | 1 |
| 2257 | - | 2412777 | 2413607 | + | queF | PA14_27850 | 7-cyano-7-deazaguanine reductase | 21 | 19 | 22 | 19 | 17 | 19 | 1 |
| 2258 | - | 2413937 | 2413674 | - | - | PA14_27870 | hypothetical protein | 110 | 166 | 140 | 86 | 128 | 110 | 1 |
| 2259 | - | 2414592 | 2414011 | - | - | PA14_27880 | hypothetical protein | 16 | 12 | 16 | 18 | 15 | 17 | 1 |
| 2260 | - | 2415332 | 2414589 | - | - | PA14_27890 | hypothetical protein | 12 | 12 | 16 | 14 | 9 | 11 | 1 |
| 2261 | - | 2415466 | 2416185 | + | - | PA14_27900 | transcriptional regulator | 43 | 42 | 48 | 47 | 37 | 36 | 1 |
| 2262 | - | 2416591 | 2416187 | - | - | PA14_27910 | hypothetical protein | 33 | 34 | 29 | 34 | 28 | 30 | 1 |
| 2263 | - | 2416694 | 2417398 | + | - | PA14_27920 | hypothetical protein | 114 | 107 | 123 | 110 | 103 | 125 | 1 |
| 2264 | - | 2417755 | 2417456 | - | - | PA14_27930 | hypothetical protein | 132 | 142 | 159 | 133 | 151 | 162 | 1 |
| 2265 | - | 2418001 | 2419185 | + | - | PA14_27940 | two-component response regulator | 200 | 170 | 205 | 199 | 175 | 184 | 1 |
| 2266 | - | 2419182 | 2419664 | + | - | PA14_27950 | hypothetical protein | 265 | 271 | 320 | 317 | 308 | 296 | 1 |
| 2267 | - | 2420673 | 2419750 | - | tal | PA14_27960 | transaldolase B | 105 | 117 | 84 | 85 | 110 | 100 | 1 |
| 2268 | - | 2421751 | 2420753 | - | - | PA14_27980 | tRNA-dihydrouridine synthase A | 21 | 17 | 21 | 18 | 18 | 19 | 1 |

| | A | B | C | D | E | F | G | H | I | J | K | L | M | N |
|---|---|---|---|---|---|---|---|---|---|---|---|---|---|---|
| 2269 | - | 2422054 | 2423370 | + | - | *PA14_27990* | sialidase | 52 | 51 | 49 | 70 | 54 | 62 | 1 |
| 2270 | - | 2423812 | 2424852 | + | - | *PA14_28000* | hypothetical protein | 16 | 15 | 19 | 15 | 14 | 13 | 1 |
| 2271 | - | 2424932 | 2425528 | + | - | *PA14_28010* | hypothetical protein | 10 | 8 | 9 | 7 | 11 | 6 | 1 |
| 2272 | - | 2425622 | 2425912 | + | - | *PA14_28020* | hypothetical protein | 15 | 10 | 16 | 11 | 12 | 14 | 1 |
| 2273 | - | 2426498 | 2426013 | - | - | *PA14_28030* | hypothetical protein | 133 | 105 | 133 | 103 | 115 | 120 | 1 |
| 2274 | - | 2428012 | 2426933 | - | - | *PA14_28040* | hypothetical protein | 6 | 7 | 7 | 5 | 4 | 4 | 1 |
| 2275 | - | 2428256 | 2429851 | + | - | *PA14_28050* | chemotaxis transducer | 327 | 239 | 283 | 339 | 409 | 428 | 1 |
| 2276 | - | 2431111 | 2429873 | - | *cpg2* | *PA14_28060* | glutamate carboxypeptidase | 109 | 78 | 90 | 132 | 113 | 137 | 1 |
| 2277 | - | 2431645 | 2431178 | - | - | *PA14_28070* | hypothetical protein | 7 | 7 | 7 | 11 | 8 | 10 | 1 |
| 2278 | - | 2431978 | 2431760 | - | - | *PA14_28080* | transcriptional regulator | 7 | 4 | 4 | 6 | 4 | 3 | 1 |
| 2279 | - | 2432544 | 2431987 | - | - | *PA14_28090* | hypothetical protein | 6 | 7 | 7 | 9 | 6 | 9 | 1 |
| 2280 | - | 2434523 | 2432715 | - | - | *PA14_28100* | hypothetical protein | 10 | 8 | 11 | 10 | 9 | 9 | 1 |
| 2281 | - | 2435260 | 2434601 | - | - | *PA14_28110* | hypothetical protein | 9 | 7 | 11 | 10 | 9 | 10 | 1 |
| 2282 | - | 2436079 | 2435738 | - | - | *PA14_28120* | hypothetical protein | 23 | 26 | 26 | 31 | 20 | 23 | 1 |
| 2283 | - | 2436420 | 2436076 | - | - | *PA14_28130* | hypothetical protein | 15 | 12 | 11 | 15 | 11 | 12 | 1 |
| 2284 | - | 2436828 | 2437226 | + | - | *PA14_28140* | hypothetical protein | 109 | 115 | 130 | 116 | 137 | 145 | 1 |
| 2285 | - | 2437310 | 2438188 | + | - | *PA14_28150* | hypothetical protein | 55 | 49 | 72 | 56 | 56 | 65 | 1 |
| 2286 | - | 2438270 | 2439226 | + | - | *PA14_28170* | formate/nitrate transporter | 72 | 84 | 97 | 96 | 83 | 79 | 1 |
| 2287 | - | 2440560 | 2439277 | - | - | *PA14_28180* | hypothetical protein | 51 | 65 | 58 | 53 | 53 | 55 | 1 |
| 2288 | - | 2440744 | 2440817 | + | - | *PA14_28190* | Val tRNA | 2 | 0 | 2 | 0 | 0 | 0 | 0.007235124 |
| 2289 | - | 2441012 | 2441449 | + | - | *PA14_28200* | hypothetical protein | 27 | 18 | 31 | 30 | 17 | 16 | 0.87182855 |
| 2290 | - | 2441454 | 2442041 | + | - | *PA14_28210* | hypothetical protein | 43 | 35 | 41 | 30 | 38 | 29 | 1 |
| 2291 | - | 2442240 | 2442761 | + | - | *PA14_28220* | hypothetical protein | 25 | 25 | 23 | 27 | 21 | 18 | 1 |
| 2292 | - | 2442951 | 2442781 | - | - | *PA14_28230* | hypothetical protein | 27 | 38 | 28 | 32 | 24 | 21 | 1 |
| 2293 | - | 2443112 | 2443336 | + | - | *PA14_28240* | hypothetical protein | 71 | 78 | 89 | 103 | 70 | 66 | 1 |
| 2294 | 2444398 | 2444509 | 2443370 | - | - | *PA14_28250* | secreted acid phosphatase | 62 | 67 | 78 | 77 | 62 | 55 | 1 |
| 2295 | 2444398 | 2444516 | 2444812 | + | - | *PA14_28260* | hypothetical protein | 113 | 91 | 98 | 144 | 107 | 164 | 1 |
| 2296 | - | 2444914 | 2445693 | + | - | *PA14_28280* | PhzF family phenazine biosynthesis protein | 59 | 59 | 53 | 53 | 61 | 52 | 1 |
| 2297 | - | 2445715 | 2446125 | + | - | *PA14_28290* | hypothetical protein | 43 | 58 | 60 | 57 | 54 | 52 | 1 |
| 2298 | - | 2446747 | 2446265 | - | - | *PA14_28300* | hypothetical protein | 8 | 9 | 6 | 8 | 7 | 7 | 1 |
| 2299 | - | 2447721 | 2446753 | - | - | *PA14_28310* | enoyl-CoA hydratase | 10 | 8 | 12 | 9 | 7 | 8 | 1 |
| 2300 | - | 2447809 | 2448390 | + | - | *PA14_28320* | TetR family transcriptional regulator | 14 | 21 | 17 | 19 | 21 | 21 | 1 |
| 2301 | - | 2448449 | 2448544 | + | - | *PA14_28330* | hypothetical protein | 13 | 14 | 13 | 16 | 15 | 15 | 1 |
| 2302 | - | 2448732 | 2449631 | + | - | *PA14_28340* | hypothetical protein | 51 | 68 | 46 | 46 | 58 | 38 | 1 |
| 2303 | - | 2450374 | 2449679 | - | - | *PA14_28350* | hypothetical protein | 24 | 33 | 28 | 33 | 27 | 27 | 1 |
| 2304 | - | 2450705 | 2450914 | + | - | *PA14_28360* | hypothetical protein | 77 | 13 | 34 | 46 | 36 | 92 | 1.69482E-25 |
| 2305 | - | 2451388 | 2451597 | + | - | *PA14_28370* | hypothetical protein | 12 | 9 | 12 | 12 | 10 | 12 | 1 |
| 2306 | - | 2451709 | 2452053 | + | - | *PA14_28380* | hypothetical protein | 20 | 18 | 19 | 21 | 23 | 21 | 1 |
| 2307 | - | 2452512 | 2452096 | - | - | *PA14_28390* | hypothetical protein | 10 | 10 | 12 | 13 | 14 | 10 | 1 |
| 2308 | - | 2453901 | 2452624 | - | - | *PA14_28400* | outer membrane OprD family porin | 179 | 252 | 185 | 134 | 160 | 137 | 1 |
| 2309 | - | 2454266 | 2454580 | + | - | *PA14_28410* | hypothetical protein | 103 | 101 | 150 | 67 | 27 | 38 | 1 |
| 2310 | - | 2455678 | 2454791 | - | - | *PA14_28420* | LysR family transcriptional regulator | 12 | 17 | 15 | 13 | 12 | 14 | 1 |
| 2311 | - | 2455781 | 2456224 | + | - | *PA14_28430* | hypothetical protein | 3 | 7 | 5 | 3 | 3 | 3 | 0.515742977 |
| 2312 | - | 2456363 | 2456797 | + | - | *PA14_28440* | hypothetical protein | 577 | 429 | 578 | 679 | 559 | 537 | 1 |
| 2313 | - | 2457319 | 2456849 | - | *eco* | *PA14_28450* | ecotin | 58 | 101 | 57 | 47 | 59 | 33 | 0.680948229 |
| 2314 | - | 2457898 | 2457668 | - | - | *PA14_28460* | hypothetical protein | 94 | 116 | 111 | 115 | 92 | 98 | 1 |
| 2315 | - | 2458077 | 2458274 | + | - | *PA14_28470* | hypothetical protein | 58 | 79 | 76 | 48 | 64 | 61 | 1 |
| 2316 | - | 2458669 | 2458340 | - | - | *PA14_28490* | hypothetical protein | 240 | 280 | 285 | 154 | 225 | 236 | 1 |
| 2317 | - | 2459154 | 2458780 | - | - | *PA14_28500* | hypothetical protein | 702 | 779 | 802 | 358 | 414 | 679 | 1 |
| 2318 | - | 2459289 | 2459732 | + | - | *PA14_28510* | hypothetical protein | 15 | 14 | 16 | 15 | 10 | 11 | 1 |
| 2319 | - | 2459780 | 2460205 | + | - | *PA14_28520* | hypothetical protein | 51 | 42 | 53 | 57 | 48 | 50 | 1 |
| 2320 | - | 2461094 | 2460195 | - | - | *PA14_28530* | hypothetical protein | 33 | 25 | 35 | 35 | 31 | 30 | 1 |
| 2321 | - | 2461317 | 2461550 | + | - | *PA14_28540* | hypothetical protein | 16 | 18 | 16 | 15 | 11 | 14 | 1 |
| 2322 | - | 2462339 | 2461821 | - | - | *PA14_28560* | hypothetical protein | 65 | 52 | 71 | 70 | 53 | 57 | 1 |
| 2323 | - | 2463071 | 2462358 | - | *endA* | *PA14_28570* | DNA-specific endonuclease I | 43 | 35 | 40 | 50 | 39 | 45 | 1 |
| 2324 | - | 2463228 | 2463449 | + | - | *PA14_28580* | hypothetical protein | 25 | 18 | 25 | 26 | 23 | 23 | 1 |
| 2325 | - | 2463446 | 2464228 | + | - | *PA14_28590* | methionine aminopeptidase | 36 | 35 | 46 | 43 | 31 | 34 | 1 |
| 2326 | - | 2464714 | 2465001 | + | - | *PA14_28600* | hypothetical protein | 373 | 286 | 336 | 288 | 210 | 280 | 1 |
| 2327 | - | 2465503 | 2465327 | - | - | *PA14_28610* | hypothetical protein | 243 | 267 | 310 | 255 | 264 | 272 | 1 |
| 2328 | - | 2465914 | 2465588 | - | - | *PA14_28620* | hypothetical protein | 65 | 79 | 108 | 77 | 76 | 63 | 1 |
| 2329 | - | 2466106 | 2466969 | + | - | *PA14_28630* | hydrolase | 29 | 19 | 29 | 24 | 18 | 19 | 1 |
| 2330 | - | 2467219 | 2469141 | + | *thrS* | *PA14_28650* | threonyl-tRNA synthetase | 148 | 176 | 148 | 133 | 145 | 124 | 1 |
| 2331 | - | 2469159 | 2469692 | + | *infC* | *PA14_28660* | translation initiation factor IF-3 | 620 | 696 | 604 | 543 | 531 | 479 | 1 |
| 2332 | - | 2469754 | 2469948 | + | *rpmI* | *PA14_28670* | 50S ribosomal protein L35 | 1416 | 1689 | 1057 | 1084 | 1267 | 900 | 1 |
| 2333 | - | 2469972 | 2470328 | + | *rplT* | *PA14_28680* | 50S ribosomal protein L20 | 1346 | 1424 | 1150 | 910 | 1089 | 731 | 1 |
| 2334 | - | 2470426 | 2471442 | + | *pheS* | *PA14_28690* | phenylalanyl-tRNA synthetase subunit alpha | 53 | 77 | 55 | 47 | 58 | 38 | 1 |
| 2335 | - | 2471477 | 2473855 | + | *pheT* | *PA14_28710* | phenylalanyl-tRNA synthetase subunit beta | 113 | 129 | 100 | 93 | 102 | 85 | 1 |
| 2336 | - | 2473859 | 2474161 | + | *ihfA* | *PA14_28720* | integration host factor subunit alpha | 3260 | 2838 | 3279 | 2459 | 2122 | 2316 | 1 |
| 2337 | - | 2474142 | 2474498 | + | - | *PA14_28730* | transcriptional regulator | 725 | 819 | 646 | 600 | 558 | 598 | 1 |
| 2338 | - | 2474588 | 2474661 | + | - | *PA14_28740* | Pro tRNA | 50 | 57 | 33 | 43 | 35 | 32 | 1 |
| 2339 | - | 2475199 | 2474825 | - | - | *PA14_28750* | hypothetical protein | 27 | 25 | 26 | 39 | 24 | 25 | 1 |
| 2340 | - | 2476088 | 2475474 | - | - | *PA14_28760* | hypothetical protein | 26 | 25 | 23 | 28 | 21 | 18 | 1 |
| 2341 | - | 2476671 | 2478314 | + | - | *PA14_28770* | hypothetical protein | 25 | 23 | 20 | 26 | 21 | 26 | 1 |
| 2342 | - | 2478590 | 2478874 | + | - | *PA14_28780* | hypothetical protein | 8 | 7 | 8 | 9 | 7 | 8 | 1 |
| 2343 | - | 2478975 | 2479136 | + | - | *PA14_28790* | hypothetical protein | 16 | 13 | 14 | 13 | 8 | 12 | 1 |
| 2344 | - | 2479279 | 2480274 | + | - | *PA14_28800* | hypothetical protein | 69 | 49 | 49 | 55 | 51 | 52 | 1 |
| 2345 | - | 2480409 | 2485532 | + | - | *PA14_28810* | DNA helicase | 203 | 169 | 164 | 195 | 182 | 192 | 1 |
| 2346 | - | 2485638 | 2488136 | + | - | *PA14_28820* | hypothetical protein | 228 | 148 | 168 | 214 | 197 | 214 | 1 |
| 2347 | - | 2488133 | 2489428 | + | - | *PA14_28830* | hypothetical protein | 415 | 311 | 301 | 377 | 342 | 383 | 1 |
| 2348 | - | 2489403 | 2491613 | + | - | *PA14_28840* | helicase | 258 | 217 | 206 | 248 | 254 | 262 | 1 |
| 2349 | - | 2491674 | 2491988 | + | - | *PA14_28850* | hypothetical protein | 1139 | 1147 | 928 | 1309 | 1112 | 1302 | 1 |
| 2350 | - | 2492021 | 2492317 | + | - | *PA14_28870* | hypothetical protein | 461 | 400 | 471 | 525 | 446 | 553 | 1 |
| 2351 | - | 2493942 | 2492593 | - | - | *PA14_28880* | hypothetical protein | 40 | 30 | 28 | 23 | 33 | 30 | 1 |
| 2352 | - | 2500235 | 2493939 | - | - | *PA14_28895* | hypothetical protein | 41 | 32 | 29 | 27 | 38 | 31 | 1 |

| | A | B | C | D | E | F | G | H | I | J | K | L | M | N |
|---|---|---|---|---|---|---|---|---|---|---|---|---|---|---|
| 2353 | - | 2500888 | 2500250 | - | - | *PA14_28910* | radical activating enzyme | 53 | 45 | 55 | 51 | 45 | 46 | 1 |
| 2354 | - | 2502773 | 2500890 | - | - | *PA14_28920* | chaperone | 43 | 39 | 36 | 33 | 41 | 33 | 1 |
| 2355 | - | 2503392 | 2502850 | - | - | *PA14_28930* | hypothetical protein | 10 | 6 | 7 | 7 | 9 | 8 | 0.895687725 |
| 2356 | - | 2503745 | 2503467 | - | - | *PA14_28940* | hypothetical protein | 63 | 54 | 61 | 36 | 54 | 67 | 1 |
| 2357 | - | 2504167 | 2503775 | - | - | *PA14_28950* | hypothetical protein | 19 | 15 | 20 | 25 | 18 | 26 | 1 |
| 2358 | - | 2504678 | 2504199 | - | - | *PA14_28960* | hypothetical protein | 28 | 26 | 37 | 37 | 25 | 33 | 1 |
| 2359 | - | 2504847 | 2505386 | + | - | *PA14_28970* | hypothetical protein | 18 | 32 | 13 | 55 | 21 | 12 | 0.828318293 |
| 2360 | - | 2505374 | 2506324 | + | - | *PA14_28980* | Fe2+-dicitrate sensor | 2 | 9 | 4 | 25 | 4 | 3 | 5.77642E-08 |
| 2361 | - | 2506753 | 2507391 | + | - | *PA14_28990* | hypothetical protein | 14 | 13 | 19 | 15 | 14 | 14 | 1 |
| 2362 | - | 2508237 | 2507551 | - | - | *PA14_29000* | hypothetical protein | 20 | 19 | 18 | 23 | 16 | 26 | 1 |
| 2363 | - | 2508825 | 2508337 | - | - | *PA14_29010* | MerR family transcriptional regulator | 47 | 38 | 50 | 46 | 45 | 45 | 1 |
| 2364 | - | 2509058 | 2509888 | + | *cpo* | *PA14_29020* | chloroperoxidase | 25 | 27 | 29 | 29 | 27 | 35 | 1 |
| 2365 | - | 2510015 | 2511250 | + | - | *PA14_29030* | FMN oxidoreductase | 20 | 17 | 19 | 18 | 17 | 19 | 1 |
| 2366 | - | 2511603 | 2511265 | - | - | *PA14_29040* | ferredoxin | 14 | 18 | 12 | 17 | 19 | 18 | 1 |
| 2367 | - | 2513929 | 2511620 | - | - | *PA14_29050* | molybdopterin oxidoreductase | 14 | 12 | 14 | 13 | 11 | 11 | 1 |
| 2368 | - | 2514283 | 2514762 | + | - | *PA14_29060* | transcriptional regulator | 71 | 59 | 62 | 67 | 62 | 58 | 1 |
| 2369 | - | 2515673 | 2514810 | - | - | *PA14_29070* | hypothetical protein | 31 | 27 | 33 | 31 | 27 | 29 | 1 |
| 2370 | - | 2517184 | 2516093 | - | - | *PA14_29090* | periplasmic spermidine/putrescine-binding protein | 11 | 9 | 10 | 9 | 10 | 11 | 1 |
| 2371 | - | 2517997 | 2517383 | - | - | *PA14_29100* | hypothetical protein | 37 | 45 | 36 | 41 | 42 | 35 | 1 |
| 2372 | - | 2519030 | 2518056 | - | *cysK* | *PA14_29110* | cysteine synthase A | 115 | 137 | 106 | 85 | 110 | 87 | 1 |
| 2373 | - | 2519151 | 2520236 | + | - | *PA14_29120* | hypothetical protein | 38 | 41 | 47 | 44 | 47 | 49 | 1 |
| 2374 | - | 2520399 | 2521244 | + | - | *PA14_29130* | ATPase | 107 | 163 | 100 | 107 | 154 | 142 | 1 |
| 2375 | - | 2521309 | 2521704 | + | - | *PA14_29150* | hypothetical protein | 93 | 124 | 81 | 91 | 103 | 116 | 1 |
| 2376 | - | 2521753 | 2522934 | + | - | *PA14_29160* | hypothetical protein | 118 | 152 | 100 | 137 | 132 | 155 | 1 |
| 2377 | - | 2523048 | 2524067 | + | - | *PA14_29180* | AraC family transcriptional regulator | 18 | 19 | 18 | 19 | 15 | 19 | 1 |
| 2378 | - | 2524582 | 2524349 | - | - | *PA14_29190* | hypothetical protein | 8 | 11 | 9 | 9 | 11 | 7 | 1 |
| 2379 | - | 2525068 | 2524592 | - | - | *PA14_29200* | hypothetical protein | 29 | 33 | 34 | 29 | 33 | 29 | 1 |
| 2380 | - | 2526804 | 2525215 | - | - | *PA14_29210* | MFS transporter | 8 | 8 | 11 | 15 | 6 | 9 | 1 |
| 2381 | - | 2528114 | 2526807 | - | - | *PA14_29220* | porin | 6 | 5 | 6 | 7 | 4 | 6 | 1 |
| 2382 | - | 2530001 | 2528163 | - | - | *PA14_29230* | hypothetical protein | 16 | 19 | 17 | 19 | 19 | 23 | 1 |
| 2383 | - | 2530609 | 2531289 | + | - | *PA14_29240* | hydrolase | 15 | 19 | 12 | 14 | 17 | 14 | 1 |
| 2384 | - | 2531346 | 2531627 | + | - | *PA14_29250* | hypothetical protein | 10 | 10 | 9 | 10 | 9 | 10 | 1 |
| 2385 | - | 2531797 | 2532690 | + | - | *PA14_29260* | transcriptional regulator | 16 | 16 | 16 | 21 | 14 | 18 | 1 |
| 2386 | - | 2532804 | 2533907 | + | - | *PA14_29270* | outer membrane lipoprotein | 23 | 22 | 22 | 25 | 19 | 19 | 1 |
| 2387 | - | 2533988 | 2534314 | + | - | *PA14_29280* | thioredoxin | 47 | 46 | 57 | 46 | 39 | 43 | 1 |
| 2388 | - | 2534842 | 2534342 | - | - | *PA14_29290* | hypothetical protein | 13 | 10 | 13 | 13 | 14 | 15 | 1 |
| 2389 | - | 2535536 | 2535012 | - | - | *PA14_29300* | transcriptional regulator | 23 | 21 | 25 | 26 | 23 | 21 | 1 |
| 2390 | - | 2535805 | 2537010 | + | - | *PA14_29320* | NADH dehydrogenase, FAD-containing subunit | 38 | 63 | 48 | 67 | 46 | 39 | 0.831807123 |
| 2391 | - | 2538037 | 2537045 | - | - | *PA14_29330* | hypothetical protein | 115 | 110 | 143 | 143 | 117 | 148 | 1 |
| 2392 | - | 2539410 | 2538496 | - | - | *PA14_29340* | hypothetical protein | 3 | 3 | 3 | 3 | 2 | 3 | 1 |
| 2393 | - | 2541670 | 2539430 | - | *pfeA* | *PA14_29350* | outer membrane receptor FepA | 14 | 12 | 15 | 15 | 11 | 12 | 1 |
| 2394 | - | 2543109 | 2541769 | - | *pfeS* | *PA14_29360* | two-component sensor PfeS | 16 | 33 | 19 | 73 | 49 | 18 | 0.292421048 |
| 2395 | - | 2544083 | 2546152 | + | - | *PA14_29390* | hypothetical protein | 124 | 95 | 106 | 107 | 117 | 89 | 1 |
| 2396 | - | 2546202 | 2550155 | + | - | *PA14_29400* | hypothetical protein | 96 | 75 | 69 | 59 | 88 | 66 | 1 |
| 2397 | - | 2550506 | 2551468 | + | - | *PA14_29410* | serine/threonine dehydratase | 55 | 48 | 44 | 33 | 47 | 41 | 1 |
| 2398 | - | 2552717 | 2551476 | - | - | *PA14_29420* | hypothetical protein | 24 | 21 | 23 | 23 | 23 | 21 | 1 |
| 2399 | - | 2553829 | 2552900 | - | - | *PA14_29440* | LysR family transcriptional regulator | 14 | 15 | 13 | 14 | 16 | 13 | 1 |
| 2400 | - | 2554826 | 2553852 | - | - | *PA14_29460* | quinone oxidoreductase | 62 | 46 | 75 | 77 | 48 | 57 | 1 |
| 2401 | - | 2555052 | 2555804 | + | - | *PA14_29470* | hypothetical protein | 217 | 109 | 153 | 113 | 92 | 97 | 1 |
| 2402 | - | 2556397 | 2557200 | + | - | *PA14_29480* | ABC-2 transporter permease | 8 | 7 | 7 | 9 | 7 | 8 | 1 |
| 2403 | - | 2557190 | 2558917 | + | - | *PA14_29490* | type II secretion protein | 9 | 11 | 10 | 13 | 10 | 10 | 1 |
| 2404 | - | 2558922 | 2560109 | + | - | *PA14_29500* | type II secretion system protein | 5 | 5 | 4 | 6 | 4 | 4 | 1 |
| 2405 | - | 2560163 | 2560597 | + | - | *PA14_29510* | type II secretion system protein | 9 | 7 | 6 | 7 | 8 | 6 | 1 |
| 2406 | - | 2560566 | 2560976 | + | - | *PA14_29520* | type II secretion system protein | 9 | 8 | 7 | 9 | 8 | 7 | 1 |
| 2407 | - | 2560976 | 2561401 | + | - | *PA14_29530* | type II secretion system protein | 12 | 9 | 10 | 12 | 9 | 9 | 1 |
| 2408 | - | 2561398 | 2561988 | + | - | *PA14_29540* | type II secretion system protein | 4 | 5 | 3 | 3 | 5 | 3 | 1 |
| 2409 | - | 2561978 | 2563057 | + | - | *PA14_29550* | hypothetical protein | 10 | 9 | 9 | 11 | 9 | 10 | 1 |
| 2410 | - | 2563101 | 2564087 | + | - | *PA14_29560* | hypothetical protein | 15 | 15 | 16 | 20 | 14 | 16 | 1 |
| 2411 | - | 2564087 | 2564671 | + | - | *PA14_29570* | hypothetical protein | 11 | 10 | 12 | 10 | 10 | 9 | 1 |
| 2412 | - | 2564914 | 2565231 | + | - | *PA14_29575* | hypothetical protein | 24 | 29 | 24 | 19 | 25 | 22 | 1 |
| 2413 | - | 2565660 | 2565307 | - | - | *PA14_29590* | transcriptional regulator | 107 | 184 | 85 | 82 | 126 | 89 | 0.793216635 |
| 2414 | - | 2566324 | 2565968 | - | - | *PA14_29600* | 6-pyruvoyl-tetrahydropterin synthase | 31 | 31 | 39 | 33 | 29 | 26 | 1 |
| 2415 | 2568048 | 2567978 | 2566425 | - | - | *PA14_29620* | anaerobic nitric oxide reductase transcriptional regulator | 45 | 62 | 60 | 52 | 44 | 37 | 1 |
| 2416 | 2568048 | 2568133 | 2569314 | + | *fhp* | *PA14_29640* | nitric oxide dioxygenase | 10 | 404 | 9 | 336 | 550 | 40 | 0 |
| 2417 | 2568048 | 2569368 | 2569625 | + | - | *PA14_29650* | hypothetical protein | 28 | 491 | 29 | 172 | 291 | 63 | 1.4453E-232 |
| 2418 | 2568048 | 2569612 | 2570805 | + | - | *PA14_29660* | hypothetical protein | 50 | 565 | 52 | 293 | 491 | 142 | 7.35411E-59 |
| 2419 | - | 2571949 | 2571056 | - | - | *PA14_29680* | hypothetical protein | 21 | 118 | 24 | 58 | 106 | 45 | 1.38852E-14 |
| 2420 | - | 2573058 | 2571976 | - | - | *PA14_29690* | hypothetical protein | 15 | 28 | 20 | 19 | 27 | 20 | 0.415011252 |
| 2421 | - | 2573273 | 2573581 | + | - | *PA14_29710* | hypothetical protein | 131 | 147 | 136 | 120 | 122 | 124 | 1 |
| 2422 | - | 2573581 | 2573895 | + | - | *PA14_29720* | hypothetical protein | 53 | 69 | 66 | 56 | 55 | 53 | 1 |
| 2423 | - | 2573895 | 2574566 | + | - | *PA14_29730* | two-component response regulator | 79 | 77 | 86 | 76 | 69 | 70 | 1 |
| 2424 | - | 2574563 | 2575900 | + | - | *PA14_29740* | two-component sensor | 65 | 63 | 80 | 72 | 63 | 63 | 1 |
| 2425 | - | 2575991 | 2576299 | + | - | *PA14_29750* | hypothetical protein | 45 | 61 | 50 | 56 | 54 | 51 | 1 |
| 2426 | - | 2578448 | 2576304 | - | - | *PA14_29760* | chemotaxis transducer | 70 | 89 | 84 | 82 | 89 | 77 | 1 |
| 2427 | - | 2578707 | 2580032 | + | - | *PA14_29770* | transporter | 20 | 19 | 23 | 20 | 16 | 20 | 1 |
| 2428 | - | 2580146 | 2581831 | + | - | *PA14_29800* | chemotaxis transducer | 69 | 88 | 80 | 73 | 83 | 72 | 1 |
| 2429 | - | 2581991 | 2583049 | + | - | *PA14_29820* | hypothetical protein | 66 | 51 | 67 | 50 | 54 | 47 | 1 |
| 2430 | - | 2584065 | 2583256 | - | - | *PA14_29830* | methyltransferase | 21 | 19 | 24 | 23 | 20 | 21 | 1 |
| 2431 | - | 2585640 | 2584180 | - | *nuoN* | *PA14_29850* | NADH dehydrogenase subunit N | 160 | 173 | 148 | 143 | 153 | 149 | 1 |
| 2432 | - | 2587177 | 2585648 | - | *nuoM* | *PA14_29860* | NADH dehydrogenase subunit M | 191 | 211 | 189 | 177 | 183 | 180 | 1 |
| 2433 | - | 2589052 | 2587205 | - | *nuoL* | *PA14_29880* | NADH dehydrogenase subunit L | 163 | 186 | 167 | 153 | 169 | 162 | 1 |
| 2434 | - | 2589357 | 2589049 | - | *nuoK* | *PA14_29890* | NADH dehydrogenase subunit K | 164 | 167 | 158 | 132 | 159 | 162 | 1 |
| 2435 | - | 2589903 | 2589403 | - | *nuoJ* | *PA14_29900* | NADH dehydrogenase subunit J | 195 | 219 | 202 | 177 | 179 | 160 | 1 |
| 2436 | - | 2590463 | 2589915 | - | *nuoI* | *PA14_29920* | NADH dehydrogenase subunit I | 226 | 265 | 205 | 220 | 227 | 233 | 1 |

| | A | B | C | D | E | F | G | H | I | J | K | L | M | N |
|---|---|---|---|---|---|---|---|---|---|---|---|---|---|---|
| 2437 | - | 2591470 | 2590475 | - | *nuoH* | *PA14_29930* | NADH dehydrogenase subunit H | 223 | 239 | 215 | 192 | 219 | 210 | 1 |
| 2438 | - | 2594184 | 2591467 | - | *nuoG* | *PA14_29940* | NADH dehydrogenase subunit G | 270 | 311 | 271 | 269 | 253 | 257 | 1 |
| 2439 | - | 2595665 | 2594316 | - | *nuoF* | *PA14_29970* | NADH dehydrogenase I subunit F | 324 | 332 | 307 | 295 | 257 | 277 | 1 |
| 2440 | - | 2596162 | 2595662 | - | *nuoE* | *PA14_29980* | NADH dehydrogenase subunit E | 305 | 344 | 291 | 315 | 300 | 289 | 1 |
| 2441 | - | 2597945 | 2596164 | - | *nuoD* | *PA14_29990* | bifunctional NADH:ubiquinone oxidoreductase subunit C/D | 204 | 247 | 199 | 214 | 214 | 200 | 1 |
| 2442 | - | 2598703 | 2598026 | - | *nuoB* | *PA14_30010* | NADH dehydrogenase subunit B | 370 | 406 | 397 | 300 | 277 | 308 | 1 |
| 2443 | - | 2599127 | 2598714 | - | *nuoA* | *PA14_30020* | NADH dehydrogenase subunit A | 174 | 124 | 200 | 109 | 116 | 128 | 1 |
| 2444 | - | 2600281 | 2599727 | - | - | *PA14_30030* | hypothetical protein | 7 | 9 | 9 | 9 | 10 | 8 | 1 |
| 2445 | - | 2602378 | 2600360 | - | - | *PA14_30040* | hypothetical protein | 20 | 18 | 22 | 24 | 16 | 18 | 1 |
| 2446 | - | 2604153 | 2602558 | - | - | *PA14_30050* | isocitrate lyase | 145 | 163 | 105 | 60 | 101 | 74 | 1 |
| 2447 | - | 2605569 | 2604748 | - | - | *PA14_30070* | hypothetical protein | 91 | 99 | 114 | 106 | 109 | 123 | 1 |
| 2448 | - | 2606186 | 2605566 | - | - | *PA14_30080* | hypothetical protein | 62 | 64 | 78 | 62 | 64 | 68 | 1 |
| 2449 | - | 2606618 | 2606193 | - | - | *PA14_30090* | acetyl transferase | 285 | 275 | 325 | 245 | 235 | 280 | 1 |
| 2450 | - | 2607780 | 2606611 | - | - | *PA14_30100* | hypothetical protein | 113 | 99 | 149 | 64 | 58 | 95 | 1 |
| 2451 | - | 2609210 | 2607840 | - | *purB* | *PA14_30110* | adenylosuccinate lyase | 66 | 62 | 81 | 75 | 54 | 62 | 1 |
| 2452 | - | 2610176 | 2609286 | - | - | *PA14_30130* | hypothetical protein | 30 | 33 | 38 | 43 | 31 | 32 | 1 |
| 2453 | - | 2610799 | 2610179 | - | - | *PA14_30140* | hypothetical protein | 79 | 76 | 90 | 82 | 79 | 65 | 1 |
| 2454 | - | 2611923 | 2610796 | - | *mnmA* | *PA14_30150* | tRNA-specific 2-thiouridylase MnmA | 103 | 83 | 122 | 92 | 80 | 84 | 1 |
| 2455 | - | 2612437 | 2611967 | - | - | *PA14_30160* | hypothetical protein | 23 | 26 | 24 | 25 | 23 | 23 | 1 |
| 2456 | - | 2614749 | 2612524 | - | *idh* | *PA14_30180* | monomeric isocitrate dehydrogenase | 138 | 126 | 108 | 71 | 101 | 73 | 1 |
| 2457 | - | 2615108 | 2616364 | + | *icd* | *PA14_30190* | isocitrate dehydrogenase | 1036 | 1457 | 940 | 673 | 987 | 701 | 1 |
| 2458 | - | 2616710 | 2616438 | - | *cspD* | *PA14_30200* | cold-shock protein CspD | 339 | 395 | 381 | 415 | 329 | 330 | 1 |
| 2459 | - | 2616936 | 2617304 | + | *clpS* | *PA14_30210* | ATP-dependent Clp protease adaptor protein ClpS | 389 | 385 | 534 | 551 | 511 | 614 | 1 |
| 2460 | - | 2617305 | 2617319 | ? | - | predicted RNA | - | 671 | 611 | 817 | 1073 | 963 | 1145 | 1 |
| 2461 | - | 2617332 | 2619608 | + | *clpA* | *PA14_30230* | ATP-dependent Clp protease, ATP-binding subunit ClpA | 787 | 766 | 827 | 901 | 862 | 980 | 1 |
| 2462 | - | 2619908 | 2619690 | - | *infA* | *PA14_30240* | translation initiation factor IF-1 | 30 | 45 | 44 | 36 | 41 | 38 | 1 |
| 2463 | - | 2620720 | 2620013 | - | - | *PA14_30260* | arginyl-tRNA-protein transferase | 80 | 98 | 105 | 73 | 71 | 96 | 1 |
| 2464 | - | 2621455 | 2620775 | - | *aat* | *PA14_30270* | leucyl/phenylalanyl-tRNA--protein transferase | 26 | 28 | 27 | 23 | 21 | 23 | 1 |
| 2465 | - | 2622443 | 2621493 | - | *trxB1* | *PA14_30280* | thioredoxin reductase 1 | 99 | 127 | 93 | 85 | 89 | 78 | 1 |
| 2466 | - | 2622671 | 2625106 | + | *ftsK* | *PA14_30290* | cell division protein FtsK | 193 | 179 | 195 | 214 | 189 | 200 | 1 |
| 2467 | - | 2625132 | 2625758 | + | *lolA* | *PA14_30310* | outer-membrane lipoprotein carrier protein | 134 | 146 | 142 | 126 | 128 | 140 | 1 |
| 2468 | - | 2625768 | 2627093 | + | - | *PA14_30320* | recombination factor protein RarA | 202 | 176 | 198 | 178 | 189 | 194 | 1 |
| 2469 | - | 2627215 | 2628495 | + | *serS* | *PA14_30330* | seryl-tRNA synthetase | 148 | 153 | 138 | 119 | 128 | 135 | 1 |
| 2470 | - | 2628497 | 2629894 | + | *cysG* | *PA14_30340* | siroheme synthase | 247 | 224 | 230 | 214 | 226 | 226 | 1 |
| 2471 | - | 2630873 | 2629899 | - | - | *PA14_30350* | hypothetical protein | 40 | 43 | 41 | 46 | 38 | 41 | 1 |
| 2472 | - | 2631944 | 2630961 | - | - | *PA14_30360* | hypothetical protein | 76 | 77 | 76 | 76 | 64 | 75 | 1 |
| 2473 | - | 2632276 | 2631941 | - | - | *PA14_30370* | hypothetical protein | 71 | 71 | 95 | 61 | 63 | 68 | 1 |
| 2474 | - | 2632578 | 2632273 | - | - | *PA14_30380* | hypothetical protein | 62 | 53 | 72 | 44 | 53 | 52 | 1 |
| 2475 | - | 2632937 | 2632578 | - | - | *PA14_30390* | sulfur relay protein TusC | 123 | 110 | 134 | 102 | 106 | 100 | 1 |
| 2476 | - | 2633329 | 2632934 | - | - | *PA14_30400* | sulfur transfer complex subunit TusD | 111 | 99 | 123 | 86 | 104 | 100 | 1 |
| 2477 | - | 2634108 | 2633440 | - | - | *PA14_30410* | hypothetical protein | 259 | 289 | 267 | 149 | 194 | 191 | 1 |
| 2478 | - | 2634238 | 2634324 | + | - | *PA14_30420* | Ser tRNA | 37 | 26 | 31 | 25 | 15 | 9 | 1 |
| 2479 | - | 2636045 | 2634462 | - | - | *PA14_30430* | thiosulfate sulfurtransferase | 29 | 33 | 29 | 30 | 30 | 32 | 1 |
| 2480 | - | 2636647 | 2636042 | - | - | *PA14_30440* | hypothetical protein | 9 | 13 | 14 | 10 | 13 | 13 | 1 |
| 2481 | - | 2636760 | 2637656 | + | - | *PA14_30450* | LysR family transcriptional regulator | 31 | 31 | 30 | 30 | 33 | 34 | 1 |
| 2482 | - | 2637996 | 2639063 | + | - | *PA14_30460* | flavin-dependent oxidoreductase | 10 | 10 | 9 | 9 | 11 | 10 | 1 |
| 2483 | - | 2639073 | 2640017 | + | - | *PA14_30470* | periplasmic aliphatic sulfonate-binding protein | 10 | 12 | 11 | 11 | 10 | 10 | 1 |
| 2484 | - | 2640027 | 2641109 | + | - | *PA14_30490* | lavin-dependent oxidoreductase | 9 | 9 | 8 | 7 | 7 | 7 | 1 |
| 2485 | - | 2641114 | 2642265 | + | - | *PA14_30500* | acyl-CoA dehydrogenase | 18 | 17 | 17 | 20 | 16 | 16 | 1 |
| 2486 | - | 2642373 | 2643359 | + | - | *PA14_30520* | periplasmic aliphatic sulfonate-binding protein | 7 | 6 | 7 | 6 | 5 | 5 | 1 |
| 2487 | - | 2643371 | 2644327 | + | - | *PA14_30540* | periplasmic aliphatic sulfonate-binding protein | 13 | 8 | 8 | 10 | 9 | 8 | 0.678622091 |
| 2488 | - | 2644483 | 2645442 | + | - | *PA14_30550* | periplasmic aliphatic sulfonate-binding protein | 28 | 23 | 23 | 27 | 25 | 23 | 1 |
| 2489 | - | 2646081 | 2645509 | - | - | *PA14_30560* | hypothetical protein | 31 | 21 | 23 | 28 | 23 | 26 | 1 |
| 2490 | 2647472 | 2647394 | 2646291 | - | - | *PA14_30570* | periplasmic spermidine/putrescine-binding protein | 143 | 81 | 100 | 139 | 110 | 127 | 1 |
| 2491 | 2647472 | 2647547 | 2648353 | + | - | *PA14_30580* | LuxR family transcriptional regulator | 219 | 279 | 277 | 363 | 278 | 235 | 1 |
| 2492 | - | 2648473 | 2651127 | + | - | *PA14_30590* | outer membrane receptor protein | 11 | 10 | 14 | 12 | 9 | 9 | 1 |
| 2493 | - | 2651138 | 2652352 | + | - | *PA14_30600* | permease | 20 | 13 | 13 | 22 | 15 | 28 | 1 |
| 2494 | - | 2653396 | 2652368 | - | - | *PA14_30620* | AraC family transcriptional regulator | 165 | 88 | 100 | 147 | 108 | 217 | 1 |
| 2495 | - | 2654014 | 2655162 | + | *pqsH* | *PA14_30630* | FAD-dependent monooxygenase | 392 | 406 | 451 | 504 | 418 | 395 | 1 |
| 2496 | - | 2655504 | 2656148 | + | *gacA* | *PA14_30650* | response regulator GacA | 430 | 332 | 533 | 487 | 392 | 487 | 1 |
| 2497 | - | 2656149 | 2657975 | + | *uvrC* | *PA14_30660* | excinuclease ABC subunit C | 85 | 72 | 90 | 82 | 74 | 84 | 1 |
| 2498 | - | 2658009 | 2658569 | + | *pgsA* | *PA14_30670* | CDP-diacylglycerol--glycerol-3-phosphate 3-phosphatidyltransferase | 33 | 34 | 36 | 30 | 27 | 25 | 1 |
| 2499 | - | 2658638 | 2658710 | + | - | *PA14_30680* | Gly tRNA | 87 | 39 | 68 | 32 | 16 | 24 | 0.032235375 |
| 2500 | - | 2660507 | 2659452 | - | - | *PA14_30690* | hypothetical protein | 30 | 25 | 24 | 35 | 31 | 27 | 1 |
| 2501 | - | 2661144 | 2664122 | + | - | *PA14_30700* | sensor/response regulator hybrid | 74 | 69 | 77 | 74 | 78 | 73 | 1 |
| 2502 | - | 2664213 | 2664746 | + | - | *PA14_30710* | osmoprotectant transporter activator protein | 240 | 236 | 232 | 231 | 201 | 233 | 1 |
| 2503 | - | 2665032 | 2665102 | + | - | *PA14_30720* | Cys tRNA | 58 | 31 | 55 | 31 | 13 | 14 | 0.503413992 |
| 2504 | - | 2666538 | 2665498 | - | - | *PA14_30730* | hypothetical protein | 53 | 44 | 46 | 51 | 55 | 41 | 1 |
| 2505 | - | 2666962 | 2667552 | + | - | *PA14_30740* | NADPH specific quinone oxidoreductase | 6 | 4 | 6 | 5 | 5 | 4 | 0.940967938 |
| 2506 | - | 2667877 | 2668743 | + | - | *PA14_30750* | tryptophan oxygenase | 37 | 32 | 39 | 41 | 34 | 37 | 1 |
| 2507 | - | 2669335 | 2668775 | - | - | *PA14_30760* | acetyltransferase | 24 | 29 | 24 | 20 | 25 | 21 | 1 |
| 2508 | - | 2669787 | 2669350 | - | - | *PA14_30770* | AsnC family transcriptional regulator | 51 | 49 | 42 | 34 | 42 | 40 | 1 |
| 2509 | - | 2669920 | 2670819 | + | - | *PA14_30790* | hypothetical protein | 39 | 29 | 38 | 31 | 28 | 32 | 1 |
| 2510 | - | 2671487 | 2670885 | - | - | *PA14_30800* | hypothetical protein | 79 | 88 | 51 | 38 | 46 | 46 | 1 |
| 2511 | - | 2671834 | 2672982 | + | *alkB1* | *PA14_30810* | alkane-1 monooxygenase | 23 | 36 | 29 | 32 | 22 | 27 | 1 |
| 2512 | - | 2673094 | 2674701 | + | - | *PA14_30820* | methyl-accepting chemotaxis transducer | 173 | 164 | 215 | 223 | 161 | 227 | 1 |
| 2513 | - | 2676055 | 2674712 | - | - | *PA14_30830* | two-component response regulator | 123 | 139 | 162 | 150 | 141 | 174 | 1 |
| 2514 | 2677542 | 2676132 | 2677544 | + | - | *PA14_30840* | signal transduction histidine kinase | 159 | 127 | 180 | 195 | 134 | 210 | 1 |
| 2515 | - | 2677870 | 2678133 | ? | - | predicted RNA | - | 1087 | 686 | 773 | 842 | 882 | 817 | 1 |
| 2516 | - | 2679706 | 2678435 | - | - | *PA14_30850* | TrbI-like protein | 7 | 5 | 9 | 7 | 6 | 10 | 1 |
| 2517 | - | 2680698 | 2679709 | - | - | *PA14_30860* | TrbG-like protein | 4 | 4 | 4 | 4 | 4 | 3 | 1 |
| 2518 | - | 2681399 | 2680695 | - | - | *PA14_30870* | conjugal transfer protein TrbF | 11 | 17 | 12 | 18 | 13 | 16 | 1 |
| 2519 | - | 2682782 | 2681412 | - | - | *PA14_30880* | conjugal transfer protein TrbL | 4 | 4 | 4 | 5 | 3 | 4 | 1 |
| 2520 | - | 2682980 | 2683123 | + | - | *PA14_30890* | hypothetical protein | 11 | 9 | 7 | 8 | 8 | 11 | 1 |

| | A | B | C | D | E | F | G | H | I | J | K | L | M | N |
|---|---|---|---|---|---|---|---|---|---|---|---|---|---|---|
| 2521 | - | 2683836 | 2683111 | - | - | *PA14_30900* | conjugal transfer protein TrbJ | 5 | 4 | 4 | 4 | 4 | 4 | 1 |
| 2522 | - | 2686286 | 2683833 | - | - | *PA14_30910* | conjugal transfer ATPase TrbE | 4 | 5 | 3 | 4 | 3 | 4 | 1 |
| 2523 | - | 2686960 | 2686568 | - | - | *PA14_30930* | TrbC-like protein | 1 | 4 | 1 | 2 | 3 | 2 | 0.055881303 |
| 2524 | - | 2688027 | 2686957 | - | - | *PA14_30940* | conjugal transfer protein | 6 | 6 | 5 | 7 | 7 | 6 | 1 |
| 2525 | - | 2688488 | 2688024 | - | - | *PA14_30950* | hypothetical protein | 17 | 16 | 17 | 14 | 14 | 12 | 1 |
| 2526 | - | 2690485 | 2688485 | - | - | *PA14_30960* | conjugal transfer coupling protein TraG | 12 | 13 | 14 | 17 | 10 | 13 | 1 |
| 2527 | - | 2692000 | 2691005 | - | - | *PA14_30970* | transcriptional regulator | 42 | 44 | 46 | 36 | 41 | 41 | 1 |
| 2528 | - | 2692432 | 2692127 | - | - | *PA14_30980* | hypothetical protein | 165 | 181 | 162 | 169 | 138 | 134 | 1 |
| 2529 | - | 2693151 | 2692483 | - | - | *PA14_30990* | hypothetical protein | 149 | 139 | 132 | 141 | 131 | 121 | 1 |
| 2530 | - | 2693907 | 2693164 | - | - | *PA14_31000* | hypothetical protein | 67 | 65 | 63 | 60 | 72 | 68 | 1 |
| 2531 | - | 2697041 | 2693904 | - | - | *PA14_31010* | cation efflux system protein | 30 | 32 | 28 | 29 | 30 | 31 | 1 |
| 2532 | - | 2698281 | 2697052 | - | - | *PA14_31030* | cation efflux system protein | 34 | 37 | 32 | 32 | 30 | 32 | 1 |
| 2533 | - | 2699525 | 2698278 | - | - | *PA14_31040* | cation efflux system protein | 24 | 26 | 25 | 24 | 23 | 24 | 1 |
| 2534 | - | 2700664 | 2699666 | - | - | *PA14_31050* | hypothetical protein | 176 | 141 | 199 | 177 | 184 | 191 | 1 |
| 2535 | - | 2701278 | 2700661 | - | - | *PA14_31060* | hypothetical protein | 111 | 101 | 148 | 97 | 119 | 121 | 1 |
| 2536 | - | 2703530 | 2701170 | - | - | *PA14_31070* | hypothetical protein | 3 | 3 | 4 | 4 | 3 | 3 | 1 |
| 2537 | - | 2704179 | 2703604 | - | - | *PA14_31080* | conjugal transfer protein | 8 | 9 | 10 | 9 | 7 | 9 | 1 |
| 2538 | - | 2705093 | 2704176 | - | - | *PA14_31090* | hypothetical protein | 5 | 6 | 6 | 5 | 5 | 6 | 1 |
| 2539 | - | 2705649 | 2705011 | - | - | *PA14_31100* | plasmid partitioning protein | 5 | 4 | 5 | 4 | 2 | 4 | 1 |
| 2540 | - | 2707231 | 2705903 | - | - | *PA14_31110* | replication initiator and transcriptional repressor protein | 11 | 8 | 11 | 9 | 8 | 10 | 1 |
| 2541 | - | 2707950 | 2707180 | - | - | *PA14_31130* | hypothetical protein | 6 | 5 | 6 | 5 | 5 | 6 | 1 |
| 2542 | - | 2708695 | 2708294 | - | - | *PA14_31150* | hypothetical protein | 204 | 154 | 169 | 155 | 151 | 266 | 1 |
| 2543 | - | 2708860 | 2709183 | + | - | *PA14_31160* | hypothetical protein | 74 | 116 | 116 | 95 | 109 | 16 | 1 |
| 2544 | - | 2709230 | 2709337 | + | - | *PA14_31170* | hypothetical protein | 129 | 161 | 155 | 140 | 149 | 28 | 1 |
| 2545 | - | 2710638 | 2709577 | - | - | *PA14_31180* | hypothetical protein | 28 | 27 | 39 | 26 | 23 | 13 | 1 |
| 2546 | - | 2712999 | 2710945 | - | - | *PA14_31190* | hypothetical protein | 3 | 2 | 3 | 3 | 3 | 2 | 1 |
| 2547 | - | 2713905 | 2713081 | - | - | *PA14_31200* | hypothetical protein | 2 | 3 | 2 | 4 | 3 | 3 | 1 |
| 2548 | - | 2714903 | 2714625 | - | - | *PA14_31220* | hypothetical protein | 5 | 7 | 8 | 11 | 7 | 8 | 1 |
| 2549 | - | 2715496 | 2714969 | - | - | *PA14_31230* | hypothetical protein | 5 | 2 | 6 | 4 | 3 | 3 | 0.405144617 |
| 2550 | - | 2716943 | 2715633 | - | - | *PA14_31240* | nrbE-like protein | 89 | 73 | 85 | 63 | 82 | 84 | 1 |
| 2551 | - | 2717738 | 2717346 | - | - | *PA14_31250* | hypothetical protein | 8 | 4 | 5 | 4 | 3 | 5 | 0.434308967 |
| 2552 | - | 2718521 | 2718015 | - | - | *PA14_31260* | RadC-like protein | 1 | 1 | 1 | 1 | 1 | 1 | 1 |
| 2553 | - | 2720011 | 2718848 | - | - | *PA14_31270* | hypothetical protein | 15 | 14 | 16 | 16 | 11 | 14 | 1 |
| 2554 | - | 2721213 | 2720008 | - | - | *PA14_31280* | integrase | 20 | 21 | 19 | 19 | 18 | 19 | 1 |
| 2555 | 2721761 | 2721861 | 2722229 | + | *pa1L* | *PA14_31290* | PA-I galactophilic lectin | 71 | 12 | 11 | 42 | 27 | 161 | 1.60308E-29 |
| 2556 | - | 2722503 | 2722844 | + | - | *PA14_31300* | hypothetical protein | 56 | 48 | 50 | 58 | 47 | 63 | 1 |
| 2557 | - | 2722963 | 2723370 | + | - | *PA14_31310* | hypothetical protein | 172 | 149 | 169 | 244 | 174 | 186 | 1 |
| 2558 | - | 2724928 | 2723429 | - | - | *PA14_31330* | EAL domain-containing protein | 22 | 19 | 24 | 13 | 17 | 27 | 1 |
| 2559 | - | 2725301 | 2725657 | + | - | *PA14_31340* | hypothetical protein | 4 | 2 | 3 | 4 | 4 | 4 | 1 |
| 2560 | - | 2726001 | 2727038 | + | - | *PA14_31350* | hypothetical protein | 240 | 71 | 167 | 327 | 118 | 340 | 0.062668448 |
| 2561 | - | 2727061 | 2727486 | + | - | *PA14_31360* | hypothetical protein | 102 | 23 | 39 | 98 | 45 | 133 | 3.49395E-10 |
| 2562 | - | 2727476 | 2728303 | + | - | *PA14_31370* | hypothetical protein | 194 | 44 | 73 | 192 | 96 | 289 | 1.5759E-05 |
| 2563 | - | 2729800 | 2728313 | - | - | *PA14_31380* | sulfate transporter | 36 | 12 | 18 | 35 | 22 | 61 | 0.005762476 |
| 2564 | - | 2730150 | 2730662 | + | - | *PA14_31390* | hypothetical protein | 43 | 48 | 47 | 59 | 40 | 56 | 1 |
| 2565 | - | 2732504 | 2730777 | - | - | *PA14_31400* | chemotaxis transducer | 20 | 22 | 26 | 28 | 19 | 19 | 1 |
| 2566 | - | 2732652 | 2732939 | + | - | *PA14_31420* | hypothetical protein | 52 | 37 | 48 | 29 | 29 | 42 | 1 |
| 2567 | - | 2733244 | 2733014 | - | - | *PA14_31430* | hypothetical protein | 55 | 56 | 78 | 57 | 65 | 51 | 1 |
| 2568 | - | 2733563 | 2734105 | + | - | *PA14_31440* | hypothetical protein | 295 | 267 | 320 | 339 | 266 | 304 | 1 |
| 2569 | - | 2734291 | 2734145 | - | - | *PA14_31450* | hypothetical protein | 91 | 90 | 109 | 115 | 87 | 107 | 1 |
| 2570 | - | 2735120 | 2734428 | - | - | *PA14_31460* | transporter | 26 | 22 | 30 | 27 | 22 | 24 | 1 |
| 2571 | - | 2735341 | 2737035 | + | - | *PA14_31470* | AMP-binding protein | 94 | 145 | 106 | 141 | 111 | 124 | 1 |
| 2572 | - | 2737104 | 2738189 | + | - | *PA14_31480* | AraC family transcriptional regulator | 72 | 80 | 74 | 96 | 76 | 86 | 1 |
| 2573 | - | 2738366 | 2740033 | + | - | *PA14_31500* | AMP-binding protein | 291 | 345 | 282 | 418 | 384 | 449 | 1 |
| 2574 | - | 2740053 | 2740820 | + | - | *PA14_31510* | short-chain dehydrogenase | 441 | 476 | 373 | 586 | 516 | 589 | 1 |
| 2575 | - | 2740838 | 2742028 | + | - | *PA14_31530* | acyl-CoA thiolase | 397 | 550 | 377 | 524 | 541 | 553 | 1 |
| 2576 | - | 2742057 | 2743184 | + | - | *PA14_31540* | acyl-CoA dehydrogenase | 285 | 431 | 272 | 367 | 373 | 378 | 1 |
| 2577 | - | 2744233 | 2743301 | - | - | *PA14_31560* | LysR family transcriptional regulator | 83 | 95 | 90 | 98 | 101 | 99 | 1 |
| 2578 | - | 2744407 | 2745636 | + | - | *PA14_31580* | acyl-CoA dehydrogenase | 40 | 44 | 43 | 45 | 54 | 57 | 1 |
| 2579 | - | 2746870 | 2745827 | - | - | *PA14_31610* | TerC family protein | 26 | 20 | 31 | 20 | 21 | 17 | 1 |
| 2580 | - | 2748414 | 2747020 | - | - | *PA14_31620* | hypothetical protein | 7 | 6 | 8 | 8 | 7 | 5 | 1 |
| 2581 | - | 2749379 | 2748474 | - | - | *PA14_31630* | LysR family transcriptional regulator | 11 | 9 | 14 | 14 | 9 | 9 | 1 |
| 2582 | - | 2749483 | 2749914 | + | - | *PA14_31640* | ring-cleaving dioxygenase | 13 | 15 | 12 | 12 | 10 | 11 | 1 |
| 2583 | - | 2750833 | 2750021 | - | *xthA* | *PA14_31650* | exonuclease III | 32 | 36 | 34 | 33 | 35 | 29 | 1 |
| 2584 | - | 2750978 | 2751691 | + | - | *PA14_31660* | hypothetical protein | 96 | 89 | 103 | 141 | 98 | 108 | 1 |
| 2585 | - | 2751825 | 2753564 | + | - | *PA14_31680* | hypothetical protein | 46 | 39 | 45 | 46 | 40 | 40 | 1 |
| 2586 | - | 2753561 | 2757226 | + | - | *PA14_31690* | hypothetical protein | 45 | 34 | 37 | 39 | 38 | 39 | 1 |
| 2587 | - | 2757361 | 2757996 | + | - | *PA14_31700* | CDP-alcohol phosphatidyltransferase | 82 | 51 | 74 | 67 | 61 | 60 | 1 |
| 2588 | - | 2758086 | 2759846 | + | - | *PA14_31720* | hypothetical protein | 27 | 22 | 20 | 17 | 17 | 18 | 1 |
| 2589 | - | 2759882 | 2761195 | + | - | *PA14_31730* | hypothetical protein | 31 | 20 | 22 | 20 | 24 | 23 | 1 |
| 2590 | - | 2761188 | 2761640 | + | - | *PA14_31740* | hypothetical protein | 36 | 20 | 28 | 17 | 25 | 25 | 0.546512063 |
| 2591 | - | 2761652 | 2762281 | + | - | *PA14_31750* | acyltransferase | 20 | 14 | 14 | 10 | 16 | 12 | 1 |
| 2592 | - | 2762283 | 2763218 | + | - | *PA14_31760* | phosphatidate cytidylyltransferase | 26 | 21 | 20 | 17 | 24 | 24 | 1 |
| 2593 | - | 2764402 | 2763407 | - | - | *PA14_31770* | oxidoreductase | 11 | 11 | 11 | 9 | 9 | 9 | 1 |
| 2594 | - | 2764521 | 2765432 | + | - | *PA14_31780* | LysR family transcriptional regulator | 34 | 26 | 42 | 29 | 22 | 29 | 1 |
| 2595 | - | 2765588 | 2766937 | + | - | *PA14_31800* | sodium:alanine symporter | 46 | 59 | 47 | 47 | 48 | 38 | 1 |
| 2596 | - | 2767072 | 2767569 | + | *tpx* | *PA14_31810* | thiol peroxidase | 157 | 168 | 81 | 127 | 171 | 180 | 1 |
| 2597 | - | 2768843 | 2767719 | - | - | *PA14_31820* | aminotransferase | 6 | 6 | 5 | 4 | 5 | 4 | 1 |
| 2598 | - | 2770365 | 2769040 | - | - | *PA14_31840* | hypothetical protein | 47 | 45 | 36 | 44 | 40 | 43 | 1 |
| 2599 | - | 2771798 | 2770365 | - | - | *PA14_31850* | hypothetical protein | 45 | 41 | 46 | 51 | 40 | 42 | 1 |
| 2600 | - | 2772055 | 2773335 | + | - | *PA14_31870* | RND efflux membrane fusion protein | 35 | 33 | 36 | 37 | 25 | 25 | 1 |
| 2601 | - | 2773332 | 2776463 | + | - | *PA14_31890* | RND efflux transporter | 32 | 34 | 33 | 37 | 28 | 28 | 1 |
| 2602 | - | 2776460 | 2779570 | + | - | *PA14_31900* | efflux transporter | 41 | 37 | 40 | 50 | 38 | 41 | 1 |
| 2603 | - | 2779567 | 2781063 | + | - | *PA14_31920* | outer membrane protein | 46 | 58 | 45 | 60 | 50 | 47 | 1 |
| 2604 | - | 2781107 | 2781430 | + | - | *PA14_31930* | hypothetical protein | 23 | 21 | 16 | 20 | 19 | 15 | 1 |

| | A | B | C | D | E | F | G | H | I | J | K | L | M | N |
|---|---|---|---|---|---|---|---|---|---|---|---|---|---|---|
| 2605 | - | 2782857 | 2781439 | - | - | PA14_31950 | two-component sensor | 19 | 19 | 19 | 19 | 17 | 16 | 1 |
| 2606 | - | 2783528 | 2782854 | - | - | PA14_31960 | two-component response regulator | 12 | 15 | 11 | 11 | 11 | 10 | 1 |
| 2607 | - | 2784041 | 2785327 | + | czcC | PA14_31970 | CzcC family cobalt/zinc/cadmium efflux transporter outer membrane protein | 1 | 1 | 1 | 1 | 0 | 1 | 1 |
| 2608 | - | 2785380 | 2786834 | + | czcB | PA14_31990 | cobalt/zinc/cadmium efflux RND transporter, membrane fusion protein, CzcB famil | 5 | 3 | 3 | 3 | 2 | 2 | 1 |
| 2609 | - | 2790219 | 2791175 | + | xylS | PA14_32060 | transcriptional regulator XylS | 18 | 15 | 16 | 16 | 13 | 16 | 1 |
| 2610 | - | 2791292 | 2792659 | + | xylX | PA14_32080 | toluate 1,2-dioxygenase subunit alpha | 8 | 9 | 9 | 9 | 7 | 8 | 1 |
| 2611 | - | 2792656 | 2793144 | + | xylY | PA14_32100 | toluate 1,2-dioxygenase subunit beta | 2 | 0 | 1 | 2 | 1 | 1 | 9.5179E-11 |
| 2612 | - | 2793178 | 2794191 | + | xylZ | PA14_32110 | toluate 1,2-dioxygenase electron transfer component | 6 | 4 | 7 | 5 | 5 | 5 | 0.381163237 |
| 2613 | - | 2794217 | 2794978 | + | benD | PA14_32130 | 1,6-dihydroxycyclohexa-2,4-diene-1-carboxylate dehydrogenase | 3 | 4 | 3 | 3 | 3 | 3 | 1 |
| 2614 | - | 2796006 | 2794984 | - | antC | PA14_32140 | anthranilate dioxygenase reductase | 2 | 2 | 2 | 2 | 0 | 2 | 1 |
| 2615 | - | 2796516 | 2796025 | - | antB | PA14_32150 | anthranilate dioxygenase small subunit | 9 | 7 | 8 | 9 | 6 | 6 | 1 |
| 2616 | - | 2797907 | 2796513 | - | antA | PA14_32160 | anthranilate dioxygenase large subunit | 6 | 5 | 4 | 5 | 4 | 3 | 1 |
| 2617 | - | 2798224 | 2799225 | + | - | PA14_32190 | transcriptional regulator | 24 | 18 | 19 | 18 | 16 | 23 | 1 |
| 2618 | - | 2800106 | 2799234 | - | catR | PA14_32200 | transcriptional regulator CatR | 10 | 9 | 13 | 11 | 9 | 11 | 1 |
| 2619 | - | 2800268 | 2801389 | + | catB | PA14_32220 | muconate cycloisomerase I | 4 | 3 | 3 | 3 | 3 | 2 | 0.831807123 |
| 2620 | - | 2801421 | 2801711 | + | catC | PA14_32230 | muconolactone delta-isomerase | 4 | 5 | 5 | 5 | 4 | 3 | 1 |
| 2621 | - | 2801756 | 2802688 | + | catA | PA14_32240 | catechol 1,2-dioxygenase | 3 | 5 | 4 | 4 | 6 | 4 | 1 |
| 2622 | - | 2803064 | 2802846 | - | - | PA14_32250 | hypothetical protein | 4 | 3 | 5 | 4 | 5 | 5 | 1 |
| 2623 | - | 2803427 | 2804773 | + | - | PA14_32270 | porin | 7 | 11 | 9 | 9 | 8 | 9 | 1 |
| 2624 | - | 2805025 | 2805642 | + | - | PA14_32280 | hypothetical protein | 172 | 194 | 210 | 288 | 175 | 244 | 1 |
| 2625 | - | 2806858 | 2808357 | + | - | PA14_32300 | kinase | 9 | 7 | 10 | 8 | 5 | 6 | 1 |
| 2626 | - | 2806899 | 2805646 | - | - | PA14_32290 | hypothetical protein | 71 | 71 | 85 | 100 | 66 | 87 | 1 |
| 2627 | - | 2808461 | 2808628 | + | - | PA14_32310 | hypothetical protein | 1720 | 2056 | 2466 | 1541 | 1824 | 2505 | 1 |
| 2628 | - | 2809898 | 2808672 | - | - | PA14_32330 | cyanate permease | 173 | 182 | 235 | 120 | 145 | 195 | 1 |
| 2629 | - | 2810350 | 2809895 | - | - | PA14_32340 | deaminase | 11 | 9 | 10 | 10 | 10 | 11 | 1 |
| 2630 | - | 2811000 | 2810368 | - | - | PA14_32350 | hypothetical protein | 14 | 11 | 16 | 14 | 10 | 13 | 1 |
| 2631 | - | 2811128 | 2812006 | + | - | PA14_32360 | lysR family transcriptional regulator | 23 | 24 | 25 | 21 | 24 | 23 | 1 |
| 2632 | - | 2812065 | 2812580 | + | - | PA14_32370 | DNA damage-inducible gene | 9 | 14 | 11 | 12 | 10 | 10 | 1 |
| 2633 | - | 2814157 | 2812739 | - | oprN | PA14_32380 | multidrug efflux outer membrane protein OprN precursor | 6 | 7 | 7 | 6 | 7 | 6 | 1 |
| 2634 | - | 2817342 | 2814154 | - | mexF | PA14_32390 | RND multidrug efflux transporter MexF | 9 | 8 | 8 | 8 | 7 | 9 | 1 |
| 2635 | - | 2818608 | 2817364 | - | mexE | PA14_32400 | RND multidrug efflux membrane fusion protein MexE | 4 | 4 | 5 | 4 | 5 | 4 | 1 |
| 2636 | - | 2819753 | 2818839 | - | mexT | PA14_32410 | transcriptional regulator MexT | 48 | 35 | 41 | 42 | 40 | 41 | 1 |
| 2637 | - | 2819974 | 2820993 | + | - | PA14_32420 | oxidoreductase | 149 | 117 | 140 | 123 | 104 | 120 | 1 |
| 2638 | - | 2821426 | 2821052 | - | - | PA14_32440 | hypothetical protein | 42 | 38 | 44 | 34 | 37 | 32 | 1 |
| 2639 | - | 2822322 | 2821507 | - | - | PA14_32450 | AraC family transcriptional regulator | 21 | 21 | 27 | 20 | 21 | 18 | 1 |
| 2640 | - | 2823116 | 2822349 | - | - | PA14_32460 | transcriptional regulator | 23 | 19 | 25 | 22 | 24 | 21 | 1 |
| 2641 | - | 2823214 | 2823486 | + | - | PA14_32470 | hypothetical protein | 11 | 15 | 14 | 14 | 14 | 13 | 1 |
| 2642 | - | 2823706 | 2823512 | - | - | PA14_32480 | hypothetical protein | 59 | 75 | 64 | 61 | 77 | 60 | 1 |
| 2643 | - | 2824001 | 2823723 | - | - | PA14_32490 | hypothetical protein | 55 | 42 | 65 | 52 | 70 | 55 | 1 |
| 2644 | - | 2824030 | 2824644 | + | - | PA14_32500 | TetR family transcriptional regulator | 94 | 67 | 91 | 67 | 63 | 66 | 1 |
| 2645 | - | 2824795 | 2825796 | + | - | PA14_32520 | hypothetical protein | 32 | 31 | 29 | 34 | 33 | 35 | 1 |
| 2646 | - | 2825911 | 2826564 | + | - | PA14_32530 | cytochrome c | 103 | 92 | 99 | 61 | 77 | 95 | 1 |
| 2647 | - | 2826561 | 2827442 | + | - | PA14_32540 | hypothetical protein | 118 | 109 | 114 | 80 | 101 | 119 | 1 |
| 2648 | - | 2828954 | 2827632 | - | - | PA14_32570 | two-component sensor | 59 | 47 | 58 | 45 | 47 | 50 | 1 |
| 2649 | - | 2829631 | 2828951 | - | - | PA14_32580 | two-component response regulator | 23 | 19 | 23 | 18 | 15 | 21 | 1 |
| 2650 | - | 2829794 | 2831563 | + | - | PA14_32590 | thiol:disulfide interchange protein | 21 | 23 | 26 | 29 | 19 | 22 | 1 |
| 2651 | - | 2831563 | 2832399 | + | - | PA14_32600 | thiol:disulfide interchange protein | 28 | 27 | 30 | 39 | 32 | 30 | 1 |
| 2652 | - | 2832396 | 2833166 | + | dsbG | PA14_32610 | disulfide isomerase/thiol-disulfide oxidase | 50 | 64 | 58 | 70 | 64 | 60 | 1 |
| 2653 | - | 2833239 | 2834573 | + | - | PA14_32630 | cytochrome P450 | 49 | 36 | 47 | 53 | 39 | 52 | 1 |
| 2654 | - | 2835466 | 2834555 | - | - | PA14_32640 | hypothetical protein | 11 | 11 | 14 | 15 | 11 | 14 | 1 |
| 2655 | - | 2836152 | 2835508 | - | - | PA14_32650 | glutathione S-transferase | 7 | 5 | 9 | 9 | 5 | 8 | 1 |
| 2656 | - | 2837511 | 2836165 | - | - | PA14_32660 | MFS family transporter | 5 | 5 | 6 | 6 | 5 | 4 | 1 |
| 2657 | - | 2838287 | 2837589 | - | - | PA14_32670 | hypothetical protein | 4 | 5 | 7 | 5 | 5 | 5 | 1 |
| 2658 | - | 2839361 | 2838300 | - | gtdA | PA14_32690 | gentisate 1,2-dioxygenase | 6 | 5 | 4 | 4 | 5 | 3 | 1 |
| 2659 | - | 2839486 | 2840400 | + | - | PA14_32700 | transcriptional regulator | 10 | 10 | 10 | 9 | 10 | 8 | 1 |
| 2660 | - | 2840489 | 2841007 | + | - | PA14_32710 | ECF subfamily RNA polymerase sigma-70 factor | 17 | 80 | 26 | 139 | 91 | 29 | 1.78038E-09 |
| 2661 | - | 2841016 | 2842002 | + | - | PA14_32720 | transmembrane sensor | 13 | 26 | 19 | 52 | 32 | 18 | 0.41668339 |
| 2662 | - | 2842146 | 2844608 | + | - | PA14_32740 | TonB-dependent receptor | 8 | 9 | 8 | 12 | 10 | 7 | 1 |
| 2663 | - | 2844726 | 2845874 | + | - | PA14_32750 | hypothetical protein | 3 | 2 | 3 | 3 | 2 | 3 | 1 |
| 2664 | - | 2845921 | 2846445 | + | - | PA14_32770 | hypothetical protein | 34 | 26 | 33 | 29 | 24 | 26 | 1 |
| 2665 | - | 2846758 | 2848455 | + | - | PA14_32780 | hypothetical protein | 34 | 30 | 39 | 35 | 31 | 28 | 1 |
| 2666 | - | 2848568 | 2864206 | + | - | PA14_32790 | hypothetical protein | 46 | 39 | 43 | 33 | 43 | 31 | 1 |
| 2667 | - | 2865179 | 2865490 | + | - | PA14_32810 | hypothetical protein | 10 | 11 | 9 | 10 | 11 | 9 | 1 |
| 2668 | - | 2868300 | 2868887 | + | - | PA14_32820 | hypothetical protein | 16 | 10 | 11 | 10 | 10 | 8 | 0.569926259 |
| 2669 | - | 2873250 | 2870581 | - | - | PA14_32830 | hypothetical protein | 37 | 41 | 34 | 47 | 43 | 37 | 1 |
| 2670 | - | 2874218 | 2873265 | - | - | PA14_32840 | hypothetical protein | 39 | 38 | 32 | 38 | 42 | 41 | 1 |
| 2671 | - | 2876003 | 2874987 | - | - | PA14_32850 | hypothetical protein | 40 | 38 | 43 | 57 | 45 | 41 | 1 |
| 2672 | - | 2877069 | 2876533 | - | - | PA14_32860 | hypothetical protein | 27 | 29 | 31 | 30 | 31 | 28 | 1 |
| 2673 | - | 2877917 | 2877066 | - | - | PA14_32880 | hypothetical protein | 17 | 15 | 16 | 14 | 15 | 13 | 1 |
| 2674 | - | 2878192 | 2877971 | - | - | PA14_32890 | hypothetical protein | 37 | 66 | 54 | 43 | 40 | 25 | 0.831807123 |
| 2675 | - | 2878349 | 2879929 | + | - | PA14_32905 | hypothetical protein | 9 | 8 | 10 | 11 | 8 | 10 | 1 |
| 2676 | - | 2880132 | 2881049 | + | - | PA14_32930 | hypothetical protein | 15 | 13 | 13 | 13 | 10 | 13 | 1 |
| 2677 | - | 2881131 | 2882666 | + | - | PA14_32940 | transcriptional regulator | 42 | 29 | 33 | 41 | 31 | 47 | 1 |
| 2678 | - | 2884416 | 2882674 | - | - | PA14_32950 | hypothetical protein | 50 | 19 | 25 | 36 | 27 | 61 | 0.137571381 |
| 2679 | - | 2884539 | 2885462 | + | - | PA14_32970 | transcriptional regulator | 7 | 8 | 7 | 10 | 7 | 6 | 1 |
| 2680 | - | 2885721 | 2886104 | + | gcvH2 | PA14_32985 | glycine cleavage system protein H | 106 | 173 | 154 | 182 | 169 | 150 | 0.885197681 |
| 2681 | - | 2886115 | 2888994 | + | gcvP2 | PA14_33000 | glycine dehydrogenase | 119 | 263 | 147 | 258 | 230 | 245 | 1 |
| 2682 | - | 2889137 | 2890393 | + | glyA2 | PA14_33010 | serine hydroxymethyltransferase | 93 | 192 | 132 | 243 | 193 | 223 | 1 |
| 2683 | - | 2890441 | 2891817 | + | sdaA | PA14_33030 | L-serine dehydratase | 124 | 200 | 183 | 244 | 220 | 244 | 1 |
| 2684 | - | 2891911 | 2893032 | + | gcvT2 | PA14_33040 | glycine cleavage system protein T2 | 181 | 308 | 256 | 398 | 364 | 382 | 1 |
| 2685 | - | 2893954 | 2893079 | - | - | PA14_33050 | hypothetical protein | 79 | 133 | 97 | 172 | 160 | 151 | 1 |
| 2686 | - | 2894922 | 2893951 | - | - | PA14_33060 | hypothetical protein | 16 | 15 | 15 | 16 | 17 | 18 | 1 |
| 2687 | - | 2895611 | 2897611 | + | - | PA14_33070 | hypothetical protein | 19 | 14 | 22 | 20 | 17 | 17 | 1 |
| 2688 | - | 2897623 | 2898636 | + | - | PA14_33080 | hypothetical protein | 22 | 19 | 21 | 23 | 21 | 21 | 1 |

| | A | B | C | D | E | F | G | H | I | J | K | L | M | N |
|---|---|---|---|---|---|---|---|---|---|---|---|---|---|---|
| 2689 | - | 2898633 | 2899673 | + | - | *PA14_33110* | hypothetical protein | 19 | 15 | 17 | 19 | 19 | 19 | 1 |
| 2690 | - | 2899827 | 2900261 | + | - | *PA14_33120* | hypothetical protein | 100 | 105 | 113 | 111 | 105 | 91 | 1 |
| 2691 | - | 2900290 | 2902275 | + | - | *PA14_33130* | cation-transporting P-type ATPase | 72 | 71 | 95 | 89 | 75 | 87 | 1 |
| 2692 | - | 2902272 | 2902817 | + | - | *PA14_33150* | hypothetical protein | 85 | 70 | 86 | 90 | 78 | 83 | 1 |
| 2693 | - | 2903221 | 2902952 | - | - | *PA14_33160* | hypothetical protein | 67 | 70 | 67 | 81 | 75 | 84 | 1 |
| 2694 | - | 2904250 | 2903336 | - | - | *PA14_33170* | transcriptional regulator | 11 | 13 | 8 | 12 | 12 | 11 | 1 |
| 2695 | - | 2904651 | 2906825 | + | - | *PA14_33190* | hypothetical protein | 13 | 12 | 17 | 19 | 15 | 16 | 1 |
| 2696 | - | 2906815 | 2907024 | + | - | *PA14_33200* | hypothetical protein | 6 | 10 | 10 | 12 | 10 | 12 | 1 |
| 2697 | - | 2907021 | 2908025 | + | - | *PA14_33220* | hypothetical protein | 15 | 21 | 19 | 26 | 19 | 21 | 1 |
| 2698 | - | 2908307 | 2908062 | - | - | *PA14_33230* | hypothetical protein | 12 | 18 | 13 | 16 | 14 | 22 | 1 |
| 2699 | - | 2908569 | 2909483 | + | - | *PA14_33240* | hypothetical protein | 4 | 4 | 4 | 4 | 4 | 3 | 1 |
| 2700 | - | 2909570 | 2910037 | + | - | *PA14_33250* | hypothetical protein | 2 | 3 | 2 | 14 | 3 | 3 | 1 |
| 2701 | - | 2910617 | 2910054 | - | *pvdS* | *PA14_33260* | extracytoplasmic-function sigma-70 factor | 3 | 10 | 4 | 32 | 9 | 5 | 0.006435902 |
| 2702 | - | 2911261 | 2912025 | + | *pvdG* | *PA14_33270* | protein PvdG | 1 | 0 | 1 | 2 | 0 | 0 | 1 |
| 2703 | - | 2912099 | 2925127 | + | *pvdL* | *PA14_33280* | peptide synthase | 4 | 4 | 5 | 5 | 4 | 4 | 1 |
| 2704 | - | 2926368 | 2925613 | - | - | *PA14_33290* | hypothetical protein | 59 | 62 | 71 | 83 | 59 | 74 | 1 |
| 2705 | - | 2928020 | 2927517 | - | - | *PA14_33300* | hypothetical protein | 184 | 240 | 157 | 129 | 183 | 146 | 1 |
| 2706 | - | 2929112 | 2928084 | - | - | *PA14_33310* | hypothetical protein | 209 | 271 | 180 | 138 | 194 | 166 | 1 |
| 2707 | - | 2930106 | 2929123 | - | - | *PA14_33320* | hypothetical protein | 189 | 221 | 183 | 164 | 185 | 176 | 1 |
| 2708 | - | 2931397 | 2930093 | - | - | *PA14_33330* | hypothetical protein | 148 | 171 | 146 | 115 | 134 | 141 | 1 |
| 2709 | - | 2934685 | 2931455 | - | - | *PA14_33340* | helicase | 74 | 68 | 65 | 75 | 65 | 66 | 1 |
| 2710 | - | 2935656 | 2934682 | - | - | *PA14_33350* | hypothetical protein | 95 | 60 | 77 | 82 | 79 | 62 | 1 |
| 2711 | - | 2935897 | 2937291 | + | - | *PA14_33360* | hypothetical protein | 30 | 17 | 27 | 36 | 17 | 19 | 1 |
| 2712 | - | 2937790 | 2937476 | - | - | *PA14_33370* | hypothetical protein | 7 | 8 | 5 | 7 | 7 | 10 | 1 |
| 2713 | - | 2939415 | 2938495 | - | - | *PA14_33380* | hypothetical protein | 3 | 3 | 4 | 4 | 2 | 3 | 1 |
| 2714 | - | 2940867 | 2939449 | - | - | *PA14_33410* | porin | 4 | 5 | 6 | 5 | 3 | 4 | 1 |
| 2715 | - | 2941629 | 2940949 | - | - | *PA14_33420* | hydrolase | 10 | 14 | 10 | 11 | 10 | 10 | 1 |
| 2716 | - | 2942587 | 2941727 | - | - | *PA14_33430* | pirin-related protein | 2 | 3 | 3 | 3 | 2 | 2 | 1 |
| 2717 | - | 2942688 | 2943626 | + | - | *PA14_33440* | LysR family transcriptional regulator | 46 | 45 | 52 | 48 | 46 | 51 | 1 |
| 2718 | - | 2945267 | 2943630 | - | *treA* | *PA14_33450* | trehalase | 18 | 22 | 21 | 32 | 22 | 28 | 1 |
| 2719 | - | 2945626 | 2946051 | + | - | *PA14_33460* | hypothetical protein | 23 | 25 | 27 | 39 | 27 | 32 | 1 |
| 2720 | - | 2946044 | 2947363 | + | *sndH* | *PA14_33480* | L-sorbosone dehydrogenase | 48 | 61 | 72 | 111 | 63 | 87 | 1 |
| 2721 | - | 2947542 | 2948951 | + | *pvdH* | *PA14_33500* | diaminobutyrate--2-oxoglutarate aminotransferase | 8 | 12 | 12 | 22 | 13 | 16 | 1 |
| 2722 | - | 2949029 | 2949247 | + | - | *PA14_33510* | hypothetical protein | 3 | 6 | 5 | 12 | 4 | 6 | 0.711763148 |
| 2723 | - | 2949248 | 2950012 | + | - | *PA14_33520* | thioesterase | 10 | 9 | 12 | 22 | 11 | 13 | 1 |
| 2724 | - | 2951029 | 2950112 | - | - | *PA14_33530* | hypothetical protein | 6 | 9 | 6 | 7 | 7 | 6 | 1 |
| 2725 | - | 2951931 | 2951026 | - | - | *PA14_33540* | ABC transporter permease | 3 | 4 | 3 | 6 | 6 | 3 | 1 |
| 2726 | - | 2952683 | 2951928 | - | - | *PA14_33550* | ABC transporter ATP-binding protein | 7 | 9 | 8 | 9 | 8 | 5 | 1 |
| 2727 | - | 2953633 | 2952680 | - | - | *PA14_33560* | adhesion protein | 6 | 7 | 6 | 7 | 5 | 4 | 1 |
| 2728 | - | 2954226 | 2953666 | - | - | *PA14_33570* | hypothetical protein | 7 | 8 | 10 | 13 | 10 | 9 | 1 |
| 2729 | - | 2954552 | 2954223 | - | - | *PA14_33580* | hypothetical protein | 13 | 15 | 14 | 22 | 18 | 20 | 1 |
| 2730 | - | 2955088 | 2954549 | - | - | *PA14_33590* | hypothetical protein | 8 | 13 | 9 | 19 | 13 | 11 | 1 |
| 2731 | - | 2956296 | 2955085 | - | - | *PA14_33600* | hypothetical protein | 13 | 13 | 15 | 15 | 14 | 13 | 1 |
| 2732 | - | 2956615 | 2972064 | + | - | *PA14_33610* | peptide synthase | 13 | 11 | 14 | 15 | 11 | 13 | 1 |
| 2733 | - | 2972068 | 2978637 | + | *pvdJ* | *PA14_33630* | protein PvdJ | 24 | 20 | 25 | 24 | 20 | 23 | 1 |
| 2734 | - | 2978649 | 2985995 | + | *pvdD* | *PA14_33650* | pyoverdine synthetase D | 27 | 25 | 30 | 32 | 24 | 28 | 1 |
| 2735 | - | 2988606 | 2986159 | - | *fpvA* | *PA14_33680* | ferripyoverdine receptor | 19 | 25 | 20 | 39 | 30 | 22 | 1 |
| 2736 | - | 2990358 | 2988709 | - | *pvdE* | *PA14_33690* | pyoverdine biosynthesis protein PvdE | 34 | 37 | 39 | 44 | 33 | 41 | 1 |
| 2737 | - | 2990736 | 2991563 | + | *pvdF* | *PA14_33700* | pyoverdine synthetase F | 18 | 18 | 17 | 20 | 14 | 16 | 1 |
| 2738 | - | 2992486 | 2991632 | - | *pvdO* | *PA14_33710* | protein PvdO | 4 | 5 | 3 | 4 | 4 | 3 | 1 |
| 2739 | - | 2993798 | 2992515 | - | *pvdN* | *PA14_33720* | protein PvdN | 3 | 4 | 4 | 5 | 3 | 4 | 1 |
| 2740 | - | 2995167 | 2993821 | - | - | *PA14_33730* | dipeptidase | 3 | 5 | 5 | 5 | 3 | 4 | 1 |
| 2741 | - | 2995383 | 2997017 | + | *pvdP* | *PA14_33740* | protein PvdP | 4 | 5 | 5 | 4 | 4 | 4 | 1 |
| 2742 | - | 2998490 | 2997066 | - | - | *PA14_33750* | outer membrane protein | 9 | 7 | 9 | 7 | 7 | 7 | 1 |
| 2743 | - | 3000487 | 2998496 | - | - | *PA14_33760* | ABC transporter ATP-binding protein/permease | 11 | 11 | 12 | 15 | 14 | 11 | 1 |
| 2744 | - | 3001662 | 3000487 | - | - | *PA14_33770* | hypothetical protein | 8 | 6 | 9 | 10 | 10 | 9 | 1 |
| 2745 | - | 3002758 | 3001763 | - | - | *PA14_33780* | transmembrane sensor | 77 | 108 | 82 | 153 | 154 | 116 | 1 |
| 2746 | - | 3002922 | 3003401 | + | - | *PA14_33800* | RNA polymerase sigma factor | 30 | 38 | 43 | 58 | 64 | 44 | 1 |
| 2747 | - | 3003534 | 3004865 | + | *pvdA* | *PA14_33810* | L-ornithine N5-oxygenase | 7 | 8 | 9 | 16 | 8 | 9 | 1 |
| 2748 | - | 3004988 | 3007276 | + | *pvdQ* | *PA14_33820* | penicillin acylase-related protein | 4 | 4 | 4 | 6 | 4 | 4 | 1 |
| 2749 | - | 3007457 | 3007780 | + | - | *PA14_33830* | hypothetical protein | 1 | 6 | 4 | 11 | 5 | 4 | 7.29817E-07 |
| 2750 | - | 3007822 | 3008742 | + | - | *PA14_33840* | transcriptional regulator | 16 | 16 | 21 | 18 | 16 | 15 | 1 |
| 2751 | - | 3008839 | 3009990 | + | *lldA* | *PA14_33860* | L-lactate dehydrogenase | 31 | 32 | 29 | 36 | 28 | 27 | 1 |
| 2752 | - | 3010299 | 3010057 | - | - | *PA14_33870* | hypothetical protein | 260 | 352 | 190 | 312 | 269 | 330 | 1 |
| 2753 | - | 3010594 | 3010830 | + | - | *PA14_33880* | hypothetical protein | 28 | 34 | 34 | 41 | 29 | 33 | 1 |
| 2754 | - | 3011096 | 3011566 | + | - | *PA14_33890* | oxidoreductase | 45 | 46 | 46 | 51 | 40 | 42 | 1 |
| 2755 | - | 3011563 | 3013878 | + | - | *PA14_33900* | aldehyde dehydrogenase | 71 | 77 | 72 | 85 | 72 | 81 | 1 |
| 2756 | - | 3015569 | 3014295 | - | - | *PA14_33910* | hypothetical protein | 8 | 10 | 9 | 9 | 9 | 9 | 1 |
| 2757 | - | 3016342 | 3015701 | - | - | *PA14_33920* | transcriptional regulator | 5 | 4 | 5 | 5 | 5 | 5 | 1 |
| 2758 | - | 3016619 | 3017014 | + | - | *PA14_33930* | hypothetical protein | 46 | 51 | 66 | 74 | 52 | 58 | 1 |
| 2759 | - | 3017573 | 3017037 | - | - | *PA14_33940* | hypothetical protein | 33 | 34 | 38 | 40 | 32 | 43 | 1 |
| 2760 | - | 3019590 | 3017584 | - | - | *PA14_33960* | hypothetical protein | 16 | 15 | 20 | 25 | 17 | 25 | 1 |
| 2761 | - | 3020694 | 3019627 | - | - | *PA14_33970* | hypothetical protein | 39 | 35 | 49 | 46 | 32 | 52 | 1 |
| 2762 | - | 3021266 | 3020733 | - | - | *PA14_33980* | hypothetical protein | 60 | 48 | 69 | 71 | 51 | 87 | 1 |
| 2763 | - | 3023889 | 3021340 | - | - | *PA14_33990* | ClpA/B-type protease | 38 | 27 | 47 | 51 | 31 | 52 | 1 |
| 2764 | - | 3024907 | 3023891 | - | - | *PA14_34000* | hypothetical protein | 25 | 19 | 30 | 32 | 20 | 32 | 1 |
| 2765 | - | 3026664 | 3024871 | - | - | *PA14_34010* | hypothetical protein | 22 | 16 | 24 | 29 | 17 | 30 | 1 |
| 2766 | - | 3027073 | 3026648 | - | - | *PA14_34020* | hypothetical protein | 30 | 24 | 31 | 36 | 23 | 39 | 1 |
| 2767 | - | 3027583 | 3027086 | - | - | *PA14_34030* | hypothetical protein | 53 | 40 | 45 | 57 | 39 | 55 | 1 |
| 2768 | - | 3029141 | 3027657 | - | - | *PA14_34050* | hypothetical protein | 42 | 35 | 42 | 47 | 28 | 41 | 1 |
| 2769 | - | 3029709 | 3029164 | - | - | *PA14_34070* | hypothetical protein | 25 | 26 | 38 | 32 | 18 | 23 | 1 |
| 2770 | - | 3029917 | 3030393 | + | - | *PA14_34080* | hypothetical protein | 106 | 122 | 174 | 145 | 104 | 113 | 1 |
| 2771 | - | 3030453 | 3031784 | + | - | *PA14_34100* | hypothetical protein | 142 | 130 | 181 | 199 | 129 | 164 | 1 |
| 2772 | - | 3031802 | 3032560 | + | - | *PA14_34110* | hypothetical protein | 53 | 39 | 52 | 49 | 40 | 50 | 1 |

| | A | B | C | D | E | F | G | H | I | J | K | L | M | N |
|---|---|---|---|---|---|---|---|---|---|---|---|---|---|---|
| 2773 | - | 3032557 | 3036372 | + | - | *PA14_34130* | hypothetical protein | 44 | 34 | 54 | 56 | 41 | 55 | 1 |
| 2774 | - | 3036369 | 3037469 | + | - | *PA14_34140* | hypothetical protein | 55 | 54 | 61 | 68 | 54 | 79 | 1 |
| 2775 | - | 3037579 | 3038664 | + | - | *PA14_34150* | transcriptional regulator | 12 | 23 | 12 | 11 | 14 | 11 | 0.415011252 |
| 2776 | - | 3038777 | 3039166 | + | - | *PA14_34170* | hypothetical protein | 3 | 4 | 4 | 4 | 4 | 3 | 1 |
| 2777 | - | 3039326 | 3039886 | + | *msuE* | *PA14_34180* | NADH-dependent FMN reductase MsuE | 1 | 1 | 1 | 1 | 1 | 1 | 1 |
| 2778 | - | 3039896 | 3041041 | + | *msuD* | *PA14_34190* | methanesulfonate sulfonatase MsuD | 2 | 2 | 2 | 3 | 2 | 2 | 1 |
| 2779 | - | 3041071 | 3042255 | + | - | *PA14_34200* | FMNH2-dependent monooxygenase | 5 | 5 | 8 | 7 | 5 | 6 | 1 |
| 2780 | - | 3042252 | 3043382 | + | - | *PA14_34210* | transcriptional regulator | 3 | 3 | 3 | 3 | 2 | 3 | 1 |
| 2781 | - | 3043554 | 3044708 | + | - | *PA14_34230* | hypothetical protein | 16 | 18 | 16 | 17 | 15 | 14 | 1 |
| 2782 | - | 3045901 | 3044774 | - | - | *PA14_34250* | glycerophosphoryl diester phosphodiesterase | 7 | 9 | 7 | 8 | 8 | 6 | 1 |
| 2783 | - | 3046732 | 3046079 | - | - | *PA14_34260* | ABC transporter permease | 6 | 5 | 7 | 6 | 5 | 6 | 1 |
| 2784 | - | 3047825 | 3046716 | - | - | *PA14_34270* | ABC transporter ATP-binding protein | 3 | 3 | 3 | 4 | 3 | 3 | 1 |
| 2785 | - | 3048616 | 3047822 | - | - | *PA14_34280* | hypothetical protein | 3 | 3 | 5 | 4 | 3 | 3 | 1 |
| 2786 | - | 3050036 | 3048648 | - | - | *PA14_34290* | DszA family monooxygenase | 3 | 2 | 3 | 2 | 2 | 3 | 1 |
| 2787 | - | 3051271 | 3050054 | - | - | *PA14_34300* | DszC family monooxygenase | 7 | 7 | 9 | 9 | 9 | 7 | 1 |
| 2788 | - | 3052517 | 3051282 | - | - | *PA14_34320* | DszC family monooxygenase | 6 | 6 | 7 | 6 | 4 | 5 | 1 |
| 2789 | - | 3052936 | 3054171 | + | - | *PA14_34330* | hypothetical protein | 78 | 97 | 96 | 117 | 95 | 105 | 1 |
| 2790 | - | 3055134 | 3054202 | - | *mtlZ* | *PA14_34340* | fructokinase | 48 | 50 | 61 | 72 | 64 | 65 | 1 |
| 2791 | - | 3056684 | 3055176 | - | *mtlY* | *PA14_34350* | xylulose kinase | 4 | 4 | 5 | 5 | 3 | 4 | 1 |
| 2792 | - | 3058156 | 3056681 | - | *mtlD* | *PA14_34360* | mannitol dehydrogenase | 5 | 5 | 5 | 5 | 4 | 5 | 1 |
| 2793 | - | 3059291 | 3058179 | - | - | *PA14_34370* | ABC maltose/mannitol transporter ATP-binding protein | 11 | 8 | 12 | 11 | 7 | 9 | 1 |
| 2794 | - | 3060164 | 3059331 | - | - | *PA14_34390* | binding-protein-dependent maltose/mannitol transport protein | 6 | 6 | 7 | 6 | 6 | 6 | 1 |
| 2795 | - | 3061107 | 3060175 | - | - | *PA14_34410* | binding-protein-dependent maltose/mannitol transport protein | 11 | 12 | 13 | 11 | 9 | 12 | 1 |
| 2796 | - | 3062493 | 3061183 | - | - | *PA14_34420* | maltose/mannitol ABC transporter substrate-binding protein | 8 | 8 | 9 | 11 | 8 | 6 | 1 |
| 2797 | - | 3063556 | 3062651 | - | *mtlR* | *PA14_34440* | transcriptional regulator MtlR | 23 | 20 | 29 | 28 | 21 | 23 | 1 |
| 2798 | - | 3064684 | 3063779 | - | - | *PA14_34450* | AraC family transcriptional regulator | 24 | 27 | 21 | 21 | 25 | 18 | 1 |
| 2799 | - | 3064789 | 3065349 | + | - | *PA14_34460* | hypothetical protein | 99 | 37 | 41 | 71 | 56 | 67 | 0.027358573 |
| 2800 | - | 3065399 | 3066466 | + | - | *PA14_34490* | hypothetical protein | 186 | 51 | 86 | 117 | 116 | 118 | 0.012682198 |
| 2801 | - | 3066463 | 3067299 | + | - | *PA14_34500* | ABC transporter ATP-binding protein | 101 | 28 | 48 | 69 | 66 | 68 | 0.000612412 |
| 2802 | - | 3067265 | 3068464 | + | - | *PA14_34510* | hypothetical protein | 61 | 19 | 31 | 47 | 39 | 45 | 0.006979836 |
| 2803 | - | 3068461 | 3069225 | + | - | *PA14_34520* | ABC transporter permease | 57 | 27 | 39 | 60 | 52 | 51 | 0.317658547 |
| 2804 | - | 3069451 | 3070860 | + | - | *PA14_34540* | xenobiotic compound DszA family monooxygenase | 22 | 8 | 13 | 18 | 17 | 20 | 0.015343712 |
| 2805 | - | 3072155 | 3070920 | - | - | *PA14_34550* | hypothetical protein | 14 | 8 | 13 | 13 | 12 | 11 | 0.679564673 |
| 2806 | - | 3073445 | 3072186 | - | - | *PA14_34580* | hypothetical protein | 3 | 4 | 5 | 5 | 4 | 4 | 1 |
| 2807 | - | 3075288 | 3073663 | - | - | *PA14_34600* | glyceraldehyde-3-phosphate dehydrogenase | 13 | 77 | 78 | 36 | 93 | 57 | 4.36376E-14 |
| 2808 | - | 3075596 | 3075453 | - | - | *PA14_34610* | hypothetical protein | 3 | 10 | 6 | 7 | 8 | 6 | 0.000532638 |
| 2809 | - | 3076974 | 3075622 | - | *gnuT* | *PA14_34630* | gluconate permease | 16 | 19 | 20 | 21 | 16 | 18 | 1 |
| 2810 | - | 3077592 | 3077071 | - | - | *PA14_34640* | gluconokinase | 12 | 19 | 16 | 17 | 19 | 13 | 1 |
| 2811 | - | 3077787 | 3078818 | + | *gntR* | *PA14_34660* | transcriptional regulator GntR | 47 | 47 | 49 | 47 | 43 | 42 | 1 |
| 2812 | - | 3079537 | 3079172 | - | - | *PA14_34670* | cupin superfamily protein | 52 | 57 | 53 | 67 | 52 | 51 | 1 |
| 2813 | - | 3080868 | 3079570 | - | - | *PA14_34680* | oxidoreductase | 22 | 26 | 27 | 31 | 29 | 23 | 1 |
| 2814 | - | 3081865 | 3080972 | - | - | *PA14_34690* | LysR family transcriptional regulator | 19 | 11 | 14 | 11 | 11 | 9 | 0.8413024 |
| 2815 | - | 3081983 | 3083158 | + | - | *PA14_34700* | beta lactamase | 2 | 2 | 2 | 2 | 2 | 2 | 1 |
| 2816 | - | 3083158 | 3084411 | + | - | *PA14_34710* | major facilitator transporter | 8 | 7 | 9 | 8 | 7 | 6 | 1 |
| 2817 | - | 3085272 | 3084472 | - | - | *PA14_34720* | hypothetical protein | 28 | 28 | 37 | 31 | 22 | 22 | 1 |
| 2818 | - | 3085870 | 3085289 | - | - | *PA14_34730* | XRE family transcriptional regulator | 8 | 20 | 9 | 7 | 10 | 8 | 0.015737353 |
| 2819 | - | 3086083 | 3085925 | - | - | *PA14_34740* | hypothetical protein | 10 | 28 | 8 | 11 | 16 | 9 | 0.013429913 |
| 2820 | - | 3086373 | 3087260 | + | - | *PA14_34750* | taurine catabolism dioxygenase | 6 | 5 | 5 | 6 | 3 | 5 | 1 |
| 2821 | - | 3087301 | 3088323 | + | - | *PA14_34770* | ABC transporter substrate-binding protein | 6 | 7 | 5 | 6 | 6 | 5 | 1 |
| 2822 | - | 3088331 | 3089179 | + | - | *PA14_34780* | ABC transporter ATP-binding protein | 2 | 4 | 4 | 4 | 2 | 1 | 0.723542393 |
| 2823 | - | 3089196 | 3090062 | + | - | *PA14_34790* | ABC transporter permease | 7 | 8 | 8 | 10 | 6 | 9 | 1 |
| 2824 | - | 3090172 | 3090789 | + | - | *PA14_34800* | amino acid transporter LysE | 57 | 43 | 58 | 42 | 44 | 47 | 1 |
| 2825 | - | 3090885 | 3094634 | + | - | *PA14_34810* | non-ribosomal peptide synthetase | 214 | 176 | 192 | 213 | 165 | 188 | 1 |
| 2826 | - | 3094663 | 3095751 | + | - | *PA14_34820* | regulatory protein | 167 | 145 | 131 | 143 | 125 | 168 | 1 |
| 2827 | - | 3095748 | 3096767 | + | - | *PA14_34830* | regulatory protein | 178 | 136 | 144 | 151 | 145 | 201 | 1 |
| 2828 | - | 3096787 | 3103164 | + | - | *PA14_34840* | non-ribosomal peptide synthetase | 335 | 230 | 300 | 278 | 272 | 347 | 1 |
| 2829 | - | 3103232 | 3103774 | + | - | *PA14_34850* | tRNA synthase | 126 | 99 | 125 | 113 | 127 | 140 | 1 |
| 2830 | - | 3103993 | 3105444 | + | *chiC* | *PA14_34870* | chitinase | 320 | 16 | 26 | 83 | 62 | 326 | 7.9991E-209 |
| 2831 | - | 3105621 | 3106370 | + | - | *PA14_34880* | GntR family transcriptional regulator | 36 | 14 | 17 | 19 | 19 | 31 | 0.01418108 |
| 2832 | - | 3106367 | 3108091 | + | - | *PA14_34900* | oxidoreductase | 31 | 17 | 15 | 19 | 21 | 35 | 0.950259107 |
| 2833 | - | 3108177 | 3108422 | + | - | *PA14_34920* | ferredoxin | 18 | 7 | 6 | 15 | 10 | 20 | 0.033575691 |
| 2834 | - | 3108760 | 3109725 | + | - | *PA14_34930* | phycobiliprotein | 5 | 2 | 3 | 2 | 4 | 5 | 0.065349589 |
| 2835 | - | 3109722 | 3110003 | + | - | *PA14_34940* | hypothetical protein | 6 | 5 | 6 | 6 | 6 | 6 | 1 |
| 2836 | - | 3110344 | 3111702 | + | - | *PA14_34960* | glucose-sensitive porin | 70 | 544 | 315 | 440 | 452 | 407 | 7.79018E-18 |
| 2837 | - | 3111856 | 3114267 | + | *gcd* | *PA14_34970* | glucose dehydrogenase | 62 | 78 | 63 | 80 | 83 | 103 | 1 |
| 2838 | - | 3114399 | 3116519 | + | - | *PA14_34990* | TonB-dependent receptor | 27 | 35 | 36 | 47 | 43 | 56 | 1 |
| 2839 | - | 3116640 | 3116927 | + | - | *PA14_35000* | hypothetical protein | 67 | 77 | 65 | 92 | 80 | 101 | 1 |
| 2840 | - | 3117575 | 3116940 | - | - | *PA14_35010* | hypothetical protein | 31 | 34 | 32 | 37 | 40 | 39 | 1 |
| 2841 | - | 3119106 | 3117562 | - | - | *PA14_35020* | hypothetical protein | 9 | 9 | 10 | 11 | 10 | 10 | 1 |
| 2842 | - | 3119769 | 3119107 | - | - | *PA14_35030* | hypothetical protein | 5 | 6 | 6 | 7 | 3 | 7 | 1 |
| 2843 | - | 3120437 | 3119766 | - | - | *PA14_35040* | hypothetical protein | 16 | 11 | 15 | 15 | 10 | 13 | 0.934705131 |
| 2844 | - | 3121891 | 3120434 | - | - | *PA14_35050* | hypothetical protein | 2 | 2 | 2 | 3 | 2 | 2 | 1 |
| 2845 | - | 3122326 | 3121898 | - | - | *PA14_35060* | hypothetical protein | 18 | 19 | 20 | 16 | 14 | 16 | 1 |
| 2846 | - | 3122614 | 3123468 | + | - | *PA14_35070* | AraC family transcriptional regulator | 26 | 27 | 32 | 28 | 25 | 24 | 1 |
| 2847 | - | 3124173 | 3123481 | - | - | *PA14_35080* | arsenical resistance protein | 16 | 19 | 17 | 15 | 17 | 15 | 1 |
| 2848 | - | 3124655 | 3124185 | - | *arsC* | *PA14_35100* | arsenate reductase | 7 | 13 | 8 | 9 | 9 | 10 | 0.679564673 |
| 2849 | - | 3125970 | 3124687 | - | *arsB* | *PA14_35110* | arsenite-antimonite efflux pump ArsB | 8 | 8 | 9 | 9 | 8 | 7 | 1 |
| 2850 | - | 3126334 | 3125984 | - | *arsR* | *PA14_35130* | arsenic resistance transcriptional regulator | 8 | 7 | 8 | 8 | 9 | 9 | 1 |
| 2851 | - | 3127303 | 3126413 | - | - | *PA14_35140* | AraC family transcriptional regulator | 8 | 7 | 9 | 9 | 6 | 7 | 1 |
| 2852 | - | 3127512 | 3128573 | + | - | *PA14_35150* | alcohol dehydrogenase | 35 | 11 | 12 | 13 | 10 | 47 | 0.000776048 |
| 2853 | - | 3128975 | 3128598 | - | - | *PA14_35160* | hypothetical protein | 114 | 10 | 7 | 8 | 5 | 116 | 1.2809E-136 |
| 2854 | - | 3129053 | 3129523 | + | - | *PA14_35170* | redox-sensing activator of soxS | 90 | 70 | 61 | 74 | 62 | 135 | 1 |
| 2855 | - | 3131228 | 3129531 | - | *pbpC* | *PA14_35190* | penicillin-binding protein 3A | 50 | 45 | 43 | 55 | 39 | 64 | 1 |
| 2856 | - | 3131865 | 3131350 | - | - | *PA14_35200* | acetyltransferase | 30 | 41 | 33 | 32 | 29 | 31 | 1 |

| | A | B | C | D | E | F | G | H | I | J | K | L | M | N |
|---|---|---|---|---|---|---|---|---|---|---|---|---|---|---|
| 2857 | - | 3132463 | 3131873 | - | - | PA14_35210 | TetR family transcriptional regulator | 24 | 25 | 23 | 21 | 19 | 20 | 1 |
| 2858 | - | 3132610 | 3133815 | + | - | PA14_35230 | hypothetical protein | 17 | 18 | 19 | 18 | 15 | 14 | 1 |
| 2859 | - | 3134849 | 3133779 | - | - | PA14_35240 | hypothetical protein | 15 | 11 | 17 | 13 | 12 | 15 | 1 |
| 2860 | - | 3134936 | 3135832 | + | - | PA14_35250 | LysR family transcriptional regulator | 38 | 45 | 40 | 44 | 37 | 41 | 1 |
| 2861 | - | 3137251 | 3135932 | - | - | PA14_35270 | cytochrome c precursor | 48 | 62 | 57 | 63 | 64 | 73 | 1 |
| 2862 | - | 3139038 | 3137263 | - | gnd | PA14_35290 | gluconate dehydrogenase | 30 | 49 | 44 | 45 | 48 | 50 | 0.940967938 |
| 2863 | - | 3139757 | 3139041 | - | - | PA14_35300 | hypothetical protein | 33 | 47 | 48 | 42 | 51 | 51 | 1 |
| 2864 | - | 3140911 | 3139925 | - | - | PA14_35320 | 2-hydroxyacid dehydrogenase | 4 | 8 | 5 | 7 | 8 | 7 | 0.381163237 |
| 2865 | - | 3142237 | 3140930 | - | - | PA14_35330 | 2-ketogluconate transporter | 4 | 6 | 7 | 7 | 7 | 5 | 1 |
| 2866 | - | 3143250 | 3142300 | - | - | PA14_35340 | 2-ketogluconate kinase | 16 | 23 | 31 | 31 | 25 | 21 | 1 |
| 2867 | - | 3144025 | 3143243 | - | - | PA14_35360 | hypothetical protein | 8 | 20 | 17 | 17 | 19 | 10 | 0.049721315 |
| 2868 | - | 3145119 | 3144097 | - | ptxS | PA14_35370 | transcriptional regulator PtxS | 14 | 31 | 25 | 27 | 29 | 19 | 0.173484736 |
| 2869 | - | 3145681 | 3146619 | + | ptxR | PA14_35380 | transcriptional regulator PtxR | 10 | 16 | 15 | 10 | 9 | 10 | 1 |
| 2870 | - | 3147369 | 3146722 | - | pvcD | PA14_35390 | pyoverdine biosynthesis protein PvcD | 5 | 6 | 7 | 6 | 4 | 4 | 1 |
| 2871 | - | 3148864 | 3147362 | - | pvcC | PA14_35400 | pyoverdine biosynthesis protein PvcC | 4 | 4 | 5 | 4 | 4 | 3 | 1 |
| 2872 | - | 3149791 | 3148916 | - | pvcB | PA14_35420 | pyoverdine biosynthesis protein PvcB | 6 | 5 | 4 | 6 | 6 | 5 | 1 |
| 2873 | - | 3150795 | 3149809 | - | pvcA | PA14_35430 | pyoverdine biosynthesis protein PvcA | 3 | 3 | 3 | 3 | 3 | 4 | 1 |
| 2874 | - | 3151997 | 3151011 | - | ansA | PA14_35440 | L-asparaginase I | 17 | 14 | 13 | 12 | 8 | 10 | 1 |
| 2875 | - | 3153542 | 3152097 | - | - | PA14_35460 | AGCS sodium/alanine/glycine symporter | 6 | 5 | 6 | 6 | 5 | 6 | 1 |
| 2876 | - | 3153711 | 3154529 | + | - | PA14_35470 | hypothetical protein | 19 | 23 | 19 | 19 | 19 | 18 | 1 |
| 2877 | - | 3156056 | 3154662 | - | lpdV | PA14_35490 | dihydrolipoamide dehydrogenase | 645 | 966 | 801 | 979 | 869 | 1119 | 1 |
| 2878 | - | 3157346 | 3156060 | - | bkdB | PA14_35500 | branched-chain alpha-keto acid dehydrogenase subunit E2 | 499 | 778 | 581 | 671 | 597 | 761 | 1 |
| 2879 | - | 3158399 | 3157347 | - | bkdA2 | PA14_35520 | 2-oxoisovalerate dehydrogenase subunit beta | 451 | 693 | 474 | 542 | 513 | 620 | 1 |
| 2880 | - | 3159628 | 3158396 | - | bkdA1 | PA14_35530 | 2-oxoisovalerate dehydrogenase subunit alpha | 323 | 534 | 313 | 320 | 363 | 394 | 1 |
| 2881 | - | 3159938 | 3160399 | + | bkdR | PA14_35540 | transcriptional regulator BkdR | 71 | 90 | 100 | 96 | 83 | 92 | 1 |
| 2882 | - | 3160737 | 3160432 | - | pslO | PA14_35550 | hypothetical protein | 45 | 56 | 45 | 48 | 45 | 44 | 1 |
| 2883 | - | 3161766 | 3160765 | - | pslN | PA14_35570 | hypothetical protein | 46 | 44 | 54 | 50 | 50 | 49 | 1 |
| 2884 | - | 3163555 | 3161822 | - | pslM | PA14_35590 | FAD-binding dehydrogenase | 4 | 4 | 4 | 5 | 4 | 4 | 1 |
| 2885 | - | 3164810 | 3163743 | - | pslL | PA14_35600 | hypothetical protein | 9 | 9 | 10 | 9 | 10 | 12 | 1 |
| 2886 | - | 3166302 | 3164893 | - | pslK | PA14_35620 | hypothetical protein | 6 | 5 | 6 | 8 | 6 | 6 | 1 |
| 2887 | - | 3167740 | 3166304 | - | pslJ | PA14_35630 | hypothetical protein | 6 | 6 | 5 | 8 | 6 | 8 | 1 |
| 2888 | - | 3168846 | 3167743 | - | pslI | PA14_35640 | transferase | 8 | 8 | 9 | 11 | 9 | 14 | 1 |
| 2889 | - | 3170045 | 3168837 | - | pslH | PA14_35650 | hypothetical protein | 10 | 9 | 10 | 14 | 11 | 15 | 1 |
| 2890 | - | 3171382 | 3170054 | - | pslG | PA14_35670 | glycosyl hydrolase | 7 | 6 | 7 | 8 | 9 | 11 | 1 |
| 2891 | - | 3172559 | 3171372 | - | pslF | PA14_35680 | hypothetical protein | 5 | 4 | 3 | 5 | 4 | 4 | 1 |
| 2892 | - | 3174547 | 3172559 | - | pslE | PA14_35690 | hypothetical protein | 8 | 6 | 7 | 10 | 6 | 7 | 1 |
| 2893 | - | 3174603 | 3175307 | + | - | PA14_35700 | hypothetical protein | 32 | 28 | 26 | 26 | 29 | 30 | 1 |
| 2894 | - | 3176147 | 3175803 | - | - | PA14_35710 | hypothetical protein | 50 | 39 | 47 | 66 | 62 | 58 | 1 |
| 2895 | - | 3176615 | 3176214 | - | - | PA14_35720 | hypothetical protein | 31 | 31 | 27 | 24 | 23 | 26 | 1 |
| 2896 | - | 3177358 | 3176984 | - | - | PA14_35730 | hypothetical protein | 8 | 5 | 9 | 7 | 6 | 6 | 1 |
| 2897 | - | 3177507 | 3180521 | + | - | PA14_35740 | transposase | 33 | 33 | 29 | 50 | 29 | 35 | 1 |
| 2898 | - | 3180518 | 3180880 | + | - | PA14_35750 | tpnA repressor protein | 133 | 111 | 107 | 145 | 103 | 123 | 1 |
| 2899 | - | 3181524 | 3181060 | - | - | PA14_35760 | hypothetical protein | 298 | 202 | 205 | 236 | 245 | 256 | 1 |
| 2900 | - | 3182437 | 3181532 | - | - | PA14_35770 | hypothetical protein | 72 | 36 | 47 | 49 | 58 | 55 | 0.670565447 |
| 2901 | - | 3183162 | 3182434 | - | - | PA14_35780 | hypothetical protein | 64 | 51 | 40 | 30 | 41 | 34 | 1 |
| 2902 | - | 3184587 | 3183178 | - | - | PA14_35790 | homospermidine synthase | 108 | 64 | 66 | 60 | 82 | 74 | 1 |
| 2903 | - | 3186315 | 3184999 | - | - | PA14_35800 | hypothetical protein | 113 | 106 | 98 | 93 | 96 | 103 | 1 |
| 2904 | - | 3187047 | 3186337 | - | - | PA14_35810 | hypothetical protein | 333 | 357 | 315 | 337 | 306 | 322 | 1 |
| 2905 | 3188223 | 3188081 | 3187110 | - | tnpS | PA14_35820 | cointegrate resolution protein S | 244 | 195 | 256 | 246 | 238 | 245 | 1 |
| 2906 | 3188223 | 3188265 | 3189263 | + | tnpT | PA14_35830 | cointegrate resolution protein T | 108 | 107 | 89 | 91 | 101 | 96 | 1 |
| 2907 | 3188223 | 3189302 | 3189946 | + | - | PA14_35840 | hypothetical protein | 257 | 221 | 211 | 248 | 262 | 280 | 1 |
| 2908 | 3188223 | 3189927 | 3190406 | + | - | PA14_35850 | hypothetical protein | 359 | 351 | 312 | 401 | 380 | 406 | 1 |
| 2909 | - | 3190988 | 3192292 | + | - | PA14_35860 | amino acid permease | 11 | 11 | 12 | 11 | 9 | 11 | 1 |
| 2910 | - | 3192371 | 3193795 | + | - | PA14_35880 | gamma-aminobutyraldehyde dehydrogenase | 20 | 14 | 18 | 21 | 16 | 22 | 1 |
| 2911 | - | 3193841 | 3195094 | + | - | PA14_35890 | diaminobutyrate--2-oxoglutarate aminotransferase | 12 | 10 | 11 | 10 | 11 | 14 | 1 |
| 2912 | - | 3195183 | 3196103 | + | - | PA14_35900 | dehydrogenase | 12 | 12 | 13 | 16 | 12 | 15 | 1 |
| 2913 | - | 3198054 | 3196396 | - | - | PA14_35920 | acetate permease | 13 | 12 | 15 | 17 | 10 | 10 | 1 |
| 2914 | - | 3198149 | 3198051 | - | - | PA14_35930 | hypothetical protein | 7 | 11 | 12 | 12 | 7 | 9 | 1 |
| 2915 | - | 3200078 | 3198432 | - | - | PA14_35940 | acyl-CoA synthetase | 9 | 9 | 10 | 11 | 7 | 9 | 1 |
| 2916 | - | 3200932 | 3200165 | - | - | PA14_35950 | dehydrogenase | 4 | 6 | 6 | 6 | 7 | 5 | 1 |
| 2917 | - | 3202104 | 3200932 | - | - | PA14_35970 | acyl-CoA dehydrogenase | 3 | 4 | 4 | 4 | 3 | 3 | 1 |
| 2918 | - | 3202511 | 3202155 | - | - | PA14_35980 | acyl-CoA dehydrogenase | 2 | 4 | 8 | 7 | 5 | 6 | 0.793216635 |
| 2919 | - | 3203704 | 3202508 | - | - | PA14_35990 | FadE36, aminoglycoside phosphotransferase | 3 | 3 | 4 | 3 | 2 | 4 | 1 |
| 2920 | - | 3204291 | 3205712 | + | prpR | PA14_36000 | propionate catabolism operon regulator | 43 | 39 | 36 | 36 | 38 | 38 | 1 |
| 2921 | - | 3206281 | 3205778 | - | - | PA14_36010 | hypothetical protein | 29 | 27 | 25 | 25 | 27 | 28 | 1 |
| 2922 | - | 3208643 | 3206340 | - | - | PA14_36020 | paraquat-inducible protein B | 38 | 35 | 31 | 39 | 32 | 33 | 1 |
| 2923 | - | 3209256 | 3208636 | - | - | PA14_36030 | paraquat-inducible protein A | 19 | 18 | 19 | 20 | 16 | 20 | 1 |
| 2924 | - | 3211497 | 3209914 | - | - | PA14_36050 | aldehyde dehydrogenase | 3 | 4 | 3 | 3 | 3 | 3 | 1 |
| 2925 | - | 3212621 | 3211626 | - | - | PA14_36060 | hypothetical protein | 6 | 6 | 6 | 6 | 6 | 6 | 1 |
| 2926 | - | 3213824 | 3212649 | - | - | PA14_36070 | hypothetical protein | 5 | 5 | 4 | 5 | 4 | 4 | 1 |
| 2927 | - | 3215177 | 3213855 | - | - | PA14_36080 | MFS transporter | 1 | 1 | 2 | 1 | 1 | 1 | 1 |
| 2928 | - | 3216613 | 3215363 | - | - | PA14_36090 | porin | 2 | 2 | 2 | 2 | 1 | 2 | 1 |
| 2929 | - | 3217799 | 3216786 | - | pdxA | PA14_36100 | 4-hydroxythreonine-4-phosphate dehydrogenase | 10 | 10 | 10 | 10 | 8 | 8 | 1 |
| 2930 | - | 3218755 | 3217796 | - | - | PA14_36110 | hypothetical protein | 6 | 8 | 6 | 6 | 6 | 6 | 1 |
| 2931 | - | 3220073 | 3218748 | - | - | PA14_36120 | MFS transporter | 3 | 3 | 4 | 3 | 3 | 4 | 1 |
| 2932 | - | 3220256 | 3221353 | + | - | PA14_36130 | hypothetical protein | 2 | 3 | 5 | 5 | 2 | 3 | 1 |
| 2933 | - | 3221381 | 3221911 | + | - | PA14_36150 | hypothetical protein | 1 | 1 | 2 | 2 | 1 | 1 | 1 |
| 2934 | - | 3221908 | 3223428 | + | - | PA14_36170 | hypothetical protein | 2 | 2 | 2 | 3 | 2 | 2 | 1 |
| 2935 | - | 3223496 | 3224443 | + | - | PA14_36180 | LysR family transcriptional regulator | 30 | 29 | 35 | 32 | 30 | 31 | 1 |
| 2936 | - | 3224525 | 3225001 | + | - | PA14_36190 | hypothetical protein | 19 | 18 | 19 | 17 | 17 | 17 | 1 |
| 2937 | - | 3225300 | 3226106 | + | - | PA14_36200 | ABC transporter substrate-binding protein | 20 | 42 | 10 | 12 | 21 | 10 | 0.225704119 |
| 2938 | - | 3226187 | 3226903 | + | - | PA14_36220 | amino acid permease | 3 | 4 | 3 | 2 | 3 | 2 | 1 |
| 2939 | - | 3226905 | 3227582 | + | - | PA14_36230 | amino acid ABC transporter permease | 13 | 16 | 13 | 12 | 15 | 11 | 1 |
| 2940 | - | 3228482 | 3227598 | - | - | PA14_36250 | hypothetical protein | 19 | 25 | 22 | 20 | 18 | 14 | 1 |

| | A | B | C | D | E | F | G | H | I | J | K | L | M | N |
|---|---|---|---|---|---|---|---|---|---|---|---|---|---|---|
| 2941 | - | 3230203 | 3228608 | - | - | *PA14_36260* | signal transduction protein | 14 | 15 | 15 | 18 | 14 | 12 | 1 |
| 2942 | - | 3231167 | 3230292 | - | - | *PA14_36270* | dehydrogenase | 52 | 61 | 64 | 47 | 37 | 37 | 1 |
| 2943 | - | 3231517 | 3231173 | - | - | *PA14_36280* | antibiotic biosynthesis monooxygenase | 13 | 26 | 28 | 14 | 16 | 10 | 0.339012339 |
| 2944 | - | 3232551 | 3231514 | - | - | *PA14_36290* | hypothetical protein | 22 | 38 | 32 | 26 | 24 | 17 | 0.723542393 |
| 2945 | - | 3233266 | 3232682 | - | - | *PA14_36300* | TetR family transcriptional regulator | 104 | 30 | 68 | 46 | 40 | 85 | 0.00024672 |
| 2946 | 3236436 | 3234672 | 3233419 | - | *hcnC* | *PA14_36310* | hydrogen cyanide synthase HcnC | 498 | 60 | 187 | 132 | 172 | 391 | 7.58162E-17 |
| 2947 | 3236436 | 3236069 | 3234675 | - | *hcnB* | *PA14_36320* | hydrogen cyanide synthase HcnB | 226 | 28 | 87 | 69 | 121 | 167 | 1.89869E-23 |
| 2948 | 3236436 | 3236380 | 3236066 | - | *hcnA* | *PA14_36330* | hydrogen cyanide synthase HcnA | 214 | 20 | 87 | 58 | 70 | 141 | 6.8676E-100 |
| 2949 | 3236436 | 3236744 | 3237988 | + | *exoY* | *PA14_36345* | adenylate cyclase | 64 | 45 | 74 | 85 | 50 | 71 | 1 |
| 2950 | - | 3238539 | 3237985 | - | - | *PA14_36350* | hypothetical protein | 69 | 57 | 80 | 100 | 66 | 86 | 1 |
| 2951 | - | 3239118 | 3238696 | - | - | *PA14_36360* | hypothetical protein | 52 | 46 | 64 | 78 | 53 | 61 | 1 |
| 2952 | - | 3240360 | 3239146 | - | - | *PA14_36370* | carboxylate-amine ligase | 25 | 28 | 39 | 40 | 34 | 33 | 1 |
| 2953 | - | 3240498 | 3241922 | + | - | *PA14_36375* | hypothetical protein | 7 | 10 | 10 | 11 | 9 | 10 | 1 |
| 2954 | - | 3241906 | 3242856 | + | - | *PA14_36390* | hypothetical protein | 12 | 15 | 17 | 17 | 14 | 14 | 1 |
| 2955 | - | 3243160 | 3242840 | - | - | *PA14_36400* | hypothetical protein | 7 | 10 | 12 | 14 | 13 | 11 | 1 |
| 2956 | - | 3243223 | 3243831 | + | - | *PA14_36410* | hypothetical protein | 18 | 27 | 25 | 33 | 22 | 33 | 1 |
| 2957 | - | 3245934 | 3243835 | - | - | *PA14_36420* | sensor/response regulator hybrid | 30 | 33 | 34 | 44 | 35 | 43 | 1 |
| 2958 | - | 3246017 | 3246607 | + | - | *PA14_36450* | hypothetical protein | 86 | 89 | 103 | 68 | 87 | 75 | 1 |
| 2959 | - | 3246620 | 3246967 | + | - | *PA14_36460* | hypothetical protein | 53 | 60 | 70 | 59 | 77 | 70 | 1 |
| 2960 | - | 3247252 | 3247560 | + | - | *PA14_36470* | hypothetical protein | 56 | 81 | 106 | 67 | 73 | 89 | 1 |
| 2961 | - | 3247812 | 3247591 | - | - | *PA14_36480* | hypothetical protein | 10 | 19 | 16 | 22 | 15 | 21 | 0.860626969 |
| 2962 | - | 3248218 | 3247868 | - | - | *PA14_36490* | hypothetical protein | 27 | 42 | 38 | 56 | 36 | 40 | 1 |
| 2963 | - | 3249321 | 3248245 | - | - | *PA14_36500* | hypothetical protein | 40 | 64 | 68 | 79 | 57 | 56 | 0.934705131 |
| 2964 | - | 3249795 | 3249325 | - | - | *PA14_36520* | hypothetical protein | 46 | 73 | 72 | 91 | 75 | 71 | 0.970587038 |
| 2965 | - | 3250449 | 3249997 | - | - | *PA14_36530* | hypothetical protein | 26 | 53 | 48 | 57 | 36 | 40 | 0.317658547 |
| 2966 | - | 3251351 | 3250575 | - | - | *PA14_36540* | hypothetical protein | 16 | 14 | 15 | 24 | 16 | 18 | 1 |
| 2967 | - | 3252445 | 3251348 | - | - | *PA14_36550* | hypothetical protein | 29 | 26 | 34 | 40 | 29 | 32 | 1 |
| 2968 | - | 3253139 | 3252774 | - | - | *PA14_36560* | hypothetical protein | 96 | 87 | 65 | 93 | 63 | 71 | 1 |
| 2969 | - | 3253507 | 3255048 | + | *glgA* | *PA14_36570* | glycogen synthase | 38 | 53 | 57 | 64 | 45 | 50 | 1 |
| 2970 | - | 3255048 | 3256799 | + | - | *PA14_36580* | glycosyl hydrolase | 27 | 43 | 48 | 64 | 49 | 47 | 0.955702824 |
| 2971 | - | 3256792 | 3258846 | + | - | *PA14_36590* | 4-alpha-glucanotransferase | 21 | 36 | 36 | 50 | 43 | 39 | 0.831807123 |
| 2972 | - | 3258839 | 3261619 | + | - | *PA14_36605* | maltooligosyl trehalose synthase | 23 | 49 | 39 | 64 | 51 | 52 | 0.436346238 |
| 2973 | - | 3261616 | 3261921 | + | - | *PA14_36620* | hypothetical protein | 9 | 29 | 23 | 36 | 31 | 29 | 0.000287258 |
| 2974 | - | 3261934 | 3264084 | + | - | *PA14_36630* | glycosyl hydrolase | 21 | 50 | 34 | 60 | 48 | 48 | 0.139802753 |
| 2975 | - | 3264184 | 3264600 | + | - | *PA14_36650* | hypothetical protein | 40 | 106 | 73 | 122 | 98 | 99 | 0.022142733 |
| 2976 | - | 3264667 | 3265914 | + | - | *PA14_36660* | alcohol dehydrogenase | 17 | 34 | 30 | 44 | 32 | 34 | 0.418246621 |
| 2977 | - | 3265918 | 3266856 | + | - | *PA14_36670* | hypothetical protein | 17 | 27 | 32 | 41 | 31 | 32 | 0.931437202 |
| 2978 | - | 3266853 | 3267590 | + | - | *PA14_36680* | hypothetical protein | 13 | 24 | 22 | 31 | 21 | 19 | 0.580848337 |
| 2979 | - | 3267587 | 3268792 | + | - | *PA14_36690* | cardiolipin synthase 2 | 20 | 29 | 29 | 47 | 34 | 37 | 1 |
| 2980 | - | 3268789 | 3269784 | + | - | *PA14_36700* | hypothetical protein | 11 | 27 | 23 | 35 | 30 | 29 | 0.0839756 |
| 2981 | - | 3271984 | 3269786 | - | *glgB* | *PA14_36710* | glycogen branching protein | 23 | 47 | 42 | 61 | 49 | 50 | 0.415011252 |
| 2982 | - | 3275283 | 3271981 | - | - | *PA14_36730* | trehalose synthase | 28 | 46 | 41 | 58 | 50 | 51 | 1 |
| 2983 | - | 3277288 | 3275294 | - | - | *PA14_36740* | hypothetical protein | 26 | 33 | 34 | 41 | 36 | 37 | 1 |
| 2984 | - | 3278313 | 3277432 | - | - | *PA14_36760* | KU domain-containing protein | 10 | 16 | 13 | 17 | 15 | 17 | 1 |
| 2985 | - | 3278578 | 3278336 | - | - | *PA14_36770* | hypothetical protein | 5 | 10 | 12 | 13 | 9 | 10 | 0.828709947 |
| 2986 | - | 3279083 | 3278592 | - | - | *PA14_36780* | hypothetical protein | 33 | 48 | 63 | 67 | 60 | 54 | 1 |
| 2987 | - | 3279214 | 3279080 | - | - | *PA14_36790* | hypothetical protein | 41 | 71 | 86 | 89 | 74 | 67 | 0.849496297 |
| 2988 | - | 3281440 | 3279311 | - | *katE* | *PA14_36810* | hydroperoxidase II | 11 | 18 | 12 | 19 | 17 | 13 | 0.967915794 |
| 2989 | - | 3281688 | 3281521 | - | - | *PA14_36820* | hypothetical protein | 16 | 66 | 48 | 157 | 47 | 90 | 1.60548E-08 |
| 2990 | - | 3282187 | 3282585 | + | - | *PA14_36830* | hypothetical protein | 16 | 27 | 30 | 42 | 32 | 31 | 0.895687725 |
| 2991 | - | 3285030 | 3282592 | - | *glgP* | *PA14_36840* | glycogen phosphorylase | 34 | 49 | 51 | 62 | 53 | 54 | 1 |
| 2992 | - | 3285370 | 3285083 | - | - | *PA14_36850* | hypothetical protein | 205 | 298 | 530 | 528 | 380 | 316 | 1 |
| 2993 | - | 3285614 | 3285802 | + | - | *PA14_36860* | hypothetical protein | 40 | 61 | 65 | 64 | 63 | 81 | 1 |
| 2994 | - | 3286682 | 3285822 | - | - | *PA14_36870* | short-chain dehydrogenase | 26 | 40 | 41 | 46 | 38 | 48 | 1 |
| 2995 | - | 3287226 | 3286708 | - | - | *PA14_36880* | ompetence-damaged protein | 4 | 9 | 12 | 11 | 9 | 10 | 0.181497819 |
| 2996 | - | 3287476 | 3287237 | - | - | *PA14_36890* | metallothionein | 9 | 15 | 17 | 14 | 15 | 15 | 1 |
| 2997 | - | 3287711 | 3287496 | - | - | *PA14_36900* | hypothetical protein | 8 | 14 | 13 | 15 | 11 | 10 | 0.907738182 |
| 2998 | - | 3290309 | 3287787 | - | *ligD* | *PA14_36910* | ATP-dependent DNA ligase | 13 | 9 | 12 | 12 | 12 | 13 | 1 |
| 2999 | - | 3290863 | 3290327 | - | - | *PA14_36920* | hypothetical protein | 14 | 6 | 10 | 7 | 5 | 8 | 0.023098865 |
| 3000 | - | 3290965 | 3291489 | + | - | *PA14_36930* | hypothetical protein | 32 | 30 | 40 | 18 | 18 | 25 | 1 |
| 3001 | - | 3291515 | 3291964 | + | - | *PA14_36940* | hypothetical protein | 4 | 8 | 10 | 9 | 7 | 8 | 0.45673241 |
| 3002 | - | 3293306 | 3291942 | - | - | *PA14_36960* | transporter | 7 | 11 | 13 | 14 | 12 | 13 | 1 |
| 3003 | - | 3293907 | 3293335 | - | - | *PA14_36980* | hypothetical protein | 23 | 38 | 46 | 42 | 35 | 34 | 1 |
| 3004 | - | 3294857 | 3294000 | - | - | *PA14_36990* | EAL domain-containing protein | 4 | 3 | 4 | 4 | 3 | 2 | 1 |
| 3005 | - | 3295567 | 3294854 | - | *cupA5* | *PA14_37000* | chaperone CupA5 | 2 | 2 | 2 | 3 | 1 | 1 | 1 |
| 3006 | - | 3296918 | 3295557 | - | *cupA4* | *PA14_37010* | fimbrial subunit CupA4 | 2 | 2 | 3 | 3 | 2 | 2 | 1 |
| 3007 | - | 3299533 | 3296915 | - | *cupA3* | *PA14_37030* | usher | 7 | 6 | 7 | 8 | 5 | 4 | 1 |
| 3008 | - | 3300263 | 3299517 | - | *cupA2* | *PA14_37040* | chaperone CupA2 | 4 | 3 | 4 | 3 | 2 | 2 | 1 |
| 3009 | - | 3300902 | 3300351 | - | *cupA1* | *PA14_37060* | fimbrial subunit CupA1 | 16 | 16 | 11 | 11 | 12 | 6 | 1 |
| 3010 | - | 3301756 | 3302982 | + | - | *PA14_37070* | hypothetical protein | 146 | 131 | 184 | 87 | 76 | 125 | 1 |
| 3011 | - | 3303409 | 3304044 | + | - | *PA14_37080* | hypothetical protein | 81 | 97 | 104 | 50 | 56 | 69 | 1 |
| 3012 | - | 3305528 | 3304080 | - | - | *PA14_37090* | aldehyde dehydrogenase | 15 | 16 | 18 | 13 | 12 | 14 | 1 |
| 3013 | - | 3307187 | 3305550 | - | - | *PA14_37100* | dehydrogenase | 6 | 5 | 7 | 5 | 5 | 5 | 1 |
| 3014 | - | 3308224 | 3307292 | - | - | *PA14_37120* | LysR family transcriptional regulator | 18 | 17 | 17 | 19 | 14 | 20 | 1 |
| 3015 | - | 3309397 | 3308261 | - | - | *PA14_37130* | hypothetical protein | 11 | 12 | 11 | 12 | 12 | 14 | 1 |
| 3016 | - | 3309523 | 3310428 | + | - | *PA14_37140* | LysR family transcriptional regulator | 10 | 12 | 11 | 10 | 11 | 10 | 1 |
| 3017 | - | 3310576 | 3311004 | + | - | *PA14_37150* | hypothetical protein | 6 | 11 | 8 | 8 | 10 | 9 | 1 |
| 3018 | - | 3311021 | 3312133 | + | - | *PA14_37170* | hypothetical protein | 27 | 25 | 28 | 32 | 28 | 25 | 1 |
| 3019 | - | 3313744 | 3312668 | - | *ada* | *PA14_37190* | O6-methylguanine-DNA methyltransferase | 20 | 28 | 24 | 24 | 19 | 16 | 1 |
| 3020 | - | 3313936 | 3314919 | + | - | *PA14_37200* | hypothetical protein | 17 | 76 | 37 | 76 | 57 | 37 | 1.49732E-07 |
| 3021 | - | 3315701 | 3314904 | - | - | *PA14_37210* | hypothetical protein | 17 | 52 | 30 | 46 | 34 | 30 | 0.003189572 |
| 3022 | - | 3316761 | 3315808 | - | - | *PA14_37220* | LysR family transcriptional regulator | 3 | 4 | 4 | 3 | 4 | 2 | 1 |
| 3023 | - | 3317016 | 3318287 | + | - | *PA14_37250* | major facilitator transporter | 4 | 26 | 11 | 17 | 16 | 10 | 3.27339E-21 |
| 3024 | - | 3318312 | 3319541 | + | - | *PA14_37260* | porin | 9 | 18 | 12 | 14 | 12 | 13 | 0.445579457 |

| | A | B | C | D | E | F | G | H | I | J | K | L | M | N |
|---|---|---|---|---|---|---|---|---|---|---|---|---|---|---|
| 3025 | - | 3319574 | 3320317 | + | - | *PA14_37270* | LamB/YcsF family protein | 6 | 40 | 16 | 25 | 19 | 14 | 2.33936E-22 |
| 3026 | - | 3320314 | 3321027 | + | - | *PA14_37290* | hypothetical protein | 3 | 25 | 11 | 16 | 15 | 9 | 1.01442E-37 |
| 3027 | - | 3321024 | 3321965 | + | - | *PA14_37310* | hypothetical protein | 6 | 29 | 14 | 25 | 19 | 13 | 4.51048E-12 |
| 3028 | - | 3322033 | 3322509 | + | - | *PA14_37320* | hypothetical protein | 10 | 19 | 18 | 20 | 17 | 13 | 0.530445973 |
| 3029 | - | 3324300 | 3322528 | - | - | *PA14_37340* | thiamine pyrophosphate protein | 9 | 13 | 13 | 16 | 14 | 13 | 1 |
| 3030 | - | 3324471 | 3324869 | + | - | *PA14_37350* | hypothetical protein | 6 | 9 | 11 | 11 | 8 | 9 | 1 |
| 3031 | - | 3326026 | 3325268 | - | - | *PA14_37360* | short chain dehydrogenase | 8 | 11 | 9 | 11 | 8 | 10 | 1 |
| 3032 | - | 3326940 | 3326011 | - | - | *PA14_37370* | esterase | 4 | 5 | 3 | 3 | 3 | 4 | 1 |
| 3033 | - | 3328426 | 3326951 | - | - | *PA14_37380* | flavin-binding monooxygenase | 3 | 4 | 4 | 4 | 3 | 4 | 1 |
| 3034 | - | 3328573 | 3329607 | + | - | *PA14_37400* | AraC family transcriptional regulator | 16 | 14 | 17 | 15 | 12 | 14 | 1 |
| 3035 | - | 3329756 | 3330601 | + | - | *PA14_37410* | hypothetical protein | 9 | 10 | 9 | 11 | 8 | 11 | 1 |
| 3036 | - | 3331559 | 3330603 | - | - | *PA14_37420* | transmembrane sensor protein | 10 | 8 | 8 | 7 | 9 | 5 | 1 |
| 3037 | - | 3332065 | 3331556 | - | - | *PA14_37430* | RNA polymerase sigma factor | 8 | 8 | 7 | 5 | 7 | 6 | 1 |
| 3038 | - | 3333364 | 3332168 | - | - | *PA14_37440* | MFS transporter | 19 | 13 | 20 | 18 | 18 | 18 | 1 |
| 3039 | - | 3334634 | 3333351 | - | - | *PA14_37460* | permease | 7 | 6 | 5 | 5 | 8 | 6 | 1 |
| 3040 | - | 3335710 | 3334631 | - | - | *PA14_37470* | flavin-dependent oxidoreductase | 8 | 6 | 5 | 4 | 5 | 5 | 1 |
| 3041 | - | 3338374 | 3335723 | - | - | *PA14_37490* | TonB-dependent receptor | 23 | 21 | 22 | 21 | 18 | 20 | 1 |
| 3042 | - | 3339314 | 3338466 | - | - | *PA14_37510* | hypothetical protein | 12 | 10 | 10 | 9 | 10 | 10 | 1 |
| 3043 | - | 3339954 | 3339298 | - | - | *PA14_37520* | hypothetical protein | 10 | 7 | 5 | 6 | 7 | 8 | 1 |
| 3044 | - | 3340853 | 3339951 | - | - | *PA14_37530* | hydrolase | 8 | 8 | 6 | 7 | 6 | 6 | 1 |
| 3045 | - | 3341373 | 3340864 | - | - | *PA14_37550* | ring-hydroxylating dioxygenase small subunit | 11 | 9 | 8 | 7 | 10 | 7 | 1 |
| 3046 | - | 3343233 | 3341401 | - | - | *PA14_37560* | asparagine synthetase, glutamine-hydrolysing | 6 | 7 | 4 | 3 | 6 | 4 | 1 |
| 3047 | - | 3344595 | 3343321 | - | - | *PA14_37570* | ring-hydroxylating dioxygenase, large terminal subunit | 11 | 13 | 12 | 12 | 16 | 13 | 1 |
| 3048 | - | 3345273 | 3344797 | - | - | *PA14_37580* | leucine-responsive regulatory protein | 41 | 39 | 42 | 37 | 45 | 51 | 1 |
| 3049 | - | 3345406 | 3346047 | + | *kynB* | *PA14_37590* | kynurenine formamidase, KynB | 28 | 43 | 35 | 32 | 36 | 29 | 1 |
| 3050 | - | 3346051 | 3347301 | + | - | *PA14_37610* | kynureninase | 38 | 58 | 40 | 41 | 45 | 37 | 1 |
| 3051 | - | 3347457 | 3348863 | + | - | *PA14_37630* | amino acid permease | 14 | 17 | 13 | 14 | 16 | 14 | 1 |
| 3052 | - | 3349149 | 3351023 | + | - | *PA14_37640* | hypothetical protein | 8 | 8 | 9 | 10 | 9 | 9 | 1 |
| 3053 | - | 3351063 | 3352967 | + | - | *PA14_37650* | hypothetical protein | 8 | 8 | 9 | 8 | 8 | 7 | 1 |
| 3054 | - | 3353906 | 3353004 | - | - | *PA14_37660* | transcriptional regulator | 14 | 19 | 20 | 19 | 18 | 15 | 1 |
| 3055 | - | 3354363 | 3354629 | + | - | *PA14_37670* | hypothetical protein | 41 | 40 | 65 | 39 | 35 | 43 | 1 |
| 3056 | - | 3354806 | 3356473 | + | - | *PA14_37680* | hypothetical protein | 17 | 16 | 22 | 16 | 18 | 21 | 1 |
| 3057 | - | 3359460 | 3356866 | - | - | *PA14_37690* | sensory box protein | 57 | 48 | 65 | 93 | 58 | 91 | 1 |
| 3058 | - | 3361740 | 3359632 | - | *fusA2* | *PA14_37710* | elongation factor G | 116 | 81 | 106 | 153 | 98 | 153 | 1 |
| 3059 | - | 3362016 | 3364658 | + | - | *PA14_37730* | TonB dependent receptor | 27 | 15 | 21 | 31 | 22 | 32 | 1 |
| 3060 | 3364761 | 3364838 | 3366562 | + | - | *PA14_37745* | carbamoyl transferase | 204 | 5 | 13 | 135 | 82 | 232 | 0 |
| 3061 | 3364761 | 3366606 | 3367769 | + | - | *PA14_37760* | MFS transporter | 115 | 7 | 12 | 71 | 57 | 127 | 3.8621E-279 |
| 3062 | 3364761 | 3367771 | 3368439 | + | - | *PA14_37770* | hydrolase | 195 | 18 | 27 | 128 | 88 | 205 | 7.70864E-77 |
| 3063 | 3364761 | 3368436 | 3369074 | + | - | *PA14_37780* | hypothetical protein | 110 | 16 | 20 | 76 | 56 | 122 | 2.0367E-25 |
| 3064 | - | 3369223 | 3371043 | + | *pcoA* | *PA14_37790* | copper resistance protein A | 9 | 7 | 8 | 10 | 7 | 9 | 1 |
| 3065 | - | 3371040 | 3372095 | + | *pcoB* | *PA14_37810* | copper resistance protein B | 8 | 4 | 4 | 5 | 6 | 6 | 0.037558237 |
| 3066 | - | 3373338 | 3372115 | - | - | *PA14_37820* | hypothetical protein | 9 | 9 | 10 | 7 | 10 | 7 | 1 |
| 3067 | - | 3374722 | 3373541 | - | - | *PA14_37830* | pyridoxal-phosphate dependent protein | 10 | 15 | 10 | 12 | 12 | 8 | 1 |
| 3068 | - | 3376478 | 3374868 | - | - | *PA14_37840* | ABC transporter ATP-binding protein | 9 | 8 | 10 | 11 | 7 | 8 | 1 |
| 3069 | - | 3377496 | 3376480 | - | - | *PA14_37850* | ABC transporter permease | 1 | 1 | 1 | 1 | 1 | 1 | 1 |
| 3070 | - | 3378571 | 3377498 | - | - | *PA14_37870* | peptide ABC transporter permease | 3 | 2 | 3 | 3 | 2 | 2 | 1 |
| 3071 | - | 3380381 | 3378573 | - | - | *PA14_37880* | ABC transporter substrate-binding protein | 3 | 3 | 2 | 3 | 3 | 3 | 1 |
| 3072 | - | 3382925 | 3380385 | - | - | *PA14_37900* | TonB-dependent receptor | 4 | 5 | 6 | 7 | 4 | 6 | 1 |
| 3073 | - | 3384506 | 3383604 | - | - | *PA14_37910* | LysR family transcriptional regulator | 5 | 8 | 5 | 6 | 8 | 4 | 1 |
| 3074 | - | 3384617 | 3386032 | + | - | *PA14_37915* | major facilitator transporter | 9 | 8 | 8 | 5 | 6 | 6 | 1 |
| 3075 | - | 3386926 | 3386039 | - | *cynR* | *PA14_37940* | DNA-binding transcriptional regulator CynR | 25 | 27 | 27 | 18 | 20 | 20 | 1 |
| 3076 | - | 3387041 | 3387703 | + | *cynT* | *PA14_37950* | carbonate dehydratase | 10 | 10 | 10 | 11 | 9 | 9 | 1 |
| 3077 | - | 3387743 | 3388213 | + | *cynS* | *PA14_37965* | cyanate hydratase | 12 | 19 | 11 | 14 | 15 | 15 | 1 |
| 3078 | - | 3389204 | 3388251 | - | - | *PA14_37980* | Fe2+-dicitrate sensor, membrane protein | 3 | 4 | 2 | 2 | 2 | 2 | 1 |
| 3079 | - | 3389707 | 3389201 | - | - | *PA14_37990* | RNA polymerase sigma factor | 1 | 2 | 2 | 2 | 2 | 3 | 1 |
| 3080 | - | 3390035 | 3390499 | + | - | *PA14_38000* | hypothetical protein | 21 | 13 | 24 | 13 | 11 | 18 | 0.434308967 |
| 3081 | - | 3390598 | 3392031 | + | - | *PA14_38010* | hypothetical protein | 42 | 38 | 39 | 41 | 38 | 37 | 1 |
| 3082 | - | 3392496 | 3392089 | - | - | *PA14_38020* | ntibiotic biosynthesis monooxygenase | 24 | 24 | 28 | 24 | 16 | 19 | 1 |
| 3083 | - | 3392910 | 3393899 | + | - | *PA14_38040* | AraC family transcriptional regulator | 28 | 23 | 22 | 24 | 20 | 23 | 1 |
| 3084 | - | 3394935 | 3395342 | + | - | *PA14_38050* | hypothetical protein | 20 | 14 | 24 | 20 | 27 | 20 | 0.958024133 |
| 3085 | - | 3395401 | 3395661 | + | - | *PA14_38060* | hypothetical protein | 21 | 18 | 20 | 18 | 18 | 17 | 1 |
| 3086 | - | 3395836 | 3397710 | + | - | *PA14_38080* | hypothetical protein | 34 | 40 | 35 | 38 | 35 | 33 | 1 |
| 3087 | - | 3398770 | 3397868 | - | - | *PA14_38090* | pseudouridylate synthase | 36 | 40 | 41 | 34 | 37 | 27 | 1 |
| 3088 | - | 3400120 | 3398891 | - | - | *PA14_38110* | serine/threonine transporter SstT | 32 | 36 | 34 | 37 | 38 | 23 | 1 |
| 3089 | - | 3401978 | 3400608 | - | - | *PA14_38130* | amino acid permease | 14 | 19 | 15 | 18 | 14 | 13 | 1 |
| 3090 | - | 3403511 | 3402135 | - | - | *PA14_38140* | glutamine synthetase | 50 | 49 | 41 | 51 | 39 | 48 | 1 |
| 3091 | - | 3404017 | 3404775 | + | - | *PA14_38160* | hypothetical protein | 17 | 13 | 17 | 15 | 13 | 11 | 1 |
| 3092 | - | 3404760 | 3405074 | + | - | *PA14_38170* | hypothetical protein | 19 | 25 | 26 | 25 | 25 | 23 | 1 |
| 3093 | - | 3406659 | 3405199 | - | - | *PA14_38180* | hypothetical protein | 9 | 11 | 9 | 11 | 13 | 10 | 1 |
| 3094 | - | 3407320 | 3406799 | - | - | *PA14_38190* | hypothetical protein | 2 | 1 | 1 | 2 | 1 | 0 | 0.853730745 |
| 3095 | - | 3408055 | 3409713 | + | - | *PA14_38200* | thiamine pyrophosphate protein | 34 | 33 | 34 | 40 | 33 | 31 | 1 |
| 3096 | - | 3410395 | 3409721 | - | - | *PA14_38210* | hypothetical protein | 27 | 37 | 26 | 83 | 31 | 22 | 1 |
| 3097 | - | 3411303 | 3410395 | - | - | *PA14_38220* | hypothetical protein | 24 | 40 | 25 | 110 | 38 | 26 | 0.831724413 |
| 3098 | - | 3412862 | 3411438 | - | - | *PA14_38250* | transcriptional regulator | 36 | 29 | 31 | 38 | 27 | 31 | 1 |
| 3099 | - | 3413042 | 3413296 | + | - | *PA14_38260* | hypothetical protein | 85 | 25 | 37 | 57 | 39 | 96 | 1.16747E-06 |
| 3100 | - | 3413293 | 3413550 | + | - | *PA14_38270* | hypothetical protein | 179 | 74 | 99 | 161 | 110 | 234 | 0.083747076 |
| 3101 | - | 3413924 | 3413625 | - | - | *PA14_38290* | hypothetical protein | 18 | 21 | 20 | 23 | 18 | 20 | 1 |
| 3102 | - | 3414421 | 3413948 | - | - | *PA14_38300* | transcriptional regulator | 11 | 8 | 10 | 7 | 7 | 10 | 1 |
| 3103 | - | 3414554 | 3414946 | + | - | *PA14_38310* | hypothetical protein | 4 | 3 | 6 | 5 | 4 | 7 | 1 |
| 3104 | - | 3415099 | 3416100 | + | - | *PA14_38320* | bile acid/Na+ symporter family transporter | 68 | 47 | 70 | 38 | 52 | 67 | 1 |
| 3105 | - | 3417510 | 3416155 | - | *gor* | *PA14_38330* | glutathione reductase | 259 | 236 | 250 | 222 | 246 | 263 | 1 |
| 3106 | - | 3417655 | 3418077 | + | - | *PA14_38340* | ring-cleaving dioxygenase | 130 | 176 | 185 | 191 | 235 | 224 | 1 |
| 3107 | - | 3419098 | 3418259 | - | *galU* | *PA14_38350* | UTP-glucose-1-phosphate uridylyltransferase | 191 | 144 | 199 | 138 | 151 | 144 | 1 |
| 3108 | - | 3420507 | 3419146 | - | - | *PA14_38360* | nucleotide sugar dehydrogenase | 12 | 12 | 19 | 17 | 15 | 14 | 1 |

| | A | B | C | D | E | F | G | H | I | J | K | L | M | N |
|---|---|---|---|---|---|---|---|---|---|---|---|---|---|---|
| 3109 | - | 3420659 | 3420883 | + | - | PA14_38370 | hypothetical protein | 24 | 28 | 31 | 33 | 33 | 34 | 1 |
| 3110 | - | 3421521 | 3420889 | - | - | PA14_38380 | transcriptional regulator | 70 | 62 | 71 | 71 | 66 | 61 | 1 |
| 3111 | - | 3421686 | 3422876 | + | - | PA14_38395 | periplasmic multidrug efflux lipoprotein | 7 | 7 | 7 | 5 | 6 | 7 | 1 |
| 3112 | - | 3422892 | 3426029 | + | amrB | PA14_38410 | multidrug efflux protein | 9 | 10 | 12 | 12 | 10 | 11 | 1 |
| 3113 | - | 3427200 | 3426271 | - | - | PA14_38420 | hypothetical protein | 46 | 43 | 47 | 51 | 51 | 54 | 1 |
| 3114 | - | 3427393 | 3427797 | + | gnyR | PA14_38430 | regulatory gene of gnyRDBHAL cluster, GnyR | 232 | 336 | 280 | 229 | 428 | 366 | 1 |
| 3115 | - | 3427846 | 3429009 | + | gnyD | PA14_38440 | citronelloyl-CoA dehydrogenase, GnyD | 432 | 689 | 446 | 537 | 637 | 617 | 1 |
| 3116 | - | 3429132 | 3430739 | + | gnyB | PA14_38460 | acyl-CoA carboxyltransferase subunit beta | 218 | 302 | 213 | 250 | 271 | 269 | 1 |
| 3117 | - | 3430753 | 3431550 | + | gnyH | PA14_38470 | gamma-carboxygeranoyl-CoA hydratase | 156 | 218 | 140 | 153 | 187 | 169 | 1 |
| 3118 | - | 3431547 | 3433514 | + | gnyA | PA14_38480 | alpha subunit of geranoyl-CoA carboxylase, GnyA | 267 | 365 | 247 | 310 | 326 | 320 | 1 |
| 3119 | - | 3433535 | 3434437 | + | gnyL | PA14_38490 | hydroxymethylglutaryl-CoA lyase | 456 | 505 | 419 | 470 | 506 | 497 | 1 |
| 3120 | - | 3435308 | 3434505 | - | - | PA14_38500 | IclR family transcriptional regulator | 87 | 103 | 126 | 90 | 125 | 109 | 1 |
| 3121 | - | 3435469 | 3436767 | + | hmgA | PA14_38510 | homogentisate 1,2-dioxygenase | 299 | 308 | 289 | 239 | 313 | 340 | 1 |
| 3122 | - | 3436772 | 3438070 | + | fahA | PA14_38530 | fumarylacetoacetase | 245 | 251 | 168 | 232 | 268 | 304 | 1 |
| 3123 | - | 3438067 | 3438705 | + | maiA | PA14_38550 | maleylacetoacetate isomerase | 283 | 273 | 201 | 264 | 273 | 320 | 1 |
| 3124 | - | 3438790 | 3440142 | + | - | PA14_38560 | MFS transporter | 16 | 10 | 11 | 12 | 12 | 16 | 1 |
| 3125 | - | 3440271 | 3441641 | + | - | PA14_38570 | transcriptional regulator | 18 | 12 | 13 | 12 | 13 | 16 | 1 |
| 3126 | - | 3441918 | 3443309 | + | - | PA14_38580 | hypothetical protein | 12 | 49 | 35 | 95 | 38 | 53 | 4.78316E-06 |
| 3127 | - | 3443343 | 3444113 | + | bdhA | PA14_38590 | 3-hydroxybutyrate dehydrogenase | 27 | 141 | 90 | 194 | 106 | 117 | 3.91329E-11 |
| 3128 | - | 3445728 | 3444304 | - | - | PA14_38610 | hypothetical protein | 23 | 40 | 31 | 42 | 36 | 42 | 0.767674398 |
| 3129 | - | 3447122 | 3445941 | - | atoB | PA14_38630 | acetyl-CoA acetyltransferase | 250 | 343 | 233 | 320 | 259 | 396 | 1 |
| 3130 | - | 3447928 | 3447272 | - | - | PA14_38640 | CoA transferase subunit B | 184 | 242 | 134 | 168 | 191 | 267 | 1 |
| 3131 | - | 3448661 | 3447963 | - | - | PA14_38660 | CoA transferase, subunit A | 382 | 401 | 273 | 309 | 323 | 436 | 1 |
| 3132 | - | 3448793 | 3449713 | + | - | PA14_38680 | LysR family transcriptional regulator | 15 | 16 | 16 | 18 | 18 | 19 | 1 |
| 3133 | - | 3449781 | 3451736 | + | - | PA14_38690 | acetoacetyl-CoA synthetase | 56 | 49 | 52 | 56 | 44 | 66 | 1 |
| 3134 | - | 3451794 | 3452072 | + | ppiC1 | PA14_38700 | peptidyl-prolyl cis-trans isomerase C1 | 59 | 57 | 63 | 73 | 48 | 55 | 1 |
| 3135 | - | 3452091 | 3452447 | + | - | PA14_38710 | hypothetical protein | 103 | 107 | 129 | 131 | 112 | 120 | 1 |
| 3136 | - | 3452444 | 3453007 | + | - | PA14_38720 | hypothetical protein | 35 | 33 | 37 | 36 | 28 | 36 | 1 |
| 3137 | - | 3453132 | 3454340 | + | - | PA14_38730 | major facilitator superfamily transporter | 13 | 10 | 14 | 11 | 8 | 14 | 1 |
| 3138 | - | 3456069 | 3454285 | - | - | PA14_38740 | two-component sensor | 33 | 39 | 27 | 34 | 31 | 35 | 1 |
| 3139 | - | 3457216 | 3456053 | - | - | PA14_38750 | iron-containing alcohol dehydrogenase | 26 | 31 | 24 | 27 | 27 | 27 | 1 |
| 3140 | - | 3459128 | 3457302 | - | - | PA14_38770 | peptidase | 67 | 38 | 55 | 53 | 44 | 53 | 1 |
| 3141 | - | 3460278 | 3459133 | - | pqqE | PA14_38780 | pyrroloquinoline quinone biosynthesis protein PqqE | 68 | 52 | 54 | 68 | 48 | 60 | 1 |
| 3142 | - | 3460528 | 3460250 | - | pqqD | PA14_38790 | pyrroloquinoline quinone biosynthesis protein PqqD | 43 | 33 | 35 | 40 | 36 | 37 | 1 |
| 3143 | - | 3461277 | 3460525 | - | pqqC | PA14_38800 | pyrroloquinoline quinone biosynthesis protein PqqC | 61 | 43 | 48 | 50 | 45 | 52 | 1 |
| 3144 | - | 3462201 | 3461287 | - | pqqB | PA14_38820 | pyrroloquinoline quinone biosynthesis protein PqqB | 37 | 26 | 28 | 29 | 25 | 25 | 1 |
| 3145 | - | 3462325 | 3462254 | - | pqqA | PA14_38825 | coenzyme PQQ synthesis protein PqqA | 56 | 52 | 53 | 54 | 38 | 58 | 1 |
| 3146 | - | 3464203 | 3462683 | - | - | PA14_38840 | NAD+ dependent acetaldehyde dehydrogenase | 80 | 150 | 108 | 94 | 94 | 88 | 1 |
| 3147 | - | 3464727 | 3464290 | - | exaB | PA14_38850 | cytochrome c550 | 3 | 3 | 4 | 4 | 3 | 5 | 1 |
| 3148 | - | 3465040 | 3466911 | + | exaA | PA14_38860 | quinoprotein alcohol dehydrogenase | 5 | 5 | 8 | 7 | 4 | 6 | 1 |
| 3149 | - | 3466965 | 3467612 | + | - | PA14_38880 | hypothetical protein | 4 | 2 | 4 | 3 | 4 | 3 | 1 |
| 3150 | - | 3468316 | 3467639 | - | - | PA14_38900 | two-component response regulator | 7 | 5 | 8 | 5 | 7 | 7 | 1 |
| 3151 | - | 3468979 | 3468329 | - | - | PA14_38910 | sensor kinase | 10 | 7 | 9 | 6 | 6 | 7 | 0.831724413 |
| 3152 | - | 3469164 | 3468988 | - | - | PA14_38920 | hypothetical protein | 10 | 8 | 11 | 9 | 7 | 10 | 1 |
| 3153 | - | 3469644 | 3470309 | + | - | PA14_38930 | transcriptional regulator | 17 | 13 | 19 | 15 | 12 | 12 | 1 |
| 3154 | - | 3470319 | 3471182 | + | - | PA14_38950 | hypothetical protein | 11 | 7 | 13 | 10 | 9 | 8 | 0.555553781 |
| 3155 | - | 3473829 | 3471184 | - | - | PA14_38970 | two-component sensor | 9 | 5 | 6 | 6 | 4 | 5 | 0.770507726 |
| 3156 | - | 3474922 | 3473759 | - | - | PA14_38990 | hypothetical protein | 10 | 7 | 8 | 8 | 4 | 5 | 0.956402005 |
| 3157 | - | 3476224 | 3475025 | - | - | PA14_39000 | hypothetical protein | 2 | 2 | 3 | 3 | 2 | 2 | 1 |
| 3158 | - | 3478818 | 3476491 | - | pqqF | PA14_39010 | pyrroloquinoline quinone biosynthesis protein F | 40 | 26 | 27 | 25 | 24 | 22 | 1 |
| 3159 | - | 3480598 | 3478895 | - | - | PA14_39020 | hypothetical protein | 5 | 5 | 5 | 5 | 3 | 5 | 1 |
| 3160 | - | 3481126 | 3482439 | + | braZ | PA14_39050 | branched-chain amino acid transport carrier | 9 | 8 | 10 | 8 | 7 | 7 | 1 |
| 3161 | - | 3482675 | 3482914 | + | - | PA14_39060 | lipoprotein | 31 | 29 | 30 | 22 | 30 | 23 | 1 |
| 3162 | - | 3483373 | 3482981 | - | - | PA14_39070 | hypothetical protein | 76 | 121 | 63 | 49 | 80 | 45 | 0.950259107 |
| 3163 | - | 3483653 | 3483420 | - | - | PA14_39080 | hypothetical protein | 26 | 24 | 25 | 17 | 20 | 14 | 1 |
| 3164 | - | 3483838 | 3484338 | + | - | PA14_39090 | hypothetical protein | 88 | 121 | 110 | 108 | 115 | 103 | 1 |
| 3165 | - | 3484730 | 3484359 | - | - | PA14_39100 | hypothetical protein | 93 | 116 | 127 | 111 | 120 | 115 | 1 |
| 3166 | - | 3485360 | 3485025 | - | - | PA14_39110 | hypothetical protein | 41 | 36 | 51 | 46 | 31 | 37 | 1 |
| 3167 | - | 3485688 | 3487253 | + | - | PA14_39130 | ABC transporter ATP-binding protein | 71 | 49 | 81 | 50 | 54 | 60 | 1 |
| 3168 | - | 3487606 | 3487340 | - | - | PA14_39140 | hypothetical protein | 163 | 142 | 148 | 163 | 153 | 188 | 1 |
| 3169 | - | 3488321 | 3487713 | - | - | PA14_39150 | azoreductase | 8 | 5 | 9 | 5 | 6 | 6 | 0.956402005 |
| 3170 | - | 3488469 | 3489404 | + | - | PA14_39160 | LysR family transcriptional regulator | 20 | 18 | 26 | 18 | 18 | 20 | 1 |
| 3171 | - | 3489849 | 3489415 | - | - | PA14_39180 | hypothetical protein | 20 | 22 | 22 | 21 | 21 | 18 | 1 |
| 3172 | - | 3491037 | 3490204 | - | bacA | PA14_39190 | UDP pyrophosphate phosphatase | 13 | 9 | 14 | 9 | 9 | 8 | 1 |
| 3173 | - | 3491319 | 3491894 | + | - | PA14_39200 | transporter | 38 | 35 | 48 | 47 | 32 | 28 | 1 |
| 3174 | - | 3491891 | 3492418 | + | - | PA14_39210 | hypothetical protein | 18 | 19 | 18 | 17 | 18 | 18 | 1 |
| 3175 | - | 3492721 | 3493212 | + | - | PA14_39220 | hypothetical protein | 2 | 1 | 3 | 3 | 2 | 3 | 1 |
| 3176 | - | 3493209 | 3493778 | + | - | PA14_39230 | hypothetical protein | 4 | 5 | 5 | 3 | 3 | 5 | 1 |
| 3177 | - | 3493835 | 3494857 | + | - | PA14_39240 | hypothetical protein | 5 | 5 | 6 | 6 | 5 | 6 | 1 |
| 3178 | - | 3494919 | 3495599 | + | - | PA14_39250 | double-glycine peptidase | 8 | 9 | 7 | 10 | 6 | 8 | 1 |
| 3179 | - | 3495613 | 3496365 | + | - | PA14_39260 | hypothetical protein | 4 | 4 | 5 | 4 | 3 | 3 | 1 |
| 3180 | - | 3496426 | 3497691 | + | - | PA14_39270 | hypothetical protein | 34 | 53 | 46 | 65 | 46 | 51 | 0.940967938 |
| 3181 | - | 3498751 | 3497825 | - | rbsK | PA14_39280 | ribokinase | 167 | 416 | 300 | 468 | 391 | 313 | 0.723542393 |
| 3182 | - | 3499818 | 3498805 | - | rbsR | PA14_39300 | ribose operon repressor RbsR | 155 | 374 | 287 | 387 | 308 | 278 | 0.80035376 |
| 3183 | - | 3500820 | 3499822 | - | rbsC | PA14_39320 | membrane protein component of ABC ribose transporter | 40 | 147 | 92 | 136 | 107 | 91 | 0.000387326 |
| 3184 | - | 3502376 | 3500844 | - | rbsA | PA14_39330 | ribose transporter | 71 | 263 | 189 | 264 | 219 | 179 | 0.016403065 |
| 3185 | - | 3503357 | 3502398 | - | rbsB | PA14_39350 | ribose ABC transporter substrate-binding protein | 104 | 512 | 343 | 533 | 479 | 349 | 1.45127E-05 |
| 3186 | - | 3504904 | 3503576 | - | - | PA14_39360 | sigma-54 dependent transcriptional regulator | 32 | 32 | 32 | 41 | 33 | 38 | 1 |
| 3187 | - | 3506527 | 3505040 | - | - | PA14_39390 | 30S ribosomal protein S6 modification protein | 42 | 40 | 41 | 54 | 42 | 51 | 1 |
| 3188 | - | 3506718 | 3507815 | + | - | PA14_39410 | hypothetical protein | 35 | 34 | 34 | 32 | 29 | 34 | 1 |
| 3189 | - | 3507877 | 3508107 | + | - | PA14_39420 | hypothetical protein | 16 | 8 | 8 | 10 | 6 | 16 | 0.401171332 |
| 3190 | - | 3508418 | 3510313 | + | - | PA14_39440 | hypothetical protein | 29 | 31 | 25 | 31 | 29 | 29 | 1 |
| 3191 | - | 3510492 | 3511631 | + | - | PA14_39460 | hypothetical protein | 47 | 41 | 44 | 53 | 45 | 48 | 1 |
| 3192 | - | 3512076 | 3512852 | + | - | PA14_39470 | hypothetical protein | 76 | 52 | 42 | 37 | 46 | 57 | 1 |

| | A | B | C | D | E | F | G | H | I | J | K | L | M | N |
|---|---|---|---|---|---|---|---|---|---|---|---|---|---|---|
| 3193 | - | 3513093 | 3515246 | + | - | *PA14_39480* | hypothetical protein | 115 | 65 | 54 | 76 | 88 | 114 | 1 |
| 3194 | - | 3516510 | 3516130 | - | - | *PA14_39500* | hypothetical protein | 21 | 23 | 21 | 20 | 20 | 20 | 1 |
| 3195 | - | 3519209 | 3517005 | - | - | *PA14_39520* | hydroxylase large subunit | 24 | 34 | 33 | 43 | 40 | 41 | 1 |
| 3196 | - | 3520195 | 3519206 | - | - | *PA14_39530* | hydroxylase molybdopterin-containing subunit | 27 | 28 | 28 | 32 | 26 | 23 | 1 |
| 3197 | - | 3520704 | 3520192 | - | - | *PA14_39540* | ferredoxin | 43 | 47 | 52 | 51 | 44 | 44 | 1 |
| 3198 | - | 3522169 | 3520874 | - | - | *PA14_39560* | chemotaxis transducer | 65 | 77 | 88 | 94 | 70 | 104 | 1 |
| 3199 | - | 3522478 | 3522774 | + | - | *PA14_39570* | hypothetical protein | 13 | 12 | 13 | 10 | 11 | 10 | 1 |
| 3200 | - | 3522821 | 3523390 | + | *rimJ* | *PA14_39580* | ribosomal protein alanine acetyltransferase | 12 | 14 | 11 | 10 | 12 | 14 | 1 |
| 3201 | - | 3525752 | 3523452 | - | *metE* | *PA14_39590* | 5-methyltetrahydropteroyltriglutamate/homocysteine S-methyltransferase | 22 | 8 | 10 | 11 | 8 | 31 | 0.049546026 |
| 3202 | - | 3527735 | 3525885 | - | - | *PA14_39610* | hypothetical protein | 34 | 38 | 47 | 36 | 34 | 29 | 1 |
| 3203 | - | 3528562 | 3528236 | - | - | *PA14_39620* | hypothetical protein | 3 | 4 | 3 | 3 | 4 | 3 | 1 |
| 3204 | - | 3529038 | 3528562 | - | - | *PA14_39630* | hypothetical protein | 1 | 1 | 1 | 1 | 1 | 1 | 1 |
| 3205 | - | 3532880 | 3529035 | - | *cobN* | *PA14_39640* | cobaltochelatase subunit CobN | 3 | 3 | 4 | 4 | 3 | 4 | 1 |
| 3206 | - | 3534841 | 3532880 | - | - | *PA14_39650* | TonB-dependent receptor | 2 | 2 | 2 | 3 | 3 | 2 | 1 |
| 3207 | - | 3535009 | 3535818 | + | - | *PA14_39660* | hypothetical protein | 7 | 7 | 5 | 5 | 6 | 5 | 1 |
| 3208 | - | 3535948 | 3535826 | - | - | *PA14_39670* | hypothetical protein | 2 | 1 | 2 | 1 | 3 | 3 | 0.239905719 |
| 3209 | - | 3535976 | 3538003 | + | - | *PA14_39690* | anaerobic ribonucleoside triphosphate reductase | 16 | 14 | 15 | 14 | 15 | 18 | 1 |
| 3210 | - | 3538047 | 3538193 | + | - | *PA14_39700* | hypothetical protein | 23 | 32 | 29 | 24 | 21 | 22 | 1 |
| 3211 | - | 3538190 | 3538888 | + | - | *PA14_39710* | radical SAM protein | 80 | 74 | 75 | 90 | 72 | 77 | 1 |
| 3212 | - | 3539082 | 3540482 | + | - | *PA14_39720* | amino acid oxidase | 5 | 5 | 5 | 6 | 5 | 6 | 1 |
| 3213 | - | 3540495 | 3540842 | + | - | *PA14_39730* | hypothetical protein | 4 | 3 | 4 | 4 | 5 | 5 | 1 |
| 3214 | - | 3540887 | 3542119 | + | - | *PA14_39750* | amino acid permease | 7 | 8 | 11 | 9 | 8 | 9 | 1 |
| 3215 | - | 3542183 | 3543730 | + | - | *PA14_39770* | regulatory protein | 50 | 48 | 55 | 59 | 50 | 54 | 1 |
| 3216 | - | 3544298 | 3545524 | + | - | *PA14_39780* | hypothetical protein | 30 | 10 | 12 | 17 | 19 | 86 | 0.003750459 |
| 3217 | - | 3545631 | 3546311 | + | - | *PA14_39790* | hypothetical protein | 46 | 37 | 48 | 44 | 41 | 33 | 1 |
| 3218 | - | 3546403 | 3546909 | + | - | *PA14_39800* | ECF subfamily RNA polymerase sigma-70 factor | 6 | 42 | 9 | 96 | 39 | 7 | 2.77209E-25 |
| 3219 | - | 3546906 | 3547856 | + | - | *PA14_39810* | transmembrane sensor | 8 | 32 | 12 | 90 | 32 | 9 | 9.20065E-07 |
| 3220 | - | 3548064 | 3550478 | + | - | *PA14_39820* | tonB-dependent receptor protein | 6 | 6 | 7 | 11 | 7 | 6 | 1 |
| 3221 | - | 3550519 | 3551670 | + | - | *PA14_39830* | hypothetical protein | 4 | 3 | 3 | 5 | 3 | 3 | 1 |
| 3222 | - | 3551857 | 3553068 | + | - | *PA14_39850* | MFS transporter | 9 | 2 | 4 | 4 | 3 | 6 | 1.56217E-07 |
| 3223 | - | 3553065 | 3554705 | + | - | *PA14_39860* | hypothetical protein | 89 | 5 | 6 | 14 | 10 | 66 | 0 |
| 3224 | - | 3554748 | 3555284 | + | - | *PA14_39870* | hydrolase | 226 | 10 | 15 | 31 | 28 | 167 | 0 |
| 3225 | 3561969 | 3556106 | 3555459 | - | *phzG2* | *PA14_39880* | pyridoxamine 5'-phosphate oxidase | 897 | 11 | 30 | 70 | 78 | 704 | 0 |
| 3226 | 3561969 | 3556965 | 3556129 | - | *phzF2* | *PA14_39890* | phenazine biosynthesis protein | 899 | 10 | 27 | 57 | 72 | 708 | 0 |
| 3227 | 3561969 | 3558862 | 3556979 | - | *phzE2* | *PA14_39910* | phenazine biosynthesis protein PhzE | 1059 | 14 | 37 | 79 | 84 | 814 | 0 |
| 3228 | 3561969 | 3559482 | 3558859 | - | *phzD2* | *PA14_39925* | phenazine biosynthesis protein PhzD | 888 | 6 | 19 | 67 | 76 | 691 | 0 |
| 3229 | 3561157 | 3560696 | 3559479 | - | *phzC2* | *PA14_39945* | phenazine biosynthesis protein PhzC | 1323 | 9 | 38 | 88 | 81 | 870 | 0 |
| 3230 | 3561157 | 3561208 | 3560720 | - | *phzB2* | *PA14_39960* | phenazine biosynthesis protein | 1000 | 9 | 24 | 119 | 78 | 797 | 0 |
| 3231 | 3561969 | 3561732 | 3561244 | - | *phzA2* | *PA14_39970* | phenazine biosynthesis protein | 176 | 1 | 5 | 20 | 9 | 134 | 0 |
| 3232 | - | 3562927 | 3562214 | - | *qscR* | *PA14_39980* | transcriptional regulator | 93 | 9 | 17 | 45 | 28 | 90 | 3.18216E-87 |
| 3233 | - | 3563689 | 3564456 | + | - | *PA14_39990* | hypothetical protein | 20 | 20 | 18 | 23 | 22 | 24 | 1 |
| 3234 | - | 3564462 | 3565631 | + | - | *PA14_40010* | hypothetical protein | 10 | 12 | 10 | 9 | 8 | 9 | 1 |
| 3235 | - | 3565650 | 3566924 | + | - | *PA14_40020* | hypothetical protein | 12 | 14 | 12 | 14 | 12 | 14 | 1 |
| 3236 | - | 3566872 | 3567564 | + | - | *PA14_40030* | hypothetical protein | 8 | 9 | 6 | 6 | 7 | 5 | 1 |
| 3237 | - | 3567590 | 3570019 | + | - | *PA14_40040* | hypothetical protein | 11 | 12 | 13 | 15 | 11 | 12 | 1 |
| 3238 | - | 3570016 | 3570753 | + | - | *PA14_40050* | hypothetical protein | 6 | 5 | 6 | 5 | 6 | 4 | 1 |
| 3239 | - | 3570750 | 3571139 | + | - | *PA14_40060* | hypothetical protein | 10 | 7 | 12 | 12 | 9 | 8 | 1 |
| 3240 | - | 3571228 | 3571851 | + | - | *PA14_40070* | glutathione S-transferase | 62 | 64 | 63 | 57 | 58 | 56 | 1 |
| 3241 | - | 3572848 | 3571865 | - | - | *PA14_40080* | hypothetical protein | 48 | 43 | 49 | 45 | 45 | 48 | 1 |
| 3242 | - | 3573078 | 3574487 | + | - | *PA14_40100* | hypothetical protein | 60 | 35 | 71 | 79 | 35 | 69 | 1 |
| 3243 | - | 3574471 | 3575139 | + | - | *PA14_40110* | hypothetical protein | 79 | 66 | 103 | 111 | 64 | 115 | 1 |
| 3244 | - | 3577506 | 3575143 | - | *polB* | *PA14_40120* | DNA polymerase II | 60 | 59 | 70 | 67 | 58 | 72 | 1 |
| 3245 | - | 3578108 | 3577575 | - | - | *PA14_40130* | hypothetical protein | 69 | 59 | 59 | 71 | 63 | 59 | 1 |
| 3246 | - | 3578194 | 3578745 | + | - | *PA14_40150* | transcriptional regulator | 27 | 22 | 23 | 22 | 22 | 18 | 1 |
| 3247 | - | 3579140 | 3578742 | - | - | *PA14_40160* | NADH-ubiquinone/plastoquinone oxidoreductase | 16 | 15 | 17 | 14 | 12 | 14 | 1 |
| 3248 | - | 3579476 | 3579153 | - | - | *PA14_40170* | transporter | 8 | 4 | 6 | 4 | 3 | 4 | 0.381163237 |
| 3249 | - | 3579807 | 3580268 | + | - | *PA14_40180* | oxidoreductase | 41 | 44 | 56 | 48 | 48 | 51 | 1 |
| 3250 | - | 3580273 | 3582468 | + | - | *PA14_40200* | oxidoreductase | 80 | 81 | 89 | 98 | 82 | 105 | 1 |
| 3251 | - | 3582626 | 3583180 | + | - | *PA14_40210* | hypothetical protein | 46 | 52 | 61 | 65 | 48 | 65 | 1 |
| 3252 | - | 3583268 | 3583846 | + | - | *PA14_40220* | hypothetical protein | 41 | 44 | 43 | 55 | 56 | 96 | 1 |
| 3253 | - | 3585142 | 3583955 | - | - | *PA14_40230* | secretion protein | 36 | 42 | 28 | 61 | 62 | 149 | 1 |
| 3254 | - | 3587303 | 3585132 | - | - | *PA14_40240* | ABC transporter ATP-binding protein/permease | 35 | 32 | 23 | 48 | 52 | 124 | 1 |
| 3255 | - | 3588570 | 3587293 | - | - | *PA14_40250* | outer membrane protein | 51 | 45 | 37 | 70 | 66 | 154 | 1 |
| 3256 | - | 3595940 | 3588570 | - | - | *PA14_40260* | hypothetical protein | 104 | 77 | 63 | 116 | 109 | 258 | 1 |
| 3257 | - | 3597214 | 3598236 | + | - | *PA14_40270* | cation transporter | 17 | 12 | 11 | 13 | 12 | 18 | 1 |
| 3258 | - | 3598297 | 3599088 | + | - | *PA14_40280* | hypothetical protein | 103 | 80 | 74 | 120 | 103 | 142 | 1 |
| 3259 | 3600666 | 3600471 | 3599215 | - | *lasA* | *PA14_40290* | LasA protease | 258 | 112 | 104 | 294 | 187 | 436 | 1 |
| 3260 | - | 3601180 | 3600755 | - | - | *PA14_40300* | hypothetical protein | 70 | 78 | 95 | 94 | 91 | 108 | 1 |
| 3261 | - | 3601703 | 3601464 | - | - | *PA14_40310* | acyl carrier protein | 377 | 25 | 208 | 423 | 232 | 410 | 2.9515E-257 |
| 3262 | - | 3604201 | 3601871 | - | *xqhA* | *PA14_40320* | secretion protein XqhA | 5 | 4 | 6 | 4 | 5 | 6 | 1 |
| 3263 | - | 3604715 | 3604188 | - | - | *PA14_40330* | hypothetical protein | 2 | 2 | 3 | 3 | 1 | 2 | 1 |
| 3264 | - | 3605066 | 3604857 | - | - | *PA14_40340* | hypothetical protein | 12 | 9 | 9 | 6 | 5 | 10 | 1 |
| 3265 | - | 3607432 | 3605156 | - | - | *PA14_40350* | hypothetical protein | 30 | 25 | 29 | 31 | 27 | 27 | 1 |
| 3266 | - | 3609101 | 3607425 | - | - | *PA14_40370* | hypothetical protein | 34 | 34 | 31 | 36 | 32 | 35 | 1 |
| 3267 | - | 3609235 | 3609885 | + | - | *PA14_40380* | TetR family transcriptional regulator | 14 | 10 | 11 | 10 | 9 | 10 | 1 |
| 3268 | - | 3610001 | 3610756 | + | *modA* | *PA14_40390* | molybdate-binding periplasmic protein precursor modA | 11 | 16 | 16 | 11 | 12 | 12 | 1 |
| 3269 | - | 3610769 | 3611455 | + | *modB* | *PA14_40410* | molybdenum transport protein ModB | 26 | 24 | 29 | 26 | 30 | 24 | 1 |
| 3270 | - | 3611457 | 3612542 | + | *modC* | *PA14_40420* | molybdenum transport protein ModC | 53 | 57 | 60 | 75 | 67 | 74 | 1 |
| 3271 | - | 3613391 | 3612564 | - | - | *PA14_40430* | hypothetical protein | 75 | 79 | 90 | 105 | 79 | 106 | 1 |
| 3272 | - | 3613477 | 3614364 | + | - | *PA14_40440* | LysR family transcriptional regulator | 11 | 13 | 14 | 12 | 14 | 11 | 1 |
| 3273 | - | 3614373 | 3615194 | + | *str* | *PA14_40450* | streptomycin 3''-phosphotransferase | 52 | 51 | 62 | 64 | 49 | 53 | 1 |
| 3274 | - | 3616649 | 3615195 | - | - | *PA14_40470* | TrkH family potassium uptake protein | 20 | 21 | 21 | 20 | 19 | 19 | 1 |
| 3275 | - | 3616827 | 3617750 | + | - | *PA14_40490* | hypothetical protein | 31 | 29 | 29 | 29 | 25 | 28 | 1 |
| 3276 | - | 3618097 | 3619539 | + | *ccoN-2* | *PA14_40510* | cbb3-type cytochrome c oxidase subunit I | 11 | 14 | 12 | 14 | 12 | 13 | 1 |

| | A | B | C | D | E | F | G | H | I | J | K | L | M | N |
|---|---|---|---|---|---|---|---|---|---|---|---|---|---|---|
| 3277 | - | 3619542 | 3619763 | + | - | *PA14_40520* | hypothetical protein | 6 | 6 | 8 | 8 | 6 | 5 | 1 |
| 3278 | - | 3619855 | 3621012 | + | - | *PA14_40540* | hypothetical protein | 19 | 16 | 21 | 22 | 13 | 17 | 1 |
| 3279 | - | 3621863 | 3621000 | - | - | *PA14_40550* | LysR family transcriptional regulator | 47 | 34 | 48 | 36 | 27 | 25 | 1 |
| 3280 | - | 3622148 | 3622423 | + | - | *PA14_40560* | hypothetical protein | 203 | 183 | 198 | 138 | 106 | 115 | 1 |
| 3281 | - | 3623677 | 3622472 | - | - | *PA14_40570* | two-component response regulator | 35 | 34 | 37 | 39 | 34 | 37 | 1 |
| 3282 | - | 3623926 | 3624930 | + | - | *PA14_40600* | transcriptional regulator | 18 | 18 | 17 | 22 | 16 | 17 | 1 |
| 3283 | - | 3624953 | 3625195 | + | - | *PA14_40610* | hypothetical protein | 3 | 4 | 3 | 4 | 5 | 5 | 1 |
| 3284 | - | 3625200 | 3626369 | + | - | *PA14_40620* | MFS transporter | 11 | 9 | 9 | 8 | 7 | 9 | 1 |
| 3285 | - | 3627012 | 3626428 | - | - | *PA14_40630* | hypothetical protein | 181 | 253 | 136 | 261 | 437 | 241 | 1 |
| 3286 | - | 3629423 | 3627135 | - | *cti* | *PA14_40640* | cis/trans isomerase | 41 | 37 | 41 | 37 | 35 | 33 | 1 |
| 3287 | - | 3629785 | 3630303 | + | - | *PA14_40650* | hypothetical protein | 4 | 3 | 5 | 4 | 6 | 3 | 1 |
| 3288 | - | 3630345 | 3630809 | + | - | *PA14_40660* | hypothetical protein | 9 | 7 | 12 | 9 | 7 | 9 | 1 |
| 3289 | - | 3631084 | 3634788 | + | *metH* | *PA14_40670* | B12-dependent methionine synthase | 104 | 68 | 80 | 69 | 57 | 97 | 1 |
| 3290 | - | 3634803 | 3635165 | + | - | *PA14_40690* | hypothetical protein | 89 | 74 | 67 | 50 | 54 | 90 | 1 |
| 3291 | - | 3635714 | 3635217 | - | - | *PA14_40700* | hypothetical protein | 67 | 50 | 53 | 46 | 44 | 57 | 1 |
| 3292 | - | 3636090 | 3635728 | - | - | *PA14_40710* | hypothetical protein | 64 | 59 | 68 | 69 | 47 | 61 | 1 |
| 3293 | - | 3636128 | 3637168 | + | - | *PA14_40730* | ribosomal RNA large subunit methyltransferase N | 27 | 27 | 34 | 34 | 28 | 30 | 1 |
| 3294 | - | 3637233 | 3637460 | + | - | *PA14_40740* | hypothetical protein | 254 | 349 | 322 | 366 | 294 | 335 | 1 |
| 3295 | - | 3637640 | 3637485 | - | - | *PA14_40750* | hypothetical protein | 211 | 322 | 291 | 331 | 276 | 326 | 1 |
| 3296 | - | 3637871 | 3639529 | + | *cysI* | *PA14_40770* | sulfite reductase | 28 | 41 | 23 | 21 | 36 | 21 | 1 |
| 3297 | - | 3639513 | 3640010 | + | - | *PA14_40780* | hypothetical protein | 46 | 53 | 31 | 28 | 41 | 26 | 1 |
| 3298 | - | 3640656 | 3640075 | - | - | *PA14_40790* | transcriptional regulator | 26 | 23 | 31 | 30 | 19 | 31 | 1 |
| 3299 | - | 3640779 | 3641216 | + | - | *PA14_40800* | hypothetical protein | 27 | 23 | 21 | 18 | 16 | 25 | 1 |
| 3300 | - | 3641311 | 3642186 | + | - | *PA14_40820* | hydrolase | 89 | 82 | 89 | 70 | 66 | 73 | 1 |
| 3301 | - | 3643165 | 3642173 | - | - | *PA14_40830* | oxidoreductase | 495 | 468 | 476 | 411 | 442 | 438 | 1 |
| 3302 | - | 3644249 | 3643224 | - | - | *PA14_40840* | periplasmic protease | 169 | 147 | 234 | 235 | 150 | 179 | 1 |
| 3303 | - | 3644484 | 3645194 | + | - | *PA14_40850* | hypothetical protein | 36 | 28 | 34 | 27 | 29 | 32 | 1 |
| 3304 | - | 3645255 | 3645569 | + | - | *PA14_40860* | hypothetical protein | 176 | 243 | 142 | 79 | 126 | 131 | 1 |
| 3305 | - | 3645827 | 3646897 | + | - | *PA14_40880* | hypothetical protein | 47 | 48 | 53 | 53 | 58 | 55 | 1 |
| 3306 | - | 3646922 | 3647689 | + | - | *PA14_40890* | short chain dehydrogenase | 65 | 82 | 49 | 52 | 69 | 63 | 1 |
| 3307 | - | 3648611 | 3647850 | - | - | *PA14_40900* | short-chain dehydrogenase | 7 | 5 | 5 | 8 | 5 | 8 | 1 |
| 3308 | - | 3648743 | 3649648 | + | - | *PA14_40910* | LysR family transcriptional regulatory protein | 57 | 56 | 69 | 70 | 54 | 60 | 1 |
| 3309 | - | 3650289 | 3649645 | - | - | *PA14_40930* | hypothetical protein | 64 | 67 | 77 | 87 | 65 | 64 | 1 |
| 3310 | - | 3650390 | 3651169 | + | - | *PA14_40940* | hypothetical protein | 45 | 48 | 55 | 51 | 37 | 37 | 1 |
| 3311 | - | 3651973 | 3651137 | - | *nudC* | *PA14_40950* | NADH pyrophosphatase | 229 | 239 | 236 | 258 | 219 | 217 | 1 |
| 3312 | - | 3653661 | 3651973 | - | - | *PA14_40960* | pilin biosynthetic protein | 243 | 229 | 230 | 258 | 188 | 204 | 1 |
| 3313 | - | 3654513 | 3653701 | - | - | *PA14_40980* | enoyl-CoA hydratase | 132 | 99 | 124 | 123 | 94 | 109 | 1 |
| 3314 | - | 3654737 | 3656239 | + | *nhaB* | *PA14_41000* | sodium/proton antiporter | 81 | 94 | 95 | 109 | 89 | 100 | 1 |
| 3315 | - | 3657578 | 3656223 | - | - | *PA14_41010* | amino acid permease | 226 | 207 | 243 | 295 | 236 | 242 | 1 |
| 3316 | - | 3659857 | 3657602 | - | - | *PA14_41020* | Orn/Arg/Lys decarboxylase | 146 | 167 | 142 | 190 | 154 | 145 | 1 |
| 3317 | - | 3660051 | 3660440 | + | - | *PA14_41030* | hypothetical protein | 118 | 99 | 113 | 119 | 96 | 109 | 1 |
| 3318 | - | 3661208 | 3660468 | - | *dnaQ* | *PA14_41050* | DNA polymerase III subunit epsilon | 89 | 81 | 79 | 88 | 70 | 75 | 1 |
| 3319 | - | 3661719 | 3661273 | - | *rnhA* | *PA14_41060* | ribonuclease H | 81 | 62 | 91 | 84 | 82 | 79 | 1 |
| 3320 | - | 3662483 | 3661722 | - | - | *PA14_41070* | hypothetical protein | 111 | 91 | 114 | 99 | 95 | 97 | 1 |
| 3321 | - | 3662879 | 3663361 | + | - | *PA14_41080* | hydroxyacylglutathione hydrolase | 30 | 34 | 41 | 31 | 31 | 30 | 1 |
| 3322 | - | 3663439 | 3665043 | + | *mltD* | *PA14_41090* | membrane-bound lytic murein transglycosylase D | 69 | 66 | 79 | 76 | 60 | 63 | 1 |
| 3323 | - | 3665234 | 3667063 | + | - | *PA14_41110* | solute-binding protein | 39 | 38 | 37 | 39 | 46 | 44 | 1 |
| 3324 | - | 3667060 | 3668907 | + | - | *PA14_41130* | ABC transporter substrate-binding protein | 118 | 127 | 105 | 109 | 118 | 118 | 1 |
| 3325 | - | 3668914 | 3669996 | + | - | *PA14_41140* | ABC transporter permease | 98 | 90 | 83 | 88 | 96 | 103 | 1 |
| 3326 | - | 3670001 | 3671020 | + | - | *PA14_41150* | ABC transporter permease | 60 | 59 | 57 | 47 | 61 | 61 | 1 |
| 3327 | - | 3671022 | 3672632 | + | - | *PA14_41160* | ABC transporter ATP-binding protein | 80 | 81 | 70 | 75 | 88 | 88 | 1 |
| 3328 | - | 3672654 | 3673451 | + | *fabI* | *PA14_41170* | NADH-dependent enoyl-ACP reductase | 107 | 121 | 77 | 81 | 102 | 90 | 1 |
| 3329 | - | 3675414 | 3673549 | - | *ppiD* | *PA14_41190* | peptidyl-prolyl cis-trans isomerase D | 104 | 125 | 96 | 86 | 102 | 80 | 1 |
| 3330 | - | 3675594 | 3675521 | - | - | *PA14_41200* | Asp tRNA | 341 | 418 | 254 | 245 | 242 | 189 | 1 |
| 3331 | - | 3675918 | 3675646 | - | *hupB* | *PA14_41210* | DNA-binding protein HU | 1422 | 2211 | 1041 | 955 | 1061 | 846 | 1 |
| 3332 | - | 3675919 | 3676041 | ? | - | predicted RNA | - | 994 | 1295 | 639 | 607 | 777 | 561 | 1 |
| 3333 | - | 3678450 | 3676054 | - | *lon* | *PA14_41220* | Lon protease | 486 | 511 | 505 | 392 | 520 | 491 | 1 |
| 3334 | - | 3679862 | 3678582 | - | *clpX* | *PA14_41230* | ATP-dependent protease ATP-binding subunit ClpX | 926 | 964 | 994 | 832 | 857 | 857 | 1 |
| 3335 | - | 3680608 | 3679967 | - | *clpP* | *PA14_41240* | ATP-dependent Clp protease proteolytic subunit | 396 | 351 | 426 | 309 | 294 | 324 | 1 |
| 3336 | - | 3682012 | 3680702 | - | *tig* | *PA14_41250* | trigger factor | 222 | 270 | 257 | 210 | 196 | 137 | 1 |
| 3337 | - | 3682244 | 3682951 | + | - | *PA14_41260* | two-component response regulator | 10 | 10 | 11 | 12 | 9 | 11 | 1 |
| 3338 | - | 3682952 | 3684238 | + | - | *PA14_41270* | two-component sensor | 12 | 11 | 11 | 12 | 10 | 11 | 1 |
| 3339 | - | 3684343 | 3686175 | + | - | *PA14_41280* | beta-lactamase | 11 | 12 | 14 | 14 | 10 | 11 | 1 |
| 3340 | - | 3686172 | 3686288 | + | - | *PA14_41290* | hypothetical protein | 31 | 35 | 40 | 32 | 35 | 34 | 1 |
| 3341 | - | 3686683 | 3686462 | - | - | *PA14_41300* | hypothetical protein | 51 | 60 | 90 | 68 | 76 | 58 | 1 |
| 3342 | - | 3687098 | 3687017 | - | - | *PA14_41320* | Leu tRNA | 72 | 60 | 95 | 60 | 37 | 40 | 1 |
| 3343 | - | 3687228 | 3687156 | - | - | *PA14_41330* | His tRNA | 106 | 78 | 112 | 84 | 48 | 53 | 1 |
| 3344 | - | 3687356 | 3687283 | - | - | *PA14_41340* | Arg tRNA | 552 | 331 | 535 | 574 | 312 | 316 | 1 |
| 3345 | - | 3687635 | 3688489 | + | *folD* | *PA14_41350* | bifunctional 5,10-methylene-tetrahydrofolate dehydrogenase/ 5,10-methylene-tetrahydrofolate cyclohydrolase | 37 | 37 | 45 | 40 | 42 | 33 | 1 |
| 3346 | - | 3689929 | 3688547 | - | *cysS* | *PA14_41360* | cysteinyl-tRNA synthetase | 244 | 250 | 255 | 250 | 230 | 221 | 1 |
| 3347 | - | 3691609 | 3689939 | - | *glnS* | *PA14_41380* | glutaminyl-tRNA synthetase | 158 | 168 | 169 | 168 | 148 | 140 | 1 |
| 3348 | - | 3691732 | 3692229 | + | *ppiB* | *PA14_41390* | peptidyl-prolyl cis-trans isomerase B | 180 | 224 | 153 | 110 | 144 | 128 | 1 |
| 3349 | - | 3692234 | 3692956 | + | - | *PA14_41400* | UDP-2,3-diacylglucosamine hydrolase | 57 | 61 | 53 | 54 | 79 | 70 | 1 |
| 3350 | - | 3693088 | 3694419 | + | - | *PA14_41420* | hypothetical protein | 11 | 10 | 11 | 9 | 9 | 7 | 1 |
| 3351 | - | 3695091 | 3694468 | - | - | *PA14_41430* | hypothetical protein | 15 | 12 | 14 | 11 | 7 | 7 | 1 |
| 3352 | - | 3695262 | 3696125 | + | - | *PA14_41440* | hypothetical protein | 436 | 371 | 562 | 213 | 201 | 386 | 1 |
| 3353 | - | 3696648 | 3696181 | - | - | *PA14_41450* | hypothetical protein | 84 | 92 | 134 | 78 | 48 | 73 | 1 |
| 3354 | - | 3697030 | 3699639 | + | *acnB* | *PA14_41470* | bifunctional aconitate hydratase 2/2-methylisocitrate dehydratase | 247 | 298 | 175 | 143 | 204 | 131 | 1 |
| 3355 | - | 3699817 | 3701025 | + | *nasS* | *PA14_41480* | hypothetical protein | 6 | 6 | 6 | 7 | 7 | 7 | 1 |
| 3356 | - | 3701042 | 3701620 | + | *nasT* | *PA14_41490* | hypothetical protein | 25 | 22 | 27 | 25 | 21 | 26 | 1 |
| 3357 | - | 3701850 | 3702545 | + | - | *PA14_41500* | lyase | 42 | 50 | 57 | 75 | 45 | 89 | 1 |
| 3358 | - | 3702898 | 3704109 | + | *nasA* | *PA14_41510* | nitrate transporter | 3 | 2 | 2 | 3 | 4 | 4 | 1 |
| 3359 | - | 3704178 | 3705779 | + | - | *PA14_41520* | serine/threonine-protein kinase | 13 | 16 | 17 | 19 | 14 | 16 | 1 |
| 3360 | - | 3706047 | 3708497 | + | *nirB* | *PA14_41530* | assimilatory nitrite reductase large subunit | 4 | 4 | 4 | 5 | 3 | 4 | 1 |

| | A | B | C | D | E | F | G | H | I | J | K | L | M | N |
|---|---|---|---|---|---|---|---|---|---|---|---|---|---|---|
| 3361 | - | 3708550 | 3708876 | + | *nirD* | *PA14_41540* | assimilatory nitrite reductase small subunit | 2 | 2 | 2 | 2 | 1 | 1 | 1 |
| 3362 | - | 3708903 | 3711629 | + | - | *PA14_41560* | assimilatory nitrate reductase | 5 | 6 | 6 | 6 | 5 | 5 | 1 |
| 3363 | - | 3711641 | 3712378 | + | *cobA* | *PA14_41563* | uroporphyrin-III C-methyltransferase | 11 | 13 | 14 | 11 | 12 | 9 | 1 |
| 3364 | - | 3713499 | 3712447 | - | *oprF* | *PA14_41570* | major porin and structural outer membrane porin OprF precursor | 3860 | 4586 | 2953 | 2511 | 3447 | 2916 | 1 |
| 3365 | - | 3714198 | 3713608 | - | *sigX* | *PA14_41575* | RNA polymerase sigma factor SigX | 706 | 773 | 867 | 793 | 766 | 743 | 1 |
| 3366 | - | 3715094 | 3714270 | - | - | *PA14_41590* | cytoplasmic membrane protein | 171 | 183 | 214 | 197 | 158 | 149 | 1 |
| 3367 | - | 3715351 | 3715097 | - | - | *PA14_41610* | hypothetical protein | 102 | 90 | 116 | 129 | 113 | 112 | 1 |
| 3368 | - | 3716398 | 3715400 | - | - | *PA14_41630* | cytoplasmic membrane-associated protein | 89 | 82 | 83 | 94 | 76 | 81 | 1 |
| 3369 | - | 3717074 | 3716586 | - | - | *PA14_41640* | ribonuclease activity regulator protein RraA | 261 | 276 | 265 | 241 | 231 | 220 | 1 |
| 3370 | - | 3718191 | 3717181 | - | - | *PA14_41650* | esterase | 15 | 16 | 18 | 10 | 14 | 11 | 1 |
| 3371 | - | 3720636 | 3718261 | - | *ppsA* | *PA14_41670* | phosphoenolpyruvate synthase | 252 | 302 | 252 | 271 | 249 | 217 | 1 |
| 3372 | - | 3720802 | 3721626 | + | - | *PA14_41680* | hypothetical protein | 82 | 79 | 105 | 98 | 90 | 93 | 1 |
| 3373 | - | 3721917 | 3722456 | + | - | *PA14_41690* | hypothetical protein | 60 | 43 | 63 | 52 | 41 | 48 | 1 |
| 3374 | - | 3722462 | 3723988 | + | - | *PA14_41710* | hypothetical protein | 114 | 122 | 114 | 114 | 113 | 104 | 1 |
| 3375 | - | 3723992 | 3724975 | + | - | *PA14_41730* | hypothetical protein | 110 | 118 | 109 | 98 | 91 | 85 | 1 |
| 3376 | - | 3726219 | 3725032 | - | - | *PA14_41740* | hypothetical protein | 16 | 13 | 17 | 20 | 13 | 19 | 1 |
| 3377 | - | 3727831 | 3726233 | - | - | *PA14_41750* | hypothetical protein | 7 | 8 | 11 | 11 | 7 | 9 | 1 |
| 3378 | - | 3728876 | 3727851 | - | - | *PA14_41760* | hypothetical protein | 10 | 13 | 13 | 15 | 11 | 15 | 1 |
| 3379 | - | 3729678 | 3728917 | - | - | *PA14_41780* | hypothetical protein | 44 | 44 | 44 | 46 | 42 | 59 | 1 |
| 3380 | - | 3730194 | 3729757 | - | - | *PA14_41790* | hypothetical protein | 79 | 90 | 78 | 82 | 67 | 103 | 1 |
| 3381 | - | 3733149 | 3730426 | - | - | *PA14_41800* | transcriptional regulator | 115 | 120 | 115 | 122 | 112 | 128 | 1 |
| 3382 | - | 3735857 | 3733152 | - | - | *PA14_41810* | transcriptional regulator | 159 | 152 | 146 | 157 | 149 | 167 | 1 |
| 3383 | - | 3736022 | 3737383 | + | *pabB* | *PA14_41820* | para-aminobenzoate synthase component I | 47 | 51 | 54 | 56 | 47 | 42 | 1 |
| 3384 | - | 3738043 | 3737426 | - | *thrH* | *PA14_41830* | phosphoserine phosphatase | 24 | 28 | 27 | 28 | 28 | 26 | 1 |
| 3385 | - | 3738178 | 3738981 | + | *cysH* | *PA14_41840* | phosphoadenosine phosphosulfate reductase | 29 | 32 | 36 | 26 | 30 | 28 | 1 |
| 3386 | - | 3739046 | 3739393 | + | - | *PA14_41860* | hypothetical protein | 20 | 26 | 24 | 21 | 21 | 23 | 1 |
| 3387 | - | 3740424 | 3739450 | - | *cysB* | *PA14_41870* | transcriptional regulator CysB | 186 | 213 | 178 | 165 | 191 | 204 | 1 |
| 3388 | - | 3741053 | 3740556 | - | - | *PA14_41880* | universal stress protein | 300 | 287 | 360 | 370 | 430 | 425 | 1 |
| 3389 | - | 3742163 | 3741216 | - | *panE* | *PA14_41900* | 2-dehydropantoate 2-reductase | 129 | 119 | 132 | 142 | 113 | 112 | 1 |
| 3390 | - | 3742540 | 3742163 | - | - | *PA14_41910* | hypothetical protein | 111 | 86 | 98 | 120 | 82 | 87 | 1 |
| 3391 | - | 3742858 | 3743934 | + | *aroF-1* | *PA14_41920* | phospho-2-dehydro-3-deoxyheptonate aldolase | 74 | 67 | 72 | 66 | 62 | 53 | 1 |
| 3392 | - | 3744477 | 3743992 | - | - | *PA14_41930* | hypothetical protein | 202 | 139 | 185 | 205 | 188 | 236 | 1 |
| 3393 | - | 3744661 | 3745350 | + | - | *PA14_41950* | enoyl-CoA hydratase | 101 | 86 | 124 | 56 | 51 | 82 | 1 |
| 3394 | - | 3745500 | 3745688 | + | - | *PA14_41960* | hypothetical protein | 147 | 135 | 189 | 52 | 63 | 128 | 1 |
| 3395 | - | 3746266 | 3745760 | - | - | *PA14_41970* | hypothetical protein | 1016 | 835 | 1191 | 507 | 484 | 644 | 1 |
| 3396 | - | 3746487 | 3746975 | + | - | *PA14_41980* | hypothetical protein | 22 | 24 | 26 | 19 | 22 | 24 | 1 |
| 3397 | - | 3747130 | 3747348 | + | - | *PA14_41990* | hypothetical protein | 8 | 9 | 9 | 7 | 6 | 6 | 1 |
| 3398 | - | 3747376 | 3747615 | + | - | *PA14_42000* | hypothetical protein | 11 | 13 | 13 | 11 | 10 | 13 | 1 |
| 3399 | - | 3748395 | 3747673 | - | - | *PA14_42010* | amidotransferase | 236 | 186 | 210 | 164 | 169 | 201 | 1 |
| 3400 | - | 3748639 | 3749010 | + | - | *PA14_42020* | hypothetical protein | 44 | 51 | 48 | 60 | 39 | 50 | 1 |
| 3401 | - | 3750017 | 3749019 | - | - | *PA14_42030* | hypothetical protein | 12 | 17 | 18 | 19 | 15 | 15 | 1 |
| 3402 | - | 3751011 | 3750040 | - | - | *PA14_42050* | oxidoreductase | 3 | 2 | 1 | 2 | 2 | 2 | 1 |
| 3403 | - | 3751162 | 3752076 | + | - | *PA14_42060* | transcriptional regulator | 11 | 12 | 11 | 10 | 10 | 12 | 1 |
| 3404 | - | 3754605 | 3752461 | - | - | *PA14_42080* | 3-hydroxyacyl-CoA dehydrogenase | 38 | 36 | 37 | 35 | 37 | 40 | 1 |
| 3405 | - | 3755824 | 3754619 | - | - | *PA14_42090* | acetyl-CoA acetyltransferase | 31 | 30 | 32 | 27 | 33 | 30 | 1 |
| 3406 | - | 3757282 | 3756395 | - | - | *PA14_42100* | hypothetical protein | 30 | 24 | 28 | 34 | 25 | 28 | 1 |
| 3407 | - | 3757395 | 3758216 | + | - | *PA14_42120* | hypothetical protein | 33 | 38 | 46 | 49 | 44 | 46 | 1 |
| 3408 | - | 3758955 | 3758224 | - | - | *PA14_42130* | hypothetical protein | 87 | 88 | 102 | 106 | 93 | 101 | 1 |
| 3409 | - | 3759837 | 3759037 | - | - | *PA14_42140* | transglutaminase | 77 | 95 | 107 | 112 | 87 | 100 | 1 |
| 3410 | - | 3760787 | 3759834 | - | - | *PA14_42150* | hypothetical protein | 39 | 46 | 57 | 63 | 49 | 57 | 1 |
| 3411 | - | 3762202 | 3760790 | - | - | *PA14_42160* | hypothetical protein | 49 | 59 | 85 | 79 | 61 | 64 | 1 |
| 3412 | - | 3763238 | 3762552 | - | - | *PA14_42180* | hypothetical protein | 154 | 142 | 137 | 112 | 122 | 131 | 1 |
| 3413 | - | 3763689 | 3763348 | - | - | *PA14_42200* | hypothetical protein | 302 | 307 | 346 | 359 | 328 | 398 | 1 |
| 3414 | - | 3764042 | 3766099 | + | - | *PA14_42220* | membrane sensor domain-containing protein | 42 | 35 | 52 | 39 | 30 | 37 | 1 |
| 3415 | - | 3766270 | 3768564 | + | *bglX* | *PA14_42230* | beta-glucosidase | 42 | 39 | 48 | 46 | 41 | 37 | 1 |
| 3416 | - | 3769311 | 3768667 | - | *pscL* | *PA14_42250* | type III secretion system protein | 55 | 44 | 59 | 47 | 50 | 44 | 1 |
| 3417 | - | 3769910 | 3769290 | - | *pscK* | *PA14_42260* | PscK type III export protein | 40 | 30 | 39 | 27 | 30 | 27 | 1 |
| 3418 | - | 3770665 | 3769919 | - | *pscJ* | *PA14_42270* | pscJ type III export protein | 86 | 66 | 91 | 70 | 84 | 79 | 1 |
| 3419 | - | 3771000 | 3770662 | - | *pscI* | *PA14_42280* | type III export protein PscI | 82 | 68 | 83 | 73 | 73 | 62 | 1 |
| 3420 | - | 3771431 | 3771000 | - | *pscH* | *PA14_42290* | type III export protein PscH | 100 | 77 | 113 | 71 | 87 | 74 | 1 |
| 3421 | - | 3771775 | 3771428 | - | *pscG* | *PA14_42300* | type III export protein PscG | 197 | 150 | 209 | 157 | 163 | 170 | 1 |
| 3422 | - | 3772035 | 3771778 | - | *pscF* | *PA14_42310* | type III export protein PscF | 156 | 157 | 168 | 150 | 155 | 153 | 1 |
| 3423 | - | 3772241 | 3772038 | - | *pscE* | *PA14_42320* | type III export protein PscE | 155 | 149 | 152 | 173 | 174 | 161 | 1 |
| 3424 | - | 3773502 | 3772204 | - | *pscD* | *PA14_42340* | type III export protein PscD | 123 | 89 | 142 | 115 | 109 | 102 | 1 |
| 3425 | - | 3775306 | 3773504 | - | *pscC* | *PA14_42350* | Type III secretion outer membrane protein PscC precursor | 79 | 60 | 87 | 74 | 65 | 64 | 1 |
| 3426 | - | 3775728 | 3775306 | - | *pscB* | *PA14_42360* | type III export apparatus protein | 129 | 87 | 139 | 119 | 87 | 95 | 1 |
| 3427 | - | 3776592 | 3775762 | - | - | *PA14_42380* | hypothetical protein | 246 | 193 | 283 | 286 | 206 | 226 | 1 |
| 3428 | - | 3777526 | 3776690 | - | *exsA* | *PA14_42390* | transcriptional regulator ExsA | 103 | 80 | 113 | 102 | 67 | 73 | 1 |
| 3429 | - | 3778237 | 3777824 | - | *exsB* | *PA14_42400* | exoenzyme S synthesis protein B | 323 | 245 | 430 | 407 | 288 | 315 | 1 |
| 3430 | - | 3778491 | 3778246 | - | - | *PA14_42410* | hypothetical protein | 1093 | 937 | 1555 | 1483 | 1045 | 1114 | 1 |
| 3431 | - | 3778937 | 3778500 | - | *exsC* | *PA14_42430* | exoenzyme S synthesis protein C | 1167 | 1083 | 1405 | 1668 | 1082 | 1189 | 1 |
| 3432 | - | 3779951 | 3779064 | - | *popD* | *PA14_42440* | translocator outer membrane protein PopD precursor | 239 | 239 | 275 | 255 | 258 | 255 | 1 |
| 3433 | - | 3781135 | 3779963 | - | *popB* | *PA14_42450* | translocator protein PopB | 323 | 300 | 372 | 356 | 354 | 335 | 1 |
| 3434 | - | 3781622 | 3781116 | - | *pcrH* | *PA14_42460* | regulatory protein PcrH | 143 | 147 | 159 | 143 | 120 | 136 | 1 |
| 3435 | - | 3782515 | 3781631 | - | *pcrV* | *PA14_42470* | type III secretion protein PcrV | 161 | 158 | 208 | 192 | 134 | 167 | 1 |
| 3436 | - | 3782821 | 3782525 | - | *pcrG* | *PA14_42480* | regulator in type III secretion | 85 | 58 | 109 | 76 | 52 | 64 | 1 |
| 3437 | - | 3783283 | 3782849 | - | *pcrR* | *PA14_42490* | transcriptional regulator protein PcrR | 28 | 24 | 42 | 31 | 33 | 22 | 1 |
| 3438 | - | 3785400 | 3783280 | - | *pcrD* | *PA14_42500* | type III secretory apparatus protein PcrD | 30 | 26 | 41 | 36 | 32 | 28 | 1 |
| 3439 | - | 3785726 | 3785397 | - | - | *PA14_42510* | hypothetical protein | 19 | 15 | 23 | 18 | 16 | 16 | 1 |
| 3440 | - | 3786096 | 3785731 | - | - | *PA14_42520* | hypothetical protein | 23 | 23 | 32 | 24 | 20 | 21 | 1 |
| 3441 | - | 3786464 | 3786093 | - | - | *PA14_42530* | type III secretion protein | 68 | 62 | 99 | 75 | 69 | 59 | 1 |
| 3442 | - | 3786729 | 3786451 | - | - | *PA14_42540* | protein in type III secretion | 41 | 38 | 58 | 56 | 42 | 36 | 1 |
| 3443 | - | 3787576 | 3786710 | - | *popN* | *PA14_42550* | Type III secretion outer membrane protein PopN precursor | 24 | 23 | 37 | 29 | 21 | 17 | 1 |
| 3444 | - | 3787767 | 3789089 | + | *pscN* | *PA14_42570* | type III secretion system ATPase | 37 | 32 | 53 | 46 | 28 | 29 | 1 |

| | A | B | C | D | E | F | G | H | I | J | K | L | M | N |
|---|---|---|---|---|---|---|---|---|---|---|---|---|---|---|
| 3445 | - | 3789086 | 3789562 | + | *pscO* | *PA14_42580* | translocation protein in type III secretion | 41 | 39 | 56 | 50 | 38 | 37 | 1 |
| 3446 | - | 3789550 | 3790668 | + | *pscP* | *PA14_42600* | translocation protein in type III secretion | 31 | 31 | 40 | 33 | 28 | 30 | 1 |
| 3447 | - | 3790665 | 3791594 | + | *pscQ* | *PA14_42610* | type III secretion system protein | 36 | 40 | 50 | 46 | 38 | 40 | 1 |
| 3448 | - | 3791591 | 3792244 | + | *pscR* | *PA14_42620* | type III secretion system protein | 28 | 29 | 36 | 32 | 31 | 30 | 1 |
| 3449 | - | 3792247 | 3792513 | + | - | *PA14_42630* | translocation protein in type III secretion | 38 | 51 | 59 | 59 | 55 | 47 | 1 |
| 3450 | - | 3792510 | 3793298 | + | *pscT* | *PA14_42640* | translocation protein in type III secretion | 15 | 15 | 22 | 22 | 19 | 15 | 1 |
| 3451 | - | 3793295 | 3794344 | + | *pscU* | *PA14_42660* | translocation protein in type III secretion | 17 | 20 | 25 | 23 | 22 | 19 | 1 |
| 3452 | - | 3796552 | 3794450 | - | - | *PA14_42670* | hypothetical protein | 48 | 59 | 48 | 50 | 56 | 44 | 1 |
| 3453 | - | 3797424 | 3796555 | - | - | *PA14_42680* | hypothetical protein | 35 | 38 | 47 | 43 | 32 | 28 | 1 |
| 3454 | - | 3798445 | 3797585 | - | *speE* | *PA14_42690* | spermidine synthase | 29 | 39 | 29 | 26 | 29 | 22 | 1 |
| 3455 | - | 3798594 | 3799487 | + | *alkA* | *PA14_42700* | DNA-3-methyladenine glycosidase II | 87 | 81 | 92 | 81 | 81 | 88 | 1 |
| 3456 | - | 3799694 | 3799503 | - | - | *PA14_42710* | hypothetical protein | 154 | 181 | 191 | 173 | 172 | 211 | 1 |
| 3457 | - | 3800420 | 3799743 | - | *masA* | *PA14_42720* | enolase-phosphatase | 63 | 65 | 61 | 54 | 61 | 52 | 1 |
| 3458 | - | 3801038 | 3800493 | - | - | *PA14_42730* | oxidase | 165 | 155 | 148 | 126 | 134 | 137 | 1 |
| 3459 | - | 3801652 | 3801035 | - | - | *PA14_42740* | methylthioribulose-1-phosphate dehydratase | 79 | 83 | 78 | 77 | 82 | 64 | 1 |
| 3460 | - | 3802800 | 3801649 | - | - | *PA14_42750* | MFS metabolite transporter | 24 | 22 | 27 | 25 | 23 | 21 | 1 |
| 3461 | - | 3803935 | 3802844 | - | *aroC* | *PA14_42760* | chorismate synthase | 92 | 68 | 111 | 92 | 79 | 83 | 1 |
| 3462 | - | 3804043 | 3805026 | + | - | *PA14_42770* | hypothetical protein | 30 | 23 | 31 | 21 | 21 | 20 | 1 |
| 3463 | - | 3805104 | 3805883 | + | - | *PA14_42780* | hypothetical protein | 85 | 91 | 114 | 86 | 100 | 97 | 1 |
| 3464 | - | 3806822 | 3805908 | - | - | *PA14_42790* | N5-glutamine S-adenosyl-L-methionine-dependent methyltransferase | 43 | 51 | 51 | 59 | 50 | 46 | 1 |
| 3465 | - | 3806993 | 3807589 | + | - | *PA14_42820* | isochorismatase family hydrolase | 193 | 171 | 220 | 135 | 179 | 167 | 1 |
| 3466 | - | 3807762 | 3808082 | + | - | *PA14_42830* | hypothetical protein | 63 | 48 | 65 | 62 | 47 | 57 | 1 |
| 3467 | - | 3808139 | 3808696 | + | - | *PA14_42840* | hypothetical protein | 47 | 33 | 49 | 40 | 31 | 42 | 1 |
| 3468 | - | 3808774 | 3809319 | + | *folE* | *PA14_42850* | GTP cyclohydrolase I | 271 | 201 | 345 | 154 | 107 | 195 | 1 |
| 3469 | - | 3809845 | 3809384 | - | - | *PA14_42860* | hypothetical protein | 1049 | 640 | 1234 | 498 | 420 | 1011 | 1 |
| 3470 | - | 3810092 | 3810472 | + | - | *PA14_42870* | hypothetical protein | 152 | 154 | 156 | 152 | 107 | 144 | 1 |
| 3471 | - | 3811489 | 3810500 | - | *stk1* | *PA14_42880* | serine-threonine kinase Stk1 | 79 | 49 | 66 | 58 | 56 | 68 | 1 |
| 3472 | - | 3812214 | 3811486 | - | *stp1* | *PA14_42890* | serine/threonine phosphoprotein phosphatase Stp1 | 94 | 32 | 38 | 46 | 53 | 80 | 0.017745004 |
| 3473 | - | 3815741 | 3812214 | - | - | *PA14_42900* | hypothetical protein | 65 | 33 | 37 | 42 | 45 | 52 | 1 |
| 3474 | - | 3816626 | 3815757 | - | - | *PA14_42910* | hypothetical protein | 62 | 23 | 24 | 33 | 45 | 59 | 0.031770039 |
| 3475 | - | 3817960 | 3816629 | - | - | *PA14_42920* | hypothetical protein | 77 | 42 | 46 | 48 | 62 | 76 | 1 |
| 3476 | - | 3818463 | 3817957 | - | - | *PA14_42940* | lipoprotein | 71 | 31 | 30 | 36 | 45 | 62 | 0.0839756 |
| 3477 | - | 3819662 | 3818469 | - | - | *PA14_42950* | hypothetical protein | 76 | 28 | 26 | 30 | 43 | 56 | 0.091217341 |
| 3478 | - | 3819820 | 3819680 | - | - | *PA14_42960* | hypothetical protein | 32 | 15 | 17 | 17 | 25 | 37 | 0.148048673 |
| 3479 | - | 3821420 | 3819909 | - | - | *PA14_42970* | transcriptional regulator | 65 | 22 | 23 | 28 | 39 | 52 | 0.05276977 |
| 3480 | - | 3824064 | 3821431 | - | - | *PA14_42980* | ClpA/B-type protease | 140 | 51 | 53 | 74 | 96 | 123 | 0.711763148 |
| 3481 | - | 3825078 | 3824071 | - | - | *PA14_42990* | hypothetical protein | 86 | 30 | 32 | 44 | 55 | 72 | 0.05276977 |
| 3482 | - | 3826622 | 3825042 | - | - | *PA14_43000* | hypothetical protein | 77 | 31 | 33 | 43 | 54 | 72 | 0.353229513 |
| 3483 | 3831541 | 3827574 | 3827167 | - | - | *PA14_43020* | hypothetical protein | 89 | 34 | 34 | 45 | 74 | 81 | 0.016396023 |
| 3484 | 3831541 | 3829062 | 3827587 | - | - | *PA14_43030* | hypothetical protein | 166 | 81 | 71 | 85 | 125 | 144 | 1 |
| 3485 | 3831541 | 3829596 | 3829090 | - | - | *PA14_43040* | hypothetical protein | 150 | 73 | 65 | 75 | 122 | 130 | 0.552627545 |
| 3486 | 3831541 | 3831188 | 3829632 | - | - | *PA14_43050* | hypothetical protein | 332 | 123 | 147 | 178 | 198 | 319 | 0.8413024 |
| 3487 | 3831541 | 3831886 | 3832404 | + | *hcpD* | *PA14_43070* | secreted protein Hcp | 47 | 21 | 17 | 26 | 27 | 32 | 0.090912888 |
| 3488 | - | 3832600 | 3834642 | + | - | *PA14_43080* | hypothetical protein | 15 | 10 | 12 | 14 | 10 | 13 | 1 |
| 3489 | - | 3834632 | 3835372 | + | - | *PA14_43090* | hypothetical protein | 17 | 17 | 16 | 17 | 15 | 17 | 1 |
| 3490 | - | 3835369 | 3840213 | + | - | *PA14_43100* | Rhs family protein | 25 | 23 | 26 | 28 | 23 | 25 | 1 |
| 3491 | - | 3841622 | 3841020 | - | - | *PA14_43110* | glutathione S-transferase | 80 | 106 | 71 | 52 | 105 | 89 | 1 |
| 3492 | - | 3842844 | 3841678 | - | - | *PA14_43130* | aminotransferase | 92 | 85 | 83 | 57 | 95 | 75 | 1 |
| 3493 | - | 3842926 | 3843402 | + | - | *PA14_43140* | MarR family transcriptional regulator | 18 | 19 | 19 | 15 | 20 | 13 | 1 |
| 3494 | - | 3844087 | 3843452 | - | - | *PA14_43150* | hypothetical protein | 28 | 25 | 21 | 21 | 19 | 14 | 1 |
| 3495 | - | 3844318 | 3845520 | + | - | *PA14_43160* | transporter | 37 | 34 | 36 | 28 | 32 | 32 | 1 |
| 3496 | - | 3846420 | 3845527 | - | - | *PA14_43170* | bile acid/Na+ symporter family transporter | 47 | 53 | 54 | 57 | 45 | 48 | 1 |
| 3497 | - | 3847340 | 3846579 | - | - | *PA14_43180* | short chain dehydrogenase | 16 | 13 | 14 | 17 | 15 | 14 | 1 |
| 3498 | - | 3847591 | 3848595 | + | - | *PA14_43190* | oxidoreductase | 29 | 25 | 27 | 30 | 24 | 29 | 1 |
| 3499 | - | 3848759 | 3850480 | + | - | *PA14_43200* | sulfate transporter | 28 | 27 | 34 | 35 | 27 | 33 | 1 |
| 3500 | - | 3852419 | 3850461 | - | - | *PA14_43220* | methyl-accepting chemotaxis transducer | 37 | 37 | 37 | 39 | 32 | 39 | 1 |
| 3501 | - | 3852621 | 3853028 | + | - | *PA14_43230* | hypothetical protein | 46 | 44 | 47 | 38 | 36 | 35 | 1 |
| 3502 | - | 3853643 | 3853035 | - | - | *PA14_43240* | hypothetical protein | 43 | 41 | 49 | 39 | 42 | 34 | 1 |
| 3503 | - | 3853781 | 3854128 | + | - | *PA14_43250* | hypothetical protein | 143 | 159 | 204 | 186 | 167 | 178 | 1 |
| 3504 | - | 3855216 | 3854107 | - | - | *PA14_43270* | tRNA 2-selenouridine synthase | 144 | 138 | 157 | 156 | 134 | 135 | 1 |
| 3505 | - | 3856250 | 3855216 | - | *selD* | *PA14_43280* | selenophosphate synthetase | 100 | 99 | 121 | 113 | 98 | 97 | 1 |
| 3506 | - | 3856612 | 3856334 | - | - | *PA14_43290* | lipoprotein | 38 | 55 | 43 | 63 | 36 | 48 | 1 |
| 3507 | - | 3856727 | 3857764 | + | - | *PA14_43300* | hypothetical protein | 46 | 45 | 47 | 45 | 46 | 41 | 1 |
| 3508 | - | 3858373 | 3857768 | - | - | *PA14_43310* | hypothetical protein | 36 | 32 | 36 | 30 | 27 | 32 | 1 |
| 3509 | - | 3859392 | 3858484 | - | - | *PA14_43320* | glutaminase | 14 | 10 | 13 | 12 | 11 | 13 | 1 |
| 3510 | - | 3860205 | 3859513 | - | *kdpE* | *PA14_43340* | two-component response regulator KdpE | 28 | 30 | 32 | 39 | 28 | 35 | 1 |
| 3511 | - | 3862963 | 3860306 | - | *kdpD* | *PA14_43350* | two-component sensor KdpD | 40 | 33 | 41 | 48 | 34 | 43 | 1 |
| 3512 | - | 3863622 | 3863071 | - | *kdpC* | *PA14_43370* | potassium-transporting ATPase subunit C | 3 | 3 | 4 | 4 | 3 | 2 | 1 |
| 3513 | - | 3865748 | 3863676 | - | *kdpB* | *PA14_43380* | potassium-transporting ATPase subunit B | 4 | 2 | 2 | 3 | 3 | 3 | 0.382662007 |
| 3514 | - | 3867454 | 3865760 | - | *kdpA* | *PA14_43400* | potassium-transporting ATPase subunit A | 2 | 2 | 3 | 2 | 2 | 3 | 1 |
| 3515 | - | 3867554 | 3867465 | - | *kdbF* | *PA14_43405* | potassium-transporting ATPase subunit F | 3 | 1 | 3 | 1 | 1 | 1 | 0.656599109 |
| 3516 | - | 3868896 | 3867742 | - | - | *PA14_43420* | acyl-CoA dehydrogenase | 25 | 27 | 27 | 27 | 24 | 24 | 1 |
| 3517 | - | 3869817 | 3868951 | - | - | *PA14_43430* | IclR family transcriptional regulator | 21 | 22 | 21 | 18 | 19 | 16 | 1 |
| 3518 | - | 3870059 | 3870844 | + | - | *PA14_43440* | enoyl-CoA hydratase | 6 | 9 | 6 | 5 | 6 | 4 | 1 |
| 3519 | - | 3870848 | 3872377 | + | - | *PA14_43460* | 3-hydroxyacyl-CoA dehydrogenase | 10 | 9 | 10 | 9 | 10 | 8 | 1 |
| 3520 | - | 3872515 | 3873162 | + | - | *PA14_43480* | GntR family transcriptional regulator | 11 | 9 | 12 | 10 | 9 | 9 | 1 |
| 3521 | - | 3873178 | 3874452 | + | - | *PA14_43490* | cyanate MFS transporter | 9 | 9 | 8 | 8 | 8 | 7 | 1 |
| 3522 | - | 3875571 | 3874504 | - | - | *PA14_43510* | hypothetical protein | 55 | 42 | 60 | 51 | 50 | 54 | 1 |
| 3523 | - | 3876543 | 3875737 | - | - | *PA14_43520* | hypothetical protein | 179 | 171 | 178 | 158 | 181 | 169 | 1 |
| 3524 | - | 3877214 | 3876552 | - | - | *PA14_43530* | hypothetical protein | 194 | 183 | 195 | 130 | 197 | 163 | 1 |
| 3525 | - | 3878154 | 3877294 | - | - | *PA14_43540* | hydrolase | 43 | 50 | 49 | 43 | 45 | 45 | 1 |
| 3526 | - | 3878963 | 3878151 | - | - | *PA14_43550* | hydrolase | 52 | 55 | 53 | 47 | 44 | 50 | 1 |
| 3527 | - | 3879706 | 3879065 | - | - | *PA14_43570* | hypothetical protein | 6 | 3 | 4 | 4 | 4 | 4 | 0.227774564 |
| 3528 | - | 3880571 | 3879747 | - | - | *PA14_43580* | transcriptional regulator | 43 | 33 | 40 | 30 | 36 | 31 | 1 |

| | A | B | C | D | E | F | G | H | I | J | K | L | M | N |
|---|---|---|---|---|---|---|---|---|---|---|---|---|---|---|
| 3529 | - | 3881066 | 3880629 | - | - | *PA14_43600* | hypothetical protein | 35 | 40 | 30 | 29 | 25 | 36 | 1 |
| 3530 | - | 3882786 | 3881119 | - | - | *PA14_43610* | AMP-binding protein | 58 | 56 | 50 | 65 | 46 | 68 | 1 |
| 3531 | - | 3883377 | 3882913 | - | - | *PA14_43620* | phosphohistidine phosphatase SixA | 97 | 94 | 95 | 84 | 89 | 81 | 1 |
| 3532 | - | 3883715 | 3883374 | - | - | *PA14_43630* | lipase | 128 | 134 | 127 | 134 | 142 | 146 | 1 |
| 3533 | - | 3884751 | 3883729 | - | *gpsA* | *PA14_43640* | NAD(P)H-dependent glycerol-3-phosphate dehydrogenase | 87 | 78 | 98 | 79 | 76 | 80 | 1 |
| 3534 | - | 3884902 | 3887019 | + | - | *PA14_43650* | hypothetical protein | 29 | 23 | 31 | 30 | 25 | 26 | 1 |
| 3535 | - | 3887191 | 3887949 | + | - | *PA14_43660* | hypothetical protein | 20 | 21 | 21 | 19 | 20 | 19 | 1 |
| 3536 | - | 3887946 | 3889901 | + | - | *PA14_43670* | sensor/response regulator hybrid | 31 | 31 | 31 | 30 | 33 | 29 | 1 |
| 3537 | - | 3890090 | 3890605 | + | *fabA* | *PA14_43680* | 3-hydroxydecanoyl-ACP dehydratase | 382 | 346 | 446 | 321 | 323 | 324 | 1 |
| 3538 | - | 3890617 | 3891834 | + | *fabB* | *PA14_43690* | 3-oxoacyl-ACP synthase | 269 | 317 | 309 | 250 | 275 | 251 | 1 |
| 3539 | - | 3891988 | 3893613 | + | - | *PA14_43710* | methyl-accepting chemotaxis transducer | 20 | 22 | 24 | 18 | 21 | 20 | 1 |
| 3540 | - | 3893718 | 3894158 | + | - | *PA14_43720* | hypothetical protein | 48 | 57 | 53 | 48 | 40 | 40 | 1 |
| 3541 | - | 3894808 | 3894332 | - | - | *PA14_43730* | hypothetical protein | 13 | 15 | 16 | 16 | 15 | 17 | 1 |
| 3542 | - | 3895910 | 3896815 | + | - | *PA14_43760* | hypothetical protein | 113 | 89 | 137 | 58 | 61 | 111 | 1 |
| 3543 | - | 3895911 | 3894877 | - | - | *PA14_43740* | hypothetical protein | 30 | 24 | 30 | 24 | 25 | 24 | 1 |
| 3544 | - | 3896854 | 3897276 | + | - | *PA14_43770* | transcriptional regulator | 19 | 15 | 23 | 9 | 10 | 16 | 1 |
| 3545 | - | 3897484 | 3897975 | + | - | *PA14_43780* | oxidoreductase | 63 | 27 | 37 | 25 | 91 | 50 | 0.071888203 |
| 3546 | - | 3897972 | 3900218 | + | - | *PA14_43790* | aldehyde dehydrogenase | 55 | 29 | 31 | 28 | 70 | 39 | 1 |
| 3547 | - | 3900268 | 3901569 | + | - | *PA14_43810* | cytochrome c | 29 | 20 | 20 | 16 | 26 | 18 | 1 |
| 3548 | - | 3902362 | 3901580 | - | - | *PA14_43820* | transcriptional regulator | 46 | 39 | 44 | 42 | 39 | 37 | 1 |
| 3549 | - | 3902443 | 3903264 | + | *panB* | *PA14_43830* | 3-methyl-2-oxobutanoate hydroxymethyltransferase | 24 | 40 | 24 | 18 | 29 | 19 | 0.807344171 |
| 3550 | - | 3904000 | 3903275 | - | - | *PA14_43840* | hypothetical protein | 140 | 241 | 135 | 91 | 213 | 110 | 1 |
| 3551 | - | 3905972 | 3904068 | - | *htpG* | *PA14_43850* | heat shock protein 90 | 207 | 404 | 237 | 97 | 321 | 164 | 1 |
| 3552 | - | 3907319 | 3906090 | - | - | *PA14_43870* | hypothetical protein | 24 | 23 | 27 | 19 | 18 | 15 | 1 |
| 3553 | - | 3907802 | 3907356 | - | - | *PA14_43880* | hypothetical protein | 14 | 22 | 16 | 16 | 14 | 13 | 1 |
| 3554 | - | 3908272 | 3907799 | - | - | *PA14_43890* | hypothetical protein | 11 | 12 | 14 | 11 | 9 | 11 | 1 |
| 3555 | - | 3908445 | 3908681 | + | - | *PA14_43900* | hypothetical protein | 232 | 469 | 228 | 198 | 272 | 277 | 0.452603163 |
| 3556 | - | 3909504 | 3908740 | - | - | *PA14_43910* | hypothetical protein | 14 | 13 | 15 | 13 | 14 | 14 | 1 |
| 3557 | - | 3910964 | 3909651 | - | *braB* | *PA14_43920* | branched chain amino acid transporter | 73 | 57 | 76 | 66 | 55 | 49 | 1 |
| 3558 | - | 3912162 | 3911275 | - | *sucD* | *PA14_43940* | succinyl-CoA synthetase subunit alpha | 665 | 851 | 489 | 362 | 624 | 364 | 1 |
| 3559 | - | 3913328 | 3912162 | - | *sucC* | *PA14_43950* | succinyl-CoA synthetase subunit beta | 710 | 852 | 613 | 376 | 626 | 366 | 1 |
| 3560 | - | 3913383 | 3913444 | ? | - | predicted RNA | - | 565 | 766 | 704 | 317 | 513 | 311 | 1 |
| 3561 | - | 3915093 | 3913657 | - | *lpdG* | *PA14_43970* | dihydrolipoamide dehydrogenase | 610 | 592 | 466 | 380 | 472 | 387 | 1 |
| 3562 | - | 3916391 | 3915162 | - | *sucB* | *PA14_44000* | dihydrolipoamide succinyltransferase | 467 | 478 | 370 | 341 | 368 | 326 | 1 |
| 3563 | - | 3919265 | 3916434 | - | *sucA* | *PA14_44010* | 2-oxoglutarate dehydrogenase E1 | 402 | 390 | 331 | 317 | 302 | 273 | 1 |
| 3564 | - | 3920229 | 3919522 | - | *sdhB* | *PA14_44020* | succinate dehydrogenase iron-sulfur subunit | 669 | 664 | 534 | 449 | 528 | 428 | 1 |
| 3565 | - | 3922013 | 3920241 | - | *sdhA* | *PA14_44030* | succinate dehydrogenase flavoprotein subunit | 598 | 596 | 509 | 501 | 476 | 367 | 1 |
| 3566 | - | 3922385 | 3922017 | - | *sdhD* | *PA14_44050* | succinate dehydrogenase (D subunit) | 452 | 385 | 390 | 318 | 337 | 246 | 1 |
| 3567 | - | 3922765 | 3922379 | - | *sdhC* | *PA14_44060* | succinate dehydrogenase, cytochrome b556 subunit | 371 | 314 | 313 | 239 | 227 | 180 | 1 |
| 3568 | - | 3923114 | 3924400 | + | *gltA* | *PA14_44070* | type II citrate synthase | 172 | 158 | 101 | 88 | 134 | 98 | 1 |
| 3569 | - | 3925123 | 3924515 | - | - | *PA14_44080* | hypothetical protein | 210 | 237 | 204 | 218 | 241 | 187 | 1 |
| 3570 | - | 3925511 | 3925257 | - | - | *PA14_44090* | Fe-S-cluster oxidoreductase | 19 | 14 | 17 | 20 | 12 | 16 | 1 |
| 3571 | - | 3926370 | 3925615 | - | - | *PA14_44100* | hypothetical protein | 69 | 53 | 80 | 76 | 61 | 63 | 1 |
| 3572 | - | 3926483 | 3926824 | + | - | *PA14_44110* | hypothetical protein | 42 | 44 | 51 | 58 | 35 | 44 | 1 |
| 3573 | - | 3927697 | 3926831 | - | - | *PA14_44120* | 3-hydroxyisobutyrate dehydrogenase | 144 | 126 | 166 | 172 | 123 | 134 | 1 |
| 3574 | - | 3927801 | 3928352 | + | - | *PA14_44130* | hypothetical protein | 29 | 28 | 40 | 36 | 28 | 29 | 1 |
| 3575 | - | 3928447 | 3928728 | + | - | *PA14_44140* | hypothetical protein | 23 | 37 | 22 | 18 | 24 | 16 | 1 |
| 3576 | - | 3929442 | 3928846 | - | - | *PA14_44150* | hypothetical protein | 46 | 56 | 51 | 46 | 47 | 46 | 1 |
| 3577 | - | 3930598 | 3929453 | - | - | *PA14_44160* | ATP-NAD kinase | 187 | 183 | 197 | 190 | 187 | 193 | 1 |
| 3578 | - | 3930808 | 3930987 | + | - | *PA14_44170* | hypothetical protein | 39 | 59 | 35 | 31 | 38 | 35 | 1 |
| 3579 | - | 3931937 | 3931056 | - | - | *PA14_44180* | LysR family transcriptional regulator | 33 | 32 | 32 | 29 | 35 | 35 | 1 |
| 3580 | - | 3932315 | 3933703 | + | - | *PA14_44190* | sugar MFS transporter | 3 | 2 | 3 | 4 | 3 | 3 | 1 |
| 3581 | - | 3933756 | 3934109 | + | - | *PA14_44200* | hypothetical protein | 13 | 10 | 16 | 12 | 12 | 13 | 1 |
| 3582 | - | 3934120 | 3935517 | + | - | *PA14_44210* | glycine/D-amino acid oxidase | 28 | 26 | 30 | 25 | 18 | 27 | 1 |
| 3583 | - | 3935669 | 3935857 | + | - | *PA14_44230* | hypothetical protein | 18 | 14 | 23 | 16 | 20 | 19 | 1 |
| 3584 | - | 3935985 | 3937349 | + | - | *PA14_44240* | glutamine synthetase | 16 | 14 | 20 | 26 | 19 | 26 | 1 |
| 3585 | - | 3937420 | 3938730 | + | - | *PA14_44260* | oxidoreductase | 24 | 34 | 37 | 50 | 56 | 55 | 1 |
| 3586 | - | 3939001 | 3938762 | - | - | *PA14_44270* | sulfur transfer protein SirA | 89 | 92 | 113 | 76 | 107 | 101 | 1 |
| 3587 | - | 3940144 | 3939086 | - | - | *PA14_44280* | RNA 2'-O-ribose methyltransferase | 40 | 33 | 36 | 35 | 32 | 30 | 1 |
| 3588 | - | 3940471 | 3943203 | + | *acnA* | *PA14_44290* | aconitate hydratase | 330 | 381 | 339 | 335 | 395 | 380 | 1 |
| 3589 | - | 3943576 | 3945141 | + | *aer* | *PA14_44300* | aerotaxis receptor Aer | 221 | 272 | 253 | 147 | 232 | 270 | 1 |
| 3590 | - | 3946336 | 3945179 | - | - | *PA14_44311* | hypothetical protein | 32 | 52 | 52 | 43 | 43 | 49 | 0.940967938 |
| 3591 | - | 3947057 | 3946425 | - | - | *PA14_44320* | hypothetical protein | 16 | 19 | 23 | 18 | 16 | 18 | 1 |
| 3592 | - | 3947245 | 3948672 | + | - | *PA14_44340* | cbb3-type cytochrome c oxidase subunit I | 533 | 550 | 578 | 379 | 452 | 254 | 1 |
| 3593 | - | 3948683 | 3949291 | + | - | *PA14_44350* | cbb3-type cytochrome c oxidase subunit II | 517 | 576 | 507 | 256 | 349 | 216 | 1 |
| 3594 | - | 3949479 | 3950405 | + | - | *PA14_44360* | cytochrome c oxidase, cbb3-type subunit III | 586 | 576 | 500 | 276 | 466 | 256 | 1 |
| 3595 | - | 3950769 | 3952196 | + | - | *PA14_44370* | cbb3-type cytochrome c oxidase subunit I | 127 | 170 | 128 | 116 | 148 | 118 | 1 |
| 3596 | - | 3952211 | 3952822 | + | - | *PA14_44380* | cbb3-type cytochrome c oxidase subunit II | 153 | 238 | 142 | 148 | 164 | 143 | 1 |
| 3597 | - | 3952819 | 3953013 | + | - | *PA14_44390* | cytochrome c oxidase subunit | 254 | 344 | 219 | 236 | 331 | 238 | 1 |
| 3598 | - | 3953010 | 3953966 | + | - | *PA14_44400* | cytochrome c oxidase, cbb3-type subunit III | 172 | 232 | 134 | 140 | 191 | 136 | 1 |
| 3599 | - | 3954185 | 3955600 | + | - | *PA14_44420* | ferredoxin | 303 | 360 | 323 | 325 | 361 | 359 | 1 |
| 3600 | - | 3955612 | 3956151 | + | - | *PA14_44430* | hypothetical protein | 195 | 237 | 220 | 237 | 207 | 247 | 1 |
| 3601 | - | 3956148 | 3958583 | + | - | *PA14_44440* | cation-transporting P-type ATPase | 67 | 71 | 68 | 72 | 71 | 77 | 1 |
| 3602 | - | 3958562 | 3958771 | + | - | *PA14_44450* | hypothetical protein | 93 | 99 | 108 | 126 | 113 | 122 | 1 |
| 3603 | - | 3958764 | 3959447 | + | - | *PA14_44460* | hypothetical protein | 44 | 42 | 45 | 44 | 44 | 43 | 1 |
| 3604 | - | 3959575 | 3960957 | + | *hemN* | *PA14_44470* | coproporphyrinogen III oxidase | 264 | 262 | 324 | 229 | 230 | 274 | 1 |
| 3605 | - | 3961473 | 3961006 | - | - | *PA14_44480* | hypothetical protein | 27 | 53 | 48 | 42 | 37 | 39 | 0.418757501 |
| 3606 | - | 3961719 | 3962453 | + | *anr* | *PA14_44490* | transcriptional regulator Anr | 286 | 256 | 302 | 213 | 191 | 232 | 1 |
| 3607 | - | 3962539 | 3963087 | + | *apt* | *PA14_44500* | adenine phosphoribosyltransferase | 46 | 41 | 40 | 32 | 33 | 36 | 1 |
| 3608 | - | 3963327 | 3964163 | + | - | *PA14_44510* | hypothetical protein | 10 | 20 | 14 | 16 | 12 | 17 | 0.542627228 |
| 3609 | - | 3964572 | 3964940 | + | - | *PA14_44520* | drug efflux transporter | 7 | 7 | 6 | 7 | 9 | 11 | 1 |
| 3610 | - | 3964934 | 3965263 | + | - | *PA14_44530* | multidrug efflux system protein MdtI | 21 | 24 | 22 | 22 | 18 | 25 | 1 |
| 3611 | - | 3965971 | 3965117 | - | - | *PA14_44540* | hypothetical protein | 22 | 27 | 27 | 27 | 23 | 27 | 1 |
| 3612 | - | 3966078 | 3967661 | + | - | *PA14_44560* | flavin-containing monooxygenase | 7 | 12 | 11 | 10 | 9 | 9 | 1 |

| | A | B | C | D | E | F | G | H | I | J | K | L | M | N |
|---|---|---|---|---|---|---|---|---|---|---|---|---|---|---|
| 3613 | - | 3967658 | 3968545 | + | - | *PA14_44570* | short-chain dehydrogenase | 17 | 21 | 18 | 14 | 14 | 15 | 1 |
| 3614 | - | 3968619 | 3969158 | + | - | *PA14_44580* | hypothetical protein | 19 | 15 | 16 | 15 | 15 | 15 | 1 |
| 3615 | - | 3970319 | 3969171 | - | - | *PA14_44590* | acyl-CoA dehydrogenase | 20 | 17 | 14 | 17 | 18 | 17 | 1 |
| 3616 | - | 3971038 | 3970442 | - | *recR* | *PA14_44610* | recombination protein RecR | 69 | 70 | 70 | 60 | 70 | 48 | 1 |
| 3617 | - | 3971443 | 3971117 | - | - | *PA14_44620* | hypothetical protein | 275 | 366 | 228 | 245 | 276 | 200 | 1 |
| 3618 | - | 3973594 | 3971489 | - | *dnaX* | *PA14_44630* | DNA polymerase III subunits gamma and tau | 90 | 89 | 84 | 93 | 89 | 81 | 1 |
| 3619 | - | 3973631 | 3974497 | + | - | *PA14_44640* | hypothetical protein | 38 | 57 | 48 | 65 | 47 | 44 | 1 |
| 3620 | - | 3975640 | 3975719 | ? | - | predicted RNA | - | 520 | 423 | 637 | 370 | 192 | 169 | 1 |
| 3621 | - | 3976224 | 3975847 | - | - | *PA14_44650* | hypothetical protein | 118 | 124 | 130 | 128 | 130 | 114 | 1 |
| 3622 | - | 3978654 | 3976270 | - | *ligA* | *PA14_44660* | NAD-dependent DNA ligase LigA | 79 | 79 | 77 | 78 | 76 | 77 | 1 |
| 3623 | - | 3979616 | 3978747 | - | *zipA* | *PA14_44670* | cell division protein ZipA | 370 | 285 | 434 | 333 | 325 | 345 | 1 |
| 3624 | - | 3983270 | 3979782 | - | - | *PA14_44680* | chromosome segregation protein | 200 | 183 | 198 | 208 | 203 | 219 | 1 |
| 3625 | - | 3983955 | 3983296 | - | - | *PA14_44690* | GntR family transcriptional regulator | 84 | 75 | 100 | 72 | 94 | 97 | 1 |
| 3626 | - | 3984136 | 3985269 | + | *alkB2* | *PA14_44700* | alkane-1 monooxygenase | 25 | 30 | 27 | 27 | 26 | 28 | 1 |
| 3627 | - | 3985489 | 3986943 | + | *xdhA* | *PA14_44710* | xanthine dehydrogenase | 28 | 34 | 37 | 33 | 32 | 39 | 1 |
| 3628 | - | 3986936 | 3989335 | + | *xdhB* | *PA14_44740* | xanthine dehydrogenase | 36 | 44 | 47 | 55 | 45 | 62 | 1 |
| 3629 | - | 3989362 | 3990201 | + | - | *PA14_44760* | xanthine dehydrogenase accessory factor X | 60 | 60 | 73 | 74 | 59 | 79 | 1 |
| 3630 | - | 3990271 | 3991575 | + | - | *PA14_44770* | guanine deaminase | 29 | 29 | 36 | 37 | 30 | 36 | 1 |
| 3631 | - | 3992408 | 3991629 | - | - | *PA14_44780* | transcriptional regulator | 109 | 80 | 102 | 80 | 80 | 84 | 1 |
| 3632 | - | 3992783 | 3994132 | + | - | *PA14_44800* | transporter | 9 | 7 | 7 | 10 | 6 | 7 | 1 |
| 3633 | - | 3994754 | 3994374 | - | - | *PA14_44820* | transthyretin family protein | 24 | 23 | 18 | 17 | 26 | 20 | 1 |
| 3634 | - | 3995127 | 3996053 | + | - | *PA14_44830* | hypothetical protein | 50 | 42 | 42 | 40 | 32 | 35 | 1 |
| 3635 | - | 3996050 | 3996565 | + | - | *PA14_44840* | hypothetical protein | 40 | 37 | 28 | 33 | 27 | 30 | 1 |
| 3636 | - | 3996583 | 3997581 | + | *alc* | *PA14_44850* | allantoicase | 77 | 58 | 83 | 62 | 51 | 64 | 1 |
| 3637 | - | 3997636 | 3998145 | + | - | *PA14_44860* | ureidoglycolate hydrolase | 46 | 42 | 38 | 43 | 39 | 44 | 1 |
| 3638 | - | 3998194 | 3999471 | + | - | *PA14_44880* | hypothetical protein | 37 | 35 | 33 | 35 | 32 | 35 | 1 |
| 3639 | - | 4000090 | 4000608 | + | *hcpA* | *PA14_44890* | secreted protein Hcp | 33 | 21 | 20 | 20 | 22 | 23 | 1 |
| 3640 | - | 4000789 | 4003317 | + | - | *PA14_44900* | hypothetical protein | 32 | 26 | 25 | 23 | 28 | 20 | 1 |
| 3641 | - | 4003314 | 4005023 | + | - | *PA14_44910* | hypothetical protein | 62 | 48 | 50 | 52 | 54 | 49 | 1 |
| 3642 | - | 4005020 | 4006162 | + | - | *PA14_44920* | hypothetical protein | 88 | 89 | 71 | 91 | 93 | 85 | 1 |
| 3643 | - | 4006175 | 4006435 | + | - | *PA14_44930* | hypothetical protein | 35 | 41 | 30 | 33 | 29 | 30 | 1 |
| 3644 | - | 4007119 | 4008486 | + | - | *PA14_44950* | transporter | 12 | 10 | 13 | 11 | 9 | 10 | 1 |
| 3645 | - | 4008907 | 4008578 | - | - | *PA14_44960* | hypothetical protein | 23 | 28 | 32 | 29 | 20 | 24 | 1 |
| 3646 | - | 4010087 | 4009092 | - | *moaA2* | *PA14_44970* | molybdenum cofactor biosynthesis protein A | 105 | 70 | 100 | 65 | 75 | 90 | 1 |
| 3647 | - | 4010323 | 4010973 | + | - | *PA14_44980* | TetR family transcriptional regulator | 23 | 24 | 22 | 23 | 19 | 23 | 1 |
| 3648 | - | 4011809 | 4011378 | - | - | *PA14_44990* | hypothetical protein | 10 | 8 | 11 | 9 | 8 | 8 | 1 |
| 3649 | - | 4012138 | 4013913 | + | *gcl* | *PA14_45000* | glyoxylate carboligase | 10 | 10 | 11 | 12 | 10 | 11 | 1 |
| 3650 | - | 4014009 | 4014791 | + | - | *PA14_45010* | hydroxypyruvate isomerase | 98 | 79 | 94 | 97 | 83 | 101 | 1 |
| 3651 | - | 4014866 | 4015756 | + | - | *PA14_45020* | oxidoreductase | 3 | 7 | 4 | 3 | 3 | 3 | 0.239905719 |
| 3652 | - | 4016053 | 4017318 | + | - | *PA14_45030* | hypothetical protein | 4 | 3 | 4 | 4 | 3 | 4 | 0.891234996 |
| 3653 | - | 4017320 | 4018753 | + | *pykF* | *PA14_45050* | pyruvate kinase | 10 | 8 | 9 | 11 | 7 | 8 | 1 |
| 3654 | - | 4019120 | 4020034 | + | - | *PA14_45060* | transporter | 5 | 3 | 5 | 6 | 4 | 4 | 1 |
| 3655 | - | 4020099 | 4020950 | + | - | *PA14_45070* | potassium channel | 11 | 9 | 14 | 13 | 10 | 12 | 1 |
| 3656 | - | 4020937 | 4021563 | + | - | *PA14_45090* | hypothetical protein | 13 | 12 | 10 | 13 | 12 | 13 | 1 |
| 3657 | - | 4021687 | 4023342 | + | - | *PA14_45100* | hypothetical protein | 11 | 12 | 11 | 10 | 13 | 9 | 1 |
| 3658 | - | 4023471 | 4024469 | + | *cysP* | *PA14_45110* | sulfate-binding protein of ABC transporter | 53 | 85 | 46 | 40 | 69 | 42 | 0.955702824 |
| 3659 | - | 4024907 | 4024551 | - | - | *PA14_45120* | hypothetical protein | 18 | 22 | 23 | 23 | 21 | 20 | 1 |
| 3660 | - | 4026202 | 4024967 | - | - | *PA14_45130* | transporter | 15 | 13 | 14 | 18 | 12 | 14 | 1 |
| 3661 | - | 4026549 | 4027325 | + | - | *PA14_45150* | transcriptional regulator | 7 | 7 | 6 | 7 | 6 | 5 | 1 |
| 3662 | - | 4027729 | 4028919 | + | - | *PA14_45170* | oxidoreductase | 6 | 8 | 8 | 8 | 8 | 7 | 1 |
| 3663 | - | 4028906 | 4030144 | + | - | *PA14_45180* | oxidoreductase | 14 | 14 | 13 | 13 | 15 | 12 | 1 |
| 3664 | - | 4030165 | 4031580 | + | - | *PA14_45190* | glycerol kinase | 17 | 19 | 21 | 19 | 19 | 17 | 1 |
| 3665 | - | 4032688 | 4031588 | - | - | *PA14_45210* | hypothetical protein | 9 | 14 | 11 | 12 | 12 | 11 | 1 |
| 3666 | - | 4034069 | 4032702 | - | - | *PA14_45240* | amino acid permease | 3 | 3 | 4 | 3 | 3 | 3 | 1 |
| 3667 | - | 4034269 | 4035075 | + | - | *PA14_45250* | transcriptional regulator | 20 | 23 | 21 | 23 | 20 | 19 | 1 |
| 3668 | - | 4035513 | 4035079 | - | - | *PA14_45260* | hypothetical protein | 199 | 176 | 194 | 175 | 196 | 139 | 1 |
| 3669 | - | 4036747 | 4035524 | - | *cycH* | *PA14_45280* | cytochrome c-type biogenesis protein | 222 | 221 | 256 | 245 | 249 | 198 | 1 |
| 3670 | - | 4037211 | 4036744 | - | *ccmH* | *PA14_45290* | cytochrome C-type biogenesis protein CcmH | 220 | 227 | 242 | 253 | 240 | 224 | 1 |
| 3671 | - | 4037750 | 4037208 | - | *ccmG* | *PA14_45300* | cytochrome C biogenesis protein CcmG | 206 | 201 | 216 | 256 | 250 | 224 | 1 |
| 3672 | - | 4039727 | 4037754 | - | *ccmF* | *PA14_45310* | cytochrome C-type biogenesis protein CcmF | 248 | 250 | 256 | 273 | 271 | 219 | 1 |
| 3673 | - | 4040216 | 4039728 | - | *ccmE* | *PA14_45330* | cytochrome c-type biogenesis protein CcmE | 375 | 341 | 484 | 467 | 358 | 332 | 1 |
| 3674 | - | 4040389 | 4040213 | - | - | *PA14_45340* | heme exporter protein | 215 | 199 | 242 | 259 | 208 | 189 | 1 |
| 3675 | - | 4041144 | 4040386 | - | *ccmC* | *PA14_45350* | heme exporter protein CcmC | 188 | 146 | 220 | 219 | 154 | 168 | 1 |
| 3676 | - | 4041938 | 4041267 | - | *ccmB* | *PA14_45370* | heme exporter protein CcmB | 70 | 64 | 83 | 102 | 69 | 64 | 1 |
| 3677 | - | 4042636 | 4041935 | - | *ccmA* | *PA14_45380* | cytochrome c biogenesis protein CcmA | 30 | 26 | 43 | 35 | 24 | 20 | 1 |
| 3678 | - | 4042922 | 4044400 | + | - | *PA14_45400* | hypothetical protein | 129 | 131 | 156 | 169 | 143 | 157 | 1 |
| 3679 | - | 4044397 | 4044732 | + | - | *PA14_45410* | FlhB domain-containing protein | 62 | 81 | 76 | 95 | 90 | 96 | 1 |
| 3680 | - | 4045993 | 4045256 | - | - | *PA14_45430* | short chain dehydrogenase | 30 | 29 | 33 | 32 | 27 | 29 | 1 |
| 3681 | - | 4046849 | 4046127 | - | - | *PA14_45440* | hypothetical protein | 26 | 23 | 26 | 26 | 21 | 20 | 1 |
| 3682 | - | 4047325 | 4046909 | - | - | *PA14_45450* | hypothetical protein | 15 | 20 | 20 | 22 | 15 | 17 | 1 |
| 3683 | - | 4048387 | 4047416 | - | - | *PA14_45460* | hypothetical protein | 23 | 24 | 25 | 23 | 22 | 22 | 1 |
| 3684 | - | 4048496 | 4049182 | + | - | *PA14_45470* | hypothetical protein | 31 | 30 | 36 | 29 | 27 | 26 | 1 |
| 3685 | - | 4049617 | 4049210 | - | - | *PA14_45480* | hypothetical protein | 564 | 492 | 660 | 532 | 528 | 511 | 1 |
| 3686 | - | 4050127 | 4049648 | - | - | *PA14_45500* | purine-binding chemotaxis protein | 616 | 757 | 688 | 856 | 844 | 856 | 1 |
| 3687 | - | 4051063 | 4050173 | - | - | *PA14_45510* | CheW domain-containing protein | 242 | 236 | 263 | 231 | 259 | 258 | 1 |
| 3688 | - | 4051943 | 4051155 | - | - | *PA14_45520* | plasmid partitioning protein | 168 | 173 | 191 | 211 | 189 | 198 | 1 |
| 3689 | - | 4052928 | 4052038 | - | *motD* | *PA14_45540* | flagellar motor protein MotD | 161 | 157 | 189 | 196 | 182 | 186 | 1 |
| 3690 | - | 4053681 | 4052941 | - | *motC* | *PA14_45560* | flagellar motor protein | 348 | 310 | 406 | 357 | 379 | 386 | 1 |
| 3691 | - | 4054876 | 4053770 | - | - | *PA14_45580* | chemotaxis-specific methylesterase | 376 | 417 | 428 | 452 | 424 | 483 | 1 |
| 3692 | - | 4057191 | 4054930 | - | - | *PA14_45590* | two-component sensor | 387 | 375 | 456 | 435 | 392 | 412 | 1 |
| 3693 | - | 4058180 | 4057392 | - | *cheZ* | *PA14_45610* | chemotaxis protein CheZ | 393 | 392 | 426 | 401 | 387 | 450 | 1 |
| 3694 | - | 4058574 | 4058200 | - | *cheY* | *PA14_45620* | two-component response regulator CheY | 394 | 396 | 411 | 310 | 286 | 375 | 1 |
| 3695 | - | 4059419 | 4058676 | - | *fliA* | *PA14_45630* | flagellar biosynthesis sigma factor | 334 | 333 | 380 | 230 | 222 | 333 | 1 |
| 3696 | - | 4060258 | 4059416 | - | *fleN* | *PA14_45640* | flagellar synthesis regulator FleN | 431 | 403 | 451 | 415 | 382 | 431 | 1 |

| | A | B | C | D | E | F | G | H | I | J | K | L | M | N |
|---|---|---|---|---|---|---|---|---|---|---|---|---|---|---|
| 3697 | - | 4061686 | 4060397 | - | *flhF* | *PA14_45660* | flagellar biosynthesis regulator FlhF | 188 | 189 | 192 | 199 | 184 | 178 | 1 |
| 3698 | - | 4063893 | 4061770 | - | *flhA* | *PA14_45680* | flagellar biosynthesis protein FlhA | 52 | 45 | 57 | 57 | 52 | 48 | 1 |
| 3699 | - | 4065393 | 4064032 | - | - | *PA14_45700* | hypothetical protein | 46 | 31 | 38 | 30 | 36 | 29 | 1 |
| 3700 | - | 4066685 | 4065426 | - | - | *PA14_45710* | hypothetical protein | 58 | 39 | 53 | 45 | 38 | 33 | 1 |
| 3701 | - | 4067894 | 4066758 | - | *flhB* | *PA14_45720* | flagellar biosynthesis protein FlhB | 36 | 35 | 48 | 49 | 47 | 49 | 1 |
| 3702 | - | 4068673 | 4067897 | - | *fliR* | *PA14_45740* | flagellar biosynthesis protein FliR | 46 | 42 | 50 | 44 | 43 | 46 | 1 |
| 3703 | - | 4068942 | 4068673 | - | *fliQ* | *PA14_45760* | flagellar biosynthesis protein FliQ | 221 | 220 | 233 | 288 | 234 | 233 | 1 |
| 3704 | - | 4069754 | 4068987 | - | *fliP* | *PA14_45770* | flagellar biosynthesis protein FliP | 63 | 58 | 68 | 66 | 56 | 61 | 1 |
| 3705 | - | 4070203 | 4069751 | - | *fliO* | *PA14_45780* | flagellar protein FliO | 134 | 116 | 149 | 131 | 116 | 114 | 1 |
| 3706 | - | 4070678 | 4070205 | - | *fliN* | *PA14_45790* | flagellar motor switch protein | 227 | 248 | 235 | 269 | 221 | 233 | 1 |
| 3707 | - | 4071677 | 4070706 | - | *fliM* | *PA14_45800* | flagellar motor switch protein FliM | 176 | 130 | 162 | 162 | 144 | 160 | 1 |
| 3708 | - | 4072206 | 4071685 | - | *fliL* | *PA14_45810* | flagellar basal body-associated protein FliL | 90 | 62 | 90 | 87 | 74 | 78 | 1 |
| 3709 | - | 4073733 | 4072450 | - | - | *PA14_45830* | hypothetical protein | 125 | 154 | 144 | 124 | 157 | 128 | 1 |
| 3710 | - | 4073921 | 4074508 | + | - | *PA14_45840* | hypothetical protein | 185 | 166 | 224 | 189 | 144 | 143 | 1 |
| 3711 | - | 4074520 | 4074927 | + | - | *PA14_45850* | hypothetical protein | 67 | 61 | 64 | 67 | 53 | 61 | 1 |
| 3712 | - | 4076381 | 4074936 | - | - | *PA14_45870* | two-component sensor | 74 | 64 | 66 | 79 | 61 | 71 | 1 |
| 3713 | - | 4077048 | 4076359 | - | - | *PA14_45880* | two-component response regulator | 62 | 42 | 56 | 56 | 55 | 54 | 1 |
| 3714 | - | 4080262 | 4077152 | - | - | *PA14_45890* | RND efflux transporter | 20 | 18 | 23 | 22 | 16 | 20 | 1 |
| 3715 | - | 4081416 | 4080259 | - | - | *PA14_45910* | RND efflux membrane fusion protein | 14 | 12 | 17 | 16 | 12 | 15 | 1 |
| 3716 | - | 4081597 | 4082310 | + | - | *PA14_45920* | hypothetical protein | 15 | 13 | 16 | 19 | 15 | 14 | 1 |
| 3717 | - | 4082311 | 4084263 | + | - | *PA14_45930* | hypothetical protein | 42 | 35 | 35 | 34 | 28 | 35 | 1 |
| 3718 | - | 4084975 | 4084370 | - | *lasI* | *PA14_45940* | autoinducer synthesis protein LasI | 722 | 511 | 486 | 390 | 413 | 535 | 1 |
| 3719 | - | 4085107 | 4085349 | + | *rsaL* | *PA14_45950* | regulatory protein RsaL | 91 | 119 | 110 | 124 | 105 | 103 | 1 |
| 3720 | - | 4086058 | 4085339 | - | *lasR* | *PA14_45960* | transcriptional regulator LasR | 614 | 643 | 699 | 869 | 703 | 762 | 1 |
| 3721 | - | 4086142 | 4086151 | ? | - | predicted RNA | - | 496 | 518 | 601 | 853 | 736 | 655 | 1 |
| 3722 | - | 4089217 | 4086509 | - | - | *PA14_45970* | cation-transporting P-type ATPase | 259 | 218 | 377 | 101 | 126 | 211 | 1 |
| 3723 | - | 4089814 | 4089383 | - | - | *PA14_45980* | acetyltransferase | 12 | 11 | 15 | 8 | 9 | 10 | 1 |
| 3724 | - | 4091588 | 4091232 | - | - | *PA14_46000* | hypothetical protein | 19 | 20 | 18 | 15 | 16 | 18 | 1 |
| 3725 | - | 4093245 | 4091629 | - | - | *PA14_46010* | ABC transporter ATP-binding protein | 7 | 6 | 6 | 6 | 6 | 5 | 1 |
| 3726 | - | 4094345 | 4093734 | - | - | *PA14_46020* | hypothetical protein | 24 | 22 | 27 | 24 | 19 | 20 | 1 |
| 3727 | - | 4094639 | 4095892 | + | - | *PA14_46030* | chemotaxis transducer | 19 | 20 | 18 | 19 | 19 | 18 | 1 |
| 3728 | - | 4096789 | 4095896 | - | *gbuR* | *PA14_46060* | transcriptional regulator | 16 | 15 | 18 | 19 | 14 | 17 | 1 |
| 3729 | - | 4096997 | 4097956 | + | *gbuA* | *PA14_46070* | guanidinobutyrase | 32 | 11 | 11 | 22 | 9 | 12 | 0.002852411 |
| 3730 | - | 4097971 | 4098390 | + | - | *PA14_46080* | hypothetical protein | 28 | 13 | 15 | 18 | 9 | 11 | 0.048133675 |
| 3731 | - | 4098409 | 4099920 | + | - | *PA14_46100* | transporter | 10 | 4 | 4 | 7 | 3 | 3 | 0.00773671 |
| 3732 | - | 4100014 | 4101405 | + | - | *PA14_46110* | sodium:solute symport protein | 27 | 13 | 14 | 21 | 14 | 13 | 0.279198294 |
| 3733 | - | 4101481 | 4103082 | + | - | *PA14_46120* | hypothetical protein | 16 | 7 | 7 | 12 | 6 | 6 | 0.122746259 |
| 3734 | - | 4103096 | 4104478 | + | - | *PA14_46140* | hypothetical protein | 21 | 12 | 12 | 13 | 9 | 10 | 0.726761916 |
| 3735 | - | 4104523 | 4105251 | + | - | *PA14_46150* | hypothetical protein | 113 | 96 | 157 | 87 | 101 | 118 | 1 |
| 3736 | - | 4105544 | 4105311 | - | - | *PA14_46160* | hypothetical protein | 365 | 510 | 478 | 360 | 350 | 363 | 1 |
| 3737 | - | 4106472 | 4105600 | - | - | *PA14_46170* | LysR family transcriptional regulator | 26 | 23 | 26 | 23 | 25 | 30 | 1 |
| 3738 | - | 4106577 | 4107770 | + | - | *PA14_46180* | hypothetical protein | 10 | 8 | 10 | 9 | 7 | 8 | 1 |
| 3739 | - | 4108641 | 4107730 | - | - | *PA14_46200* | hypothetical protein | 16 | 18 | 20 | 16 | 16 | 13 | 1 |
| 3740 | - | 4109928 | 4108837 | - | - | *PA14_46220* | periplasmic spermidine/putrescine-binding protein | 18 | 13 | 19 | 12 | 12 | 15 | 1 |
| 3741 | - | 4110979 | 4109939 | - | *aphA* | *PA14_46230* | acetylpolyamine aminohydrolase | 14 | 12 | 14 | 14 | 13 | 10 | 1 |
| 3742 | - | 4111189 | 4113612 | + | - | *PA14_46240* | hypothetical protein | 41 | 33 | 44 | 45 | 40 | 49 | 1 |
| 3743 | - | 4113679 | 4114620 | + | - | *PA14_46250* | hypothetical protein | 16 | 18 | 18 | 19 | 18 | 24 | 1 |
| 3744 | - | 4114656 | 4115414 | + | - | *PA14_46260* | hypothetical protein | 34 | 37 | 36 | 34 | 38 | 44 | 1 |
| 3745 | - | 4117026 | 4115434 | - | - | *PA14_46270* | helicase | 44 | 54 | 41 | 42 | 48 | 39 | 1 |
| 3746 | - | 4117560 | 4117787 | + | - | *PA14_46280* | hypothetical protein | 50 | 77 | 70 | 77 | 67 | 62 | 1 |
| 3747 | - | 4118551 | 4117919 | - | - | *PA14_46290* | TetR family transcriptional regulator | 37 | 40 | 42 | 46 | 35 | 38 | 1 |
| 3748 | - | 4118644 | 4119309 | + | - | *PA14_46300* | hypothetical protein | 31 | 34 | 37 | 32 | 31 | 36 | 1 |
| 3749 | - | 4119387 | 4119794 | + | - | *PA14_46310* | hypothetical protein | 40 | 38 | 35 | 40 | 42 | 43 | 1 |
| 3750 | - | 4123109 | 4119822 | - | - | *PA14_46320* | pyruvate carboxylase | 12 | 12 | 14 | 15 | 11 | 12 | 1 |
| 3751 | - | 4123314 | 4124234 | + | - | *PA14_46330* | transcriptional regulator | 25 | 23 | 30 | 25 | 23 | 20 | 1 |
| 3752 | - | 4124591 | 4124250 | - | - | *PA14_46340* | hypothetical protein | 36 | 33 | 33 | 32 | 29 | 23 | 1 |
| 3753 | - | 4125304 | 4124672 | - | - | *PA14_46360* | two-component response regulator | 22 | 28 | 31 | 26 | 21 | 17 | 1 |
| 3754 | - | 4125450 | 4127072 | + | - | *PA14_46370* | two-component sensor | 17 | 14 | 13 | 12 | 13 | 10 | 1 |
| 3755 | - | 4127515 | 4127099 | - | - | *PA14_46380* | hypothetical protein | 18 | 14 | 15 | 15 | 16 | 14 | 1 |
| 3756 | - | 4127785 | 4127531 | - | - | *PA14_46390* | hypothetical protein | 20 | 19 | 24 | 20 | 18 | 15 | 1 |
| 3757 | - | 4129094 | 4128129 | - | - | *PA14_46400* | transcriptional regulator | 24 | 21 | 18 | 18 | 18 | 18 | 1 |
| 3758 | - | 4129186 | 4130019 | + | - | *PA14_46420* | short chain dehydrogenase | 20 | 21 | 18 | 16 | 17 | 15 | 1 |
| 3759 | - | 4130471 | 4130109 | - | - | *PA14_46430* | hypothetical protein | 70 | 56 | 64 | 56 | 59 | 54 | 1 |
| 3760 | - | 4131147 | 4130614 | - | - | *PA14_46440* | acetyltransferase | 77 | 70 | 68 | 67 | 62 | 52 | 1 |
| 3761 | - | 4131283 | 4133016 | + | *aceK* | *PA14_46450* | bifunctional isocitrate dehydrogenase kinase/phosphatase | 72 | 74 | 71 | 83 | 70 | 71 | 1 |
| 3762 | - | 4134890 | 4133700 | - | - | *PA14_46460* | hypothetical protein | 40 | 27 | 34 | 24 | 39 | 32 | 1 |
| 3763 | - | 4136281 | 4135139 | - | *pdxB* | *PA14_46470* | erythronate-4-phosphate dehydrogenase | 53 | 53 | 77 | 48 | 65 | 51 | 1 |
| 3764 | - | 4137002 | 4136496 | - | - | *PA14_46480* | transcriptional regulator | 5 | 9 | 5 | 7 | 6 | 7 | 1 |
| 3765 | - | 4137111 | 4138379 | + | *fabF2* | *PA14_46490* | 3-oxoacyl-ACP synthase | 10 | 11 | 12 | 12 | 11 | 11 | 1 |
| 3766 | - | 4140104 | 4138746 | - | - | *PA14_46510* | hypothetical protein | 172 | 155 | 182 | 266 | 160 | 245 | 1 |
| 3767 | - | 4141240 | 4140101 | - | - | *PA14_46520* | hypothetical protein | 205 | 186 | 185 | 356 | 160 | 267 | 1 |
| 3768 | - | 4142251 | 4141826 | - | - | *PA14_46530* | hypothetical protein | 86 | 59 | 80 | 108 | 49 | 67 | 1 |
| 3769 | - | 4142401 | 4144101 | + | - | *PA14_46540* | hypothetical protein | 77 | 54 | 67 | 102 | 53 | 93 | 1 |
| 3770 | - | 4144573 | 4145301 | + | - | *PA14_46550* | ribonuclease | 76 | 61 | 60 | 86 | 67 | 83 | 1 |
| 3771 | - | 4145865 | 4145464 | - | - | *PA14_46560* | outer membrane protein | 15 | 15 | 14 | 17 | 12 | 19 | 1 |
| 3772 | - | 4146137 | 4146754 | + | - | *PA14_46570* | transcriptional regulator | 66 | 54 | 54 | 56 | 43 | 46 | 1 |
| 3773 | - | 4146826 | 4147941 | + | - | *PA14_46580* | multidrug efflux gene product | 7 | 4 | 4 | 4 | 4 | 4 | 0.831807123 |
| 3774 | - | 4147938 | 4151027 | + | - | *PA14_46590* | AcrB/AcrD/AcrF family protein | 13 | 11 | 11 | 12 | 9 | 9 | 1 |
| 3775 | - | 4151058 | 4151864 | + | - | *PA14_46600* | short chain dehydrogenase | 49 | 50 | 65 | 55 | 42 | 49 | 1 |
| 3776 | - | 4152822 | 4152349 | - | - | *PA14_46610* | hypothetical protein | 20 | 20 | 18 | 17 | 16 | 21 | 1 |
| 3777 | - | 4154124 | 4152844 | - | - | *PA14_46620* | pyridine nucleotide-disulfide oxidoreductase | 28 | 21 | 27 | 25 | 21 | 26 | 1 |
| 3778 | - | 4154793 | 4155629 | + | - | *PA14_46630* | hypothetical protein | 8 | 10 | 9 | 9 | 9 | 7 | 1 |
| 3779 | - | 4158230 | 4155789 | - | - | *PA14_46640* | siderophore receptor | 8 | 9 | 10 | 12 | 9 | 7 | 1 |
| 3780 | - | 4159150 | 4158308 | - | - | *PA14_46650* | transmembrane sensor | 10 | 12 | 11 | 21 | 13 | 8 | 1 |

| | A | B | C | D | E | F | G | H | I | J | K | L | M | N |
|---|---|---|---|---|---|---|---|---|---|---|---|---|---|---|
| 3781 | - | 4159887 | 4159147 | - | - | *PA14_46660* | RNA polymerase ECF-subfamily sigma-70 factor | 6 | 8 | 7 | 15 | 10 | 6 | 1 |
| 3782 | - | 4160223 | 4159900 | - | - | *PA14_46670* | hypothetical protein | 14 | 14 | 14 | 12 | 10 | 13 | 1 |
| 3783 | - | 4160669 | 4162102 | + | - | *PA14_46680* | transporter | 39 | 37 | 49 | 41 | 35 | 28 | 1 |
| 3784 | - | 4162977 | 4163627 | + | - | *PA14_46710* | transcriptional regulator | 25 | 21 | 32 | 23 | 20 | 16 | 1 |
| 3785 | - | 4162978 | 4162085 | - | - | *PA14_46700* | hypothetical protein | 40 | 34 | 46 | 37 | 34 | 32 | 1 |
| 3786 | - | 4163665 | 4164063 | + | - | *PA14_46720* | hypothetical protein | 56 | 70 | 62 | 60 | 67 | 57 | 1 |
| 3787 | - | 4164555 | 4164079 | - | - | *PA14_46740* | hypothetical protein | 54 | 67 | 58 | 66 | 64 | 55 | 1 |
| 3788 | - | 4165730 | 4164624 | - | - | *PA14_46750* | hypothetical protein | 28 | 24 | 27 | 37 | 23 | 34 | 1 |
| 3789 | - | 4166133 | 4165753 | - | - | *PA14_46760* | hypothetical protein | 19 | 17 | 22 | 27 | 17 | 22 | 1 |
| 3790 | - | 4166280 | 4166705 | + | - | *PA14_46770* | hypothetical protein | 19 | 19 | 20 | 26 | 15 | 16 | 1 |
| 3791 | - | 4166742 | 4167155 | + | - | *PA14_46780* | hypothetical protein | 9 | 11 | 9 | 11 | 12 | 8 | 1 |
| 3792 | - | 4167327 | 4168538 | + | - | *PA14_46800* | hypothetical protein | 8 | 8 | 9 | 11 | 10 | 11 | 1 |
| 3793 | - | 4169769 | 4168546 | - | - | *PA14_46810* | RNA polymerase ECF-subfamily sigma-70 factor | 33 | 34 | 48 | 49 | 37 | 47 | 1 |
| 3794 | - | 4170536 | 4169787 | - | - | *PA14_46820* | hypothetical protein | 36 | 36 | 41 | 56 | 34 | 51 | 1 |
| 3795 | - | 4171061 | 4170711 | - | - | *PA14_46830* | hypothetical protein | 17 | 19 | 21 | 23 | 17 | 22 | 1 |
| 3796 | - | 4171951 | 4171244 | - | - | *PA14_46840* | hypothetical protein | 64 | 66 | 68 | 101 | 66 | 75 | 1 |
| 3797 | - | 4172017 | 4172709 | + | - | *PA14_46850* | transcriptional regulator | 7 | 5 | 5 | 6 | 3 | 5 | 1 |
| 3798 | - | 4172793 | 4174340 | + | - | *PA14_46860* | lysine decarboxylase | 4 | 4 | 5 | 6 | 4 | 4 | 1 |
| 3799 | - | 4175940 | 4174351 | - | - | *PA14_46880* | glutathione synthase | 13 | 14 | 12 | 14 | 11 | 12 | 1 |
| 3800 | - | 4176918 | 4176124 | - | - | *PA14_46890* | short-chain dehydrogenase | 120 | 126 | 123 | 129 | 120 | 131 | 1 |
| 3801 | - | 4177472 | 4177014 | - | - | *PA14_46900* | hypothetical protein | 33 | 35 | 27 | 21 | 28 | 25 | 1 |
| 3802 | - | 4177924 | 4178832 | + | - | *PA14_46910* | ABC transporter substrate-binding protein | 307 | 573 | 307 | 267 | 413 | 306 | 1 |
| 3803 | - | 4179035 | 4179781 | + | - | *PA14_46920* | ABC transporter permease | 68 | 101 | 61 | 64 | 91 | 71 | 1 |
| 3804 | - | 4179778 | 4180446 | + | - | *PA14_46930* | ABC transporter permease | 124 | 180 | 101 | 134 | 157 | 138 | 1 |
| 3805 | - | 4180443 | 4181177 | + | - | *PA14_46950* | ABC transporter ATP-binding protein | 206 | 315 | 150 | 198 | 274 | 212 | 1 |
| 3806 | - | 4181255 | 4182928 | + | *ggt* | *PA14_46960* | gamma-glutamyltranspeptidase | 119 | 143 | 94 | 109 | 131 | 124 | 1 |
| 3807 | - | 4183014 | 4184102 | + | *ansB* | *PA14_46970* | glutaminase-asparaginase | 332 | 534 | 227 | 257 | 455 | 318 | 1 |
| 3808 | - | 4184198 | 4186099 | + | - | *PA14_46980* | two-component sensor | 33 | 39 | 32 | 33 | 48 | 38 | 1 |
| 3809 | - | 4186144 | 4187421 | + | - | *PA14_46990* | two-component response regulator | 58 | 66 | 49 | 55 | 61 | 58 | 1 |
| 3810 | - | 4187705 | 4188817 | + | - | *PA14_47000* | oxidoreductase | 10 | 9 | 10 | 10 | 10 | 9 | 1 |
| 3811 | - | 4189012 | 4189218 | + | - | *PA14_47010* | hypothetical protein | 118 | 71 | 114 | 76 | 73 | 81 | 0.86477425 |
| 3812 | - | 4189368 | 4189919 | + | - | *PA14_47030* | hypothetical protein | 63 | 51 | 70 | 58 | 65 | 72 | 1 |
| 3813 | - | 4190188 | 4191735 | + | - | *PA14_47040* | TerC family protein | 45 | 35 | 45 | 31 | 30 | 24 | 1 |
| 3814 | - | 4192570 | 4191791 | - | - | *PA14_47060* | short chain dehydrogenase | 23 | 22 | 23 | 18 | 19 | 15 | 1 |
| 3815 | - | 4193007 | 4192576 | - | - | *PA14_47070* | hypothetical protein | 2 | 2 | 2 | 1 | 0 | 1 | 1 |
| 3816 | - | 4193092 | 4194000 | + | - | *PA14_47080* | transcriptional regulator | 26 | 26 | 33 | 32 | 27 | 36 | 1 |
| 3817 | - | 4195932 | 4194004 | - | - | *PA14_47090* | protease | 75 | 77 | 90 | 110 | 77 | 103 | 1 |
| 3818 | - | 4197566 | 4196019 | - | *ilvA2* | *PA14_47100* | threonine dehydratase | 11 | 13 | 13 | 11 | 14 | 10 | 1 |
| 3819 | - | 4197970 | 4197563 | - | - | *PA14_47110* | hypothetical protein | 15 | 13 | 15 | 13 | 16 | 14 | 1 |
| 3820 | - | 4198748 | 4198236 | - | - | *PA14_47120* | hypothetical protein | 393 | 579 | 426 | 371 | 571 | 446 | 1 |
| 3821 | - | 4199140 | 4198808 | - | - | *PA14_47130* | hypothetical protein | 434 | 586 | 554 | 391 | 478 | 427 | 1 |
| 3822 | - | 4201467 | 4199269 | - | - | *PA14_47140* | TonB-dependent receptor | 7 | 7 | 8 | 7 | 7 | 7 | 1 |
| 3823 | - | 4202596 | 4201706 | - | *cyoE* | *PA14_47150* | protoheme IX farnesyltransferase | 25 | 7 | 8 | 10 | 7 | 13 | 1.44415E-06 |
| 3824 | - | 4202942 | 4202607 | - | *cyoD* | *PA14_47160* | cytochrome o ubiquinol oxidase subunit IV | 27 | 6 | 6 | 7 | 8 | 22 | 1.62282E-10 |
| 3825 | - | 4203571 | 4202942 | - | *cyoC* | *PA14_47180* | cytochrome o ubiquinol oxidase subunit III | 27 | 7 | 7 | 8 | 9 | 19 | 1.12702E-10 |
| 3826 | - | 4205551 | 4203575 | - | *cyoB* | *PA14_47190* | cytochrome o ubiquinol oxidase subunit I | 30 | 13 | 17 | 17 | 14 | 26 | 0.401409914 |
| 3827 | - | 4206553 | 4205558 | - | *cyoA* | *PA14_47210* | cytochrome o ubiquinol oxidase subunit II | 25 | 14 | 14 | 17 | 12 | 20 | 0.573569409 |
| 3828 | - | 4208721 | 4207180 | - | - | *PA14_47230* | MFS transporter | 17 | 28 | 23 | 26 | 21 | 20 | 0.873872452 |
| 3829 | - | 4208879 | 4209493 | + | - | *PA14_47240* | transcriptional regulator | 21 | 23 | 22 | 24 | 21 | 21 | 1 |
| 3830 | - | 4209887 | 4209501 | - | - | *PA14_47250* | hypothetical protein | 50 | 45 | 48 | 46 | 36 | 42 | 1 |
| 3831 | - | 4211259 | 4209901 | - | - | *PA14_47260* | MFS transporter | 9 | 8 | 11 | 9 | 6 | 6 | 1 |
| 3832 | - | 4211367 | 4212278 | + | - | *PA14_47270* | LysR family transcriptional regulator | 17 | 16 | 18 | 16 | 15 | 17 | 1 |
| 3833 | - | 4213156 | 4212329 | - | *phnX* | *PA14_47280* | phosphonoacetaldehyde hydrolase | 18 | 16 | 18 | 21 | 18 | 17 | 1 |
| 3834 | - | 4214307 | 4213192 | - | *phnW* | *PA14_47300* | 2-aminoethylphosphonate--pyruvate transaminase | 6 | 6 | 6 | 7 | 6 | 6 | 1 |
| 3835 | - | 4214475 | 4215335 | + | - | *PA14_47310* | LysR family transcriptional regulator | 46 | 49 | 44 | 44 | 42 | 39 | 1 |
| 3836 | - | 4215869 | 4215345 | - | - | *PA14_47320* | hypothetical protein | 116 | 130 | 135 | 112 | 121 | 121 | 1 |
| 3837 | - | 4216669 | 4215896 | - | - | *PA14_47330* | hypothetical protein | 180 | 181 | 176 | 165 | 210 | 196 | 1 |
| 3838 | - | 4217134 | 4216673 | - | - | *PA14_47340* | HIT family protein | 191 | 170 | 188 | 175 | 210 | 199 | 1 |
| 3839 | - | 4217671 | 4217198 | - | - | *PA14_47350* | hypothetical protein | 95 | 64 | 134 | 65 | 63 | 69 | 1 |
| 3840 | - | 4217820 | 4219871 | + | - | *PA14_47360* | oligopeptidase | 25 | 28 | 25 | 24 | 23 | 22 | 1 |
| 3841 | - | 4219948 | 4220487 | + | - | *PA14_47370* | signal peptidase | 10 | 11 | 11 | 12 | 10 | 8 | 1 |
| 3842 | - | 4223049 | 4220494 | - | - | *PA14_47380* | heme utilization protein | 19 | 15 | 22 | 21 | 14 | 15 | 1 |
| 3843 | - | 4224161 | 4223178 | - | - | *PA14_47390* | transmembrane sensor | 3 | 15 | 2 | 62 | 13 | 3 | 3.42958E-13 |
| 3844 | - | 4224685 | 4224158 | - | - | *PA14_47400* | RNA polymerase ECF-subfamily sigma-70 factor | 2 | 34 | 2 | 125 | 48 | 3 | 1.2645E-182 |
| 3845 | - | 4225284 | 4224835 | - | - | *PA14_47410* | hypothetical protein | 8 | 6 | 6 | 7 | 7 | 6 | 1 |
| 3846 | - | 4225636 | 4225361 | - | - | *PA14_47420* | hypothetical protein | 19 | 24 | 24 | 21 | 19 | 23 | 1 |
| 3847 | - | 4225704 | 4226654 | + | - | *PA14_47430* | metal transporter | 18 | 21 | 19 | 14 | 15 | 13 | 1 |
| 3848 | - | 4227598 | 4226666 | - | - | *PA14_47440* | 2-hydroxyacid dehydrogenase | 76 | 67 | 77 | 76 | 73 | 68 | 1 |
| 3849 | - | 4228146 | 4227853 | - | - | *PA14_47450* | hypothetical protein | 31 | 37 | 34 | 34 | 30 | 26 | 1 |
| 3850 | - | 4229267 | 4228143 | - | *rnd* | *PA14_47460* | ribonuclease D | 51 | 39 | 54 | 47 | 41 | 41 | 1 |
| 3851 | - | 4229408 | 4230469 | + | - | *PA14_47490* | hypothetical protein | 24 | 42 | 26 | 24 | 34 | 21 | 0.767922893 |
| 3852 | - | 4230769 | 4231623 | + | *sseA* | *PA14_47500* | 3-mercaptopyruvate sulfurtransferase | 65 | 76 | 63 | 56 | 68 | 61 | 1 |
| 3853 | - | 4232484 | 4231645 | - | - | *PA14_47510* | hydrolase | 62 | 69 | 59 | 57 | 56 | 49 | 1 |
| 3854 | - | 4232596 | 4233189 | + | - | *PA14_47520* | transcriptional regulator | 62 | 59 | 56 | 40 | 53 | 56 | 1 |
| 3855 | - | 4233411 | 4233890 | + | - | *PA14_47530* | hypothetical protein | 48 | 51 | 61 | 48 | 47 | 43 | 1 |
| 3856 | - | 4234096 | 4235370 | + | - | *PA14_47540* | outer membrane protein | 221 | 365 | 202 | 169 | 249 | 176 | 1 |
| 3857 | - | 4235979 | 4235425 | - | - | *PA14_47550* | glutathione peroxidase | 82 | 74 | 80 | 79 | 61 | 93 | 1 |
| 3858 | - | 4237343 | 4236144 | - | - | *PA14_47560* | MFS transporter | 4 | 4 | 4 | 4 | 3 | 3 | 1 |
| 3859 | - | 4237866 | 4237417 | - | - | *PA14_47580* | MarR family transcriptional regulator | 32 | 43 | 37 | 34 | 23 | 27 | 1 |
| 3860 | - | 4237997 | 4239817 | + | - | *PA14_47600* | acyl-CoA dehydrogenase | 25 | 25 | 24 | 25 | 27 | 24 | 1 |
| 3861 | - | 4240408 | 4239848 | - | - | *PA14_47610* | transcriptional regulator | 187 | 197 | 162 | 197 | 289 | 264 | 1 |
| 3862 | - | 4240527 | 4242032 | + | - | *PA14_47640* | major facilitator transporter | 10 | 13 | 11 | 8 | 17 | 11 | 1 |
| 3863 | - | 4242732 | 4241995 | - | *cobS* | *PA14_47650* | cobalamin synthase | 31 | 29 | 25 | 21 | 34 | 21 | 1 |
| 3864 | - | 4243319 | 4242735 | - | - | *PA14_47660* | hypothetical protein | 39 | 37 | 32 | 27 | 37 | 30 | 1 |

| | A | B | C | D | E | F | G | H | I | J | K | L | M | N |
|---|---|---|---|---|---|---|---|---|---|---|---|---|---|---|
| 3865 | - | 4244371 | 4243316 | - | *cobT* | *PA14_47670* | nicotinate-nucleotide--dimethylbenzimidazole phosphoribosyltransferase | 81 | 76 | 68 | 54 | 88 | 60 | 1 |
| 3866 | - | 4244889 | 4244368 | - | *cobU* | *PA14_47680* | adenosylcobinamide kinase/adenosylcobinamide-phosphate guanylyltransferase | 82 | 73 | 67 | 59 | 82 | 61 | 1 |
| 3867 | - | 4246354 | 4244882 | - | *cobQ* | *PA14_47690* | cobyric acid synthase | 59 | 55 | 48 | 44 | 59 | 41 | 1 |
| 3868 | - | 4247342 | 4246347 | - | *cobC* | *PA14_47720* | threonine-phosphate decarboxylase | 36 | 38 | 31 | 27 | 40 | 29 | 1 |
| 3869 | - | 4248273 | 4247335 | - | *cobD* | *PA14_47730* | cobalamin biosynthesis protein | 33 | 30 | 32 | 26 | 37 | 25 | 1 |
| 3870 | - | 4248926 | 4248270 | - | - | *PA14_47750* | hypothetical protein | 48 | 44 | 42 | 35 | 47 | 38 | 1 |
| 3871 | - | 4250235 | 4248928 | - | *cobB* | *PA14_47760* | cobyrinic acid a,c-diamide synthase | 48 | 48 | 46 | 41 | 61 | 41 | 1 |
| 3872 | - | 4250878 | 4250267 | - | *cobO* | *PA14_47790* | cob(I)yrinic acid a,c-diamide adenosyltransferase | 90 | 101 | 84 | 78 | 99 | 81 | 1 |
| 3873 | - | 4252792 | 4250942 | - | - | *PA14_47800* | tonB-dependent receptor | 29 | 38 | 34 | 28 | 37 | 25 | 1 |
| 3874 | - | 4255183 | 4253144 | - | - | *PA14_47810* | hypothetical protein | 5 | 4 | 6 | 6 | 4 | 5 | 1 |
| 3875 | - | 4255428 | 4256096 | + | - | *PA14_47820* | transcriptional regulator | 28 | 21 | 32 | 20 | 25 | 24 | 1 |
| 3876 | - | 4256213 | 4257157 | + | - | *PA14_47840* | hypothetical protein | 12 | 10 | 11 | 12 | 9 | 9 | 1 |
| 3877 | - | 4257154 | 4258269 | + | - | *PA14_47850* | hypothetical protein | 8 | 7 | 8 | 8 | 6 | 7 | 1 |
| 3878 | - | 4258499 | 4259752 | + | - | *PA14_47860* | oxidoreductase | 10 | 10 | 10 | 12 | 9 | 9 | 1 |
| 3879 | - | 4260651 | 4259749 | - | - | *PA14_47870* | hypothetical protein | 10 | 10 | 11 | 10 | 7 | 9 | 1 |
| 3880 | - | 4260751 | 4261617 | + | - | *PA14_47880* | transcriptional regulator | 27 | 26 | 28 | 23 | 27 | 24 | 1 |
| 3881 | - | 4262858 | 4261914 | - | - | *PA14_47890* | hypothetical protein | 23 | 31 | 24 | 28 | 28 | 28 | 1 |
| 3882 | - | 4262897 | 4264339 | + | - | *PA14_47900* | MFS transporter | 4 | 3 | 4 | 4 | 3 | 3 | 1 |
| 3883 | - | 4264504 | 4265178 | + | - | *PA14_47910* | transcriptional regulator | 27 | 26 | 26 | 24 | 18 | 26 | 1 |
| 3884 | - | 4265435 | 4266256 | + | - | *PA14_47920* | ABC transporter substrate-binding protein | 2 | 2 | 3 | 3 | 1 | 1 | 1 |
| 3885 | - | 4266315 | 4268057 | + | - | *PA14_47930* | hypothetical protein | 12 | 11 | 12 | 12 | 10 | 11 | 1 |
| 3886 | - | 4268057 | 4268722 | + | - | *PA14_47940* | ABC transporter permease | 3 | 3 | 3 | 3 | 2 | 2 | 1 |
| 3887 | - | 4268733 | 4269377 | + | - | *PA14_47950* | ABC transporter permease | 4 | 5 | 6 | 3 | 3 | 3 | 1 |
| 3888 | - | 4269374 | 4270096 | + | - | *PA14_47960* | ABC transporter ATP-binding protein | 8 | 7 | 8 | 9 | 7 | 6 | 1 |
| 3889 | - | 4270122 | 4271156 | + | - | *PA14_47970* | hypothetical protein | 31 | 27 | 36 | 36 | 26 | 28 | 1 |
| 3890 | - | 4271307 | 4272224 | + | - | *PA14_48000* | dihydrodipicolinate synthetase | 4 | 4 | 4 | 4 | 5 | 4 | 1 |
| 3891 | - | 4272317 | 4273897 | + | - | *PA14_48010* | semialdehyde dehydrogenase | 18 | 16 | 21 | 18 | 16 | 19 | 1 |
| 3892 | - | 4275005 | 4274001 | - | - | *PA14_48020* | L-malate dehydrogenase | 59 | 50 | 51 | 57 | 51 | 58 | 1 |
| 3893 | - | 4275245 | 4276870 | + | - | *PA14_48030* | methyl-accepting chemotaxis transducer | 158 | 146 | 155 | 155 | 152 | 161 | 1 |
| 3894 | - | 4277278 | 4276883 | - | *aprI* | *PA14_48040* | alkaline proteinase inhibitor AprI | 385 | 428 | 350 | 400 | 391 | 387 | 1 |
| 3895 | - | 4278964 | 4277525 | - | *aprA* | *PA14_48060* | alkaline metalloproteinase | 1009 | 1199 | 768 | 1095 | 1355 | 1466 | 1 |
| 3896 | 4285523 | 4280768 | 4279323 | - | *aprF* | *PA14_48090* | alkaline protease secretion outer membrane protein AprF precursor | 163 | 181 | 170 | 213 | 182 | 213 | 1 |
| 3897 | 4285523 | 4282070 | 4280772 | - | *aprE* | *PA14_48100* | alkaline protease secretion protein AprE | 268 | 279 | 294 | 422 | 315 | 402 | 1 |
| 3898 | 4285523 | 4283848 | 4282067 | - | *aprD* | *PA14_48115* | alkaline protease secretion protein AprD | 179 | 187 | 190 | 257 | 190 | 225 | 1 |
| 3899 | 4285523 | 4285179 | 4283935 | - | - | *PA14_48140* | hypothetical protein | 184 | 195 | 203 | 267 | 180 | 194 | 1 |
| 3900 | - | 4286195 | 4286536 | + | - | *PA14_48150* | hypothetical protein | 18 | 20 | 17 | 20 | 14 | 14 | 1 |
| 3901 | - | 4286772 | 4289348 | + | - | *PA14_48160* | sensor/response regulator hybrid | 51 | 55 | 62 | 73 | 69 | 66 | 1 |
| 3902 | - | 4291143 | 4289371 | - | - | *PA14_48170* | hypothetical protein | 39 | 39 | 47 | 40 | 44 | 42 | 1 |
| 3903 | - | 4291783 | 4291223 | - | - | *PA14_48190* | transcriptional regulator | 12 | 7 | 11 | 9 | 7 | 8 | 0.447575716 |
| 3904 | - | 4291948 | 4292745 | + | - | *PA14_48200* | enoyl-CoA hydratase | 6 | 5 | 5 | 6 | 6 | 8 | 1 |
| 3905 | - | 4292802 | 4293686 | + | - | *PA14_48210* | hydrolase | 6 | 9 | 7 | 9 | 7 | 9 | 1 |
| 3906 | - | 4293859 | 4293960 | + | - | *PA14_48230* | hypothetical protein | 2 | 2 | 5 | 5 | 4 | 5 | 1 |
| 3907 | - | 4294069 | 4295517 | + | - | *PA14_48240* | outer membrane component of multidrug efflux pump | 82 | 70 | 99 | 82 | 75 | 82 | 1 |
| 3908 | - | 4295514 | 4296665 | + | - | *PA14_48280* | multidrug resistance efflux pump | 6 | 7 | 6 | 6 | 6 | 6 | 1 |
| 3909 | - | 4296655 | 4298166 | + | - | *PA14_48300* | MFS transporter | 5 | 5 | 5 | 6 | 5 | 5 | 1 |
| 3910 | - | 4298951 | 4298169 | - | - | *PA14_48310* | transcriptional regulator | 35 | 34 | 28 | 37 | 32 | 34 | 1 |
| 3911 | - | 4299143 | 4299646 | + | - | *PA14_48320* | hypothetical protein | 52 | 47 | 49 | 48 | 42 | 42 | 1 |
| 3912 | - | 4299925 | 4299668 | - | - | *PA14_48330* | hypothetical protein | 40 | 38 | 39 | 39 | 36 | 35 | 1 |
| 3913 | - | 4301961 | 4299922 | - | - | *PA14_48340* | hypothetical protein | 10 | 8 | 9 | 8 | 7 | 8 | 1 |
| 3914 | - | 4302869 | 4301961 | - | - | *PA14_48350* | hypothetical protein | 3 | 4 | 3 | 3 | 2 | 2 | 1 |
| 3915 | - | 4303079 | 4302882 | - | - | *PA14_48380* | hypothetical protein | 1 | 1 | 1 | 1 | 1 | 0 | 1 |
| 3916 | - | 4303213 | 4304019 | + | - | *PA14_48390* | AraC family transcriptional regulator | 29 | 23 | 27 | 20 | 22 | 25 | 1 |
| 3917 | - | 4304129 | 4304479 | + | - | *PA14_48400* | hypothetical protein | 13 | 16 | 15 | 12 | 16 | 15 | 1 |
| 3918 | - | 4305404 | 4304556 | - | - | *PA14_48410* | hypothetical protein | 56 | 54 | 57 | 49 | 60 | 50 | 1 |
| 3919 | - | 4306051 | 4305443 | - | - | *PA14_48420* | transcriptional regulator | 35 | 45 | 42 | 36 | 36 | 33 | 1 |
| 3920 | - | 4306157 | 4306762 | + | - | *PA14_48440* | NAD(P)H dehydrogenase | 16 | 21 | 17 | 13 | 18 | 14 | 1 |
| 3921 | - | 4307897 | 4306779 | - | - | *PA14_48450* | peptidyl-arginine deiminase | 19 | 17 | 18 | 18 | 19 | 19 | 1 |
| 3922 | - | 4309009 | 4307897 | - | - | *PA14_48460* | polyamine ABC transporter substrate-binding protein | 5 | 4 | 5 | 6 | 5 | 6 | 1 |
| 3923 | - | 4309917 | 4309006 | - | - | *PA14_48470* | N-carbamoylputrescine amidohydrolase | 7 | 6 | 6 | 6 | 5 | 5 | 1 |
| 3924 | - | 4310961 | 4309921 | - | - | *PA14_48490* | peptidylarginine deiminase | 7 | 5 | 5 | 4 | 4 | 6 | 1 |
| 3925 | - | 4311093 | 4311980 | + | - | *PA14_48500* | transcriptional regulator | 10 | 12 | 10 | 13 | 11 | 13 | 1 |
| 3926 | - | 4312055 | 4312390 | + | - | *PA14_48510* | hypothetical protein | 20 | 15 | 19 | 20 | 15 | 19 | 1 |
| 3927 | - | 4312387 | 4313544 | + | - | *PA14_48520* | membrane-bound lytic murein transglycosylase A | 48 | 47 | 46 | 48 | 40 | 44 | 1 |
| 3928 | 4313855 | 4314228 | 4316084 | + | - | *PA14_48530* | AMP-binding protein | 15 | 5 | 11 | 15 | 12 | 28 | 0.004810474 |
| 3929 | 4313855 | 4316081 | 4317343 | + | - | *PA14_48540* | hypothetical protein | 20 | 3 | 5 | 9 | 5 | 40 | 1.09298E-23 |
| 3930 | 4313855 | 4317337 | 4317951 | + | - | *PA14_48550* | hypothetical protein | 21 | 9 | 9 | 15 | 9 | 39 | 0.012478882 |
| 3931 | 4313855 | 4317948 | 4318826 | + | - | *PA14_48560* | hypothetical protein | 37 | 11 | 17 | 22 | 16 | 66 | 2.34134E-05 |
| 3932 | 4313855 | 4318823 | 4320190 | + | - | *PA14_48570* | 2-isopropylmalate synthase | 63 | 20 | 24 | 32 | 27 | 99 | 0.016569495 |
| 3933 | 4313855 | 4320187 | 4320933 | + | - | *PA14_48590* | hypothetical protein | 62 | 7 | 7 | 18 | 16 | 124 | 3.98951E-84 |
| 3934 | - | 4321135 | 4322421 | + | - | *PA14_48600* | AMP-binding protein | 33 | 5 | 8 | 11 | 12 | 71 | 1.83255E-31 |
| 3935 | - | 4322421 | 4324019 | + | - | *PA14_48610* | sparagine synthase | 24 | 3 | 6 | 6 | 6 | 48 | 2.59705E-66 |
| 3936 | - | 4324016 | 4324975 | + | - | *PA14_48620* | clavaminic acid synthetase | 23 | 4 | 6 | 8 | 8 | 44 | 1.15967E-30 |
| 3937 | - | 4324972 | 4326201 | + | - | *PA14_48630* | MFS transporter | 18 | 3 | 4 | 8 | 7 | 42 | 2.77756E-23 |
| 3938 | - | 4326198 | 4326833 | + | - | *PA14_48640* | hypothetical protein | 41 | 11 | 15 | 18 | 17 | 77 | 2.20041E-08 |
| 3939 | - | 4326951 | 4327649 | + | - | *PA14_48650* | hypothetical protein | 18 | 17 | 17 | 19 | 12 | 18 | 1 |
| 3940 | - | 4327830 | 4328765 | + | - | *PA14_48660* | hypothetical protein | 32 | 32 | 34 | 33 | 35 | 33 | 1 |
| 3941 | - | 4330281 | 4328782 | - | - | *PA14_48680* | hypothetical protein | 28 | 30 | 30 | 33 | 33 | 38 | 1 |
| 3942 | - | 4332289 | 4330448 | - | *kefB* | *PA14_48700* | glutathione-regulated potassium-efflux system protein KefB | 65 | 63 | 61 | 66 | 69 | 66 | 1 |
| 3943 | - | 4332765 | 4332292 | - | - | *PA14_48710* | hypothetical protein | 33 | 29 | 36 | 32 | 27 | 34 | 1 |
| 3944 | - | 4333872 | 4332925 | - | - | *PA14_48730* | hypothetical protein | 45 | 41 | 37 | 33 | 37 | 32 | 1 |
| 3945 | - | 4334456 | 4333899 | - | - | *PA14_48740* | hypothetical protein | 60 | 56 | 51 | 44 | 54 | 54 | 1 |
| 3946 | - | 4334892 | 4334485 | - | - | *PA14_48750* | hypothetical protein | 26 | 27 | 24 | 17 | 21 | 16 | 1 |
| 3947 | - | 4335652 | 4335035 | - | - | *PA14_48760* | hydrolase | 48 | 60 | 44 | 54 | 61 | 69 | 1 |
| 3948 | - | 4335953 | 4336867 | + | - | *PA14_48770* | transcriptional regulator | 50 | 51 | 51 | 64 | 49 | 48 | 1 |

| | A | B | C | D | E | F | G | H | I | J | K | L | M | N |
|---|---|---|---|---|---|---|---|---|---|---|---|---|---|---|
| 3949 | - | 4337537 | 4336875 | - | - | *PA14_48780* | hypothetical protein | 71 | 84 | 72 | 102 | 76 | 71 | 1 |
| 3950 | - | 4338090 | 4337557 | - | - | *PA14_48790* | lipoprotein | 100 | 107 | 103 | 104 | 89 | 80 | 1 |
| 3951 | - | 4338790 | 4338173 | - | - | *PA14_48800* | hypothetical protein | 225 | 231 | 258 | 260 | 160 | 161 | 1 |
| 3952 | - | 4339626 | 4338856 | - | - | *PA14_48810* | NAD-dependent deacetylase | 76 | 88 | 87 | 51 | 39 | 71 | 1 |
| 3953 | - | 4341039 | 4339639 | - | - | *PA14_48830* | transcriptional regulator | 85 | 84 | 98 | 38 | 28 | 80 | 1 |
| 3954 | - | 4341250 | 4342014 | + | - | *PA14_48840* | hypothetical protein | 11 | 12 | 10 | 12 | 12 | 12 | 1 |
| 3955 | - | 4342119 | 4343537 | + | - | *PA14_48850* | amino acid permease | 8 | 9 | 10 | 10 | 9 | 7 | 1 |
| 3956 | - | 4344210 | 4343539 | - | - | *PA14_48860* | hypothetical protein | 35 | 37 | 36 | 37 | 35 | 32 | 1 |
| 3957 | - | 4345081 | 4344257 | - | - | *PA14_48870* | C32 tRNA thiolase | 33 | 24 | 34 | 27 | 20 | 23 | 1 |
| 3958 | - | 4346203 | 4345187 | - | - | *PA14_48880* | bacteriophage integrase | 21 | 15 | 17 | 17 | 17 | 16 | 1 |
| 3959 | - | 4347501 | 4346203 | - | - | *PA14_48890* | hypothetical protein | 8 | 7 | 9 | 7 | 7 | 9 | 1 |
| 3960 | - | 4347733 | 4347632 | - | - | *PA14_48900* | hypothetical protein | 5 | 5 | 4 | 2 | 4 | 7 | 1 |
| 3961 | - | 4349013 | 4347730 | - | - | *PA14_48910* | hypothetical protein | 9 | 8 | 9 | 8 | 9 | 8 | 1 |
| 3962 | - | 4349373 | 4349017 | - | - | *PA14_48920* | bacteriophage protein | 3 | 4 | 4 | 3 | 3 | 2 | 1 |
| 3963 | - | 4350664 | 4349378 | - | - | *PA14_48930* | coat protein A of bacteriophage Pf1 | 5 | 5 | 5 | 7 | 6 | 5 | 1 |
| 3964 | - | 4351099 | 4350851 | - | *coaB* | *PA14_48940* | coat protein B of bacteriophage Pf1) | 85 | 83 | 63 | 64 | 86 | 84 | 1 |
| 3965 | - | 4351363 | 4351112 | - | - | *PA14_48950* | hypothetical protein | 22 | 21 | 14 | 17 | 19 | 16 | 1 |
| 3966 | - | 4351468 | 4351376 | - | - | *PA14_48960* | hypothetical protein | 10 | 7 | 5 | 6 | 6 | 4 | 1 |
| 3967 | - | 4351919 | 4351485 | - | - | *PA14_48970* | helix destabilizing protein of bacteriophage Pf1 | 3 | 5 | 3 | 3 | 3 | 1 | 1 |
| 3968 | - | 4352431 | 4352054 | - | - | *PA14_48980* | hypothetical protein | 4 | 4 | 4 | 4 | 3 | 2 | 1 |
| 3969 | - | 4352725 | 4352435 | - | - | *PA14_48990* | hypothetical protein | 3 | 4 | 3 | 4 | 2 | 3 | 1 |
| 3970 | - | 4352941 | 4352729 | - | - | *PA14_49000* | hypothetical protein | 3 | 5 | 5 | 4 | 5 | 5 | 1 |
| 3971 | - | 4353159 | 4352944 | - | - | *PA14_49010* | hypothetical protein | 7 | 5 | 3 | 5 | 5 | 5 | 1 |
| 3972 | - | 4353287 | 4353553 | + | - | *PA14_49020* | hypothetical protein | 42 | 45 | 45 | 35 | 35 | 33 | 1 |
| 3973 | - | 4353842 | 4354873 | + | - | *PA14_49030* | hypothetical protein | 60 | 80 | 44 | 42 | 53 | 38 | 1 |
| 3974 | - | 4355849 | 4356442 | + | - | *PA14_49040* | hypothetical protein | 19 | 19 | 24 | 22 | 19 | 22 | 1 |
| 3975 | - | 4356601 | 4357200 | + | - | *PA14_49050* | hypothetical protein | 28 | 27 | 33 | 30 | 22 | 30 | 1 |
| 3976 | - | 4357368 | 4357865 | + | - | *PA14_49060* | hypothetical protein | 20 | 22 | 25 | 30 | 21 | 25 | 1 |
| 3977 | - | 4359048 | 4357867 | - | - | *PA14_49070* | hypothetical protein | 36 | 42 | 42 | 51 | 38 | 44 | 1 |
| 3978 | - | 4360276 | 4359137 | - | - | *PA14_49080* | acyl-CoA dehydrogenase | 6 | 7 | 6 | 8 | 6 | 6 | 1 |
| 3979 | - | 4361397 | 4360378 | - | - | *PA14_49090* | hypothetical protein | 9 | 8 | 8 | 9 | 8 | 8 | 1 |
| 3980 | - | 4362023 | 4361394 | - | - | *PA14_49100* | glutathione S-transferase | 6 | 6 | 9 | 6 | 5 | 7 | 1 |
| 3981 | - | 4362225 | 4363115 | + | - | *PA14_49110* | transcriptional regulator | 25 | 20 | 20 | 21 | 15 | 17 | 1 |
| 3982 | - | 4364500 | 4363190 | - | *dctA* | *PA14_49130* | C4-dicarboxylate transporter DctA | 118 | 65 | 44 | 51 | 66 | 50 | 1 |
| 3983 | - | 4364836 | 4365837 | + | - | *PA14_49150* | transcriptional regulator | 47 | 55 | 64 | 75 | 61 | 67 | 1 |
| 3984 | - | 4369203 | 4365841 | - | - | *PA14_49160* | sensor protein | 62 | 58 | 73 | 97 | 76 | 85 | 1 |
| 3985 | - | 4370651 | 4369305 | - | *phoQ* | *PA14_49170* | two-component sensor PhoQ | 136 | 120 | 134 | 124 | 154 | 145 | 1 |
| 3986 | - | 4371325 | 4370648 | - | *phoP* | *PA14_49180* | two-component response regulator PhoP | 272 | 267 | 239 | 180 | 286 | 245 | 1 |
| 3987 | - | 4372007 | 4371405 | - | *oprH* | *PA14_49200* | PhoP/Q and low Mg2+ inducible outer membrane prote | 418 | 528 | 309 | 275 | 468 | 369 | 1 |
| 3988 | - | 4372229 | 4372396 | + | *napE* | *PA14_49210* | periplasmic nitrate reductase NapE | 733 | 681 | 910 | 847 | 663 | 711 | 1 |
| 3989 | - | 4372405 | 4372896 | + | *napF* | *PA14_49220* | ferredoxin protein NapF | 362 | 329 | 436 | 388 | 308 | 350 | 1 |
| 3990 | - | 4372889 | 4373218 | + | *napD* | *PA14_49230* | NapD protein of periplasmic nitrate reductase | 85 | 84 | 91 | 88 | 75 | 99 | 1 |
| 3991 | - | 4373199 | 4375703 | + | *napA* | *PA14_49250* | nitrate reductase catalytic subunit | 236 | 245 | 264 | 334 | 238 | 321 | 1 |
| 3992 | - | 4375714 | 4376205 | + | *napB* | *PA14_49260* | cytochrome c-type protein NapB precursor | 83 | 97 | 97 | 106 | 91 | 117 | 1 |
| 3993 | - | 4376216 | 4376812 | + | *napc* | *PA14_49270* | cytochrome c-type protein NapC | 88 | 95 | 99 | 109 | 109 | 136 | 1 |
| 3994 | - | 4378040 | 4376844 | - | - | *PA14_49280* | transglycosylase | 48 | 54 | 54 | 55 | 57 | 50 | 1 |
| 3995 | - | 4378497 | 4379231 | + | - | *PA14_49290* | hypothetical protein | 11 | 7 | 11 | 8 | 11 | 7 | 0.711763148 |
| 3996 | - | 4381332 | 4379275 | - | - | *PA14_49300* | lipoxygenase | 21 | 17 | 17 | 21 | 17 | 18 | 1 |
| 3997 | 4382169 | 4381730 | 4381410 | - | - | *PA14_49310* | hypothetical protein | 17 | 18 | 14 | 21 | 20 | 22 | 1 |
| 3998 | - | 4382901 | 4382230 | - | - | *PA14_49320* | hypothetical protein | 77 | 70 | 90 | 81 | 85 | 92 | 1 |
| 3999 | - | 4383077 | 4383865 | + | - | *PA14_49330* | hypothetical protein | 29 | 24 | 33 | 37 | 23 | 27 | 1 |
| 4000 | - | 4384093 | 4384821 | + | *pcpS* | *PA14_49340* | protein Pcprotein PS | 27 | 30 | 32 | 31 | 28 | 26 | 1 |
| 4001 | - | 4385634 | 4384822 | - | - | *PA14_49350* | hypothetical protein | 110 | 88 | 122 | 121 | 99 | 96 | 1 |
| 4002 | - | 4385846 | 4388455 | + | - | *PA14_49360* | glucosyl transferase | 21 | 20 | 28 | 30 | 22 | 29 | 1 |
| 4003 | - | 4388570 | 4389721 | + | *dapE* | *PA14_49380* | succinyl-diaminopimelate desuccinylase | 25 | 19 | 23 | 20 | 21 | 18 | 1 |
| 4004 | - | 4389721 | 4390524 | + | *rrmA* | *PA14_49390* | rRNA methyltransferase | 20 | 22 | 24 | 19 | 20 | 15 | 1 |
| 4005 | - | 4390542 | 4390925 | + | - | *PA14_49400* | hypothetical protein | 75 | 74 | 87 | 69 | 61 | 64 | 1 |
| 4006 | - | 4391240 | 4391031 | - | - | *PA14_49410* | cold-shock protein | 1108 | 1866 | 1051 | 956 | 652 | 532 | 1 |
| 4007 | - | 4392918 | 4391560 | - | - | *PA14_49420* | two-component sensor | 27 | 24 | 26 | 29 | 27 | 22 | 1 |
| 4008 | - | 4393970 | 4393260 | - | - | *PA14_49440* | two-component response regulator | 39 | 35 | 41 | 38 | 32 | 30 | 1 |
| 4009 | - | 4394703 | 4397594 | + | *nrdA* | *PA14_49460* | ribonucleotide-diphosphate reductase subunit alpha | 111 | 168 | 108 | 96 | 115 | 79 | 1 |
| 4010 | - | 4397858 | 4399105 | + | *nrdB* | *PA14_49470* | ribonucleotide-diphosphate reductase subunit beta | 157 | 279 | 155 | 116 | 186 | 119 | 1 |
| 4011 | - | 4399221 | 4401008 | + | - | *PA14_49480* | hypothetical protein | 24 | 35 | 18 | 15 | 24 | 14 | 1 |
| 4012 | - | 4401791 | 4402345 | + | - | *PA14_49500* | hypothetical protein | 72 | 60 | 57 | 62 | 72 | 57 | 1 |
| 4013 | - | 4403159 | 4402698 | - | *pyoS3I* | *PA14_49510* | immunity protein S3I structureal gene | 216 | 278 | 223 | 153 | 256 | 184 | 1 |
| 4014 | - | 4405534 | 4403159 | - | *pyoS3A* | *PA14_49520* | pyocin killing protein | 115 | 175 | 109 | 137 | 169 | 128 | 1 |
| 4015 | - | 4405954 | 4406322 | + | - | *PA14_49530* | hypothetical protein | 16 | 12 | 12 | 14 | 13 | 12 | 1 |
| 4016 | - | 4406350 | 4407129 | + | - | *PA14_49540* | hypothetical protein | 38 | 33 | 27 | 30 | 28 | 31 | 1 |
| 4017 | - | 4407441 | 4409348 | + | *toxA* | *PA14_49560* | exotoxin A | 6 | 6 | 6 | 6 | 6 | 5 | 1 |
| 4018 | - | 4409608 | 4411098 | + | - | *PA14_49570* | amino acid permease | 3 | 2 | 3 | 3 | 3 | 3 | 1 |
| 4019 | - | 4411200 | 4412381 | + | - | *PA14_49580* | iron-containing alcohol dehydrogenase | 9 | 7 | 10 | 10 | 8 | 8 | 1 |
| 4020 | - | 4413288 | 4412386 | - | - | *PA14_49590* | transcriptional regulator | 13 | 11 | 12 | 14 | 12 | 11 | 1 |
| 4021 | - | 4413782 | 4415116 | + | - | *PA14_49610* | MFS family transporter | 2 | 2 | 2 | 2 | 2 | 2 | 1 |
| 4022 | - | 4415184 | 4415837 | + | - | *PA14_49620* | phosphoserine phosphatase | 2 | 1 | 1 | 2 | 2 | 2 | 0.033575691 |
| 4023 | - | 4415919 | 4416635 | + | - | *PA14_49630* | transcriptional regulator | 13 | 13 | 13 | 11 | 12 | 10 | 1 |
| 4024 | - | 4416801 | 4417703 | + | - | *PA14_49640* | transcriptional regulator | 16 | 18 | 16 | 14 | 13 | 12 | 1 |
| 4025 | - | 4417840 | 4418676 | + | - | *PA14_49650* | hypothetical protein | 30 | 25 | 23 | 19 | 19 | 17 | 1 |
| 4026 | - | 4419622 | 4418774 | - | - | *PA14_49660* | hypothetical protein | 31 | 23 | 26 | 20 | 21 | 25 | 1 |
| 4027 | - | 4420588 | 4419713 | - | - | *PA14_49680* | transcriptional regulator | 15 | 12 | 12 | 11 | 11 | 13 | 1 |
| 4028 | - | 4420734 | 4421720 | + | - | *PA14_49690* | oxidoreductase | 34 | 121 | 52 | 113 | 190 | 144 | 0.000555972 |
| 4029 | - | 4422458 | 4421727 | - | - | *PA14_49700* | transcriptional regulator | 42 | 56 | 41 | 56 | 69 | 63 | 1 |
| 4030 | - | 4422630 | 4423505 | + | - | *PA14_49710* | chaperone protein HchA | 19 | 22 | 17 | 15 | 19 | 18 | 1 |
| 4031 | - | 4423761 | 4424174 | + | - | *PA14_49720* | hypothetical protein | 13 | 12 | 15 | 10 | 8 | 9 | 1 |
| 4032 | - | 4424241 | 4424591 | + | - | *PA14_49730* | hypothetical protein | 29 | 20 | 25 | 25 | 23 | 22 | 1 |

| | A | B | C | D | E | F | G | H | I | J | K | L | M | N |
|---|---|---|---|---|---|---|---|---|---|---|---|---|---|---|
| 4033 | 4425570 | 4425290 | 4424601 | - | - | *PA14_49740* | hypothetical protein | 56 | 61 | 74 | 58 | 59 | 49 | 1 |
| 4034 | 4425570 | 4425603 | 4426871 | + | - | *PA14_49750* | MFS family transporter | 26 | 3 | 12 | 29 | 21 | 28 | 1.14807E-74 |
| 4035 | 4425570 | 4426859 | 4427836 | + | *rhlC* | *PA14_49760* | rhamnosyltransferase 2 | 73 | 16 | 28 | 71 | 49 | 72 | 5.63449E-09 |
| 4036 | - | 4428240 | 4427833 | - | - | *PA14_49780* | fosfomycin resistance protein | 46 | 35 | 34 | 50 | 45 | 49 | 1 |
| 4037 | - | 4429186 | 4428275 | - | - | *PA14_49790* | transcriptional regulator | 20 | 19 | 21 | 23 | 21 | 17 | 1 |
| 4038 | - | 4429321 | 4430310 | + | - | *PA14_49800* | oxidoreductase | 123 | 95 | 94 | 97 | 104 | 108 | 1 |
| 4039 | - | 4430405 | 4430992 | + | - | *PA14_49810* | hypothetical protein | 32 | 30 | 28 | 30 | 30 | 32 | 1 |
| 4040 | - | 4431751 | 4430999 | - | - | *PA14_49820* | cobalamin biosynthetic protein | 90 | 80 | 78 | 94 | 85 | 82 | 1 |
| 4041 | - | 4433261 | 4431765 | - | *dgt* | *PA14_49840* | deoxyguanosinetriphosphate triphosphohydrolase | 78 | 73 | 77 | 85 | 75 | 78 | 1 |
| 4042 | - | 4433492 | 4433812 | + | - | *PA14_49850* | hypothetical protein | 219 | 324 | 257 | 263 | 240 | 168 | 1 |
| 4043 | - | 4433921 | 4434076 | + | - | *PA14_49860* | hypothetical protein | 121 | 163 | 136 | 125 | 105 | 87 | 1 |
| 4044 | - | 4434649 | 4434110 | - | - | *PA14_49870* | peptide deformylase | 225 | 215 | 243 | 224 | 214 | 194 | 1 |
| 4045 | - | 4434859 | 4435431 | + | - | *PA14_49880* | hypothetical protein | 83 | 69 | 121 | 93 | 65 | 77 | 1 |
| 4046 | - | 4435431 | 4436738 | + | - | *PA14_49890* | hypothetical protein | 43 | 41 | 54 | 53 | 41 | 47 | 1 |
| 4047 | - | 4436710 | 4437216 | + | - | *PA14_49900* | hypothetical protein | 123 | 171 | 131 | 165 | 139 | 153 | 1 |
| 4048 | - | 4437362 | 4438045 | + | - | *PA14_49910* | hypothetical protein | 84 | 97 | 111 | 111 | 93 | 115 | 1 |
| 4049 | - | 4438152 | 4438526 | + | - | *PA14_49920* | hypothetical protein | 102 | 93 | 134 | 109 | 104 | 111 | 1 |
| 4050 | - | 4439375 | 4438536 | - | - | *PA14_49930* | hypothetical protein | 23 | 23 | 34 | 25 | 21 | 21 | 1 |
| 4051 | - | 4439630 | 4441954 | + | - | *PA14_49940* | hypothetical protein | 59 | 64 | 66 | 62 | 63 | 65 | 1 |
| 4052 | - | 4442058 | 4442393 | + | - | *PA14_49960* | hypothetical protein | 64 | 81 | 94 | 90 | 78 | 77 | 1 |
| 4053 | - | 4444162 | 4442396 | - | - | *PA14_49970* | ABC transporter ATP-binding protein/permease | 27 | 28 | 30 | 33 | 28 | 27 | 1 |
| 4054 | - | 4444633 | 4444824 | + | - | *PA14_49990* | hypothetical protein | 513 | 492 | 644 | 490 | 517 | 529 | 1 |
| 4055 | - | 4444925 | 4445146 | + | - | *PA14_50000* | hypothetical protein | 109 | 114 | 114 | 141 | 104 | 100 | 1 |
| 4056 | - | 4446298 | 4445150 | - | - | *PA14_50010* | dehydrogenase | 56 | 51 | 74 | 63 | 57 | 57 | 1 |
| 4057 | - | 4447014 | 4446391 | - | - | *PA14_50020* | hypothetical protein | 52 | 46 | 63 | 48 | 52 | 48 | 1 |
| 4058 | - | 4447127 | 4447837 | + | - | *PA14_50030* | hypothetical protein | 41 | 37 | 41 | 41 | 34 | 42 | 1 |
| 4059 | - | 4448621 | 4447842 | - | - | *PA14_50040* | transcriptional regulator | 50 | 45 | 49 | 50 | 42 | 46 | 1 |
| 4060 | - | 4448711 | 4449868 | + | - | *PA14_50050* | MFS family transporter | 7 | 7 | 9 | 9 | 6 | 7 | 1 |
| 4061 | - | 4449985 | 4451181 | + | - | *PA14_50060* | hypothetical protein | 27 | 27 | 32 | 29 | 30 | 30 | 1 |
| 4062 | - | 4451380 | 4452093 | + | - | *PA14_50070* | hypothetical protein | 64 | 62 | 69 | 66 | 57 | 58 | 1 |
| 4063 | - | 4452541 | 4452098 | - | *fliJ* | *PA14_50080* | flagellar biosynthesis chaperone | 70 | 79 | 79 | 86 | 72 | 71 | 1 |
| 4064 | - | 4453910 | 4452555 | - | *fliI* | *PA14_50100* | flagellum-specific ATP synthase | 131 | 144 | 138 | 139 | 145 | 120 | 1 |
| 4065 | - | 4454706 | 4453900 | - | *fliH* | *PA14_50110* | flagellar assembly protein H | 219 | 232 | 230 | 210 | 210 | 199 | 1 |
| 4066 | - | 4455724 | 4454708 | - | *fliG* | *PA14_50130* | flagellar motor switch protein G | 212 | 209 | 240 | 227 | 229 | 219 | 1 |
| 4067 | - | 4457526 | 4455730 | - | *fliF* | *PA14_50140* | flagellar MS-ring protein | 95 | 102 | 108 | 107 | 108 | 95 | 1 |
| 4068 | - | 4457878 | 4457549 | - | *fliE* | *PA14_50160* | flagellar hook-basal body protein FliE | 138 | 126 | 193 | 172 | 202 | 161 | 1 |
| 4069 | - | 4459546 | 4458125 | - | *fleR* | *PA14_50180* | two-component response regulator | 67 | 79 | 65 | 67 | 65 | 59 | 1 |
| 4070 | - | 4460759 | 4459551 | - | *fleS* | *PA14_50200* | two-component sensor | 65 | 58 | 67 | 68 | 57 | 56 | 1 |
| 4071 | - | 4462344 | 4460872 | - | *fleQ* | *PA14_50220* | transcriptional regulator FleQ | 331 | 315 | 301 | 327 | 258 | 269 | 1 |
| 4072 | - | 4462922 | 4462626 | - | - | *PA14_50240* | hypothetical protein | 276 | 324 | 313 | 194 | 327 | 283 | 1 |
| 4073 | - | 4463325 | 4462945 | - | *fliS* | *PA14_50250* | flagellar protein FliS | 699 | 689 | 996 | 491 | 776 | 699 | 1 |
| 4074 | - | 4464871 | 4463447 | - | *fliD* | *PA14_50270* | flagellar capping protein FliD | 351 | 446 | 319 | 327 | 393 | 402 | 1 |
| 4075 | - | 4465328 | 4464957 | - | - | *PA14_50280* | hypothetical protein | 196 | 326 | 173 | 160 | 250 | 189 | 1 |
| 4076 | - | 4466873 | 4465407 | - | *fliC* | *PA14_50290* | flagellin type B | 1139 | 1651 | 1035 | 879 | 1538 | 1203 | 1 |
| 4077 | - | 4472315 | 4467231 | - | - | *PA14_50300* | hypothetical protein | 226 | 215 | 232 | 242 | 240 | 227 | 1 |
| 4078 | - | 4472970 | 4472308 | - | - | *PA14_50310* | sugar nucleotidyltransferase | 167 | 149 | 175 | 177 | 167 | 184 | 1 |
| 4079 | - | 4473548 | 4472946 | - | - | *PA14_50320* | hypothetical protein | 150 | 153 | 163 | 158 | 160 | 163 | 1 |
| 4080 | - | 4474314 | 4473553 | - | - | *PA14_50330* | hypothetical protein | 199 | 182 | 228 | 213 | 218 | 228 | 1 |
| 4081 | - | 4475688 | 4474369 | - | *flgL* | *PA14_50340* | flagellar hook-associated protein FlgL | 171 | 193 | 169 | 171 | 188 | 196 | 1 |
| 4082 | - | 4477765 | 4475714 | - | *flgK* | *PA14_50360* | flagellar hook-associated protein FlgK | 167 | 178 | 156 | 155 | 164 | 165 | 1 |
| 4083 | - | 4478986 | 4477784 | - | *flgJ* | *PA14_50380* | flagellar rod assembly protein/muramidase FlgJ | 117 | 129 | 118 | 112 | 113 | 105 | 1 |
| 4084 | - | 4480106 | 4478997 | - | *flgI* | *PA14_50410* | flagellar basal body P-ring protein | 154 | 156 | 149 | 163 | 169 | 149 | 1 |
| 4085 | - | 4480813 | 4480118 | - | *flgH* | *PA14_50420* | flagellar basal body L-ring protein | 84 | 100 | 88 | 99 | 103 | 91 | 1 |
| 4086 | - | 4481644 | 4480859 | - | *flgG* | *PA14_50430* | flagellar basal body rod protein FlgG | 110 | 133 | 112 | 99 | 136 | 115 | 1 |
| 4087 | - | 4482440 | 4481691 | - | *flgF* | *PA14_50440* | flagellar basal body rod protein FlgF | 109 | 96 | 114 | 113 | 140 | 123 | 1 |
| 4088 | - | 4484045 | 4482657 | - | *flgE* | *PA14_50450* | flagellar hook protein FlgE | 148 | 207 | 138 | 118 | 189 | 144 | 1 |
| 4089 | - | 4484786 | 4484073 | - | *flgD* | *PA14_50460* | flagellar basal body rod modification protein | 110 | 162 | 132 | 123 | 153 | 134 | 1 |
| 4090 | - | 4485239 | 4484799 | - | *flgC* | *PA14_50470* | flagellar basal body rod protein FlgC | 80 | 106 | 97 | 84 | 117 | 94 | 1 |
| 4091 | - | 4485652 | 4485245 | - | *flgB* | *PA14_50480* | flagellar basal body rod protein FlgB | 86 | 99 | 100 | 99 | 107 | 104 | 1 |
| 4092 | - | 4486267 | 4485905 | - | - | *PA14_50500* | hypothetical protein | 194 | 177 | 192 | 83 | 102 | 199 | 1 |
| 4093 | - | 4486651 | 4486334 | - | - | *PA14_50510* | hypothetical protein | 54 | 40 | 57 | 43 | 35 | 38 | 1 |
| 4094 | - | 4486950 | 4488071 | + | *braC* | *PA14_50520* | branched-chain amino acid transport protein BraC | 369 | 572 | 378 | 427 | 423 | 362 | 1 |
| 4095 | - | 4488332 | 4489255 | + | *braD* | *PA14_50530* | branched-chain amino acid transport protein BraD | 41 | 79 | 48 | 43 | 48 | 44 | 0.580848337 |
| 4096 | - | 4489252 | 4490505 | + | *livM* | *PA14_50540* | leucine/isoleucine/valine transporter permease subunit | 85 | 147 | 96 | 109 | 105 | 92 | 1 |
| 4097 | - | 4490502 | 4491269 | + | *livG* | *PA14_50550* | leucine/isoleucine/valine transporter ATP-binding subunit | 108 | 169 | 111 | 135 | 134 | 129 | 1 |
| 4098 | - | 4491272 | 4491973 | + | *braG* | *PA14_50560* | branched-chain amino acid transport protein BraG | 118 | 187 | 124 | 148 | 144 | 142 | 1 |
| 4099 | - | 4494344 | 4492050 | - | - | *PA14_50570* | hypothetical protein | 93 | 82 | 75 | 66 | 90 | 66 | 1 |
| 4100 | - | 4496291 | 4494381 | - | - | *PA14_50590* | HSP90 family protein | 29 | 29 | 27 | 25 | 32 | 21 | 1 |
| 4101 | - | 4497304 | 4496390 | - | - | *PA14_50600* | transcriptional regulator | 7 | 9 | 8 | 8 | 8 | 6 | 1 |
| 4102 | - | 4497338 | 4498009 | + | - | *PA14_50610* | short chain dehydrogenase | 25 | 32 | 27 | 38 | 23 | 31 | 1 |
| 4103 | - | 4498493 | 4498020 | - | - | *PA14_50620* | hypothetical protein | 61 | 55 | 73 | 87 | 65 | 81 | 1 |
| 4104 | - | 4498513 | 4499178 | + | - | *PA14_50630* | hypothetical protein | 49 | 50 | 52 | 45 | 45 | 41 | 1 |
| 4105 | - | 4499458 | 4499207 | - | - | *PA14_50640* | hypothetical protein | 89 | 92 | 82 | 75 | 57 | 74 | 1 |
| 4106 | - | 4499976 | 4499455 | - | - | *PA14_50650* | hypothetical protein | 112 | 113 | 113 | 96 | 104 | 111 | 1 |
| 4107 | - | 4501084 | 4499987 | - | - | *PA14_50660* | hypothetical protein | 109 | 102 | 111 | 87 | 93 | 100 | 1 |
| 4108 | - | 4502000 | 4501095 | - | - | *PA14_50670* | hypothetical protein | 53 | 42 | 63 | 50 | 53 | 56 | 1 |
| 4109 | - | 4502461 | 4502114 | - | - | *PA14_50680* | monovalent cation/H+ antiporter subunit G | 86 | 107 | 99 | 108 | 122 | 122 | 1 |
| 4110 | - | 4502715 | 4502458 | - | - | *PA14_50690* | monovalent cation/H+ antiporter subunit F | 60 | 75 | 56 | 55 | 68 | 68 | 1 |
| 4111 | - | 4503236 | 4502712 | - | - | *PA14_50700* | monovalent cation/H+ antiporter subunit E | 134 | 147 | 137 | 157 | 173 | 177 | 1 |
| 4112 | - | 4504732 | 4503233 | - | - | *PA14_50710* | monovalent cation/H+ antiporter subunit D | 86 | 80 | 89 | 91 | 95 | 91 | 1 |
| 4113 | - | 4505067 | 4504729 | - | - | *PA14_50720* | monovalent cation/H+ antiporter subunit C | 101 | 103 | 90 | 115 | 122 | 116 | 1 |
| 4114 | - | 4507868 | 4505067 | - | - | *PA14_50730* | monovalent cation/H+ antiporter subunit A | 89 | 78 | 91 | 89 | 89 | 84 | 1 |
| 4115 | - | 4508661 | 4508197 | - | - | *PA14_50740* | hypothetical protein | 632 | 696 | 630 | 426 | 573 | 515 | 1 |
| 4116 | - | 4508849 | 4509091 | + | - | *PA14_50750* | hypothetical protein | 39 | 34 | 37 | 30 | 31 | 34 | 1 |

| | A | B | C | D | E | F | G | H | I | J | K | L | M | N |
|---|---|---|---|---|---|---|---|---|---|---|---|---|---|---|
| 4117 | - | 4510215 | 4509070 | - | - | *PA14_50760* | hypothetical protein | 56 | 42 | 56 | 42 | 42 | 40 | 1 |
| 4118 | - | 4511615 | 4510257 | - | - | *PA14_50770* | transporter | 6 | 8 | 8 | 6 | 6 | 6 | 1 |
| 4119 | - | 4512887 | 4511775 | - | - | *PA14_50790* | hypothetical protein | 122 | 89 | 135 | 82 | 80 | 85 | 1 |
| 4120 | - | 4513542 | 4512895 | - | *pdxH* | *PA14_50800* | pyridoxamine 5'-phosphate oxidase | 129 | 116 | 167 | 93 | 98 | 112 | 1 |
| 4121 | - | 4514522 | 4513641 | - | - | *PA14_50810* | hypothetical protein | 185 | 213 | 234 | 208 | 166 | 155 | 1 |
| 4122 | - | 4515906 | 4514728 | - | - | *PA14_50820* | esterase | 72 | 64 | 71 | 69 | 64 | 72 | 1 |
| 4123 | - | 4518256 | 4515983 | - | - | *PA14_50830* | hypothetical protein | 26 | 22 | 28 | 24 | 20 | 20 | 1 |
| 4124 | - | 4520479 | 4518335 | - | *dinG* | *PA14_50840* | ATP-dependent DNA helicase DinG | 35 | 35 | 37 | 36 | 34 | 28 | 1 |
| 4125 | - | 4520609 | 4521073 | + | - | *PA14_50850* | hypothetical protein | 47 | 52 | 52 | 55 | 49 | 50 | 1 |
| 4126 | - | 4521921 | 4521112 | - | - | *PA14_50860* | hypothetical protein | 119 | 119 | 141 | 147 | 118 | 121 | 1 |
| 4127 | - | 4522213 | 4521914 | - | - | *PA14_50870* | hypothetical protein | 119 | 128 | 147 | 174 | 123 | 132 | 1 |
| 4128 | - | 4522961 | 4522329 | - | - | *PA14_50880* | hypothetical protein | 619 | 721 | 1091 | 861 | 594 | 885 | 1 |
| 4129 | - | 4523092 | 4523589 | + | - | *PA14_50890* | hypothetical protein | 52 | 43 | 61 | 45 | 41 | 45 | 1 |
| 4130 | - | 4523592 | 4524065 | + | - | *PA14_50900* | hypothetical protein | 47 | 44 | 45 | 37 | 45 | 46 | 1 |
| 4131 | - | 4524065 | 4524382 | + | - | *PA14_50910* | hypothetical protein | 52 | 60 | 55 | 71 | 50 | 61 | 1 |
| 4132 | - | 4524465 | 4525085 | + | - | *PA14_50920* | hypothetical protein | 62 | 68 | 78 | 82 | 59 | 78 | 1 |
| 4133 | - | 4525763 | 4525203 | - | - | *PA14_50930* | hypothetical protein | 217 | 194 | 253 | 199 | 171 | 189 | 1 |
| 4134 | - | 4526078 | 4526572 | + | - | *PA14_50940* | hypothetical protein | 59 | 68 | 55 | 74 | 77 | 55 | 1 |
| 4135 | - | 4526580 | 4526780 | + | - | *PA14_50950* | hypothetical protein | 31 | 35 | 34 | 22 | 24 | 20 | 1 |
| 4136 | - | 4526947 | 4527588 | + | - | *PA14_50970* | glutathione S-transferase | 47 | 57 | 44 | 43 | 52 | 42 | 1 |
| 4137 | - | 4527711 | 4530254 | + | *pac* | *PA14_50980* | penicillin amidase | 70 | 76 | 74 | 78 | 65 | 65 | 1 |
| 4138 | - | 4531770 | 4530409 | - | - | *PA14_51000* | hypothetical protein | 78 | 80 | 81 | 86 | 69 | 68 | 1 |
| 4139 | - | 4533295 | 4532540 | - | - | *PA14_51010* | hypothetical protein | 128 | 122 | 122 | 124 | 136 | 119 | 1 |
| 4140 | - | 4533717 | 4533430 | - | - | *PA14_51020* | hypothetical protein | 163 | 180 | 139 | 138 | 161 | 151 | 1 |
| 4141 | - | 4535157 | 4533871 | - | - | *PA14_51040* | oxidoreductase | 26 | 22 | 24 | 28 | 23 | 28 | 1 |
| 4142 | - | 4536802 | 4535213 | - | - | *PA14_51050* | aldehyde dehydrogenase | 24 | 28 | 26 | 35 | 24 | 29 | 1 |
| 4143 | - | 4536881 | 4537357 | + | - | *PA14_51060* | hypothetical protein | 11 | 17 | 10 | 11 | 11 | 9 | 1 |
| 4144 | - | 4538984 | 4537734 | - | - | *PA14_51070* | porin | 5 | 5 | 5 | 4 | 4 | 3 | 1 |
| 4145 | - | 4540049 | 4539063 | - | - | *PA14_51080* | dioxygenase | 12 | 12 | 11 | 11 | 11 | 11 | 1 |
| 4146 | - | 4540982 | 4540065 | - | - | *PA14_51090* | short-chain dehydrogenase | 14 | 13 | 9 | 9 | 12 | 10 | 1 |
| 4147 | - | 4542168 | 4541023 | - | - | *PA14_51100* | acyl-CoA dehydrogenase | 12 | 11 | 9 | 10 | 9 | 10 | 1 |
| 4148 | - | 4542931 | 4542170 | - | - | *PA14_51110* | enoyl-CoA hydratase/isomerase | 16 | 13 | 13 | 12 | 12 | 15 | 1 |
| 4149 | - | 4544043 | 4542931 | - | - | *PA14_51120* | acyl-CoA dehydrogenase | 15 | 12 | 11 | 9 | 11 | 11 | 1 |
| 4150 | - | 4545922 | 4544567 | - | *mucK* | *PA14_51150* | cis,cis-muconate transporter MucK | 7 | 6 | 8 | 6 | 7 | 8 | 1 |
| 4151 | - | 4546870 | 4546004 | - | - | *PA14_51160* | hypothetical protein | 15 | 14 | 14 | 15 | 14 | 15 | 1 |
| 4152 | - | 4549001 | 4546854 | - | *pauA* | *PA14_51170* | pimeloyl-CoA synthetase | 20 | 16 | 17 | 16 | 13 | 15 | 1 |
| 4153 | - | 4550262 | 4549111 | - | - | *PA14_51190* | thiolase | 28 | 28 | 26 | 24 | 23 | 28 | 1 |
| 4154 | - | 4550645 | 4550259 | - | - | *PA14_51200* | hypothetical protein | 39 | 34 | 33 | 35 | 34 | 40 | 1 |
| 4155 | - | 4550803 | 4551603 | + | - | *PA14_51205* | transcriptional regulator | 74 | 75 | 80 | 75 | 92 | 81 | 1 |
| 4156 | - | 4552635 | 4551643 | - | - | *PA14_51220* | glycosyl transferase family protein | 77 | 76 | 83 | 79 | 83 | 75 | 1 |
| 4157 | - | 4553064 | 4552978 | - | - | *PA14_51230* | Ser tRNA | 258 | 234 | 308 | 224 | 128 | 108 | 1 |
| 4158 | - | 4553868 | 4553158 | - | *purC* | *PA14_51240* | phosphoribosylaminoimidazole-succinocarboxamide synthase | 158 | 214 | 175 | 142 | 148 | 129 | 1 |
| 4159 | - | 4554655 | 4553897 | - | - | *PA14_51250* | hypothetical protein | 137 | 148 | 154 | 155 | 132 | 118 | 1 |
| 4160 | - | 4555784 | 4554594 | - | - | *PA14_51260* | hypothetical protein | 207 | 237 | 214 | 213 | 193 | 167 | 1 |
| 4161 | - | 4556680 | 4555802 | - | *dapA* | *PA14_51270* | dihydrodipicolinate synthase | 273 | 204 | 286 | 272 | 263 | 238 | 1 |
| 4162 | - | 4556864 | 4557421 | + | - | *PA14_51280* | hypothetical protein | 51 | 37 | 51 | 35 | 37 | 34 | 1 |
| 4163 | - | 4557433 | 4557906 | + | *bcp* | *PA14_51290* | bacterioferritin comigratory protein | 48 | 67 | 44 | 41 | 47 | 38 | 1 |
| 4164 | - | 4559040 | 4557967 | - | - | *PA14_51300* | hypothetical protein | 25 | 23 | 25 | 25 | 21 | 19 | 1 |
| 4165 | - | 4559325 | 4559074 | - | - | *PA14_51310* | hypothetical protein | 33 | 22 | 37 | 29 | 25 | 27 | 0.844537532 |
| 4166 | - | 4559488 | 4560921 | + | - | *PA14_51320* | hypothetical protein | 55 | 60 | 56 | 62 | 51 | 54 | 1 |
| 4167 | - | 4562088 | 4561030 | - | *nadA* | *PA14_51330* | quinolinate synthetase | 127 | 104 | 139 | 109 | 106 | 120 | 1 |
| 4168 | - | 4562837 | 4563835 | + | *mvfR* | *PA14_51340* | transcriptional regulator MvfR | 165 | 242 | 211 | 215 | 238 | 189 | 1 |
| 4169 | - | 4564390 | 4563788 | - | *phnB* | *PA14_51350* | anthranilate synthase component II | 663 | 1014 | 762 | 780 | 937 | 557 | 1 |
| 4170 | - | 4565960 | 4564368 | - | *phnA* | *PA14_51360* | anthranilate synthase component I | 1143 | 1767 | 1670 | 1464 | 1616 | 905 | 1 |
| 4171 | - | 4566982 | 4566077 | - | *pqsE* | *PA14_51380* | quinolone signal response protein | 1139 | 1967 | 1680 | 448 | 924 | 894 | 1 |
| 4172 | - | 4567989 | 4566976 | - | *pqsD* | *PA14_51390* | 3-oxoacyl-ACP synthase | 1222 | 2281 | 1461 | 1289 | 1861 | 955 | 1 |
| 4173 | - | 4569078 | 4568032 | - | *pqsC* | *PA14_51410* | PqsC | 865 | 1715 | 945 | 1022 | 1419 | 676 | 1 |
| 4174 | - | 4569922 | 4569071 | - | *pqsB* | *PA14_51420* | PqsB | 886 | 1902 | 1018 | 1070 | 1481 | 675 | 1 |
| 4175 | - | 4571469 | 4569916 | - | *pqsA* | *PA14_51430* | coenzyme A ligase | 379 | 716 | 512 | 504 | 629 | 261 | 1 |
| 4176 | - | 4572028 | 4572549 | + | *ogt* | *PA14_51440* | methylated-DNA--protein-cysteine methyltransferase | 41 | 36 | 51 | 35 | 36 | 36 | 1 |
| 4177 | - | 4575148 | 4572629 | - | *cupC3* | *PA14_51450* | usher CupC3 | 32 | 32 | 34 | 48 | 27 | 28 | 1 |
| 4178 | - | 4575957 | 4575244 | - | *cupC2* | *PA14_51460* | chaperone CupC | 1 | 0 | 0 | 0 | 0 | 0 | 0.162072635 |
| 4179 | - | 4576632 | 4576015 | - | *cupC1* | *PA14_51470* | fimbrial subunit CupC1 | 6 | 5 | 4 | 4 | 5 | 4 | 1 |
| 4180 | - | 4577454 | 4577077 | - | - | *PA14_51480* | hypothetical protein | 22 | 31 | 21 | 19 | 25 | 22 | 1 |
| 4181 | - | 4578053 | 4578691 | + | - | *PA14_51490* | hypothetical protein | 27 | 26 | 25 | 26 | 24 | 25 | 1 |
| 4182 | - | 4579069 | 4579629 | + | - | *PA14_51500* | hypothetical protein | 19 | 21 | 21 | 23 | 18 | 19 | 1 |
| 4183 | - | 4580154 | 4579750 | - | - | *PA14_51510* | hypothetical protein | 72 | 60 | 63 | 63 | 59 | 58 | 1 |
| 4184 | - | 4580960 | 4580547 | - | *spcU* | *PA14_51520* | SpcU | 258 | 250 | 234 | 256 | 253 | 262 | 1 |
| 4185 | - | 4583020 | 4580957 | - | *exoU* | *PA14_51530* | ExoU | 203 | 203 | 210 | 211 | 190 | 193 | 1 |
| 4186 | - | 4583726 | 4583157 | - | - | *PA14_51540* | transposase | 177 | 174 | 210 | 185 | 132 | 141 | 1 |
| 4187 | - | 4583761 | 4584861 | + | - | *PA14_51550* | transposase | 34 | 33 | 34 | 41 | 28 | 32 | 1 |
| 4188 | - | 4585439 | 4584990 | - | - | *PA14_51560* | acetyltransferase | 95 | 78 | 75 | 60 | 73 | 72 | 1 |
| 4189 | - | 4585871 | 4585653 | - | - | *PA14_51570* | hypothetical protein | 603 | 561 | 496 | 530 | 693 | 626 | 1 |
| 4190 | - | 4586130 | 4585927 | - | - | *PA14_51580* | hypothetical protein | 3 | 2 | 1 | 3 | 2 | 2 | 1 |
| 4191 | - | 4586242 | 4586559 | + | - | *PA14_51590* | hypothetical protein | 83 | 93 | 82 | 57 | 53 | 76 | 1 |
| 4192 | - | 4587288 | 4586665 | - | - | *PA14_51600* | hypothetical protein | 24 | 25 | 21 | 24 | 25 | 23 | 1 |
| 4193 | - | 4587609 | 4587325 | - | - | *PA14_51610* | hypothetical protein | 10 | 9 | 8 | 6 | 10 | 10 | 1 |
| 4194 | - | 4587841 | 4588149 | + | - | *PA14_51620* | transposase | 299 | 288 | 296 | 301 | 272 | 283 | 1 |
| 4195 | - | 4588146 | 4588991 | + | - | *PA14_51630* | transposase | 79 | 73 | 76 | 75 | 71 | 69 | 1 |
| 4196 | - | 4589078 | 4589455 | + | - | *PA14_51640* | hypothetical protein | 7 | 7 | 6 | 6 | 6 | 7 | 1 |
| 4197 | - | 4589452 | 4590732 | + | - | *PA14_51650* | integrase | 8 | 7 | 6 | 7 | 7 | 6 | 1 |
| 4198 | - | 4591178 | 4591106 | - | - | *PA14_51660* | Lys tRNA | 85 | 117 | 123 | 58 | 46 | 112 | 1 |
| 4199 | - | 4591912 | 4591238 | - | - | *PA14_51670* | hypothetical protein | 30 | 29 | 32 | 28 | 24 | 31 | 1 |
| 4200 | - | 4592722 | 4591928 | - | - | *PA14_51680* | radical activating enzyme | 20 | 20 | 20 | 17 | 16 | 20 | 1 |

| | A | B | C | D | E | F | G | H | I | J | K | L | M | N |
|---|---|---|---|---|---|---|---|---|---|---|---|---|---|---|
| 4201 | - | 4593619 | 4592795 | - | - | PA14_51690 | hypothetical protein | 165 | 172 | 145 | 126 | 125 | 113 | 1 |
| 4202 | - | 4594135 | 4593629 | - | oprL | PA14_51710 | peptidoglycan associated lipoprotein OprL precursor | 1620 | 1837 | 1313 | 1101 | 1454 | 1010 | 1 |
| 4203 | - | 4595486 | 4594188 | - | tolB | PA14_51720 | translocation protein TolB | 223 | 253 | 211 | 195 | 209 | 189 | 1 |
| 4204 | - | 4596526 | 4595483 | - | tolA | PA14_51730 | TolA protein | 256 | 215 | 288 | 259 | 270 | 245 | 1 |
| 4205 | - | 4596969 | 4596529 | - | tolR | PA14_51740 | TolR protein | 200 | 232 | 180 | 204 | 222 | 187 | 1 |
| 4206 | - | 4597687 | 4596992 | - | tolQ | PA14_51750 | TolQ protein | 212 | 237 | 204 | 198 | 212 | 190 | 1 |
| 4207 | - | 4598135 | 4597689 | - | - | PA14_51770 | hypothetical protein | 361 | 307 | 349 | 365 | 407 | 340 | 1 |
| 4208 | - | 4599246 | 4598188 | - | ruvB | PA14_51780 | Holliday junction DNA helicase RuvB | 133 | 119 | 137 | 131 | 136 | 125 | 1 |
| 4209 | - | 4599862 | 4599257 | - | ruvA | PA14_51790 | Holliday junction DNA helicase RuvA | 92 | 87 | 97 | 99 | 96 | 92 | 1 |
| 4210 | - | 4600403 | 4599879 | - | ruvC | PA14_51800 | Holliday junction resolvase | 112 | 92 | 117 | 88 | 96 | 96 | 1 |
| 4211 | - | 4601234 | 4600488 | - | - | PA14_51810 | hypothetical protein | 391 | 415 | 364 | 385 | 304 | 275 | 1 |
| 4212 | - | 4603076 | 4601301 | - | aspS | PA14_51820 | aspartyl-tRNA synthetase | 184 | 180 | 199 | 189 | 140 | 135 | 1 |
| 4213 | - | 4603516 | 4603986 | + | - | PA14_51830 | DNA-binding stress protein | 617 | 525 | 317 | 185 | 375 | 637 | 1 |
| 4214 | - | 4604200 | 4604814 | + | - | PA14_51840 | cold-shock protein | 23 | 19 | 17 | 15 | 14 | 15 | 1 |
| 4215 | - | 4605072 | 4604863 | - | - | PA14_51850 | hypothetical protein | 52 | 60 | 63 | 58 | 64 | 52 | 1 |
| 4216 | - | 4605702 | 4605073 | - | - | PA14_51860 | hypothetical protein | 110 | 122 | 134 | 123 | 124 | 123 | 1 |
| 4217 | - | 4606220 | 4607551 | + | oprD | PA14_51880 | basic amino acid, basic peptide and impenem outer membrane porin OprD precursor | 589 | 1219 | 451 | 391 | 741 | 431 | 1 |
| 4218 | - | 4608022 | 4607615 | - | - | PA14_51890 | hypothetical protein | 49 | 51 | 52 | 43 | 42 | 42 | 1 |
| 4219 | - | 4608130 | 4609845 | + | proS | PA14_51900 | prolyl-tRNA synthetase | 88 | 98 | 82 | 84 | 80 | 71 | 1 |
| 4220 | - | 4609854 | 4610810 | + | - | PA14_51910 | hypothetical protein | 73 | 82 | 74 | 78 | 75 | 73 | 1 |
| 4221 | - | 4611102 | 4610827 | - | - | PA14_51920 | acylphosphatase | 108 | 105 | 105 | 155 | 124 | 119 | 1 |
| 4222 | - | 4611566 | 4611102 | - | - | PA14_51930 | thioredoxin | 122 | 108 | 131 | 151 | 126 | 135 | 1 |
| 4223 | - | 4611911 | 4612294 | + | - | PA14_51940 | hypothetical protein | 85 | 124 | 135 | 97 | 90 | 70 | 1 |
| 4224 | - | 4612499 | 4612305 | - | - | PA14_51950 | hypothetical protein | 251 | 321 | 394 | 330 | 367 | 286 | 1 |
| 4225 | - | 4613884 | 4612649 | - | - | PA14_51960 | ribonuclease | 51 | 54 | 56 | 60 | 43 | 47 | 1 |
| 4226 | - | 4614009 | 4614362 | + | - | PA14_51980 | arsenate reductase | 59 | 60 | 71 | 48 | 50 | 46 | 1 |
| 4227 | - | 4614359 | 4614955 | + | wrbA | PA14_51990 | Trp repressor binding protein WrbA | 44 | 51 | 53 | 46 | 44 | 44 | 1 |
| 4228 | - | 4614957 | 4615376 | + | - | PA14_52000 | hypothetical protein | 40 | 42 | 41 | 46 | 43 | 32 | 1 |
| 4229 | - | 4616280 | 4615576 | - | - | PA14_52010 | DNA replication initiation factor | 32 | 25 | 32 | 24 | 21 | 24 | 1 |
| 4230 | - | 4617519 | 4616482 | - | - | PA14_52020 | hypothetical protein | 86 | 88 | 86 | 82 | 75 | 69 | 1 |
| 4231 | - | 4617733 | 4618794 | + | purM | PA14_52040 | phosphoribosylaminoimidazole synthetase | 63 | 70 | 75 | 60 | 52 | 42 | 1 |
| 4232 | - | 4618794 | 4619462 | + | purN | PA14_52050 | phosphoribosylglycinamide formyltransferase | 71 | 81 | 86 | 61 | 67 | 50 | 1 |
| 4233 | - | 4619455 | 4620171 | + | - | PA14_52060 | hypothetical protein | 123 | 118 | 104 | 80 | 97 | 85 | 1 |
| 4234 | - | 4620785 | 4620225 | - | - | PA14_52070 | transcriptional regulator | 109 | 117 | 109 | 108 | 109 | 98 | 1 |
| 4235 | - | 4620901 | 4621134 | + | - | PA14_52080 | hypothetical protein | 41 | 54 | 41 | 31 | 35 | 40 | 1 |
| 4236 | - | 4621146 | 4621400 | + | - | PA14_52090 | hypothetical protein | 63 | 73 | 70 | 60 | 54 | 61 | 1 |
| 4237 | - | 4621406 | 4621732 | + | - | PA14_52110 | hypothetical protein | 72 | 86 | 95 | 83 | 69 | 69 | 1 |
| 4238 | - | 4621825 | 4622016 | + | - | PA14_52120 | hypothetical protein | 92 | 58 | 55 | 78 | 90 | 101 | 1 |
| 4239 | - | 4623385 | 4622057 | - | - | PA14_52130 | hypothetical protein | 216 | 88 | 126 | 159 | 153 | 211 | 0.885197681 |
| 4240 | - | 4624112 | 4623573 | - | - | PA14_52140 | hypothetical protein | 35 | 33 | 40 | 35 | 32 | 27 | 1 |
| 4241 | - | 4624345 | 4625283 | + | lpxO2 | PA14_52150 | lipopolysaccharide biosynthetic protein LpxO2 | 73 | 60 | 75 | 60 | 60 | 62 | 1 |
| 4242 | - | 4626156 | 4625326 | - | mazG | PA14_52160 | nucleoside triphosphate pyrophosphohydrolase | 55 | 61 | 60 | 50 | 56 | 55 | 1 |
| 4243 | - | 4628504 | 4626261 | - | relA | PA14_52180 | GTP pyrophosphokinase | 291 | 288 | 277 | 307 | 283 | 275 | 1 |
| 4244 | - | 4629941 | 4628589 | - | rumA | PA14_52190 | 23S rRNA 5-methyluridine methyltransferase | 140 | 127 | 153 | 139 | 125 | 118 | 1 |
| 4245 | - | 4630849 | 4629950 | - | cysM | PA14_52210 | cysteine synthase B | 129 | 104 | 136 | 114 | 109 | 113 | 1 |
| 4246 | - | 4633327 | 4631099 | - | - | PA14_52230 | outer membrane receptor FepA | 8 | 9 | 10 | 13 | 10 | 9 | 1 |
| 4247 | - | 4634900 | 4633563 | - | - | PA14_52240 | two-component sensor | 20 | 33 | 30 | 69 | 52 | 28 | 0.391278558 |
| 4248 | - | 4635619 | 4634900 | - | - | PA14_52250 | two-component response regulator | 15 | 41 | 21 | 86 | 130 | 26 | 0.022359242 |
| 4249 | - | 4635805 | 4638582 | + | - | PA14_52260 | sensor/response regulator hybrid | 64 | 55 | 64 | 69 | 56 | 67 | 1 |
| 4250 | - | 4638585 | 4639574 | + | ldhA | PA14_52270 | D-lactate dehydrogenase | 41 | 47 | 46 | 50 | 49 | 53 | 1 |
| 4251 | - | 4639610 | 4640035 | + | - | PA14_52280 | hypothetical protein | 51 | 56 | 65 | 65 | 55 | 61 | 1 |
| 4252 | - | 4640103 | 4640510 | + | - | PA14_52290 | hypothetical protein | 41 | 45 | 54 | 55 | 34 | 52 | 1 |
| 4253 | - | 4640623 | 4642551 | + | - | PA14_52300 | hypothetical protein | 11 | 13 | 15 | 16 | 12 | 13 | 1 |
| 4254 | - | 4643607 | 4642558 | - | dinP | PA14_52310 | DNA polymerase IV | 62 | 51 | 61 | 60 | 51 | 49 | 1 |
| 4255 | - | 4643754 | 4643827 | + | - | PA14_52320 | Met tRNA | 49 | 33 | 56 | 29 | 11 | 18 | 1 |
| 4256 | - | 4644009 | 4644323 | + | - | PA14_52330 | hypothetical protein | 101 | 111 | 95 | 100 | 113 | 87 | 1 |
| 4257 | - | 4644697 | 4644338 | - | - | PA14_52340 | hypothetical protein | 244 | 231 | 236 | 274 | 351 | 272 | 1 |
| 4258 | - | 4644946 | 4647591 | + | - | PA14_52350 | hypothetical protein | 39 | 36 | 37 | 34 | 40 | 38 | 1 |
| 4259 | - | 4647593 | 4648876 | + | - | PA14_52370 | hypothetical protein | 91 | 81 | 96 | 88 | 94 | 93 | 1 |
| 4260 | - | 4649035 | 4649583 | + | - | PA14_52380 | cytochrome b561 | 32 | 37 | 31 | 27 | 26 | 34 | 1 |
| 4261 | - | 4651542 | 4649638 | - | kup | PA14_52400 | potassium uptake protein Kup | 76 | 73 | 92 | 104 | 78 | 85 | 1 |
| 4262 | - | 4651789 | 4653111 | + | rimO | PA14_52420 | ribosomal protein S12 methylthiotransferase | 21 | 21 | 23 | 26 | 20 | 24 | 1 |
| 4263 | - | 4653174 | 4653635 | + | - | PA14_52430 | hypothetical protein | 8 | 8 | 6 | 10 | 10 | 6 | 1 |
| 4264 | - | 4653628 | 4654020 | + | - | PA14_52440 | hypothetical protein | 13 | 15 | 14 | 17 | 22 | 16 | 1 |
| 4265 | - | 4655518 | 4654070 | - | mgtE | PA14_52460 | Mg transporter MgtE | 35 | 36 | 38 | 35 | 43 | 39 | 1 |
| 4266 | - | 4655845 | 4656360 | + | - | PA14_52465 | hypothetical protein | 20 | 30 | 19 | 32 | 37 | 23 | 1 |
| 4267 | - | 4656891 | 4656406 | - | - | PA14_52480 | hypothetical protein | 60 | 116 | 74 | 118 | 131 | 80 | 0.452109894 |
| 4268 | - | 4657447 | 4656944 | - | - | PA14_52490 | hypothetical protein | 55 | 97 | 64 | 94 | 98 | 65 | 0.676260136 |
| 4269 | - | 4657814 | 4657539 | - | - | PA14_52500 | hypothetical protein | 45 | 60 | 40 | 87 | 89 | 49 | 1 |
| 4270 | - | 4658181 | 4657807 | - | - | PA14_52510 | hypothetical protein | 22 | 39 | 27 | 46 | 41 | 23 | 0.85148132 |
| 4271 | - | 4658876 | 4658346 | - | - | PA14_52520 | hypothetical protein | 48 | 56 | 58 | 57 | 63 | 55 | 1 |
| 4272 | - | 4659047 | 4659760 | + | - | PA14_52530 | transcriptional regulator | 20 | 22 | 21 | 19 | 17 | 21 | 1 |
| 4273 | - | 4659935 | 4659862 | - | - | PA14_52540 | Arg tRNA | 63 | 45 | 73 | 53 | 45 | 38 | 1 |
| 4274 | - | 4660123 | 4660050 | - | - | PA14_52550 | Arg tRNA | 152 | 98 | 165 | 101 | 103 | 85 | 0.723542393 |
| 4275 | - | 4660331 | 4660244 | - | - | PA14_52560 | Ser tRNA | 507 | 386 | 625 | 341 | 397 | 324 | 1 |
| 4276 | - | 4660578 | 4660393 | - | rsmA | PA14_52570 | carbon storage regulator | 342 | 420 | 439 | 275 | 345 | 309 | 1 |
| 4277 | - | 4662001 | 4660763 | - | lysC | PA14_52580 | aspartate kinase | 119 | 127 | 109 | 100 | 123 | 99 | 1 |
| 4278 | - | 4664773 | 4662149 | - | alaS | PA14_52600 | alanyl-tRNA synthetase | 73 | 83 | 76 | 62 | 74 | 61 | 1 |
| 4279 | - | 4665899 | 4664895 | - | - | PA14_52610 | hypothetical protein | 33 | 28 | 35 | 28 | 28 | 25 | 1 |
| 4280 | - | 4667042 | 4666044 | - | aruE | PA14_52630 | succinylglutamate desuccinylase | 304 | 326 | 355 | 366 | 285 | 296 | 1 |
| 4281 | - | 4667346 | 4667056 | - | - | PA14_52640 | hypothetical protein | 268 | 355 | 271 | 292 | 333 | 319 | 1 |
| 4282 | - | 4668706 | 4667360 | - | aruB | PA14_52660 | succinylarginine dihydrolase | 316 | 401 | 305 | 356 | 394 | 360 | 1 |
| 4283 | - | 4670169 | 4668703 | - | astD | PA14_52670 | succinylglutamic semialdehyde dehydrogenase | 259 | 299 | 257 | 270 | 286 | 266 | 1 |
| 4284 | - | 4671234 | 4670212 | - | aruG | PA14_52690 | arginine/ornithine succinyltransferase AII subunit | 146 | 175 | 136 | 161 | 150 | 153 | 1 |

| | A | B | C | D | E | F | G | H | I | J | K | L | M | N |
|---|---|---|---|---|---|---|---|---|---|---|---|---|---|---|
| 4285 | - | 4672262 | 4671246 | - | *aruF* | *PA14_52700* | arginine/ornithine succinyltransferase AI subunit | 229 | 214 | 224 | 243 | 199 | 220 | 1 |
| 4286 | - | 4673681 | 4672461 | - | *argD* | *PA14_52720* | bifunctional N-succinyldiaminopimelate-aminotransferase/acetylornithine transaminase | 459 | 572 | 485 | 641 | 558 | 487 | 1 |
| 4287 | - | 4673848 | 4674171 | + | - | *PA14_52730* | hypothetical protein | 76 | 69 | 89 | 90 | 76 | 80 | 1 |
| 4288 | - | 4675181 | 4674192 | - | *argR* | *PA14_52740* | transcriptional regulator ArgR | 115 | 117 | 129 | 110 | 122 | 124 | 1 |
| 4289 | - | 4675978 | 4675214 | - | *aotP* | *PA14_52750* | arginine/ornithine transport protein AotP | 223 | 243 | 264 | 253 | 268 | 279 | 1 |
| 4290 | - | 4677109 | 4675997 | - | - | *PA14_52760* | hypothetical protein | 91 | 110 | 112 | 111 | 116 | 116 | 1 |
| 4291 | - | 4677821 | 4677123 | - | *aotM* | *PA14_52770* | arginine/ornithine transport protein AotM | 66 | 72 | 79 | 74 | 67 | 78 | 1 |
| 4292 | - | 4678527 | 4677838 | - | *aotQ* | *PA14_52780* | arginine/ornithine transport protein AotQ | 61 | 56 | 69 | 63 | 62 | 72 | 1 |
| 4293 | - | 4679425 | 4678646 | - | *aotJ* | *PA14_52790* | arginine/ornithine binding protein AotJ | 118 | 175 | 184 | 144 | 167 | 158 | 1 |
| 4294 | - | 4681921 | 4679966 | - | *acsA* | *PA14_52800* | acetyl-CoA synthetase | 901 | 558 | 744 | 515 | 405 | 405 | 1 |
| 4295 | - | 4683386 | 4682103 | - | - | *PA14_52810* | C4-dicarboxylate transporter | 14 | 11 | 14 | 15 | 12 | 12 | 1 |
| 4296 | - | 4684021 | 4683383 | - | - | *PA14_52820* | C4-dicarboxylate transporter | 21 | 18 | 23 | 21 | 12 | 16 | 1 |
| 4297 | - | 4685065 | 4684070 | - | - | *PA14_52840* | C4-dicarboxylate-binding periplasmic protein | 10 | 11 | 12 | 12 | 8 | 12 | 1 |
| 4298 | - | 4686006 | 4685179 | - | - | *PA14_52850* | acyl-CoA lyase subunit beta | 6 | 4 | 5 | 4 | 4 | 6 | 0.978560082 |
| 4299 | - | 4687219 | 4686017 | - | - | *PA14_52870* | hypothetical protein | 4 | 4 | 3 | 3 | 3 | 3 | 1 |
| 4300 | - | 4688627 | 4687275 | - | - | *PA14_52880* | hypothetical protein | 2 | 2 | 3 | 2 | 2 | 2 | 1 |
| 4301 | - | 4689046 | 4688666 | - | - | *PA14_52890* | ring-cleaving dioxygenase | 2 | 1 | 2 | 2 | 2 | 2 | 0.956402005 |
| 4302 | - | 4690221 | 4689061 | - | - | *PA14_52900* | acyl-CoA dehydrogenase | 9 | 8 | 8 | 9 | 6 | 6 | 1 |
| 4303 | - | 4691094 | 4690267 | - | - | *PA14_52910* | hypothetical protein | 5 | 3 | 5 | 3 | 4 | 3 | 1 |
| 4304 | - | 4691211 | 4692107 | + | - | *PA14_52920* | transcriptional regulator | 27 | 29 | 28 | 29 | 28 | 31 | 1 |
| 4305 | - | 4693185 | 4692241 | - | - | *PA14_52930* | transcriptional regulator | 45 | 38 | 47 | 48 | 39 | 40 | 1 |
| 4306 | - | 4693405 | 4695600 | + | - | *PA14_52940* | hypothetical protein | 22 | 28 | 26 | 39 | 26 | 28 | 1 |
| 4307 | - | 4695869 | 4696135 | + | - | *PA14_52960* | hypothetical protein | 24 | 27 | 23 | 26 | 19 | 22 | 1 |
| 4308 | - | 4697912 | 4696353 | - | *phhR* | *PA14_52980* | transcriptional regulator PhhR | 85 | 82 | 76 | 102 | 78 | 76 | 1 |
| 4309 | - | 4698197 | 4698985 | + | *phhA* | *PA14_52990* | phenylalanine 4-monooxygenase | 486 | 896 | 598 | 756 | 808 | 861 | 1 |
| 4310 | - | 4699090 | 4699446 | + | *phhB* | *PA14_53000* | pterin-4-alpha-carbinolamine dehydratase | 319 | 637 | 373 | 362 | 408 | 447 | 1 |
| 4311 | - | 4699443 | 4700642 | + | *phhC* | *PA14_53010* | aromatic amino acid aminotransferase | 308 | 506 | 315 | 359 | 398 | 398 | 1 |
| 4312 | - | 4700957 | 4701889 | + | *pbpG* | *PA14_53020* | D-alanyl-D-alanine endopeptidase | 76 | 55 | 69 | 53 | 66 | 62 | 1 |
| 4313 | - | 4702325 | 4701912 | - | - | *PA14_53030* | peptidyl-tRNA hydrolase domain-containing protein | 94 | 84 | 92 | 80 | 78 | 71 | 1 |
| 4314 | - | 4702829 | 4702446 | - | - | *PA14_53040* | lysozyme inhibitor | 510 | 647 | 375 | 299 | 392 | 308 | 1 |
| 4315 | - | 4704400 | 4702982 | - | *aroP2* | *PA14_53050* | aromatic amino acid transport protein AroP2 | 25 | 22 | 23 | 23 | 17 | 27 | 1 |
| 4316 | - | 4705771 | 4704698 | - | *hpd* | *PA14_53070* | 4-hydroxyphenylpyruvate dioxygenase | 336 | 610 | 269 | 435 | 500 | 557 | 1 |
| 4317 | - | 4706093 | 4706881 | + | - | *PA14_53090* | transcriptional regulator | 35 | 32 | 33 | 34 | 40 | 42 | 1 |
| 4318 | - | 4706899 | 4707138 | + | - | *PA14_53100* | hypothetical protein | 35 | 39 | 32 | 28 | 49 | 50 | 1 |
| 4319 | - | 4707164 | 4708123 | + | - | *PA14_53110* | oxidoreductase | 41 | 45 | 37 | 38 | 48 | 43 | 1 |
| 4320 | - | 4708956 | 4708174 | - | - | *PA14_53120* | hypothetical protein | 292 | 223 | 238 | 209 | 252 | 264 | 1 |
| 4321 | - | 4711502 | 4709031 | - | - | *PA14_53140* | hypothetical protein | 109 | 108 | 118 | 128 | 120 | 137 | 1 |
| 4322 | - | 4713571 | 4711781 | - | - | *PA14_53150* | ABC transporter ATP-binding protein/permease | 92 | 82 | 100 | 111 | 94 | 98 | 1 |
| 4323 | - | 4714166 | 4713564 | - | - | *PA14_53160* | hypothetical protein | 146 | 120 | 137 | 150 | 113 | 137 | 1 |
| 4324 | - | 4715144 | 4714206 | - | - | *PA14_53180* | hypothetical protein | 76 | 60 | 75 | 70 | 51 | 57 | 1 |
| 4325 | - | 4715614 | 4715309 | - | *bolA* | *PA14_53190* | morphogene protein BolA | 161 | 148 | 143 | 141 | 164 | 134 | 1 |
| 4326 | - | 4716176 | 4715628 | - | - | *PA14_53200* | hypothetical protein | 140 | 142 | 160 | 112 | 137 | 130 | 1 |
| 4327 | - | 4716344 | 4717363 | + | - | *PA14_53210* | hypothetical protein | 24 | 27 | 31 | 32 | 24 | 20 | 1 |
| 4328 | - | 4717492 | 4718886 | + | *fumC2* | *PA14_53220* | fumarate hydratase | 95 | 101 | 83 | 84 | 103 | 93 | 1 |
| 4329 | - | 4718879 | 4719502 | + | - | *PA14_53230* | oxidoreductase | 56 | 61 | 53 | 52 | 61 | 62 | 1 |
| 4330 | - | 4719782 | 4720951 | + | *cpbD* | *PA14_53250* | chitin-binding protein CbpD | 764 | 142 | 114 | 306 | 306 | 773 | 0.000552202 |
| 4331 | - | 4721128 | 4722090 | + | - | *PA14_53260* | hypothetical protein | 47 | 52 | 48 | 43 | 50 | 50 | 1 |
| 4332 | - | 4722520 | 4722101 | - | - | *PA14_53270* | hypothetical protein | 32 | 34 | 32 | 28 | 31 | 30 | 1 |
| 4333 | - | 4723724 | 4722774 | - | *trxB2* | *PA14_53290* | thioredoxin reductase 2 | 289 | 62 | 85 | 74 | 43 | 263 | 0.000160291 |
| 4334 | - | 4724457 | 4723858 | - | - | *PA14_53300* | alkyl hydroperoxide reductase | 371 | 26 | 18 | 23 | 16 | 257 | 1.6225E-146 |
| 4335 | - | 4724553 | 4726598 | + | - | *PA14_53310* | hypothetical protein | 24 | 25 | 31 | 31 | 25 | 29 | 1 |
| 4336 | - | 4726683 | 4727423 | + | - | *PA14_53330* | sulfate transport protein CysZ | 63 | 63 | 66 | 63 | 58 | 55 | 1 |
| 4337 | - | 4727802 | 4729814 | + | - | *PA14_53340* | hypothetical protein | 14 | 10 | 15 | 11 | 11 | 13 | 1 |
| 4338 | - | 4730156 | 4732348 | + | *plcH* | *PA14_53360* | hemolytic phospholipase C | 5 | 6 | 6 | 7 | 5 | 5 | 1 |
| 4339 | - | 4732365 | 4732988 | + | *plcR* | *PA14_53370* | phospholipase accessory protein PlcR | 6 | 4 | 6 | 6 | 4 | 5 | 1 |
| 4340 | - | 4733076 | 4734296 | + | - | *PA14_53380* | glycosyl transferase family protein | 11 | 11 | 12 | 10 | 9 | 11 | 1 |
| 4341 | - | 4735259 | 4734300 | - | - | *PA14_53390* | hypothetical protein | 16 | 25 | 18 | 18 | 22 | 17 | 1 |
| 4342 | - | 4736562 | 4735450 | - | - | *PA14_53400* | oxidoreductase | 22 | 20 | 19 | 17 | 22 | 21 | 1 |
| 4343 | - | 4737193 | 4736603 | - | - | *PA14_53410* | transcriptional regulator | 14 | 12 | 16 | 13 | 19 | 16 | 1 |
| 4344 | - | 4737826 | 4737344 | - | - | *PA14_53420* | glutathione peroxidase | 242 | 201 | 246 | 188 | 233 | 232 | 1 |
| 4345 | - | 4738517 | 4738032 | - | *slyD* | *PA14_53430* | peptidyl-prolyl cis-trans isomerase SlyD | 662 | 541 | 796 | 548 | 541 | 572 | 1 |
| 4346 | - | 4738624 | 4738824 | + | - | *PA14_53450* | hypothetical protein | 241 | 236 | 259 | 141 | 190 | 259 | 1 |
| 4347 | - | 4738824 | 4740008 | + | - | *PA14_53470* | acetate kinase | 306 | 278 | 327 | 173 | 246 | 316 | 1 |
| 4348 | - | 4740071 | 4742185 | + | *pta* | *PA14_53480* | phosphate acetyltransferase | 226 | 243 | 248 | 166 | 212 | 238 | 1 |
| 4349 | - | 4742312 | 4743226 | + | - | *PA14_53490* | acyltransferase | 25 | 25 | 34 | 23 | 24 | 27 | 1 |
| 4350 | - | 4744009 | 4743296 | - | - | *PA14_53500* | hypothetical protein | 107 | 132 | 85 | 79 | 124 | 95 | 1 |
| 4351 | - | 4744756 | 4744115 | - | - | *PA14_53510* | hypothetical protein | 20 | 20 | 17 | 18 | 19 | 18 | 1 |
| 4352 | - | 4745877 | 4744858 | - | *oruR* | *PA14_53520* | transcriptional regulator OruR | 31 | 35 | 25 | 31 | 26 | 23 | 1 |
| 4353 | - | 4746002 | 4746835 | + | - | *PA14_53530* | hypothetical protein | 86 | 203 | 124 | 85 | 136 | 125 | 0.283677495 |
| 4354 | - | 4747911 | 4746970 | - | - | *PA14_53540* | hydrolase | 7 | 9 | 6 | 9 | 7 | 7 | 1 |
| 4355 | - | 4748703 | 4748020 | - | - | *PA14_53550* | transcriptional regulator | 24 | 22 | 30 | 42 | 29 | 24 | 1 |
| 4356 | - | 4748791 | 4749669 | + | - | *PA14_53560* | hypothetical protein | 9 | 9 | 9 | 11 | 8 | 8 | 1 |
| 4357 | - | 4749726 | 4750083 | ? | - | predicted RNA | - | 1360173 | 1075686 | 1233957 | 1662863 | 1359183 | 1397545 | 0.770507726 |
| 4358 | - | 4751322 | 4750546 | - | - | *PA14_53570* | integrase | 19 | 22 | 19 | 21 | 17 | 21 | 1 |
| 4359 | - | 4751681 | 4752283 | + | - | *PA14_53580* | hypothetical protein | 34 | 25 | 24 | 16 | 28 | 17 | 1 |
| 4360 | - | 4752288 | 4753559 | + | - | *PA14_53590* | hypothetical protein | 92 | 104 | 63 | 60 | 73 | 60 | 1 |
| 4361 | - | 4753546 | 4754154 | + | - | *PA14_53600* | hypothetical protein | 43 | 42 | 32 | 32 | 45 | 35 | 1 |
| 4362 | - | 4754190 | 4754795 | + | - | *PA14_53610* | hypothetical protein | 8 | 6 | 8 | 7 | 12 | 8 | 1 |
| 4363 | - | 4756330 | 4756968 | + | - | *PA14_53620* | hypothetical protein | 4 | 4 | 2 | 4 | 6 | 4 | 1 |
| 4364 | - | 4757211 | 4757546 | + | - | *PA14_53630* | hypothetical protein | 5 | 7 | 5 | 9 | 10 | 5 | 1 |
| 4365 | - | 4757720 | 4758238 | + | - | *PA14_53640* | hypothetical protein | 27 | 24 | 26 | 32 | 29 | 24 | 1 |
| 4366 | - | 4758490 | 4758936 | + | - | *PA14_53650* | hypothetical protein | 55 | 47 | 53 | 63 | 61 | 58 | 1 |
| 4367 | - | 4758953 | 4760041 | + | - | *PA14_53660* | hypothetical protein | 64 | 57 | 51 | 60 | 68 | 50 | 1 |
| 4368 | - | 4760960 | 4760148 | - | - | *PA14_53670* | hypothetical protein | 148 | 152 | 119 | 137 | 137 | 128 | 1 |

| | A | B | C | D | E | F | G | H | I | J | K | L | M | N |
|---|---|---|---|---|---|---|---|---|---|---|---|---|---|---|
| 4369 | - | 4761493 | 4761260 | - | - | *PA14_53680* | hypothetical protein | 15 | 17 | 13 | 19 | 22 | 10 | 1 |
| 4370 | - | 4761706 | 4761978 | + | - | *PA14_53690* | hypothetical protein | 17 | 22 | 16 | 16 | 23 | 12 | 1 |
| 4371 | - | 4762434 | 4762009 | - | - | *PA14_53700* | ring-cleaving dioxygenase | 28 | 37 | 19 | 33 | 48 | 29 | 1 |
| 4372 | - | 4762535 | 4763419 | + | - | *PA14_53720* | transcriptional regulator | 60 | 57 | 51 | 58 | 62 | 54 | 1 |
| 4373 | - | 4764345 | 4763392 | - | - | *PA14_53730* | transcriptional regulator | 143 | 141 | 124 | 137 | 141 | 140 | 1 |
| 4374 | - | 4764566 | 4765000 | + | - | *PA14_53740* | hypothetical protein | 31 | 44 | 25 | 48 | 62 | 28 | 1 |
| 4375 | - | 4764997 | 4766244 | + | - | *PA14_53750* | hypothetical protein | 48 | 68 | 47 | 78 | 83 | 49 | 1 |
| 4376 | - | 4766533 | 4767828 | + | - | *PA14_53770* | hypothetical protein | 62 | 87 | 55 | 103 | 133 | 63 | 1 |
| 4377 | - | 4767825 | 4769072 | + | - | *PA14_53780* | MFS family transporter | 61 | 92 | 61 | 111 | 127 | 66 | 1 |
| 4378 | - | 4769103 | 4769804 | + | - | *PA14_53790* | haloacid dehalogenase | 93 | 119 | 97 | 145 | 141 | 93 | 1 |
| 4379 | - | 4769911 | 4771227 | + | *mntH2* | *PA14_53800* | manganese transport protein MntH | 49 | 78 | 50 | 90 | 96 | 50 | 1 |
| 4380 | - | 4771752 | 4771279 | - | - | *PA14_53810* | hypothetical protein | 55 | 77 | 68 | 102 | 77 | 55 | 1 |
| 4381 | - | 4772588 | 4771821 | - | - | *PA14_53820* | hypothetical protein | 176 | 322 | 189 | 326 | 311 | 217 | 1 |
| 4382 | - | 4773610 | 4773215 | - | - | *PA14_53830* | hypothetical protein | 12 | 16 | 19 | 24 | 15 | 20 | 1 |
| 4383 | - | 4774171 | 4774398 | + | - | *PA14_53840* | hypothetical protein | 744 | 716 | 544 | 477 | 550 | 585 | 1 |
| 4384 | - | 4775429 | 4774608 | - | - | *PA14_53850* | oxidoreductase | 38 | 38 | 42 | 43 | 37 | 40 | 1 |
| 4385 | - | 4775901 | 4775461 | - | - | *PA14_53860* | hypothetical protein | 52 | 50 | 60 | 49 | 45 | 58 | 1 |
| 4386 | - | 4776310 | 4775987 | - | - | *PA14_53870* | hypothetical protein | 12 | 21 | 18 | 30 | 29 | 14 | 0.728079346 |
| 4387 | - | 4777830 | 4776310 | - | - | *PA14_53880* | hypothetical protein | 15 | 21 | 23 | 27 | 23 | 18 | 1 |
| 4388 | - | 4778360 | 4777827 | - | - | *PA14_53890* | hypothetical protein | 2 | 2 | 4 | 4 | 3 | 2 | 1 |
| 4389 | - | 4778779 | 4780770 | + | - | *PA14_53900* | helicase | 30 | 34 | 38 | 38 | 34 | 26 | 1 |
| 4390 | - | 4781434 | 4780784 | - | *pmtA* | *PA14_53910* | phospholipid methyltransferase | 67 | 75 | 95 | 117 | 67 | 78 | 1 |
| 4391 | - | 4781552 | 4782274 | + | - | *PA14_53920* | transcriptional regulator | 62 | 26 | 37 | 30 | 20 | 29 | 0.113236431 |
| 4392 | - | 4782285 | 4783181 | + | *prpB* | *PA14_53940* | 2-methylisocitrate lyase | 145 | 196 | 126 | 159 | 159 | 149 | 1 |
| 4393 | - | 4783274 | 4784401 | + | *prpC* | *PA14_53950* | methylcitrate synthase | 267 | 355 | 200 | 231 | 273 | 229 | 1 |
| 4394 | - | 4784532 | 4787138 | + | - | *PA14_53970* | aconitate hydratase | 175 | 196 | 161 | 171 | 183 | 172 | 1 |
| 4395 | - | 4787257 | 4788444 | + | - | *PA14_53980* | hypothetical protein | 82 | 95 | 78 | 82 | 92 | 87 | 1 |
| 4396 | - | 4788579 | 4790063 | + | *prpD* | *PA14_54000* | 2-methylcitrate dehydratase | 91 | 102 | 82 | 107 | 104 | 111 | 1 |
| 4397 | - | 4790829 | 4790041 | - | - | *PA14_54010* | transcriptional regulator | 96 | 85 | 84 | 99 | 82 | 93 | 1 |
| 4398 | - | 4790936 | 4791751 | + | - | *PA14_54030* | hypothetical protein | 6 | 6 | 7 | 4 | 5 | 4 | 1 |
| 4399 | - | 4793292 | 4791877 | - | - | *PA14_54040* | amino acid permease | 47 | 57 | 48 | 27 | 45 | 23 | 1 |
| 4400 | - | 4794280 | 4793615 | - | - | *PA14_54050* | hypothetical protein | 71 | 62 | 81 | 63 | 56 | 65 | 1 |
| 4401 | - | 4794494 | 4794285 | - | - | *PA14_54070* | hypothetical protein | 117 | 117 | 145 | 135 | 118 | 132 | 1 |
| 4402 | - | 4794649 | 4797774 | + | - | *PA14_54080* | hypothetical protein | 54 | 60 | 71 | 68 | 71 | 81 | 1 |
| 4403 | - | 4798942 | 4797779 | - | - | *PA14_54090* | hypothetical protein | 30 | 29 | 37 | 29 | 25 | 28 | 1 |
| 4404 | - | 4799303 | 4798998 | - | - | *PA14_54110* | transporter | 3 | 5 | 5 | 6 | 3 | 2 | 1 |
| 4405 | - | 4799951 | 4799313 | - | - | *PA14_54120* | ACP phosphodieterase | 3 | 5 | 3 | 3 | 3 | 4 | 1 |
| 4406 | - | 4800078 | 4801007 | + | - | *PA14_54130* | transcriptional regulator | 21 | 21 | 22 | 18 | 18 | 19 | 1 |
| 4407 | - | 4802672 | 4801152 | - | *putP* | *PA14_54150* | sodium/proline symporter PutP | 150 | 150 | 68 | 52 | 114 | 71 | 1 |
| 4408 | - | 4806166 | 4802984 | - | *putA* | *PA14_54170* | bifunctional proline dehydrogenase/pyrroline-5-carboxylate dehydrogenase | 369 | 313 | 234 | 175 | 270 | 204 | 1 |
| 4409 | - | 4806630 | 4808693 | + | - | *PA14_54180* | hypothetical protein | 8 | 6 | 7 | 8 | 7 | 8 | 1 |
| 4410 | - | 4809567 | 4808815 | - | *pruR* | *PA14_54190* | proline utilization regulator | 35 | 28 | 44 | 37 | 28 | 27 | 1 |
| 4411 | - | 4809758 | 4812157 | + | - | *PA14_54210* | ATP-dependent protease | 167 | 214 | 207 | 96 | 218 | 131 | 1 |
| 4412 | - | 4812268 | 4812672 | + | *icp* | *PA14_54220* | inhibitor of cysteine peptidase | 40 | 48 | 41 | 25 | 47 | 29 | 1 |
| 4413 | - | 4812771 | 4813523 | + | - | *PA14_54230* | hypothetical protein | 21 | 13 | 19 | 10 | 13 | 11 | 0.831807123 |
| 4414 | - | 4813655 | 4814227 | + | - | *PA14_54240* | hypothetical protein | 19 | 16 | 21 | 21 | 16 | 20 | 1 |
| 4415 | - | 4814332 | 4815072 | + | - | *PA14_54260* | hypothetical protein | 11 | 10 | 11 | 11 | 9 | 9 | 1 |
| 4416 | - | 4815069 | 4816037 | + | - | *PA14_54270* | hypothetical protein | 37 | 42 | 48 | 40 | 33 | 29 | 1 |
| 4417 | - | 4816874 | 4816128 | - | *pdxJ* | *PA14_54290* | pyridoxine 5'-phosphate synthase | 93 | 96 | 94 | 91 | 98 | 89 | 1 |
| 4418 | - | 4817568 | 4816867 | - | *recO* | *PA14_54300* | DNA repair protein RecO | 21 | 17 | 21 | 20 | 19 | 18 | 1 |
| 4419 | - | 4818546 | 4817629 | - | *era* | *PA14_54320* | GTP-binding protein Era | 127 | 157 | 148 | 156 | 149 | 152 | 1 |
| 4420 | - | 4819228 | 4818539 | - | *rnc* | *PA14_54330* | ribonuclease III | 205 | 188 | 225 | 198 | 183 | 191 | 1 |
| 4421 | - | 4819602 | 4819225 | - | - | *PA14_54340* | hypothetical protein | 209 | 224 | 264 | 266 | 233 | 232 | 1 |
| 4422 | - | 4820626 | 4819772 | - | *lepB* | *PA14_54350* | signal peptidase I | 171 | 181 | 214 | 179 | 180 | 157 | 1 |
| 4423 | - | 4822431 | 4820632 | - | *lepA* | *PA14_54370* | GTP-binding protein LepA | 96 | 95 | 118 | 103 | 105 | 99 | 1 |
| 4424 | - | 4824005 | 4822581 | - | *mucD* | *PA14_54390* | serine protease MucD | 574 | 606 | 496 | 409 | 551 | 442 | 1 |
| 4425 | - | 4824500 | 4824045 | - | *mucC* | *PA14_54400* | positive regulator for alginate biosynthesis MucC | 263 | 288 | 359 | 321 | 351 | 322 | 1 |
| 4426 | - | 4825447 | 4824497 | - | *mucB* | *PA14_54410* | negative regulator for alginate biosynthesis MucB | 388 | 456 | 455 | 463 | 453 | 451 | 1 |
| 4427 | - | 4826040 | 4825456 | - | *mucA* | *PA14_54420* | anti-sigma factor MucA | 976 | 901 | 1103 | 801 | 861 | 922 | 1 |
| 4428 | - | 4826653 | 4826072 | - | *algU* | *PA14_54430* | RNA polymerase sigma factor AlgU | 1051 | 858 | 1292 | 899 | 1137 | 1049 | 1 |
| 4429 | - | 4827062 | 4828678 | + | *nadB* | *PA14_54450* | L-aspartate oxidase | 124 | 128 | 134 | 136 | 120 | 124 | 1 |
| 4430 | - | 4829099 | 4828647 | - | - | *PA14_54460* | hypothetical protein | 129 | 132 | 144 | 139 | 114 | 104 | 1 |
| 4431 | - | 4829337 | 4829083 | - | - | *PA14_54470* | hypothetical protein | 68 | 86 | 78 | 77 | 68 | 44 | 1 |
| 4432 | - | 4829610 | 4830554 | + | - | *PA14_54480* | hypothetical protein | 78 | 57 | 71 | 60 | 52 | 58 | 1 |
| 4433 | - | 4830655 | 4831491 | + | - | *PA14_54490* | hypothetical protein | 40 | 48 | 52 | 44 | 46 | 40 | 1 |
| 4434 | - | 4832881 | 4831499 | - | - | *PA14_54500* | two-component sensor | 33 | 31 | 37 | 39 | 28 | 27 | 1 |
| 4435 | - | 4833545 | 4832874 | - | - | *PA14_54510* | two-component response regulator | 34 | 24 | 33 | 34 | 26 | 31 | 1 |
| 4436 | - | 4833756 | 4835039 | + | - | *PA14_54520* | porin | 13 | 11 | 13 | 11 | 10 | 11 | 1 |
| 4437 | - | 4835069 | 4836052 | + | - | *PA14_54540* | hypothetical protein | 30 | 33 | 24 | 23 | 23 | 24 | 1 |
| 4438 | - | 4836101 | 4836571 | + | - | *PA14_54550* | hypothetical protein | 19 | 16 | 13 | 16 | 12 | 16 | 1 |
| 4439 | - | 4836582 | 4838099 | + | - | *PA14_54570* | hypothetical protein | 24 | 21 | 20 | 21 | 18 | 22 | 1 |
| 4440 | - | 4838092 | 4839129 | + | - | *PA14_54580* | hypothetical protein | 38 | 33 | 36 | 36 | 35 | 36 | 1 |
| 4441 | - | 4839950 | 4839255 | - | *ung* | *PA14_54590* | uracil-DNA glycosylase | 82 | 88 | 87 | 92 | 75 | 70 | 1 |
| 4442 | - | 4840871 | 4840050 | - | - | *PA14_54600* | hypothetical protein | 27 | 25 | 27 | 32 | 22 | 23 | 1 |
| 4443 | - | 4841839 | 4840928 | - | - | *PA14_54610* | transcriptional regulator | 16 | 15 | 18 | 19 | 14 | 17 | 1 |
| 4444 | - | 4841942 | 4843450 | + | - | *PA14_54620* | aldehyde dehydrogenase | 280 | 323 | 289 | 398 | 292 | 391 | 1 |
| 4445 | - | 4843462 | 4844625 | + | - | *PA14_54630* | acyl-CoA dehydrogenase | 222 | 263 | 229 | 274 | 206 | 302 | 1 |
| 4446 | - | 4844691 | 4845509 | + | - | *PA14_54640* | enoyl-CoA hydratase | 373 | 476 | 408 | 462 | 384 | 507 | 1 |
| 4447 | - | 4845568 | 4846671 | + | - | *PA14_54660* | enoyl-CoA hydratase/isomerase | 219 | 281 | 231 | 287 | 236 | 332 | 1 |
| 4448 | - | 4846783 | 4847679 | + | - | *PA14_54670* | 3-hydroxyisobutyrate dehydrogenase | 133 | 129 | 144 | 151 | 123 | 170 | 1 |
| 4449 | - | 4848072 | 4848380 | + | - | *PA14_54680* | hypothetical protein | 13 | 12 | 11 | 13 | 14 | 15 | 1 |
| 4450 | - | 4848496 | 4849137 | + | - | *PA14_54690* | hypothetical protein | 22 | 18 | 19 | 17 | 18 | 16 | 1 |
| 4451 | - | 4851148 | 4849172 | - | - | *PA14_54700* | beta-lactamase | 7 | 7 | 6 | 8 | 7 | 7 | 1 |
| 4452 | - | 4851251 | 4852165 | + | - | *PA14_54710* | transcriptional regulator | 20 | 20 | 20 | 18 | 15 | 15 | 1 |

| | A | B | C | D | E | F | G | H | I | J | K | L | M | N |
|---|---|---|---|---|---|---|---|---|---|---|---|---|---|---|
| 4453 | - | 4852426 | 4852169 | - | - | *PA14_54720* | hypothetical protein | 32 | 30 | 35 | 29 | 25 | 23 | 1 |
| 4454 | - | 4852725 | 4852480 | - | - | *PA14_54730* | hypothetical protein | 7 | 10 | 6 | 10 | 8 | 7 | 1 |
| 4455 | - | 4853291 | 4852842 | - | - | *PA14_54740* | hypothetical protein | 7 | 7 | 8 | 8 | 9 | 8 | 1 |
| 4456 | - | 4853550 | 4853383 | - | - | *PA14_54750* | hypothetical protein | 58 | 55 | 81 | 84 | 67 | 68 | 1 |
| 4457 | - | 4853789 | 4854865 | + | - | *PA14_54760* | hypothetical protein | 26 | 26 | 22 | 22 | 26 | 22 | 1 |
| 4458 | - | 4854862 | 4855677 | + | - | *PA14_54770* | hypothetical protein | 21 | 30 | 22 | 21 | 20 | 19 | 1 |
| 4459 | - | 4855698 | 4855970 | + | - | *PA14_54790* | hypothetical protein | 30 | 28 | 34 | 31 | 29 | 28 | 1 |
| 4460 | - | 4855970 | 4856662 | + | - | *PA14_54800* | pseudouridylate synthase | 13 | 10 | 15 | 14 | 10 | 11 | 1 |
| 4461 | - | 4856798 | 4857841 | + | - | *PA14_54810* | hypothetical protein | 57 | 51 | 57 | 55 | 53 | 56 | 1 |
| 4462 | - | 4857921 | 4858658 | + | - | *PA14_54820* | hypothetical protein | 30 | 36 | 41 | 29 | 19 | 19 | 1 |
| 4463 | - | 4859110 | 4860012 | + | - | *PA14_54830* | transferase | 55 | 47 | 56 | 58 | 50 | 51 | 1 |
| 4464 | - | 4860143 | 4860213 | + | - | *PA14_54840* | Gly tRNA | 49 | 25 | 40 | 26 | 19 | 19 | 0.320506198 |
| 4465 | - | 4860413 | 4860634 | + | - | *PA14_54850* | hypothetical protein | 26 | 21 | 31 | 24 | 25 | 23 | 1 |
| 4466 | - | 4861445 | 4860837 | - | - | *PA14_54860* | hypothetical protein | 19 | 24 | 22 | 23 | 19 | 18 | 1 |
| 4467 | - | 4862516 | 4861608 | - | - | *PA14_54870* | hypothetical protein | 33 | 82 | 30 | 202 | 72 | 35 | 0.095047946 |
| 4468 | - | 4863513 | 4862635 | - | - | *PA14_54880* | serine acetyltransferase | 33 | 34 | 37 | 39 | 38 | 40 | 1 |
| 4469 | - | 4865318 | 4863414 | - | - | *PA14_54890* | hypothetical protein | 48 | 44 | 50 | 51 | 48 | 51 | 1 |
| 4470 | - | 4866469 | 4865315 | - | - | *PA14_54900* | hypothetical protein | 30 | 29 | 36 | 36 | 31 | 31 | 1 |
| 4471 | - | 4867507 | 4866692 | - | - | *PA14_54910* | thioesterase | 26 | 35 | 32 | 37 | 26 | 34 | 1 |
| 4472 | - | 4871967 | 4867504 | - | - | *PA14_54920* | non-ribosomal peptide synthetase | 33 | 37 | 33 | 36 | 33 | 35 | 1 |
| 4473 | - | 4875437 | 4872096 | - | - | *PA14_54930* | non-ribosomal peptide synthetase | 26 | 27 | 27 | 30 | 26 | 30 | 1 |
| 4474 | - | 4878535 | 4875551 | - | - | *PA14_54940* | siderophore biosynthesis enzyme | 17 | 17 | 17 | 18 | 16 | 18 | 1 |
| 4475 | - | 4880135 | 4878789 | - | - | *PA14_54950* | hypothetical protein | 12 | 9 | 12 | 11 | 9 | 11 | 1 |
| 4476 | - | 4881058 | 4880090 | - | - | *PA14_54960* | hypothetical protein | 8 | 5 | 6 | 7 | 6 | 4 | 1 |
| 4477 | - | 4882919 | 4881162 | - | - | *PA14_54970* | ABC transporter | 10 | 11 | 9 | 10 | 9 | 9 | 1 |
| 4478 | - | 4884677 | 4882923 | - | - | *PA14_54980* | poossible ABC-type transporter protein | 11 | 9 | 9 | 10 | 8 | 9 | 1 |
| 4479 | - | 4884920 | 4885870 | + | - | *PA14_55000* | ABC transporter periplasmic protein | 10 | 11 | 10 | 14 | 11 | 12 | 1 |
| 4480 | - | 4885877 | 4886899 | + | - | *PA14_55020* | ABC transporter permease | 16 | 16 | 16 | 15 | 13 | 15 | 1 |
| 4481 | - | 4886892 | 4887956 | + | - | *PA14_55030* | ABC transporter permease | 12 | 11 | 15 | 13 | 10 | 10 | 1 |
| 4482 | - | 4887953 | 4888810 | + | - | *PA14_55040* | ferric enterobactin transporter ATP-binding protein | 12 | 13 | 13 | 19 | 11 | 16 | 1 |
| 4483 | - | 4888957 | 4891011 | + | - | *PA14_55050* | TonB-dependent receptor | 87 | 79 | 81 | 82 | 81 | 77 | 1 |
| 4484 | - | 4892103 | 4891261 | - | - | *PA14_55060* | hypothetical protein | 49 | 52 | 44 | 56 | 51 | 48 | 1 |
| 4485 | - | 4892399 | 4892136 | - | - | *PA14_55070* | hypothetical protein | 72 | 80 | 75 | 96 | 82 | 81 | 1 |
| 4486 | - | 4893060 | 4894265 | + | - | *PA14_55080* | hypothetical protein | 152 | 107 | 109 | 144 | 135 | 177 | 1 |
| 4487 | - | 4894348 | 4894746 | + | - | *PA14_55090* | hypothetical protein | 88 | 76 | 84 | 89 | 68 | 84 | 1 |
| 4488 | - | 4895700 | 4895224 | - | - | *PA14_55100* | hypothetical protein | 94 | 96 | 123 | 106 | 100 | 95 | 1 |
| 4489 | - | 4896225 | 4895932 | - | - | *PA14_55110* | hypothetical protein | 920 | 1107 | 1026 | 973 | 634 | 554 | 1 |
| 4490 | - | 4896471 | 4897388 | + | - | *PA14_55117* | hypothetical protein | 9 | 10 | 8 | 9 | 8 | 8 | 1 |
| 4491 | - | 4897827 | 4897432 | - | *gloA2* | *PA14_55130* | lactoylglutathione lyase | 8 | 11 | 7 | 10 | 8 | 9 | 1 |
| 4492 | - | 4898168 | 4897872 | - | - | *PA14_55140* | hypothetical protein | 8 | 7 | 6 | 6 | 5 | 5 | 1 |
| 4493 | - | 4898284 | 4899138 | + | - | *PA14_55150* | transcriptional regulator | 6 | 6 | 6 | 7 | 8 | 6 | 1 |
| 4494 | - | 4899424 | 4900203 | + | *toxR* | *PA14_55160* | transcriptional regulator ToxR | 6 | 6 | 7 | 8 | 5 | 6 | 1 |
| 4495 | - | 4900296 | 4900829 | + | *cat* | *PA14_55170* | chloramphenicol acetyltransferase | 36 | 37 | 46 | 38 | 27 | 26 | 1 |
| 4496 | - | 4901189 | 4902088 | + | - | *PA14_55180* | glycosyl transferase family protein | 84 | 67 | 93 | 70 | 63 | 72 | 1 |
| 4497 | - | 4902216 | 4903610 | + | - | *PA14_55200* | amidase | 32 | 31 | 38 | 49 | 30 | 41 | 1 |
| 4498 | - | 4903681 | 4904997 | + | - | *PA14_55220* | MFS family transporter | 12 | 10 | 11 | 13 | 11 | 13 | 1 |
| 4499 | - | 4905076 | 4905981 | + | - | *PA14_55230* | hypothetical protein | 12 | 12 | 11 | 13 | 12 | 11 | 1 |
| 4500 | - | 4906629 | 4905964 | - | - | *PA14_55240* | hypothetical protein | 50 | 51 | 51 | 52 | 45 | 40 | 1 |
| 4501 | - | 4907687 | 4906701 | - | - | *PA14_55250* | transcriptional regulator | 8 | 10 | 8 | 9 | 9 | 9 | 1 |
| 4502 | - | 4908122 | 4907697 | - | - | *PA14_55260* | hypothetical protein | 10 | 11 | 12 | 14 | 10 | 9 | 1 |
| 4503 | - | 4909066 | 4908122 | - | - | *PA14_55280* | peptidyl-prolyl cis-trans isomerase, PpiC-type | 10 | 10 | 9 | 10 | 7 | 8 | 1 |
| 4504 | - | 4909550 | 4909071 | - | - | *PA14_55290* | hypothetical protein | 6 | 6 | 3 | 5 | 3 | 2 | 1 |
| 4505 | - | 4910241 | 4909579 | - | - | *PA14_55300* | hypothetical protein | 1 | 1 | 1 | 2 | 2 | 2 | 1 |
| 4506 | - | 4911982 | 4910276 | - | - | *PA14_55320* | hypothetical protein | 7 | 7 | 6 | 8 | 6 | 6 | 1 |
| 4507 | - | 4912796 | 4912050 | - | - | *PA14_55330* | hypothetical protein | 1 | 1 | 2 | 2 | 2 | 2 | 1 |
| 4508 | - | 4913224 | 4912805 | - | *exbD2* | *PA14_55340* | transport protein ExbD | 2 | 1 | 2 | 1 | 0 | 2 | 1 |
| 4509 | - | 4915082 | 4913235 | - | *exbB2* | *PA14_55360* | transport protein ExbB2 | 4 | 4 | 5 | 4 | 3 | 3 | 1 |
| 4510 | - | 4916740 | 4915106 | - | - | *PA14_55380* | hypothetical protein | 4 | 3 | 3 | 4 | 4 | 4 | 1 |
| 4511 | - | 4917507 | 4916905 | - | - | *PA14_55390* | hypothetical protein | 0 | 0 | 0 | 0 | 0 | 0 | 1 |
| 4512 | - | 4930432 | 4917899 | - | - | *PA14_55400* | hypothetical protein | 15 | 14 | 18 | 17 | 14 | 15 | 1 |
| 4513 | - | 4931810 | 4930632 | - | - | *PA14_55410* | hypothetical protein | 109 | 96 | 121 | 124 | 100 | 122 | 1 |
| 4514 | - | 4933538 | 4932324 | - | - | *PA14_55430* | type II secretion system protein | 8 | 6 | 9 | 7 | 6 | 7 | 1 |
| 4515 | - | 4934947 | 4933538 | - | - | *PA14_55440* | type II secretion system protein | 9 | 11 | 9 | 9 | 7 | 7 | 1 |
| 4516 | - | 4937355 | 4934944 | - | - | *PA14_55450* | type II secretion system protein | 4 | 4 | 5 | 5 | 5 | 5 | 1 |
| 4517 | - | 4937945 | 4937352 | - | - | *PA14_55460* | type II secretion system protein | 4 | 5 | 7 | 6 | 3 | 4 | 1 |
| 4518 | - | 4939074 | 4937932 | - | - | *PA14_55470* | type II secretion system protein | 3 | 3 | 4 | 4 | 2 | 3 | 1 |
| 4519 | - | 4940105 | 4939140 | - | - | *PA14_55480* | type II secretion system protein | 3 | 4 | 4 | 4 | 3 | 3 | 1 |
| 4520 | - | 4940562 | 4940113 | - | - | *PA14_55490* | HxcT pseudopilin | 7 | 5 | 5 | 7 | 4 | 4 | 1 |
| 4521 | - | 4940796 | 4941170 | + | - | *PA14_55500* | HxcV pseudopilin | 7 | 7 | 8 | 6 | 5 | 5 | 1 |
| 4522 | - | 4941604 | 4941173 | - | - | *PA14_55510* | hypothetical protein | 18 | 16 | 21 | 17 | 15 | 17 | 1 |
| 4523 | - | 4941752 | 4942201 | + | - | *PA14_55520* | type II secretion system protein | 4 | 4 | 3 | 4 | 4 | 4 | 1 |
| 4524 | - | 4942185 | 4942826 | + | - | *PA14_55530* | type II secretion system protein | 30 | 26 | 28 | 25 | 21 | 26 | 1 |
| 4525 | - | 4943795 | 4942836 | - | - | *PA14_55540* | transmembrane sensor | 10 | 9 | 11 | 11 | 7 | 9 | 1 |
| 4526 | - | 4944410 | 4943865 | - | - | *PA14_55550* | ECF subfamily RNA polymerase sigma-70 factor | 4 | 5 | 5 | 7 | 5 | 5 | 1 |
| 4527 | - | 4945123 | 4944407 | - | - | *PA14_55560* | hypothetical protein | 10 | 9 | 8 | 14 | 12 | 8 | 1 |
| 4528 | - | 4945289 | 4945609 | + | - | *PA14_55570* | hypothetical protein | 15 | 27 | 13 | 72 | 46 | 15 | 0.8413024 |
| 4529 | - | 4946322 | 4945726 | - | *nemO* | *PA14_55580* | heme oxygenase | 11 | 37 | 11 | 113 | 61 | 11 | 5.66496E-05 |
| 4530 | - | 4946628 | 4947179 | + | - | *PA14_55590* | hypothetical protein | 20 | 19 | 18 | 26 | 21 | 23 | 1 |
| 4531 | - | 4947187 | 4948602 | + | - | *PA14_55600* | hypothetical protein | 12 | 13 | 11 | 13 | 14 | 13 | 1 |
| 4532 | - | 4948578 | 4951673 | + | *dnaE2* | *PA14_55610* | error-prone DNA polymerase | 11 | 11 | 11 | 13 | 12 | 13 | 1 |
| 4533 | - | 4952151 | 4952020 | - | - | *PA14_55620* | 5S ribosomal RNA | 11 | 10 | 26 | 11 | 5 | 1 | 1 |
| 4534 | - | 4955171 | 4952288 | - | - | *PA14_55631* | 23S ribosomal RNA | 610 | 869 | 717 | 414 | 745 | 121 | 1 |
| 4535 | - | 4955473 | 4955401 | - | - | *PA14_55634* | Ala tRNA | 696 | 956 | 932 | 643 | 489 | 240 | 1 |
| 4536 | - | 4955578 | 4955505 | - | - | *PA14_55635* | Ile tRNA | 710 | 1071 | 928 | 568 | 435 | 216 | 1 |

| | A | B | C | D | E | F | G | H | I | J | K | L | M | N |
|---|---|---|---|---|---|---|---|---|---|---|---|---|---|---|
| 4537 | - | 4957169 | 4955644 | - | - | *PA14_55637* | 16S ribosomal RNA | 237 | 360 | 900 | 310 | 553 | 68 | 1 |
| 4538 | - | 4957850 | 4959079 | + | *sbcD* | *PA14_55640* | exonuclease SbcD | 20 | 17 | 15 | 13 | 15 | 14 | 1 |
| 4539 | - | 4959088 | 4962723 | + | *sbcC* | *PA14_55650* | exonuclease | 38 | 34 | 35 | 37 | 35 | 38 | 1 |
| 4540 | - | 4964910 | 4962745 | - | *recD* | *PA14_55660* | exodeoxyribonuclease V subunit alpha | 146 | 136 | 138 | 145 | 155 | 164 | 1 |
| 4541 | - | 4968644 | 4964907 | - | *recB* | *PA14_55670* | exodeoxyribonuclease V subunit beta | 135 | 127 | 129 | 144 | 144 | 150 | 1 |
| 4542 | - | 4972156 | 4968641 | - | *recC* | *PA14_55690* | exodeoxyribonuclease V subunit gamma | 100 | 91 | 93 | 102 | 96 | 95 | 1 |
| 4543 | - | 4972880 | 4972197 | - | - | *PA14_55710* | hypothetical protein | 104 | 90 | 109 | 110 | 102 | 87 | 1 |
| 4544 | - | 4973883 | 4972993 | - | - | *PA14_55720* | hypothetical protein | 4 | 3 | 5 | 5 | 4 | 5 | 1 |
| 4545 | - | 4973983 | 4974786 | + | - | *PA14_55730* | AraC family transcriptional regulator | 19 | 17 | 16 | 20 | 17 | 19 | 1 |
| 4546 | - | 4974841 | 4976046 | + | - | *PA14_55740* | transporter | 64 | 39 | 50 | 36 | 34 | 34 | 1 |
| 4547 | - | 4977658 | 4978344 | + | - | *PA14_55760* | hypothetical protein | 22 | 17 | 32 | 20 | 14 | 15 | 1 |
| 4548 | - | 4977659 | 4976043 | - | - | *PA14_55750* | chemotaxis transducer | 130 | 76 | 101 | 62 | 48 | 51 | 1 |
| 4549 | - | 4978489 | 4979958 | + | - | *PA14_55770* | phosphate transporter | 42 | 36 | 46 | 34 | 34 | 39 | 1 |
| 4550 | - | 4982794 | 4980026 | - | - | *PA14_55780* | two-component sensor | 117 | 64 | 87 | 90 | 92 | 168 | 1 |
| 4551 | - | 4983310 | 4982804 | - | - | *PA14_55790* | hypothetical protein | 188 | 142 | 148 | 133 | 172 | 245 | 1 |
| 4552 | - | 4983374 | 4983856 | + | - | *PA14_55800* | hypothetical protein | 9 | 6 | 6 | 8 | 7 | 12 | 1 |
| 4553 | - | 4983977 | 4984804 | + | - | *PA14_55810* | two-component response regulator | 154 | 115 | 100 | 130 | 123 | 170 | 1 |
| 4554 | - | 4986500 | 4984830 | - | - | *PA14_55820* | hypothetical protein | 54 | 39 | 42 | 73 | 61 | 117 | 1 |
| 4555 | - | 4986809 | 4986525 | - | - | *PA14_55840* | hypothetical protein | 95 | 66 | 63 | 124 | 92 | 245 | 1 |
| 4556 | - | 4987569 | 4986832 | - | - | *PA14_55850* | pilus assembly protein | 86 | 56 | 57 | 110 | 84 | 204 | 1 |
| 4557 | - | 4988477 | 4987566 | - | - | *PA14_55860* | hypothetical protein | 61 | 45 | 55 | 78 | 61 | 127 | 1 |
| 4558 | - | 4989371 | 4988487 | - | - | *PA14_55880* | hypothetical protein | 96 | 66 | 85 | 124 | 89 | 203 | 1 |
| 4559 | - | 4990633 | 4989368 | - | - | *PA14_55890* | type II secretion system protein | 58 | 40 | 40 | 76 | 47 | 120 | 1 |
| 4560 | - | 4991814 | 4990630 | - | - | *PA14_55900* | hypothetical protein | 76 | 56 | 56 | 96 | 59 | 143 | 1 |
| 4561 | - | 4993074 | 4991824 | - | - | *PA14_55920* | type II secretion system protein | 80 | 57 | 66 | 115 | 66 | 155 | 1 |
| 4562 | - | 4993999 | 4993088 | - | - | *PA14_55930* | pilus assembly protein | 87 | 69 | 83 | 119 | 83 | 155 | 1 |
| 4563 | - | 4994370 | 4994392 | ? | - | predicted RNA | - | 414 | 325 | 339 | 497 | 418 | 847 | 1 |
| 4564 | - | 4994409 | 4994621 | + | - | *PA14_55940* | hypothetical protein | 1082 | 915 | 940 | 1499 | 1276 | 2519 | 1 |
| 4565 | - | 4994810 | 4996708 | + | *pctC* | *PA14_55960* | chemotactic transducer PctC | 39 | 63 | 43 | 50 | 52 | 49 | 1 |
| 4566 | - | 4998302 | 4996812 | - | - | *PA14_55980* | hypothetical protein | 90 | 80 | 74 | 71 | 66 | 71 | 1 |
| 4567 | - | 4998702 | 5000591 | + | *pctA* | *PA14_56000* | chemotactic transducer PctA | 61 | 79 | 68 | 67 | 73 | 80 | 1 |
| 4568 | - | 5000826 | 5002715 | + | *pctB* | *PA14_56010* | chemotactic transducer PctB | 62 | 92 | 73 | 71 | 77 | 69 | 1 |
| 4569 | - | 5003033 | 5004148 | + | - | *PA14_56030* | hypothetical protein | 222 | 220 | 258 | 297 | 258 | 302 | 1 |
| 4570 | - | 5004141 | 5004902 | + | - | *PA14_56040* | hypothetical protein | 110 | 110 | 116 | 151 | 128 | 152 | 1 |
| 4571 | - | 5004899 | 5005900 | + | - | *PA14_56050* | hypothetical protein | 84 | 84 | 85 | 94 | 80 | 86 | 1 |
| 4572 | - | 5006988 | 5006137 | - | *purU* | *PA14_56060* | formyltetrahydrofolate deformylase | 84 | 85 | 77 | 67 | 70 | 57 | 1 |
| 4573 | - | 5007248 | 5007622 | + | *mvaT* | *PA14_56070* | transcriptional regulator MvaT, P16 subunit | 345 | 431 | 357 | 212 | 301 | 261 | 1 |
| 4574 | - | 5009149 | 5007707 | - | *sbcB* | *PA14_56080* | exonuclease I | 66 | 67 | 67 | 64 | 57 | 58 | 1 |
| 4575 | - | 5009374 | 5010111 | + | - | *PA14_56090* | hypothetical protein | 40 | 29 | 33 | 21 | 27 | 20 | 1 |
| 4576 | - | 5010149 | 5010946 | + | - | *PA14_56100* | hypothetical protein | 21 | 18 | 24 | 16 | 19 | 13 | 1 |
| 4577 | - | 5010943 | 5011923 | + | - | *PA14_56110* | hypothetical protein | 39 | 35 | 41 | 34 | 34 | 26 | 1 |
| 4578 | - | 5011910 | 5013481 | + | - | *PA14_56130* | hypothetical protein | 49 | 43 | 46 | 42 | 47 | 37 | 1 |
| 4579 | - | 5013478 | 5014755 | + | - | *PA14_56140* | hypothetical protein | 27 | 30 | 29 | 26 | 34 | 24 | 1 |
| 4580 | - | 5014752 | 5015759 | + | - | *PA14_56160* | hypothetical protein | 90 | 102 | 83 | 73 | 117 | 72 | 1 |
| 4581 | - | 5015756 | 5017087 | + | - | *PA14_56170* | hypothetical protein | 112 | 110 | 139 | 115 | 119 | 111 | 1 |
| 4582 | - | 5017451 | 5017092 | - | - | *PA14_56180* | hypothetical protein | 280 | 351 | 328 | 293 | 307 | 347 | 1 |
| 4583 | - | 5017934 | 5017512 | - | - | *PA14_56190* | hypothetical protein | 153 | 139 | 201 | 211 | 140 | 153 | 1 |
| 4584 | - | 5018418 | 5018035 | - | - | *PA14_56200* | lipoprotein | 105 | 116 | 126 | 119 | 119 | 125 | 1 |
| 4585 | - | 5018512 | 5019318 | + | - | *PA14_56210* | hypothetical protein | 26 | 27 | 32 | 31 | 24 | 26 | 1 |
| 4586 | - | 5019403 | 5020317 | + | - | *PA14_56220* | hypothetical protein | 206 | 172 | 233 | 118 | 116 | 158 | 1 |
| 4587 | - | 5020400 | 5021851 | + | *pykA* | *PA14_56240* | pyruvate kinase | 191 | 259 | 210 | 165 | 227 | 166 | 1 |
| 4588 | - | 5022814 | 5022041 | - | - | *PA14_56250* | enoyl-CoA hydratase/isomerase | 71 | 77 | 67 | 65 | 65 | 70 | 1 |
| 4589 | - | 5023806 | 5022880 | - | - | *PA14_56260* | hypothetical protein | 152 | 132 | 139 | 146 | 124 | 131 | 1 |
| 4590 | - | 5024926 | 5023799 | - | - | *PA14_56280* | hypothetical protein | 116 | 111 | 107 | 118 | 103 | 109 | 1 |
| 4591 | - | 5025202 | 5026725 | + | - | *PA14_56300* | fumarase | 71 | 88 | 72 | 58 | 69 | 45 | 1 |
| 4592 | - | 5028078 | 5026822 | - | *mntH1* | *PA14_56340* | manganese transport protein MntH | 28 | 31 | 29 | 28 | 30 | 26 | 1 |
| 4593 | - | 5028523 | 5028215 | - | - | *PA14_56360* | hypothetical protein | 67 | 87 | 69 | 56 | 65 | 61 | 1 |
| 4594 | - | 5029128 | 5028544 | - | - | *PA14_56370* | hypothetical protein | 96 | 121 | 100 | 74 | 81 | 88 | 1 |
| 4595 | - | 5029843 | 5029232 | - | - | *PA14_56380* | hypothetical protein | 16 | 25 | 25 | 25 | 26 | 26 | 1 |
| 4596 | - | 5031217 | 5029937 | - | - | *PA14_56390* | hypothetical protein | 58 | 62 | 59 | 67 | 64 | 66 | 1 |
| 4597 | - | 5032396 | 5031317 | - | - | *PA14_56410* | phospholipase | 36 | 36 | 36 | 33 | 40 | 30 | 1 |
| 4598 | - | 5033043 | 5032471 | - | - | *PA14_56420* | hypothetical protein | 101 | 88 | 115 | 93 | 71 | 81 | 1 |
| 4599 | - | 5033878 | 5033108 | - | - | *PA14_56430* | transcriptional regulator | 66 | 73 | 85 | 67 | 74 | 63 | 1 |
| 4600 | - | 5035462 | 5033978 | - | - | *PA14_56450* | amidase | 25 | 23 | 23 | 21 | 19 | 22 | 1 |
| 4601 | - | 5036782 | 5035463 | - | - | *PA14_56470* | MFS transporter | 4 | 3 | 4 | 3 | 3 | 3 | 1 |
| 4602 | - | 5038040 | 5036820 | - | - | *PA14_56480* | hydrolase | 32 | 36 | 36 | 43 | 41 | 46 | 1 |
| 4603 | - | 5038795 | 5038205 | - | - | *PA14_56510* | hypothetical protein | 127 | 123 | 129 | 130 | 143 | 132 | 1 |
| 4604 | - | 5039191 | 5038856 | - | - | *PA14_56520* | hypothetical protein | 60 | 61 | 56 | 62 | 49 | 54 | 1 |
| 4605 | - | 5040342 | 5039188 | - | - | *PA14_56530* | hypothetical protein | 38 | 35 | 36 | 33 | 33 | 37 | 1 |
| 4606 | - | 5041156 | 5040371 | - | - | *PA14_56540* | hypothetical protein | 341 | 245 | 454 | 239 | 189 | 313 | 1 |
| 4607 | - | 5042137 | 5041241 | - | - | *PA14_56550* | hypothetical protein | 83 | 61 | 80 | 67 | 64 | 81 | 1 |
| 4608 | - | 5042340 | 5043095 | + | - | *PA14_56560* | hypothetical protein | 238 | 205 | 300 | 222 | 193 | 250 | 1 |
| 4609 | - | 5043095 | 5043871 | + | - | *PA14_56570* | acyltransferase | 559 | 427 | 613 | 426 | 430 | 603 | 1 |
| 4610 | - | 5044728 | 5043868 | - | - | *PA14_56590* | hypothetical protein | 1983 | 1296 | 1876 | 1064 | 1131 | 2049 | 1 |
| 4611 | - | 5045450 | 5044878 | - | - | *PA14_56600* | hypothetical protein | 55 | 35 | 46 | 34 | 39 | 53 | 1 |
| 4612 | - | 5045603 | 5045905 | + | - | *PA14_56620* | hypothetical protein | 24 | 28 | 31 | 26 | 24 | 20 | 1 |
| 4613 | - | 5045954 | 5047120 | + | - | *PA14_56640* | MFS transporter | 11 | 25 | 23 | 20 | 16 | 16 | 0.147672258 |
| 4614 | - | 5047157 | 5048209 | + | - | *PA14_56660* | xenobiotic reductase | 38 | 136 | 108 | 89 | 92 | 70 | 0.000975615 |
| 4615 | - | 5048511 | 5048269 | - | - | *PA14_56670* | hypothetical protein | 60 | 207 | 138 | 163 | 133 | 131 | 0.000229501 |
| 4616 | - | 5050811 | 5048511 | - | *feoB* | *PA14_56680* | ferrous iron transport protein B | 35 | 74 | 48 | 57 | 82 | 79 | 0.598517837 |
| 4617 | - | 5051056 | 5050829 | - | *feoA* | *PA14_56690* | ferrous iron transport protein A | 21 | 59 | 22 | 20 | 120 | 86 | 0.002562618 |
| 4618 | - | 5051746 | 5051273 | - | - | *PA14_56700* | hypothetical protein | 34 | 24 | 45 | 25 | 23 | 31 | 1 |
| 4619 | - | 5053653 | 5052664 | - | - | *PA14_56720* | oxidoreductase | 45 | 42 | 42 | 40 | 43 | 41 | 1 |
| 4620 | - | 5053838 | 5054896 | + | - | *PA14_56730* | hypothetical protein | 52 | 47 | 53 | 60 | 50 | 58 | 1 |

| | A | B | C | D | E | F | G | H | I | J | K | L | M | N |
|---|---|---|---|---|---|---|---|---|---|---|---|---|---|---|
| 4621 | - | 5055799 | 5054897 | - | *iciA* | *PA14_56740* | chromosome replication initiation inhibitor protein | 62 | 59 | 67 | 65 | 62 | 65 | 1 |
| 4622 | - | 5055897 | 5056298 | + | - | *PA14_56750* | hypothetical protein | 4 | 1 | 5 | 3 | 1 | 3 | 0.059601067 |
| 4623 | - | 5056300 | 5056902 | + | - | *PA14_56770* | transporter | 8 | 10 | 12 | 10 | 7 | 9 | 1 |
| 4624 | - | 5057249 | 5057830 | + | *sodB* | *PA14_56780* | superoxide dismutase | 744 | 884 | 473 | 354 | 659 | 680 | 1 |
| 4625 | - | 5058011 | 5060074 | + | - | *PA14_56790* | diguanylate cyclase | 80 | 83 | 81 | 83 | 79 | 90 | 1 |
| 4626 | - | 5060977 | 5060075 | - | - | *PA14_56800* | hypothetical protein | 133 | 147 | 169 | 161 | 137 | 153 | 1 |
| 4627 | - | 5061551 | 5060979 | - | - | *PA14_56810* | lemA-like protein | 101 | 108 | 122 | 101 | 113 | 104 | 1 |
| 4628 | - | 5061745 | 5063085 | + | *icmP* | *PA14_56830* | metalloproteinase outer membrane | 66 | 120 | 66 | 90 | 102 | 67 | 1 |
| 4629 | - | 5063248 | 5064669 | + | - | *PA14_56840* | hypothetical protein | 15 | 23 | 19 | 31 | 35 | 21 | 1 |
| 4630 | - | 5064679 | 5065743 | + | - | *PA14_56850* | lipoprotein | 39 | 66 | 42 | 71 | 74 | 50 | 0.831807123 |
| 4631 | - | 5065754 | 5066848 | + | - | *PA14_56870* | hypothetical protein | 64 | 71 | 60 | 82 | 70 | 55 | 1 |
| 4632 | - | 5067018 | 5068148 | + | - | *PA14_56880* | membrane fusion protein | 41 | 38 | 43 | 44 | 38 | 37 | 1 |
| 4633 | - | 5068199 | 5071255 | + | - | *PA14_56890* | multidrug efflux protein | 51 | 44 | 45 | 45 | 47 | 41 | 1 |
| 4634 | - | 5071393 | 5072589 | + | *pncB2* | *PA14_56900* | nicotinate phosphoribosyltransferase | 35 | 35 | 37 | 34 | 30 | 35 | 1 |
| 4635 | - | 5072956 | 5072714 | - | - | *PA14_56910* | hypothetical protein | 133 | 201 | 245 | 275 | 112 | 108 | 1 |
| 4636 | - | 5073657 | 5072953 | - | *inaA* | *PA14_56920* | InaA protein | 59 | 44 | 60 | 43 | 35 | 46 | 1 |
| 4637 | - | 5074343 | 5073660 | - | - | *PA14_56930* | hypothetical protein | 56 | 56 | 73 | 67 | 38 | 49 | 1 |
| 4638 | - | 5075703 | 5074423 | - | - | *PA14_56940* | two-component sensor | 26 | 25 | 25 | 27 | 24 | 24 | 1 |
| 4639 | - | 5076376 | 5075693 | - | - | *PA14_56950* | two-component response regulator | 57 | 47 | 51 | 66 | 46 | 54 | 1 |
| 4640 | - | 5076593 | 5077387 | + | - | *PA14_56960* | hypothetical protein | 27 | 30 | 37 | 41 | 27 | 41 | 1 |
| 4641 | - | 5077659 | 5077330 | - | - | *PA14_56970* | hypothetical protein | 60 | 48 | 51 | 74 | 47 | 70 | 1 |
| 4642 | - | 5078062 | 5077679 | - | - | *PA14_56980* | camphor resistance protein CrcB | 41 | 33 | 36 | 48 | 36 | 47 | 1 |
| 4643 | - | 5078292 | 5078978 | + | - | *PA14_56990* | hypothetical protein | 37 | 15 | 14 | 19 | 20 | 40 | 0.05276977 |
| 4644 | - | 5080678 | 5079035 | - | *groEL* | *PA14_57010* | chaperonin GroEL | 1286 | 2653 | 1231 | 505 | 1897 | 894 | 1 |
| 4645 | - | 5081022 | 5080729 | - | *groES* | *PA14_57020* | co-chaperonin GroES | 856 | 1819 | 1067 | 406 | 1063 | 552 | 1 |
| 4646 | - | 5081685 | 5081218 | - | *fxsA* | *PA14_57030* | FxsA protein | 168 | 138 | 260 | 74 | 173 | 106 | 1 |
| 4647 | - | 5082488 | 5081754 | - | - | *PA14_57040* | hypothetical protein | 55 | 52 | 42 | 36 | 42 | 35 | 1 |
| 4648 | - | 5082597 | 5083355 | + | *fabG* | *PA14_57050* | 3-ketoacyl-ACP reductase | 273 | 307 | 239 | 250 | 243 | 201 | 1 |
| 4649 | - | 5084421 | 5083414 | - | - | *PA14_57060* | hypothetical protein | 42 | 34 | 52 | 38 | 38 | 33 | 1 |
| 4650 | - | 5085536 | 5085919 | + | - | *PA14_57080* | hypothetical protein | 62 | 55 | 75 | 72 | 53 | 56 | 1 |
| 4651 | - | 5085537 | 5084536 | - | - | *PA14_57070* | hypothetical protein | 19 | 21 | 25 | 24 | 16 | 15 | 1 |
| 4652 | - | 5085962 | 5087746 | + | - | *PA14_57100* | permease | 31 | 25 | 31 | 28 | 28 | 26 | 1 |
| 4653 | - | 5088639 | 5087803 | - | - | *PA14_57110* | hypothetical protein | 97 | 126 | 112 | 82 | 122 | 88 | 1 |
| 4654 | - | 5089351 | 5088872 | - | - | *PA14_57130* | nucleotide-binding protein | 118 | 130 | 111 | 82 | 96 | 86 | 1 |
| 4655 | - | 5089455 | 5090555 | + | - | *PA14_57140* | two-component response regulator | 17 | 17 | 18 | 17 | 16 | 16 | 1 |
| 4656 | - | 5090592 | 5091503 | + | *apbA* | *PA14_57160* | 2-dehydropantoate 2-reductase | 29 | 29 | 38 | 31 | 27 | 38 | 1 |
| 4657 | - | 5091554 | 5093650 | + | - | *PA14_57170* | two-component sensor | 22 | 22 | 24 | 24 | 22 | 25 | 1 |
| 4658 | - | 5093655 | 5094233 | + | - | *PA14_57180* | hypothetical protein | 77 | 83 | 77 | 75 | 76 | 71 | 1 |
| 4659 | - | 5095250 | 5094303 | - | - | *PA14_57190* | hypothetical protein | 111 | 88 | 92 | 93 | 89 | 80 | 1 |
| 4660 | - | 5095939 | 5095304 | - | - | *PA14_57200* | glutathione S-transferase | 148 | 130 | 144 | 142 | 133 | 117 | 1 |
| 4661 | - | 5097277 | 5096060 | - | *argJ* | *PA14_57210* | bifunctional ornithine acetyltransferase/N-acetylglutamate synthase | 115 | 140 | 120 | 106 | 124 | 86 | 1 |
| 4662 | - | 5100174 | 5097424 | - | *secA* | *PA14_57220* | preprotein translocase subunit SecA | 352 | 367 | 451 | 399 | 328 | 334 | 1 |
| 4663 | - | 5101231 | 5100308 | - | - | *PA14_57240* | hypothetical protein | 118 | 123 | 144 | 157 | 116 | 108 | 1 |
| 4664 | - | 5101378 | 5101773 | + | - | *PA14_57250* | hypothetical protein | 51 | 43 | 57 | 48 | 45 | 40 | 1 |
| 4665 | - | 5102745 | 5101834 | - | *lpxC* | *PA14_57260* | UDP-3-O-[3-hydroxymyristoyl] N-acetylglucosamine deacetylase | 554 | 569 | 494 | 398 | 453 | 454 | 1 |
| 4666 | - | 5104042 | 5102858 | - | *ftsZ* | *PA14_57275* | cell division protein FtsZ | 1068 | 1265 | 996 | 927 | 1093 | 1087 | 1 |
| 4667 | - | 5105346 | 5104093 | - | *ftsA* | *PA14_57290* | cell division protein FtsA | 1175 | 997 | 1250 | 1099 | 1133 | 1204 | 1 |
| 4668 | - | 5106231 | 5105368 | - | *ftsQ* | *PA14_57300* | cell division protein FtsQ | 403 | 327 | 385 | 372 | 404 | 480 | 1 |
| 4669 | - | 5107194 | 5106235 | - | *ddl* | *PA14_57320* | D-alanine--D-alanine ligase | 791 | 681 | 804 | 787 | 822 | 932 | 1 |
| 4670 | - | 5108633 | 5107191 | - | *murC* | *PA14_57330* | UDP-N-acetylmuramate--L-alanine ligase | 603 | 562 | 568 | 550 | 598 | 622 | 1 |
| 4671 | - | 5109699 | 5108626 | - | *murG* | *PA14_57340* | UDPdiphospho-muramoylpentapeptide beta-N- acetylglucosaminyltransferase | 709 | 599 | 646 | 606 | 605 | 672 | 1 |
| 4672 | - | 5110888 | 5109689 | - | *ftsW* | *PA14_57360* | cell division protein FtsW | 482 | 444 | 519 | 563 | 515 | 580 | 1 |
| 4673 | - | 5112234 | 5110888 | - | *murD* | *PA14_57370* | UDP-N-acetylmuramoyl-L-alanyl-D-glutamate synthetase | 411 | 373 | 373 | 395 | 399 | 440 | 1 |
| 4674 | - | 5113330 | 5112248 | - | *mraY* | *PA14_57380* | phospho-N-acetylmuramoyl-pentapeptide-transferase | 267 | 261 | 244 | 234 | 266 | 276 | 1 |
| 4675 | - | 5114706 | 5113330 | - | *murF* | *PA14_57390* | UDP-N-acetylmuramoylalanyl-D-glutamyl-2,6- diaminopimelate--D-alanyl-D-alanine ligase | 446 | 415 | 410 | 456 | 484 | 498 | 1 |
| 4676 | - | 5116162 | 5114699 | - | *murE* | *PA14_57410* | UDP-N-acetylmuramoylalanyl-D-glutamate--2,6-diaminopimelate ligase | 473 | 454 | 462 | 474 | 487 | 510 | 1 |
| 4677 | - | 5117901 | 5116162 | - | *ftsI* | *PA14_57425* | penicillin-binding protein 3 | 493 | 453 | 480 | 558 | 523 | 520 | 1 |
| 4678 | - | 5118191 | 5117898 | - | *ftsL* | *PA14_57440* | cell division protein FtsL | 357 | 354 | 323 | 350 | 316 | 354 | 1 |
| 4679 | - | 5119129 | 5118188 | - | *mraW* | *PA14_57450* | S-adenosyl-methyltransferase MraW | 349 | 343 | 338 | 366 | 412 | 405 | 1 |
| 4680 | - | 5119587 | 5119132 | - | - | *PA14_57460* | cell division protein MraZ | 590 | 499 | 736 | 539 | 698 | 654 | 1 |
| 4681 | - | 5121140 | 5120292 | - | - | *PA14_57470* | hypothetical protein | 43 | 31 | 38 | 41 | 25 | 30 | 1 |
| 4682 | - | 5121267 | 5123081 | + | - | *PA14_57480* | lipoprotein | 76 | 69 | 76 | 76 | 67 | 69 | 1 |
| 4683 | - | 5123078 | 5123455 | + | - | *PA14_57490* | hypothetical protein | 135 | 114 | 112 | 127 | 110 | 109 | 1 |
| 4684 | - | 5123484 | 5124077 | + | *gmhA* | *PA14_57500* | phosphoheptose isomerase | 148 | 145 | 145 | 128 | 106 | 128 | 1 |
| 4685 | - | 5124074 | 5124652 | + | - | *PA14_57510* | secreted lipoprotein | 87 | 100 | 82 | 89 | 94 | 91 | 1 |
| 4686 | - | 5125116 | 5124709 | - | *sspB* | *PA14_57520* | ClpXP protease specificity-enhancing factor | 104 | 150 | 126 | 117 | 118 | 97 | 1 |
| 4687 | - | 5125745 | 5125128 | - | *sspA* | *PA14_57530* | stringent starvation protein A | 61 | 73 | 77 | 66 | 83 | 60 | 1 |
| 4688 | - | 5126613 | 5125831 | - | - | *PA14_57540* | cytochrome c1 | 442 | 802 | 428 | 473 | 501 | 330 | 1 |
| 4689 | - | 5127824 | 5126613 | - | - | *PA14_57560* | cytochrome b | 397 | 683 | 389 | 487 | 572 | 340 | 1 |
| 4690 | - | 5128417 | 5127824 | - | - | *PA14_57570* | cytochrome c reductase, iron-sulfur subun | 406 | 513 | 460 | 383 | 476 | 330 | 1 |
| 4691 | - | 5129059 | 5128667 | - | *rpsI* | *PA14_57580* | 30S ribosomal protein S9 | 322 | 396 | 364 | 227 | 239 | 163 | 1 |
| 4692 | - | 5129502 | 5129074 | - | *rplM* | *PA14_57590* | 50S ribosomal protein L13 | 303 | 309 | 321 | 205 | 204 | 156 | 1 |
| 4693 | - | 5129748 | 5130785 | + | - | *PA14_57600* | oxidoreductase | 58 | 75 | 60 | 69 | 64 | 64 | 1 |
| 4694 | - | 5132303 | 5133202 | + | - | *PA14_57630* | AraC family transcriptional regulator | 39 | 41 | 49 | 50 | 39 | 44 | 1 |
| 4695 | - | 5133361 | 5134323 | + | - | *PA14_57640* | hypothetical protein | 14 | 15 | 14 | 16 | 10 | 10 | 1 |
| 4696 | - | 5135482 | 5134388 | - | - | *PA14_57650* | hypothetical protein | 27 | 17 | 25 | 18 | 13 | 14 | 1 |
| 4697 | - | 5136923 | 5135577 | - | *trpS* | *PA14_57670* | tryptophanyl-tRNA synthetase | 49 | 57 | 45 | 36 | 43 | 39 | 1 |
| 4698 | - | 5137620 | 5136991 | - | - | *PA14_57680* | hypothetical protein | 57 | 47 | 57 | 45 | 52 | 42 | 1 |
| 4699 | - | 5137765 | 5138211 | + | - | *PA14_57690* | hypothetical protein | 133 | 155 | 124 | 107 | 144 | 145 | 1 |
| 4700 | - | 5140186 | 5138285 | - | *cysN* | *PA14_57710* | bifunctional sulfate adenylyltransferase subunit 1/adenylylsulfate kinase | 90 | 126 | 54 | 61 | 102 | 55 | 1 |
| 4701 | - | 5141115 | 5140198 | - | *cysD* | *PA14_57720* | sulfate adenylyltransferase subunit 2 | 42 | 58 | 43 | 32 | 49 | 35 | 1 |
| 4702 | - | 5141460 | 5142563 | + | *mltB1* | *PA14_57730* | murein hydrolase B | 76 | 74 | 92 | 87 | 66 | 71 | 1 |
| 4703 | - | 5143308 | 5142550 | - | - | *PA14_57740* | hypothetical protein | 122 | 123 | 127 | 141 | 102 | 118 | 1 |
| 4704 | - | 5143425 | 5144594 | + | *algW* | *PA14_57760* | AlgW protein | 80 | 73 | 88 | 98 | 68 | 63 | 1 |

| | A | B | C | D | E | F | G | H | I | J | K | L | M | N |
|---|---|---|---|---|---|---|---|---|---|---|---|---|---|---|
| 4705 | - | 5145692 | 5144637 | - | *hisC1* | *PA14_57770* | histidinol-phosphate aminotransferase | 95 | 98 | 80 | 86 | 87 | 77 | 1 |
| 4706 | - | 5147017 | 5145695 | - | *hisD* | *PA14_57780* | histidinol dehydrogenase | 82 | 65 | 67 | 64 | 63 | 60 | 1 |
| 4707 | - | 5147778 | 5147143 | - | *hisG* | *PA14_57800* | ATP phosphoribosyltransferase | 145 | 161 | 149 | 143 | 155 | 127 | 1 |
| 4708 | - | 5149061 | 5147796 | - | *murA* | *PA14_57810* | UDP-N-acetylglucosamine 1-carboxyvinyltransferase | 205 | 209 | 205 | 206 | 181 | 164 | 1 |
| 4709 | - | 5149321 | 5149082 | - | - | *PA14_57820* | hypothetical protein | 290 | 222 | 372 | 328 | 262 | 253 | 1 |
| 4710 | - | 5149746 | 5149438 | - | - | *PA14_57830* | hypothetical protein | 251 | 238 | 293 | 255 | 255 | 206 | 1 |
| 4711 | - | 5150390 | 5149743 | - | - | *PA14_57840* | hypothetical protein | 192 | 220 | 207 | 179 | 195 | 187 | 1 |
| 4712 | - | 5150875 | 5150402 | - | - | *PA14_57850* | hypothetical protein | 120 | 132 | 116 | 121 | 120 | 105 | 1 |
| 4713 | - | 5151673 | 5150876 | - | - | *PA14_57870* | ABC transporter permease | 94 | 88 | 95 | 88 | 86 | 73 | 1 |
| 4714 | - | 5152482 | 5151673 | - | - | *PA14_57880* | ABC transporter ATP-binding protein | 118 | 88 | 106 | 91 | 95 | 90 | 1 |
| 4715 | - | 5152896 | 5153876 | + | - | *PA14_57890* | hypothetical protein | 100 | 81 | 124 | 88 | 83 | 89 | 1 |
| 4716 | - | 5153876 | 5154415 | + | - | *PA14_57900* | hypothetical protein | 144 | 147 | 171 | 162 | 147 | 145 | 1 |
| 4717 | - | 5154424 | 5154996 | + | - | *PA14_57910* | hypothetical protein | 178 | 175 | 198 | 208 | 172 | 180 | 1 |
| 4718 | - | 5154983 | 5155510 | + | - | *PA14_57920* | hypothetical protein | 206 | 224 | 212 | 191 | 190 | 175 | 1 |
| 4719 | - | 5155510 | 5156235 | + | - | *PA14_57930* | ABC transporter ATP-binding protein | 146 | 153 | 158 | 136 | 144 | 136 | 1 |
| 4720 | - | 5156461 | 5157954 | + | *rpoN* | *PA14_57940* | RNA polymerase factor sigma-54 | 83 | 77 | 88 | 85 | 80 | 82 | 1 |
| 4721 | - | 5157989 | 5158026 | ? | - | predicted RNA | - | 1217 | 1185 | 1023 | 1130 | 1146 | 1395 | 1 |
| 4722 | - | 5158032 | 5158340 | + | - | *PA14_57950* | hypothetical protein | 2891 | 3039 | 2284 | 2085 | 2447 | 2783 | 1 |
| 4723 | - | 5158354 | 5158818 | + | *ptsN* | *PA14_57960* | nitrogen regulatory IIA protein | 565 | 706 | 591 | 418 | 552 | 545 | 1 |
| 4724 | 5159548 | 5158820 | 5159680 | + | - | *PA14_57970* | hypothetical protein | 235 | 261 | 249 | 192 | 221 | 243 | 1 |
| 4725 | 5159548 | 5159710 | 5159982 | + | - | *PA14_57980* | phosphoryl carrier protein | 106 | 111 | 98 | 73 | 79 | 81 | 1 |
| 4726 | - | 5161011 | 5160079 | - | - | *PA14_57990* | hypothetical protein | 18 | 36 | 21 | 113 | 31 | 16 | 0.347673727 |
| 4727 | - | 5161641 | 5161030 | - | *sodM* | *PA14_58000* | superoxide dismutase | 16 | 41 | 20 | 115 | 25 | 18 | 0.03981134 |
| 4728 | - | 5162103 | 5161654 | - | - | *PA14_58010* | hypothetical protein | 6 | 38 | 11 | 123 | 25 | 11 | 5.07146E-25 |
| 4729 | - | 5163507 | 5162131 | - | *fumC* | *PA14_58030* | fumarate hydratase | 17 | 69 | 17 | 279 | 41 | 20 | 4.3504E-06 |
| 4730 | - | 5163895 | 5163500 | - | - | *PA14_58040* | hypothetical protein | 2 | 94 | 4 | 245 | 77 | 10 | 0 |
| 4731 | - | 5165392 | 5164043 | - | *pmbA* | *PA14_58050* | PmbA protein | 111 | 116 | 115 | 108 | 114 | 118 | 1 |
| 4732 | - | 5165413 | 5166015 | + | - | *PA14_58060* | hypothetical protein | 236 | 238 | 322 | 271 | 267 | 288 | 1 |
| 4733 | - | 5167488 | 5166046 | - | - | *PA14_58070* | hypothetical protein | 270 | 253 | 309 | 252 | 251 | 284 | 1 |
| 4734 | - | 5168339 | 5167491 | - | - | *PA14_58080* | hypothetical protein | 274 | 270 | 419 | 255 | 315 | 279 | 1 |
| 4735 | - | 5172218 | 5168388 | - | - | *PA14_58090* | hypothetical protein | 112 | 88 | 111 | 115 | 107 | 118 | 1 |
| 4736 | - | 5173690 | 5172233 | - | *cafA* | *PA14_58100* | cytoplasmic axial filament protein | 166 | 142 | 175 | 180 | 180 | 189 | 1 |
| 4737 | - | 5174329 | 5173724 | - | - | *PA14_58110* | Maf-like protein | 139 | 106 | 161 | 157 | 120 | 149 | 1 |
| 4738 | - | 5174913 | 5174419 | - | *mreD* | *PA14_58120* | rod shape-determining protein MreD | 77 | 82 | 89 | 79 | 105 | 63 | 1 |
| 4739 | - | 5175905 | 5174913 | - | *mreC* | *PA14_58130* | rod shape-determining protein MreC | 54 | 47 | 57 | 48 | 56 | 42 | 1 |
| 4740 | - | 5177022 | 5175985 | - | *mreB* | *PA14_58150* | rod shape-determining protein MreB | 105 | 99 | 109 | 86 | 73 | 63 | 1 |
| 4741 | - | 5177262 | 5177549 | + | *gatC* | *PA14_58170* | aspartyl/glutamyl-tRNA amidotransferase subunit C | 67 | 74 | 74 | 53 | 56 | 43 | 1 |
| 4742 | - | 5177565 | 5179019 | + | *gatA* | *PA14_58180* | aspartyl/glutamyl-tRNA amidotransferase subunit A | 102 | 123 | 106 | 108 | 104 | 83 | 1 |
| 4743 | - | 5179126 | 5180571 | + | *gatB* | *PA14_58190* | aspartyl/glutamyl-tRNA amidotransferase subunit B | 123 | 139 | 112 | 105 | 111 | 89 | 1 |
| 4744 | - | 5180631 | 5181008 | + | - | *PA14_58210* | hypothetical protein | 42 | 39 | 39 | 29 | 36 | 30 | 1 |
| 4745 | - | 5181040 | 5181426 | + | - | *PA14_58220* | hypothetical protein | 71 | 70 | 64 | 56 | 69 | 63 | 1 |
| 4746 | - | 5182332 | 5181541 | - | - | *PA14_58230* | hypothetical protein | 230 | 204 | 180 | 173 | 218 | 191 | 1 |
| 4747 | - | 5183986 | 5182337 | - | - | *PA14_58240* | hypothetical protein | 108 | 72 | 95 | 75 | 96 | 86 | 1 |
| 4748 | - | 5188533 | 5183983 | - | - | *PA14_58250* | hypothetical protein | 110 | 87 | 93 | 88 | 93 | 88 | 1 |
| 4749 | - | 5189198 | 5188560 | - | - | *PA14_58260* | hypothetical protein | 80 | 67 | 71 | 74 | 78 | 71 | 1 |
| 4750 | - | 5190964 | 5189195 | - | - | *PA14_58270* | hypothetical protein | 165 | 126 | 129 | 133 | 141 | 116 | 1 |
| 4751 | - | 5191812 | 5191003 | - | - | *PA14_58290* | hypothetical protein | 78 | 65 | 71 | 74 | 55 | 61 | 1 |
| 4752 | - | 5192535 | 5191975 | - | - | *PA14_58300* | two-component response regulator | 307 | 360 | 303 | 273 | 329 | 268 | 1 |
| 4753 | - | 5193829 | 5192561 | - | - | *PA14_58320* | two-component sensor | 136 | 128 | 145 | 137 | 138 | 122 | 1 |
| 4754 | - | 5194106 | 5194816 | + | - | *PA14_58330* | hypothetical protein | 149 | 178 | 149 | 124 | 155 | 151 | 1 |
| 4755 | - | 5195069 | 5196682 | + | - | *PA14_58350* | ABC transporter substrate-binding protein | 75 | 125 | 103 | 125 | 103 | 105 | 1 |
| 4756 | - | 5196776 | 5198374 | + | - | *PA14_58360* | ABC transporter substrate-binding protein | 42 | 57 | 46 | 50 | 47 | 47 | 1 |
| 4757 | - | 5199659 | 5198442 | - | - | *PA14_58375* | metallopeptidase | 168 | 169 | 138 | 185 | 161 | 142 | 1 |
| 4758 | - | 5199828 | 5200391 | + | - | *PA14_58380* | transcriptional regulator | 221 | 180 | 265 | 311 | 220 | 276 | 1 |
| 4759 | - | 5200657 | 5202258 | + | - | *PA14_58390* | ABC transporter substrate-binding protein | 439 | 417 | 387 | 385 | 387 | 362 | 1 |
| 4760 | - | 5202483 | 5203937 | + | - | *PA14_58410* | porin | 95 | 185 | 125 | 90 | 127 | 88 | 1 |
| 4761 | - | 5203987 | 5205588 | + | - | *PA14_58420* | dipeptide ABC transporter substrate-binding protein | 106 | 200 | 111 | 66 | 113 | 79 | 1 |
| 4762 | - | 5205656 | 5206666 | + | - | *PA14_58440* | ABC transporter permease | 131 | 179 | 114 | 95 | 133 | 104 | 1 |
| 4763 | - | 5206678 | 5207589 | + | *dppC* | *PA14_58450* | ABC transporter permease | 125 | 164 | 100 | 87 | 113 | 93 | 1 |
| 4764 | - | 5207642 | 5208616 | + | *dppD* | *PA14_58470* | ABC transporter ATP-binding protein | 134 | 181 | 108 | 95 | 128 | 112 | 1 |
| 4765 | - | 5208616 | 5209587 | + | *dppF* | *PA14_58490* | dipeptide transporter ATP-binding subunit | 181 | 214 | 157 | 145 | 155 | 149 | 1 |
| 4766 | - | 5210252 | 5209620 | - | - | *PA14_58500* | hypothetical protein | 139 | 146 | 139 | 155 | 117 | 108 | 1 |
| 4767 | - | 5210394 | 5210867 | + | - | *PA14_58510* | AsnC family transcriptional regulator | 27 | 28 | 30 | 26 | 22 | 23 | 1 |
| 4768 | - | 5211801 | 5210872 | - | - | *PA14_58515* | hypothetical protein | 61 | 57 | 56 | 61 | 57 | 56 | 1 |
| 4769 | - | 5212475 | 5211798 | - | - | *PA14_58530* | hypothetical protein | 38 | 40 | 40 | 40 | 42 | 38 | 1 |
| 4770 | - | 5213227 | 5212472 | - | - | *PA14_58540* | hypothetical protein | 160 | 160 | 171 | 167 | 158 | 153 | 1 |
| 4771 | - | 5213357 | 5214256 | + | *lpxO1* | *PA14_58550* | lipopolysaccharide biosynthetic protein LpxO1 | 54 | 45 | 53 | 49 | 43 | 45 | 1 |
| 4772 | - | 5217106 | 5214554 | - | - | *PA14_58560* | oxidoreductase | 11 | 17 | 16 | 17 | 15 | 10 | 1 |
| 4773 | - | 5219583 | 5217322 | - | - | *PA14_58570* | outer membrane ferric siderophore receptor | 11 | 15 | 13 | 19 | 15 | 11 | 1 |
| 4774 | - | 5219801 | 5220481 | + | - | *PA14_58580* | hydroxylase | 18 | 63 | 25 | 76 | 74 | 25 | 0.000155454 |
| 4775 | - | 5220484 | 5221299 | + | - | *PA14_58600* | hypothetical protein | 29 | 49 | 32 | 89 | 54 | 34 | 0.76759159 |
| 4776 | - | 5221422 | 5223224 | + | - | *PA14_58610* | hypothetical protein | 19 | 15 | 19 | 17 | 14 | 14 | 1 |
| 4777 | - | 5223705 | 5223280 | - | - | *PA14_58620* | hypothetical protein | 20 | 14 | 17 | 18 | 17 | 14 | 1 |
| 4778 | - | 5224984 | 5223821 | - | - | *PA14_58630* | ornithine decarboxylase | 51 | 39 | 39 | 38 | 39 | 54 | 1 |
| 4779 | - | 5227457 | 5225436 | - | - | *PA14_58650* | chemotaxis transducer | 82 | 97 | 88 | 118 | 95 | 98 | 1 |
| 4780 | - | 5228483 | 5227647 | - | - | *PA14_58660* | hypothetical protein | 76 | 73 | 79 | 72 | 67 | 62 | 1 |
| 4781 | - | 5229046 | 5228480 | - | *ampD* | *PA14_58670* | N-acetyl-anhydromuranmyl-L-alanine amidase | 29 | 27 | 25 | 23 | 27 | 27 | 1 |
| 4782 | - | 5231452 | 5229197 | - | - | *PA14_58690* | hypothetical protein | 418 | 457 | 485 | 589 | 466 | 528 | 1 |
| 4783 | - | 5231738 | 5232586 | + | *nadC* | *PA14_58700* | nicotinate-nucleotide pyrophosphorylase | 51 | 47 | 48 | 45 | 50 | 42 | 1 |
| 4784 | - | 5232712 | 5232640 | - | - | *PA14_58710* | Thr tRNA | 8 | 11 | 15 | 8 | 11 | 8 | 1 |
| 4785 | - | 5233456 | 5232749 | - | - | *PA14_58720* | hypothetical protein | 18 | 19 | 26 | 16 | 27 | 12 | 1 |
| 4786 | - | 5234050 | 5233511 | - | *pilA* | *PA14_58730* | type IV pilin structural subunit | 882 | 2047 | 877 | 796 | 1568 | 662 | 1 |
| 4787 | - | 5234253 | 5234143 | - | - | *PA14_58740* | hypothetical protein | 2 | 4 | 3 | 3 | 4 | 2 | 1 |
| 4788 | - | 5234279 | 5235982 | + | *pilB* | *PA14_58750* | type 4 fimbrial biogenesis protein PilB | 87 | 101 | 79 | 78 | 82 | 75 | 1 |

| | A | B | C | D | E | F | G | H | I | J | K | L | M | N |
|---|---|---|---|---|---|---|---|---|---|---|---|---|---|---|
| 4789 | - | 5236082 | 5237203 | + | *pilC* | *PA14_58760* | type 4 fimbrial biogenesis protein pilC | 240 | 231 | 202 | 195 | 243 | 219 | 1 |
| 4790 | - | 5237206 | 5238078 | + | *pilD* | *PA14_58770* | type 4 prepilin peptidase PilD | 131 | 128 | 118 | 149 | 134 | 120 | 1 |
| 4791 | - | 5238075 | 5238686 | + | *coaE* | *PA14_58780* | dephospho-CoA kinase | 151 | 124 | 146 | 133 | 126 | 120 | 1 |
| 4792 | - | 5238683 | 5238883 | + | - | *PA14_58790* | zinc-binding protein | 128 | 125 | 127 | 146 | 120 | 133 | 1 |
| 4793 | - | 5239129 | 5238920 | - | - | *PA14_58800* | hypothetical protein | 103 | 80 | 101 | 92 | 72 | 91 | 1 |
| 4794 | - | 5239924 | 5239235 | - | - | *PA14_58810* | hypothetical protein | 41 | 50 | 40 | 46 | 48 | 44 | 1 |
| 4795 | - | 5240391 | 5239921 | - | - | *PA14_58820* | hypothetical protein | 39 | 48 | 39 | 43 | 49 | 41 | 1 |
| 4796 | - | 5240813 | 5240388 | - | - | *PA14_58830* | hypothetical protein | 49 | 51 | 50 | 50 | 56 | 44 | 1 |
| 4797 | - | 5240946 | 5241575 | + | - | *PA14_58840* | hypothetical protein | 131 | 133 | 138 | 154 | 150 | 168 | 1 |
| 4798 | - | 5241572 | 5242021 | + | - | *PA14_58850* | hypothetical protein | 118 | 115 | 126 | 123 | 127 | 140 | 1 |
| 4799 | - | 5242047 | 5242217 | + | - | *PA14_58860* | hypothetical protein | 48 | 44 | 44 | 33 | 42 | 48 | 1 |
| 4800 | - | 5242281 | 5243588 | + | *ndh* | *PA14_58870* | NADH dehydrogenase | 65 | 75 | 62 | 70 | 77 | 75 | 1 |
| 4801 | - | 5244719 | 5243628 | - | - | *PA14_58880* | ATPase | 40 | 39 | 36 | 43 | 35 | 34 | 1 |
| 4802 | - | 5245268 | 5246905 | + | - | *PA14_58890* | hemolysin activation/secretion protein | 5 | 5 | 6 | 7 | 5 | 6 | 1 |
| 4803 | - | 5246954 | 5251207 | + | - | *PA14_58900* | large exoprotein | 19 | 19 | 19 | 20 | 18 | 18 | 1 |
| 4804 | - | 5251580 | 5252446 | + | - | *PA14_58910* | chromosome partitioning-like protein | 9 | 7 | 7 | 5 | 6 | 5 | 1 |
| 4805 | - | 5252505 | 5253185 | + | - | *PA14_58920* | hypothetical protein | 13 | 11 | 10 | 11 | 11 | 9 | 1 |
| 4806 | - | 5253182 | 5253679 | + | - | *PA14_58930* | hypothetical protein | 14 | 11 | 10 | 11 | 10 | 9 | 1 |
| 4807 | - | 5253676 | 5253825 | + | - | *PA14_58940* | hypothetical protein | 14 | 15 | 19 | 15 | 13 | 16 | 1 |
| 4808 | - | 5253822 | 5254508 | + | - | *PA14_58950* | hypothetical protein | 11 | 9 | 10 | 8 | 9 | 8 | 1 |
| 4809 | - | 5254508 | 5255209 | + | - | *PA14_58960* | hypothetical protein | 13 | 12 | 15 | 15 | 12 | 12 | 1 |
| 4810 | - | 5255206 | 5255913 | + | - | *PA14_58970* | hypothetical protein | 8 | 9 | 7 | 8 | 7 | 8 | 1 |
| 4811 | - | 5255897 | 5256121 | + | - | *PA14_58980* | hypothetical protein | 9 | 6 | 6 | 8 | 7 | 5 | 1 |
| 4812 | - | 5256118 | 5257464 | + | - | *PA14_58990* | DNA helicase | 12 | 11 | 9 | 10 | 10 | 12 | 1 |
| 4813 | - | 5257636 | 5258163 | + | - | *PA14_59000* | hypothetical protein | 9 | 10 | 7 | 8 | 7 | 10 | 1 |
| 4814 | - | 5258160 | 5258417 | + | - | *PA14_59010* | hypothetical protein | 6 | 12 | 10 | 10 | 7 | 8 | 0.831724413 |
| 4815 | - | 5258410 | 5258904 | + | - | *PA14_59020* | hypothetical protein | 22 | 20 | 19 | 22 | 15 | 17 | 1 |
| 4816 | - | 5258897 | 5259130 | + | - | *PA14_59030* | hypothetical protein | 28 | 27 | 18 | 31 | 29 | 30 | 1 |
| 4817 | - | 5259130 | 5260146 | + | - | *PA14_59050* | nucleoid-associated protein NdpA | 56 | 53 | 56 | 60 | 59 | 61 | 1 |
| 4818 | - | 5260143 | 5260397 | + | - | *PA14_59060* | hypothetical protein | 27 | 31 | 23 | 31 | 30 | 24 | 1 |
| 4819 | - | 5260394 | 5262133 | + | - | *PA14_59070* | hypothetical protein | 27 | 24 | 21 | 24 | 25 | 24 | 1 |
| 4820 | - | 5262161 | 5262928 | + | - | *PA14_59090* | hypothetical protein | 28 | 32 | 26 | 30 | 30 | 30 | 1 |
| 4821 | - | 5262925 | 5264250 | + | - | *PA14_59100* | hypothetical protein | 17 | 16 | 15 | 15 | 16 | 15 | 1 |
| 4822 | - | 5264247 | 5264522 | + | - | *PA14_59110* | hypothetical protein | 22 | 24 | 23 | 22 | 28 | 23 | 1 |
| 4823 | - | 5264513 | 5264752 | + | - | *PA14_59120* | hypothetical protein | 26 | 23 | 24 | 22 | 23 | 17 | 1 |
| 4824 | - | 5265022 | 5265750 | + | - | *PA14_59130* | hypothetical protein | 21 | 22 | 17 | 16 | 15 | 16 | 1 |
| 4825 | - | 5265756 | 5266289 | + | - | *PA14_59140* | hypothetical protein | 22 | 21 | 16 | 17 | 18 | 15 | 1 |
| 4826 | - | 5266303 | 5266773 | + | - | *PA14_59150* | single-stranded DNA-binding protein | 104 | 103 | 73 | 74 | 87 | 91 | 1 |
| 4827 | - | 5267237 | 5267482 | + | - | *PA14_59160* | hypothetical protein | 41 | 40 | 35 | 36 | 38 | 36 | 1 |
| 4828 | - | 5267854 | 5267621 | - | - | *PA14_59170* | hypothetical protein | 206 | 220 | 173 | 158 | 160 | 165 | 1 |
| 4829 | 5268881 | 5268174 | 5270099 | + | - | *PA14_59180* | topoisomerase I - like protein | 63 | 74 | 87 | 129 | 87 | 91 | 1 |
| 4830 | - | 5270173 | 5270643 | + | - | *PA14_59190* | hypothetical protein | 147 | 139 | 134 | 126 | 119 | 133 | 1 |
| 4831 | - | 5270777 | 5272666 | + | - | *PA14_59200* | hypothetical protein | 18 | 16 | 17 | 18 | 15 | 14 | 1 |
| 4832 | - | 5272663 | 5274636 | + | - | *PA14_59210* | DNA helicase | 14 | 13 | 11 | 13 | 11 | 12 | 1 |
| 4833 | - | 5275324 | 5276820 | + | - | *PA14_59220* | pyocin S5 | 114 | 263 | 149 | 207 | 243 | 216 | 0.925024982 |
| 4834 | - | 5277171 | 5276845 | - | - | *PA14_59230* | colicin immunity protein | 10 | 8 | 10 | 11 | 20 | 14 | 1 |
| 4835 | - | 5277443 | 5278567 | + | *pilL2* | *PA14_59240* | type IV B pilus protein | 16 | 11 | 15 | 14 | 13 | 13 | 1 |
| 4836 | - | 5278639 | 5280276 | + | *pilN2* | *PA14_59250* | Type IV B pilus protein | 11 | 9 | 9 | 11 | 8 | 10 | 1 |
| 4837 | - | 5280280 | 5281605 | + | *pilO2* | *PA14_59270* | type IV b pilus protein | 15 | 14 | 15 | 16 | 15 | 14 | 1 |
| 4838 | - | 5281595 | 5282128 | + | *pilP2* | *PA14_59280* | type IV B pilus protein | 9 | 10 | 10 | 9 | 9 | 9 | 1 |
| 4839 | - | 5282137 | 5283717 | + | *pilQ2* | *PA14_59290* | type IV B pilus protein | 10 | 9 | 8 | 7 | 7 | 9 | 1 |
| 4840 | - | 5283717 | 5284796 | + | *pilR2* | *PA14_59310* | type IV B pilus protein | 11 | 9 | 10 | 12 | 9 | 11 | 1 |
| 4841 | - | 5284818 | 5285348 | + | *pilS2* | *PA14_59320* | type IV B pilus protein | 31 | 33 | 27 | 32 | 27 | 26 | 1 |
| 4842 | - | 5285345 | 5286286 | + | *pilT2* | *PA14_59340* | type IV B pilus protein | 20 | 15 | 17 | 15 | 14 | 16 | 1 |
| 4843 | - | 5286291 | 5287619 | + | *pilV2* | *PA14_59350* | Type IV B pilus protein | 30 | 29 | 30 | 28 | 26 | 28 | 1 |
| 4844 | - | 5287648 | 5288085 | + | *pilM2* | *PA14_59360* | type IV B pilus protein | 43 | 44 | 49 | 39 | 49 | 45 | 1 |
| 4845 | - | 5289000 | 5289566 | + | - | *PA14_59380* | hypothetical protein | 257 | 206 | 132 | 175 | 190 | 253 | 1 |
| 4846 | - | 5289001 | 5288357 | - | - | *PA14_59370* | hypothetical protein | 84 | 71 | 57 | 76 | 76 | 81 | 1 |
| 4847 | - | 5289818 | 5289952 | + | - | *PA14_59390* | hypothetical protein | 173 | 240 | 130 | 128 | 207 | 159 | 1 |
| 4848 | - | 5290227 | 5290766 | + | - | *PA14_59400* | hypothetical protein | 410 | 284 | 202 | 234 | 236 | 348 | 1 |
| 4849 | - | 5291081 | 5290815 | - | - | *PA14_59410* | hypothetical protein | 288 | 191 | 117 | 163 | 170 | 236 | 1 |
| 4850 | - | 5291101 | 5291337 | + | - | *PA14_59430* | hypothetical protein | 126 | 104 | 58 | 70 | 73 | 102 | 1 |
| 4851 | - | 5291822 | 5292529 | + | - | *PA14_59440* | hypothetical protein | 32 | 21 | 19 | 25 | 24 | 38 | 1 |
| 4852 | - | 5292765 | 5293115 | + | - | *PA14_59470* | hypothetical protein | 271 | 198 | 190 | 185 | 160 | 265 | 1 |
| 4853 | - | 5293173 | 5293808 | + | - | *PA14_59480* | hypothetical protein | 57 | 51 | 39 | 45 | 43 | 57 | 1 |
| 4854 | - | 5293805 | 5294080 | + | - | *PA14_59490* | hypothetical protein | 67 | 65 | 49 | 56 | 51 | 70 | 1 |
| 4855 | - | 5294150 | 5294512 | + | - | *PA14_59500* | hypothetical protein | 40 | 35 | 25 | 30 | 29 | 38 | 1 |
| 4856 | - | 5294580 | 5294834 | + | - | *PA14_59510* | hypothetical protein | 34 | 34 | 19 | 32 | 33 | 36 | 1 |
| 4857 | - | 5294926 | 5295531 | + | - | *PA14_59520* | hypothetical protein | 19 | 18 | 16 | 18 | 17 | 23 | 1 |
| 4858 | - | 5295561 | 5297012 | + | - | *PA14_59530* | hypothetical protein | 13 | 10 | 12 | 11 | 12 | 12 | 1 |
| 4859 | - | 5297117 | 5299366 | + | - | *PA14_59540* | hypothetical protein | 12 | 9 | 10 | 8 | 7 | 8 | 1 |
| 4860 | - | 5299690 | 5300916 | + | - | *PA14_59550* | hypothetical protein | 35 | 40 | 23 | 17 | 30 | 21 | 1 |
| 4861 | - | 5301173 | 5301439 | + | - | *PA14_59560* | transposase | 3 | 4 | 5 | 5 | 3 | 4 | 1 |
| 4862 | - | 5301490 | 5301825 | + | - | *PA14_59570* | transposase | 84 | 72 | 68 | 88 | 63 | 70 | 1 |
| 4863 | - | 5301889 | 5303424 | + | - | *PA14_59580* | transposase | 115 | 98 | 95 | 108 | 98 | 96 | 1 |
| 4864 | - | 5303485 | 5303844 | + | - | *PA14_59590* | hypothetical protein | 50 | 41 | 36 | 36 | 34 | 30 | 1 |
| 4865 | - | 5304162 | 5303917 | - | - | *PA14_59600* | hypothetical protein | 70 | 41 | 54 | 38 | 35 | 34 | 0.739269516 |
| 4866 | - | 5304920 | 5304246 | - | - | *PA14_59610* | hypothetical protein | 49 | 54 | 37 | 46 | 51 | 48 | 1 |
| 4867 | - | 5305668 | 5305003 | - | - | *PA14_59620* | hypothetical protein | 103 | 105 | 88 | 92 | 85 | 93 | 1 |
| 4868 | - | 5306710 | 5305679 | - | - | *PA14_59630* | hypothetical protein | 15 | 18 | 13 | 10 | 13 | 12 | 1 |
| 4869 | - | 5307321 | 5308010 | + | - | *PA14_59640* | hypothetical protein | 22 | 18 | 22 | 22 | 19 | 17 | 1 |
| 4870 | - | 5308021 | 5308776 | + | - | *PA14_59650* | hypothetical protein | 15 | 14 | 11 | 13 | 12 | 9 | 1 |
| 4871 | - | 5308761 | 5309342 | + | - | *PA14_59660* | hypothetical protein | 36 | 40 | 32 | 39 | 32 | 34 | 1 |
| 4872 | - | 5309339 | 5309839 | + | - | *PA14_59670* | hypothetical protein | 10 | 9 | 9 | 9 | 9 | 11 | 1 |

| | A | B | C | D | E | F | G | H | I | J | K | L | M | N |
|---|---|---|---|---|---|---|---|---|---|---|---|---|---|---|
| 4873 | - | 5309848 | 5310105 | + | - | *PA14_59680* | hypothetical protein | 9 | 9 | 10 | 9 | 11 | 7 | 1 |
| 4874 | - | 5310109 | 5312343 | + | - | *PA14_59690* | hypothetical protein | 13 | 12 | 11 | 15 | 12 | 14 | 1 |
| 4875 | - | 5312343 | 5313089 | + | - | *PA14_59700* | hypothetical protein | 22 | 16 | 17 | 23 | 16 | 18 | 1 |
| 4876 | - | 5313527 | 5314075 | + | *cupD1* | *PA14_59710* | fimbrial protein | 15 | 19 | 13 | 12 | 13 | 9 | 1 |
| 4877 | - | 5314164 | 5314910 | + | *cupD2* | *PA14_59720* | pili assembly chaperone | 5 | 6 | 5 | 6 | 6 | 4 | 1 |
| 4878 | - | 5317504 | 5318850 | + | *cupD4* | *PA14_59750* | fimbrial subunit | 14 | 13 | 11 | 12 | 12 | 12 | 1 |
| 4879 | - | 5318840 | 5319556 | + | *cupD5* | *PA14_59760* | pili assembly chaperone | 40 | 37 | 47 | 40 | 30 | 32 | 1 |
| 4880 | - | 5320295 | 5319600 | - | *rcsB* | *PA14_59770* | two component response regulator | 99 | 109 | 112 | 76 | 88 | 88 | 1 |
| 4881 | - | 5323549 | 5320295 | - | *rcsC* | *PA14_59780* | kinase sensor protein | 111 | 105 | 153 | 71 | 68 | 90 | 1 |
| 4882 | - | 5324829 | 5323630 | - | *pvrR* | *PA14_59790* | two component response regulator | 332 | 414 | 414 | 139 | 127 | 248 | 1 |
| 4883 | - | 5327621 | 5324826 | - | - | *PA14_59800* | kinase sensor protein | 37 | 31 | 34 | 38 | 31 | 35 | 1 |
| 4884 | - | 5327953 | 5328603 | + | - | *PA14_59820* | hypothetical protein | 28 | 22 | 21 | 31 | 20 | 24 | 1 |
| 4885 | - | 5328614 | 5330095 | + | - | *PA14_59830* | helicase | 28 | 35 | 30 | 39 | 33 | 30 | 1 |
| 4886 | - | 5330226 | 5331335 | + | - | *PA14_59840* | hypothetical protein | 344 | 383 | 341 | 130 | 198 | 303 | 1 |
| 4887 | - | 5331690 | 5331379 | - | - | *PA14_59845* | hypothetical protein | 89 | 108 | 100 | 94 | 100 | 79 | 1 |
| 4888 | - | 5331863 | 5332168 | + | - | *PA14_59850* | hypothetical protein | 5 | 7 | 4 | 6 | 4 | 6 | 1 |
| 4889 | - | 5332241 | 5332600 | + | - | *PA14_59860* | type III effector Hop protein | 12 | 9 | 14 | 12 | 11 | 12 | 1 |
| 4890 | - | 5332597 | 5332836 | + | - | *PA14_59870* | hypothetical protein | 12 | 10 | 13 | 11 | 9 | 11 | 1 |
| 4891 | - | 5332854 | 5333210 | + | - | *PA14_59880* | hypothetical protein | 10 | 10 | 8 | 10 | 6 | 8 | 1 |
| 4892 | - | 5333221 | 5333607 | + | - | *PA14_59890* | hypothetical protein | 24 | 17 | 19 | 19 | 18 | 23 | 1 |
| 4893 | - | 5333604 | 5334263 | + | - | *PA14_59900* | hypothetical protein | 18 | 16 | 14 | 16 | 15 | 16 | 1 |
| 4894 | - | 5334260 | 5335144 | + | - | *PA14_59910* | hypothetical protein | 28 | 24 | 31 | 37 | 24 | 26 | 1 |
| 4895 | - | 5335128 | 5336633 | + | - | *PA14_59920* | hypothetical protein | 13 | 11 | 10 | 13 | 12 | 12 | 1 |
| 4896 | - | 5336611 | 5337054 | + | - | *PA14_59930* | hypothetical protein | 20 | 15 | 11 | 17 | 15 | 18 | 1 |
| 4897 | - | 5337054 | 5339996 | + | - | *PA14_59940* | hypothetical protein | 20 | 18 | 18 | 19 | 17 | 19 | 1 |
| 4898 | - | 5339993 | 5340277 | + | - | *PA14_59950* | hypothetical protein | 17 | 13 | 11 | 11 | 9 | 12 | 1 |
| 4899 | - | 5340274 | 5340933 | + | - | *PA14_59960* | protein-disulfide isomerase | 111 | 96 | 95 | 84 | 108 | 85 | 1 |
| 4900 | - | 5341148 | 5340930 | - | - | *PA14_59970* | hypothetical protein | 267 | 230 | 248 | 219 | 290 | 205 | 1 |
| 4901 | - | 5341357 | 5341788 | + | - | *PA14_59980* | hypothetical protein | 4 | 3 | 4 | 3 | 4 | 3 | 1 |
| 4902 | - | 5341788 | 5342726 | + | - | *PA14_59990* | hypothetical protein | 7 | 5 | 7 | 5 | 6 | 6 | 0.80035376 |
| 4903 | - | 5342744 | 5344126 | + | - | *PA14_60000* | hypothetical protein | 11 | 9 | 10 | 10 | 10 | 9 | 1 |
| 4904 | - | 5344126 | 5344467 | + | - | *PA14_60010* | hypothetical protein | 11 | 12 | 10 | 11 | 9 | 9 | 1 |
| 4905 | - | 5344464 | 5345975 | + | - | *PA14_60020* | hypothetical protein | 19 | 15 | 15 | 14 | 20 | 19 | 1 |
| 4906 | - | 5346345 | 5346001 | - | - | *PA14_60030* | hypothetical protein | 178 | 161 | 170 | 183 | 247 | 226 | 1 |
| 4907 | - | 5346464 | 5346736 | + | - | *PA14_60040* | hypothetical protein | 81 | 85 | 64 | 64 | 65 | 65 | 1 |
| 4908 | - | 5346740 | 5347090 | + | - | *PA14_60050* | plasmid stablization protein | 107 | 101 | 90 | 114 | 95 | 93 | 1 |
| 4909 | - | 5347532 | 5347834 | + | - | *PA14_60060* | hypothetical protein | 6 | 9 | 5 | 6 | 7 | 6 | 1 |
| 4910 | - | 5348012 | 5350435 | + | - | *PA14_60070* | hypothetical protein | 98 | 81 | 75 | 98 | 84 | 90 | 1 |
| 4911 | - | 5350432 | 5352363 | + | - | *PA14_60080* | hypothetical protein | 208 | 176 | 158 | 225 | 183 | 226 | 1 |
| 4912 | - | 5352790 | 5353305 | + | - | *PA14_60090* | hypothetical protein | 106 | 91 | 85 | 78 | 115 | 93 | 1 |
| 4913 | - | 5354049 | 5353510 | - | *dtd* | *PA14_60100* | deoxycytidine triphosphate deaminase | 100 | 97 | 77 | 68 | 82 | 68 | 1 |
| 4914 | - | 5354756 | 5354046 | - | - | *PA14_60110* | hypothetical protein | 108 | 78 | 80 | 87 | 92 | 96 | 1 |
| 4915 | - | 5355289 | 5354756 | - | *dcd2* | *PA14_60120* | deoxycytidine deaminase | 121 | 86 | 95 | 90 | 85 | 89 | 1 |
| 4916 | - | 5355824 | 5357743 | + | - | *PA14_60130* | hypothetical protein | 41 | 28 | 33 | 28 | 24 | 29 | 1 |
| 4917 | - | 5357740 | 5358660 | + | - | *PA14_60140* | xerD-like integrase | 54 | 34 | 33 | 30 | 32 | 30 | 1 |
| 4918 | - | 5359468 | 5359396 | - | - | *PA14_60150* | Lys tRNA | 87 | 68 | 122 | 64 | 36 | 38 | 1 |
| 4919 | - | 5359555 | 5359482 | - | - | *PA14_60160* | Pro tRNA | 86 | 60 | 119 | 73 | 38 | 30 | 0.87182855 |
| 4920 | - | 5359640 | 5359568 | - | - | *PA14_60180* | Asn tRNA | 118 | 98 | 187 | 110 | 53 | 42 | 1 |
| 4921 | - | 5362439 | 5359875 | - | *clpB* | *PA14_60190* | clpB protein | 531 | 526 | 721 | 250 | 477 | 386 | 1 |
| 4922 | - | 5363323 | 5362595 | - | - | *PA14_60200* | hypothetical protein | 87 | 71 | 103 | 85 | 78 | 70 | 1 |
| 4923 | - | 5364282 | 5363320 | - | *rluD* | *PA14_60210* | pseudouridine synthase | 62 | 48 | 65 | 58 | 56 | 51 | 1 |
| 4924 | - | 5364428 | 5365453 | + | *comL* | *PA14_60230* | competence protein ComL | 33 | 22 | 33 | 22 | 18 | 22 | 1 |
| 4925 | - | 5365571 | 5365855 | + | - | *PA14_60240* | hypothetical protein | 17 | 18 | 17 | 15 | 8 | 12 | 1 |
| 4926 | - | 5365845 | 5367437 | + | *pilS* | *PA14_60250* | two-component sensor PilS | 28 | 25 | 30 | 32 | 25 | 26 | 1 |
| 4927 | - | 5367452 | 5368789 | + | *pilR* | *PA14_60260* | two-component response regulator PilR | 52 | 66 | 52 | 68 | 56 | 50 | 1 |
| 4928 | - | 5369929 | 5368835 | - | - | *PA14_60270* | glycine/D-amino acid oxidases | 104 | 93 | 106 | 89 | 101 | 91 | 1 |
| 4929 | - | 5370692 | 5371180 | + | *fimU* | *PA14_60280* | type 4 fimbrial biogenesis protein FimU | 70 | 54 | 59 | 67 | 58 | 56 | 1 |
| 4930 | - | 5371711 | 5372538 | + | *pilW* | *PA14_60290* | type 4 fimbrial biogenesis protein PilW | 51 | 50 | 47 | 48 | 44 | 45 | 1 |
| 4931 | - | 5372541 | 5373140 | + | *pilX* | *PA14_60300* | type 4 fimbrial biogenesis protein PilX | 71 | 73 | 65 | 70 | 66 | 66 | 1 |
| 4932 | - | 5373154 | 5376630 | + | *pilY1* | *PA14_60310* | type 4 fimbrial biogenesis protein PilY1 | 137 | 151 | 119 | 139 | 145 | 144 | 1 |
| 4933 | - | 5376956 | 5377381 | + | *pilE* | *PA14_60320* | type 4 fimbrial biogenesis protein PilE | 66 | 76 | 57 | 47 | 58 | 59 | 1 |
| 4934 | - | 5378394 | 5377450 | - | *ispH* | *PA14_60330* | 4-hydroxy-3-methylbut-2-enyl diphosphate reductase | 100 | 78 | 88 | 76 | 84 | 78 | 1 |
| 4935 | - | 5378920 | 5378480 | - | - | *PA14_60350* | peptidyl-prolyl cis-trans isomerase, FkbP-type | 172 | 169 | 148 | 135 | 145 | 118 | 1 |
| 4936 | - | 5379422 | 5378913 | - | *lspA* | *PA14_60360* | lipoprotein signal peptidase | 150 | 150 | 139 | 116 | 122 | 107 | 1 |
| 4937 | - | 5382246 | 5379415 | - | *ileS* | *PA14_60370* | isoleucyl-tRNA synthetase | 205 | 218 | 197 | 201 | 203 | 178 | 1 |
| 4938 | - | 5383208 | 5382270 | - | *ribF* | *PA14_60380* | bifunctional riboflavin kinase/FMN adenylyltransferase | 74 | 54 | 74 | 56 | 50 | 56 | 1 |
| 4939 | - | 5384843 | 5383305 | - | - | *PA14_60390* | virulence factor, membrane protein | 16 | 13 | 17 | 14 | 15 | 13 | 1 |
| 4940 | - | 5385127 | 5385402 | + | *rpsT* | *PA14_60400* | 30S ribosomal protein S20 | 65 | 74 | 75 | 46 | 44 | 41 | 1 |
| 4941 | - | 5385933 | 5385469 | - | - | *PA14_60410* | hypothetical protein | 248 | 245 | 263 | 273 | 274 | 225 | 1 |
| 4942 | - | 5387063 | 5385945 | - | *proB* | *PA14_60420* | gamma-glutamyl kinase | 424 | 413 | 543 | 489 | 466 | 425 | 1 |
| 4943 | - | 5388354 | 5387134 | - | *obgE* | *PA14_60445* | GTPase ObgE | 137 | 117 | 147 | 109 | 112 | 104 | 1 |
| 4944 | - | 5388754 | 5388497 | - | *rpmA* | *PA14_60450* | 50S ribosomal protein L27 | 147 | 193 | 154 | 92 | 100 | 68 | 1 |
| 4945 | - | 5389089 | 5388778 | - | *rplU* | *PA14_60460* | 50S ribosomal protein L21 | 219 | 236 | 247 | 140 | 168 | 130 | 1 |
| 4946 | - | 5389330 | 5390298 | + | *ispB* | *PA14_60470* | octaprenyl-diphosphate synthase | 47 | 46 | 54 | 49 | 39 | 37 | 1 |
| 4947 | - | 5390436 | 5390660 | + | - | *PA14_60480* | hypothetical protein | 6 | 41 | 6 | 129 | 23 | 7 | 4.82069E-19 |
| 4948 | - | 5391003 | 5393030 | + | - | *PA14_60490* | cytochrome c | 110 | 91 | 120 | 62 | 88 | 83 | 1 |
| 4949 | - | 5393717 | 5393100 | - | *fklB* | *PA14_60500* | peptidyl-prolyl cis-trans isomerase FklB | 98 | 104 | 128 | 113 | 103 | 102 | 1 |
| 4950 | - | 5394107 | 5393802 | - | - | *PA14_60520* | hypothetical protein | 198 | 242 | 409 | 341 | 245 | 305 | 1 |
| 4951 | - | 5394852 | 5394364 | - | - | *PA14_60530* | hypothetical protein | 12 | 12 | 13 | 9 | 10 | 12 | 1 |
| 4952 | - | 5395367 | 5395026 | - | - | *PA14_60540* | hypothetical protein | 226 | 228 | 196 | 199 | 212 | 226 | 1 |
| 4953 | - | 5397973 | 5395520 | - | - | *PA14_60550* | ATP-dependent protease | 127 | 145 | 134 | 121 | 132 | 121 | 1 |
| 4954 | - | 5398398 | 5398072 | - | - | *PA14_60560* | hypothetical protein | 548 | 517 | 687 | 384 | 480 | 546 | 1 |
| 4955 | - | 5398633 | 5399121 | + | - | *PA14_60570* | hypothetical protein | 506 | 869 | 331 | 200 | 568 | 273 | 1 |
| 4956 | - | 5399180 | 5401036 | + | - | *PA14_60580* | hypothetical protein | 49 | 66 | 42 | 41 | 53 | 41 | 1 |

| | A | B | C | D | E | F | G | H | I | J | K | L | M | N |
|---|---|---|---|---|---|---|---|---|---|---|---|---|---|---|
| 4957 | - | 5401584 | 5401030 | - | - | *PA14_60590* | hypothetical protein | 75 | 72 | 67 | 72 | 70 | 68 | 1 |
| 4958 | - | 5403235 | 5401640 | - | *rtcR* | *PA14_60600* | transcriptional regulator RtcR | 37 | 37 | 34 | 39 | 35 | 32 | 1 |
| 4959 | - | 5403550 | 5403623 | + | - | *PA14_60620* | Arg tRNA | 62 | 125 | 97 | 75 | 79 | 82 | 0.263777066 |
| 4960 | - | 5403895 | 5405040 | + | - | *PA14_60630* | hypothetical protein | 9 | 10 | 9 | 9 | 9 | 8 | 1 |
| 4961 | - | 5405096 | 5406310 | + | - | *PA14_60650* | hypothetical protein | 16 | 16 | 16 | 16 | 15 | 13 | 1 |
| 4962 | - | 5406347 | 5407159 | + | - | *PA14_60660* | nucleotidyltransferase | 36 | 34 | 39 | 36 | 27 | 32 | 1 |
| 4963 | - | 5407156 | 5408181 | + | *rtcA* | *PA14_60670* | RNA 3'-terminal-phosphate cyclase | 27 | 23 | 24 | 23 | 22 | 24 | 1 |
| 4964 | - | 5408283 | 5408660 | + | - | *PA14_60690* | lipoprotein | 8 | 7 | 10 | 9 | 6 | 6 | 1 |
| 4965 | - | 5408784 | 5409824 | + | *ccpR* | *PA14_60700* | cytochrome c551 peroxidase | 563 | 618 | 457 | 167 | 353 | 251 | 1 |
| 4966 | - | 5411222 | 5409885 | - | *gdhA* | *PA14_60710* | glutamate dehydrogenase | 43 | 24 | 26 | 23 | 25 | 27 | 1 |
| 4967 | - | 5412909 | 5411518 | - | - | *PA14_60730* | outer membrane protein | 22 | 27 | 19 | 26 | 25 | 28 | 1 |
| 4968 | - | 5413657 | 5413169 | - | *pra* | *PA14_60750* | protein activator | 198 | 174 | 103 | 174 | 215 | 273 | 1 |
| 4969 | - | 5415532 | 5414255 | - | - | *PA14_60760* | hypothetical protein | 49 | 49 | 47 | 61 | 58 | 62 | 1 |
| 4970 | - | 5417003 | 5415522 | - | - | *PA14_60770* | outer membrane protein | 45 | 48 | 41 | 58 | 57 | 55 | 1 |
| 4971 | - | 5418189 | 5416996 | - | - | *PA14_60780* | ABC transporter permease | 37 | 42 | 39 | 48 | 41 | 37 | 1 |
| 4972 | - | 5418872 | 5418186 | - | - | *PA14_60790* | ABC transporter ATP-binding protein | 71 | 73 | 72 | 82 | 69 | 75 | 1 |
| 4973 | - | 5421040 | 5419376 | - | - | *PA14_60800* | ABC transporter ATP-binding protein | 191 | 240 | 168 | 169 | 204 | 168 | 1 |
| 4974 | - | 5421533 | 5422051 | + | - | *PA14_60810* | transcriptional regulator NfxB | 139 | 159 | 174 | 147 | 202 | 230 | 1 |
| 4975 | - | 5423541 | 5422102 | - | *oprJ* | *PA14_60820* | outer membrane protein OprJ | 29 | 32 | 31 | 24 | 33 | 29 | 1 |
| 4976 | - | 5426678 | 5423547 | - | *mexD* | *PA14_60830* | multidrug efflux RND transporter MexD | 23 | 26 | 24 | 19 | 24 | 15 | 1 |
| 4977 | - | 5427869 | 5426706 | - | *mexC* | *PA14_60850* | multidrug efflux RND membrane fusion protein | 45 | 54 | 58 | 37 | 54 | 31 | 1 |
| 4978 | - | 5428030 | 5428593 | + | *nfxB* | *PA14_60860* | transcriptional regulator NfxB | 33 | 54 | 46 | 34 | 55 | 36 | 0.802744579 |
| 4979 | - | 5428837 | 5433084 | + | *morA* | *PA14_60870* | motility regulator | 73 | 66 | 79 | 76 | 81 | 75 | 1 |
| 4980 | - | 5433207 | 5434460 | + | *glyA* | *PA14_60890* | serine hydroxymethyltransferase | 42 | 42 | 43 | 33 | 44 | 31 | 1 |
| 4981 | - | 5434918 | 5434538 | - | - | *PA14_60900* | hypothetical protein | 152 | 159 | 184 | 215 | 197 | 186 | 1 |
| 4982 | - | 5435933 | 5434929 | - | - | *PA14_60920* | hypothetical protein | 92 | 139 | 122 | 138 | 128 | 153 | 1 |
| 4983 | - | 5436196 | 5435993 | - | - | *PA14_60930* | hypothetical protein | 244 | 279 | 242 | 281 | 311 | 343 | 1 |
| 4984 | - | 5438311 | 5436245 | - | - | *PA14_60950* | hypothetical protein | 442 | 462 | 436 | 477 | 547 | 540 | 1 |
| 4985 | - | 5439155 | 5438649 | - | - | *PA14_60960* | hypothetical protein | 5957 | 6590 | 6005 | 8958 | 6174 | 9428 | 1 |
| 4986 | - | 5439358 | 5439735 | + | - | *PA14_60970* | hypothetical protein | 312 | 336 | 369 | 343 | 364 | 354 | 1 |
| 4987 | - | 5441103 | 5439742 | - | *radA* | *PA14_60990* | DNA repair protein RadA | 80 | 85 | 89 | 88 | 83 | 78 | 1 |
| 4988 | - | 5441213 | 5441647 | + | - | *PA14_61000* | hypothetical protein | 220 | 178 | 173 | 247 | 345 | 239 | 1 |
| 4989 | - | 5441963 | 5441709 | - | - | *PA14_61010* | hypothetical protein | 2142 | 2432 | 2595 | 1450 | 1361 | 2382 | 1 |
| 4990 | - | 5442589 | 5442038 | - | - | *PA14_61020* | hypothetical protein | 80 | 28 | 29 | 37 | 28 | 69 | 0.006435902 |
| 4991 | - | 5444184 | 5442643 | - | *katB* | *PA14_61040* | catalase | 88 | 15 | 14 | 19 | 13 | 68 | 1.51881E-11 |
| 4992 | - | 5444712 | 5445125 | + | *mscL* | *PA14_61050* | large-conductance mechanosensitive channel | 445 | 543 | 517 | 431 | 456 | 476 | 1 |
| 4993 | - | 5446006 | 5445230 | - | - | *PA14_61060* | oxidoreductase | 58 | 65 | 47 | 68 | 86 | 99 | 1 |
| 4994 | - | 5447111 | 5446113 | - | - | *PA14_61080* | c4-dicarboxylate-binding protein | 10 | 12 | 10 | 11 | 8 | 8 | 1 |
| 4995 | - | 5447439 | 5448563 | + | - | *PA14_61090* | hypothetical protein | 46 | 42 | 48 | 40 | 44 | 39 | 1 |
| 4996 | - | 5449142 | 5448564 | - | - | *PA14_61110* | hypothetical protein | 86 | 77 | 93 | 80 | 81 | 83 | 1 |
| 4997 | - | 5450110 | 5449139 | - | - | *PA14_61120* | hypothetical protein | 181 | 150 | 181 | 149 | 155 | 165 | 1 |
| 4998 | - | 5451361 | 5450114 | - | - | *PA14_61130* | c-type cytochrome | 44 | 39 | 43 | 35 | 38 | 48 | 1 |
| 4999 | - | 5451903 | 5451364 | - | - | *PA14_61140* | hypothetical protein | 33 | 37 | 37 | 33 | 39 | 48 | 1 |
| 5000 | - | 5454727 | 5451896 | - | - | *PA14_61150* | oxidoreductase | 51 | 50 | 60 | 51 | 50 | 63 | 1 |
| 5001 | - | 5455038 | 5456249 | + | - | *PA14_61170* | transmembrane protein | 3 | 3 | 4 | 3 | 4 | 3 | 1 |
| 5002 | - | 5456798 | 5456409 | - | - | *PA14_61180* | hypothetical protein | 3 | 5 | 3 | 6 | 6 | 5 | 1 |
| 5003 | - | 5458807 | 5457101 | - | - | *PA14_61190* | hypothetical protein | 96 | 63 | 86 | 78 | 76 | 101 | 1 |
| 5004 | - | 5464854 | 5458870 | - | - | *PA14_61200* | hypothetical protein | 58 | 20 | 29 | 21 | 20 | 27 | 0.615940863 |
| 5005 | - | 5465175 | 5466146 | + | *hprA* | *PA14_61210* | glycerate dehydrogenase | 15 | 13 | 16 | 13 | 12 | 10 | 1 |
| 5006 | - | 5466256 | 5467254 | + | - | *PA14_61220* | hypothetical protein | 25 | 21 | 29 | 24 | 19 | 21 | 1 |
| 5007 | - | 5467349 | 5468812 | + | *lysP* | *PA14_61250* | APC family lysine-specific permease | 23 | 22 | 27 | 19 | 22 | 18 | 1 |
| 5008 | - | 5469165 | 5469743 | + | - | *PA14_61260* | hypothetical protein | 3 | 4 | 4 | 4 | 5 | 4 | 1 |
| 5009 | - | 5469860 | 5470303 | + | - | *PA14_61270* | hypothetical protein | 45 | 39 | 39 | 40 | 28 | 41 | 1 |
| 5010 | - | 5471338 | 5470313 | - | - | *PA14_61280* | epimerase | 56 | 55 | 54 | 55 | 47 | 40 | 1 |
| 5011 | - | 5472303 | 5471482 | - | - | *PA14_61290* | lipoprotein | 40 | 51 | 35 | 35 | 51 | 27 | 1 |
| 5012 | - | 5472466 | 5474604 | + | - | *PA14_61300* | chemotaxis transducer | 171 | 190 | 181 | 178 | 204 | 205 | 1 |
| 5013 | - | 5475221 | 5474607 | - | - | *PA14_61320* | hypothetical protein | 115 | 125 | 121 | 133 | 129 | 128 | 1 |
| 5014 | - | 5475647 | 5476351 | + | - | *PA14_61330* | magnesium transporter, MgtC family | 31 | 32 | 33 | 35 | 27 | 35 | 1 |
| 5015 | - | 5476627 | 5476364 | - | - | *PA14_61340* | hypothetical protein | 88 | 75 | 86 | 109 | 84 | 91 | 1 |
| 5016 | - | 5476809 | 5476627 | - | - | *PA14_61350* | hypothetical protein | 81 | 84 | 97 | 123 | 89 | 94 | 1 |
| 5017 | - | 5476906 | 5478066 | + | - | *PA14_61360* | hypothetical protein | 31 | 30 | 38 | 42 | 30 | 32 | 1 |
| 5018 | - | 5478444 | 5478166 | - | - | *PA14_61370* | hypothetical protein | 101 | 65 | 127 | 88 | 76 | 82 | 1 |
| 5019 | - | 5478677 | 5478934 | + | - | *PA14_61380* | hypothetical protein | 149 | 253 | 198 | 234 | 232 | 235 | 0.831807123 |
| 5020 | - | 5479579 | 5478992 | - | - | *PA14_61390* | lipoprotein | 81 | 100 | 91 | 81 | 81 | 72 | 1 |
| 5021 | - | 5480421 | 5481944 | + | *mqoB* | *PA14_61400* | malate:quinone oxidoreductase | 134 | 111 | 103 | 83 | 91 | 77 | 1 |
| 5022 | - | 5482409 | 5481996 | - | - | *PA14_61410* | hypothetical protein | 156 | 178 | 208 | 211 | 179 | 240 | 1 |
| 5023 | - | 5482956 | 5482666 | - | - | *PA14_61430* | hypothetical protein | 17 | 18 | 15 | 12 | 12 | 9 | 1 |
| 5024 | - | 5483524 | 5483039 | - | - | *PA14_61440* | hypothetical protein | 19 | 15 | 23 | 15 | 18 | 21 | 1 |
| 5025 | - | 5484060 | 5483587 | - | - | *PA14_61450* | hypothetical protein | 43 | 45 | 52 | 62 | 38 | 38 | 1 |
| 5026 | - | 5484619 | 5484062 | - | - | *PA14_61460* | hypoxanthine-guanine phosphoribosyltransferase | 23 | 26 | 31 | 29 | 24 | 23 | 1 |
| 5027 | - | 5484787 | 5485425 | + | *upp* | *PA14_61470* | uracil phosphoribosyltransferase | 43 | 52 | 55 | 46 | 45 | 47 | 1 |
| 5028 | - | 5485428 | 5486711 | + | *uraA* | *PA14_61480* | uracil permease | 86 | 91 | 91 | 90 | 77 | 80 | 1 |
| 5029 | - | 5487031 | 5487579 | + | - | *PA14_61500* | hypothetical protein | 42 | 26 | 45 | 36 | 31 | 50 | 1 |
| 5030 | - | 5487610 | 5488143 | + | - | *PA14_61510* | hypothetical protein | 49 | 32 | 65 | 49 | 40 | 51 | 1 |
| 5031 | - | 5488143 | 5488685 | + | - | *PA14_61520* | hypothetical protein | 18 | 13 | 14 | 12 | 12 | 15 | 1 |
| 5032 | - | 5488704 | 5489492 | + | - | *PA14_61530* | pili assembly chaperone | 23 | 15 | 17 | 19 | 14 | 29 | 1 |
| 5033 | - | 5489509 | 5491881 | + | - | *PA14_61540* | hypothetical protein | 14 | 12 | 11 | 15 | 12 | 18 | 1 |
| 5034 | - | 5491878 | 5492825 | + | - | *PA14_61550* | hypothetical protein | 32 | 29 | 30 | 38 | 26 | 38 | 1 |
| 5035 | - | 5494200 | 5492827 | - | - | *PA14_61560* | MFS transporter | 6 | 6 | 5 | 6 | 6 | 8 | 1 |
| 5036 | - | 5495502 | 5494480 | - | *hemH* | *PA14_61580* | ferrochelatase | 85 | 71 | 62 | 65 | 78 | 90 | 1 |
| 5037 | - | 5496416 | 5495499 | - | - | *PA14_61590* | hypothetical protein | 157 | 124 | 162 | 138 | 148 | 161 | 1 |
| 5038 | - | 5496830 | 5497813 | + | - | *PA14_61600* | hypothetical protein | 157 | 152 | 189 | 143 | 163 | 175 | 1 |
| 5039 | - | 5497966 | 5498925 | + | - | *PA14_61610* | hypothetical protein | 41 | 35 | 36 | 28 | 53 | 55 | 1 |
| 5040 | - | 5498935 | 5499834 | + | - | *PA14_61620* | MerR family transcriptional regulator | 19 | 19 | 18 | 15 | 28 | 26 | 1 |

| | A | B | C | D | E | F | G | H | I | J | K | L | M | N |
|---|---|---|---|---|---|---|---|---|---|---|---|---|---|---|
| 5041 | - | 5499831 | 5501276 | + | *phr* | *PA14_61640* | deoxyribodipyrimidine photolyase | 30 | 30 | 30 | 33 | 42 | 37 | 1 |
| 5042 | - | 5501923 | 5501402 | - | *pagL* | *PA14_61650* | Lipid A 3-O-deacylase | 423 | 741 | 452 | 432 | 725 | 427 | 1 |
| 5043 | - | 5502854 | 5502057 | - | *murI* | *PA14_61660* | glutamate racemase | 117 | 99 | 115 | 102 | 104 | 88 | 1 |
| 5044 | - | 5503602 | 5502844 | - | *moeB* | *PA14_61670* | molybdopterin biosynthesis protein MoeB | 266 | 238 | 252 | 231 | 221 | 207 | 1 |
| 5045 | - | 5504426 | 5503596 | - | - | *PA14_61680* | methyl transferase | 119 | 112 | 115 | 110 | 105 | 85 | 1 |
| 5046 | - | 5505510 | 5504428 | - | *prfA* | *PA14_61700* | peptide chain release factor 1 | 105 | 79 | 98 | 88 | 85 | 76 | 1 |
| 5047 | - | 5506796 | 5505528 | - | *hemA* | *PA14_61710* | glutamyl-tRNA reductase | 46 | 30 | 43 | 32 | 23 | 27 | 1 |
| 5048 | - | 5506940 | 5508712 | + | - | *PA14_61720* | hypothetical protein | 83 | 83 | 79 | 65 | 78 | 72 | 1 |
| 5049 | - | 5508717 | 5509334 | + | *lolB* | *PA14_61740* | outer membrane lipoprotein LolB | 80 | 78 | 82 | 79 | 84 | 79 | 1 |
| 5050 | - | 5509336 | 5510184 | + | *ipk* | *PA14_61750* | 4-diphosphocytidyl-2-C-methyl-D-erythritol kinase | 87 | 86 | 81 | 73 | 69 | 61 | 1 |
| 5051 | - | 5510221 | 5510292 | + | - | *PA14_61760* | Gln tRNA | 123 | 97 | 119 | 75 | 52 | 47 | 1 |
| 5052 | - | 5510351 | 5511292 | + | *prs* | *PA14_61770* | ribose-phosphate pyrophosphokinase | 116 | 101 | 126 | 81 | 79 | 66 | 1 |
| 5053 | - | 5511409 | 5512023 | + | - | *PA14_61780* | 50S ribosomal protein L25 | 239 | 358 | 210 | 219 | 252 | 156 | 1 |
| 5054 | - | 5512065 | 5512649 | + | *pth* | *PA14_61790* | peptidyl-tRNA hydrolase | 41 | 46 | 45 | 34 | 34 | 25 | 1 |
| 5055 | - | 5512690 | 5513790 | + | - | *PA14_61820* | GTP-dependent nucleic acid-binding protein EngD | 74 | 72 | 86 | 57 | 72 | 51 | 1 |
| 5056 | - | 5513969 | 5514042 | + | - | *PA14_61830* | Met tRNA | 21 | 23 | 20 | 28 | 24 | 36 | 1 |
| 5057 | - | 5514501 | 5514196 | - | - | *PA14_61840* | virulence-associated protein | 124 | 108 | 86 | 83 | 99 | 101 | 1 |
| 5058 | - | 5515120 | 5517348 | + | - | *PA14_61850* | TonB-dependent receptor | 24 | 29 | 26 | 31 | 27 | 24 | 1 |
| 5059 | - | 5518065 | 5517418 | - | - | *PA14_61860* | carbonic anhydrase | 80 | 97 | 80 | 104 | 92 | 90 | 1 |
| 5060 | - | 5519365 | 5518127 | - | - | *PA14_61870* | hypothetical protein | 64 | 69 | 72 | 85 | 63 | 73 | 1 |
| 5061 | - | 5520016 | 5519564 | - | *rimI* | *PA14_61880* | peptide n-acetyltransferase RimI | 19 | 18 | 22 | 19 | 16 | 18 | 1 |
| 5062 | - | 5520714 | 5520013 | - | - | *PA14_61890* | hypothetical protein | 45 | 36 | 44 | 40 | 29 | 32 | 1 |
| 5063 | - | 5521510 | 5520974 | - | - | *PA14_61910* | hypothetical protein | 19 | 21 | 19 | 25 | 18 | 29 | 1 |
| 5064 | - | 5522560 | 5521541 | - | - | *PA14_61920* | hypothetical protein | 18 | 19 | 20 | 22 | 17 | 21 | 1 |
| 5065 | - | 5523608 | 5522562 | - | - | *PA14_61940* | hypothetical protein | 21 | 24 | 22 | 26 | 24 | 29 | 1 |
| 5066 | - | 5524406 | 5523804 | - | - | *PA14_61950* | hypothetical protein | 54 | 45 | 61 | 46 | 47 | 42 | 1 |
| 5067 | - | 5524691 | 5525989 | + | - | *PA14_61960* | hypothetical protein | 18 | 18 | 20 | 17 | 17 | 14 | 1 |
| 5068 | - | 5525979 | 5526674 | + | - | *PA14_61980* | hypothetical protein | 19 | 23 | 23 | 19 | 22 | 15 | 1 |
| 5069 | - | 5526671 | 5529517 | + | - | *PA14_61990* | hypothetical protein | 48 | 59 | 48 | 40 | 51 | 33 | 1 |
| 5070 | - | 5529629 | 5530636 | + | *hitA* | *PA14_62000* | ferric iron-binding periplasmic protein HitA | 34 | 78 | 39 | 69 | 91 | 58 | 0.221092106 |
| 5071 | - | 5530657 | 5532195 | + | *hitB* | *PA14_62010* | iron ABC transporter, permease | 19 | 28 | 24 | 28 | 36 | 27 | 1 |
| 5072 | - | 5534582 | 5532276 | - | - | *PA14_62020* | paraquat-inducible protein B-like protein | 86 | 78 | 70 | 91 | 74 | 79 | 1 |
| 5073 | - | 5535195 | 5534575 | - | - | *PA14_62030* | paraquat-inducible protein A-like protein | 52 | 40 | 50 | 50 | 49 | 49 | 1 |
| 5074 | - | 5535889 | 5535182 | - | - | *PA14_62040* | paraquat-inducible protein | 39 | 33 | 38 | 33 | 36 | 32 | 1 |
| 5075 | - | 5536159 | 5536028 | - | - | *PA14_62050* | 5S ribosomal RNA | 10 | 10 | 25 | 10 | 5 | 1 | 1 |
| 5076 | - | 5539185 | 5536295 | - | - | *PA14_62060* | 23S ribosomal RNA | 610 | 870 | 721 | 416 | 749 | 121 | 1 |
| 5077 | - | 5539487 | 5539415 | - | - | *PA14_62070* | Ala tRNA | 696 | 956 | 932 | 643 | 489 | 240 | 1 |
| 5078 | - | 5539592 | 5539519 | - | - | *PA14_62080* | Ile tRNA | 710 | 1071 | 928 | 568 | 435 | 216 | 1 |
| 5079 | - | 5541183 | 5539658 | - | - | *PA14_62090* | 16S ribosomal RNA | 237 | 360 | 900 | 310 | 553 | 68 | 1 |
| 5080 | - | 5542343 | 5541735 | - | - | *PA14_62100* | sulfite oxidase subunit YedZ | 42 | 32 | 35 | 30 | 27 | 30 | 1 |
| 5081 | - | 5543356 | 5542343 | - | - | *PA14_62110* | sulfite oxidase subunit YedY | 40 | 33 | 39 | 26 | 26 | 29 | 1 |
| 5082 | - | 5544232 | 5543417 | - | *pssA* | *PA14_62120* | phosphatidylserine synthase | 95 | 78 | 72 | 56 | 72 | 61 | 1 |
| 5083 | - | 5545403 | 5544387 | - | *ilvC* | *PA14_62130* | ketol-acid reductoisomerase | 193 | 275 | 148 | 113 | 202 | 116 | 1 |
| 5084 | - | 5545937 | 5545446 | - | *ilvH* | *PA14_62150* | acetolactate synthase 3 regulatory subunit | 98 | 135 | 80 | 83 | 106 | 76 | 1 |
| 5085 | - | 5547664 | 5545940 | - | *ilvI* | *PA14_62160* | acetolactate synthase 3 catalytic subunit | 171 | 193 | 150 | 168 | 179 | 135 | 1 |
| 5086 | - | 5548215 | 5548667 | + | - | *PA14_62170* | hypothetical protein | 26 | 28 | 28 | 21 | 22 | 25 | 1 |
| 5087 | - | 5549104 | 5548775 | - | - | *PA14_62180* | hypothetical protein | 245 | 226 | 263 | 241 | 239 | 206 | 1 |
| 5088 | - | 5549883 | 5549104 | - | - | *PA14_62190* | hypothetical protein | 145 | 153 | 146 | 139 | 147 | 127 | 1 |
| 5089 | - | 5552224 | 5549900 | - | *mrcB* | *PA14_62200* | penicillin-binding protein 1B | 69 | 60 | 77 | 66 | 61 | 54 | 1 |
| 5090 | - | 5552437 | 5553999 | + | - | *PA14_62230* | hypothetical protein | 64 | 68 | 59 | 63 | 60 | 60 | 1 |
| 5091 | - | 5554091 | 5554438 | + | - | *PA14_62240* | hypothetical protein | 218 | 289 | 289 | 252 | 250 | 318 | 1 |
| 5092 | - | 5554664 | 5554936 | + | - | *PA14_62250* | hypothetical protein | 63 | 68 | 89 | 90 | 82 | 84 | 1 |
| 5093 | - | 5555041 | 5555838 | + | - | *PA14_62260* | hypothetical protein | 33 | 39 | 43 | 40 | 33 | 37 | 1 |
| 5094 | - | 5557189 | 5556302 | - | - | *PA14_62270* | hypothetical protein | 46 | 44 | 39 | 43 | 33 | 37 | 1 |
| 5095 | - | 5557967 | 5557200 | - | *hmuV* | *PA14_62280* | hemin importer ATP-binding subunit | 70 | 71 | 71 | 97 | 71 | 62 | 1 |
| 5096 | - | 5558950 | 5557967 | - | - | *PA14_62290* | ABC transporter permease | 25 | 24 | 24 | 41 | 25 | 18 | 1 |
| 5097 | - | 5559894 | 5559001 | - | - | *PA14_62300* | hypothetical protein | 50 | 59 | 55 | 146 | 72 | 47 | 1 |
| 5098 | - | 5560968 | 5559904 | - | - | *PA14_62330* | hemin degrading factor | 14 | 47 | 15 | 124 | 75 | 17 | 0.000303735 |
| 5099 | - | 5561148 | 5563442 | + | - | *PA14_62350* | heme/hemoglobin uptake outer membrane receptor PhuR | 16 | 19 | 21 | 36 | 22 | 17 | 1 |
| 5100 | - | 5563545 | 5563892 | + | - | *PA14_62360* | Rieske family iron-sulfur cluster-binding protein | 51 | 48 | 47 | 33 | 27 | 38 | 1 |
| 5101 | - | 5563973 | 5564485 | + | - | *PA14_62370* | hypothetical protein | 37 | 40 | 51 | 42 | 41 | 45 | 1 |
| 5102 | - | 5564938 | 5564489 | - | - | *PA14_62380* | hypothetical protein | 254 | 255 | 353 | 442 | 305 | 316 | 1 |
| 5103 | - | 5565188 | 5565637 | + | - | *PA14_62390* | hypothetical protein | 521 | 621 | 499 | 366 | 442 | 369 | 1 |
| 5104 | - | 5566931 | 5565696 | - | - | *PA14_62400* | aminotransferase | 116 | 159 | 126 | 138 | 136 | 104 | 1 |
| 5105 | - | 5567101 | 5567955 | + | - | *PA14_62410* | hypothetical protein | 54 | 41 | 53 | 48 | 41 | 39 | 1 |
| 5106 | - | 5568023 | 5568916 | + | - | *PA14_62420* | hypothetical protein | 239 | 189 | 291 | 242 | 213 | 231 | 1 |
| 5107 | - | 5569414 | 5568938 | - | - | *PA14_62430* | hypothetical protein | 57 | 51 | 63 | 48 | 47 | 44 | 1 |
| 5108 | - | 5569828 | 5571123 | + | - | *PA14_62440* | transporter | 26 | 26 | 31 | 30 | 28 | 28 | 1 |
| 5109 | - | 5571120 | 5572211 | + | *trmA* | *PA14_62450* | tRNA (uracil-5-)-methyltransferase | 47 | 49 | 58 | 54 | 50 | 47 | 1 |
| 5110 | - | 5572952 | 5572245 | - | - | *PA14_62470* | sugar fermentation stimulation protein A | 86 | 87 | 88 | 98 | 89 | 86 | 1 |
| 5111 | - | 5574124 | 5572952 | - | - | *PA14_62480* | hypothetical protein | 81 | 80 | 94 | 88 | 73 | 86 | 1 |
| 5112 | - | 5574317 | 5574763 | + | *dksA* | *PA14_62490* | suppressor protein DksA | 194 | 264 | 197 | 168 | 153 | 134 | 1 |
| 5113 | - | 5574831 | 5575712 | + | - | *PA14_62510* | glutamyl-Q tRNA(Asp) synthetase | 54 | 55 | 47 | 46 | 53 | 40 | 1 |
| 5114 | - | 5575782 | 5575958 | + | - | *PA14_62520* | hypothetical protein | 28 | 20 | 29 | 24 | 22 | 27 | 1 |
| 5115 | - | 5575942 | 5578893 | + | *cbrA* | *PA14_62530* | two-component sensor CbrA | 39 | 37 | 40 | 40 | 40 | 35 | 1 |
| 5116 | - | 5578920 | 5580356 | + | *cbrB* | *PA14_62540* | two-component response regulator CbrB | 324 | 297 | 334 | 264 | 283 | 290 | 1 |
| 5117 | - | 5580536 | 5580865 | ? | - | predicted RNA | - | 29365 | 45934 | 42934 | 47182 | 39306 | 53290 | 1 |
| 5118 | - | 5581256 | 5582659 | + | *pcnB* | *PA14_62560* | poly(A) polymerase | 100 | 91 | 98 | 99 | 84 | 84 | 1 |
| 5119 | - | 5582656 | 5583144 | + | *folK* | *PA14_62570* | 2-amino-4-hydroxy-6-hydroxymethyldihydropteridine pyrophosphokinase | 64 | 72 | 65 | 71 | 61 | 50 | 1 |
| 5120 | - | 5583394 | 5584194 | + | *panB* | *PA14_62580* | 3-methyl-2-oxobutanoate hydroxymethyltransferase | 48 | 58 | 52 | 50 | 56 | 39 | 1 |
| 5121 | - | 5584191 | 5585042 | + | *panC* | *PA14_62590* | pantoate--beta-alanine ligase | 46 | 63 | 41 | 42 | 57 | 38 | 1 |
| 5122 | - | 5585136 | 5585516 | + | *panD* | *PA14_62600* | aspartate alpha-decarboxylase | 37 | 44 | 42 | 33 | 34 | 29 | 1 |
| 5123 | - | 5585606 | 5587270 | + | *pgi* | *PA14_62620* | glucose-6-phosphate isomerase | 107 | 96 | 84 | 78 | 87 | 89 | 1 |
| 5124 | - | 5587464 | 5589401 | + | *acsB* | *PA14_62630* | acetyl-CoA synthetase | 63 | 69 | 63 | 78 | 68 | 83 | 1 |

| | A | B | C | D | E | F | G | H | I | J | K | L | M | N |
|---|---|---|---|---|---|---|---|---|---|---|---|---|---|---|
| 5125 | - | 5589498 | 5590379 | + | - | *PA14_62640* | hypothetical protein | 54 | 44 | 43 | 50 | 40 | 41 | 1 |
| 5126 | - | 5590504 | 5593770 | + | - | *PA14_62650* | hypothetical protein | 117 | 117 | 122 | 123 | 118 | 149 | 1 |
| 5127 | - | 5593775 | 5594068 | + | - | *PA14_62660* | hypothetical protein | 228 | 252 | 223 | 325 | 265 | 318 | 1 |
| 5128 | - | 5594161 | 5594379 | + | - | *PA14_62670* | hypothetical protein | 144 | 132 | 162 | 153 | 140 | 142 | 1 |
| 5129 | - | 5594636 | 5594439 | - | - | *PA14_62680* | hypothetical protein | 1919 | 1838 | 1775 | 1549 | 1871 | 1897 | 1 |
| 5130 | - | 5595030 | 5594686 | - | - | *PA14_62690* | hypothetical protein | 1975 | 1750 | 1299 | 1149 | 1221 | 1528 | 1 |
| 5131 | - | 5597408 | 5595303 | - | *pnp* | *PA14_62710* | polynucleotide phosphorylase | 223 | 296 | 202 | 164 | 265 | 161 | 1 |
| 5132 | - | 5597851 | 5597582 | - | *rpsO* | *PA14_62720* | 30S ribosomal protein S15 | 875 | 1540 | 939 | 533 | 621 | 337 | 1 |
| 5133 | - | 5598861 | 5597947 | - | *truB* | *PA14_62730* | tRNA pseudouridine synthase B | 199 | 193 | 221 | 191 | 176 | 147 | 1 |
| 5134 | - | 5599253 | 5598864 | - | *rbfA* | *PA14_62740* | ribosome-binding factor A | 113 | 112 | 118 | 94 | 115 | 90 | 1 |
| 5135 | - | 5601878 | 5599356 | - | *infB* | *PA14_62760* | translation initiation factor IF-2 | 426 | 489 | 434 | 411 | 449 | 335 | 1 |
| 5136 | - | 5603387 | 5601906 | - | *nusA* | *PA14_62770* | transcription elongation factor NusA | 334 | 379 | 401 | 385 | 359 | 288 | 1 |
| 5137 | - | 5603890 | 5603432 | - | - | *PA14_62780* | hypothetical protein | 83 | 39 | 96 | 71 | 46 | 60 | 0.197700537 |
| 5138 | - | 5604093 | 5604020 | - | - | *PA14_62790* | Met tRNA | 263 | 95 | 275 | 267 | 163 | 176 | 0.000461274 |
| 5139 | - | 5604269 | 5604187 | - | - | *PA14_62800* | Leu tRNA | 397 | 315 | 409 | 334 | 296 | 230 | 1 |
| 5140 | - | 5604672 | 5604283 | - | *secG* | *PA14_62810* | preprotein translocase subunit SecG | 536 | 530 | 611 | 446 | 471 | 415 | 1 |
| 5141 | - | 5605430 | 5604675 | - | *tpiA* | *PA14_62830* | triosephosphate isomerase | 577 | 457 | 673 | 548 | 510 | 440 | 1 |
| 5142 | - | 5606833 | 5605496 | - | *glmM* | *PA14_62840* | phosphoglucosamine mutase | 348 | 379 | 310 | 281 | 365 | 323 | 1 |
| 5143 | - | 5607701 | 5606850 | - | *folP* | *PA14_62850* | dihydropteroate synthase | 215 | 183 | 212 | 172 | 201 | 206 | 1 |
| 5144 | - | 5609630 | 5607711 | - | *ftsH* | *PA14_62860* | cell division protein FtsH | 803 | 596 | 671 | 513 | 594 | 688 | 1 |
| 5145 | - | 5610452 | 5609829 | - | *rrmJ* | *PA14_62870* | cell division protein FtsJ | 176 | 167 | 245 | 144 | 192 | 155 | 1 |
| 5146 | - | 5610547 | 5610861 | + | - | *PA14_62880* | hypothetical protein | 138 | 114 | 163 | 125 | 147 | 119 | 1 |
| 5147 | - | 5611309 | 5610902 | - | - | *PA14_62890* | hypothetical protein | 348 | 280 | 326 | 276 | 318 | 266 | 1 |
| 5148 | - | 5611796 | 5611320 | - | *greA* | *PA14_62900* | transcription elongation factor GreA | 293 | 340 | 242 | 225 | 276 | 230 | 1 |
| 5149 | - | 5615014 | 5611793 | - | *carB* | *PA14_62910* | carbamoyl phosphate synthase large subunit | 218 | 261 | 202 | 188 | 256 | 184 | 1 |
| 5150 | - | 5615684 | 5615034 | - | - | *PA14_62920* | leucine export protein LeuE | 125 | 128 | 116 | 117 | 152 | 108 | 1 |
| 5151 | - | 5616832 | 5615696 | - | *carA* | *PA14_62930* | carbamoyl phosphate synthase small subunit | 163 | 147 | 165 | 135 | 146 | 131 | 1 |
| 5152 | - | 5617821 | 5617015 | - | *dapB* | *PA14_62940* | dihydrodipicolinate reductase | 156 | 187 | 160 | 80 | 198 | 133 | 1 |
| 5153 | - | 5619011 | 5617878 | - | *dnaJ* | *PA14_62960* | chaperone protein DnaJ | 232 | 283 | 256 | 124 | 288 | 185 | 1 |
| 5154 | - | 5621040 | 5619127 | - | *dnaK* | *PA14_62970* | molecular chaperone DnaK | 751 | 1031 | 905 | 383 | 891 | 558 | 1 |
| 5155 | - | 5621690 | 5621130 | - | *grpE* | *PA14_62990* | heat shock protein GrpE | 404 | 533 | 551 | 222 | 530 | 307 | 1 |
| 5156 | - | 5621858 | 5623534 | + | *recN* | *PA14_63010* | DNA repair protein RecN | 49 | 54 | 48 | 55 | 48 | 55 | 1 |
| 5157 | - | 5624008 | 5623604 | - | *fur* | *PA14_63020* | ferric uptake regulation protein | 513 | 497 | 532 | 459 | 450 | 477 | 1 |
| 5158 | - | 5624106 | 5624636 | + | *omlA* | *PA14_63030* | outer membrane lipoprotein OmlA precursor | 52 | 36 | 67 | 39 | 43 | 49 | 1 |
| 5159 | - | 5625009 | 5624704 | - | - | *PA14_63040* | hypothetical protein | 113 | 91 | 121 | 117 | 103 | 102 | 1 |
| 5160 | - | 5625436 | 5625002 | - | - | *PA14_63050* | hypothetical protein | 235 | 203 | 305 | 294 | 226 | 263 | 1 |
| 5161 | - | 5625711 | 5626190 | + | *smpB* | *PA14_63060* | SsrA-binding protein | 173 | 165 | 173 | 160 | 125 | 121 | 1 |
| 5162 | - | 5626986 | 5626213 | - | - | *PA14_63070* | GntR family transcriptional regulator | 96 | 94 | 120 | 108 | 84 | 99 | 1 |
| 5163 | - | 5627315 | 5628997 | + | *lldP* | *PA14_63080* | L-lactate permease | 16 | 17 | 24 | 20 | 13 | 41 | 1 |
| 5164 | - | 5629153 | 5630298 | + | *lldD* | *PA14_63090* | L-lactate dehydrogenase | 19 | 16 | 18 | 12 | 13 | 29 | 1 |
| 5165 | - | 5630365 | 5633181 | + | - | *PA14_63100* | ferredoxin | 25 | 21 | 26 | 20 | 17 | 48 | 1 |
| 5166 | - | 5633510 | 5633992 | + | - | *PA14_63110* | S-adenosylmethionine decarboxylase | 19 | 62 | 50 | 58 | 49 | 47 | 0.000275419 |
| 5167 | - | 5634070 | 5635119 | + | - | *PA14_63120* | hypothetical protein | 22 | 78 | 67 | 77 | 71 | 72 | 0.000282896 |
| 5168 | - | 5635122 | 5635982 | + | - | *PA14_63130* | hypothetical protein | 23 | 53 | 60 | 64 | 50 | 55 | 0.110188923 |
| 5169 | - | 5635995 | 5636660 | + | *pmrA* | *PA14_63150* | two-component response regulator | 28 | 43 | 54 | 66 | 51 | 65 | 1 |
| 5170 | - | 5636684 | 5638117 | + | *pmrB* | *PA14_63160* | PmrB: two-component regulator system signal sensor kinase PmrB | 15 | 27 | 28 | 40 | 29 | 37 | 0.540975013 |
| 5171 | - | 5638181 | 5638579 | + | - | *PA14_63170* | transcriptional regulator | 246 | 336 | 289 | 379 | 350 | 328 | 1 |
| 5172 | - | 5640004 | 5639111 | - | - | *PA14_63190* | hypothetical protein | 60 | 73 | 62 | 87 | 71 | 70 | 1 |
| 5173 | - | 5640987 | 5640094 | - | - | *PA14_63200* | hypothetical protein | 64 | 60 | 73 | 77 | 64 | 75 | 1 |
| 5174 | - | 5642292 | 5641111 | - | - | *PA14_63210* | two-component response regulator | 150 | 147 | 202 | 216 | 157 | 222 | 1 |
| 5175 | - | 5642645 | 5642400 | - | - | *PA14_63220* | hypothetical protein | 74 | 172 | 200 | 370 | 206 | 306 | 0.0849596 |
| 5176 | - | 5643569 | 5642679 | - | - | *PA14_63230* | hypothetical protein | 67 | 41 | 61 | 39 | 54 | 47 | 1 |
| 5177 | - | 5643724 | 5644191 | + | - | *PA14_63240* | transcriptional regulator | 13 | 17 | 12 | 14 | 21 | 14 | 1 |
| 5178 | - | 5645637 | 5644360 | - | - | *PA14_63250* | acetyl-CoA acetyltransferase | 39 | 41 | 41 | 47 | 47 | 51 | 1 |
| 5179 | - | 5645892 | 5647247 | + | *fabG* | *PA14_63270* | 3-ketoacyl-ACP reductase | 76 | 75 | 75 | 82 | 83 | 86 | 1 |
| 5180 | - | 5648325 | 5647324 | - | - | *PA14_63280* | transcriptional regulator | 185 | 227 | 161 | 242 | 196 | 219 | 1 |
| 5181 | - | 5648586 | 5649443 | + | *phaJ3* | *PA14_63290* | hypothetical protein | 12 | 9 | 13 | 11 | 10 | 13 | 1 |
| 5182 | - | 5649525 | 5649830 | + | - | *PA14_63300* | hypothetical protein | 13 | 9 | 12 | 9 | 12 | 8 | 1 |
| 5183 | - | 5649827 | 5650576 | + | - | *PA14_63310* | hypothetical protein | 21 | 19 | 18 | 19 | 18 | 16 | 1 |
| 5184 | - | 5650642 | 5651262 | + | - | *PA14_63320* | hypothetical protein | 40 | 40 | 34 | 35 | 38 | 35 | 1 |
| 5185 | - | 5652182 | 5651247 | - | - | *PA14_63330* | glycerolphosphodiesterase | 24 | 25 | 26 | 29 | 22 | 23 | 1 |
| 5186 | - | 5652314 | 5652877 | + | - | *PA14_63340* | lipoprotein | 88 | 94 | 94 | 80 | 86 | 82 | 1 |
| 5187 | - | 5652879 | 5653361 | + | - | *PA14_63350* | GNAT family acetyltransferase | 64 | 59 | 52 | 47 | 62 | 51 | 1 |
| 5188 | - | 5653351 | 5653728 | + | - | *PA14_63360* | hypothetical protein | 30 | 28 | 28 | 27 | 28 | 25 | 1 |
| 5189 | - | 5653725 | 5654192 | + | - | *PA14_63370* | hypothetical protein | 71 | 66 | 66 | 69 | 65 | 65 | 1 |
| 5190 | - | 5654309 | 5655037 | + | - | *PA14_63380* | hypothetical protein | 46 | 41 | 51 | 54 | 50 | 46 | 1 |
| 5191 | - | 5655638 | 5655060 | - | - | *PA14_63390* | adenylate kinase | 33 | 38 | 30 | 28 | 35 | 24 | 1 |
| 5192 | - | 5656435 | 5655635 | - | - | *PA14_63410* | hypothetical protein | 33 | 36 | 32 | 27 | 36 | 25 | 1 |
| 5193 | - | 5656610 | 5656494 | - | - | *PA14_63420* | hypothetical protein | 30 | 34 | 32 | 21 | 25 | 25 | 1 |
| 5194 | - | 5657099 | 5656731 | - | - | *PA14_63430* | hypothetical protein | 63 | 54 | 62 | 61 | 57 | 45 | 1 |
| 5195 | - | 5657320 | 5657700 | + | - | *PA14_63440* | bacteriophage integrase | 29 | 29 | 34 | 30 | 30 | 24 | 1 |
| 5196 | - | 5657975 | 5658310 | + | - | *PA14_63450* | hypothetical protein | 49 | 45 | 61 | 57 | 56 | 53 | 1 |
| 5197 | - | 5658498 | 5658407 | - | - | *PA14_63460* | Sec tRNA | 36 | 25 | 36 | 18 | 21 | 20 | 1 |
| 5198 | - | 5659247 | 5658630 | - | - | *PA14_63470* | methyltransferase | 84 | 82 | 80 | 76 | 93 | 85 | 1 |
| 5199 | - | 5660739 | 5659342 | - | - | *PA14_63480* | amino acid permease | 11 | 13 | 12 | 12 | 9 | 10 | 1 |
| 5200 | - | 5662194 | 5660788 | - | - | *PA14_63500* | hypothetical protein | 11 | 10 | 10 | 10 | 10 | 9 | 1 |
| 5201 | - | 5662476 | 5663159 | + | - | *PA14_63520* | transcriptional regulator | 27 | 36 | 37 | 34 | 31 | 34 | 1 |
| 5202 | - | 5665088 | 5663163 | - | *selB* | *PA14_63530* | selenocysteine-specific elongation factor | 89 | 88 | 88 | 74 | 76 | 66 | 1 |
| 5203 | - | 5666491 | 5665085 | - | *selA* | *PA14_63540* | selenocysteine synthase | 98 | 96 | 93 | 80 | 89 | 79 | 1 |
| 5204 | - | 5667499 | 5666570 | - | *fdhE* | *PA14_63550* | formate dehydrogenase accessory protein FdhE | 59 | 67 | 72 | 56 | 67 | 60 | 1 |
| 5205 | - | 5668258 | 5667632 | - | *fdnI* | *PA14_63570* | nitrate-inducible formate dehydrogenase subunit gamma | 194 | 234 | 269 | 213 | 241 | 222 | 1 |
| 5206 | - | 5669259 | 5668330 | - | *fdnH* | *PA14_63580* | nitrate-inducible formate dehydrogenase subunit beta | 204 | 268 | 276 | 239 | 275 | 285 | 1 |
| 5207 | - | 5672348 | 5669268 | - | *fdnG* | *PA14_63605* | formate dehydrogenase-O, major subunit | 266 | 351 | 337 | 292 | 308 | 320 | 1 |
| 5208 | - | 5673585 | 5672656 | - | *lipC* | *PA14_63620* | lipase LipC | 7 | 10 | 9 | 10 | 8 | 11 | 1 |

| | A | B | C | D | E | F | G | H | I | J | K | L | M | N |
|---|---|---|---|---|---|---|---|---|---|---|---|---|---|---|
| 5209 | - | 5673945 | 5675990 | + | *fadH2* | *PA14_63640* | 2,4-dienoyl-CoA reductase | 7 | 7 | 6 | 6 | 6 | 6 | 1 |
| 5210 | - | 5676199 | 5676648 | + | - | *PA14_63650* | hypothetical protein | 14 | 13 | 16 | 14 | 13 | 12 | 1 |
| 5211 | - | 5676805 | 5678040 | + | - | *PA14_63660* | hypothetical protein | 15 | 14 | 15 | 14 | 14 | 12 | 1 |
| 5212 | - | 5678016 | 5678510 | + | - | *PA14_63680* | hypothetical protein | 9 | 7 | 6 | 8 | 6 | 6 | 1 |
| 5213 | - | 5679949 | 5678513 | - | - | *PA14_63700* | hypothetical protein | 7 | 6 | 7 | 7 | 6 | 7 | 1 |
| 5214 | - | 5680920 | 5679946 | - | - | *PA14_63710* | glycosyl transferase family protein | 2 | 3 | 2 | 2 | 1 | 1 | 1 |
| 5215 | - | 5681279 | 5680917 | - | - | *PA14_63720* | hypothetical protein | 3 | 4 | 5 | 4 | 5 | 4 | 1 |
| 5216 | - | 5681516 | 5682877 | + | - | *PA14_63730* | transporter | 10 | 13 | 13 | 13 | 13 | 9 | 1 |
| 5217 | - | 5683221 | 5682853 | - | - | *PA14_63740* | hypothetical protein | 11 | 18 | 17 | 16 | 16 | 14 | 1 |
| 5218 | - | 5684876 | 5683209 | - | - | *PA14_63750* | hypothetical protein | 12 | 13 | 14 | 12 | 12 | 12 | 1 |
| 5219 | - | 5685128 | 5684919 | - | - | *PA14_63770* | hypothetical protein | 5 | 4 | 5 | 5 | 3 | 4 | 1 |
| 5220 | - | 5685955 | 5685179 | - | - | *PA14_63780* | hypothetical protein | 5 | 6 | 4 | 6 | 5 | 4 | 1 |
| 5221 | - | 5688764 | 5686053 | - | *mgtA* | *PA14_63800* | Mg(2+) transport ATPase, P-type 2 | 9 | 9 | 9 | 7 | 8 | 7 | 1 |
| 5222 | - | 5689209 | 5688991 | - | - | *PA14_63820* | hypothetical protein | 7 | 7 | 8 | 10 | 7 | 8 | 1 |
| 5223 | - | 5690378 | 5689539 | - | - | *PA14_63830* | N-hydroxyarylamine O-acetyltransferase | 41 | 30 | 43 | 30 | 28 | 31 | 1 |
| 5224 | - | 5690503 | 5690958 | + | - | *PA14_63840* | hypothetical protein | 23 | 24 | 26 | 27 | 21 | 28 | 1 |
| 5225 | - | 5692366 | 5690963 | - | *lpd3* | *PA14_63850* | dihydrolipoamide dehydrogenase | 52 | 51 | 62 | 68 | 54 | 58 | 1 |
| 5226 | - | 5692539 | 5693078 | + | - | *PA14_63860* | hypothetical protein | 16 | 17 | 13 | 15 | 11 | 14 | 1 |
| 5227 | - | 5693146 | 5693706 | + | - | *PA14_63880* | transcriptional regulator | 38 | 37 | 31 | 27 | 31 | 31 | 1 |
| 5228 | - | 5694508 | 5693711 | - | - | *PA14_63890* | short-chain dehydrogenase | 23 | 24 | 23 | 23 | 20 | 18 | 1 |
| 5229 | - | 5694739 | 5695356 | + | - | *PA14_63900* | hemolyin III | 68 | 72 | 63 | 61 | 79 | 63 | 1 |
| 5230 | - | 5696256 | 5695402 | - | - | *PA14_63910* | hypothetical protein | 11 | 11 | 9 | 12 | 13 | 13 | 1 |
| 5231 | - | 5697539 | 5696238 | - | - | *PA14_63920* | hypothetical protein | 16 | 14 | 15 | 16 | 12 | 14 | 1 |
| 5232 | - | 5698327 | 5697536 | - | - | *PA14_63940* | hypothetical protein | 13 | 12 | 12 | 14 | 12 | 14 | 1 |
| 5233 | - | 5700483 | 5698357 | - | - | *PA14_63960* | uter membrane protein | 13 | 12 | 11 | 16 | 16 | 26 | 1 |
| 5234 | - | 5700625 | 5701779 | + | - | *PA14_63970* | hypothetical protein | 8 | 9 | 8 | 9 | 8 | 8 | 1 |
| 5235 | - | 5703942 | 5702032 | - | *speA* | *PA14_63990* | arginine decarboxylase | 70 | 63 | 70 | 78 | 69 | 72 | 1 |
| 5236 | - | 5704564 | 5704193 | - | - | *PA14_64000* | translation initiation factor Sui1 | 31 | 24 | 32 | 32 | 23 | 26 | 1 |
| 5237 | - | 5705304 | 5704768 | - | - | *PA14_64010* | hypothetical protein | 82 | 79 | 78 | 63 | 73 | 81 | 1 |
| 5238 | - | 5706378 | 5705308 | - | - | *PA14_64030* | hypothetical protein | 365 | 276 | 387 | 290 | 335 | 339 | 1 |
| 5239 | - | 5706731 | 5708359 | + | - | *PA14_64050* | two-component response regulator | 371 | 349 | 383 | 333 | 386 | 399 | 1 |
| 5240 | - | 5708437 | 5710410 | + | - | *PA14_64060* | chemotaxis transducer | 11 | 10 | 13 | 9 | 9 | 9 | 1 |
| 5241 | - | 5710570 | 5712345 | + | *dipZ* | *PA14_64080* | thiol:disulfide interchange protein | 35 | 30 | 42 | 43 | 34 | 28 | 1 |
| 5242 | - | 5712489 | 5712932 | + | *aroQ1* | *PA14_64090* | 3-dehydroquinate dehydratase | 73 | 58 | 90 | 69 | 74 | 69 | 1 |
| 5243 | - | 5712956 | 5713426 | + | *accB* | *PA14_64100* | acetyl-CoA carboxylase biotin carboxyl carrier protein subunit | 320 | 489 | 311 | 345 | 393 | 259 | 1 |
| 5244 | - | 5713444 | 5714793 | + | *accC* | *PA14_64110* | acetyl-CoA carboxylase biotin carboxylase subunit | 268 | 400 | 237 | 270 | 344 | 223 | 1 |
| 5245 | - | 5714904 | 5715770 | + | - | *PA14_64120* | hypothetical protein | 7 | 7 | 7 | 9 | 6 | 6 | 1 |
| 5246 | - | 5715852 | 5716736 | + | *prmA* | *PA14_64140* | 50S ribosomal protein L11 methyltransferase | 42 | 36 | 41 | 37 | 38 | 35 | 1 |
| 5247 | - | 5716832 | 5718109 | + | - | *PA14_64170* | hypothetical protein | 31 | 27 | 32 | 25 | 21 | 22 | 1 |
| 5248 | - | 5718313 | 5719311 | + | - | *PA14_64180* | hypothetical protein | 91 | 95 | 129 | 92 | 86 | 63 | 1 |
| 5249 | - | 5719308 | 5719631 | + | *fis* | *PA14_64190* | DNA-binding protein Fis | 58 | 80 | 68 | 59 | 73 | 45 | 1 |
| 5250 | - | 5719711 | 5721318 | + | *purH* | *PA14_64200* | bifunctional phosphoribosylaminoimidazolecarboxamide formyltransferase/IMP cyclohydrolase | 62 | 69 | 69 | 50 | 62 | 38 | 1 |
| 5251 | - | 5721422 | 5722711 | + | *purD* | *PA14_64220* | phosphoribosylamine--glycine ligase | 70 | 64 | 70 | 50 | 60 | 43 | 1 |
| 5252 | - | 5722815 | 5725643 | + | *retS* | *PA14_64230* | RetS (regulator of exopolysaccharide and Type III secretion) | 68 | 60 | 65 | 72 | 63 | 63 | 1 |
| 5253 | - | 5726098 | 5726691 | + | - | *PA14_64240* | MarC family protein | 9 | 11 | 13 | 9 | 11 | 9 | 1 |
| 5254 | - | 5726982 | 5727074 | + | - | *PA14_64260* | hypothetical protein | 2 | 2 | 2 | 1 | 2 | 2 | 1 |
| 5255 | - | 5727116 | 5728381 | + | - | *PA14_64270* | hypothetical protein | 39 | 31 | 53 | 44 | 29 | 27 | 1 |
| 5256 | - | 5728528 | 5730117 | + | - | *PA14_64280* | branched-chain amino acid ABC transporter permease | 2 | 1 | 3 | 2 | 3 | 2 | 0.934705131 |
| 5257 | - | 5730114 | 5731193 | + | - | *PA14_64290* | ABC transporter permease | 4 | 4 | 6 | 6 | 5 | 5 | 1 |
| 5258 | - | 5731190 | 5732047 | + | - | *PA14_64300* | ABC transporter ATP-binding protein | 6 | 6 | 7 | 6 | 6 | 4 | 1 |
| 5259 | - | 5732184 | 5732882 | + | - | *PA14_64310* | ABC transporter ATP-binding protein | 8 | 7 | 9 | 7 | 7 | 6 | 1 |
| 5260 | - | 5732966 | 5733418 | + | - | *PA14_64320* | hypothetical protein | 72 | 66 | 72 | 76 | 55 | 59 | 1 |
| 5261 | - | 5733427 | 5734269 | + | *ureD* | *PA14_64335* | urease accessory protein | 29 | 29 | 26 | 28 | 29 | 26 | 1 |
| 5262 | - | 5734271 | 5734573 | + | *ureA* | *PA14_64350* | urease subunit gamma | 74 | 110 | 69 | 81 | 114 | 68 | 1 |
| 5263 | - | 5734582 | 5735100 | + | - | *PA14_64360* | hypothetical protein | 94 | 101 | 76 | 73 | 104 | 67 | 1 |
| 5264 | - | 5735117 | 5735422 | + | *ureB* | *PA14_64370* | urease subunit beta | 93 | 101 | 75 | 92 | 123 | 90 | 1 |
| 5265 | - | 5735485 | 5737185 | + | *ureC* | *PA14_64390* | urease subunit alpha | 114 | 116 | 89 | 95 | 119 | 91 | 1 |
| 5266 | - | 5737454 | 5738671 | + | - | *PA14_64400* | hypothetical protein | 39 | 39 | 42 | 37 | 43 | 40 | 1 |
| 5267 | - | 5738947 | 5738681 | - | - | *PA14_64410* | hypothetical protein | 161 | 141 | 244 | 139 | 219 | 196 | 1 |
| 5268 | - | 5739073 | 5739687 | + | - | *PA14_64420* | hypothetical protein | 24 | 22 | 23 | 22 | 20 | 20 | 1 |
| 5269 | - | 5739839 | 5740534 | + | - | *PA14_64430* | hypothetical protein | 82 | 76 | 70 | 63 | 82 | 70 | 1 |
| 5270 | - | 5741404 | 5740541 | - | - | *PA14_64440* | hypothetical protein | 118 | 118 | 113 | 104 | 108 | 94 | 1 |
| 5271 | - | 5741612 | 5742877 | + | - | *PA14_64450* | heat-shock protein | 60 | 67 | 63 | 69 | 63 | 68 | 1 |
| 5272 | - | 5743449 | 5743033 | - | - | *PA14_64460* | hypothetical protein | 246 | 309 | 264 | 317 | 257 | 321 | 1 |
| 5273 | - | 5743847 | 5744143 | + | - | *PA14_64470* | hypothetical protein | 45 | 56 | 53 | 56 | 52 | 53 | 1 |
| 5274 | - | 5744232 | 5744576 | + | *osmE* | *PA14_64480* | DNA-binding transcriptional activator OsmE | 504 | 708 | 533 | 556 | 572 | 690 | 1 |
| 5275 | - | 5745027 | 5744620 | - | - | *PA14_64490* | hypothetical protein | 151 | 226 | 212 | 226 | 234 | 228 | 1 |
| 5276 | - | 5745174 | 5745986 | + | - | *PA14_64500* | transcriptional regulator | 53 | 82 | 54 | 72 | 79 | 66 | 0.967785381 |
| 5277 | - | 5748057 | 5745988 | - | - | *PA14_64510* | hypothetical protein | 77 | 77 | 83 | 87 | 84 | 93 | 1 |
| 5278 | - | 5748353 | 5748886 | + | - | *PA14_64520* | bacterioferritin | 354 | 412 | 347 | 247 | 429 | 384 | 1 |
| 5279 | - | 5749160 | 5749501 | + | - | *PA14_64530* | hypothetical protein | 15 | 13 | 19 | 9 | 13 | 14 | 1 |
| 5280 | - | 5750382 | 5749615 | - | - | *PA14_64540* | hypothetical protein | 18 | 16 | 15 | 19 | 16 | 19 | 1 |
| 5281 | - | 5750993 | 5750379 | - | - | *PA14_64550* | hypothetical protein | 2 | 3 | 3 | 2 | 2 | 1 | 1 |
| 5282 | - | 5751670 | 5751047 | - | - | *PA14_64560* | hypothetical protein | 1 | 2 | 3 | 1 | 3 | 2 | 0.449233963 |
| 5283 | - | 5751808 | 5752497 | + | *irlR* | *PA14_64570* | two-component response regulator | 16 | 11 | 14 | 11 | 14 | 14 | 1 |
| 5284 | - | 5752476 | 5753867 | + | - | *PA14_64580* | two-component sensor | 14 | 14 | 13 | 14 | 12 | 11 | 1 |
| 5285 | - | 5755174 | 5753858 | - | - | *PA14_64590* | MFS transporter | 35 | 29 | 36 | 31 | 31 | 21 | 1 |
| 5286 | - | 5756506 | 5755394 | - | - | *PA14_64610* | hypothetical protein | 62 | 66 | 75 | 54 | 72 | 57 | 1 |
| 5287 | - | 5757603 | 5756503 | - | - | *PA14_64620* | oxidoreductase | 163 | 151 | 209 | 140 | 207 | 158 | 1 |
| 5288 | - | 5757762 | 5758391 | + | - | *PA14_64640* | TetR family transcriptional regulator | 27 | 19 | 26 | 22 | 16 | 18 | 1 |
| 5289 | - | 5758621 | 5759124 | + | *ureE* | *PA14_64650* | urease accessory protein UreE | 14 | 12 | 14 | 10 | 12 | 8 | 1 |
| 5290 | - | 5759121 | 5759792 | + | *ureF* | *PA14_64660* | urease accessory protein UreF | 15 | 9 | 11 | 12 | 9 | 12 | 0.759690501 |
| 5291 | - | 5759803 | 5760417 | + | *ureG* | *PA14_64670* | urease accessory protein UreG | 20 | 18 | 19 | 20 | 19 | 20 | 1 |
| 5292 | - | 5760447 | 5761019 | + | *ureJ* | *PA14_64680* | hypothetical protein | 19 | 20 | 18 | 33 | 22 | 20 | 1 |

| | A | B | C | D | E | F | G | H | I | J | K | L | M | N |
|---|---|---|---|---|---|---|---|---|---|---|---|---|---|---|
| 5293 | - | 5762057 | 5761035 | - | - | *PA14_64690* | transmembrane sensor | 9 | 26 | 10 | 73 | 22 | 10 | 0.003675948 |
| 5294 | - | 5762586 | 5762050 | - | - | *PA14_64700* | RNA polymerase sigma factor | 3 | 25 | 3 | 59 | 21 | 5 | 1.61472E-31 |
| 5295 | - | 5762752 | 5765721 | + | - | *PA14_64710* | extracellular heme-binding protein | 13 | 13 | 16 | 21 | 15 | 14 | 1 |
| 5296 | - | 5767119 | 5765866 | - | - | *PA14_64720* | porin | 12 | 14 | 15 | 15 | 11 | 12 | 1 |
| 5297 | - | 5768696 | 5767227 | - | - | *PA14_64740* | aldehyde dehydrogenase | 34 | 33 | 30 | 24 | 26 | 26 | 1 |
| 5298 | - | 5770059 | 5768719 | - | - | *PA14_64750* | MFS transporter | 10 | 7 | 7 | 8 | 8 | 6 | 1 |
| 5299 | - | 5771732 | 5770146 | - | *mdlC* | *PA14_64770* | benzoylformate decarboxylase | 7 | 8 | 7 | 8 | 6 | 6 | 1 |
| 5300 | - | 5771833 | 5772729 | + | - | *PA14_64780* | transcriptional regulator | 11 | 12 | 11 | 9 | 11 | 9 | 1 |
| 5301 | - | 5774078 | 5772744 | - | - | *PA14_64790* | MFS transporter | 4 | 4 | 4 | 4 | 4 | 4 | 1 |
| 5302 | - | 5774457 | 5775512 | + | *vanA* | *PA14_64800* | vanillate O-demethylase oxygenase | 6 | 8 | 8 | 7 | 7 | 7 | 1 |
| 5303 | - | 5775527 | 5776480 | + | *vanB* | *PA14_64810* | vanillate O-demethylase | 19 | 21 | 23 | 27 | 20 | 24 | 1 |
| 5304 | - | 5777190 | 5776477 | - | - | *PA14_64820* | transcriptional regulator | 59 | 67 | 63 | 51 | 51 | 50 | 1 |
| 5305 | - | 5778015 | 5777254 | - | - | *PA14_64840* | short-chain dehydrogenase | 47 | 68 | 44 | 38 | 53 | 42 | 1 |
| 5306 | - | 5779266 | 5778373 | - | - | *PA14_64850* | ornithine cyclodeaminase | 13 | 12 | 14 | 17 | 12 | 20 | 1 |
| 5307 | - | 5780053 | 5779337 | - | - | *PA14_64860* | ABC transporter ATP-binding protein | 31 | 27 | 25 | 29 | 22 | 40 | 1 |
| 5308 | - | 5780922 | 5780050 | - | - | *PA14_64870* | ABC transporter ATP-binding protein | 20 | 14 | 17 | 21 | 16 | 25 | 1 |
| 5309 | - | 5782196 | 5780919 | - | - | *PA14_64880* | branched chain amino acid ABC transporter permease | 6 | 6 | 6 | 6 | 5 | 8 | 1 |
| 5310 | - | 5783121 | 5782207 | - | - | *PA14_64890* | branched chain amino acid ABC transporter permease | 12 | 13 | 13 | 14 | 10 | 13 | 1 |
| 5311 | - | 5784480 | 5783356 | - | - | *PA14_64900* | ABC transporter substrate-binding protein | 92 | 83 | 91 | 94 | 76 | 106 | 1 |
| 5312 | - | 5785100 | 5786038 | + | - | *PA14_64910* | LysR family transcriptional regulator | 23 | 23 | 22 | 21 | 22 | 26 | 1 |
| 5313 | - | 5786095 | 5787720 | + | - | *PA14_64920* | methyl-accepting chemotaxis protein | 159 | 168 | 204 | 227 | 153 | 214 | 1 |
| 5314 | - | 5788462 | 5787767 | - | - | *PA14_64930* | hypothetical protein | 273 | 190 | 198 | 316 | 264 | 332 | 1 |
| 5315 | - | 5789079 | 5788477 | - | - | *PA14_64940* | hypothetical protein | 345 | 59 | 147 | 245 | 255 | 305 | 4.94249E-11 |
| 5316 | - | 5789189 | 5789848 | + | *pncA* | *PA14_64950* | hypothetical protein | 16 | 50 | 32 | 19 | 46 | 12 | 0.000820036 |
| 5317 | - | 5789851 | 5791050 | + | *pncB1* | *PA14_64960* | nicotinate phosphoribosyltransferase | 43 | 81 | 56 | 47 | 57 | 43 | 0.605197743 |
| 5318 | - | 5791079 | 5791906 | + | *nadE* | *PA14_64980* | NAD synthetase | 55 | 94 | 57 | 45 | 64 | 42 | 0.790655172 |
| 5319 | - | 5792041 | 5792964 | + | - | *PA14_64990* | hypothetical protein | 60 | 67 | 64 | 56 | 63 | 51 | 1 |
| 5320 | - | 5793479 | 5793033 | - | *azu* | *PA14_65000* | azurin | 1730 | 3090 | 1881 | 1138 | 1980 | 1297 | 1 |
| 5321 | - | 5793755 | 5794342 | + | - | *PA14_65010* | hypothetical protein | 64 | 72 | 67 | 88 | 83 | 80 | 1 |
| 5322 | - | 5795050 | 5794355 | - | - | *PA14_65030* | hypothetical protein | 86 | 91 | 90 | 112 | 92 | 112 | 1 |
| 5323 | - | 5796088 | 5795237 | - | - | *PA14_65040* | hypothetical protein | 49 | 38 | 57 | 85 | 45 | 66 | 1 |
| 5324 | - | 5797274 | 5796339 | - | - | *PA14_65050* | hypothetical protein | 39 | 38 | 36 | 48 | 41 | 44 | 1 |
| 5325 | - | 5799772 | 5797274 | - | - | *PA14_65060* | hypothetical protein | 33 | 32 | 33 | 39 | 32 | 33 | 1 |
| 5326 | - | 5802277 | 5800034 | - | - | *PA14_65080* | hypothetical protein | 36 | 38 | 37 | 38 | 35 | 35 | 1 |
| 5327 | - | 5802370 | 5804391 | + | - | *PA14_65090* | hypothetical protein | 115 | 98 | 121 | 140 | 97 | 135 | 1 |
| 5328 | - | 5805469 | 5804393 | - | *alr* | *PA14_65110* | biosynthetic alanine racemase | 100 | 109 | 135 | 136 | 112 | 128 | 1 |
| 5329 | - | 5806932 | 5805538 | - | *dnaB* | *PA14_65130* | replicative DNA helicase | 63 | 62 | 61 | 57 | 56 | 51 | 1 |
| 5330 | - | 5807507 | 5807061 | - | *rplI* | *PA14_65150* | 50S ribosomal protein L9 | 473 | 657 | 455 | 334 | 525 | 240 | 1 |
| 5331 | - | 5808398 | 5807529 | - | - | *PA14_65160* | hypothetical protein | 481 | 533 | 584 | 287 | 439 | 215 | 1 |
| 5332 | - | 5808665 | 5808435 | - | *rpsR* | *PA14_65170* | 30S ribosomal protein S18 | 350 | 621 | 335 | 229 | 302 | 171 | 0.950259107 |
| 5333 | - | 5809114 | 5808695 | - | *rpsF* | *PA14_65180* | 30S ribosomal protein S6 | 380 | 629 | 408 | 271 | 363 | 200 | 1 |
| 5334 | - | 5810079 | 5809333 | - | - | *PA14_65190* | TrmH family RNA methyltransferase , group 3 | 215 | 212 | 186 | 210 | 227 | 211 | 1 |
| 5335 | - | 5812799 | 5810076 | - | *rnr* | *PA14_65200* | exoribonuclease RNase R | 340 | 304 | 296 | 286 | 313 | 308 | 1 |
| 5336 | - | 5813004 | 5813087 | + | - | *PA14_65210* | Leu tRNA | 384 | 269 | 465 | 321 | 135 | 166 | 1 |
| 5337 | - | 5813220 | 5813303 | + | - | *PA14_65220* | Leu tRNA | 268 | 201 | 333 | 205 | 93 | 108 | 1 |
| 5338 | - | 5814755 | 5813463 | - | *purA* | *PA14_65230* | adenylosuccinate synthetase | 327 | 351 | 293 | 293 | 312 | 272 | 1 |
| 5339 | - | 5815991 | 5814807 | - | *hisZ* | *PA14_65250* | ATP phosphoribosyltransferase | 305 | 227 | 284 | 266 | 255 | 247 | 1 |
| 5340 | - | 5816211 | 5816026 | - | - | *PA14_65260* | hypothetical protein | 233 | 176 | 285 | 262 | 253 | 239 | 1 |
| 5341 | - | 5817176 | 5816307 | - | *hflC* | *PA14_65270* | protease subunit HflC | 659 | 764 | 617 | 632 | 769 | 649 | 1 |
| 5342 | - | 5818378 | 5817176 | - | *hflK* | *PA14_65280* | protease subunit HflK | 696 | 702 | 736 | 714 | 713 | 666 | 1 |
| 5343 | - | 5819774 | 5818473 | - | *hflX* | *PA14_65300* | GTP-binding protein | 521 | 577 | 593 | 510 | 590 | 481 | 1 |
| 5344 | - | 5820035 | 5819787 | - | *hfq* | *PA14_65310* | RNA-binding protein Hfq | 1079 | 1620 | 1264 | 958 | 1205 | 989 | 1 |
| 5345 | - | 5821112 | 5820141 | - | *miaA* | *PA14_65320* | tRNA delta(2)-isopentenylpyrophosphate transferase | 612 | 563 | 786 | 542 | 497 | 473 | 1 |
| 5346 | - | 5823071 | 5821170 | - | *mutL* | *PA14_65350* | DNA mismatch repair protein | 304 | 278 | 308 | 329 | 277 | 310 | 1 |
| 5347 | - | 5824498 | 5823071 | - | *amiB* | *PA14_65370* | N-acetylmuramoyl-L-alanine amidase | 199 | 167 | 203 | 260 | 191 | 225 | 1 |
| 5348 | - | 5824974 | 5824507 | - | - | *PA14_65380* | hypothetical protein | 131 | 124 | 126 | 143 | 122 | 125 | 1 |
| 5349 | - | 5826470 | 5824962 | - | - | *PA14_65390* | hypothetical protein | 144 | 116 | 150 | 131 | 116 | 110 | 1 |
| 5350 | - | 5826552 | 5827637 | + | - | *PA14_65400* | iron-sulfur cluster-binding protein | 95 | 83 | 102 | 97 | 89 | 99 | 1 |
| 5351 | - | 5828213 | 5827671 | - | *orn* | *PA14_65410* | oligoribonuclease | 175 | 179 | 221 | 185 | 183 | 189 | 1 |
| 5352 | - | 5828325 | 5829344 | + | - | *PA14_65420* | ribosome-associated GTPase | 183 | 169 | 205 | 238 | 186 | 200 | 1 |
| 5353 | - | 5830391 | 5829348 | - | *motB* | *PA14_65430* | flagellar motor protein MotB | 144 | 160 | 153 | 166 | 143 | 144 | 1 |
| 5354 | - | 5831262 | 5830411 | - | *motA* | *PA14_65450* | flagellar motor protein MotA | 116 | 83 | 130 | 91 | 108 | 110 | 1 |
| 5355 | - | 5831403 | 5832914 | + | - | *PA14_65470* | hypothetical protein | 58 | 63 | 68 | 63 | 56 | 66 | 1 |
| 5356 | - | 5832989 | 5833804 | + | *rhdA* | *PA14_65480* | thiosulfate sulfurtransferase | 35 | 30 | 33 | 27 | 30 | 34 | 1 |
| 5357 | - | 5833808 | 5834677 | + | *psd* | *PA14_65500* | phosphatidylserine decarboxylase | 63 | 67 | 60 | 55 | 66 | 54 | 1 |
| 5358 | - | 5835355 | 5836818 | + | - | *PA14_65520* | hypothetical protein | 155 | 159 | 150 | 148 | 141 | 158 | 1 |
| 5359 | - | 5836883 | 5838958 | + | - | *PA14_65540* | hypothetical protein | 126 | 137 | 132 | 132 | 123 | 132 | 1 |
| 5360 | - | 5840334 | 5839045 | - | *serB* | *PA14_65560* | phosphoserine phosphatase | 67 | 60 | 77 | 64 | 61 | 64 | 1 |
| 5361 | - | 5840480 | 5842018 | + | - | *PA14_65570* | hypothetical protein | 86 | 74 | 96 | 72 | 62 | 78 | 1 |
| 5362 | - | 5842015 | 5842551 | + | - | *PA14_65580* | membrane-bound metal-dependent hydrolase | 16 | 23 | 21 | 18 | 15 | 17 | 1 |
| 5363 | - | 5843319 | 5842609 | - | - | *PA14_65590* | hypothetical protein | 152 | 116 | 172 | 149 | 105 | 133 | 1 |
| 5364 | - | 5845876 | 5843612 | - | *parC* | *PA14_65605* | DNA topoisomerase IV subunit A | 187 | 202 | 180 | 173 | 217 | 168 | 1 |
| 5365 | - | 5846408 | 5845884 | - | - | *PA14_65630* | hypothetical protein | 203 | 193 | 210 | 180 | 203 | 185 | 1 |
| 5366 | - | 5847418 | 5846405 | - | - | *PA14_65640* | hypothetical protein | 195 | 201 | 169 | 170 | 194 | 164 | 1 |
| 5367 | - | 5849307 | 5847418 | - | *parE* | *PA14_65660* | DNA topoisomerase IV subunit B | 250 | 262 | 235 | 236 | 275 | 227 | 1 |
| 5368 | - | 5849939 | 5849319 | - | - | *PA14_65670* | hypothetical protein | 73 | 64 | 79 | 78 | 64 | 63 | 1 |
| 5369 | - | 5850890 | 5850072 | - | - | *PA14_65690* | hypothetical protein | 55 | 49 | 60 | 54 | 44 | 42 | 1 |
| 5370 | - | 5851502 | 5851044 | - | - | *PA14_65700* | hypothetical protein | 107 | 109 | 117 | 87 | 128 | 104 | 1 |
| 5371 | - | 5852110 | 5851493 | - | *aspP* | *PA14_65710* | adenosine diphosphate sugar pyrophosphatase | 226 | 209 | 269 | 159 | 221 | 189 | 1 |
| 5372 | - | 5852353 | 5853099 | + | - | *PA14_65720* | lipoprotein | 71 | 53 | 82 | 61 | 64 | 65 | 1 |
| 5373 | - | 5855041 | 5853158 | - | *thiC* | *PA14_65740* | thiamine biosynthesis protein ThiC | 22 | 27 | 23 | 23 | 22 | 22 | 1 |
| 5374 | - | 5855497 | 5856945 | + | - | *PA14_65750* | outer membrane efflux protein | 80 | 95 | 106 | 99 | 96 | 97 | 1 |
| 5375 | - | 5857715 | 5857011 | - | - | *PA14_65760* | NAD(P)H quinone oxidoreductase | 65 | 67 | 69 | 73 | 65 | 76 | 1 |
| 5376 | - | 5858947 | 5857766 | - | *aspC* | *PA14_65770* | aspartate transaminase | 40 | 37 | 40 | 39 | 40 | 43 | 1 |

| | A | B | C | D | E | F | G | H | I | J | K | L | M | N |
|---|---|---|---|---|---|---|---|---|---|---|---|---|---|---|
| 5377 | - | 5860649 | 5858970 | - | - | PA14_65795 | hypothetical protein | 21 | 23 | 22 | 22 | 21 | 24 | 1 |
| 5378 | - | 5862782 | 5860692 | - | - | PA14_65810 | hypothetical protein | 10 | 12 | 10 | 10 | 10 | 11 | 1 |
| 5379 | - | 5864009 | 5862849 | - | - | PA14_65820 | acyl-CoA dehydrogenase | 10 | 13 | 10 | 11 | 11 | 10 | 1 |
| 5380 | - | 5864821 | 5864030 | - | - | PA14_65840 | enoyl-CoA hydratase/isomerase | 9 | 13 | 13 | 9 | 12 | 10 | 1 |
| 5381 | - | 5866492 | 5865080 | - | - | PA14_65850 | amino acid ABC transporter permease | 8 | 8 | 9 | 9 | 10 | 8 | 1 |
| 5382 | - | 5868988 | 5866676 | - | - | PA14_65860 | two-component sensor | 32 | 33 | 33 | 36 | 30 | 30 | 1 |
| 5383 | - | 5869701 | 5868925 | - | - | PA14_65870 | hypothetical protein | 31 | 35 | 31 | 38 | 29 | 32 | 1 |
| 5384 | - | 5870439 | 5869705 | - | - | PA14_65880 | two-component response regulator | 73 | 79 | 65 | 80 | 71 | 70 | 1 |
| 5385 | - | 5871204 | 5870557 | - | - | PA14_65900 | TetR family transcriptional regulator | 92 | 76 | 86 | 76 | 72 | 75 | 1 |
| 5386 | - | 5872372 | 5871281 | - | - | PA14_65920 | hypothetical protein | 15 | 15 | 11 | 13 | 11 | 20 | 1 |
| 5387 | - | 5872546 | 5874492 | + | - | PA14_65940 | oxidoreductase | 30 | 33 | 29 | 31 | 26 | 36 | 1 |
| 5388 | - | 5874570 | 5875169 | + | - | PA14_65950 | transcriptional regulator | 66 | 54 | 64 | 72 | 63 | 61 | 1 |
| 5389 | - | 5876495 | 5875218 | - | waaA | PA14_65960 | 3-deoxy-D-manno-octulosonic-acid transferase | 62 | 63 | 65 | 65 | 53 | 52 | 1 |
| 5390 | - | 5877415 | 5876531 | - | - | PA14_65970 | transcriptional regulator | 22 | 19 | 21 | 18 | 17 | 17 | 1 |
| 5391 | - | 5877500 | 5877832 | + | qacH | PA14_65990 | SMR multidrug efflux transporter | 15 | 13 | 15 | 15 | 16 | 12 | 1 |
| 5392 | - | 5877893 | 5879068 | + | - | PA14_66000 | hypothetical protein | 51 | 42 | 53 | 47 | 45 | 43 | 1 |
| 5393 | - | 5879065 | 5879877 | + | - | PA14_66010 | hypothetical protein | 48 | 50 | 45 | 49 | 42 | 41 | 1 |
| 5394 | - | 5880046 | 5880966 | + | - | PA14_66020 | hypothetical protein | 43 | 38 | 44 | 48 | 40 | 42 | 1 |
| 5395 | - | 5882199 | 5880991 | - | - | PA14_66040 | acyl-CoA dehydrogenase | 17 | 17 | 18 | 19 | 19 | 20 | 1 |
| 5396 | - | 5883537 | 5882248 | - | - | PA14_66050 | acyl-CoA dehydrogenase | 18 | 20 | 19 | 19 | 17 | 19 | 1 |
| 5397 | - | 5885089 | 5883665 | - | rfaE | PA14_66060 | bifunctional heptose 7-phosphate kinase/heptose 1-phosphate adenyltransferase | 260 | 247 | 302 | 230 | 266 | 225 | 1 |
| 5398 | - | 5886941 | 5885130 | - | msbA | PA14_66080 | transport protein MsbA | 110 | 98 | 113 | 95 | 103 | 83 | 1 |
| 5399 | - | 5887049 | 5887699 | + | - | PA14_66090 | hypothetical protein | 54 | 47 | 65 | 50 | 44 | 51 | 1 |
| 5400 | - | 5887707 | 5888906 | + | - | PA14_66100 | hypothetical protein | 45 | 42 | 45 | 41 | 41 | 39 | 1 |
| 5401 | - | 5889815 | 5888931 | - | - | PA14_66110 | glycosyl transferase family protein | 332 | 276 | 348 | 271 | 308 | 311 | 1 |
| 5402 | - | 5890932 | 5889976 | - | - | PA14_66120 | hypothetical protein | 401 | 403 | 410 | 427 | 472 | 390 | 1 |
| 5403 | - | 5892423 | 5891005 | - | - | PA14_66140 | hypothetical protein | 136 | 119 | 141 | 128 | 148 | 124 | 1 |
| 5404 | - | 5893323 | 5892427 | - | - | PA14_66150 | hypothetical protein | 256 | 251 | 278 | 250 | 268 | 235 | 1 |
| 5405 | - | 5894453 | 5893317 | - | - | PA14_66160 | glycosyl transferase family protein | 352 | 378 | 356 | 337 | 322 | 294 | 1 |
| 5406 | - | 5896194 | 5894437 | - | - | PA14_66170 | carbamoyl transferase | 409 | 390 | 410 | 422 | 430 | 406 | 1 |
| 5407 | - | 5897776 | 5896298 | - | - | PA14_66190 | hypothetical protein | 233 | 222 | 226 | 218 | 229 | 186 | 1 |
| 5408 | - | 5898531 | 5897773 | - | - | PA14_66200 | hypothetical protein | 144 | 153 | 148 | 132 | 150 | 121 | 1 |
| 5409 | - | 5899262 | 5898528 | - | - | PA14_66210 | hypothetical protein | 259 | 241 | 257 | 251 | 277 | 246 | 1 |
| 5410 | - | 5900068 | 5899262 | - | waaP | PA14_66220 | lipopolysaccharide kinase WaaP | 158 | 152 | 146 | 154 | 173 | 142 | 1 |
| 5411 | - | 5901186 | 5900065 | - | waaG | PA14_66230 | UDP-glucose:(heptosyl) LPS alpha 1,3-glucosyltransferase WaaG | 230 | 237 | 226 | 231 | 266 | 220 | 1 |
| 5412 | - | 5902250 | 5901186 | - | waaC | PA14_66240 | lipopolysaccharide heptosyltransferase I | 246 | 240 | 263 | 251 | 265 | 228 | 1 |
| 5413 | - | 5903284 | 5902247 | - | waaF | PA14_66250 | heptosyltransferase II | 300 | 265 | 308 | 302 | 289 | 257 | 1 |
| 5414 | - | 5904267 | 5903344 | - | ilvE | PA14_66260 | branched-chain amino acid aminotransferase | 455 | 464 | 498 | 486 | 498 | 414 | 1 |
| 5415 | - | 5907271 | 5904323 | - | glnE | PA14_66270 | bifunctional glutamine-synthetase adenylyltransferase/deadenyltransferase | 114 | 110 | 108 | 116 | 110 | 106 | 1 |
| 5416 | - | 5907553 | 5910201 | + | aceE | PA14_66290 | pyruvate dehydrogenase subunit E1 | 302 | 367 | 283 | 334 | 312 | 312 | 1 |
| 5417 | - | 5910346 | 5911989 | + | aceF | PA14_66310 | dihydrolipoamide acetyltransferase | 228 | 272 | 211 | 240 | 241 | 231 | 1 |
| 5418 | - | 5912407 | 5915106 | + | - | PA14_66320 | hypothetical protein | 91 | 89 | 92 | 99 | 101 | 104 | 1 |
| 5419 | - | 5915201 | 5915848 | + | msrA | PA14_66330 | peptide methionine sulfoxide reductase | 154 | 188 | 157 | 159 | 160 | 162 | 1 |
| 5420 | - | 5916688 | 5915852 | - | - | PA14_66340 | hypothetical protein | 112 | 116 | 129 | 115 | 112 | 139 | 1 |
| 5421 | - | 5916990 | 5918792 | + | - | PA14_66350 | acyl-CoA dehydrogenase | 29 | 28 | 33 | 31 | 28 | 32 | 1 |
| 5422 | - | 5919211 | 5920827 | + | - | PA14_66380 | potassium/proton antiporter | 35 | 34 | 36 | 29 | 34 | 32 | 1 |
| 5423 | - | 5920919 | 5924275 | + | - | PA14_66400 | potassium efflux protein KefA | 44 | 51 | 46 | 45 | 48 | 42 | 1 |
| 5424 | - | 5924342 | 5925802 | + | - | PA14_66410 | hypothetical protein | 32 | 42 | 20 | 37 | 53 | 45 | 1 |
| 5425 | - | 5926208 | 5926927 | + | - | PA14_66420 | hypothetical protein | 31 | 25 | 28 | 31 | 24 | 27 | 1 |
| 5426 | - | 5927048 | 5928325 | + | metY | PA14_66440 | O-acetylhomoserine aminocarboxypropyltransferase | 25 | 35 | 29 | 22 | 37 | 28 | 1 |
| 5427 | - | 5928416 | 5928868 | + | - | PA14_66450 | hypothetical protein | 21 | 19 | 19 | 17 | 18 | 18 | 1 |
| 5428 | - | 5928981 | 5929796 | + | - | PA14_66460 | hypothetical protein | 605 | 373 | 525 | 217 | 237 | 614 | 1 |
| 5429 | - | 5930585 | 5929818 | - | - | PA14_66480 | hypothetical protein | 215 | 168 | 205 | 190 | 169 | 253 | 1 |
| 5430 | - | 5931581 | 5930676 | - | - | PA14_66490 | LysR family transcriptional regulator | 22 | 17 | 17 | 21 | 20 | 21 | 1 |
| 5431 | - | 5931759 | 5933075 | + | - | PA14_66510 | MFS transporter | 25 | 8 | 20 | 13 | 9 | 13 | 0.003830572 |
| 5432 | - | 5934032 | 5933103 | - | - | PA14_66520 | short chain dehydrogenase | 13 | 13 | 11 | 14 | 12 | 12 | 1 |
| 5433 | - | 5934144 | 5935217 | + | - | PA14_66530 | transcriptional regulator | 17 | 18 | 19 | 22 | 19 | 19 | 1 |
| 5434 | - | 5935291 | 5936271 | + | - | PA14_66540 | hypothetical protein | 11 | 17 | 15 | 20 | 15 | 14 | 1 |
| 5435 | - | 5937459 | 5936392 | - | hemE | PA14_66550 | uroporphyrinogen decarboxylase | 34 | 34 | 38 | 28 | 32 | 26 | 1 |
| 5436 | - | 5939069 | 5937636 | - | gltD | PA14_66560 | glutamate synthase subunit beta | 94 | 72 | 70 | 63 | 74 | 65 | 1 |
| 5437 | - | 5943543 | 5939098 | - | gltB | PA14_66570 | glutamate synthase subunit alpha | 76 | 58 | 67 | 61 | 67 | 60 | 1 |
| 5438 | - | 5945423 | 5943768 | - | - | PA14_66580 | hypothetical protein | 256 | 253 | 225 | 241 | 215 | 214 | 1 |
| 5439 | - | 5946536 | 5945430 | - | aroB | PA14_66600 | 3-dehydroquinate synthase | 153 | 144 | 144 | 144 | 140 | 137 | 1 |
| 5440 | - | 5947104 | 5946586 | - | aroK | PA14_66610 | shikimate kinase | 249 | 246 | 257 | 248 | 230 | 233 | 1 |
| 5441 | - | 5949248 | 5947116 | - | pilQ | PA14_66620 | type 4 fimbrial biogenesis outer membrane protein PilQ precursor | 371 | 510 | 385 | 446 | 444 | 426 | 1 |
| 5442 | - | 5949826 | 5949302 | - | pilP | PA14_66630 | type 4 fimbrial biogenesis protein PilP | 195 | 261 | 200 | 224 | 224 | 212 | 1 |
| 5443 | - | 5950446 | 5949823 | - | pilO | PA14_66640 | type 4 fimbrial biogenesis protein PilO | 309 | 373 | 322 | 354 | 370 | 344 | 1 |
| 5444 | - | 5951039 | 5950443 | - | pilN | PA14_66650 | type 4 fimbrial biogenesis protein PilN | 482 | 534 | 523 | 510 | 537 | 534 | 1 |
| 5445 | - | 5952103 | 5951039 | - | pilM | PA14_66660 | type 4 fimbrial biogenesis protein PilM | 577 | 606 | 629 | 575 | 600 | 563 | 1 |
| 5446 | - | 5952288 | 5954759 | + | ponA | PA14_66670 | penicillin-binding protein 1A | 69 | 65 | 86 | 84 | 73 | 74 | 1 |
| 5447 | - | 5956123 | 5954855 | - | maeB | PA14_66680 | malic enzyme | 117 | 236 | 134 | 118 | 172 | 107 | 1 |
| 5448 | - | 5957665 | 5956226 | - | - | PA14_66690 | protease | 116 | 115 | 149 | 139 | 134 | 144 | 1 |
| 5449 | - | 5958429 | 5957662 | - | - | PA14_66700 | nuclease | 97 | 97 | 132 | 113 | 111 | 108 | 1 |
| 5450 | - | 5958704 | 5958489 | - | rpmE | PA14_66710 | 50S ribosomal protein L31 | 225 | 272 | 227 | 177 | 204 | 129 | 1 |
| 5451 | - | 5958880 | 5961099 | + | priA | PA14_66720 | primosome assembly protein PriA | 20 | 24 | 20 | 21 | 20 | 18 | 1 |
| 5452 | - | 5961348 | 5963111 | + | argS | PA14_66750 | arginyl-tRNA synthetase | 142 | 149 | 166 | 117 | 136 | 134 | 1 |
| 5453 | - | 5963146 | 5963841 | + | - | PA14_66760 | hypothetical protein | 88 | 98 | 101 | 76 | 88 | 79 | 1 |
| 5454 | - | 5963960 | 5964493 | + | hslV | PA14_66770 | ATP-dependent protease peptidase subunit | 243 | 225 | 388 | 116 | 233 | 172 | 1 |
| 5455 | - | 5964522 | 5965865 | + | hslU | PA14_66790 | ATP-dependent protease ATP-binding subunit HslU | 153 | 196 | 195 | 79 | 187 | 120 | 1 |
| 5456 | - | 5965960 | 5966331 | + | - | PA14_66800 | hypothetical protein | 55 | 71 | 67 | 52 | 56 | 44 | 1 |
| 5457 | - | 5966750 | 5968429 | + | phaC1 | PA14_66820 | poly(3-hydroxyalkanoic acid) synthase 1 | 99 | 118 | 90 | 134 | 108 | 148 | 1 |
| 5458 | - | 5968582 | 5969439 | + | phaD | PA14_66830 | poly(3-hydroxyalkanoic acid) depolymerase | 131 | 152 | 122 | 182 | 142 | 207 | 1 |
| 5459 | - | 5969743 | 5971425 | + | phaC2 | PA14_66840 | poly(3-hydroxyalkanoic acid) synthase 2 | 69 | 65 | 74 | 123 | 68 | 102 | 1 |
| 5460 | - | 5971481 | 5972098 | + | - | PA14_66850 | TetR family transcriptional regulator | 64 | 72 | 73 | 118 | 77 | 116 | 1 |

| | A | B | C | D | E | F | G | H | I | J | K | L | M | N |
|---|---|---|---|---|---|---|---|---|---|---|---|---|---|---|
| 5461 | - | 5973071 | 5972142 | - | *phaF* | *PA14_66875* | polyhydroxyalkanoate synthesis protein PhaF | 790 | 852 | 713 | 655 | 784 | 774 | 1 |
| 5462 | - | 5973498 | 5973082 | - | - | *PA14_66880* | hypothetical protein | 129 | 117 | 111 | 119 | 112 | 140 | 1 |
| 5463 | - | 5973917 | 5973642 | - | - | *PA14_66890* | hypothetical protein | 53 | 46 | 65 | 47 | 50 | 45 | 1 |
| 5464 | - | 5974053 | 5974823 | + | *ubiE* | *PA14_66900* | ubiquinone/menaquinone biosynthesis methyltransferase | 137 | 114 | 141 | 104 | 115 | 109 | 1 |
| 5465 | - | 5974838 | 5975464 | + | - | *PA14_66910* | hypothetical protein | 93 | 91 | 86 | 99 | 97 | 91 | 1 |
| 5466 | - | 5975461 | 5977062 | + | *ubiB* | *PA14_66920* | ubiquinone biosynthesis protein UbiB | 82 | 76 | 81 | 90 | 80 | 77 | 1 |
| 5467 | - | 5977178 | 5977582 | + | *hisI* | *PA14_66940* | phosphoribosyl-AMP cyclohydrolase | 247 | 180 | 278 | 208 | 236 | 249 | 1 |
| 5468 | - | 5977575 | 5977910 | + | *hisE* | *PA14_66950* | phosphoribosyl-ATP pyrophosphatase | 189 | 158 | 217 | 165 | 173 | 183 | 1 |
| 5469 | - | 5977936 | 5978184 | + | *tatA* | *PA14_66960* | twin arginine translocase A | 176 | 228 | 190 | 190 | 182 | 191 | 1 |
| 5470 | - | 5978198 | 5978623 | + | *tatB* | *PA14_66970* | sec-independent translocase | 145 | 143 | 136 | 135 | 148 | 130 | 1 |
| 5471 | - | 5978620 | 5979423 | + | *tatC* | *PA14_66980* | sec-independent protein translocase TatC | 74 | 64 | 69 | 76 | 80 | 69 | 1 |
| 5472 | - | 5979420 | 5980127 | + | - | *PA14_66990* | 16S ribosomal RNA methyltransferase RsmE | 37 | 31 | 33 | 32 | 30 | 28 | 1 |
| 5473 | - | 5980339 | 5982282 | + | - | *PA14_67010* | chemotaxis transducer | 16 | 18 | 18 | 15 | 16 | 13 | 1 |
| 5474 | - | 5982392 | 5982853 | + | - | *PA14_67020* | hypothetical protein | 33 | 37 | 39 | 36 | 31 | 33 | 1 |
| 5475 | - | 5983595 | 5982861 | - | - | *PA14_67030* | ABC transporter ATP-binding protein | 53 | 64 | 51 | 49 | 61 | 43 | 1 |
| 5476 | - | 5984550 | 5983588 | - | - | *PA14_67040* | ABC transporter permease | 24 | 30 | 23 | 21 | 24 | 17 | 1 |
| 5477 | - | 5985416 | 5984616 | - | - | *PA14_67050* | ABC transporter substrate-binding protein | 39 | 47 | 40 | 33 | 42 | 32 | 1 |
| 5478 | - | 5988239 | 5985654 | - | *mdoH* | *PA14_67065* | glucosyltransferase MdoH | 145 | 145 | 137 | 152 | 153 | 149 | 1 |
| 5479 | - | 5989809 | 5988232 | - | *mdoG* | *PA14_67090* | glucan biosynthesis protein G | 175 | 190 | 155 | 159 | 180 | 151 | 1 |
| 5480 | - | 5990703 | 5990266 | - | - | *PA14_67100* | D-tyrosyl-tRNA(Tyr) deacylase | 137 | 126 | 110 | 138 | 128 | 116 | 1 |
| 5481 | - | 5991671 | 5990700 | - | - | *PA14_67110* | prolyl aminopeptidase | 78 | 79 | 66 | 70 | 74 | 68 | 1 |
| 5482 | - | 5991853 | 5992329 | + | - | *PA14_67120* | hypothetical protein | 29 | 27 | 26 | 30 | 26 | 26 | 1 |
| 5483 | - | 5993233 | 5992334 | - | - | *PA14_67130* | ABC transporter substrate-binding protein | 17 | 21 | 15 | 20 | 22 | 19 | 1 |
| 5484 | - | 5993626 | 5993276 | - | - | *PA14_67140* | hypothetical protein | 107 | 90 | 96 | 94 | 77 | 78 | 1 |
| 5485 | - | 5994901 | 5993651 | - | - | *PA14_67150* | oxidoreductase | 34 | 32 | 32 | 33 | 32 | 32 | 1 |
| 5486 | - | 5994966 | 5995922 | + | - | *PA14_67170* | LysR family transcriptional regulator | 67 | 52 | 62 | 56 | 64 | 54 | 1 |
| 5487 | - | 5996854 | 5995985 | - | - | *PA14_67180* | hypothetical protein | 27 | 22 | 20 | 19 | 28 | 21 | 1 |
| 5488 | - | 5997744 | 5996863 | - | - | *PA14_67190* | hypothetical protein | 15 | 9 | 9 | 10 | 12 | 11 | 0.73766974 |
| 5489 | - | 5998610 | 5997741 | - | - | *PA14_67200* | hypothetical protein | 28 | 18 | 17 | 19 | 22 | 21 | 1 |
| 5490 | - | 5999494 | 5998619 | - | - | *PA14_67210* | hypothetical protein | 38 | 23 | 28 | 20 | 34 | 28 | 1 |
| 5491 | - | 6001735 | 5999498 | - | - | *PA14_67220* | hypothetical protein | 34 | 23 | 24 | 27 | 31 | 26 | 1 |
| 5492 | - | 6004104 | 6001732 | - | - | *PA14_67230* | hypothetical protein | 19 | 14 | 16 | 16 | 16 | 14 | 1 |
| 5493 | - | 6005174 | 6004374 | - | *hutG* | *PA14_67240* | N-formylglutamate amidohydrolase | 54 | 71 | 54 | 81 | 60 | 77 | 1 |
| 5494 | - | 6006375 | 6005167 | - | *hutI* | *PA14_67250* | imidazolonepropionase | 66 | 73 | 67 | 91 | 70 | 85 | 1 |
| 5495 | - | 6007904 | 6006372 | - | - | *PA14_67260* | histidine/phenylalanine ammonia-lyase | 58 | 64 | 62 | 85 | 65 | 75 | 1 |
| 5496 | - | 6008731 | 6007901 | - | - | *PA14_67270* | ABC transporter ATP-binding protein | 45 | 46 | 40 | 57 | 47 | 57 | 1 |
| 5497 | - | 6009579 | 6008728 | - | - | *PA14_67280* | ABC transporter permease | 27 | 30 | 30 | 41 | 31 | 35 | 1 |
| 5498 | - | 6010572 | 6009604 | - | - | *PA14_67300* | ABC transporter substrate-binding protein | 20 | 24 | 28 | 35 | 29 | 28 | 1 |
| 5499 | - | 6012093 | 6010690 | - | - | *PA14_67310* | amino acid permease | 10 | 18 | 13 | 21 | 16 | 15 | 0.721087348 |
| 5500 | - | 6013685 | 6012156 | - | *hutH* | *PA14_67320* | histidine ammonia-lyase | 10 | 19 | 15 | 23 | 18 | 15 | 0.453215346 |
| 5501 | - | 6015224 | 6013782 | - | - | *PA14_67340* | cytosine/purines uracil thiamine allantoin permease | 12 | 18 | 16 | 18 | 18 | 17 | 1 |
| 5502 | - | 6017011 | 6015332 | - | *hutU* | *PA14_67350* | urocanate hydratase | 51 | 64 | 67 | 83 | 83 | 105 | 1 |
| 5503 | - | 6018133 | 6017336 | - | - | *PA14_67370* | hypothetical protein | 61 | 54 | 81 | 91 | 66 | 82 | 1 |
| 5504 | - | 6019090 | 6018152 | - | - | *PA14_67380* | fatty acid desaturase | 8 | 8 | 8 | 8 | 6 | 7 | 1 |
| 5505 | - | 6020128 | 6019106 | - | - | *PA14_67400* | ABC transporter substrate-binding protein | 56 | 56 | 64 | 63 | 65 | 73 | 1 |
| 5506 | - | 6020860 | 6020270 | - | - | *PA14_67410* | hypothetical protein | 115 | 123 | 141 | 132 | 129 | 154 | 1 |
| 5507 | - | 6021609 | 6020857 | - | *hutC* | *PA14_67420* | histidine utilization genes repressor protein | 119 | 122 | 140 | 117 | 150 | 155 | 1 |
| 5508 | - | 6021710 | 6023071 | + | - | *PA14_67440* | N-formimino-L-glutamate deiminase | 93 | 117 | 109 | 121 | 177 | 196 | 1 |
| 5509 | - | 6023783 | 6023214 | - | *blc* | *PA14_67450* | outer membrane lipoprotein Blc | 117 | 117 | 147 | 126 | 137 | 125 | 1 |
| 5510 | - | 6024037 | 6023780 | - | - | *PA14_67460* | lipoprotein | 151 | 144 | 164 | 127 | 152 | 154 | 1 |
| 5511 | - | 6024712 | 6024110 | - | - | *PA14_67470* | hypothetical protein | 163 | 137 | 181 | 129 | 135 | 128 | 1 |
| 5512 | - | 6025727 | 6024717 | - | *fbp* | *PA14_67490* | fructose-1,6-bisphosphatase | 214 | 160 | 240 | 167 | 175 | 192 | 1 |
| 5513 | - | 6026397 | 6025867 | - | *gloA3* | *PA14_67500* | lactoylglutathione lyase | 83 | 99 | 81 | 76 | 109 | 100 | 1 |
| 5514 | - | 6028491 | 6026551 | - | *estA* | *PA14_67510* | esterase EstA | 185 | 198 | 220 | 209 | 191 | 216 | 1 |
| 5515 | - | 6029994 | 6028600 | - | - | *PA14_67520* | hypothetical protein | 133 | 108 | 107 | 104 | 122 | 109 | 1 |
| 5516 | - | 6033596 | 6029991 | - | - | *PA14_67530* | hypothetical protein | 77 | 61 | 64 | 77 | 70 | 60 | 1 |
| 5517 | - | 6034319 | 6033735 | - | - | *PA14_67540* | hypothetical protein | 102 | 85 | 79 | 108 | 97 | 88 | 1 |
| 5518 | - | 6034403 | 6034828 | + | - | *PA14_67550* | transcriptional regulator | 58 | 54 | 69 | 73 | 52 | 62 | 1 |
| 5519 | - | 6036729 | 6034912 | - | *typA* | *PA14_67560* | GTP-binding protein TypA | 72 | 80 | 86 | 65 | 81 | 53 | 1 |
| 5520 | - | 6038400 | 6036946 | - | *thiI* | *PA14_67580* | thiamine biosynthesis protein ThiI | 36 | 28 | 59 | 36 | 27 | 24 | 1 |
| 5521 | - | 6038737 | 6040146 | + | *glnA* | *PA14_67600* | glutamine synthetase | 212 | 319 | 215 | 198 | 286 | 243 | 1 |
| 5522 | - | 6040294 | 6040698 | + | - | *PA14_67620* | hypothetical protein | 69 | 54 | 61 | 56 | 51 | 52 | 1 |
| 5523 | - | 6042902 | 6040695 | - | - | *PA14_67630* | hypothetical protein | 35 | 30 | 31 | 35 | 31 | 30 | 1 |
| 5524 | - | 6043190 | 6043711 | + | - | *PA14_67640* | hypothetical protein | 77 | 68 | 89 | 84 | 72 | 74 | 1 |
| 5525 | - | 6043708 | 6044280 | + | - | *PA14_67650* | hypothetical protein | 33 | 34 | 44 | 35 | 32 | 34 | 1 |
| 5526 | - | 6044551 | 6045627 | + | *ntrB* | *PA14_67670* | two-component sensor NtrB | 41 | 34 | 39 | 42 | 31 | 34 | 1 |
| 5527 | - | 6045630 | 6047060 | + | *ntrC* | *PA14_67680* | two-component response regulator NtrC | 31 | 36 | 35 | 39 | 33 | 36 | 1 |
| 5528 | - | 6048249 | 6048710 | + | - | *PA14_67710* | rRNA methylase | 21 | 28 | 26 | 21 | 23 | 16 | 1 |
| 5529 | - | 6048250 | 6047783 | - | - | *PA14_67700* | hypothetical protein | 46 | 45 | 50 | 41 | 51 | 39 | 1 |
| 5530 | - | 6049260 | 6048769 | - | *secB* | *PA14_67720* | preprotein translocase subunit SecB | 81 | 154 | 74 | 46 | 89 | 54 | 0.569926259 |
| 5531 | - | 6049551 | 6049297 | - | *grx* | *PA14_67740* | glutaredoxin | 37 | 34 | 44 | 28 | 31 | 27 | 1 |
| 5532 | - | 6049972 | 6049553 | - | - | *PA14_67750* | rhodanese-like domain-containing protein | 52 | 33 | 51 | 36 | 30 | 34 | 1 |
| 5533 | - | 6050297 | 6051844 | + | *pgm* | *PA14_67770* | phosphoglyceromutase | 144 | 115 | 116 | 107 | 114 | 127 | 1 |
| 5534 | - | 6052158 | 6052976 | + | - | *PA14_67780* | hypothetical protein | 9 | 7 | 8 | 10 | 6 | 6 | 1 |
| 5535 | - | 6053126 | 6054412 | + | - | *PA14_67790* | membrane-bound metallopeptidase | 100 | 84 | 102 | 97 | 87 | 81 | 1 |
| 5536 | - | 6054441 | 6055751 | + | - | *PA14_67810* | carboxyl-terminal protease | 180 | 203 | 168 | 201 | 211 | 186 | 1 |
| 5537 | - | 6055751 | 6056524 | + | - | *PA14_67820* | hypothetical protein | 63 | 62 | 57 | 55 | 65 | 54 | 1 |
| 5538 | - | 6056593 | 6058074 | + | - | *PA14_67830* | hypothetical protein | 38 | 39 | 34 | 31 | 45 | 27 | 1 |
| 5539 | - | 6058870 | 6058115 | - | - | *PA14_67840* | ABC-type amino acid transporter | 40 | 61 | 41 | 43 | 45 | 35 | 1 |
| 5540 | - | 6059772 | 6059020 | - | - | *PA14_67850* | ABC-type amino acid transport protein, periplasmic component | 16 | 21 | 19 | 16 | 15 | 13 | 1 |
| 5541 | - | 6060609 | 6059863 | - | - | *PA14_67860* | ABC-type amino acid transporter | 27 | 34 | 34 | 28 | 24 | 17 | 1 |
| 5542 | - | 6061551 | 6060781 | - | *hisF1* | *PA14_67880* | imidazole glycerol phosphate synthase subunit HisF | 83 | 103 | 91 | 87 | 104 | 70 | 1 |
| 5543 | - | 6062299 | 6061562 | - | *hisA* | *PA14_67890* | 1-(5-phosphoribosyl)-5-[(5-phosphoribosylamino)methylideneamino] imidazole-4-carboxamide isomerase | 96 | 106 | 98 | 103 | 107 | 81 | 1 |
| 5544 | - | 6062608 | 6062348 | - | - | *PA14_67900* | hypothetical protein | 58 | 59 | 78 | 73 | 68 | 53 | 1 |

| | A | B | C | D | E | F | G | H | I | J | K | L | M | N |
|---|---|---|---|---|---|---|---|---|---|---|---|---|---|---|
| 5545 | - | 6063253 | 6062612 | - | *hisH* | *PA14_67920* | imidazole glycerol phosphate synthase subunit HisH | 187 | 177 | 223 | 211 | 176 | 158 | 1 |
| 5546 | - | 6063843 | 6063250 | - | *hisB* | *PA14_67930* | imidazoleglycerol-phosphate dehydratase | 144 | 118 | 167 | 144 | 130 | 136 | 1 |
| 5547 | - | 6064004 | 6064402 | + | - | *PA14_67940* | hypothetical protein | 11 | 8 | 9 | 10 | 8 | 9 | 1 |
| 5548 | - | 6064439 | 6064675 | + | - | *PA14_67960* | hypothetical protein | 25 | 32 | 25 | 24 | 23 | 22 | 1 |
| 5549 | - | 6064672 | 6065778 | + | - | *PA14_67970* | hypothetical protein | 26 | 27 | 27 | 26 | 24 | 23 | 1 |
| 5550 | - | 6065872 | 6068124 | + | - | *PA14_67975* | hypothetical protein | 77 | 86 | 78 | 78 | 81 | 77 | 1 |
| 5551 | - | 6068121 | 6069188 | + | *mutY* | *PA14_67990* | A/G-specific adenine glycosylase | 109 | 107 | 100 | 103 | 102 | 98 | 1 |
| 5552 | - | 6069232 | 6069504 | + | - | *PA14_68000* | hypothetical protein | 219 | 271 | 223 | 228 | 296 | 251 | 1 |
| 5553 | - | 6070731 | 6070803 | + | - | *PA14_68030* | Phe tRNA | 7 | 8 | 10 | 3 | 3 | 3 | 1 |
| 5554 | - | 6071600 | 6070863 | - | - | *PA14_68040* | short-chain dehydrogenase | 89 | 79 | 86 | 93 | 87 | 92 | 1 |
| 5555 | - | 6071759 | 6072448 | + | - | *PA14_68050* | hypothetical protein | 22 | 20 | 28 | 21 | 15 | 18 | 1 |
| 5556 | - | 6072697 | 6073470 | + | - | *PA14_68060* | ABC transporter ATP-binding protein | 115 | 140 | 134 | 130 | 116 | 124 | 1 |
| 5557 | - | 6073485 | 6074237 | + | - | *PA14_68070* | periplasmic binding protein | 135 | 243 | 121 | 136 | 157 | 124 | 1 |
| 5558 | - | 6074299 | 6074994 | + | - | *PA14_68080* | ABC transporter permease | 27 | 36 | 27 | 25 | 36 | 24 | 1 |
| 5559 | - | 6074991 | 6075683 | + | - | *PA14_68090* | ABC transporter permease | 42 | 63 | 42 | 45 | 48 | 39 | 1 |
| 5560 | - | 6076037 | 6076972 | + | - | *PA14_68100* | hypothetical protein | 16 | 16 | 17 | 16 | 14 | 15 | 1 |
| 5561 | - | 6077376 | 6077846 | + | - | *PA14_68110* | transcriptional regulator | 8 | 7 | 7 | 7 | 6 | 10 | 1 |
| 5562 | - | 6077850 | 6079328 | + | - | *PA14_68120* | outer membrane protein | 26 | 22 | 27 | 27 | 20 | 28 | 1 |
| 5563 | - | 6079343 | 6080527 | + | - | *PA14_68130* | multidrug resistance protein | 29 | 26 | 23 | 28 | 22 | 36 | 1 |
| 5564 | - | 6080538 | 6082067 | + | - | *PA14_68140* | drug efflux transporter | 21 | 15 | 18 | 21 | 17 | 30 | 1 |
| 5565 | - | 6082141 | 6082213 | + | - | *PA14_68150* | Thr tRNA | 9 | 9 | 22 | 10 | 4 | 2 | 1 |
| 5566 | 6082366 | 6082452 | 6083510 | + | *rmlB* | *PA14_68170* | dTDP-D-glucose 4,6-dehydratase | 214 | 92 | 118 | 236 | 192 | 256 | 0.87194538 |
| 5567 | 6082366 | 6083507 | 6084415 | + | *rmlD* | *PA14_68190* | dTDP-4-dehydrorhamnose reductase | 269 | 116 | 140 | 271 | 208 | 258 | 0.924641456 |
| 5568 | 6082366 | 6084412 | 6085293 | + | *rmlA* | *PA14_68200* | glucose-1-phosphate thymidylyltransferase | 347 | 169 | 168 | 369 | 301 | 371 | 1 |
| 5569 | 6082366 | 6085293 | 6085838 | + | *rmlC* | *PA14_68210* | dTDP-4-dehydrorhamnose 3,5-epimerase | 465 | 237 | 197 | 507 | 361 | 499 | 1 |
| 5570 | 6082366 | 6085898 | 6087736 | + | - | *PA14_68230* | two-component sensor | 37 | 27 | 30 | 37 | 40 | 42 | 1 |
| 5571 | 6082366 | 6087733 | 6089121 | + | - | *PA14_68250* | two-component response regulator | 91 | 79 | 73 | 87 | 79 | 83 | 1 |
| 5572 | - | 6089362 | 6090357 | + | - | *PA14_68260* | c4-dicarboxylate-binding protein | 170 | 145 | 97 | 131 | 125 | 144 | 1 |
| 5573 | - | 6090376 | 6091008 | + | - | *PA14_68280* | dicarboxylate transporter | 61 | 40 | 25 | 36 | 36 | 38 | 1 |
| 5574 | - | 6091005 | 6092288 | + | - | *PA14_68290* | C4-dicarboxylate transporter | 76 | 58 | 39 | 54 | 60 | 58 | 1 |
| 5575 | 6093032 | 6093086 | 6094534 | + | *arcD* | *PA14_68300* | arginine/ornithine antiporter | 879 | 1012 | 1228 | 333 | 430 | 952 | 1 |
| 5576 | 6093032 | 6094556 | 6095812 | + | *arcA* | *PA14_68330* | arginine deiminase | 445 | 679 | 637 | 241 | 438 | 468 | 1 |
| 5577 | 6093032 | 6095892 | 6096902 | + | *arcB* | *PA14_68340* | ornithine carbamoyltransferase | 341 | 530 | 482 | 220 | 406 | 347 | 1 |
| 5578 | 6093032 | 6096963 | 6097895 | + | *arcC* | *PA14_68350* | carbamate kinase | 258 | 368 | 344 | 190 | 316 | 230 | 1 |
| 5579 | - | 6098345 | 6100249 | + | - | *PA14_68360* | beta-ketoacyl synthase | 121 | 137 | 120 | 126 | 136 | 113 | 1 |
| 5580 | - | 6101120 | 6100299 | - | *cysQ* | *PA14_68370* | 3'(2'),5'-bisphosphate nucleotidase | 210 | 189 | 189 | 216 | 230 | 220 | 1 |
| 5581 | - | 6101683 | 6101117 | - | *nudE* | *PA14_68380* | ADP-ribose diphosphatase NudE | 180 | 151 | 141 | 176 | 194 | 173 | 1 |
| 5582 | - | 6101809 | 6102474 | + | - | *PA14_68390* | hydrolase | 100 | 110 | 113 | 123 | 105 | 109 | 1 |
| 5583 | - | 6102964 | 6102527 | - | - | *PA14_68400* | LysM domain/BON superfamily protein | 1933 | 1727 | 1674 | 2397 | 2346 | 2194 | 1 |
| 5584 | - | 6103138 | 6104058 | + | - | *PA14_68420* | LysR family transcriptional regulator | 30 | 33 | 33 | 33 | 33 | 31 | 1 |
| 5585 | - | 6104117 | 6104956 | + | - | *PA14_68430* | formate dehydrogenase accessory protein FdhD | 51 | 34 | 53 | 49 | 37 | 63 | 1 |
| 5586 | - | 6104964 | 6107285 | + | - | *PA14_68440* | oxidoreductase | 21 | 14 | 24 | 18 | 16 | 39 | 1 |
| 5587 | - | 6107697 | 6108110 | + | - | *PA14_68450* | hypothetical protein | 258 | 140 | 292 | 216 | 205 | 195 | 1 |
| 5588 | - | 6108208 | 6108615 | + | - | *PA14_68460* | hypothetical protein | 52 | 58 | 76 | 38 | 45 | 44 | 1 |
| 5589 | - | 6108899 | 6108684 | - | - | *PA14_68470* | hypothetical protein | 150 | 151 | 218 | 106 | 116 | 170 | 1 |
| 5590 | - | 6109123 | 6109680 | + | - | *PA14_68480* | chorismate mutase | 50 | 41 | 54 | 47 | 41 | 43 | 1 |
| 5591 | - | 6110105 | 6109662 | - | - | *PA14_68490* | hypothetical protein | 62 | 65 | 66 | 80 | 61 | 81 | 1 |
| 5592 | - | 6111291 | 6110128 | - | - | *PA14_68500* | iron-containing alcohol dehydrogenase | 39 | 45 | 40 | 51 | 44 | 48 | 1 |
| 5593 | - | 6113108 | 6111318 | - | - | *PA14_68510* | acyl-CoA dehydrogenase | 19 | 21 | 17 | 24 | 21 | 22 | 1 |
| 5594 | - | 6114345 | 6113110 | - | - | *PA14_68530* | 3-hydroxyacyl-CoA dehydrogenase | 23 | 25 | 21 | 26 | 22 | 24 | 1 |
| 5595 | - | 6114488 | 6115396 | + | - | *PA14_68550* | LysR family transcriptional regulator | 42 | 43 | 39 | 40 | 39 | 42 | 1 |
| 5596 | - | 6116013 | 6115411 | - | - | *PA14_68560* | nitroreductase | 45 | 65 | 46 | 45 | 49 | 42 | 1 |
| 5597 | - | 6116265 | 6116621 | + | - | *PA14_68570* | hypothetical protein | 24 | 28 | 24 | 31 | 24 | 29 | 1 |
| 5598 | - | 6118219 | 6116678 | - | *pckA* | *PA14_68580* | phosphoenolpyruvate carboxykinase | 132 | 139 | 147 | 97 | 129 | 98 | 1 |
| 5599 | - | 6119228 | 6118335 | - | *hslO* | *PA14_68610* | Hsp33-like chaperonin | 35 | 28 | 39 | 29 | 27 | 25 | 1 |
| 5600 | - | 6119341 | 6120144 | + | - | *PA14_68620* | hypothetical protein | 65 | 65 | 66 | 72 | 69 | 66 | 1 |
| 5601 | - | 6120577 | 6120182 | - | - | *PA14_68630* | heat shock protein | 35 | 46 | 40 | 39 | 36 | 36 | 1 |
| 5602 | - | 6120787 | 6121221 | + | - | *PA14_68640* | hypothetical protein | 36 | 37 | 40 | 37 | 36 | 33 | 1 |
| 5603 | - | 6121353 | 6122258 | + | *rimK* | *PA14_68660* | ribosomal protein S6 modification protein | 49 | 51 | 53 | 49 | 48 | 48 | 1 |
| 5604 | - | 6122449 | 6123372 | + | - | *PA14_68670* | carboxypeptidase | 67 | 78 | 72 | 72 | 71 | 72 | 1 |
| 5605 | - | 6124768 | 6123449 | - | *envZ* | *PA14_68680* | two-component sensor EnvZ | 101 | 102 | 97 | 93 | 106 | 86 | 1 |
| 5606 | - | 6125606 | 6124863 | - | *ompR* | *PA14_68700* | osmolarity response regulator | 151 | 135 | 146 | 102 | 127 | 107 | 1 |
| 5607 | - | 6125795 | 6128134 | + | - | *PA14_68710* | hypothetical protein | 41 | 52 | 45 | 37 | 45 | 31 | 1 |
| 5608 | - | 6128146 | 6128535 | + | - | *PA14_68720* | hypothetical protein | 27 | 38 | 30 | 23 | 27 | 18 | 1 |
| 5609 | - | 6128743 | 6130326 | + | *gshA* | *PA14_68730* | glutamate--cysteine ligase | 71 | 75 | 69 | 60 | 71 | 67 | 1 |
| 5610 | - | 6131667 | 6130369 | - | *argA* | *PA14_68740* | N-acetylglutamate synthase | 80 | 52 | 60 | 52 | 49 | 51 | 1 |
| 5611 | - | 6132004 | 6132705 | + | - | *PA14_68750* | multiple drug resistance protein MarC | 59 | 76 | 58 | 63 | 64 | 70 | 1 |
| 5612 | - | 6133832 | 6132678 | - | *argE* | *PA14_68770* | acetylornithine deacetylase | 141 | 144 | 144 | 134 | 155 | 145 | 1 |
| 5613 | - | 6135200 | 6133932 | - | - | *PA14_68780* | phosphate transporter | 83 | 54 | 95 | 54 | 51 | 70 | 1 |
| 5614 | - | 6135924 | 6135247 | - | - | *PA14_68800* | hypothetical protein | 334 | 197 | 494 | 190 | 164 | 265 | 1 |
| 5615 | - | 6136038 | 6137402 | + | - | *PA14_68810* | hypothetical protein | 67 | 50 | 62 | 56 | 58 | 56 | 1 |
| 5616 | - | 6137440 | 6139224 | + | - | *PA14_68820* | secretion pathway ATPase | 151 | 154 | 174 | 197 | 178 | 195 | 1 |
| 5617 | - | 6139446 | 6139898 | + | - | *PA14_68830* | hypothetical protein | 62 | 60 | 80 | 87 | 68 | 82 | 1 |
| 5618 | - | 6140030 | 6140359 | + | - | *PA14_68840* | hypothetical protein | 280 | 379 | 377 | 336 | 483 | 377 | 1 |
| 5619 | - | 6143270 | 6140394 | - | *gcvP1* | *PA14_68850* | glycine dehydrogenase | 108 | 105 | 112 | 155 | 104 | 155 | 1 |
| 5620 | - | 6143833 | 6143444 | - | *gcvH1* | *PA14_68860* | glycine cleavage system protein H | 100 | 152 | 91 | 87 | 95 | 86 | 1 |
| 5621 | - | 6144962 | 6143880 | - | *gcvT* | *PA14_68870* | glycine cleavage system aminomethyltransferase T | 134 | 134 | 164 | 145 | 124 | 98 | 1 |
| 5622 | - | 6146732 | 6145113 | - | - | *PA14_68890* | iron ABC transporter, permease | 21 | 23 | 22 | 29 | 32 | 22 | 1 |
| 5623 | - | 6147798 | 6146800 | - | - | *PA14_68900* | iron ABC transporter substrate-binding protein | 35 | 73 | 37 | 65 | 93 | 49 | 0.366758507 |
| 5624 | - | 6148780 | 6147866 | - | - | *PA14_68920* | LysR family transcriptional regulator | 21 | 17 | 20 | 19 | 20 | 21 | 1 |
| 5625 | - | 6148877 | 6150070 | + | - | *PA14_68930* | permease | 31 | 18 | 27 | 26 | 25 | 28 | 0.954437084 |
| 5626 | - | 6150109 | 6150930 | + | - | *PA14_68940* | hypothetical protein | 225 | 44 | 57 | 178 | 144 | 264 | 8.07305E-08 |
| 5627 | - | 6152184 | 6150967 | - | - | *PA14_68955* | 2-octaprenyl-3-methyl-6-methoxy-1,4-benzoquinol hydroxylase | 294 | 179 | 187 | 270 | 250 | 284 | 1 |
| 5628 | - | 6152695 | 6152213 | - | - | *PA14_68970* | hypothetical protein | 124 | 112 | 126 | 139 | 127 | 120 | 1 |

| | A | B | C | D | E | F | G | H | I | J | K | L | M | N |
|---|---|---|---|---|---|---|---|---|---|---|---|---|---|---|
| 5629 | - | 6153884 | 6152700 | - | *ubiH* | *PA14_68980* | 2-octaprenyl-6-methoxyphenyl hydroxylase | 445 | 385 | 444 | 441 | 434 | 402 | 1 |
| 5630 | - | 6155226 | 6153892 | - | *pepP* | *PA14_69000* | aminopeptidase | 505 | 424 | 493 | 529 | 504 | 476 | 1 |
| 5631 | - | 6155790 | 6155236 | - | - | *PA14_69010* | hypothetical protein | 160 | 128 | 176 | 138 | 144 | 136 | 1 |
| 5632 | - | 6155877 | 6156188 | + | - | *PA14_69020* | hypothetical protein | 91 | 73 | 96 | 68 | 106 | 91 | 1 |
| 5633 | - | 6156185 | 6156499 | + | - | *PA14_69030* | hypothetical protein | 202 | 229 | 209 | 178 | 248 | 202 | 1 |
| 5634 | - | 6156721 | 6157332 | + | - | *PA14_69040* | 5-formyltetrahydrofolate cyclo-ligase | 91 | 91 | 110 | 84 | 94 | 88 | 1 |
| 5635 | - | 6157699 | 6158157 | + | - | *PA14_69050* | hypothetical protein | 383 | 380 | 511 | 340 | 370 | 366 | 1 |
| 5636 | - | 6159291 | 6158167 | - | - | *PA14_69060* | ABC transporter permease | 331 | 350 | 446 | 353 | 381 | 373 | 1 |
| 5637 | - | 6162045 | 6159295 | - | - | *PA14_69070* | ABC transporter ATP-binding protein/permease | 368 | 376 | 475 | 270 | 280 | 358 | 1 |
| 5638 | - | 6163115 | 6162042 | - | - | *PA14_69090* | hypothetical protein | 1054 | 871 | 1366 | 547 | 427 | 964 | 1 |
| 5639 | - | 6163741 | 6163322 | - | - | *PA14_69100* | flagellar basal body-associated protein FliL-like protein | 93 | 113 | 121 | 100 | 82 | 102 | 1 |
| 5640 | - | 6163896 | 6164873 | + | - | *PA14_69110* | oxidoreductase | 37 | 40 | 43 | 42 | 38 | 41 | 1 |
| 5641 | - | 6165123 | 6166469 | + | *glpT* | *PA14_69130* | sn-glycerol-3-phosphate transporter | 46 | 48 | 47 | 54 | 52 | 49 | 1 |
| 5642 | - | 6167474 | 6166506 | - | - | *PA14_69140* | CDP-6-deoxy-delta-3,4-glucoseen reductase | 214 | 203 | 198 | 214 | 208 | 180 | 1 |
| 5643 | - | 6168940 | 6167474 | - | - | *PA14_69150* | hypothetical protein | 156 | 160 | 153 | 163 | 159 | 143 | 1 |
| 5644 | - | 6171007 | 6169019 | - | - | *PA14_69170* | O-antigen acetylase | 17 | 18 | 20 | 25 | 17 | 18 | 1 |
| 5645 | - | 6172337 | 6171078 | - | *rho* | *PA14_69190* | transcription termination factor Rho | 112 | 71 | 104 | 69 | 74 | 80 | 1 |
| 5646 | - | 6172908 | 6172582 | - | *trxA* | *PA14_69200* | thioredoxin | 1192 | 1011 | 935 | 650 | 800 | 827 | 1 |
| 5647 | - | 6173092 | 6174612 | + | *ppx* | *PA14_69220* | exopolyphosphatase | 166 | 168 | 185 | 182 | 152 | 163 | 1 |
| 5648 | - | 6176809 | 6174599 | - | *ppk* | *PA14_69230* | polyphosphate kinase | 396 | 456 | 393 | 458 | 433 | 442 | 1 |
| 5649 | - | 6177840 | 6176827 | - | *hemB* | *PA14_69240* | delta-aminolevulinic acid dehydratase | 451 | 474 | 528 | 429 | 473 | 448 | 1 |
| 5650 | - | 6178084 | 6178674 | + | - | *PA14_69250* | hypothetical protein | 30 | 21 | 35 | 31 | 25 | 34 | 1 |
| 5651 | - | 6178839 | 6179507 | + | - | *PA14_69260* | isoprenoid biosynthesis protein with amidotransferase-like domain | 93 | 112 | 82 | 61 | 80 | 75 | 1 |
| 5652 | - | 6179603 | 6180076 | + | - | *PA14_69270* | hypothetical protein | 53 | 56 | 46 | 39 | 47 | 48 | 1 |
| 5653 | - | 6180543 | 6180061 | - | - | *PA14_69280* | hypothetical protein | 203 | 164 | 202 | 164 | 151 | 157 | 1 |
| 5654 | - | 6182511 | 6180610 | - | - | *PA14_69300* | hypothetical protein | 56 | 57 | 66 | 67 | 57 | 53 | 1 |
| 5655 | - | 6183245 | 6182616 | - | - | *PA14_69310* | LysE family efflux protein | 70 | 44 | 81 | 49 | 53 | 55 | 1 |
| 5656 | - | 6184233 | 6183475 | - | - | *PA14_69320* | integral membrane transport protein | 144 | 133 | 147 | 129 | 138 | 138 | 1 |
| 5657 | - | 6184818 | 6184240 | - | - | *PA14_69330* | hypothetical protein | 121 | 92 | 112 | 123 | 108 | 93 | 1 |
| 5658 | - | 6186734 | 6184818 | - | - | *PA14_69340* | ABC transporter ATP-binding protein | 91 | 73 | 85 | 83 | 66 | 69 | 1 |
| 5659 | - | 6186780 | 6187259 | + | - | *PA14_69350* | hypothetical protein | 326 | 306 | 338 | 272 | 327 | 321 | 1 |
| 5660 | - | 6188314 | 6187256 | - | *algP* | *PA14_69370* | alginate regulatory protein AlgP | 1347 | 1562 | 1203 | 1174 | 1347 | 1225 | 1 |
| 5661 | - | 6188440 | 6189069 | + | - | *PA14_69380* | peptidyl-prolyl cis-trans isomerase, FkbP-type | 27 | 24 | 30 | 25 | 24 | 24 | 1 |
| 5662 | - | 6189613 | 6189131 | - | *algQ* | *PA14_69390* | anti-RNA polymerase sigma 70 factor | 350 | 339 | 330 | 360 | 288 | 413 | 1 |
| 5663 | - | 6190384 | 6189893 | - | *dsbH* | *PA14_69400* | disulfide bond formation protein | 38 | 41 | 45 | 47 | 42 | 42 | 1 |
| 5664 | - | 6191802 | 6190564 | - | - | *PA14_69420* | enzyme of heme biosynthesis | 461 | 429 | 436 | 498 | 467 | 478 | 1 |
| 5665 | - | 6192929 | 6191799 | - | - | *PA14_69430* | hypothetical protein | 265 | 234 | 262 | 250 | 252 | 253 | 1 |
| 5666 | - | 6193711 | 6192956 | - | *hemD* | *PA14_69440* | uroporphyrinogen-III synthase | 229 | 174 | 252 | 205 | 202 | 209 | 1 |
| 5667 | - | 6194649 | 6193708 | - | *hemC* | *PA14_69450* | porphobilinogen deaminase | 180 | 155 | 184 | 150 | 162 | 163 | 1 |
| 5668 | - | 6195504 | 6194758 | - | *algR* | *PA14_69470* | alginate biosynthesis regulatory protein AlgR | 609 | 585 | 614 | 638 | 513 | 661 | 1 |
| 5669 | - | 6196585 | 6195509 | - | *algZ* | *PA14_69480* | alginate biosynthesis protein AlgZ/FimS | 201 | 213 | 219 | 234 | 187 | 207 | 1 |
| 5670 | - | 6196810 | 6198204 | + | *argH* | *PA14_69500* | argininosuccinate lyase | 149 | 132 | 159 | 161 | 141 | 149 | 1 |
| 5671 | - | 6199408 | 6198335 | - | - | *PA14_69510* | hypothetical protein | 100 | 87 | 93 | 101 | 119 | 109 | 1 |
| 5672 | - | 6202985 | 6199461 | - | - | *PA14_69520* | hypothetical protein | 51 | 48 | 53 | 48 | 41 | 51 | 1 |
| 5673 | - | 6203863 | 6202982 | - | - | *PA14_69540* | hypothetical protein | 12 | 10 | 11 | 12 | 11 | 10 | 1 |
| 5674 | - | 6205895 | 6203853 | - | - | *PA14_69550* | hypothetical protein | 15 | 10 | 12 | 14 | 10 | 13 | 1 |
| 5675 | - | 6206610 | 6206092 | - | *hcpB* | *PA14_69560* | secreted protein Hcp | 46 | 25 | 21 | 26 | 32 | 30 | 0.490513327 |
| 5676 | - | 6207883 | 6206903 | - | *corA* | *PA14_69570* | magnesium/cobalt transport protein | 38 | 39 | 40 | 38 | 39 | 34 | 1 |
| 5677 | - | 6207985 | 6208260 | + | - | *PA14_69580* | hypothetical protein | 156 | 166 | 157 | 107 | 141 | 119 | 1 |
| 5678 | - | 6208279 | 6209160 | + | - | *PA14_69590* | ABC-type amino acid transporter | 69 | 59 | 65 | 46 | 62 | 55 | 1 |
| 5679 | - | 6209203 | 6209436 | + | - | *PA14_69600* | hypothetical protein | 128 | 208 | 121 | 137 | 168 | 150 | 0.879101916 |
| 5680 | - | 6209585 | 6212437 | + | *cyaA* | *PA14_69610* | adenylate cyclase | 65 | 71 | 64 | 67 | 79 | 71 | 1 |
| 5681 | - | 6212478 | 6213176 | + | - | *PA14_69620* | hypothetical protein | 49 | 50 | 46 | 44 | 54 | 45 | 1 |
| 5682 | - | 6213624 | 6213220 | - | *rnk* | *PA14_69630* | nucleoside diphosphate kinase regulator | 123 | 107 | 130 | 113 | 114 | 116 | 1 |
| 5683 | - | 6214282 | 6213947 | - | *cyaY* | *PA14_69640* | frataxin-like protein | 108 | 112 | 68 | 133 | 222 | 172 | 1 |
| 5684 | - | 6214536 | 6214676 | + | *lppL* | *PA14_69660* | lipopeptide LppL | 38 | 51 | 49 | 43 | 46 | 43 | 1 |
| 5685 | - | 6214687 | 6215934 | + | *lysA* | *PA14_69670* | diaminopimelate decarboxylase | 159 | 149 | 146 | 147 | 140 | 143 | 1 |
| 5686 | - | 6215945 | 6216775 | + | *dapF* | *PA14_69690* | diaminopimelate epimerase | 154 | 159 | 137 | 147 | 139 | 135 | 1 |
| 5687 | - | 6216806 | 6217507 | + | - | *PA14_69700* | hypothetical protein | 199 | 177 | 194 | 193 | 217 | 198 | 1 |
| 5688 | - | 6217527 | 6218438 | + | *xerC* | *PA14_69710* | site-specific tyrosine recombinase XerC | 82 | 69 | 67 | 64 | 75 | 70 | 1 |
| 5689 | - | 6218435 | 6219133 | + | - | *PA14_69720* | hydrolase | 61 | 65 | 54 | 62 | 68 | 56 | 1 |
| 5690 | - | 6220333 | 6219164 | - | - | *PA14_69740* | MFS transporter | 9 | 7 | 8 | 9 | 9 | 7 | 1 |
| 5691 | - | 6220496 | 6221872 | + | - | *PA14_69750* | transcriptional regulator | 67 | 62 | 72 | 70 | 64 | 73 | 1 |
| 5692 | - | 6222811 | 6221897 | - | - | *PA14_69760* | fimbrial protein | 5 | 5 | 9 | 8 | 8 | 8 | 1 |
| 5693 | - | 6223565 | 6223248 | - | - | *PA14_69770* | hypothetical protein | 238 | 273 | 269 | 268 | 168 | 180 | 1 |
| 5694 | - | 6224068 | 6223643 | - | - | *PA14_69780* | hypothetical protein | 70 | 69 | 86 | 73 | 50 | 46 | 1 |
| 5695 | - | 6225657 | 6224329 | - | *amtB* | *PA14_69795* | ammonium transporter | 72 | 71 | 67 | 72 | 75 | 77 | 1 |
| 5696 | - | 6226035 | 6225697 | - | *glnK* | *PA14_69810* | nitrogen regulatory protein P-II 2 | 2683 | 2552 | 2942 | 3152 | 3073 | 3289 | 1 |
| 5697 | - | 6226475 | 6226735 | + | - | *PA14_69820* | hypothetical protein | 47 | 44 | 45 | 34 | 33 | 34 | 1 |
| 5698 | - | 6226776 | 6228269 | + | - | *PA14_69840* | magnesium chelatase | 35 | 33 | 33 | 32 | 35 | 30 | 1 |
| 5699 | - | 6228394 | 6230379 | + | - | *PA14_69850* | choline transporter | 62 | 69 | 58 | 63 | 71 | 62 | 1 |
| 5700 | - | 6231471 | 6230422 | - | *pchP* | *PA14_69870* | phosphorylcholine phosphatase | 34 | 38 | 35 | 36 | 34 | 32 | 1 |
| 5701 | - | 6232539 | 6231622 | - | - | *PA14_69880* | LysR family transcriptional regulator | 9 | 12 | 12 | 11 | 10 | 10 | 1 |
| 5702 | - | 6232688 | 6234154 | + | - | *PA14_69890* | multidrug efflux protein NorA | 24 | 37 | 29 | 36 | 35 | 27 | 1 |
| 5703 | - | 6235742 | 6234066 | - | - | *PA14_69900* | hypothetical protein | 100 | 140 | 110 | 130 | 126 | 112 | 1 |
| 5704 | - | 6235930 | 6237939 | + | *rep* | *PA14_69910* | ATP-dependent DNA helicase Rep | 33 | 28 | 35 | 30 | 27 | 24 | 1 |
| 5705 | - | 6238073 | 6239791 | + | *poxB* | *PA14_69925* | pyruvate dehydrogenase (cytochrome) | 30 | 24 | 27 | 26 | 27 | 27 | 1 |
| 5706 | - | 6239955 | 6240527 | + | *xpt* | *PA14_69940* | xanthine phosphoribosyltransferase | 73 | 81 | 88 | 95 | 77 | 88 | 1 |
| 5707 | - | 6242394 | 6240535 | - | - | *PA14_69950* | hypothetical protein | 52 | 52 | 52 | 64 | 61 | 55 | 1 |
| 5708 | - | 6243014 | 6242604 | - | *cycB* | *PA14_69970* | cytochrome c5 | 1779 | 1412 | 1800 | 1403 | 1375 | 1042 | 1 |
| 5709 | - | 6243784 | 6243236 | - | - | *PA14_69980* | transcriptional regulator | 254 | 261 | 251 | 315 | 236 | 267 | 1 |
| 5710 | - | 6245008 | 6243935 | - | *dadX* | *PA14_69990* | alanine racemase | 159 | 136 | 150 | 130 | 140 | 131 | 1 |
| 5711 | - | 6245452 | 6245099 | - | - | *PA14_70010* | hypothetical protein | 195 | 247 | 189 | 178 | 211 | 169 | 1 |
| 5712 | - | 6246725 | 6245427 | - | *dadA* | *PA14_70040* | D-amino acid dehydrogenase small subunit | 279 | 293 | 268 | 307 | 315 | 238 | 1 |

| | A | B | C | D | E | F | G | H | I | J | K | L | M | N |
|---|---|---|---|---|---|---|---|---|---|---|---|---|---|---|
| 5713 | - | 6247426 | 6247082 | - | - | PA14_70050 | hypothetical protein | 154 | 181 | 124 | 144 | 133 | 161 | 1 |
| 5714 | - | 6247630 | 6247439 | - | - | PA14_70060 | lipoprotein | 143 | 155 | 128 | 132 | 150 | 155 | 1 |
| 5715 | - | 6250215 | 6247651 | - | - | PA14_70070 | hypothetical protein | 119 | 95 | 115 | 123 | 97 | 99 | 1 |
| 5716 | - | 6250369 | 6250857 | + | lrp | PA14_70080 | leucine-responsive regulatory protein | 68 | 74 | 69 | 71 | 80 | 78 | 1 |
| 5717 | - | 6251036 | 6252355 | + | - | PA14_70100 | oxidoreductase | 23 | 22 | 23 | 24 | 18 | 19 | 1 |
| 5718 | - | 6252441 | 6254030 | + | - | PA14_70110 | cardiolipin synthase | 29 | 29 | 31 | 38 | 24 | 28 | 1 |
| 5719 | - | 6255197 | 6254034 | - | - | PA14_70120 | MFS transporter | 24 | 26 | 25 | 31 | 23 | 25 | 1 |
| 5720 | - | 6256837 | 6255344 | - | - | PA14_70140 | aldehyde dehydrogenase | 180 | 186 | 143 | 164 | 152 | 176 | 1 |
| 5721 | - | 6257067 | 6258401 | + | - | PA14_70160 | omega amino acid--pyruvate transaminase | 45 | 37 | 38 | 49 | 34 | 51 | 1 |
| 5722 | - | 6258467 | 6258829 | + | - | PA14_70170 | hypothetical protein | 22 | 26 | 20 | 26 | 22 | 36 | 1 |
| 5723 | - | 6259121 | 6258966 | - | rpmG | PA14_70180 | 50S ribosomal protein L33 | 221 | 385 | 253 | 115 | 162 | 97 | 0.73131877 |
| 5724 | - | 6259369 | 6259133 | - | rpmB | PA14_70190 | 50S ribosomal protein L28 | 310 | 494 | 364 | 203 | 234 | 146 | 1 |
| 5725 | - | 6259725 | 6261305 | + | - | PA14_70200 | dipeptide ABC transporter substrate-binding protein | 68 | 60 | 63 | 67 | 68 | 62 | 1 |
| 5726 | - | 6261360 | 6261551 | + | - | PA14_70220 | hypothetical protein | 54 | 70 | 54 | 67 | 58 | 58 | 1 |
| 5727 | - | 6262242 | 6261568 | - | radC | PA14_70230 | DNA repair protein RadC | 188 | 156 | 201 | 234 | 166 | 186 | 1 |
| 5728 | - | 6262382 | 6263590 | + | coaC | PA14_70240 | bifunctional phosphopantothenoylcysteine decarboxylase/phosphopantothenate synthase | 75 | 50 | 73 | 53 | 57 | 54 | 1 |
| 5729 | - | 6263598 | 6264053 | + | dut | PA14_70260 | deoxyuridine 5'-triphosphate nucleotidohydrolase | 98 | 103 | 101 | 82 | 96 | 84 | 1 |
| 5730 | - | 6264129 | 6266693 | + | algC | PA14_70270 | phosphomannomutase | 242 | 240 | 228 | 225 | 253 | 211 | 1 |
| 5731 | - | 6266710 | 6267615 | + | argB | PA14_70280 | acetylglutamate kinase | 417 | 395 | 399 | 389 | 484 | 393 | 1 |
| 5732 | - | 6268729 | 6267659 | - | - | PA14_70290 | transcriptional regulator | 43 | 46 | 39 | 44 | 40 | 43 | 1 |
| 5733 | - | 6268955 | 6269905 | + | - | PA14_70300 | hypothetical protein | 9 | 8 | 9 | 9 | 8 | 9 | 1 |
| 5734 | - | 6271184 | 6269976 | - | - | PA14_70310 | hypothetical protein | 6 | 5 | 5 | 5 | 4 | 5 | 1 |
| 5735 | - | 6272595 | 6271267 | - | - | PA14_70330 | oxidoreductase | 7 | 6 | 9 | 9 | 8 | 8 | 1 |
| 5736 | - | 6273062 | 6272673 | - | - | PA14_70340 | cytochrome c(mono-heme type) | 1 | 1 | 1 | 1 | 1 | 2 | 1 |
| 5737 | - | 6273236 | 6273700 | + | - | PA14_70350 | hypothetical protein | 66 | 80 | 83 | 86 | 72 | 79 | 1 |
| 5738 | - | 6274297 | 6273680 | - | - | PA14_70360 | hypothetical protein | 345 | 352 | 396 | 492 | 375 | 375 | 1 |
| 5739 | - | 6274959 | 6274318 | - | pyrE | PA14_70370 | orotate phosphoribosyltransferase | 143 | 160 | 170 | 162 | 153 | 132 | 1 |
| 5740 | - | 6275040 | 6275819 | + | crc | PA14_70390 | catabolite repression control protein | 233 | 230 | 273 | 218 | 241 | 236 | 1 |
| 5741 | - | 6276249 | 6275878 | - | - | PA14_70400 | hypothetical protein | 253 | 248 | 269 | 262 | 197 | 211 | 1 |
| 5742 | - | 6277015 | 6276296 | - | rph | PA14_70420 | ribonuclease PH | 111 | 109 | 125 | 109 | 95 | 92 | 1 |
| 5743 | - | 6277196 | 6278059 | + | - | PA14_70430 | hypothetical protein | 80 | 57 | 89 | 59 | 54 | 60 | 1 |
| 5744 | - | 6278118 | 6278729 | + | gmk | PA14_70440 | guanylate kinase | 29 | 24 | 32 | 31 | 28 | 30 | 1 |
| 5745 | - | 6278818 | 6279084 | + | rpoZ | PA14_70450 | DNA-directed RNA polymerase subunit omega | 628 | 616 | 717 | 597 | 456 | 490 | 1 |
| 5746 | - | 6279150 | 6281255 | + | spoT | PA14_70470 | guanosine-3',5'-bis(diphosphate) 3'-pyrophosphohydrolase | 199 | 212 | 234 | 220 | 195 | 194 | 1 |
| 5747 | - | 6281317 | 6281697 | + | - | PA14_70480 | hypothetical protein | 346 | 445 | 279 | 289 | 373 | 302 | 1 |
| 5748 | - | 6281752 | 6282483 | + | - | PA14_70490 | lipoprotein | 151 | 158 | 146 | 140 | 159 | 129 | 1 |
| 5749 | - | 6283110 | 6282490 | - | - | PA14_70510 | hypothetical protein | 24 | 22 | 19 | 22 | 22 | 17 | 1 |
| 5750 | - | 6283977 | 6283177 | - | - | PA14_70530 | AraC family transcriptional regulator | 58 | 59 | 57 | 55 | 51 | 51 | 1 |
| 5751 | - | 6284828 | 6283977 | - | - | PA14_70550 | hypothetical protein | 81 | 87 | 79 | 79 | 79 | 72 | 1 |
| 5752 | - | 6284966 | 6285898 | + | - | PA14_70560 | LysR family transcriptional regulator | 100 | 84 | 97 | 68 | 85 | 87 | 1 |
| 5753 | - | 6285895 | 6287970 | + | recG | PA14_70570 | ATP-dependent DNA helicase RecG | 126 | 135 | 111 | 129 | 135 | 126 | 1 |
| 5754 | - | 6288060 | 6289469 | + | - | PA14_70580 | hypothetical protein | 388 | 416 | 428 | 382 | 397 | 453 | 1 |
| 5755 | - | 6289933 | 6289541 | - | - | PA14_70590 | hypothetical protein | 59 | 78 | 82 | 71 | 76 | 76 | 1 |
| 5756 | - | 6290341 | 6290069 | - | - | PA14_70600 | HU family DNA-binding protein | 249 | 415 | 290 | 295 | 258 | 314 | 1 |
| 5757 | - | 6291697 | 6290543 | - | - | PA14_70620 | rubredoxin reductase | 98 | 92 | 90 | 87 | 94 | 90 | 1 |
| 5758 | - | 6291916 | 6291749 | - | rubA2 | PA14_70630 | rubredoxin 2 | 108 | 106 | 125 | 108 | 126 | 90 | 1 |
| 5759 | - | 6292267 | 6292100 | - | rubA1 | PA14_70640 | rubredoxin 1 | 41 | 59 | 46 | 54 | 48 | 34 | 1 |
| 5760 | - | 6292802 | 6292401 | - | - | PA14_70650 | GlcG protein | 147 | 81 | 114 | 158 | 67 | 74 | 0.970587038 |
| 5761 | - | 6294033 | 6292807 | - | glcF | PA14_70670 | glycolate oxidase iron-sulfur subunit | 165 | 90 | 121 | 182 | 81 | 87 | 1 |
| 5762 | - | 6295122 | 6294043 | - | glcE | PA14_70680 | glycolate oxidase FAD binding subunit | 252 | 119 | 173 | 213 | 118 | 105 | 1 |
| 5763 | - | 6296621 | 6295122 | - | glcD | PA14_70690 | glycolate oxidase subunit GlcD | 339 | 195 | 230 | 319 | 209 | 166 | 1 |
| 5764 | - | 6296826 | 6297581 | + | glcC | PA14_70710 | DNA-binding transcriptional regulator GlcC | 100 | 85 | 80 | 97 | 95 | 99 | 1 |
| 5765 | - | 6297661 | 6298197 | + | ubiC | PA14_70720 | hypothetical protein | 54 | 54 | 64 | 80 | 55 | 54 | 1 |
| 5766 | - | 6298224 | 6299114 | + | ubiA | PA14_70730 | 4-hydroxybenzoate octaprenyltransferase | 124 | 138 | 159 | 218 | 147 | 190 | 1 |
| 5767 | - | 6299593 | 6299138 | - | - | PA14_70740 | hypothetical protein | 162 | 200 | 222 | 260 | 184 | 241 | 1 |
| 5768 | - | 6299698 | 6300387 | + | phoB | PA14_70750 | two-component response regulator PhoB | 46 | 49 | 50 | 43 | 51 | 53 | 1 |
| 5769 | - | 6300460 | 6301791 | + | phoR | PA14_70760 | two-component sensor PhoR | 24 | 22 | 24 | 27 | 24 | 26 | 1 |
| 5770 | - | 6301894 | 6303234 | + | - | PA14_70770 | hypothetical protein | 71 | 65 | 70 | 64 | 59 | 58 | 1 |
| 5771 | - | 6304168 | 6303269 | - | - | PA14_70780 | hypothetical protein | 75 | 70 | 73 | 77 | 73 | 63 | 1 |
| 5772 | - | 6305187 | 6304285 | - | - | PA14_70790 | two-component response regulator | 162 | 118 | 161 | 113 | 139 | 158 | 1 |
| 5773 | - | 6306034 | 6305306 | - | phoU | PA14_70800 | phosphate uptake regulatory protein PhoU | 104 | 108 | 119 | 72 | 81 | 73 | 1 |
| 5774 | - | 6306963 | 6306130 | - | pstB | PA14_70810 | phosphate transporter ATP-binding protein | 46 | 58 | 62 | 40 | 56 | 46 | 1 |
| 5775 | - | 6308655 | 6306979 | - | pstA | PA14_70830 | phosphate ABC transporter permease | 39 | 35 | 50 | 27 | 30 | 24 | 1 |
| 5776 | - | 6310708 | 6308675 | - | pstC | PA14_70850 | membrane protein component of ABC phosphate transporter | 15 | 19 | 20 | 12 | 14 | 12 | 1 |
| 5777 | - | 6312102 | 6311131 | - | - | PA14_70860 | hypothetical protein | 28 | 42 | 31 | 19 | 29 | 18 | 1 |
| 5778 | - | 6312629 | 6312498 | - | - | PA14_70870 | 5S ribosomal RNA | 10 | 10 | 26 | 10 | 5 | 2 | 1 |
| 5779 | - | 6315656 | 6312766 | - | - | PA14_70880 | 23S ribosomal RNA | 612 | 872 | 726 | 418 | 753 | 121 | 1 |
| 5780 | - | 6315958 | 6315886 | - | - | PA14_70890 | Ala tRNA | 696 | 956 | 932 | 643 | 489 | 240 | 1 |
| 5781 | - | 6316063 | 6315990 | - | - | PA14_70900 | Ile tRNA | 710 | 1071 | 928 | 568 | 435 | 216 | 1 |
| 5782 | - | 6317654 | 6316129 | - | - | PA14_70910 | 16S ribosomal RNA | 237 | 360 | 900 | 310 | 553 | 68 | 1 |
| 5783 | - | 6319545 | 6318229 | - | - | PA14_70920 | major facilitator transporter | 22 | 20 | 19 | 18 | 18 | 17 | 1 |
| 5784 | - | 6319833 | 6320237 | + | - | PA14_70930 | hypothetical protein | 121 | 149 | 111 | 113 | 100 | 98 | 1 |
| 5785 | - | 6321968 | 6320283 | - | betA | PA14_70940 | choline dehydrogenase | 143 | 98 | 82 | 68 | 93 | 102 | 1 |
| 5786 | - | 6323576 | 6322104 | - | betB | PA14_70950 | betaine aldehyde dehydrogenase | 339 | 264 | 224 | 229 | 346 | 351 | 1 |
| 5787 | - | 6324230 | 6323637 | - | betI | PA14_70970 | transcriptional regulator BetI | 322 | 181 | 268 | 219 | 284 | 347 | 1 |
| 5788 | - | 6324563 | 6326113 | + | betT1 | PA14_70980 | choline transporter BetT | 13 | 6 | 5 | 9 | 6 | 12 | 0.05276977 |
| 5789 | - | 6327508 | 6326330 | - | - | PA14_71000 | lycine betaine/L-proline ABC transporter, ATP-binding subunit | 75 | 82 | 68 | 76 | 75 | 91 | 1 |
| 5790 | - | 6328351 | 6327512 | - | - | PA14_71020 | BC-type proline/glycine betaine transport system, permease component | 53 | 56 | 48 | 50 | 54 | 53 | 1 |
| 5791 | - | 6329331 | 6328393 | - | - | PA14_71030 | hypothetical protein | 86 | 78 | 82 | 74 | 74 | 87 | 1 |
| 5792 | - | 6329795 | 6331171 | + | sdaB | PA14_71060 | L-serine dehydratase | 6 | 6 | 8 | 7 | 6 | 8 | 1 |
| 5793 | - | 6331592 | 6332695 | + | - | PA14_71070 | AraC family transcriptional regulator | 69 | 75 | 66 | 71 | 66 | 78 | 1 |
| 5794 | - | 6333118 | 6332741 | - | - | PA14_71080 | hypothetical protein | 94 | 99 | 86 | 86 | 76 | 112 | 1 |
| 5795 | - | 6334151 | 6333258 | - | - | PA14_71090 | LysR family transcriptional regulator | 21 | 24 | 22 | 25 | 17 | 21 | 1 |
| 5796 | - | 6334259 | 6335326 | + | - | PA14_71100 | hypothetical protein | 9 | 7 | 5 | 15 | 17 | 14 | 1 |

| | A | B | C | D | E | F | G | H | I | J | K | L | M | N |
|---|---|---|---|---|---|---|---|---|---|---|---|---|---|---|
| 5797 | - | 6336265 | 6335270 | - | - | *PA14_71110* | lipolytic protein | 2 | 1 | 1 | 3 | 3 | 2 | 0.831807123 |
| 5798 | - | 6336750 | 6336271 | - | - | *PA14_71120* | hypothetical protein | 4 | 4 | 2 | 3 | 3 | 1 | 1 |
| 5799 | - | 6337766 | 6336801 | - | - | *PA14_71140* | 3-hydroxybutyryl-CoA dehydrogenase | 1 | 2 | 1 | 2 | 1 | 2 | 0.503413992 |
| 5800 | - | 6338701 | 6337817 | - | - | *PA14_71150* | hypothetical protein | 1 | 1 | 1 | 1 | 1 | 0 | 1 |
| 5801 | - | 6339712 | 6338774 | - | - | *PA14_71160* | hypothetical protein | 5 | 4 | 4 | 5 | 5 | 5 | 1 |
| 5802 | - | 6339923 | 6340933 | + | - | *PA14_71170* | AraC family transcriptional regulator | 29 | 28 | 23 | 37 | 31 | 29 | 1 |
| 5803 | - | 6342017 | 6340863 | - | - | *PA14_71180* | acetylornithine deacetylase | 24 | 23 | 20 | 24 | 20 | 23 | 1 |
| 5804 | - | 6342748 | 6342074 | - | - | *PA14_71190* | hypothetical protein | 7 | 5 | 5 | 6 | 6 | 5 | 1 |
| 5805 | - | 6343184 | 6342762 | - | - | *PA14_71200* | YjgF family translation initiation inhibitor | 7 | 5 | 5 | 5 | 5 | 6 | 1 |
| 5806 | - | 6344536 | 6343184 | - | - | *PA14_71210* | hypothetical protein | 6 | 6 | 6 | 7 | 6 | 7 | 1 |
| 5807 | - | 6346302 | 6344830 | - | *cls* | *PA14_71220* | cardiolipin synthetase | 49 | 49 | 54 | 62 | 47 | 48 | 1 |
| 5808 | - | 6346455 | 6346925 | + | - | *PA14_71230* | hypothetical protein | 18 | 18 | 14 | 21 | 21 | 19 | 1 |
| 5809 | - | 6347147 | 6348124 | + | - | *PA14_71240* | hypothetical protein | 28 | 28 | 24 | 28 | 26 | 30 | 1 |
| 5810 | - | 6348251 | 6348781 | + | - | *PA14_71250* | hypothetical protein | 11 | 10 | 11 | 10 | 6 | 13 | 1 |
| 5811 | - | 6348797 | 6350857 | + | - | *PA14_71260* | FMN oxidoreductase | 16 | 18 | 17 | 17 | 17 | 19 | 1 |
| 5812 | - | 6350959 | 6352920 | + | - | *PA14_71280* | ferredoxin | 11 | 10 | 10 | 12 | 11 | 13 | 1 |
| 5813 | - | 6353285 | 6354271 | + | - | *PA14_71300* | electron transfer flavoprotein alpha subunit | 9 | 7 | 8 | 8 | 9 | 10 | 1 |
| 5814 | - | 6354307 | 6355080 | + | - | *PA14_71310* | hypothetical protein | 5 | 4 | 4 | 4 | 4 | 6 | 1 |
| 5815 | - | 6355205 | 6355774 | + | - | *PA14_71320* | hypothetical protein | 24 | 23 | 18 | 24 | 20 | 24 | 1 |
| 5816 | - | 6355782 | 6355988 | + | - | *PA14_71330* | transcriptional regulator | 24 | 14 | 20 | 20 | 25 | 20 | 0.625658913 |
| 5817 | - | 6356020 | 6356424 | + | - | *PA14_71340* | hypothetical protein | 14 | 11 | 12 | 12 | 10 | 11 | 1 |
| 5818 | - | 6356438 | 6356671 | + | - | *PA14_71350* | hypothetical protein | 22 | 18 | 20 | 21 | 20 | 18 | 1 |
| 5819 | - | 6356668 | 6357000 | + | - | *PA14_71360* | hypothetical protein | 27 | 24 | 29 | 21 | 21 | 23 | 1 |
| 5820 | - | 6356997 | 6357287 | + | - | *PA14_71370* | hypothetical protein | 128 | 100 | 122 | 113 | 118 | 127 | 1 |
| 5821 | - | 6357460 | 6357284 | - | - | *PA14_71380* | hypothetical protein | 463 | 394 | 484 | 485 | 459 | 533 | 1 |
| 5822 | - | 6358025 | 6357465 | - | - | *PA14_71390* | hypothetical protein | 302 | 244 | 272 | 245 | 270 | 289 | 1 |
| 5823 | - | 6358262 | 6358152 | - | - | *PA14_71400* | hypothetical protein | 67 | 53 | 72 | 46 | 39 | 56 | 1 |
| 5824 | - | 6359599 | 6358310 | - | - | *PA14_71410* | ring hydroxylating dioxygenase, alpha-subunit | 14 | 17 | 15 | 13 | 14 | 15 | 1 |
| 5825 | - | 6360027 | 6361127 | + | - | *PA14_71420* | ferredoxin | 12 | 13 | 13 | 16 | 13 | 14 | 1 |
| 5826 | - | 6363562 | 6361121 | - | - | *PA14_71430* | hypothetical protein | 53 | 50 | 51 | 57 | 48 | 47 | 1 |
| 5827 | - | 6366007 | 6364967 | - | *ltaA* | *PA14_71440* | low specificity l-threonine aldolase | 57 | 84 | 50 | 42 | 61 | 47 | 1 |
| 5828 | - | 6366732 | 6366091 | - | - | *PA14_71450* | hypothetical protein | 31 | 38 | 35 | 33 | 36 | 26 | 1 |
| 5829 | - | 6366948 | 6368201 | + | *glyA1* | *PA14_71460* | serine hydroxymethyltransferase | 34 | 49 | 40 | 56 | 41 | 53 | 1 |
| 5830 | - | 6368287 | 6369537 | + | *soxB* | *PA14_71470* | sarcosine oxidase beta subunit | 10 | 10 | 9 | 11 | 8 | 13 | 1 |
| 5831 | - | 6369631 | 6369951 | + | *soxD* | *PA14_71490* | sarcosine oxidase delta subunit | 6 | 5 | 3 | 6 | 3 | 5 | 1 |
| 5832 | - | 6369948 | 6372965 | + | *soxA* | *PA14_71500* | sarcosine oxidase alpha subunit | 19 | 17 | 17 | 19 | 15 | 23 | 1 |
| 5833 | - | 6373059 | 6373694 | + | *soxG* | *PA14_71510* | sarcosine oxidase gamma subunit | 15 | 19 | 17 | 17 | 16 | 23 | 1 |
| 5834 | - | 6373744 | 6374601 | + | *purU2* | *PA14_71530* | formyltetrahydrofolate deformylase | 13 | 13 | 15 | 15 | 16 | 23 | 1 |
| 5835 | - | 6374808 | 6376007 | + | *fdhA* | *PA14_71560* | glutathione-independent formaldehyde dehydrogenase | 17 | 16 | 16 | 17 | 15 | 23 | 1 |
| 5836 | - | 6376091 | 6377047 | + | - | *PA14_71570* | hypothetical protein | 62 | 59 | 67 | 53 | 68 | 61 | 1 |
| 5837 | - | 6377660 | 6377112 | - | - | *PA14_71580* | hypothetical protein | 76 | 73 | 85 | 65 | 82 | 79 | 1 |
| 5838 | - | 6377967 | 6377722 | - | - | *PA14_71590* | hypothetical protein | 873 | 1081 | 1135 | 1127 | 1002 | 1134 | 1 |
| 5839 | - | 6379164 | 6378082 | - | *purK* | *PA14_71600* | phosphoribosylaminoimidazole carboxylase ATPase subunit | 64 | 76 | 64 | 54 | 74 | 45 | 1 |
| 5840 | - | 6379686 | 6379195 | - | *purE* | *PA14_71620* | phosphoribosylaminoimidazole carboxylase catalytic subunit | 59 | 64 | 76 | 49 | 54 | 44 | 1 |
| 5841 | - | 6380034 | 6381062 | + | *adhA* | *PA14_71630* | alcohol dehydrogenase | 514 | 581 | 692 | 281 | 358 | 450 | 1 |
| 5842 | - | 6382007 | 6381099 | - | - | *PA14_71640* | LysR family transcriptional regulator | 90 | 102 | 116 | 96 | 83 | 85 | 1 |
| 5843 | - | 6382183 | 6383607 | + | *aspA* | *PA14_71650* | aspartate ammonia-lyase | 140 | 226 | 122 | 213 | 189 | 115 | 1 |
| 5844 | - | 6385185 | 6383971 | - | - | *PA14_71670* | hypothetical protein | 47 | 47 | 45 | 46 | 47 | 35 | 1 |
| 5845 | - | 6386697 | 6385222 | - | - | *PA14_71680* | GntR family transcriptional regulator | 11 | 11 | 12 | 13 | 12 | 9 | 1 |
| 5846 | - | 6386825 | 6387274 | + | - | *PA14_71690* | GNAT family acetyltransferase | 13 | 17 | 11 | 13 | 16 | 11 | 1 |
| 5847 | - | 6387283 | 6387957 | + | - | *PA14_71700* | GNAT family acetyltransferase | 24 | 37 | 35 | 28 | 31 | 22 | 1 |
| 5848 | - | 6389235 | 6387982 | - | *mtr* | *PA14_71710* | tryptophan permease | 23 | 28 | 26 | 23 | 23 | 21 | 1 |
| 5849 | - | 6391239 | 6389416 | - | - | *PA14_71720* | pyruvate carboxylase subunit B | 97 | 246 | 155 | 130 | 198 | 223 | 0.703239983 |
| 5850 | - | 6392670 | 6391255 | - | - | *PA14_71740* | pyruvate carboxylase subunit A | 83 | 205 | 134 | 98 | 173 | 192 | 0.605197743 |
| 5851 | - | 6392884 | 6393819 | + | - | *PA14_71750* | LysR family transcriptional regulator | 9 | 13 | 13 | 10 | 13 | 13 | 1 |
| 5852 | - | 6393988 | 6393773 | - | - | *PA14_71760* | hypothetical protein | 27 | 26 | 34 | 18 | 32 | 33 | 1 |
| 5853 | - | 6394976 | 6394095 | - | - | *PA14_71780* | RpiR family transcriptional regulator | 172 | 117 | 192 | 116 | 142 | 147 | 1 |
| 5854 | - | 6395062 | 6396528 | + | - | *PA14_71800* | glucose-6-phosphate 1-dehydrogenase | 96 | 94 | 99 | 93 | 106 | 118 | 1 |
| 5855 | - | 6396609 | 6398003 | + | - | *PA14_71820* | peptidase | 38 | 41 | 52 | 32 | 41 | 26 | 1 |
| 5856 | - | 6398788 | 6397997 | - | - | *PA14_71830* | hypothetical protein | 89 | 76 | 85 | 67 | 83 | 64 | 1 |
| 5857 | - | 6400920 | 6398719 | - | - | *PA14_71840* | hypothetical protein | 69 | 69 | 63 | 60 | 70 | 55 | 1 |
| 5858 | - | 6403831 | 6400979 | - | - | *PA14_71850* | hypothetical protein | 12 | 17 | 13 | 13 | 13 | 11 | 1 |
| 5859 | - | 6404008 | 6406194 | + | *uvrD* | *PA14_71870* | DNA-dependent helicase II | 94 | 98 | 96 | 108 | 98 | 91 | 1 |
| 5860 | - | 6406232 | 6406660 | + | - | *PA14_71880* | hypothetical protein | 39 | 47 | 56 | 52 | 46 | 48 | 1 |
| 5861 | - | 6406758 | 6408251 | + | - | *PA14_71890* | coenzyme A transferase | 78 | 71 | 77 | 65 | 54 | 54 | 1 |
| 5862 | - | 6408629 | 6408832 | + | - | *PA14_71900* | hypothetical protein | 1146 | 1323 | 1227 | 740 | 980 | 1143 | 1 |
| 5863 | - | 6408839 | 6408863 | ? | - | predicted RNA | - | 836 | 935 | 867 | 573 | 713 | 846 | 1 |
| 5864 | - | 6410033 | 6408888 | - | *wbpZ* | *PA14_71910* | glycosyltransferase WbpZ | 218 | 241 | 208 | 281 | 268 | 279 | 1 |
| 5865 | - | 6411161 | 6410034 | - | *wbpY* | *PA14_71920* | glycosyltransferase WbpY | 101 | 116 | 94 | 113 | 120 | 113 | 1 |
| 5866 | - | 6412527 | 6411145 | - | *wbpX* | *PA14_71930* | glycosyltransferase WbpX | 133 | 151 | 122 | 151 | 141 | 152 | 1 |
| 5867 | - | 6413789 | 6412524 | - | *wzt* | *PA14_71940* | ABC subunit of A-band LPS efflux transporter | 162 | 174 | 145 | 194 | 189 | 176 | 1 |
| 5868 | - | 6414586 | 6413789 | - | *wzm* | *PA14_71960* | membrane subunit of A-band LPS efflux transporter | 108 | 111 | 98 | 134 | 146 | 125 | 1 |
| 5869 | - | 6416025 | 6414586 | - | *wbpW* | *PA14_71970* | GDP-mannose pyrophosphorylase | 168 | 199 | 169 | 200 | 177 | 173 | 1 |
| 5870 | - | 6417000 | 6416029 | - | *gmd* | *PA14_71990* | GDP-mannose 4,6-dehydratase | 250 | 305 | 247 | 296 | 266 | 253 | 1 |
| 5871 | - | 6417911 | 6416997 | - | *rmd* | *PA14_72000* | oxidoreductase Rmd | 113 | 118 | 121 | 126 | 127 | 116 | 1 |
| 5872 | - | 6418319 | 6419947 | + | - | *PA14_72010* | glycosyltransferase | 92 | 78 | 84 | 85 | 91 | 95 | 1 |
| 5873 | - | 6419941 | 6421242 | + | - | *PA14_72020* | hypothetical protein | 80 | 105 | 80 | 94 | 101 | 96 | 1 |
| 5874 | - | 6421239 | 6422102 | + | - | *PA14_72030* | hypothetical protein | 54 | 57 | 49 | 50 | 56 | 51 | 1 |
| 5875 | - | 6422099 | 6423241 | + | - | *PA14_72040* | acyltransferase | 33 | 36 | 36 | 36 | 38 | 38 | 1 |
| 5876 | - | 6423235 | 6424071 | + | - | *PA14_72050* | hypothetical protein | 51 | 67 | 45 | 44 | 55 | 48 | 1 |
| 5877 | - | 6424234 | 6424446 | + | - | *PA14_72060* | hypothetical protein | 23 | 21 | 23 | 15 | 24 | 34 | 1 |
| 5878 | - | 6424542 | 6424859 | + | - | *PA14_72070* | hypothetical protein | 124 | 221 | 114 | 106 | 141 | 109 | 0.723542393 |
| 5879 | - | 6424984 | 6425277 | + | - | *PA14_72080* | hypothetical protein | 37 | 28 | 46 | 42 | 32 | 31 | 1 |
| 5880 | - | 6425306 | 6425638 | + | - | *PA14_72090* | hypothetical protein | 51 | 44 | 46 | 36 | 37 | 33 | 1 |

| | A | B | C | D | E | F | G | H | I | J | K | L | M | N |
|---|---|---|---|---|---|---|---|---|---|---|---|---|---|---|
| 5881 | - | 6425635 | 6427593 | + | - | *PA14_72110* | hypothetical protein | 50 | 51 | 55 | 52 | 47 | 46 | 1 |
| 5882 | - | 6428120 | 6427701 | - | - | *PA14_72130* | hypothetical protein | 107 | 119 | 129 | 124 | 119 | 114 | 1 |
| 5883 | - | 6428188 | 6429132 | + | - | *PA14_72140* | hypothetical protein | 16 | 15 | 17 | 15 | 16 | 14 | 1 |
| 5884 | - | 6429129 | 6429491 | + | - | *PA14_72150* | hypothetical protein | 26 | 26 | 26 | 30 | 27 | 23 | 1 |
| 5885 | - | 6429771 | 6431075 | + | - | *PA14_72170* | citrate transporter | 21 | 18 | 22 | 22 | 19 | 21 | 1 |
| 5886 | - | 6431096 | 6431860 | + | - | *PA14_72180* | membrane protein TerC | 8 | 6 | 9 | 5 | 8 | 7 | 1 |
| 5887 | - | 6432480 | 6431866 | - | *prfH* | *PA14_72200* | peptide chain release factor-like protein | 33 | 21 | 30 | 21 | 26 | 28 | 1 |
| 5888 | - | 6433616 | 6432477 | - | - | *PA14_72210* | hypothetical protein | 39 | 25 | 34 | 28 | 28 | 30 | 1 |
| 5889 | - | 6434784 | 6433984 | - | - | *PA14_72220* | hypothetical protein | 34 | 25 | 36 | 45 | 34 | 30 | 1 |
| 5890 | - | 6435138 | 6436886 | + | - | *PA14_72230* | hypothetical protein | 114 | 101 | 117 | 98 | 129 | 129 | 1 |
| 5891 | - | 6436955 | 6438328 | + | - | *PA14_72250* | metalloprotease | 249 | 245 | 328 | 218 | 174 | 220 | 1 |
| 5892 | - | 6438894 | 6438334 | - | - | *PA14_72260* | hypothetical protein | 1311 | 1316 | 1762 | 643 | 559 | 1322 | 1 |
| 5893 | - | 6440326 | 6439037 | - | *citA* | *PA14_72280* | citrate transporter | 23 | 21 | 21 | 29 | 20 | 24 | 1 |
| 5894 | - | 6440701 | 6441741 | + | - | *PA14_72300* | hypothetical protein | 25 | 25 | 31 | 33 | 26 | 27 | 1 |
| 5895 | - | 6442989 | 6441760 | - | - | *PA14_72320* | hypothetical protein | 54 | 42 | 55 | 60 | 51 | 43 | 1 |
| 5896 | - | 6444428 | 6443094 | - | *gltP* | *PA14_72340* | glutamate/aspartate:proton symporter | 38 | 33 | 46 | 35 | 29 | 33 | 1 |
| 5897 | - | 6444651 | 6444845 | + | - | *PA14_72350* | hypothetical protein | 89 | 48 | 73 | 56 | 48 | 47 | 0.391278558 |
| 5898 | - | 6444984 | 6444994 | ? | - | predicted RNA | - | 364 | 269 | 492 | 786 | 610 | 442 | 1 |
| 5899 | - | 6445608 | 6445141 | - | - | *PA14_72360* | hypothetical protein | 741 | 370 | 498 | 388 | 386 | 347 | 1 |
| 5900 | - | 6445791 | 6445630 | - | - | *PA14_72370* | hypothetical protein | 1035 | 578 | 649 | 507 | 532 | 575 | 1 |
| 5901 | - | 6446259 | 6447608 | + | *algB* | *PA14_72380* | two-component response regulator AlgB | 47 | 39 | 39 | 40 | 42 | 36 | 1 |
| 5902 | - | 6447605 | 6449392 | + | - | *PA14_72390* | two-component sensor | 53 | 57 | 55 | 62 | 68 | 52 | 1 |
| 5903 | - | 6450206 | 6449427 | - | - | *PA14_72400* | N-acetylmuramoyl-L-alanine amidase family protein | 70 | 70 | 77 | 75 | 70 | 75 | 1 |
| 5904 | - | 6450829 | 6450236 | - | - | *PA14_72410* | hypothetical protein | 34 | 29 | 33 | 30 | 30 | 33 | 1 |
| 5905 | - | 6452962 | 6450947 | - | - | *PA14_72420* | diguanylate cyclase | 264 | 266 | 266 | 337 | 279 | 275 | 1 |
| 5906 | - | 6453837 | 6452959 | - | - | *PA14_72430* | hypothetical protein | 219 | 238 | 230 | 207 | 213 | 206 | 1 |
| 5907 | - | 6454488 | 6453853 | - | *dsbA* | *PA14_72450* | thiol:disulfide interchange protein DsbA | 303 | 287 | 306 | 206 | 246 | 218 | 1 |
| 5908 | - | 6455261 | 6454656 | - | *cc4* | *PA14_72460* | cytochrome c4 | 282 | 318 | 311 | 262 | 276 | 237 | 1 |
| 5909 | - | 6455600 | 6455307 | - | - | *PA14_72470* | cytochrome | 196 | 139 | 210 | 172 | 107 | 120 | 1 |
| 5910 | - | 6455784 | 6456431 | + | *engB* | *PA14_72480* | ribosome biogenesis GTP-binding protein YsxC | 184 | 183 | 234 | 155 | 168 | 165 | 1 |
| 5911 | - | 6459436 | 6456695 | - | *polA* | *PA14_72490* | DNA polymerase I | 84 | 86 | 88 | 86 | 77 | 72 | 1 |
| 5912 | - | 6459514 | 6459804 | + | - | *PA14_72500* | hypothetical protein | 1183 | 1196 | 1290 | 901 | 999 | 1269 | 1 |
| 5913 | - | 6459838 | 6460788 | + | *thrB* | *PA14_72510* | homoserine kinase | 278 | 309 | 313 | 231 | 327 | 350 | 1 |
| 5914 | - | 6460908 | 6461375 | ? | - | predicted RNA | - | 1127 | 1102 | 1450 | 1565 | 1406 | 1990 | 1 |
| 5915 | - | 6462093 | 6461404 | - | - | *PA14_72520* | hypothetical protein | 219 | 238 | 202 | 259 | 260 | 309 | 1 |
| 5916 | - | 6464314 | 6462110 | - | - | *PA14_72540* | ribonucleotide reductase | 160 | 153 | 135 | 188 | 185 | 211 | 1 |
| 5917 | - | 6465347 | 6464424 | - | - | *PA14_72550* | adhesin | 82 | 49 | 67 | 56 | 62 | 71 | 1 |
| 5918 | - | 6465417 | 6465920 | + | *np20* | *PA14_72560* | transcriptional regulator np20 | 279 | 180 | 249 | 264 | 324 | 391 | 1 |
| 5919 | - | 6465920 | 6466729 | + | *znuC* | *PA14_72580* | zinc transporter | 168 | 110 | 143 | 138 | 156 | 175 | 1 |
| 5920 | - | 6466722 | 6467510 | + | *znuB* | *PA14_72590* | ABC zinc transporter permease ZnuB | 72 | 51 | 71 | 64 | 70 | 73 | 1 |
| 5921 | - | 6467552 | 6468340 | + | - | *PA14_72600* | lipoprotein | 49 | 43 | 50 | 42 | 40 | 42 | 1 |
| 5922 | - | 6468547 | 6469554 | + | *metN* | *PA14_72620* | DL-methionine transporter ATP-binding subunit | 33 | 29 | 38 | 28 | 25 | 31 | 1 |
| 5923 | - | 6469554 | 6470231 | + | - | *PA14_72630* | ABC transporter permease | 50 | 65 | 65 | 44 | 59 | 47 | 1 |
| 5924 | - | 6470308 | 6471090 | + | - | *PA14_72640* | TonB-dependent receptor | 79 | 121 | 68 | 51 | 102 | 57 | 1 |
| 5925 | - | 6471293 | 6472150 | + | - | *PA14_72650* | hypothetical protein | 35 | 5 | 5 | 4 | 3 | 4 | 2.57052E-52 |
| 5926 | - | 6472155 | 6472808 | + | - | *PA14_72660* | hypothetical protein | 109 | 11 | 9 | 12 | 10 | 11 | 3.74949E-74 |
| 5927 | - | 6472805 | 6474136 | + | - | *PA14_72690* | glutamine synthetase | 87 | 19 | 19 | 23 | 16 | 17 | 1.74921E-06 |
| 5928 | - | 6474204 | 6474872 | + | - | *PA14_72700* | hypothetical protein | 88 | 38 | 46 | 49 | 35 | 38 | 0.252416432 |
| 5929 | - | 6474974 | 6476323 | + | - | *PA14_72710* | transporter | 63 | 16 | 14 | 15 | 16 | 15 | 1.97337E-05 |
| 5930 | - | 6477685 | 6476342 | - | - | *PA14_72720* | two-component response regulator | 88 | 72 | 86 | 67 | 71 | 63 | 1 |
| 5931 | - | 6479448 | 6477682 | - | - | *PA14_72740* | two-component sensor | 32 | 26 | 31 | 27 | 28 | 25 | 1 |
| 5932 | - | 6479530 | 6480408 | + | - | *PA14_72750* | hypothetical protein | 23 | 15 | 21 | 20 | 21 | 18 | 1 |
| 5933 | - | 6480480 | 6481268 | + | - | *PA14_72760* | beta-lactamase | 46 | 43 | 51 | 51 | 48 | 49 | 1 |
| 5934 | - | 6481781 | 6481281 | - | - | *PA14_72770* | hypothetical protein | 89 | 78 | 91 | 85 | 78 | 86 | 1 |
| 5935 | - | 6481814 | 6482680 | + | *pdxY* | *PA14_72780* | pyridoxamine kinase | 52 | 45 | 57 | 52 | 48 | 51 | 1 |
| 5936 | - | 6483302 | 6482718 | - | - | *PA14_72790* | hypothetical protein | 36 | 38 | 42 | 42 | 39 | 36 | 1 |
| 5937 | - | 6485007 | 6483304 | - | - | *PA14_72800* | potassium efflux transporter | 41 | 36 | 44 | 41 | 41 | 40 | 1 |
| 5938 | - | 6485952 | 6485386 | - | - | *PA14_72810* | hypothetical protein | 96 | 87 | 116 | 101 | 101 | 114 | 1 |
| 5939 | - | 6486029 | 6486730 | + | - | *PA14_72820* | hypothetical protein | 80 | 69 | 92 | 112 | 79 | 85 | 1 |
| 5940 | - | 6486543 | 6486767 | + | - | *PA14_72830* | hypothetical protein | 168 | 144 | 189 | 272 | 180 | 185 | 1 |
| 5941 | - | 6487573 | 6486815 | - | - | *PA14_72840* | short-chain dehydrogenase | 146 | 140 | 138 | 160 | 135 | 152 | 1 |
| 5942 | - | 6489035 | 6487668 | - | - | *PA14_72850* | glutamine synthetase | 43 | 39 | 34 | 49 | 31 | 38 | 1 |
| 5943 | - | 6490389 | 6489037 | - | - | *PA14_72870* | aminotransferase | 39 | 39 | 37 | 53 | 34 | 37 | 1 |
| 5944 | - | 6490547 | 6491329 | + | - | *PA14_72880* | short-chain dehydrogenase | 24 | 27 | 25 | 26 | 25 | 23 | 1 |
| 5945 | - | 6491340 | 6492074 | + | - | *PA14_72890* | transcriptional regulator | 87 | 94 | 93 | 78 | 66 | 73 | 1 |
| 5946 | - | 6492303 | 6492091 | - | - | *PA14_72900* | lipoprotein | 216 | 214 | 229 | 223 | 193 | 216 | 1 |
| 5947 | - | 6492922 | 6492521 | - | - | *PA14_72920* | hypothetical protein | 212 | 228 | 255 | 227 | 206 | 230 | 1 |
| 5948 | - | 6493975 | 6493118 | - | - | *PA14_72930* | hypothetical protein | 234 | 185 | 176 | 105 | 128 | 118 | 1 |
| 5949 | - | 6494339 | 6496096 | + | - | *PA14_72940* | sodium/proton antiporter | 43 | 42 | 39 | 37 | 37 | 38 | 1 |
| 5950 | - | 6496412 | 6497719 | + | - | *PA14_72960* | MFS dicarboxylate transporter | 13 | 14 | 18 | 15 | 15 | 9 | 1 |
| 5951 | - | 6499163 | 6498135 | - | *tonB* | *PA14_72970* | TonB protein | 55 | 221 | 65 | 475 | 464 | 149 | 4.74072E-05 |
| 5952 | - | 6500224 | 6499256 | - | - | *PA14_72980* | G3E family GTPase | 39 | 26 | 34 | 31 | 31 | 31 | 1 |
| 5953 | - | 6500327 | 6500695 | + | - | *PA14_72990* | hypothetical protein | 44 | 43 | 53 | 48 | 41 | 50 | 1 |
| 5954 | - | 6501489 | 6500848 | - | - | *PA14_73000* | hypothetical protein | 15 | 14 | 13 | 16 | 15 | 17 | 1 |
| 5955 | - | 6502688 | 6501486 | - | - | *PA14_73010* | hypothetical protein | 14 | 13 | 14 | 15 | 18 | 18 | 1 |
| 5956 | - | 6503107 | 6502703 | - | - | *PA14_73020* | DksA/TraR family C4-type zinc finger protein | 7 | 6 | 7 | 7 | 9 | 13 | 1 |
| 5957 | - | 6503220 | 6503636 | + | - | *PA14_73030* | hypothetical protein | 49 | 56 | 54 | 53 | 52 | 54 | 1 |
| 5958 | - | 6504807 | 6503614 | - | *amiA* | *PA14_73040* | N-acetylmuramoyl-L-alanine amidase | 15 | 19 | 19 | 19 | 18 | 17 | 1 |
| 5959 | - | 6504907 | 6505803 | + | - | *PA14_73050* | GTP cyclohydrolase | 11 | 11 | 14 | 15 | 11 | 14 | 1 |
| 5960 | - | 6505800 | 6506360 | + | - | *PA14_73060* | hypothetical protein | 14 | 14 | 15 | 15 | 13 | 15 | 1 |
| 5961 | - | 6506363 | 6507700 | + | *pyrQ* | *PA14_73070* | dihydroorotase | 34 | 44 | 42 | 52 | 37 | 48 | 1 |
| 5962 | - | 6508989 | 6507745 | - | - | *PA14_73090* | hypothetical protein | 55 | 75 | 57 | 78 | 55 | 76 | 1 |
| 5963 | - | 6509474 | 6509076 | - | - | *PA14_73100* | hypothetical protein | 56 | 61 | 63 | 79 | 55 | 74 | 1 |
| 5964 | - | 6511495 | 6509471 | - | - | *PA14_73110* | hypothetical protein | 37 | 40 | 37 | 50 | 38 | 45 | 1 |

| | A | B | C | D | E | F | G | H | I | J | K | L | M | N |
|---|---|---|---|---|---|---|---|---|---|---|---|---|---|---|
| 5965 | - | 6512529 | 6511570 | - | - | *PA14_73120* | hypothetical protein | 75 | 87 | 69 | 80 | 86 | 99 | 1 |
| 5966 | - | 6513877 | 6512693 | - | - | *PA14_73140* | hypothetical protein | 355 | 190 | 183 | 210 | 214 | 371 | 1 |
| 5967 | - | 6514115 | 6514738 | + | - | *PA14_73150* | hypothetical protein | 64 | 71 | 68 | 63 | 57 | 61 | 1 |
| 5968 | - | 6515098 | 6516303 | + | - | *PA14_73160* | permease | 14 | 15 | 11 | 11 | 12 | 10 | 1 |
| 5969 | - | 6518202 | 6516367 | - | *glmS* | *PA14_73170* | glucosamine--fructose-6-phosphate aminotransferase | 56 | 66 | 45 | 35 | 51 | 36 | 1 |
| 5970 | - | 6518991 | 6518218 | - | *glmR* | *PA14_73190* | GlmR transcriptional regulator | 14 | 13 | 13 | 12 | 13 | 14 | 1 |
| 5971 | - | 6519568 | 6519059 | - | - | *PA14_73200* | hypothetical protein | 117 | 131 | 113 | 135 | 147 | 115 | 1 |
| 5972 | - | 6520946 | 6519582 | - | *glmU* | *PA14_73220* | glucosamine-1-phosphate acetyltransferase/N-acetylglucosamine-1-phosphate uridyltransferase | 162 | 156 | 157 | 150 | 149 | 150 | 1 |
| 5973 | - | 6521492 | 6521067 | - | *atpC* | *PA14_73230* | F0F1 ATP synthase subunit epsilon | 368 | 574 | 341 | 309 | 349 | 273 | 1 |
| 5974 | - | 6522910 | 6521534 | - | *atpD* | *PA14_73240* | F0F1 ATP synthase subunit beta | 1062 | 1298 | 896 | 864 | 1074 | 811 | 1 |
| 5975 | - | 6523801 | 6522941 | - | *atpG* | *PA14_73250* | F0F1 ATP synthase subunit gamma | 569 | 763 | 496 | 477 | 593 | 430 | 1 |
| 5976 | - | 6525396 | 6523852 | - | *atpA* | *PA14_73260* | F0F1 ATP synthase subunit alpha | 750 | 1053 | 677 | 663 | 796 | 543 | 1 |
| 5977 | - | 6525951 | 6525415 | - | *atpH* | *PA14_73280* | F0F1 ATP synthase subunit delta | 659 | 966 | 633 | 620 | 761 | 507 | 1 |
| 5978 | - | 6525952 | 6525961 | ? | - | predicted RNA | - | 1059 | 1472 | 1005 | 867 | 930 | 747 | 1 |
| 5979 | - | 6526433 | 6525963 | - | *atpF* | *PA14_73290* | F0F1 ATP synthase subunit B | 1221 | 1648 | 1172 | 996 | 1041 | 791 | 1 |
| 5980 | - | 6526748 | 6526491 | - | *atpE* | *PA14_73300* | F0F1 ATP synthase subunit C | 1755 | 2830 | 1895 | 2307 | 2605 | 1945 | 1 |
| 5981 | - | 6527667 | 6526798 | - | *atpB* | *PA14_73310* | F0F1 ATP synthase subunit A | 250 | 301 | 265 | 281 | 278 | 220 | 1 |
| 5982 | - | 6528064 | 6527684 | - | *atpI* | *PA14_73320* | F0F1 ATP synthase subunit I | 57 | 42 | 71 | 63 | 48 | 50 | 1 |
| 5983 | - | 6529100 | 6528228 | - | *parB* | *PA14_73330* | chromosome partitioning protein Spo0J | 210 | 252 | 214 | 197 | 202 | 182 | 1 |
| 5984 | - | 6529898 | 6529110 | - | *soj* | *PA14_73350* | chromosome partitioning protein Soj | 139 | 137 | 132 | 117 | 124 | 122 | 1 |
| 5985 | - | 6530561 | 6529917 | - | *gidB* | *PA14_73360* | 16S rRNA methyltransferase GidB | 106 | 137 | 132 | 132 | 120 | 102 | 1 |
| 5986 | - | 6532453 | 6530561 | - | *gidA* | *PA14_73370* | tRNA uridine 5-carboxymethylaminomethyl modification protein GidA | 87 | 90 | 104 | 111 | 87 | 85 | 1 |
| 5987 | - | 6533310 | 6532927 | - | - | *PA14_73390* | hypothetical protein | 24 | 26 | 29 | 31 | 20 | 26 | 1 |
| 5988 | - | 6535001 | 6533634 | - | *trmE* | *PA14_73400* | tRNA modification GTPase TrmE | 43 | 42 | 43 | 44 | 39 | 39 | 1 |
| 5989 | - | 6536808 | 6535072 | - | - | *PA14_73410* | inner membrane protein translocase component YidC | 120 | 146 | 118 | 104 | 133 | 78 | 1 |
| 5990 | - | 6537456 | 6537049 | - | *rnpA* | *PA14_73420* | ribonuclease P | 764 | 849 | 783 | 698 | 634 | 504 | 1 |

[a]Associated binding site refers to the genomic position of the base in the center of a given ChIP peak of a confirmed RhlR binding site from Table 1.

[b,c]Gene start and gene end indicate the transcription start and end. In the case of untranslated genes, this indicates the transcription start and end as predicted by Rockhopper [66].

[d]Strand indicates whether gene is present on the + or – strand.

Table S3

| | A | B | C | D | E |
|---|---|---|---|---|---|
| 1 | **Strain Name** | **Genotype** | **Plasmid** | **Antibiotic Resistance** | **Source** |
| 2 | JPS0112 | *E. coli* DH5a | pBADA-*rhlR* | Amp | Paczkowski |
| 3 | JPS0114 | *E. coli* DH5a | pCS26-p*rhlA*-*luxCDABE* | Kan | Paczkowski |
| 4 | JPS0115 | *E. coli* DH5a | pCS26-p*lasB*-*luxCDABE* | Kan | Paczkowski |
| 5 | JPS0151 | *P. aeruginosa* Δ*rhlR* | | | Mukherjee |
| 6 | JPS0154 | *P. aeruginosa* Δ*rhlI* | | | Mukherjee |
| 7 | JPS0278 | *P. aeruginosa* Δ*rhlI* Δ*pqsE* | | | Simanek |
| 8 | JPS0476 | *E. coli* Top10 | pBADA-*rhlR*, pCS26-p*rhlA*-*luxCDABE*, pACYC-*pqsE* | Amp, Kan, Tet | Taylor |
| 9 | JPS0477 | *E. coli* Top10 | pBADA-*rhlR*, pCS26-p*rhlA*-*luxCDABE*, pACYC-empty | Amp, Kan, Tet | Taylor |
| 10 | JPS0489 | *E. coli* DH5a | pACYC-*pqsE* | Tet | Taylor |
| 11 | JPS0544 | *P. aeruginosa pqsE*- NI (R243A/R246A/R247A) | | | Simanek |
| 12 | JPS0561 | *P. aeruginosa* Δ*pqsE* | | | Simanek |
| 13 | JPS0568 | *P. aeruginosa pqsE* (D73A) | | | Taylor |
| 14 | JPS0808 | *E. coli* DH5a | pACYC-empty | Tet | Taylor |
| 15 | JPS0920 | *E. coli* Top10 | pBAD-*rhlR*, pCS26-p*hcnA*-*luxCDABE*, pACYC-empty | Amp, Kan, Tet | This study |
| 16 | JPS0921 | *E. coli* Top10 | pBAD-*rhlR*, pCS26-p*hcnA*-*luxCDABE*, pACYC-*pqsE* | Amp, Kan, Tet | This study |
| 17 | JPS0924 | *E. coli* Top10 | pBAD-*rhlR*, pCS26-p*lasB*-*luxCDABE*, pACYC-empty | Amp, Kan, Tet | This study |
| 18 | JPS0925 | *E. coli* Top10 | pBAD-*rhlR*, pCS26-p*lasB*-*luxCDABE*, pACYC-*pqsE* | Amp, Kan, Tet | This study |
| 19 | JPS0948 | *E. coli* Top10 | pBAD-*rhlR*, pCS26-p*lecB*-*luxCDABE*, pACYC-empty | Amp, Kan, Tet | This study |
| 20 | JPS0949 | *E. coli* Top10 | pBAD-*rhlR*, pCS26-p*lecB*-*luxCDABE*, pACYC-*pqsE* | Amp, Kan, Tet | This study |

Table S4

| | A | B | C |
|---|---|---|---|
| 1 | **Primer Name** | **Primer Sequence** | **Description** |
| 2 | oJP967 | CAACCAGAAGATCGGCAAGTA | *lasB* RT-PCR primer |
| 3 | oJP968 | GTTCATGTCGACGGTGATGA | lasB RT-PCR primer |
| 4 | oJP1326 | CAACACGATATCCAGCCCCT | *hcnA* RT-PCR primer |
| 5 | oJP1327 | CATTGAGCACGTTGAGCACG | *hcnA* RT-PCR primer |
| 6 | oJP1328 | CCTGGCCGAACATTTCAACG | *rhlA* RT-PCR primer |
| 7 | oJP1329 | TTTCCACCTCGTCGTCCTTG | *rhlA* RT-PCR primer |
| 8 | oJP1330 | GAGGAACTGGAAGCGGTCAA | *gyrA* RT-PCR primer |
| 9 | oJP1331 | CTTCCTCGGTGATCAGGTCG | *gyrA* RT-PCR primer |
| 10 | oJP1358 | ATAAGCCTCACCCCCTGTCA | *sigX* RT-PCR primer |
| 11 | oJP1359 | AACGCCGCATCAATTCTTCA | *sigX* RT-PCR primer |
| 12 | oJP1436 | AATGATACGGCGACCACCGAGATCTACAC | ChIP primer |
| 13 | oJP1437 | CAAGCAGAAGACGGCATACGAGAT | ChIP primer |
| 14 | oJP1474 | CAGGTCAACAACGAAACCGTC | *lecB* RT-PCR primer |
| 15 | oJP1475 | GGCGAAGTTCAGCTCATTGG | *lecB* RT-PCR primer |
| 16 | oJP1506 | ATTACAGACAGCGTAACAATCG | *lecB* promoter 220 bp fragment |
| 17 | oJP1507 | TATGGCGTGTCGGGAGAGAT | *lecB* promoter 220 bp fragment |
| 18 | oJP1514 | CTGGGGTAGGCGCGGGTTCGA | *hcnA* promoter 220 bp fragment |
| 19 | oJP1515 | TGGATGGTCATGTCTGCCCGC | *hcnA* promoter 220 bp fragment |
| 20 | oJP1522 | TTTTCATGCCTTTTCCGCCAACC | *rhlA* promoter 220 bp fragment |
| 21 | oJP1523 | GGGGCTTGTGTGGGTCTTGC | *rhlA* promoter 220 bp fragment |
| 22 | oJP1592 | GAGAACCACATGGGGCCTCC | *hcnA* promoter fused to luxCDABE (5'-P) |
| 23 | oJP1593 | tatta <u>ggatcc</u> ATCCGTGAGAGAGAGTCGGG | *hcnA* promoter fused to luxCDABE (<u>BamHI</u>) |
| 24 | oJP1594 | AGTATATTTCGAGACAATCGCCCC | *lecB* promoter fused to luxCDABE (5'-P) |
| 25 | oJP1595 | tatta <u>ggatcc</u> GAATACCTGGCCTGCCTTTTC | *lecB* promoter fused to luxCDABE (<u>BamHI</u>) |
